# TRACER navigates rearrangement-driven sesterterpene chemical space via multimodal enzyme-product representation learning

**DOI:** 10.64898/2026.08.16.745124

**Authors:** Cuiping Xing, Kangjie Lv, Weiyan Zhang, Yuwei Chen, Keying Lan, Guoliang Zhu, Bin Zhu, Shou-Mao Shen, Xinjun Zhang, Yucheng Gu, Yue-Wei Guo, Hideaki Oikawa, Tom Hsiang, Lixin Zhang, Youyuan Li, Lan Jiang, Xueting Liu

**Author notes:** Correspondence (X. Liu), (L. Jiang), (Y. Li). These authors contributed equally (Cuiping Xing, Kangjie Lv, Weiyan Zhang).

## Abstract

Skeletal rearrangement drives the immense structural complexity of terpene, yet predicting it remains a formidable challenge due to sequence-function decoupling in terpene synthases. Here, we established TRACER (terpene rearrangement annotation via co-attentive enzyme-product representation), a multimodal framework mapping the latent associations between sequence-derived enzyme representations and product chemotypes. Retrospective validation proved TRACER’s exceptional precision in predicting compound classes and discriminating skeletal rearrangement (SR) from non-skeletal rearrangement (NSR) pathways. TRACER-guided genome mining characterized two bifunctional synthases, FsPS and AcPS, uncovering four unprecedented carbon skeletons. Density functional theory calculations deciphered these cyclization cascades, pinpointing a critical 5/6/11 tricyclic intermediate as the key branching node for scaffold diversification. Mutagenesis and molecular dynamics simulations suggested that E305 in FsPS enables rearrangement by maintaining active-site water exclusion, whereas its alanine mutation causes premature carbocation quenching. Collectively, this work establishes a predictive paradigm for the rational discovery and mechanistic elucidation of complex terpene architectures.

## Introduction

Terpenoids represent a pinnacle of natural product complexity, where structural diversity is primarily orchestrated by the variation in their carbon skeletons.^1, 2^ Terpene synthases (TPSs) are the enzymatic engines of this diversification, directing intricate carbocation-mediated cyclization cascades that involve high-energy skeletal rearrangements.^1, 3, 4^ However, the rational discovery of unprecedented skeletons is increasingly hindered by a fundamental sequence-function decoupling.^5, 6, 7, 8^ The sensitivity of these cascades is so pronounced that subtle, single-residue variations in the active site can dramatically redirect reaction trajectories, resulting in entirely different scaffolds.^1, 9, 10, 11, 12^ This interplay between intrinsic chemical complexity and sensitive enzyme regulation makes TPS function inherently unpredictable via traditional homology-based mining, leading to a “discovery plateau” characterized by high experimental burden and low hit rates.^6, 12^

To overcome this, recent efforts have incorporated biochemical context, such as mechanism-based docking and machine learning (ML), into functional inference. While docking and virtual screening provide substrate-level constraints, their structure-based nature and computational cost preclude large-scale genomic screening.^13^ Recent studies, including EC-number annotation from protein sequences by CLEAN,^14^ compatibility prediction between α-ketoglutarate-dependent non-heme iron enzymes and their substrates by CATNIP,^15^ and substrate specificity prediction across representative enzyme families by EZSpecificity,^16^ have demonstrated the growing impact of ML in enzyme function annotation and substrate specificity inference.^17^ ML has also been increasingly applied to TPS functional prediction, enabling rapid identification of TPSs from genomic data,^18^ inference of substrate preference,^19^ and detection of key active-site determinants underlying catalytic function.^8^ However, these models remain protein-only input frameworks, limiting their ability to learn the nonlinear enzyme-product dependencies and generally restricting predictions to specific types of carbon skeleton.^8, 19, 20^ Bridging this gap requires a multimodal framework capable of co-aligning protein-level evolutionary information with the structural grammar of chemical scaffolds.^15, 16^

To address the challenge of predicting skeletal rearrangement potential, fungal bifunctional terpene synthases (BFTPSs) serve as an ideal system because they comprise two catalytic categories with distinct rearrangement propensities: Type A primarily diversifies products through stereoisomerization, whereas Type B frequently generates structural complexity via skeletal rearrangements (Scheme 1A and Supplementary Table 1).^11, 21, 22, 23, 24, 25, 26^ However, the extreme sensitivity of these cascades to subtle active-site variations makes traditional sequence-based annotation ineffective.^8^ The paraphyletic distribution of skeletal-rearranging (SR) and non-skeletal-rearranging (NSR) BFTPSs highlights a profound sequence-function decoupling, where primary sequences fail to dictate catalytic outcomes (Scheme 1B and Supplementary Fig. 1). This “functional opacity,” exacerbated by the scarcity of experimental data, precludes the development of high-quality supervised models. Nevertheless, large language models (LLMs) pre-trained on vast biochemical landscapes can capture deep contextual representations and single-residue determinants, enabling robust prediction even in limited training data.^27, 28^ These advances provide a transformative opportunity to unlock the rearrangement-driven diversity hidden within uncharted genomic resources.^29, 30^

**Fig. 1.**
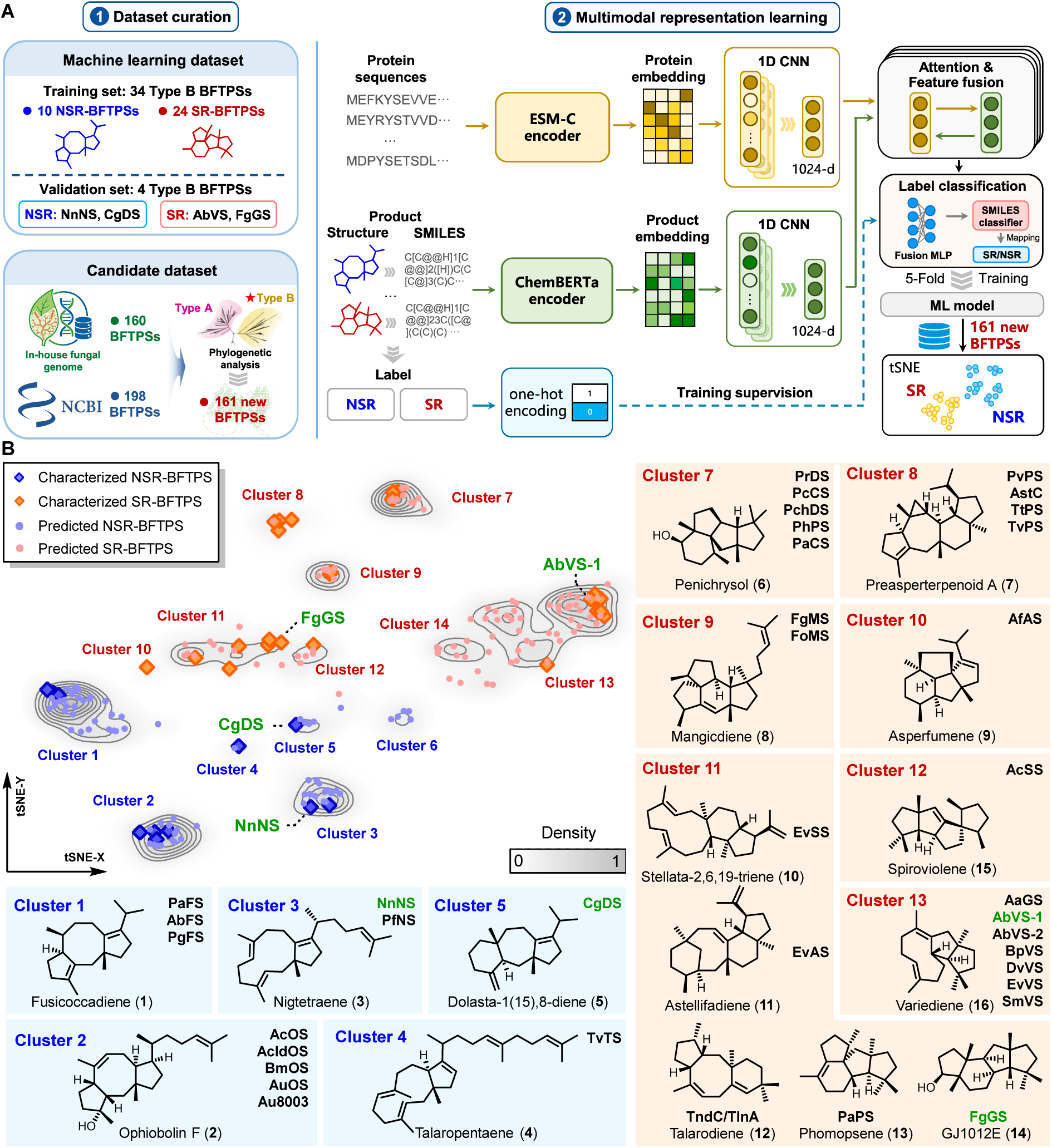
Multimodal enzyme-product representation learning framework and functional landscape of BFTPS. (**A**) Dataset construction and TRACER architecture. Thirty-four experimentally characterized Type B BFTPSs were curated from the literature to build the machine-learning dataset, while 161 uncharacterized BFTPS candidates were obtained through genome mining from NCBI and an in-house plant endophytic fungal genome library. In TRACER, protein sequences inputs and SMILES representations are first embedded using pre-trained encoders, followed by convolution-based local feature extraction. A cross-modal attention mechanism decodes latent enzyme-product relationships in the shared embedding space, and the fused representations drive both functional classification and downstream visualization. (**B**) t-SNE projection of the learned latent space reveals chemotype-based clustering of characterized and predicted BFTPSs. SR-BFTPSs (red) and NSR-BFTPSs (blue) resolve into distinct functional groups (Clusters 1–14), each corresponding to representative terpene skeletons. Density contours highlight cluster boundaries inferred from kernel density estimation. The four BFTPSs utilized for model validation are highlighted in green.

**Scheme 1.**
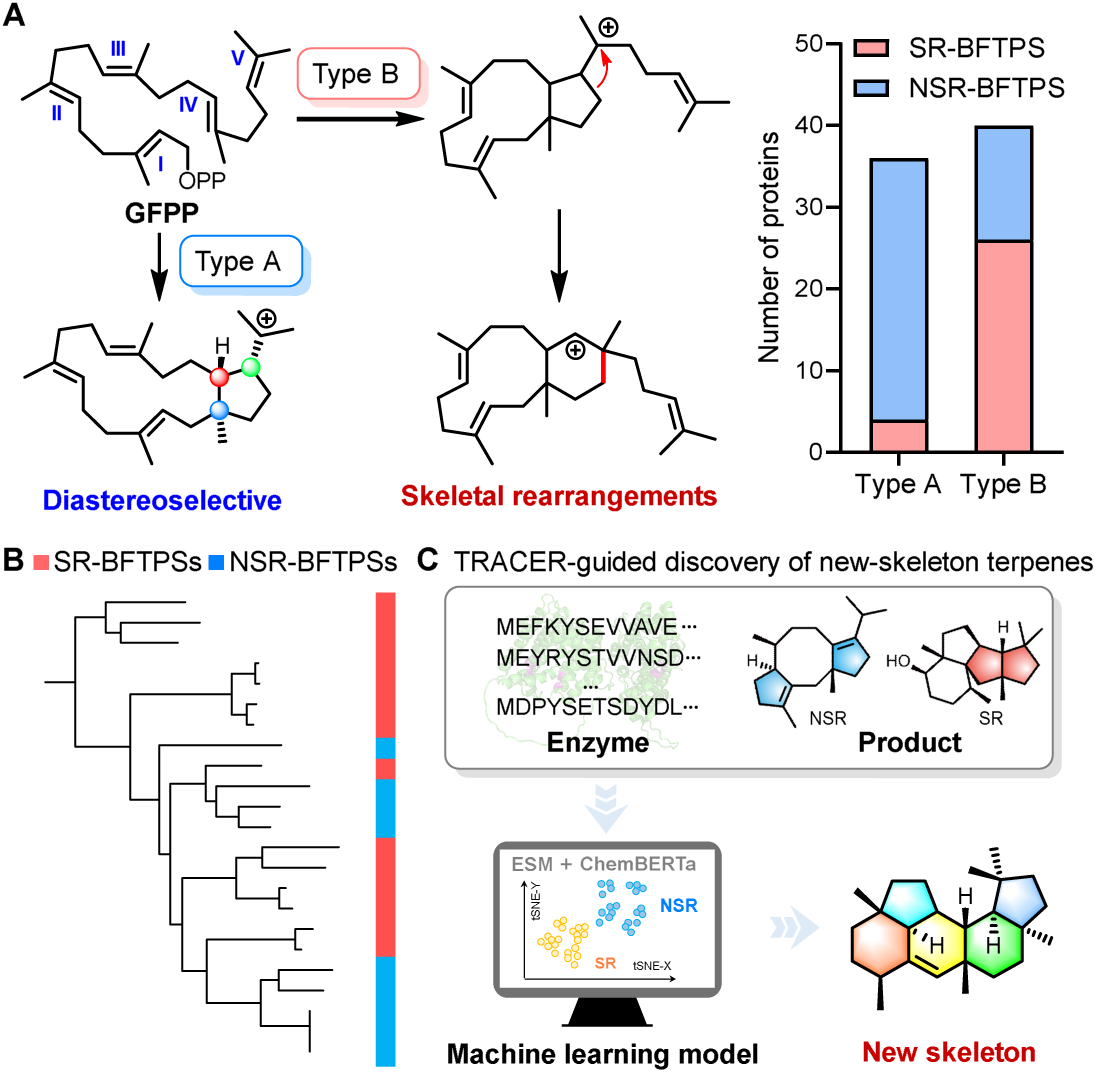
Sequence-function decoupling in BFTPSs and TRACER-enabled identification of rearrangement-capable enzymes. (**A**) Mechanistic divergence between Type A and Type B BFTPSs. Type A enzymes primarily expand structural diversity through stereoisomerization, whereas 68% of Type B enzymes undergo complex skeletal rearrangements to construct unique polycyclic frameworks. (**B**) Functional opacity in the BFTPSs phylogenetic landscape. The paraphyletic distribution of skeletal-rearranging (SR, red) and non-skeletal-rearranging (NSR, blue) BFTPSs underscores a profound sequence-function decoupling, where primary sequence similarity fails to dictate catalytic outcomes. (**C**) TRACER-enabled enzyme-product representation learning for discovery terpene with new ckeleton. By integrating protein sequence (ESM-C) and product chemical (ChemBERTa) embeddings, TRACER captures latent enzyme-product relationships, enabling high-precision classification and prioritization of SR-capable enzymes for the exploration of uncharted chemical space.

Here, we present a multimodal enzyme-product representation learning framework named TRACER (terpene rearrangement annotation via co-attentive enzyme-product representation) that bridges protein sequences and product structure by coupling ESM-C-derived^31^ protein embeddings with ChemBERTa-derived^32^ molecular representations. Rather than relying solely on sequence similarity, this framework learns latent associations between BFTPS sequences and terpene chemotypes, enabling the prioritization of enzymes with skeletal-rearranging potential (Scheme 1C). Guided by this predictive prioritization, we identified two previously uncharacterized enzymes, FsPS and AcPS, which collectively produced 10 undescribed sesterterpenes, including six compounds bearing four unprecedented carbon skeletons. Density functional theory (DFT) calculations and molecular dynamics (MD) simulations further elucidated the branching nodes of these complex cascades and identified E305 enables rearrangement by maintaining active-site water exclusion, whereas its alanine mutation causes premature carbocation quenching. Conclusively, this study establishes an operative paradigm that reconciles sequence-function decoupling, effectively transforming expanding genomic resources from an unannotated sequence space into a predictable repository of unprecedented molecular architectures.

## Results

### TRACER: A multimodal enzyme-product representation learning framework for predicting rearrangement-capable BFTPSs

To establish a systematic basis for functional prediction, we curated a dataset of 75 experimentally verified BFTPSs from the literature, comprising 34 Type A, 38 Type B, and 3 atypical BFTPSs (Supplementary Table 1). A pronounced divergence in catalytic outcomes was observed: while only 12% (4/34) of Type A BFTPSs directed skeletal rearrangement, a substantial 68% (26/38) of Type B enzymes generated rearranged scaffolds as their major products. Due to this higher propensity for skeletal rearrangement, we focused subsequent model development on Type B BFTPSs, which were categorized as either skeletal-rearranging (SR-BFTPSs, n = 26) or non-skeletal-rearranging (NSR-BFTPSs, n = 12). A sequence similarity network (SSN) revealed that SR- and NSR-BFTPSs are phylogenetically interspersed (Supplementary Fig. 2A), indicating that primary sequence identity alone is insufficient to distinguish their functional divergence. Given that skeletal rearrangement is governed by subtle active-site features that shape, stabilize, and shield carbocation intermediates, we reasoned that a multimodal approach integrating both protein sequences and chemical features would be essential to capture these cryptic catalytic differences.

To resolve these functional discrepancies, we developed TRACER, a multimodal enzyme-product representation learning framework (Fig. 1A and Supplementary Fig. 3). The architecture employs a dual-encoder strategy in which full-length protein embeddings are generated using the pre-trained ESM-C model,^31^ while corresponding terpene products are captured via ChemBERTa.^32^ A multi-branch 1D convolutional neural network (CNN) is then utilized to extract local features associated with protein sequence motifs and product substructure patterns.^33, 34^ These cross-modal features are subsequently integrated via a bi-directional co-attention module designed to capture residue-fragment interactions associated with skeletal rearrangements (Supplementary Fig. 3).^16, 35^ TRACER is optimized using a composite loss function combining product-level cross-entropy loss, label-level (SR/NSR) supervision, and supervised contrastive loss to promote enzyme-product representation alignment (Supplementary Fig. 3).^36^ For inference, each uncharacterized BFTPS is paired with all compounds in the learned chemical library to compute a compatibility score. The SR/NSR classification of the query protein is then assigned via the predefined product-to-label mapping, and the resulting product-label pairs facilitate downstream clustering. To mitigate overfitting under limited training data, we implement a robust regularization strategy encompassing dropout,^37^ weight decay,^38^ and early stopping, which effectively stabilized model convergence (Supplementary Fig. 4).

The curated Type B BFTPS dataset was further partitioned into a training subset and an independent validation subset at an approximately 9:1 ratio. The model was trained on 34 experimentally verified enzyme-product pairs (24 SR- and 10 NSR-BFTPSs) across 14 unique product chemotypes and was evaluated on a validation set of four sequences representing diverse rearrangement types and distinct product scaffold classes. Performance was benchmarked using accuracy, recall, F1-score, and precision (Supplementary Fig. 5). In five-fold cross-validation, the model converged within 500 epochs and achieved an overall label accuracy of 92.5%–95.2% (Supplementary Tables 2 and 3), significantly outperforming BLAST-based classification methods (84.2%) (Supplementary Table 4). This alignment indicates that integrating protein and product representations extracts critical catalytic insights beyond primary sequence similarity. In support of this, t-distributed stochastic neighbor embedding (t-SNE) visualization of the learned latent space demonstrated clear separation between SR- and NSR-BFTPSs, with enzymes yielding structurally related scaffolds forming cohesive chemical clusters. Together, these results suggest that TRACER learns enzyme-product representations that are informative for SR/NSR classification and scaffold-level organization, providing a basis for machine-learning-guided prioritization in genome mining.

### Retrospective benchmarking resolves rearrangement-capable BFTPSs beyond sequence homology

To evaluate the predictive capacity of TRACER, we constructed a candidate library combining 160 putative BFTPSs from our in-house fungal genome collection^39, 40, 41, 42, 43, 44, 45^ and 198 sequences from the ClusteredNR database (Supplementary Table 5). After manual curation, including reading frame correction via 2ndFind (https://biosyn.nih.go.jp/2ndfind/) and canonical motif-based filtering, 161 high-confidence Type B candidates were selected for functional prioritization (Supplementary Table 6). The model-derived embedding landscape resolved these candidates and 38 known BFTPSs into 14 distinct functional clusters (Clusters 1–14), alongside 13 isolated uncharacterized enzymes located outside any cluster (Fig. 1B). Additionally, two clusters, Clusters 6 and 14, represented “dark clusters” devoid of previously characterized representatives (Fig. 1B). Analysis of the cyclization pathways and product scaffolds represented in the remaining 12 clusters demonstrated that 26 SR-BFTPSs (red) and 12 NSR-BFTPSs (blue) segregated clearly into distinct regions, suggesting that the TRACER-derived latent space captures structural features beyond primary sequence similarity, markedly resolving the intermixed organization observed in SSN analysis (Supplementary Fig. 2B). Crucially, the four validation-set enzymes reserved for the validation set (FgGS,^46^ AbVS-1,^47^ NnNS,^42^ and CgDS^48^) were correctly mapped to their expected SR or NSR categories (Fig. 1B).

Retrospective analysis showed that the 38 characterized BFTPSs were distributed across 12 TRACER-derived clusters, comprised eight multi-reference clusters (MRCs) containing at least two characterized BFTPSs and four singleton-reference clusters (SRCs) featuring only single characterized member (Fig. 1B and Supplementary Table 7). Seven of the eight MRCs (Clusters 1‒3, 7‒9, and 13) exhibited high scaffold consistency, highlighting the model’s ability to organize BFTPSs according to product beyond SR/NSR assignment (Fig. 1B). Cluster 11 emerged as a unique case where all six enzymes were correctly classified as SR, yet they yielded five distinct scaffolds (Fig. 2A and Supplementary Table 7). Specifically, four enzymes (PaPS,^22^ FgGS,^46^ TndC,^49^ and TlnA^50^) utilized a Type B path II early-stage cyclization pathway^21^ to form complex ring systems, whereas EvSS^51^ and EvAS^52, 53^ followed a Type B_path III route^21^ to generate 5/6/11 and 5/6/8/6 systems, respectively (Fig. 2B and Supplementary Fig. 6). While the model generally aligned with terminal product identity, this co-clustering of divergent scaffolds in Cluster 11 likely stems from the inherent scarcity of experimental training data in this chemical space. Further expanding the training set with more experimentally characterized BFTPSs will improve the accuracy of product-related clustering.^54^

**Fig. 2.**
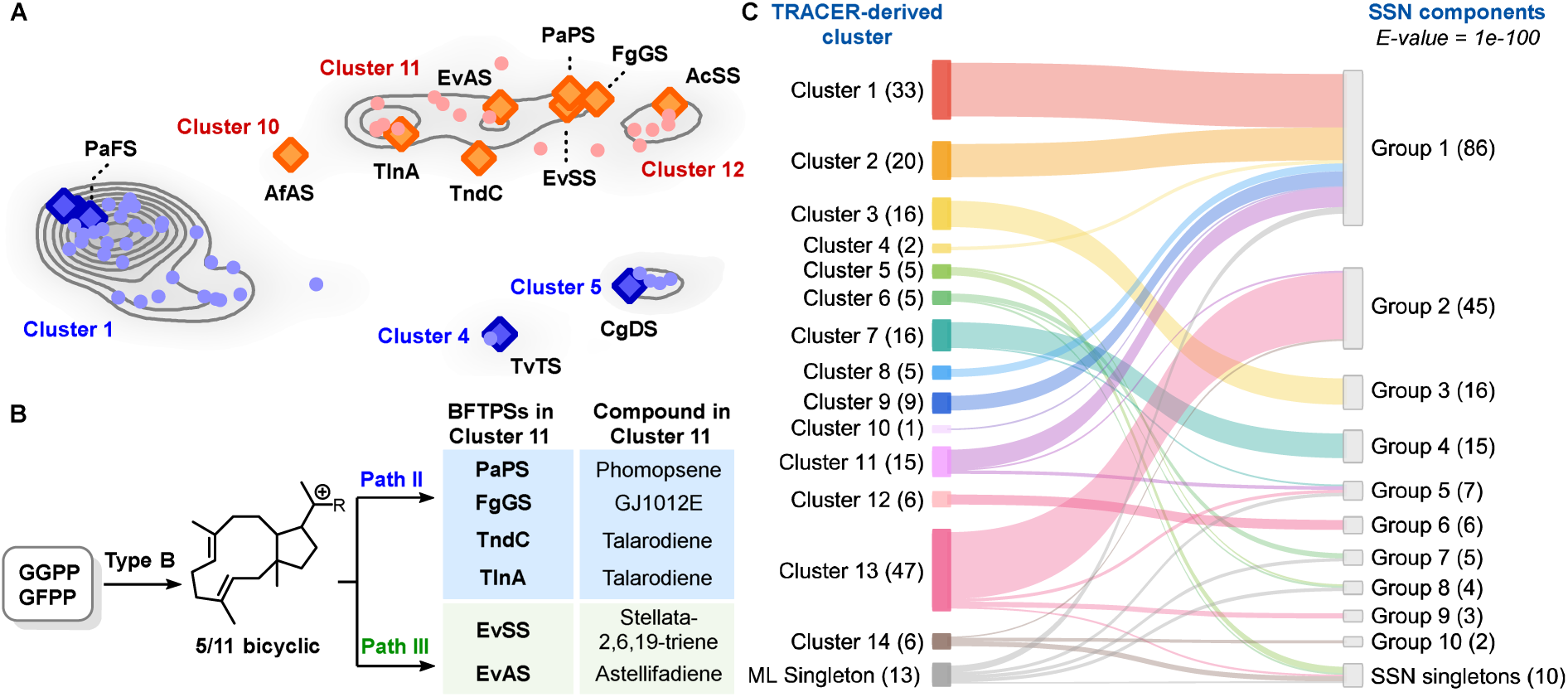
Evaluation of TRACER-derived clustering across atypical and sequence-discordant BFTPS regions. (**A**) Magnified latent-space views of Cluster 11 and singleton-anchored clusters, including Clusters 4, 5, 10, and 12. (**B**) Schematic comparison of divergent cyclization pathway and product outcomes among Cluster 11 BFTPSs. (**C**) Alluvial diagram comparing TRACER-derived clusters with SSN components. The SSN components were generated at E-value cutoff of 1e−100. Left-side nodes represent clusters defined by the TRACER-derived latent space, whereas right-side nodes represent connected components in the SSN. Flow widths reflect the number of proteins shared between each TRACER-derived cluster and SSN component.

The remaining four SRCs captured atypical catalytic logic within the BFTPS landscape (Fig. 2A and Supplementary Table 7). For instance, Cluster 4 was anchored by TvTS, the only Type B BFTPS reported to produce a triterpene scaffold,^23^ while Cluster 5 corresponded to CgDS,^48^ which generated a 5/7/6 tricyclic framework, the most structurally complex scaffold among known NSR outcomes. A particularly insightful case was Cluster 10, anchored by AfAS. Despite sharing 76.0% sequence similarity with the Cluster 1 member PaFS (a fusicoccadiene producer yielding an NSR product^22^), AfAS was positioned at the boundary between the SR and NSR regions and generated the rearranged product asperfumene.^55^ Interestingly, a single I65Y mutation redirected AfAS from the SR product to a NSR product fusicoccadiene.^55^ Similarly, the isolation of the spirocyclic terpene producer AcSS from Cluster 12 underscored the model’s discriminative resolution for unconventional cyclization logic.^25^ Together, these SRCs indicate that TRACER can resolve distinct scaffold-associated features from single-reference product groups within the BFTPS landscape.

To evaluate whether TRACER simply recapitulated conventional sequence-based grouping, we compared the TRACER-derived clusters with SSN components derived from pairwise sequence similarity.^56^ The results showed that proteins grouped within the same SSN component were separated into multiple TRACER-derived clusters (Supplementary Fig. 2B and Supplementary Table 8). For example, SSN Group 1 grouped together proteins with a relatively high average pairwise sequence similarity of 57.0%, whereas the TRACER further resolved this sequence-connected component into product-divergent Clusters 1, 2, 8, 9, and 11. (Supplementary Fig. 7). Conversely, proteins distributed across multiple SSN groups or singleton nodes were consolidated by the TRACER into coherent clusters, such as Clusters 5, 13, and 14. A hallmark of this discriminative power was DvVS, which was placed into SSN Group 5 rather than the variediene-associated Group 2, a known challenge for conventional sequence-based prediction.^6, 57, 58^ In contrast, DvVS was assigned into Cluster 13, the variediene-producer cluster, despite its low average similarity to other Cluster 13 members (47.4%) (Supplementary Table 8).^41^ These results demonstrated that the model effectively maps the enzyme-product landscape, enabling high-fidelity SR/NSR discrimination and product-coherent clustering. This robust representation provides the prerequisite for the model-guided discovery of uncharacterized BFTPSs.

We further performed an ablation analysis in which product structural information was removed to evaluate its contribution to skeletal rearrangement prediction (Supplementary Fig. 8). Under this setting, the known NSR-BFTPS CgDS^48^ in the validation set was incorrectly classified as SR, indicating a decline in predictive accuracy. The model’s ability to maintain product-coherent clustering was also reduced. For example, proteins producing fusicoccadiene and ophiobolin F were grouped into the same cluster, and the variediene-producing BpVS^25^ was positioned far from the other variediene-producing BFTPSs. In addition, singleton-anchored clusters that were clearly resolved before ablation were no longer well distinguished, with AfAS^55^ clustered together with EvAS^52^ and EvSS^51^ in the ablated model. These results clearly demonstrated that the integration of product information substantially improves both SR/NSR discrimination and product-related clustering.

### Experimental validation of TRACER prioritization enables the discovery of unprecedented sesterterpene skeleton

The retrospective validation established that TRACER captures the underlying biosynthetic logic of BFTPSs, even across low sequence-homology regimes. Building upon these insights, the TRACER-derived landscape was deployed as a strategic framework to prioritize candidates for experimental validation, aiming to identify enzymes capable of producing terpenes bearing new carbon skeletons. While most clusters were anchored by characterized enzymes, Clusters 6 and 14 emerged as “dark clusters” composed entirely of uncharacterized enzymes, corresponding to predicted SR- and NSR-BFTPSs, respectively (Fig. 1A). To systematically interrogate these predictions and their utility in functional bioprospecting, a structured prioritization strategy was implemented with three objectives: (i) to verify functional fidelity in discriminating SR from NSR activities, (ii) to evaluate cluster reliability by testing whether candidates within product-coherent clusters yield consistent scaffolds despite low primary sequence identity, and (iii) to identify enzymes capable of producing unprecedented scaffolds. Guided by this prioritization, 18 uncharacterized BFTPS candidates were selected for experimental validation, including 11 for assessing product-coherent clustering, three showing discordance between TRACER- and SSN-derived relationships, five SRC-associated candidates representing atypical catalytic logic, and two from the “dark clusters” (Cluster 6 and 14) (Fig. 3A and Supplementary Table 9).

**Fig. 3.**
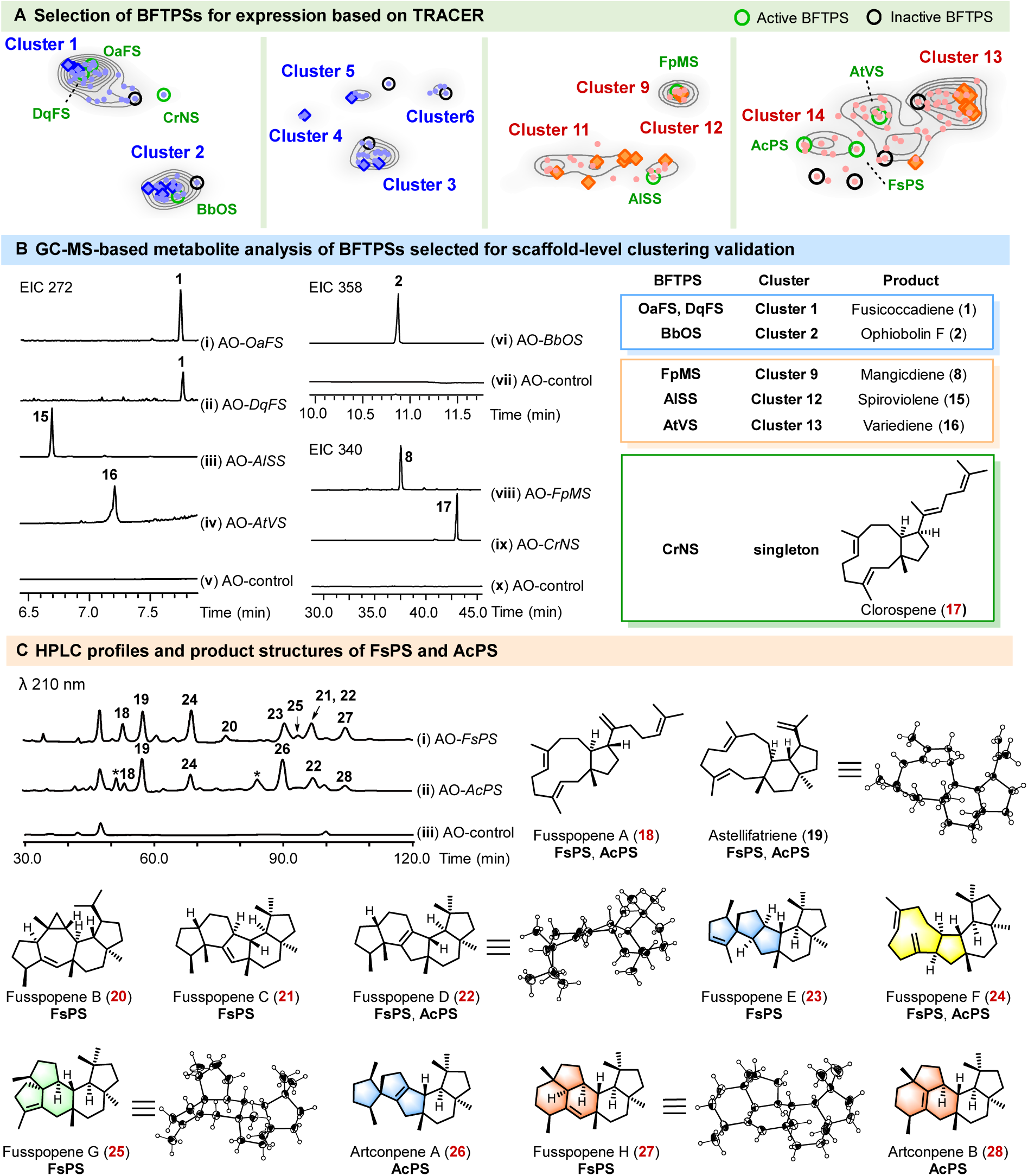
TRACER-guided selection of novel BFTPS candidates and experimental characterization of their products. (**A**) Distribution of the selected BFTPS candidates in the TRACER-derived embedding space. Enzymes chosen for experimental characterization are highlighted within the predictive functional landscape. (**B**) GC-MS analysis of metabolites produced by A. oryzae transformants. Extracted ion chromatograms (EICs) of m/z 272, 340 and 358 trace the metabolic profiles of expressing OaFS (i), DqFS (ii), AlSS (iii), AtVS (iv), BbOS (vi), FpMS (viii), and CrNS (ix), alongside their elucidated chemical structures. (**C**) HPLC profiles and product structures of FsPS and AcPS. Metabolites were produced by A. oryzae transformants expressing FsPS (i) or AcPS (ii), and detected by DAD at 210 nm. Low-abundance products that could not be structurally characterized are marked with an asterisk (*). Newly identified compounds are highlighted in red, and compounds bearing newly discovered skeletons are indicated by colored structural fills.

All 18 candidates were expressed in *Aspergillus oryzae* NSAR1, with nine yielding detectable terpene products via gas chromatography-mass spectrometry (GC-MS) analysis. Remarkably, structural characterization revealed that all these nine enzymes perfectly matched the model’s SR/NSR prediction (achieving a 100% concordance rate), underscoring the robustness of the embedding space (Fig. 3). Notably, five enzymes selected from MRCs were evaluated and yielded expected products. Within the NSR-BFTPS region, DqFS and OaFS from Cluster 1 produced fusicoccadiene (**1**),^22, 25, 59^ while BbOS from Cluster 2 yielded ophiobolin F (**2**),^25, 60^ matching the product profiles of the known enzymes in their respective clusters. Concurrently, the predictive accuracy of the SR-BFTPS region was confirmed, as FpMS (Cluster 9) generated mangicdiene (**8**),^45, 46, 61^ AlSS (Cluster 12) yielded spiroviolene (**15**),^25^ and AtVS (Cluster 13) produced variediene (**16**) (Fig. 3B, Supplementary Figs. 9−13; Supplementary Tables 10−14).^25, 41, 47, 62^

We further focused on candidates within “dark clusters” and isolated singleton nodes, reasoning that these distinct candidates were the most likely to harbor unexplored chemistry. This strategy led directly to the characterization of three enzymes: CrNS (a singleton adjacent to Cluster 1), FsPS (a singleton adjacent to Cluster 14), and AcPS (from Cluster 14). As an NSR-BFTPS, CrNS produced a new, unrearranged bicyclic sesterterpenoid (**17**) (Fig. 3B, Supplementary Figs. 14 and 28−29; Supplementary Table 15). In contrast, the SR-BFTPSs FsPS and AcPS displayed pronounced catalytic plasticity, generating a complex mixture of 11 distinct sesterterpenes (**18**–**28**, Fig. 3C), including ten newly discovered compounds. All structures were first elucidated by comprehensive NMR analysis (Supplementary Figs. 15–25; Supplementary Tables 16–26), and their absolute configurations were assigned based on electronic circular dichroism (ECD) spectroscopy and single-crystal X-ray diffraction (Supplementary Figs. 30–51 and 58; Supplementary Table 27). Consistent with the ablation analysis, removal of product information disrupted this prioritization logic, misplacing CrNS in the SR region and separating AtVS from the variediene-producing cluster (Supplementary Fig. 8). Moreover, incorporating the newly characterized BFTPSs into a retrained model further refined the functional landscape, with Cluster 11 redistributed into four product-related clusters. (Supplementary Fig. 59). This indicated that expansion of the dataset can improve the resolution of product-related clustering.

Compounds **18**–**22** represented complex bicyclic, tricyclic, or pentacyclic frameworks structurally related to known compounds like ophiobora-2,6,15(23),18-tetraene,^10^ astellifatriene,^53^ preasperterpenoid A,^63, 64^ and peniroquesines.^65^ Notably, the remaining six compounds (**23**–**28**) possessed four entirely unprecedented carbon skeletons. Among them, compounds **23** and **26** displayed a distinctive 5/6/5/5/5 spirocyclic scaffold, with a spiro-ring connection reminiscent of spiroviolene,^66, 67, 68^ marking the first spirocyclic scaffold reported among BFTPS-derived sesterterpenes. The 5/6/5/9 tetracyclic framework of **24** provided key structural support for the proposed downstream cyclization cascade. In contrast, **25** exhibited a structurally unique 5/6/6/5/5 polycyclic architecture, incorporating substructures analogous to those of phomopsene and mangicdiene.^45, 46, 61, 67^ Finally, **27** and **28** represented a newly observed 5/6/6/5/6 fused pentacyclic ring system with an unrecorded cyclization topology.

ChemBERTa-based molecular embedding placed these new skeleton-containing compounds near the edge of sesterterpenoid chemical space (Supplementary Fig. 60),^69^ with four compounds falling below the low-similarity threshold of Tanimoto similarity (T < 0.4) in the Morgan fingerprint-based Tanimoto analysis (Supplementary Table 28),^70^ indicating that TRACER-guided prioritization can uncover structurally underexplored scaffold space. Moreover, compounds **23** and **26** share a spiro-tetracyclic substructure with the anti-allergic metabolite spirograterpene A,^71^ while compound **25** contains substructures related to anti-inflammatory mangicdiene-derived mangicol analogues.^45, 61^ These structural similarities suggest that the new scaffolds of **23**, **26**, and **25** may provide promising frameworks with bioactivity potentials. The discovery of four novel scaffolds from a minimal set of targets demonstrated the superiority of TRACER-guided prioritization to effectively mine enzymes with high innovation vectors, highlighting its promise as a predictive strategy for terpene synthase discovery beyond conventional sequence-based approaches.^7, 72^

### DFT elucidation of cryptic rearrangement cascades

The product profiles of FsPS and AcPS revealed a remarkable structural diversity, encompassing eight distinct skeleton types ranging from bicyclic to complex pentacyclic frameworks. The identical stereochemistry and shared A/B ring substructures ubiquitous in the metabolic profiles of FsPS and AcPS point to a unified carbocation-driven trajectory (Fig. 4A). This shared trajectory diverges at advanced branching intermediates to generate the structurally diverse frameworks of **18**–**22** (Fig. 4 and Supplementary Fig. 61). To probe the thermodynamic and kinetic origins of this divergence in cyclization routes, DFT calculations were performed to map the intricate cyclization pathways leading to these products.

**Fig. 4.**
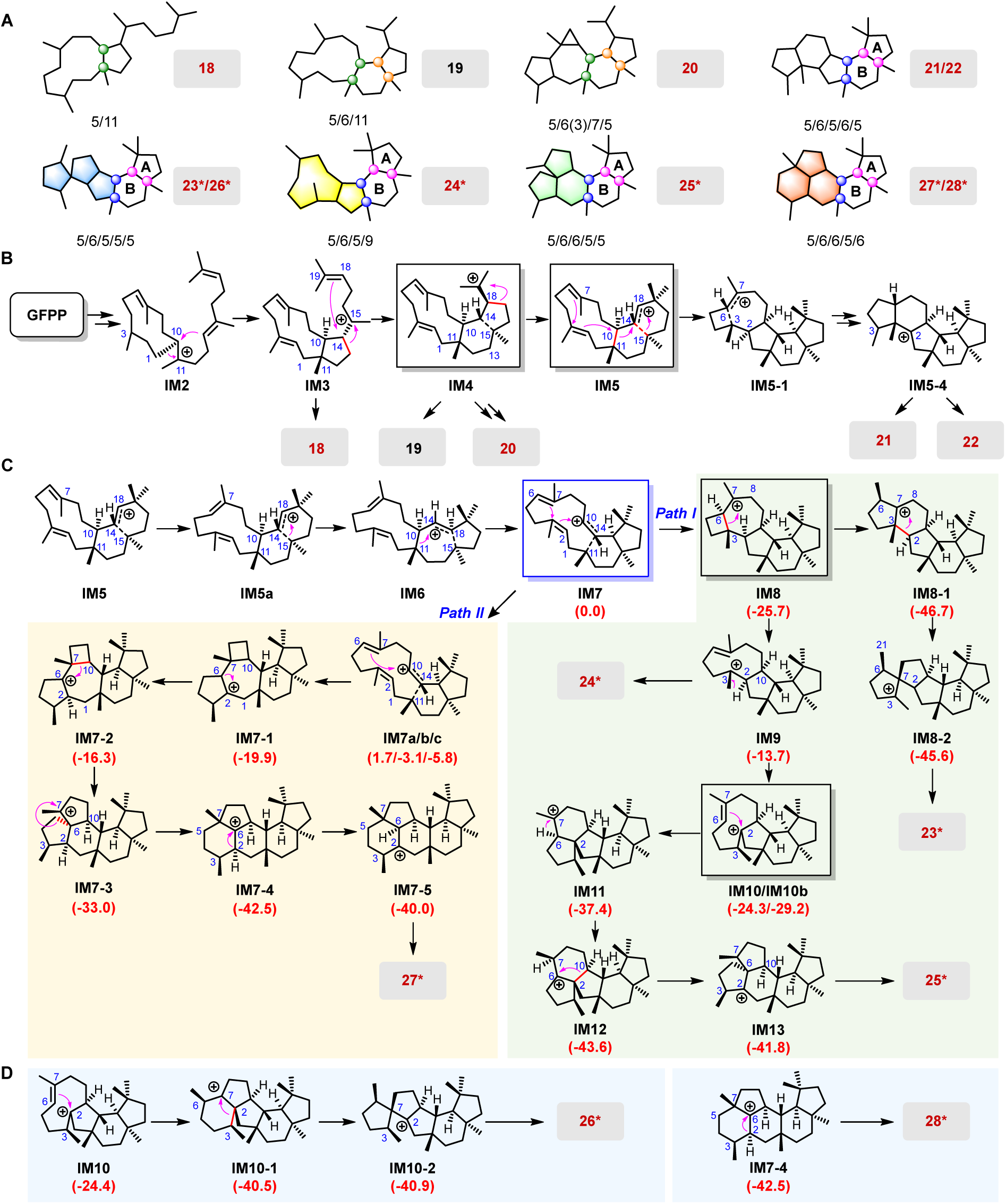
Proposed cyclization pathways for compounds 18–28 produced by FsPS and AcPS by DFT calculations. **(A**) Skeletons of FsPS and AcPS-derived products. Chiral centers with identical configurations are highlighted by circles of the same color on the corresponding skeletons. (**B**) Peniroquesine-like cyclization pathway leading to the formation of compounds **18**–**22**. (**C**) DFT-supported mechanisms for the formation of new skeleton compounds produced by FsPS. (**D**) DFT-supported mechanisms for the formation of new skeleton compounds **26** and **28** produced by AcPS. Gibbs free energies (kcal/mol, Gibbs free energies calculated at the M06-2X/6-31+G(d,p) level) relative to **IM7** are indicated in parentheses. **IM** denotes intermediate, **TS** denotes transition state. Newly characterized compounds are shown in red, and new skeletons are denoted by “*”.

The cascade begins with GFPP ionization and C1–C10 cyclization to form the carbocation intermediate **IM2**, followed by a 1,2-alkyl shift and subsequent C10–C14 cyclization to generate **IM3**, which undergo facile deprotonation to afford **18**.^73^ Alternatively, **IM3** undergoes ring expansion through terminal π-bond participation to yield the 5/6/11-fused intermediate **IM4**, which produced **19** after deprotonation.^10, 73^ **IM4** further served as a key branching intermediate: a 1,5-hydride shift and annulation cascade generated a preasperterpenoid A-like carbocation that afforded **20** (Fig. 4B and Supplementary Fig. 61),^74^ whereas reverse Wagner–Meerwein rearrangement and successive alkyl shifts produced the peniroquesine-like pentacyclic intermediate **IM5-4**, yielding **21** and **22** after deprotonation (Fig. 4B and Supplementary Fig. 61).^65, 73^

Notably, **IM5** undergoes a conformational rearrangement followed by two consecutive asynchronous 1,2-alkyl shifts to generate the secondary carbocation **IM7**,^73^ in which the C6=C7 and C2=C3 double bonds adopted a favorable orientation for subsequent divergent cyclization pathways. This geometry positioned **IM7** as a key branching intermediate, enabling two alternative cyclization modes that diverge through distinct bond-forming events. In ***Path I***, **IM7** undergoes asynchronous concerted annulation through the simultaneous formation of C2–C10 and C3–C6 bonds to generate the stabilized tertiary carbocation **IM8**. Subsequent Wagner–Meerwein methyl migration and ring contraction produced the 5/6/5/5/5 spirocyclic intermediate **IM8-2**, which yielded compound **23** after deprotonation (Fig. 4C and Supplementary Figs. 62 and 63).^75^ Alternatively, **IM8** undergoes ring expansion to yield the 5/6/5/9 tetracyclic intermediate **IM9**, whose direct deprotonation produced **24**. **IM9** further undergoes a 1,2-hydride shift, conformational rearrangement, and cation-mediated C2–C6 cyclization to generate **IM11**, followed by another hydride shift to form **IM12**. Subsequent C2–C10 bond cleavage and C6–C10 bond formation generated **IM13** with a 5/6/6/5/5 pentacyclic framework, whose undergoes deprotonation at C3 to yield **25** (Fig. 4C and Supplementary Figs. 64 and 65). This pathway strongly mirrors the proposed biosynthetic cascades of phomopsene and mangicdiene.^76, 77^ In ***Path II***, **IM7** undergoes conformational rearrangement and asynchronous annulation via C7–C10 and C2–C6 bond formation to generate **IM7-1**. Subsequent hydride migration, C7–C10 bond cleavage, and C6–C10 bond formation (**IM7-1 → IM7-2 → IM7-3**) yielded the 5/6/6/5/5 pentacyclic intermediate I**M7-3**. Further ring expansion and a final 1,2-hydride shift produced **IM7-5**, which undergoes deprotonation to afford **27** (Fig. 4C and Supplementary Figs. 66 and 67). Two additional C6–C10 cyclization routes from **IM7** toward **27** were also evaluated but appeared energetically unfavorable (Supplementary Figs. 66, 68 and 69).

AcPS additionally generated compounds **26** and **28** as pathway-specific products. Although **26** shared the same skeleton as **23**, DFT calculations excluded the previously proposed formation route^75^ from **IM8-2** due to a kinetically prohibitive activation barrier of an exocyclic 1,3-hydride shift (Δ*G*^ǂ^ = 54.7 kcal/mol) (Supplementary Fig. 63). Instead, **IM10** was proposed as an alternative branching intermediate that undergoes concomitant C2–C7 bond formation and 1,2-methyl migration to generate **IM10-1**. **IM10-1** then rearranges to the spirocyclic intermediate **IM10-2**,^78^ which subsequently undergoes deprotonation at C10 to yield **26** (Fig. 4D and Supplementary Figs. 70 and 71). Compound **28** was generated from **IM7-4** via deprotonation at C2 (Fig. 4D).

In summary, these quantum chemical simulations provide robust, energetically feasible pathways for four unprecedented sesterterpene frameworks, including a novel 5/6/5/9 tetracyclic scaffold, the first BFTPS-derived spirocyclic 5/6/5/5/5 scaffold in sesterterpenoids, a structurally unusual 5/6/6/5/5 pentacyclic framework, and a previously unreported 5/6/6/5/6 fused pent acyclic ring system. The divergence between our proposed pathway and previous cyclization hypotheses^75^ may arise from the broad conformational flexibility of carbocation intermediates,^73^ and our DFT results represent one plausible route among multiple possible trajectories. Collectively, these findings underscore the intrinsic complexity of terpene synthase-mediated cyclization.^79, 80^

### Structural and mutational interrogation of active-site residues directing carbocation flux

Previous studies have shown that terpene synthase active-site residues can precisely regulate carbocation migrations, with subtle single-site substitutions frequently pivot product chemotypes.^9, 53, 81^ We therefore reasoned that FsPS and AcPS leverage a similar residue-directed regime to control skeletal rearrangement. To identify the responsible structural determinants, we attempted to express both enzymes in *Saccharomyces cerevisiae* for site-directed mutagenesis. Given that AcPS did not yield detectable products in *S. cerevisiae* even after codon optimization, we focused subsequent mutational and mechanistic studies exclusively on FsPS.

AlphaFold3 modeling revealed that the TC domain of FsPS adopts a canonical class I terpene cyclase fold.^82^ Its catalytic cavity is circumscribed by six aromatic residues (F69, F92, W159, F188, Y217, and W312), four aliphatic residues (I68, V88, A185, and I189), and seven polar or charged residues (S60, H73, D95, D183, N224, E305, and N309) (Supplementary Fig. 72). A USalign-based structural alignment against 38 characterized Type B BFTPSs identified six invariant pocket residues (F92, D95, D183, N224, N309, and W312), while the position corresponding to A185 was occupied exclusively by either Ala or Gly (Supplementary Fig. 73).^83^ Given that these conserved residues are likely associated with general catalytic competence, alanine-scanning mutagenesis was performed on the remaining pocket-lining residues, together with an A185G substitution to assess the role of this naturally variable position.

The majority of these variants completely lost catalytic activity, with the sole exceptions of A185G and E305A, underscoring the acute sensitivity of the FsPS catalytic pocket to spatial perturbations (Fig. 5A and Supplementary Fig. 74). The A185G substitution enhanced total product titers and shifted the major metabolic outcome toward compound **18** (Figs. 5A-i, 5A-ii; and Supplementary Fig. 75), thereby modulating product selectivity without impairing the downstream carbocation rearrangement. Concurrently, the E305A variant exclusively yield **29**, a sesterterpene (*m/z* 358, Figs. 5A-iv and 5A-vi), displaying a GC-MS fragmentation profile highly reminiscent of **18**. Purification and structural elucidation confirmed that **29** shared the same 5/11 ring system as **18** (Fig. 5B, Supplementary Fig. 26; and Supplementary Table 29), confirming its derivation from the un-rearranged bicyclic intermediate **IM3**. These results identified A185 and E305 as key residues affecting FsPS activity, with E305 critically gating downstream skeletal rearrangement.

**Fig. 5.**
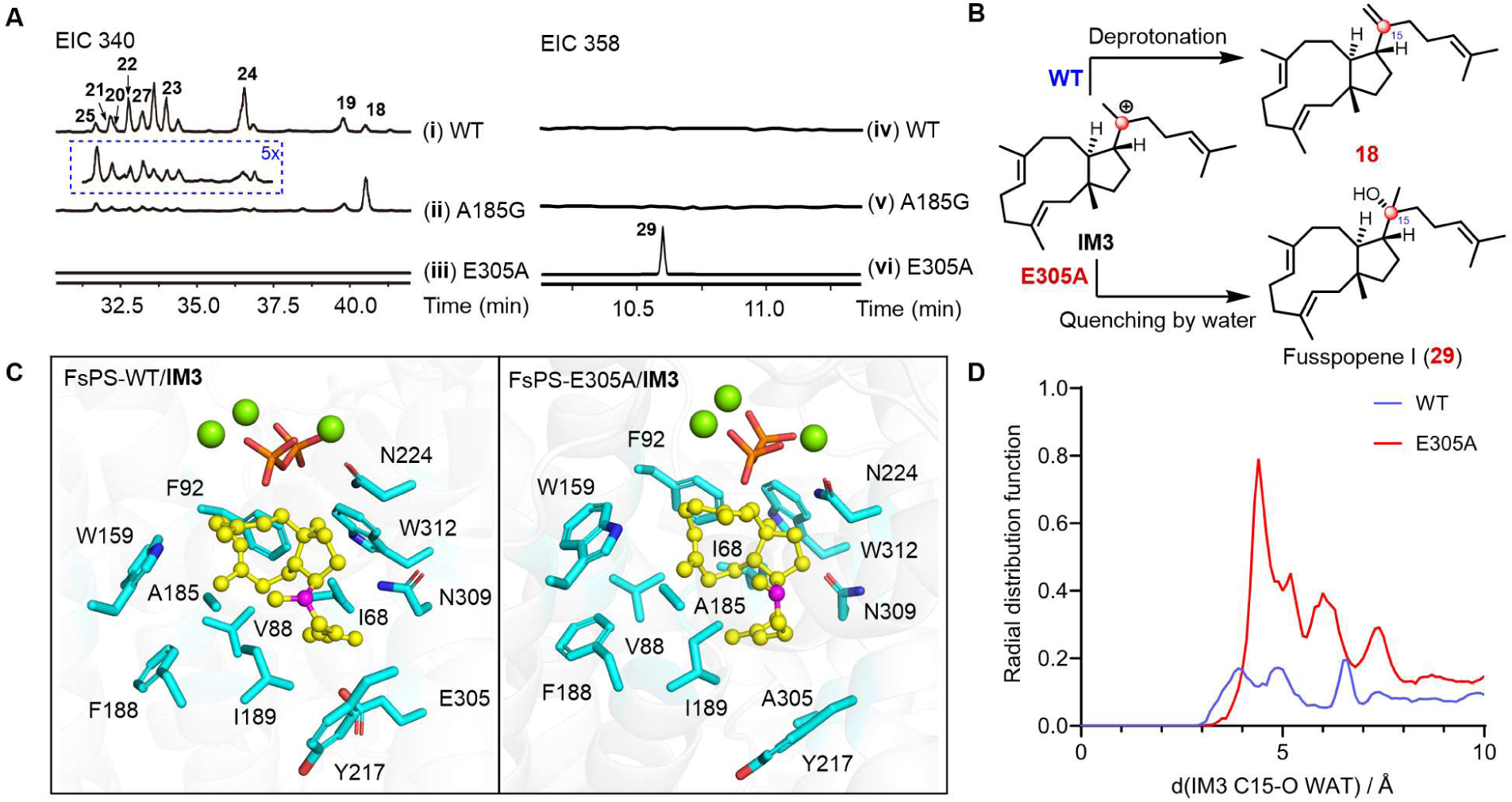
Molecular dynamics simulations and mutational analysis identifying active-site residues that governing carbocation rearrangement in FsPS. (**A**) GC-MS profiles of the wild-type FsPS (FsPS-WT), and its A185G and E305A variants. (**B**) Structural characterization of the newly trapped compound **29**. Structural divergence relative to **18** is demarcated by red circles. (**C**) Representative conformational snapshots extracted from the 100 ns MD trajectories, illustrating the spatial orientation of **IM3** within the active sites of FsPS-WT and FsPS-E305A. The substrate is rendered as a yellow ball-and-stick model, while the key residues are shown as cyan sticks. The Mg^2+^ ions and pyrophosphate (PPi) groups are depicted as green spheres and orange sticks, respectively. (**D**) RDF profiles quantifying water accessibility around the C15 catalytic center in FsPS-WT and the FsPS-E305A variant.

To elucidate the molecular mechanism by which E305 dictates this reaction outcome, **IM3** was docked into the FsPS active site, followed by dual parallel 100 ns MD simulations (Fig. 5C and Supplementary Fig. 76). Per-residue free-energy decomposition by MM-PBSA showed that E305 made the largest favorable contribution to **IM3** binding, primarily driven by its electrostatic component (Supplementary Fig. 77).^84^ We further docked **IM3** into the E305A mutant and performed the same MD simulation analyses (Fig. 5C and Supplementary Figs. 76–78). Compared with the wild type, the E305A variant exhibited significantly enhanced conformational fluctuations for **IM3** (Supplementary Fig. 76), consistent with the loss of the E305-mediated electrostatic stabilization. Radial distribution function (RDF) analysis of ambient water molecules surrounding C15 center revealed a dense solvent population at ∼4 Å in the E305A architecture, whereas the wild-type pocket remained largely water-excluded (Fig. 5D).^85, 86^

These results suggested that E305 anchors **IM3** within a protective hydrophobic shield, limiting deleterious solvent intrusion.^7, 87^ The substitution of this residue destabilizes the target conformation and permits water-mediated quenching of the allylic carbocation, yielding the non-rearranged hydroxylated metabolite. Therefore, E305 does not function as a classical electronic “switch” that actively steers carbocation flux toward rearranged or non-rearranged products. Instead, its disruption fundamentally alters local hydration dynamics, precipitating the premature nucleophilic quenching of transient carbocation intermediates by invading solvent molecules.

## Discussion

Terpenoids represent a major source of structurally diverse and functionally valuable natural products, and their carbon-skeleton diversity is largely generated by terpene synthases through carbocation-mediated cyclization cascades.^5, 6, 7, 88^ Fungal BFTPSs constitute distinctive biosynthetic platforms in which integrated prenyltransferase and terpene cyclase domains couple precursor chain elongation with carbocation-mediated cyclization, enabling the formation of elaborate polycyclic scaffolds.^21^ Despite the rapid expansion of fungal genomic databases, identifying candidates capable of forging novel carbon scaffold remains a primary bottleneck in natural product discovery.^12, 25^ This bottleneck arises because terpene cyclization proceeds over highly sensitive carbocation potential energy surfaces (PES),^89^ where subtle spatial or electrostatic variations within the active site dramatically redirect reaction trajectories without requiring macro-level sequence divergence.^1, 7^ Consequently, traditional sequence-similarity-based approaches frequently fail to resolve these functionally opaque regimes.^90^

While high-throughput heterologous expression and activity screening have expanded the characterized TPS landscape, increasing experimental throughput has yielded diminishing returns in scaffold novelty.^91, 92^ Recent large-scale mining efforts across bacterial and fungal TPSs have yielded vast numbers of terpene products, yet new carbon scaffolds remain exceedingly rare, highlighting that unguided genomic search scales with linear experimental burden rather than structural innovation.^6, 25^ To transcend this screening plateau, machine learning approaches must move beyond sequence-only classification and capture the implicit nonlinear coupling between protein evolutionary constraints and product molecular syntax.^31, 34, 93^ TRACER addresses this challenge by jointly embedding BFTPS sequences (via ESM-C) and product topologies (via ChemBERTa) into a shared multimodal space. By leveraging bi-directional cross-attention, product chemistry guides the interpretation of protein sequence features, enabling TRACER to resolve functional variation associated with product chemotypes and to prioritize, before experimental validation, candidates with high skeletal-rearranging potential. In our experimental validation, TRACER-guided mining of just 18 candidates yielded four unprecedented scaffold classes from FsPS and AcPS, representing a remarkable, ∼25-fold increase in scaffold discovery efficiency compared to non-prioritized empirical screening campaigns. The improvement over sequence-similarity-based assignment and the agreement between predicted and experimentally observed SR/NSR outcomes collectively establish TRACER as a chemically resolved functional framework and enables the rational prioritization of enzymes for exploring underrepresented scaffold space.

Enzyme-function prediction has progressed from sequence-only classification to integrated protein-molecule modeling. DeepECtransformer and CLEAN infer EC functions from individual protein sequences,^14, 94^ whereas ESP predicts enzyme-substrate compatibility from independently encoded protein and molecular representations,^95^ and EZSpecificity incorporates structure-based cross-attention to improve substrate-specificity prediction.^16^ These general-purpose models provide broad coverage across enzyme families but are not optimized to resolve the fine-scale, within-family product divergence characteristic of terpene synthases. In terpene synthase analysis, Terzyme identifies TPSs and prenyltransferases and assigns broad precursor classes,^96^ while EnzymeExplorer combines structural-domain analysis with protein language models to detect sequence-divergent TPSs.^19^ The carbocation-specificity model developed by Durairaj et al. further predicts precursor-cation classes and candidate residues involved in pathway selection in plant sesquiterpene synthases.^8^ These TPS-specific approaches advance enzyme recognition, precursor assignment, and early mechanistic classification, but do not resolve final-product chemotypes or skeletal-rearrangement outcomes. TRACER advances TPS functional prediction to the product level by jointly modeling BFTPSs and their corresponding products, thereby resolving within-family differences in product chemotype and skeletal-rearrangement potential for chemically informed candidate selection. This work extends established protein machine-learning frameworks from general enzyme annotation and substrate compatibility to fine-grained inference of product-defined catalytic outcomes in specialized enzyme families.

Beyond expanding scaffold diversity, our integrated quantum chemical calculations elevate these structural discoveries into a unified mechanistic framework for sesterterpene cyclization. DFT calculations revealed that the structural multiplicity of FsPS and AcPS originates from a central, rearrangement-driven branching node (the 5/6/11-fused carbocation intermediate **IM4** and its downstream isomer **IM7**). Rather than employing distinct, independently evolved cyclization programs, FsPS and AcPS navigate chemical space through exquisite thermodynamic partitioning across this shared intermediate manifold. In particular, the energetically accessible C7–C10/C2–C6 annulation introduces a bond-construction logic distinct from the previously proposed C6–C10 cyclization routes,^67^ whereas the prohibitive barriers of the alternative pathways constrain the chemically accessible branches. Specifically, the energetically favorable asynchronous annulation and ring-rearrangement cascades (Path I and Path II) bypass previously assumed pathways involving prohibitive high energy barriers, providing a clear thermodynamic rationale for the observed molecular topologies. Furthermore, our MD simulations and site-directed mutagenesis demonstrated that active-site residue E305 in FsPS acts as a crucial “hydrophobic gatekeeper.” Instead of directly steering carbocation migration through electrostatic guidance, E305 maintains an active-site water-exclusion zone; its alanine substitution disrupts this protective microenvironment, inducing premature nucleophilic quenching of the transient carbocation by invading solvent molecules.

To rigorously evaluate the boundaries of this paradigm, several critical machine learning and biological limitations must be acknowledged. First, the small size of the curated training dataset (N = 38 Type B BFTPSs) poses an inherent risk of overfitting and limits the model’s ability to predict entirely novel, out-of-distribution chemotypes that lack representation in the training chemical space. While cross-validation metrics were robust, TRACER’s current inference logic relies on pairing query sequences with known product libraries, which may bias predictions toward known scaffold clusters rather than explicitly generating novel molecular geometries. Second, TRACER operates as a functional prioritization tool and does not predict heterologous expression efficiency, catalytic turnover (*k*cat), or quantitative product ratios in host chassis—factors heavily influenced by host-specific metabolic fluxes and protein solubility. Finally, as with many deep learning architectures, TRACER’s latent representations currently lack direct structural interpretability, failing to explicitly pinpoint the precise active-site pocket residues or conformational dynamics associated with specific catalytic outcomes. Future developments should prioritize mitigating these limitations through iterative dataset expansion and structural integration. As demonstrated by our retraining analysis, incorporating newly validated BFTPSs provides functional anchors for unresolved “dark clusters” and sharpens cluster boundaries. To bridge the gap between latent embeddings and structural mechanics, next-generation frameworks should incorporate three-dimensional active-pocket descriptors— specifically targeting the Catalytic Functional Plasticity Zone (CAPZ)—and explicit carbocation potential energy surface parameters into the co-attention module. Combining multimodal representation learning with active-site spatial geometry and quantum mechanical constraints will transform enzyme discovery from predictive classification toward the rational, atomistic engineering of terpene synthases for bespoke chemical architectures.

## Methods

### Dataset construction and genome mining of BFTPS candidates

Putative bifunctional terpene synthase (BFTPS) sequences were retrieved from the NCBI protein database (https://www.ncbi.nlm.nih.gov/protein/?term=fungi) and an in-house fungal genome database using a Hidden Markov Model (HMM)-based search. The in-house database contains draft genome sequences of more than 500 plant endophytic fungi, along with their predicted genes, and was constructed by Prof. Tom Hsiang (University of Guelph, Canada). The HMM profile corresponding to the terpene synthase family (PF01397) was retrieved from the PFAM database (http://pfam.xfam.org) and used as the query, and homologous sequences were extracted using hmmsearch (HMMER v3.1b2)^97^ at an E-value threshold of 1e-5. All retrieved sequences were then filtered for redundancy using CD-HIT,^98^ merging entries with greater than 90% pairwise sequence identity. To eliminate false positives, the remaining sequences were examined for the presence and completeness of the four conserved motifs characteristic of BFTPSs, including DDXXD/E, NSE/DTE, DDXXD, and DDXXN.^1^

### Sequence correction and sequence similarity analyses

Multiple sequences were aligned using the Clustal Omega web server (https://www.ebi.ac.uk/ jdispatcher/msa/clustalo?stype=protein). Alignment figures were generated with ESPript 3.2 (https://espript.ibcp.fr/ESPript/cgi-bin/ESPript.cgi). Putative BFTPS sequences exhibiting annotation abnormalities were corrected through a standardized re-annotation workflow developed locally. First, multiple sequence alignment was used to identify candidates with irregular regions, and for each sequence we determined whether the anomaly arose from redundancy or truncation and recorded the exact amino-acid boundaries of the abnormal fragment. The full biosynthetic gene clusters containing these sequences were then submitted to 2ndFind (https://biosyn.nih.go.jp/2ndfind/) for ORF re-prediction, and the resulting candidate ORFs were compared across several fungal model organisms, including *Aspergillus nidulans*, *A. oryzae*, *Botrytis cinerea*, and *Fusarium* sp., to improve prediction reliability. Based on the nature and position of the anomaly identified during alignment, we selected the ORF that best matched the expected length and preserved the characteristic BFTPS motifs. The corrected sequence was used to replace the original entry and subsequently re-aligned to confirm that the abnormality had been eliminated. Finally, each corrected sequence was evaluated using AlphaFold3 structural prediction to ensure that the resulting protein model displayed complete and coherent BFTPS domain architecture before inclusion in downstream analyses. The sequence identity of BFTPS protein sequences was assessed using BLAST to generate SSN profiles. SSNs were then visualized using Cytoscape.^99^ Maximum-likelihood phylogenetic trees were constructed with MEGA v6.065 with 1,000 bootstrap replicates and subsequently annotated, refined, and visualized using the iTOL web server (https://itol.embl.de/). Three TPSs from the plant (accession number Q9LUE0.1, Q9LIA1.2, and Q9LH31.1) was used as outgroups.

### TRACER architecture

TRACER was designed as a multimodal enzyme-product representation learning model to associate Type B BFTPS sequences with their corresponding product chemotypes. Each enzyme–product pair was represented by two input modalities: the full-length BFTPS amino acid sequence and the SMILES string of the corresponding terpene product. Protein sequences were encoded using the pre-trained ESM-C model (ESMC-600M-202412),^31^ and product SMILES strings were encoded using ChemBERTa-77M.^32^ To standardize the inputs, protein sequences were truncated to a maximum length of 350 residues, and SMILES strings were tokenized with a maximum length of 128 tokens. The ESM-C- and ChemBERTa-derived embeddings were independently processed through parallel multi-branch one-dimensional convolutional neural networks. Convolutional kernels of different sizes were used to capture local feature patterns from protein and product representations, and the resulting features were projected into a shared embedding dimension. The normalized protein and product embeddings were then passed into a bi-directional co-attention module. In this module, protein features and product features were alternately used as queries, keys, and values, allowing TRACER to learn cross-modal associations between enzyme sequence features and product structural features. The co-attention-enhanced protein and product embeddings were fused into a joint representation for downstream prediction. Product identity was first predicted through a fully connected classification head, and the corresponding SR/NSR functional label was assigned using a predefined product-to-label mapping. In parallel, label-level supervision was used to guide SR/NSR classification, and supervised contrastive learning was applied to align enzyme-product pairs with related functional labels in the shared latent space. The final training objective combined product classification loss, label classification loss, and supervised contrastive loss.

### Model training and inference

The curated Type B BFTPS dataset was divided into a training subset and an independent validation subset at an approximately 9:1 ratio. TRACER was trained on 34 experimentally verified enzyme–product pairs and evaluated using four validation enzymes that represented distinct product scaffold classes and rearrangement types. Model performance was assessed using five-fold cross-validation, with accuracy, precision, recall, and F1-score used as classification metrics. Dropout, weight decay, adaptive learning-rate reduction, and early stopping were applied to reduce overfitting. For inference, each uncharacterized BFTPS candidate was paired with all product SMILES representations in the training product library. Candidate enzyme–product pairs were scored using both embedding similarity and the output of the trained joint model. The highest-scoring product was selected as the predicted product chemotype, and its associated SR or NSR label was assigned as the predicted functional class. The learned joint representations were further projected into two-dimensional space using t-SNE for visualization and cluster-based candidate prioritization. Detailed preprocessing procedures, model equations, hyperparameters, and visualization settings are provided in the Supplementary Methods.

### Strains, media, and growth conditions

All wild-type filamentous fungi used in this study were obtained from Prof. Hsiang at the University of Guelph (Ontario, Canada). Detailed species information for these strains is provided in Supplementary Table 9. The fungi were initially cultivated on 2% potato dextrose agar (PDA, Fisher Scientific) at 28 °C for 5 days, and used as a source for BFTPS genes. *A. oryzae* NSAR1 (*niaD*^−^ *sC*^−^ *adeA*^−^ Δ*argB*::*adeA*^−^) (AO) was used as the host for gene cluster expression, and cultured at 30 °C, 200 rpm in DPY (dextrin– polypeptone–yeast extract: 2% dextrin, 1% polypeptone, 0.5% yeast extract, 100 mL) broth supplemented with appropriate nutrients. Transformants of the AO were grown in MPY broth (containing maltose 3%, polypeptone 1%, yeast extract 0.5%, (NH4)2SO4 0.1%, methionine 0.15%, and adenine 0.01%) at 30 °C and 220 rpm for 3 days. *Escherichia coli* DH10b was used for plasmid cloning. *Saccharomyces cerevisiae* mva-y10^100^ (SC) provided by Prof. Xiaozhou Luo at Shenzhen Key Laboratory for the Intelligent Microbial Manufacturing of Medicines (Shenzhen, China) was used as the host for mutants’ expression, and cultured at 30 °C, 220 rpm in YPD broth (20 g/L tryptone, 10 g/L yeast extract, 20 g/L glucose).

### Genomic DNA isolation, plasmid construction, and transformation of AO

Genomic DNA was extracted using the Rapid Fungal Genomic DNA Isolation Kit (Sangon Biotech). All genes encoding BFTPS from wild-type filamentous fungi in this study were amplified from genomic DNA using KOD FX NEO (TOYOBO Life Science) with primer pairs shown in Supplementary Table 30. Each PCR product was inserted into the appropriate restriction site of pUARA4 using ClonExpress^®^II One Step Cloning Kit (Vazyme Biotech) to construct expression plasmids. BFTPS sequences obtained from NCBI were directly synthesized and cloned into the pUARA4 vector by BGI Genomics Co., Ltd. (Shenzhen, China). All plasmids used in this experiment are listed in Supplementary Table 30. To construct transformant AO-*BFTPS*, protoplasts of AO were prepared using the protoplast-polyethylene glycol method as described in a previous study.^39^ The AO transformants listed in Supplementary Table 31 were cultivated on MPY medium following a previously described method.^39^

### Transformation of *S. cerevisiae* expressing FsPS and its mutants

*FsPS* and its mutants were constructed into plasmids pXW55 and introduced into *S. cerevisiae* mva-y10 (Supplementary Table 30 and 31). Subsequently, the resulting transformants were cultured in 50 mL of uracil-dropout broth (20 g/L glucose, 5.0 g/L casamino acids, 0.02 g/L adenine, 6.7 g/L YNB, 0.02 g/L tryptophan, pH 7.5) for 48 hours at 30 °C with shaking at 220 rpm. And then, 10 mL of culture were inoculated into each 3 L flasks containing 1L of YPD medium and fermentation proceeded at 30°C with shaking at 220 rpm for 3 days.

### GC-MS analysis of metabolites derived from AO transformants and SC transformants

To analyze the metabolites produced by each AO transformant and each SC transformant. After fermentation, the cells were collected by centrifugation at 3,000 rpm and subjected to acetone extraction assisted by ultrasonication for 30 min. The resulting mixture was centrifuged again at 3,000 rpm to separate the supernatant, which was then concentrated under reduced pressure. The resulting residue was reconstituted in a mixture of water and ethyl acetate. The ethyl acetate-soluble fraction was collected and evaporated under vacuum to obtain the crude extract. The crude extracts were dissolved in ethyl acetate at a final concentration of 50 mg/mL and subjected to GC-MS analysis using a SHIMADZU DB-5 ms capillary column (0.25 mm × 30.0 m, 0.25 μm film thickness). The initial temperature was 60 °C, then increased at a rate of 25 °C/min to 280 °C, followed by a ramp of 10 °C/min to 310 °C. The MS settings included a source temperature of 230 °C and an electron energy of 0.4 kV. To achieve improved separation of structurally diverse terpene products generated by FsPS and AcPS, an optimized GC-MS temperature program was additionally employed. The oven temperature was initially set at 60 °C, increased at 10 °C/min to 165 °C, then ramped at 1.5 °C/min to 200 °C and held for 6 min. Subsequently, the temperature was raised at 4 °C/min to 240 °C, followed by a final ramp of 12 °C/min to 310 °C, where it was maintained for 5 min. This multi-step program enabled enhanced resolution of closely eluting terpene isomers and was therefore used for detailed product profiling of FsPS and AcPS crude extracts.

### TDDFT-ECD, ^13^C NMR and DFT mechanistic calculations

The electronic circular dichroism (ECD) calculations were conducted using the Gaussian 09 program package.^101^ The conformational analyses were carried out via random searching in the Sybyl-X 2.0 software using the MMFF94 force field with an energy window of 15 kcal/mol. The geometries were optimized at the B3LYP/6-31G(d) level in Gaussian 09 software to afford the energy minimized conformers. The ECD spectra for each stable conformer was then computed using time-dependent density functional theory (TDDFT) at the B3LYP/6-31G(d,p) level, incorporating the IEFPCM solvation model with acetonitrile as solvent. The simulated ECD curves were weighted based on the Boltzmann distribution of individual conformers.

Optimized conformers within an energy window of 5 kcal/mol, with no imaginary frequencies, were selected for gauge-independent atomic orbital (GIAO) calculations of ^13^C NMR chemical shifts using density functional theory (DFT) at the B3LYP/6-311+G(d,p) level, incorporating the PCM solvation model. The resulting ^13^C NMR shielding constants of individual conformers were Boltzmann weighted based on their relative Gibbs free energies. Agreement between experimental and calculated ^13^C NMR data was assessed using the improved probability DP4^+^ method.^102^

The cyclization pathways of compounds **18**−**28** were investigated via DFT calculations using the Gaussian 09 program.^101^ Structure optimizations were done at the M06-2X level in the gas phase using the 6-31+G(d,p) basis set.^103^ The vibrational frequency calculations at the same level of theory with optimization were performed to verify that each local minimum has no imaginary frequency and that each TS has only a single imaginary frequency. Single-point energies were calculated at the mPW1PW91/6-311+G(d,p) level based on the optimized structure by using the M06-2X method.^104^ The Gibbs free energy used for discussion in this study was calculated by adding the gas-phase Gibbs free energy correction.

### Molecular docking and dynamic simulation

The TC domains of FsPS and FsPS-E305A models were reconstructed with the substrate and the 3Mg^2+^ coordination shell. The protein structures of TC domains of FsPS and FsPS-E305A were predicted using AlphaFold3.^82^ Subsequent evaluation using the SAVES server showed that the Ramachandran plots of the FsPS and FsPS-E305A models contained 94.2% and 92.4% of residues in the most favored regions, respectively, supporting the overall reliability of the predicted structures (Supplementary Figs. 72 and 78).^105^ The intermediate **IM3** was docked into FsPS and FsPS-E305A using AutoDock Vina.^106^ Partial charges of all **IM3** atoms were derived using the restrained electrostatic potential (RESP) fitting scheme based on electrostatic potentials calculated at the HF/6-31G(d) level with the Gaussian 09 software package.^101^ Ligand topology and parameter files were constructed using the sobtop program,^107^ while protein structures were protonated with the aid of the pdb2pqr web server.^108^ Molecular dynamics simulations were carried out employing the amber99sb-ildn force field to describe interatomic interactions. The simulated system was placed in a periodic rectangular box with periodic boundary conditions applied along all three spatial dimensions and explicitly solvated using TIP3P water molecules. System neutrality was achieved by the addition of appropriate counterions (Na^+^ and Cl^-^). Prior to equilibration, energy minimization was conducted to remove unfavorable steric contacts within the system. Subsequently, equilibration simulations were performed under both NVT and NPT ensembles for 200 ps at a target temperature of 298.15 K. During these stages, covalent bonds involving hydrogen atoms were constrained using the LINCS algorithm. Long-range electrostatic interactions were treated using the Particle Mesh Ewald (PME) approach with a real-space cutoff of 10 Å. Temperature control was maintained via the V-rescale thermostat, while pressure regulation was achieved using the Parrinello-Rahman barostat.^109^ Following equilibration, two independent 100 ns production molecular dynamics simulations were performed for each system with an integration time step of 2 fs. Trajectory coordinates were saved at 100-step intervals. All simulations were executed using the GPU-accelerated version of GROMACS 2020.3 (http://manual.gromacs.org/2020.3/index.html).

### Reporting summary

Further information on research design is available in the nature portfolio reporting summary linked to this article.

## Data availability

Detailed information on reagents, methodologies, experimental procedures, GC traces, NMR spectra, mass spectrometric data, and DFT data is provided in the Supporting Information. Crystallographic data for compounds **19**, **22**, **25**, and **27** have been deposited with the Cambridge Crystallographic Data Centre (CCDC) under deposition numbers 2435241, 2504628, 2504621, and 2504622, respectively. Protein sequences reported in this study are provided in the Supplemental Information. The custom code developed for this study is publicly available on GitHub at https://github.com/Lvkangjie/TRACER-BFTPS.

## Acknowledgments

This work was supported by the National Key Research and Development Program of China (2025YFA0923200), the National Natural Science Foundation of China (82504636, 21977029 and 32121005), Shanghai Sci-Tech Inno Center for Infection & Immunity (SSIII-2024A01 and SSIII-2024A02), Shanghai Municipal Science and Technology Major Project, the 111 Project (B18022), and Syngenta Ph.D. fellowship for Kangjie Lv (SPF182).

## Author contributions

Conceptualization, X.L., L.J., and Y.L.; methodology, K.L. W.Z.; investigation, C.X., K.L., Y.C., K.L. and B.Z.; writing – original draft, K.L.; writing – review & editing, X.L., L.J., Y.L., C.X., W.Z., G.Z., S.M.S., X.Z., and T.H.; funding acquisition, X.L., L.J., L.Z., Y.G., and Y.W.G.; resources, H.O., and T.H.; supervision, X.L., L.J., and Y.L.

## Competing interests

The authors declare no competing interests.

## Notes

### Competing Interest Statement

The authors have declared no competing interest.

