## Supplemental Data 1 for "TRACER navigates rearrangement-driven sesterterpene chemical space via multimodal enzyme-product representation learning"

### Supplemental Materials and Methods

#### 1 General

NMR spectra were acquired on an Agilent DD2 (Santa Clara, CA, USA) 600 MHz spectrometer. Chemical shifts were calibrated internally against the residual signal of the solvent in which the sample was dissolved (benzene-$d_6$ $\delta_H$ 7.16 and $\delta_C$ 128.06; pyridine-$d_5$ $\delta_H$ 8.74, 7.58, 7.22 and $\delta_C$ 150.35, 135.91, 123.87), and all deuterated solvents were from Cambridge Isotope Laboratories (CIL). HR-EI-MS measurements were obtained on Waters GCT Premier mass spectrometer or Waters ACQUITY QDa. RP-HPLC was performed on an Agilent 1260 Series separation module with a diode array detector. Semipreparative HPLC was carried out using an ACE C18-PFP column (10 × 250 mm, 5 μm), a Cosmosil cholester column (10 × 250 mm, 5 μm), and a Phenomenex C18 column (10 × 250 mm, 5 μm). GC-MS measurements were performed on a QP2010SE GC-MS (Shimadzu, Kyoto, Japan). The CD spectra were recorded on a Chirascan circular dichroism spectrometer using acetonitrile (ACN) as solvent. Optical rotation was measured using a Rudolph Research Analytical Autopol IV Automatic Polarimeter. X-ray crystallographic analysis was carried out on a Bruker SMART APEX II diffractometer employing Cu *Kα* radiation. Biological reagents, chemicals, and media were purchased from standard commercial sources unless stated otherwise.

#### 2 Machine learning model architecture

##### 2.1 General computational details

All machine-learning experiments were conducted using Python 3.10 and the PyTorch 2.2 deep-learning framework, with CUDA-enabled GPU acceleration. Protein sequence representations were generated with the ESM C model (ESMC-600M-202412),[1] implemented via the official esm package, while small-molecule embeddings were obtained with ChemBERTa-77M[2] from the HuggingFace Transformers library. Data handling and preprocessing were performed using pandas 2.2, NumPy 1.26, and scikit-learn 1.5. The supervised contrastive loss was implemented using the pytorch-metric-learning package (v2.3). Visualization and clustering analyses utilized matplotlib 3.8, scikit-learn, and KernelDensity for density estimation. All model training and inference were executed on a workstation equipped with an NVIDIA GeForce RTX 3070 Ti GPU (8 GB), a 13th Gen Intel(R) Core(TM) i5-13490F CPU @ 2.50 GHz, and 32 GB system memory. All GPU-accelerated operations, including forward and backward passes, gradient updates, and convolution computations, were performed on the RTX 3070 Ti, leveraging CUDA (v12.2) for GPU parallelization.

##### 2.2 Data loading and preprocessing for machine learning

The curated dataset included four essential fields: protein name, amino acid sequence, small molecule SMILES strings, and functional category labels. Raw CSV data were read using the pandas library, with missing SMILES entries replaced by empty strings. Protein sequences were subjected to stringent cleaning: non-standard amino acid characters were removed, only uppercase letters A–Z were retrained, sequences were truncated to 350 residues to standardize input length and maintain computational efficiency. Small molecule SMILES were tokenized using the ChemBERTa tokenizer, with a maximum token length of 128; longer sequences were truncated, and shorter sequences padded to ensure uniform input dimensions. Functional category labels were transformed into numerical form using Scikit-learn's LabelEncoder. In addition, a comprehensive mapping system was constructed to facilitate label-based tasks: (i) a one-to-one mapping from small molecules to functional categories, ensuring each molecule uniquely corresponds to a label; (ii) a one-to-many mapping from labels to small molecules, reflecting that each functional category may include multiple molecules; and (iii) a small molecule-to-index mapping to support downstream classification and embedding retrieval operations.

##### 2.3 Protein and small molecule feature input and extraction

A dual-stream architecture independently extracted features from proteins and small molecules while maintaining alignment in a shared embedding space. Protein sequences were embedded using the ESM C pretrained model (ESMC_600M_202412, 600M parameters), yielding hidden states of dimension $(B, L, 1152)$, where $B$ is batch size and $L$ sequence length, then transposed to $(B,\ 1152,\ L)$ for convolutional processing. Multi-scale 1D convolutional neural networks with kernel sizes 3, 5, and 7 were applied, each containing two convolution layers with batch normalization and ReLU activations, followed by adaptive max pooling. Outputs were concatenated and projected to a fusion dimension of 1024 via a linear transformation and L2 normalization. Small molecule SMILES embeddings were processed analogously using the ChemBERTa-77M model, generating normalized embeddings through the same multi-scale CNN pipeline. This symmetric design ensured that both modalities reside in the same semantic embedding space, enabling meaningful cross-modal interactions in subsequent attention modules.

#### 2.4 Co-attention mechanism

Input to the co-attention module consisted of normalized protein and small molecule feature vectors, each with a dimensionality of 1024. Multi-head attention with eight parallel attention heads was applied to capture bidirectional interactions between the two modalities. Protein and small molecule embeddings were independently projected into a shared hidden space of size 1024 using linear transformation layers:

$$\tilde{P} = W_P P + b_P, \tilde{M} = W_M M + b_M$$

where $W_P$, $W_M \in \mathbb{R}^{1024\times1024}$ and $b_P$, $b_M \in \mathbb{R}^{1024}$. For each attention head, queries $Q$, keys $K$, and values $V$ are computed, and scaled dot-product attention is applied:

$$\text{Attention}(Q, K, V) = \text{Softmax}\left(\frac{QK^\top}{\sqrt{128}}\right)V$$

with a key dimension of 128 (1024 features divided across eight heads). Two directional flows were calculated: protein features as queries with small molecule features as keys and values, and small molecule features as queries with protein features as keys and values. Residual connections and layer normalization (with epsilon = 1e-5) were applied to stabilize training, and a dropout rate of 0.4 is used within attention layers to prevent overfitting. Global pooling was performed before co-attention to compress sequences into single-vector representations, reducing computational complexity for pairwise interactions. The output embeddings retained a dimensionality of 1024 for both proteins and small molecules, enriched with cross-modal contextual information while preserving the original feature size, ensuring seamless integration with downstream fusion and classification layers.

#### 2.5 Feature fusion and classification

Co-attention enhanced embeddings were concatenated along the feature dimension ($\mathrm{F}_{\text{joint}} = [\widehat{\mathrm{P}}; \widehat{\mathrm{M}}]$) and passed through a two-layer fully connected network with intermediate ReLU activations and Dropout (p = 0.3) to prevent overfitting. The joint embedding was adaptively weighted:

$$E_{\text{joint}} = \alpha \cdot F_{\text{protein}} + (1 - \alpha) \cdot F_{\text{molecule}}$$

where $\alpha \in [0,1]$ is a learnable parameter constrained via Sigmoid. Small molecule classification was performed using a two-layer fully connected network generating logits for each category. Protein function predictions were indirectly obtained by mapping small molecule classification probabilities through a pre-constructed label matrix, enabling protein functional inference without direct protein-to-label supervision.

#### 2.6 Multi-objective loss function

We employed a multi-objective optimization framework, combining small molecule classification loss $\mathcal{L}_{\text{smiles}}$, label-level auxiliary supervision loss $\mathcal{L}_{\text{label}}$, and supervised contrastive loss $\mathcal{L}_{\text{supcon}}$. The total loss is defined as:

$$\mathcal{L}_{\text{total}} = \lambda_1 \mathcal{L}_{\text{smiles}} + \lambda_2 \mathcal{L}_{\text{label}} + \lambda_3 \mathcal{L}_{\text{supcon}}$$

where $\lambda_1$, $\lambda_2$, and $\lambda_3$ are tunable weight coefficients controlling the relative influence of each task. These hyperparameters were empirically tuned ($\lambda_1$ = 0.5, $\lambda_2$ = 0.2, $\lambda_3$ = 1.0) to balance the contributions of structural recognition, functional categorization, and representation alignment, preventing any single objective from dominating the optimization.

Small-molecule classification employs standard categorical cross-entropy:

$$\mathcal{L}_{\text{smiles}} = -\sum_{c=1}^{C} \mathrm{y_c} \log \mathrm{p_c}$$

with $C$ denoting the number of unique molecules, $\mathrm{y_c}$ the one-hot ground truth vector, and $p_c$ the predicted probability for class $c$.

Protein functional prediction was computed indirectly by projecting small-molecule logits through a fixed molecule–label mapping matrix, followed by cross-entropy against the true protein label distribution:

$$\mathcal{L}_{label} = -\sum_{k=1}^{K} y_k \log p_k$$

where $K$ denotes the number of functional categories, $y_k$ is the one-hot encoded ground-truth label, and $p_k$ is the predicted class probability.

To align protein and small-molecule representations in a shared embedding space, we applied a supervised contrastive objective. For each batch containing $N$ samples, protein embeddings and their paired molecular embeddings were concatenated to form a set of $2N$ contrastive samples. The loss was calculated as follows:

$$\mathcal{L}_{\text{supcon}} = \sum_{n=1}^{2N} \frac{-1}{|P(n)|} \sum_{p\in P(n)} \log \frac{\exp(\text{sim}(z_n, z_p)/\tau)}{\sum_{a\in A(n)} \exp(\text{sim}(z_n, z_a)/\tau)}$$

where $z_n$ denotes the embedding of sample $n$, $z_p$ is a *positive* embedding sharing the same functional label, $P(n)$ is the set of *positive* samples for $n$, $A(n)$ is the set of all contrastive samples excluding $n$, and $\tau$ is the temperature hyperparameter. This formulation explicitly pulls protein–molecule pairs of the same class closer while pushing apart pairs of different classes, strengthening the discriminability of the integrated representation space.

#### 2.7 Training optimization strategy

Model training used a five-fold stratified cross-validation scheme to preserve category distributions across folds, ensuring robust evaluation under class imbalance. Optimization was performed using the AdamW optimizer with an initial learning rate of $1\times10^{-5}$ and weight decay $1\times10^{-4}$, which decouples weight decay from gradient updates to control model complexity and mitigate overfitting. Dynamic learning rate decay halved the learning rate when validation loss stagnated for 10 epochs, guiding convergence toward flatter minima that can better generalize. Additional regularization was provided by Dropout layers and L2 normalization in feature extraction modules, reducing neuron co-adaptation and constraining feature magnitude.

Early stopping was applied with a patience of 100 epochs after an initial 100-epoch training period. Validation loss was monitored throughout training, and the patience counter was reset whenever a new minimum validation loss was reached. Training was stopped if no further improvement in validation loss occurred for 100 consecutive epochs after epoch 100, and the model parameters corresponding to the global minimum validation loss were retained. This strategy helps reduce overfitting by preventing continued optimization after validation performance has plateaued or begun to deteriorate. A batch size of 32 balanced gradient stability with computational efficiency. This combination of stratified cross-validation, adaptive learning rate scheduling, regularization, and calibrated early stopping effectively prevented overfitting while ensuring stable convergence in the multi-task, high-dimensional embedding space.

#### 2.8 Prediction inference

During inference, functional label prediction was performed using a training-library-restricted two-stage reranking strategy. The candidate small-molecule library was constructed from the SMILES entries observed during model training. Each candidate SMILES was first tokenized and passed through the ChemBERTa-based molecular encoder followed by the trained molecular CNN adapter to obtain a normalized molecular embedding, denoted as $\mathbf{f}_{SM}$. For a query protein sequence, the ESM-CNN protein encoder was used to extract a normalized protein embedding, denoted as $\mathbf{f}_{\mathrm{Pr}}$ .

For each query protein, all candidate SMILES in the training library were evaluated. First, the query protein embedding was compared with each candidate molecular embedding in the shared representation space. The similarity between the query protein and each candidate SMILES was calculated using cosine similarity:

$$\mathrm{sim}(\mathbf{f}_{Prot}, \mathbf{f}_{\mathrm{SM}}) = \frac{\mathbf{f}_{Prot} \cdot \mathbf{f}_{\mathrm{SM}}}{\parallel \mathbf{f}_{Prot} \parallel \parallel \mathbf{f}_{SM} \parallel}$$

This similarity score reflects the proximity between the query protein and each candidate molecule in the learned protein–molecule embedding space.

Next, each candidate SMILES was individually paired with the query protein and passed through the complete protein–molecule joint model. Specifically, the protein and candidate molecule representations were processed by the co-attention module and feature fusion network, followed by the trained SMILES classification head. For each candidate SMILES, a self-consistency score was calculated as the softmax probability assigned by the classifier to that candidate's own SMILES class. This score measures the extent to which the full joint model supports the compatibility of the query protein–candidate SMILES pair.

The final score for each candidate SMILES was computed by integrating the embedding-based similarity and the classifier-derived self-consistency score:

$$\mathrm{S}_{joint} = \mathrm{w}_r \cdot \mathrm{sim}(\mathbf{f}_{Prot}, \mathbf{f}_{SM}) + w_c \cdot \mathrm{P}_{self}(\mathrm{SMILES}_i)$$

where sim $(\mathbf{f}_{\mathrm{Prot}}, \mathbf{f}_{SM})$ denotes the embedding similarity between the query protein and candidate $SMILES_i$, $P_{self}(SMILES_i)$ denotes the self-consistency probability assigned to the candidate's own SMILES class, and $w_r$ , and $w_c$ are the weighting coefficients for retrieval similarity and self-consistency, respectively. The candidate SMILES with the highest joint score across the entire training-library candidate set was selected as the predicted product for the query protein.

The final functional label was not produced by an independent label classification head. Instead, after the optimal SMILES candidate was selected, its functional label was assigned according to the predefined SMILES-to-label mapping established during training. Thus, the inference process first identified the most compatible training-library SMILES through exhaustive candidate scoring and then derived the functional label from the selected product identity. This design kept functional label assignment directly coupled to the predicted product SMILES and its associated functional category.

#### 2.9 Visual analysis

Feature space quality was quantitatively assessed using three classical clustering metrics. The silhouette coefficient measures intra-cluster compactness relative to inter-cluster separation, providing an intuitive estimate of cluster cohesion. The Davies-Bouldin index evaluates the average ratio of intra-cluster distances to inter-cluster separation, where lower values indicate better clustering. The Calinski-Harabasz index quantifies the ratio of between-cluster dispersion to within-cluster dispersion, favoring well-separated and dense clusters. Together, these metrics offered complementary perspectives on the organization of high-dimensional embeddings.

High-dimensional feature embeddings were projected to two-dimensional space using t-SNE (perplexity 30, early exaggeration 8, learning rate 200, 3000 iterations) to facilitate visualization of complex multimodal relationships. To reduce point overlap, adaptive jittering added small perturbations sampled from a normal distribution proportional to local density. Density estimation is performed with a Gaussian kernel (bandwidth 0.5) to compute probability distributions across the projected space. Low-density regions below a threshold of 0.25 were highlighted as class boundaries, often corresponding to samples with higher classification uncertainty. Final visualizations integrate scatter plots, contour lines, and boundary annotations, enabling intuitive evaluation of both cluster structure and classification reliability while illustrating the discriminative capability of the joint protein-product embeddings.

#### 3 Isolation of compounds 1−2, 8 and 15−29

To isolate individual metabolites, AO transformants were cultured in MPY medium at 30 °C for 3 days in 3 L Erlenmeyer flasks, each containing 1 L of medium. After fermentation, the culture broth from each flask was transferred to a 1 L centrifuge bottle and centrifuged at 9,000 rpm for 10 minutes to collect the cell pellets. The pellets were extracted three times with acetone at room temperature, and the combined extracts were concentrated *in vacuo*. The resulting residues were further extracted with ethyl acetate. The crude extracts were fractionated by silica gel column chromatography using hexane as the eluent and subsequently purified by preparative HPLC. Detailed purification procedures for each compound are described below.

After cultivation of transformants AO-*OaFS* in 5 L of MPY and AO-*DqFS* in 10 L of MPY, crude extracts (0.7 g and 2.0 g, respectively) were obtained. Each extract was subjected to silica gel column chromatography using hexane as the mobile phase. A fraction (23.7 mg) that contained compound **1** was further purified by semipreparative HPLC on a Cholester column (10 × 250 mm, 5 µm), using ACN-$H_2O$ (95:5, *v*/*v*) as the mobile phase at a flow rate of 4.0 mL/min to yield compound **1** (2.6 mg).

The 20 L MPY fermentation broth of the AO-*BbOS* transformant afforded 1.6 g of crude extract. This extract was subjected to silica gel column chromatography using hexane as the mobile phase to yield fractions Fr.1–Fr.3. Fr.1 (36.1 mg) was further purified by semipreparative HPLC on an ACE C18-PFP column (10 × 250 mm, 5 µm), eluted with 100% ACN at a flow rate of 4.0 mL/min to afford compound **2** (2.7 mg).

A total of 2.0 g of crude extract was obtained from AO-*FpMS* cultured in 20 L of MPY medium. The extract was subjected to silica gel column chromatography with hexane as the eluent, yielding eight fractions designated Fr.1 to Fr.8. Fr.1 (19.3 mg), which were further purified by semipreparative HPLC using an ACE C18-PFP column (10 × 250 mm, 5 µm) eluted with 100% ACN at a flow rate of 4.0 mL/min, resulting in the isolation of compound **8** (1.8 mg).

The AO-*AlSS* transformant was fermented in 20 L of MPY to afford a crude extract (1.3 g). The extract was first subjected to silica gel column chromatography using hexane as the mobile phase to achieve preliminary fractionation. The fractions (85.7 mg) containing the target metabolite were further purified by semipreparative HPLC on an ACE C18-PFP column (10 × 250 mm, 5 µm) with isocratic elution of ACN-$H_2O$ (90:10, *v*/*v*) at a flow rate of 4.0 mL/min to obtain compound **15** (6.6 mg).

The AO-*AtVS* strain was cultured in 18 L of MPY, affording a crude extract (1.5 g), which was initially fractionated by silica gel chromatography with hexane as the eluent. The fractions (21.2 mg) containing **16** were subsequently subjected to semipreparative HPLC on an ACE C18-PFP column (10 × 250 mm, 5 µm) using 100% ACN as the mobile phase at a flow rate of 4.0 mL/min to yield compound **16** (1.7 mg).

The AO-*CrNS* strain was fermented in 10 L of MPY medium to yield a crude extract (2.3 g). This extract was first separated by silica gel chromatography using hexane as the mobile phase. The fractions (18.9 mg) containing **17** were further purified by semipreparative HPLC on a Phenomenex C18 column (10 × 250 mm, 5 µm) using a MeOH-$H_2O$ gradient (0–60 min, 85:15 to 99:1, *v*/*v*) at a flow rate of 3.8 mL/min, affording compound **17** (2.6 mg).

The crude extract (30.5 g) from the AO-*FsPS* strain cultured in 100 L of MPY was subjected to silica gel column chromatography using hexane as the eluent. Based on GC-MS analysis, the target fractions were combined (262.4 mg) and further separated by semipreparative HPLC on an ACE C18-PFP column (10 × 250 mm, 5 µm), eluted with ACN-$H_2O$ at a flow rate of 3.8 mL/min using the following gradient program: 0–60 min, 85:15 to 99:1; 60–120 min, 99:1 (*v*/*v*), yielding five fractions (Fr.1–Fr.5). Fr.2 (11.2 mg) was further purified on a Phenomenex C18 column (10 × 250 mm, 5 µm) using MeOH-$H_2O$ (93:7, *v*/*v*) at a flow rate of 3.8 mL/min to obtain compound **18** (1.1 mg). Fr.3 (9.3 mg) was purified on an ACE C18-PFP column (10 × 250 mm, 5 µm) with ACN-$H_2O$ (85:15, *v*/*v*), affording compound **19** (1.8 mg). Fr.4 (20.7 mg) was separated on a Phenomenex C18 column (10 × 250 mm, 5 µm) using a MeOH-$H_2O$ gradient (0–60 min, 90:10 to 99:1, *v*/*v*) at a flow rate of 3.8 mL/min, yielding compound **20** (0.5 mg) and compound **24** (2.2 mg). Fr.5 (35.0 mg) was fractionated on a Phenomenex C18 column (10 × 250 mm, 5 µm) with MeOH-$H_2O$ (93:7, *v*/*v*) at 3.8 mL/min into four subfractions (Fr.5.1–Fr.5.4). Fr.5.1 (3.5 mg) was further purified by semipreparative HPLC on a Phenomenex C18 column

(10 × 250 mm, 5 μm) using ACN-$H_2O$ at a flow rate of 1.0 mL/min with the following gradient program: 0–60 min, 92:8 to 96:4; 60–90 min, 96:4 to 99:1; 90–140 min, 99:1 (*v*/*v*), affording compound **23** (0.6 mg). Fr.5.2 (2.1 mg) was also purified on a Phenomenex C18 column (10 × 250 mm, 5 μm) using the method as Fr.5.1, affording compound **25** (0.4 mg). Fr.5.3 (7.4 mg) was separated on a Phenomenex C18 column (10 × 250 mm, 5 μm) using ACN-$H_2O$ at a flow rate of 1.0 mL/min with the following gradient program: 0–60 min, 92:8 to 96:4; 60–90 min, 96:4 to 99:1; 90–160 min, 99:1 (*v*/*v*), yielding compound **21** (1.0 mg) and compound **22** (0.1 mg). Fr.5.4 (3.3 mg) was purified on a Phenomenex C18 column (10 × 250 mm, 5 μm) using MeOH-$H_2O$ (92:8, *v*/*v*) at a flow rate of 1.0 mL/min, affording compound **27** (1.1 mg).

Fermentation of the AO-*AcPS* transformant in 40 L of MPY afforded 13.5 g of crude extract. The extract was initially fractionated on a neutral silica gel column using hexane as the mobile phase, and fractions containing the target metabolites were identified by GC-MS analysis. These fractions were subsequently subjected to semipreparative HPLC on a Phenomenex C18 column (10 × 250 mm, 5 μm), using ACN-$H_2O$ at a flow rate of 3.8 mL/min with the following gradient program: 0–60 min, 96:4 to 98:2; 60–120 min, 98:2 (*v*/*v*), to yield eight fractions (Fr.1–Fr.8). GC–MS analysis confirmed that the target compounds were enriched in Fr.2–Fr.8. Fr.2 (3.9 mg) and Fr.3 (23.3 mg) were separately purified by semipreparative HPLC using a gradient of MeOH-$H_2O$ (0–60 min, 90:10 to 95:5, *v*/*v*) at a flow rate of 3.8 mL/min on a Phenomenex C18 column and an ACE C18-PFP column, respectively, affording compound **18** (0.8 mg) and compound **19** (4.8 mg). Fr.5 (5.8 mg) was purified on an ACE C18-PFP column (10 × 250 mm, 5 μm) under isocratic elution with MeOH-$H_2O$ (87:13, *v*/*v*) at a flow rate of 3.8 mL/min, providing compound **24** (2.0 mg). Fr.9 (74.3 mg) was purified on an ACE C18-PFP column (10 × 250 mm, 5 μm) using MeOH-$H_2O$ (92:8, *v*/*v*) as the mobile phase at a flow rate of 3.8 mL/min, affording compound **22** (2.0 mg), compound **26** (4.0 mg), and compound **28** (1.6 mg).

FsPS-E305A was expressed in *S. cerevisiae* mva-y10 and subjected to 80 L fermentation in YPD. After extraction of the fermentation broth, 6.53 g of crude extract was obtained and initially fractionated on a neutral silica gel column using hexane as the mobile phase. Target fraction (Fr.1) was collected based on GC-MS analysis. Fr.1 (2.3 g) was further separated on an ACE C18-PFP semi-preparative column at a flow rate of 3.5 mL/min using a gradient of MeOH-$H_2O$ (0–60 min, 85:15 to 100:0, *v*/*v*), affording fractions Fr.1.1–Fr.1.6. Fraction Fr.1.1 (64.5 mg) was subsequently purified by semi-preparative HPLC using a Phenomenex C18 column with isocratic elution of ACN-$H_2O$ (85:15, *v*/*v*), yielding compound **29** (1.8 mg).

#### 4 Structural elucidation of compounds 1−2, 8 and 15−29

Compounds **1, 2, 8, 15, 16** and **19** were respectively identified as fusicoccadiene (**1**),[3, 4, 5, 6] ophiobolin F (**2**),[7] mangicdiene (**8**),[8, 9, 10] spiroviolene (**15**),[11, 12, 13] variediene (**16**),[14, 15, 16] and astelifatriene (**19**)[17] by comparison of their GC-MS and 1D NMR spectra with literature data (Figures S9−S13 and S16, Tables S10−S14 and S17). A single crystal of **19** suitable for X-ray diffraction analysis was obtained by repeated crystallization from MeOH and Ethyl acetate (1:1, *v*/*v*) at -20 °C. The absolute configuration of **19** was subsequently established by single-crystal X-ray diffraction using Cu *Kα* radiation [Flack parameter of 0.1(2)] (Figure S58 and Table S27, CCDC 2435241). Consequently, the absolute configuration of **19** was definitively established as 10*S*,11*R*,14*S*,15*S*,18*S* by comparing the experimental and calculated ECD spectra (Figures S32 and S33), confirming its identity as astellifatriene.

Compound **17** was obtained as a colorless oil. Its molecular formula was established as $C_{25}H_{40}$ based on HR-EI-MS data, which exhibited a molecular ion peak at *m*/*z* 340.3131 $[M]^+$ (calcd for $[C_{25}H_{40}]^+$, 340.3125), corresponding to six degrees of unsaturation. Comprehensive analysis of the $^1H$ NMR, $^{13}C$ NMR, and HSQC spectra revealed the presence of 25 carbon signals, including six methyl groups, eight methylene groups, two methine carbons, eight olefinic carbons, and one *sp*$^3$ quaternary carbon. Considering the presence of four double bonds accounting for four degrees of unsaturation, the remaining two degrees indicated that compound **17** possesses a bicyclic ring system. The $^1H$–$^1H$ COSY spectrum established four distinct spin systems: H-1−H-2 (**I**), H-4−H-5−H-6 (**II**), H-8−H-9−H-10−H-14−H-13−H-12 (**III**), and H-16−H-17−H-18 (**IV**). These fragments were further connected through key HMBC correlations, including those from $H_3$-20 to C-2/C-3/C-4, $H_3$-21 to C-6/C-7/C-8, $H_3$-22 to C-1/C-10/C-11/C-12, $H_3$-23 to C-14/C-15/C-16, $H_3$-24 to C-18/C-19/C-25, as well as from $H_3$-25 to C-18/C-19/C-24. Collectively, these correlations allowed the construction of the planar structure of **17**, revealing a previously unreported sesterterpene featuring an 5/11 fused bicyclic skeleton. The $\Delta^{2,3}$, $\Delta^{6,7}$ and $\Delta^{15,16}$ double bonds were assigned as *E* configuration based on the NOESY correlations of $H_3$-20/H-1a, $H_3$-21/H-5a, and $H_3$-23/H-17. The key NOESY correlations of H-10/H-12b, H-12b/H-14, and $H_3$-22/H12a/$H_3$-21 revealed that H-10 and H-14 were on the same side whereas $H_3$-22 was located on the other side, further establishing the relative configuration of **17** as 10*S**,11*R**,14*S** (Figure S14 and Table S15). Finally, TD-DFT calculations of the ECD spectra enabled the determination of the absolute configuration of compound **17** as 10*S*,11*R*,14*S* (Figures S28 and S29), which was designated clorospene.

The molecular formula of **18** was assigned as $C_{25}H_{40}$ based on its HR-EI-MS data (*m*/*z* 340. 3133 $[M]^+$; calcd for $[C_{25}H_{40}]^+$, 340.3125). Comparison of its 1D and 2D NMR data with those of ophiobora-2,6,15(23),18-tetraene,[18] indicated that the two compounds shared the same planar structure. In the NOESY spectrum, the observed correlations of $H_3$-22/H-14, $H_3$-22/H-1a, and H-1b/H-10, indicated that $H_3$-22 and H-14 were on the same side of the cyclopentane unit, while H-10 was on the other side, thus establishing the relative configuration of **18** as 10*S**,11*R**,14*R** (Figure S15 and Table S16). Finally, **18** was determined as 14-*epi*-ophiobora-2,6,15(23),18-tetraene by comparing the experimental and calculated ECD spectra (Figures S30 and S31), and it was designated fusspopene A.

The molecular formula of compound **20** was determined to be $C_{25}H_{40}$ based on its HR-EI-MS data, which showed a molecular ion peak at *m/z* 340.3129 $[M]^+$ (calcd for $[C_{25}H_{40}]^+$, 340.3125). Analysis of the $^1H$ and $^{13}C$ NMR spectra data revealed that compound **20** was similar to preasperterpenoid A, previously produced by AstC and PvPS.[19, 20] The principal structural difference between the two compounds was the position of the olefinic bond. In preasperterpenoid A, the double bond is located between C-2 and C-3, whereas in compound **20**, the double bond was found at positions C-1 and C-2, confirmed by the key HMBC correlation of $H_3$-20 to C-2/C-3/C-4, and $H_3$-22 to C-1/C-11/C-12/C-10. The relative configuration of **20** could be further assigned on the basis of its key NOESY correlations of H-3/H-5b, H-5a/$H_3$-21, H-9/$H_3$-21, H-9/H-14, H-9/$H_3$-22, H-12b/$H_3$-22, H-6/H-10, H-6/H-12a, H-12a/$H_3$-23, and H-18/$H_3$-23 (Figure S17 and Table S18). Subsequently, comparison of the experimental and calculated $^{13}C$ NMR chemical shifts further supported assignment of the relative configuration at C-3, establishing the relative configuration of compound **20** as 3*S**,6*S**,7*R**,9*R**,10*S**,11*S**,14*S**,15*S**,18*R**(Figures S54 and S55). Finally, the absolute configuration of **20** was assigned as 3*S*,6*S*,7*R*,9*R*,10*S*,11*S*,14*S*,15*S*,18*R* by comparing the experimental and calculated ECD spectra (Figures S34 and S35), and compound **20** was named fusspopene B.

Compound **21** exhibited a molecular ion peak at *m/z* 340.3128 $[M]^+$ (calcd for $[C_{25}H_{40}]^+$, 340.3125) in the HR-EI-MS spectrum, consistent with a molecular formula of $C_{25}H_{40}$ and six degrees of unsaturation. Analysis of the $^1H$, $^{13}C$ NMR, and HSQC spectra revealed 25 carbon signals, including six methyl groups, eight methylene groups, five methine carbons, two olefinic carbons, and four $sp^3$ quaternary carbons. The degree of unsaturation and NMR features suggested a pentacyclic framework. The $^1H$–$^1H$ COSY spectrum established four distinct spin systems: $H_3$-20−H-3−H-4 (**I**), H-5−H-6−H-8−H-9−H-10−H-14−H-18 (**II**), H-12−H-13 (**III**), and H-16−H-17 (**IV**). Key HMBC correlations of $H_3$-20 to C-3/C-4/C-7; $H_3$-21 to C-2/C-3/C-6/C-7; $H_3$-22 to C-1/C-11/C-12/C-14; $H_3$-23 to C-13/C-15/C-16/C-18; $H_3$-24 to C-17/C-18/C-19/C-25; $H_3$-25 to C-17/C-18/C-19/C-24, H-1 to C-7/C-10, and H-10 to C-2 further established the planar structure of **21**, which was similar to peniroquesines A−C[21] with a novel 5/6/5/6/5 fused pentacyclic skeleton. The NOESY correlations of H-3/H-10, H-10/H-18/$H_3$-23/$H_3$-25, H-14/$H_3$-22/$H_3$-24, $H_3$-21/$H_3$-22, H-6/$H_3$-22, and $H_3$-20/$H_3$-22 indicated that H-3, H-10, H-18, $H_3$-23, and $H_3$-25 were on the same side, whereas H-6, H-14, $H_3$-20, $H_3$-21, $H_3$-22, and $H_3$-24 were located on the opposite side (Figure S18 and Table S19), thus establishing the relative configuration of **21** as 3*R**,6*R**,7*R**,10*S**,11*S**,14*R**,15*S**,18*S**. The absolute configuration of **21** was established as 3*R*,6*R*,7*R*,10*S*,11*S*,14*R*,15*S*,18*S* by comparison of the experimental and calculated ECD spectra (Figures S36 and S37). Accordingly, compound **21** was named fusspopene C.

The molecular formula of compound **22** was determined as $C_{25}H_{40}$ from HR-EI-MS, which displayed a molecular ion peak at *m/z* 340.3129 $[M]^+$ (calcd for $[C_{25}H_{40}]^+$, 340.3125), corresponding to six degrees of unsaturation. The NMR spectroscopic data of compound **22** closely resembled those of compound **21**, suggesting that **22** is a structural analog of **21** that differs primarily in the position of the double bond. The HMBC correlations from $H_3$-21 to C-2/C-3/C-6/C-7 and from H-14 to C-2/C-10 established the presence of a $\Delta^{2,10}$ double bond in **22**, rather than the $\Delta^{1,2}$ double bond observed in **21**. In the NOESY spectrum, correlations of H-6/H-14, H-14/$H_3$-21/$H_3$-22/$H_3$-24, H-12a/$H_3$-22, H-3/H-12b, H-12b/$H_3$-23, H-18/$H_3$-23, and H-18/$H_3$-25 were observed (Figure S19 and Table S20), single-crystal X-ray diffraction analysis using Cu *Kα* radiation further established the relative configuration of **22** as 3*S**,6*R**,7*R**,11*R**,14*R**,15*S**,18*S** [Flack parameter = 0.1(4)] (Figure S58 and Table S27, CCDC 2504628). Finally, comparison of the experimental and calculated ECD spectra assigned the absolute configuration of **22** as 3*S*,6*R*,7*R*,11*R*,14*R*,15*S*,18*S* (Figures S38 and S39), and the compound was designated fusspopene D.

Compound **23** displayed a molecular ion peak at *m/z* 340.3131 $[M]^+$ (calcd for $[C_{25}H_{40}]^+$, 340.3125), indicating six degrees of unsaturation. Analysis of the $^1H$, $^{13}C$ NMR, and HSQC spectra revealed 25 carbon signals, comprising six methyl groups, eight methylene groups, five methine groups, two olefinic carbons, and four $sp^3$ quaternary carbons, suggesting that **23** possesses a pentacyclic structure. The $^1H$–$^1H$ COSY spectrum of **23** revealed four contiguous spin systems H-4−H-5−H-6−$H_3$-21 (**I**), H-1−H-2−H-10−[H-9−H-8]−[H-14−H-18] (**II**), H-12−H-13 (**III**), and H-16−H-17 (**IV**) respectively. Detailed analysis of the $^{13}C$ NMR data led to the existence of two identical rings (A and B) with compounds **22** and **23**, which was further confirmed by the spin systems **III** and **IV** and the HMBC correlations of $H_3$-22 to C-1/C-11/C-12/C-14, $H_3$-23 to C-13/C-15/C-16/C-18, $H_3$-24 to C-17/C-18/C-19/C-25, and $H_3$-25 to C-17/C-18/C-19/C-24. The spin systems **II** indicated the presence of a cyclopentane ring (ring C) and further confirmed the fusion of rings B and C through the C-11 and C-14 bond. The HMBC correlations from H-2 to C-1/C-7/C-8/C-9, and from H-8a to C-7/C-9 indicated that **23** possesses a cyclopentene ring (ring D). In addition, spin system **I**, together with the HMBC correlations from $H_3$-20 to C-

3/C-4/C-7, $H_3$-21 to C-5/C-6/C-7, H-2 to C-6, and H-8a to C-6 indicated the presence of a cyclopentane ring (ring E), which is spiro-fused with ring D by sharing the C-7 bond. Accordingly, the planar structure of compound **23** was defined as a novel 5/6/5/5/5 pentacyclic sesterterpene. The relative configuration of **23** could be further assigned on basis of its key NOESY correlations of H-8/$H_3$-21, H-8b//$H_3$-22, H-14/$H_3$-22, H-14/$H_3$-24, H-2/H-6, H-6/H-10 and H-10/H-18/$H_3$-23/$H_3$-25 indicated that H-8, H-14, $H_3$-21, $H_3$-22, and $H_3$-24 were on the same side whereas H-2, H-6, H-10, $H_3$-23, and $H_3$-25 were located on the opposite. Moreover, NOESY correlations of H-6/H-2/H-10 established both the C-6 and C-7 relative stereochemistry as well as the position of the double bond. Thus, the relative configurations of all chiral carbons in the ring systems were determined to be 2*R**,6*R**,7*R**,10*R**,11*R**,14*R**,15*S**,18*S** (Figure S20 and Table S21). Finally, comparison of experimental and calculated ECD spectra (Figures S40 and S41) confirmed the absolute configuration of **23** as 2*R*,6*R*,7*R*,10*R*,11*R*,14*R*,15*S*,18*S*, and the compound was designated fusspopene E.

The molecular formula of **24** was determined to be $C_{25}H_{40}$ based on its HR-EI-MS data, which showed a molecular ion peak at *m/z* 340.3132 $[M]^+$ (calcd for $[C_{25}H_{40}]^+$, 340.3125). Analysis of the $^1$H, $^{13}$C NMR, and HSQC spectra of compound **24** revealed the presence of 25 carbon signals, including five methyl groups, nine methylene groups, four methine groups, four olefinic carbons, and three $sp^3$ quaternary carbons. Considering that compound **24** exhibits six degrees of unsaturation but contains only two double bonds, it is inferred to possess a tetracyclic structure. Analysis of the COSY spectrum of **24** discovered the proton connectivity for four structure fragments: H-4−H-5−H-6 (**I**), H-1−H-2−H-10−[H-9−H-8]−[H-14−H-18] (**II**), H-12−H-13 (**III**), and H-16−H-17 (**IV**). Analysis of the $^{13}$C NMR data led to the existence of three identical rings (A−C) with **23,** which was further confirmed by the spin systems **II**, **III** and **IV** and the HMBC correlations of $H_3$-22 to C-1/C-11/C-12/C-14; $H_3$-23 to C-13/C-15/C-16/C-18; $H_3$-24 to C-17/C-18/C-19/C-25; $H_3$-25 to C-17/C-18/C-19/C-24; and H-12 to C-11/C-13/C-14/C-15. Fragments **I** and **II** could be connected by the clear HMBC correlations from $H_2$-20 to C-2 /C-4 and $H_3$-21 to C-6/C-7/C-8. Thus, the planar structure of **24** could be fully constructed and was shown to possess a unique 5/6/5/9 tetracyclic pentacyclic ring system. The $\Delta^{6,7}$ was determined to be *E* configuration based on the NOESY correlations between H-5a/$H_3$-21. The relative configuration of **24** was further determined based on its key NOESY correlations of H-2/H-18, H-10/H-18, H-10/$H_3$-23, $H_3$-23/$H_3$-25, H-14/$H_3$-22, and H-14/$H_3$-24 (Figure S21 and Table S22). By comparing experimental and calculated ECD spectra (Figures S42 and S43), the absolute configurations of 2*R*,10*R*,11*R*,14*R*,15*S*,18*S*-**24** were determined and named fusspopene F.

Compound **25**, which the molecular formula was determined as $C_{25}H_{40}$ from HR-EI-MS, displayed a molecular ion peak at *m/z* 340.3131 $[M]^+$ (calcd for $[C_{25}H_{40}]^+$, 340.3125), corresponding to six degrees of unsaturation. Analysis of the $^1$H, $^{13}$C, and HSQC spectra revealed 25 carbon signals, including six methyl groups, nine methylene groups, four methine groups, two olefinic carbons, and four $sp^3$ quaternary carbons, indicating a pentacyclic structure. Analysis of the $^1$H–$^1$H COSY correlation revealed the presence of four independent spin systems: H-4−H-5 (**I**), $H_3$-21−H-7−H-8−H-9−H-10−H-14 (**II**), H-12−H-13 (**III**), and H-16−H-17 (**IV**). Detailed analysis of the $^{13}$C NMR data led to the existence of two identical rings (A and B) with compounds **22−24**, which was further confirmed by the spin systems **III**, **IV**, and the HMBC correlations of $H_3$-23 to C-13/C-15/C-16/C-18, $H_3$-24 to C-17/C-18/C-19/C-25, and $H_3$-25 to C-17/C-18/C-19/C-24. Spin system **II** and the HMBC correlations of H-1 to C-2/C-3/C-6/C-11/C-12/C-14 indicated the presence of a cyclohexane ring (ring C). The spin system **II** combined with the HMBC correlations of $H_3$-21 to C-6/C-7/C-8; and H-10/C-8 indicated that **25** possesses a cyclopentene ring (ring D). The remaining one ring (cyclopentene, ring E) was deduced from spin system **I** and the HMBC correlations of $H_3$-21 to C-6/C-7, H-7 to C-5/C-6, and $H_3$-20 to C-2/C-3/C-4. Accordingly, the planar structure of compound **25** was defined as a unique 5/6/6/5/5 pentacyclic sesterterpene. The relative configurations of **25** was determined by the key NOESY correlation of H-14/ $H_3$-21, H-14/ $H_3$-22, H-14/ $H_3$-24, H-5a/H-10, H-10/H-18, H-18/$H_3$-23, and H-18/$H_3$-25 (Figure S22 and Table S23). The relative configuration of **25** was determined as 6*S**,7*S**,10*S**,11*R**,14*R**,15*S**,18*S** by single-crystal X-ray diffraction analysis using Cu *Kα* radiation [Flack parameter of -0.1(4)] (Figure S58 and Table S27, CCDC 2504621), and the absolute configuration of **25** was established as 6*S*,7*S*,10*S*,11*R*,14*R*,15*S*,18*S* by comparing of the experimental and calculated ECD spectrum (Figures S44 and S45). Compound **25** was a previously unreported sesterterpene and was named fusspopene G.

The HR-EI-MS analysis of compound **26** exhibited a molecular ion peak at *m/z* 340.3132 $[M]^+$ (calcd for $[C_{25}H_{40}]^+$, 340.3125), consistent with six degrees of unsaturation. The NMR spectroscopic data of **26** closely resembled those of compound **23**, suggesting that **26** is a structural analogue of **23**. The major difference was the position of the olefinic bond. Key HMBC correlations from H-3 to C-2, H-6 to C-2, H-8 to C-2/C-10, H-9 to C-2, and H-14 to C-2/C-10 established the presence of a $\Delta^{2,10}$ double bond in **26**, instead of the $\Delta^{3,4}$ double bond observed in **23**. The NOESY correlations of H-8a/$H_3$-20/$H_3$-22, H-8b/$H_3$-21, H-14/$H_3$-21, H-14/$H_3$-22, H-14/$H_3$-24, $H_3$-20/$H_3$-22, H-18/$H_3$-23, and H-18/$H_3$-25 allowed assignment of the relative stereochemistry (Figure S23 and Table S24). Finally, comparison of the experimental and calculated ECD spectra (Figures S46 and S47) established the absolute configuration of **26** as 3*S*,6*R*,7*R*,11*R*,14*R*,15*S*,18*S*, and the compound was designated artconpene A.

The molecular formula of **27** was inferred to be $C_{25}H_{40}$ based on HR-EI-MS data, which exhibited a molecular ion peak at *m/z* 340.3132 $[M]^+$ (calcd for $[C_{25}H_{40}]^+$, 340.3125), indicating six degrees of unsaturation. Analysis of the $^1H$, $^{13}C$ NMR, and HSQC spectra of compound **27** revealed 25 carbon signals, including six methyl groups, eight methylene groups, five methine groups, two olefinic carbons, and four $sp^3$ quaternary carbons, suggesting a pentacyclic structure. Analysis of the COSY spectrum of **27** revealed the proton connectivity for three structure fragments: $H_3$-20−H-4−H-5 (**I**), H-8−H-9−H-10−[H-2]−H-14−H-18 (**II**), and H-16−H-17 (**III**). Detailed analysis of the NMR spectroscopic data of compound **27** indicated that it possesses the same A–D tetracyclic core as compound **25**. This assignment was supported by the observed spin systems **II−III** together with the HMBC correlations from $H_3$-21 to C-2/C-6/C-7/C-8, $H_3$-22 to C-1/C-11/C-12/C-14, $H_3$-23 to C-13/C-15/C-16/C-18, $H_3$-24 to C-17/C-18/C-19/C-25, $H_3$-25 to C-17/C-18/C-19/C-24, H-18 to C-11/C-14/C-17/C-19, H-13 to C-12/C-15, H-1 to C-2/C-14, H-2 to C-3, and H-10 to C-7, thereby establishing the conserved A–D ring framework. In addition, spin system **I**, together with the HMBC correlations from $H_3$-20 to C-3/C-4/C-5, $H_3$-21 to C-2/C-6/C-7/C-8, and H-5 to C-6 indicated the presence of ring E. Accordingly, the planar structure of compound **27** was defined as a novel 5/6/6/5/6 pentacyclic sesterterpene. The NOESY correlations of H-2/H-10, H-10/H-18, H-18/$H_3$-23, $H_3$-23/$H_3$-25, H-14/$H_3$-21, H-14/$H_3$-22, H-14/$H_3$-24, and $H_3$-20/$H_3$-22 established the relative configuration of **27** as 2*R**,4*S**,7*S**,10*S**,11*S**,14*R**,15*S**,18*S** (Figure S24 and Table S25), which was further supported by single-crystal X-ray diffraction analysis using Cu *Kα* radiation with a Flack parameter of 0.1(5) (Figure S58 and Table S27, CCDC 2504622). Finally, comparison of the experimental and calculated ECD spectra established the absolute configuration of **27** as 2*R*,4*S*,7*S*,10*S*,11*S*,14*R*,15*S*,18*S* (Figures S48 and S49), and the compound was designated fusspopene H.

The molecular formula of **28** was determined as $C_{25}H_{40}$ based on HR-EI-MS, which displayed a molecular ion peak at *m/z* 340.3130 $[M]^+$ (calcd for $[C_{25}H_{40}]^+$, 340.3125), corresponding to six degrees of unsaturation. The GC-MS and NMR spectroscopic data of **28** closely resembled those of compound **27**, suggesting that the two compounds share a closely related skeleton. The major structural difference between **28** and **27** was the position of the olefinic bond, which was established by the HMBC correlations from H-1 to C-2/C-3, H-4 to C-3, H-6 to C-2, $H_3$-20 to C-3, and $H_3$-21 to C-2, confirming the presence of a $\Delta^{2,3}$ double bond in **28**. The NOESY correlations of H-4/H-10, H-10/$H_3$-23, H-18/$H_3$-23, H-18/$H_3$-25, H-1b/$H_3$-20, H-1b/$H_3$-21, H-1b/$H_3$-22, H-14/$H_3$-24, $H_3$-22/$H_3$-24 provided information on the relative configuration of stereocenters (Figure S25 and Table S26). Finally, comparison with calculated ECD spectra established the absolute configuration of **28** as 4*S*,7*R*,10*S*,11*R*,14*R*,15*S*,18*S* (Figures S50 and S51), and the compound was designated artconpene B.

The molecular formula of **29** was established as $C_{25}H_{40}O$ based on HR-EI-MS, which showed a molecular ion peak at *m/z* 358.3239 $[M]^+$ (calcd for $[C_{25}H_{40}O]^+$, 358.3236), corresponding to six degrees of unsaturation. Detailed analysis of the NMR spectroscopic data revealed that compound **29** is closely related to compound **18**. The major differences were the introduction of a hydroxyl group at C-15 and the absence of one double bond in **29**. Further evidence was obtained from the HMBC correlations of $H_3$-23 to C-14/C-15/C-16, and H-10 to C-15, which established **29** as a hydroxylated 5/11 bicyclic sesterterpene.[22] In addition, NOESY correlations of H-1a/$H_3$-20 and H-5/$H_3$-21 further supported the $\Delta^{2,3}$, $\Delta^{6,7}$ double bonds were assigned as *E* configuration. The relative configuration of C-10, C-11, and C-14 were assigned as 10*S**,11*R**, and 14*R** according to the key NOESY correlations of H-1a/H-10, H-1b/$H_3$-22, and H-14/$H_3$-22 (Figure S26 and Table S29). Because $H_3$-23 is positioned on a flexible side chain, the NOE correlation involving $H_3$-23 were insufficient to unambiguously establish the relative configuration at this position. Therefore, comparison of the experimental and calculated $^{13}C$ NMR chemical shifts was performed (Figures S56 and S57), which allowed assignment of the configuration at C-15, thereby establishing the relative configuration of compound **29** as 10*S**,11*R**,14*R**,15*R**. Finally, comparison with calculated ECD spectra established the absolute configuration of **29** as 10*S*,11*R*,14*R*,15*R* (Figures S52 and S53), and this compound was accordingly named fusspopene I.

fusicoccadiene (**1**): colorless oil; $[\alpha]_D^{20}$ = +18.0 (*c* 0.1, $CHCl_3$); $^1H$ (pyridine-$d_5$, 600 MHz) and $^{13}C$ (pyridine-$d_5$, 150 MHz) NMR spectra, see Figure S9 and Table S10; HR-EI-MS: $[M]^+$ *m/z* 272.2499 (calcd for $[C_{20}H_{32}]^+$, 272.2499)

ophiobolin F (**2**): colorless needles; $[\alpha]_D^{30}$ = +41.0 (*c* 0.1, $CHCl_3$); $^1H$ (benzene-$d_6$, 600 MHz) and $^{13}C$ (benzene-$d_6$, 150 MHz) NMR spectra, see Figure S10 and Table S11; HR-EI-MS: $[M]^+$ *m/z* 358.3239 (calcd for $[C_{25}H_{40}O]^+$, 358.3230)

mangicdiene (**8**): colorless oil; $^1H$ (benzene-$d_6$, 600 MHz) and $^{13}C$ (benzene-$d_6$, 150 MHz) NMR spectra, see Figure S11 and Table S12; HR-EI-MS: $[M]^+$ *m/z* 340.3136 (calcd for $[C_{25}H_{40}]^+$, 340.3125)

spiroviolene (**15**): colorless oil; $[\alpha]_D^{30}$ = -4.3 (*c* 0.1, $CHCl_3$); $^1H$ (benzene-$d_6$, 600 MHz) and $^{13}C$ (benzene-$d_6$, 150

MHz) NMR spectra, see Figure S12 and Table S13; HR-EI-MS: $[M]^+$ *m/z* 272.2502 (calcd for $[C_{20}H_{32}]^+$, 272.2499)

variediene (**16**): colorless oil; $[\alpha]_D^{30}$ = -127.1 (*c* 0.17, $CHCl_3$); $^1H$ (benzene-$d_6$, 600 MHz) and $^{13}C$ (benzene-$d_6$, 150 MHz) NMR spectra, see Figure S13 and Table S14; HR-EI-MS: $[M]^+$ *m/z* 272.2506 (calcd for $[C_{20}H_{32}]^+$, 272.2499)

clorospene (**17**): colorless oil; $[\alpha]_D^{30}$ = +75.4 (*c* 0.26, $CHCl_3$); $^1H$ (benzene-$d_6$, 600 MHz) and $^{13}C$ (benzene-$d_6$, 150 MHz) NMR data, see Figure S14 and Table S15; HR-EI-MS: $[M]^+$ *m/z* 340.3131 (calcd for $[C_{25}H_{40}]^+$, 340.3125)

fusspopene A (**18**): colorless oil; $[\alpha]_D^{30}$ = -68.7 (*c* 0.16, $CHCl_3$); $^1H$ (benzene-$d_6$, 600 MHz) and $^{13}C$ (benzene-$d_6$, 150 MHz) NMR data, see Figure S15 and Table S16; HR-EI-MS: $[M]^+$ *m/z* 340.3133 (calcd for $[C_{25}H_{40}]^+$, 340.3125)

astelifatriene (**19**): colorless oil; $^1H$ (benzene-$d_6$, 600 MHz) and $^{13}C$ (benzene-$d_6$, 150 MHz) NMR spectra, see Figure S16 and Table S17; HR-EI-MS: $[M]^+$ *m/z* 340.3133 (calcd for $[C_{25}H_{40}]^+$, 340.3125)

fusspopene B (**20**): colorless oil; $[\alpha]_D^{30}$ = -224.0 (*c* 0.05, $CHCl_3$); $^1H$ (benzene-$d_6$, 600 MHz) and $^{13}C$ (benzene-$d_6$, 150 MHz) NMR data, see Figure S17 and Table S18; HR-EI-MS: $[M]^+$ *m/z* 340.3129 (calcd for $[C_{25}H_{40}]^+$, 340.3125)

fusspopene C (**21**): colorless oil; $[\alpha]_D^{30}$ = -216.0 (*c* 0.05, $CHCl_3$); $^1H$ (benzene-$d_6$, 600 MHz) and $^{13}C$ (benzene-$d_6$, 150 MHz) NMR data, see Figure S18 and Table S19; HR-EI-MS: $[M]^+$ *m/z* 340.3128 (calcd for $[C_{25}H_{40}]^+$, 340.3125)

fusspopene D (**22**): colorless needles; $[\alpha]_D^{30}$ = +46 (*c* 0.1, $CHCl_3$); $^1H$ (benzene-$d_6$, 600 MHz) and $^{13}C$ (benzene-$d_6$, 150 MHz) NMR data, see Figure S19 and Table S20; HR-EI-MS: $[M]^+$ *m/z* 340.3129 (calcd for $[C_{25}H_{40}]^+$, 340.3125)

fusspopene E (**23**): colorless oil; $[\alpha]_D^{30}$ = -186.0 (*c* 0.06, $CHCl_3$); $^1H$ (benzene-$d_6$, 600 MHz) and $^{13}C$ (benzene-$d_6$, 150 MHz) NMR data, see Figure S20 and Table S21; HR-EI-MS: $[M]^+$ *m/z* 340.3131 (calcd for $[C_{25}H_{40}]^+$, 340.3125)

fusspopene F (**24**): colorless oil; $[\alpha]_D^{30}$ = +97.9 (*c* 0.23, $CHCl_3$); $^1H$ (benzene-$d_6$, 600 MHz) and $^{13}C$ (benzene-$d_6$, 150 MHz) NMR data, see Figure S21 and Table S22; HR-EI-MS: $[M]^+$ *m/z* 340.3132 (calcd for $[C_{25}H_{40}]^+$, 340.3125)

fusspopene G (**25**): colorless needles; $[\alpha]_D^{30}$ = -365.0 (*c* 0.04, $CHCl_3$); $^1H$ (benzene-$d_6$, 600 MHz) and $^{13}C$ (benzene-$d_6$, 150 MHz) NMR data, see Figure S22 and Table S23; HR-EI-MS: $[M]^+$ *m/z* 340.3131 (calcd for $[C_{25}H_{40}]^+$, 340.3125)

artconpene A (**26**): colorless oil; $[\alpha]_D^{30}$ = +52.0 (*c* 0.1, $CHCl_3$); $^1H$ (benzene-$d_6$, 600 MHz) and $^{13}C$ (benzene-$d_6$, 150 MHz) NMR data, see Figure S23 and Table S24; HR-EI-MS: $[M]^+$ *m/z* 340.3131 (calcd for $[C_{25}H_{40}]^+$, 340.3125)

fusspopene H (**27**): colorless needles; $[\alpha]_D^{30}$ = +61.8 (*c* 0.11, $CHCl_3$); $^1H$ (benzene-$d_6$, 600 MHz) and $^{13}C$ (benzene-$d_6$, 150 MHz) NMR data, see Figure S24 and Table S25; HR-EI-MS: $[M]^+$ *m/z* 340.3132 (calcd for $[C_{25}H_{40}]^+$, 340.3125)

artconpene B (**28**): colorless needles; $[\alpha]_D^{30}$ = +75.0 (*c* 0.08, $CHCl_3$); $^1H$ (benzene-$d_6$, 600 MHz) and $^{13}C$ (benzene-$d_6$, 150 MHz) NMR data, see Figure S25 and Table S26; HR-EI-MS: $[M]^+$ *m/z* 340.3130 (calcd for $[C_{25}H_{40}]^+$, 340.3125)

fusspopene I (**29**): colorless oil; $[\alpha]_D^{30}$ = +349.3 (*c* 0.15, $CHCl_3$); $^1H$ (benzene-$d_6$, 600 MHz) and $^{13}C$ (benzene-$d_6$, 150 MHz) NMR data, see Figure S26 and Table S29; HR-EI-MS: $[M]^+$ *m/z* 358.3239 (calcd for $[C_{25}H_{40}O]^+$,

358.3230)

### 5 Proposed cyclization pathways for FsPS- and AcPS-derived compounds 18–28

Both FsPS and AcPS catalyze the cyclization of geranylfarnesyl pyrophosphate (GFPP) to yield the sesterterpenes **21** and **22**, which share the same carbon skeleton as peniroquesines A–C, previously reported sesterterpenes identified from *Penicillium roqueforti.*[21] Detailed structural analysis further revealed that the A and B rings of the newly identified compounds possess identical connectivity and absolute configurations as those of compounds **21** and **22** (Figure 4A), indicating a conserved early-stage cyclization sequence. Based on this structural similarity, we hypothesized that the novel skeletons arise from divergent downstream cyclization pathways that branch from the common core intermediates leading to **21** and **22**. Building on this shared structural foundation, we reconstructed a comprehensive cyclization network and investigated how FsPS and AcPS generates multiple structurally complex and highly congested sesterterpenes (**18**–**28**) through retrosynthetic analysis and DFT calculations.

The cyclization mechanism of peniroquesine has been investigated through isotopic labeling and DFT calculations.[21, 23] We therefore proposed the cyclization pathway of the FsPS-derived products based on the established peniroquesine pathway. The dissociation of pyrophosphate from GFPP to generate an allylic carbocation (**IM1**) initiates a multistep carbocation cascade. A through-space cation–*π* interaction between the allylic cation in **IM1** and the *π*-electrons of the C10–C11 double bond promotes smooth annulation to give **IM2**, a tertiary (3°) carbocation stabilized by hyperconjugation with the C1–C10 *σ*-bond. This intermediate subsequently undergoes a reverse (3° → secondary (2°)) Wagner–Meerwein rearrangement to afford a 2° carbocation, which is immediately trapped by the *π*-electrons of the nearby C14–C15 double bond to generate a 5/11 bicyclic 3° carbocation (**IM3**) (Figure 4B). After a conformational change from **IM3** to **IM3a**, C–C σ-bond formation proceeds in a manner analogous to the **IM2** → **IM3** step, yielding a 5/6/11 tricyclic 3° carbocation (**IM4**), which follows the same biosynthetic pathway as peniroquesines described by Taro et al. The deprotonation ($-H^+$) at C23 of **IM3** and C24 of **IM4** yields compounds **18** and **19**, respectively (Figure S61).

**IM4** serves as the branch point and leading to the formation **20**, which shares the same carbon skeleton as the known sesterterpene preasperterpenoid A.[19, 24] **IM4** undergoes a 1,5-hydrogen shift to generate a *μ*-hydrogen–bridged cycloalkonium intermediate (**IM4-1**), which rapidly evolves into the relatively stable 2° carbocation **IM4-2**. Subsequently, electron transfer from $\Delta^{6,7}$ to C9, followed by transfer from $\Delta^{2,3}$ to C6, ultimately forming **IM4-3,** a 5/6/7/3/5 pentacyclic 3° carbocation intermediate. In the biosynthesis of preasperterpenoid A, the carbocation intermediate **IM4-3** undergoes direct deprotonation to yield the final product. In contrast, in FsPS, **IM4-3** undergoes a 1,2-hydride shift that transfers the positive charge from C3 to C2, generating **IM4-4** with a calculated energy barrier of 5.0 kcal/mol. Subsequent deprotonation at C1 then forms compound **20** (Figure S61). This mechanistic divergence explains the structural difference between preasperterpenoid A and compound **20**, and further illustrates how subtle modulation of carbocation migration can direct terpene scaffold diversification.

In another pathway, a third reverse Wagner–Meerwein rearrangement of **IM4**, involving cleavage of the C17–C18 bond and formation of a new C17–C19 bond accompanied by ring expansion, yields **IM5**. **IM5** could first undergoes a rearrangement reaction involves two consecutive 1,2-alkyl shifts, with cleavage of the C14–C15 and C10–C11 bonds and formation of new C15–C18 and C11–C14 bonds, followed by cyclization through C2–C10 and C3–C6 bond formation to generate **IM5-1**, a 5/6/5/7/4 pentacyclic carbocation intermediate. **IM5-1** then undergoes a conformational adjustment in which the C6–C7 double bond geometry changes from *trans* to *cis*, affording **IM5-1a**. Further conformational reorganization converts the B ring from a boat to a chair conformation, generating **IM5-1b**. **IM5-1b** subsequently undergoes annulation accompanied by a Wagner–Meerwein methyl shift, followed by a reverse Wagner–Meerwein rearrangement and additional skeletal reorganization to give **IM5-2**, featuring a *cis*-fused bicyclo [4.2.1]nonane framework. From **IM5-2**, a facile ring-contraction occurs, likely driven by the strained bicyclic structure and favorable orbital overlap between the carbocation p-orbital and the C2–C3 σ-bond, affording the more stable tertiary carbocation **IM5-3**. **IM5-3** then undergoes a ring-spanning 1,3-hydride shift to generate **IM5-4**. Finally, **IM5-4** undergoes deprotonation at C1 and C10 to produce compounds **21** and **22**, respectively (Figure S61).

Alternatively, **IM5** undergoes a conformational change to afford **IM5a**, in which the *Si* face of the C6=C7 double bond is oriented toward the *Re* face of the C2=C3 double bond. **IM5a** then proceed through two successive transformations analogous to 1,2-alkyl shifts, involving cleavage of the C14–C15 and C10–C11 bonds and formation of new C15–C18 and C11–C14 bonds, to afford intermediates **IM6** and **IM7**. At this stage, **IM7** establishes the common A/B-ring framework shared by all downstream products, whereas the structural divergence among the subsequent compounds arises from different cyclization modes involved in the construction of the C/D/E rings. **IM7** serves as a second key branching intermediate derived two distinct pathways (***Path I*** and ***Path II***), affording compounds **23**–**26** and **27**–**28**, respectively.

In ***Path I***, **IM7** undergoes a smooth asynchronous concerted annulation via C2–C10 and C3–C6 bond formation, yielding **IM8**, a 3°carbocation intermediate with substantial stabilization energy (−25.7 kcal/mol). **IM8** subsequently undergoes a Wagner− Meerwein methyl shift, involved in the cleavage of the C3–C6 bond and formation of a new C3–C7 bond to afford **IM8-1**, a 5/6/5/6/5 polycyclic intermediate, with an energy barrier of 9.4 kcal/mol. This is followed by cleavage of the C2–C3 bond and formation of a new C2–C7 bond, generating **IM8-2**, a 5/6/5/5/5 spirocyclic framework. This ring-contraction process occurs with a very low activation energy of 3.4 kcal/mol, and deprotonation at C4 of **IM8-2** directly yields **23** (Figures 4C, S62 and S63)**.** Although the carbon skeleton of compound **23** is reported here for the first time, this structural scaffold appears in the cyclization pathway proposed for roquefornine by Wang and co-workers.[25] Alternatively, Cleavage of the C3–C6 *σ* bond within **IM8** generates a 3° tetracyclic carbocation (**IM9**) possessing a 5/6/5/9 fused framework, which is further stabilized by a cation–*π* interaction with the newly formed C6–C7 double bond. **IM9** can undergo direct deprotonation at C20 to yield **24**. Alternatively, **IM9** subsequently undergoes a 1,2-hydride shift that transfers the positive charge from C2 to C3 to afford **IM10**, with an activation energy of 6.6 kcal/mol. **IM10** then experiences further conformational reorganization to generate **IM10a**, followed by an electron transfer from $\Delta^{6,7}$ to C2 that smoothly constructs a 5/6/5/6/5 pentacyclic skeleton (**IM11**). This intermediate undergoes a 1,2-hydride shift to generate **IM12** with an exceptionally low energy barrier of 1.2 kcal/mol. Subsequently, a C–C bond rearrangement takes place after overcoming an 11.4 kcal/mol barrier, leading to a 5/6/6/5/5 pentacyclic skeleton intermediate **IM13**, which upon final deprotonation at C3 generates compound **25**. DFT calculations confirm that this pathway is energetically feasible, with the highest barrier (11.4 kcal/mol) arising during the ring-contraction step from **IM12** to **IM13** (Figures 4C, S64 and S65).

In ***Path II***, **IM7** undergoes conformational changes and then experiences a smooth asynchronous concerted annulation via C7–C10 and C2–C6 bond formation and yielding **IM7-1**, which features a 5/6/4/7/5 pentacyclic scaffold. This step represents the highest activation barrier along this pathway (25.3 kcal/mol). A subsequent 1,2-hydride shift to generate **IM7-2**, followed by a 1,2-alkyl migration gets carbocation **IM7-3** with 5/6/6/5/5 pentacyclic scaffold, both steps proceed with comparatively low activation barriers of 5.4 and 0.7 kcal/mol, respectively. Further skeletal reorganization via a 1,2-alkyl shift cleaves the C5–C6 bond and forms a new C5–C7 bond, affording **IM7-4** with the requisite 5/6/6/5/6 framework, crossing an exceptionally low energy barrier of 2.1 kcal/mol. Finally, a 1,2-hydride shift from C2 to C6 (3.6 kcal/mol) of **IM7-5**, followed by deprotonation at C1, delivers compound **27** (Figures 4C, S66 and S67). Given that C6–C10 cyclization has been proposed in related terpene cyclization pathways such as phomopsene,[26] two additional alternative C6–C10 cyclization pathways (***Path II'*** and ***Path II''***) from **IM7** were also evaluated.

In ***Path II'***, after two consecutive conformational adjustments, **IM7b** undergoes electron transfer from $\Delta^{6,7}$ to C10 to generate **IM7i-1**, a 5/6/5/9 tetracyclic intermediate, with a relatively high energy barrier of 19.8 kcal/mol. A subsequent conformational shift, mediated by rotation about the C8–C9–C10–C14 dihedral angle, enables a concerted 1,2- alkyl migration and electron transfer from $\Delta^{6,7}$ to C10, involing cleavage of C5-C6 bond and formation of new C5-C7 and C2-C6 bond, to afford **IM7i-3** with the characteristic 5/6/6/5/6 ring architecture. This concerted asynchronous process avoids the 2° cation (**TS_ IM7i-2–IM7i-3**) as intermediate. A subsequent 1,2-hydride shift of **IM7i-3** transfers the positive charge from C2 to C3 and forms the 3° carbocation **IM7i-4**, whose deprotonation at C1 would yield compounds **27** (Figure S66, ***Path II'***). However, DFT calculations revealed that the **IM7i-2** to **IM7i-3** transformation requires an insurmountable activation barrier of 91.1 kcal/mol, which suggests that the carbocation cascade is unlikely to proceed spontaneously at ambient temperature (Figure S68). ***Path II''*** was inspired by a previously proposed cyclization pathway of mangicdiene.[8, 26] Starting from **IM7i-2**, a smooth annulation involving a 1,2-hydride shift to form a bridgehead 3° carbocation, a conformational rearrangement of the nine-membered ring, and a subsequent 1,2-hydride shift, thereby constructing the 5/6/6/5/5 pentacyclic skeleton **IM13** in a single step. Two sequential 1,3-hydride shifts then migrate the cation first to C10 and subsequently to C7, generating **IM13-1** and **IM13-2**. A following 1,2-alkyl shift cleaves the C5–C6 bond and forms a new C5–C7 bond, yielding **IM13-3** with the requisite 5/6/6/5/6 skeleton (Figure S66, ***Path II''***). Nevertheless, DFT results showed that the **IM13** to **IM13-1** 1,3-hydride shift, has a prohibitively high activation barrier of 56.4 kcal/mol, again rendering this pathway energetically implausible (Figure S69).

AcPS additionally produce compounds **26** and **28** as pathway-specific products. Compound **26** possesses the same carbon skeleton as compound **23**. We initially hypothesized that **IM8-2** could further undergo a 1,3-hydride shift , followed by deprotonation to produce **26**, which was proposed by Wang et al. to explain the cyclization of roquefornine.[25] However, DFT calculations revealed that this transformation requires surmounting a prohibitively high energy barrier of 54.7 kcal/mol, indicating that this pathway is not viable (Figures S62 and S63). These results prompted us to reconsider the cyclization pathway of **26** from a scaffold-oriented perspective. Structurally, **26** features a C/D/E tricyclic 5/5/5 spirocyclic framework that closely resembles that of the diterpene spiroviolene.[12, 13, 27] Guided by the established spiroviolene cyclization mechanism, we proposed a thermodynamically feasible pathway to **26**, in which the intermediate **IM10** acts as a branching point and undergoes a spiroviolene-like carbocation cascade. A distinctive methyl shift, assisted by the C2 carbocation, then occurs to form a nonclassical transition state (**TS_10–10-1**) with an activation barrier of 5.5 kcal/mol, affording the 5/6/5/5/6 pentacyclic intermediate **IM10-1**. Subsequent migration of the C2–C3 σ bond toward the C7 carbocation (ring contraction) gives **IM10-2**, featuring a 5/6/5/5/5 spirocyclic framework. Finally, deprotonation at C10 affords final product compound **26** (Figures 4D, S70 and S71). Moreover, deprotonation at C2 of **IM7-4** would give **28**, generated by AcPS and sharing the same skeleton as **27**. (Figures 4D and S66).

### Tables S1 – S32

**Table S1.** Curated list of previously characterized fungal bifunctional terpene synthases producing cyclized terpene products. BFTPSs yielding non-cyclized linear products were excluded. The retained enzymes and their corresponding products were classified into Type A, Type B, and Others according to their catalytic outcomes.

| No. | Year | Name | Major product | Type | Classification | Ref. |
|---|---|---|---|---|---|---|
| 1 | 2007 | PaFS | Fusicoccadiene | Type B | NSR | [6] |
| 2 | 2008 | AbFS | Fusicoccadiene | Type B | NSR | [28] |
| 3 | 2008 | PaPS | Phomopsene | Type B | SR | [29] |
| 4 | 2013 | AcOS | Ophiobolin F | Type B | NSR | [7] |
| 5 | 2015 | NfSS | Sesterfisherol | Type A | NSR | [30] |
| 6 | 2015 | EvSS | Stellata-2,6,19-triene | Type B | SR | [31] |
| 7 | 2016 | EvQS | Quiannulatene | Type A | SR | [32] |
| 8 | 2016 | Au8003 | Ophiobolin F | Type B | NSR | [33] |
| 9 | 2016 | EvAS | Astellifadiene | Type B | SR | [34] |
| 10 | 2016 | EvVS | Variediene | Type B | SR | [15] |
| 11 | 2016 | FgMS | Mangicdiene | Type B | SR | [8] |
| 12 | 2017 | BmTS1 | Bm1 | Type A | NSR | [35] |
| 13 | 2017 | BmTS2 | Bm2 | Type A | NSR | [35] |
| 14 | 2017 | BmTS3 | Bm3 | Type A | NSR | [35] |
| 15 | 2017 | PbSS | Sesterbrasiliatriene | Type A | NSR | [19] |
| 16 | 2017 | PbTS | Pb1 | Type A | NSR | [35] |
| 17 | 2017 | FgGS | GJ1012E | Type B | SR | [8] |
| 18 | 2017 | PvPS | Preasperterpeniod A | Type B | SR | [19] |
| 19 | 2018 | CgDS | Dolasta-1(15),8-diene | Type B | NSR | [36] |
| 20 | 2018 | PcCS | Penichrysol | Type B | SR | [37] |
| 21 | 2019 | PrDS | Penichrysol | Type B | SR | [38] |
| 22 | 2019 | AbVS-1 | Variediene | Type B | SR | [16] |
| 23 | 2019 | AstC | Preasperterpenoid A | Type B | SR | [20] |
| 24 | 2019 | PchDS | Penichrysol | Type B | SR | [38] |
| 25 | 2020 | AcldOS | Ophiobolin F | Type B | NSR | [39] |
| 26 | 2021 | AuAS | Aspergildiene A | Type A | NSR | [40] |
| 27 | 2021 | BsPS | Bm3 | Type A | NSR | [41] |
| 28 | 2021 | FoFS | Fusoxypenes A | Type A | NSR | [42] |
| 29 | 2021 | CiGS | Pb1 | Type A | NSR | [11] |
| 30 | 2021 | CiSS | Sesterorbiculene | Type A | NSR | [11] |
| 31 | 2021 | ZbSS | Sesterevisene | Type A | NSR | [11] |
| 32 | 2021 | AaSS | Sesterfisherol | Type A | NSR | [11] |
| 33 | 2021 | CfBS | Bm3 | Type A | NSR | [11] |
| 34 | 2021 | CoFS | Pb1 | Type A | NSR | [11] |
| 35 | 2021 | CoSS | Sesterorbiculene | Type A | NSR | [11] |
| 36 | 2021 | ChVS | (-)-Variculatriene B | Type A | NSR | [11] |
| 37 | 2021 | ChPS | Pb1 | Type A | NSR | [11] |
| 38 | 2021 | PoVS | (-)-Variculatriene B | Type A | NSR | [11] |
| 39 | 2021 | CgSS | Sesterorbiculene | Type A | NSR | [11] |
| 40 | 2021 | CsPS | Pb1 | Type A | NSR | [11] |
| 41 | 2021 | LmVS | (-)-Variculatriene B | Type A | NSR | [11] |
| 42 | 2021 | PaSS | Sesterbrasiliatriene | Type A | NSR | [11] |
| 43 | 2021 | CsVS | (-)-Variculatriene B | Type A | NSR | [11] |
| 44 | 2021 | PoVS | (-)-Variculatriene B | Type A | NSR | [11] |
| 45 | 2021 | MpBS | Bm3 | Type A | NSR | [11] |
| 46 | 2021 | ChBS | Brassiteraene A | Type A | NSR | [11] |
| 47 | 2021 | AtAS | Aspterpenacids A | Type A | SR | [11] |

| | | | | | | |
|---|---|---|---|---|---|---|
| 48 | 2021 | AcldAS | Asperterpenol A | Type A | SR | [43] |
| 49 | 2021 | AuOS | Ophiobolin F | Type B | NSR | [11] |
| 50 | 2021 | PgFS | Fusicoccadiene | Type B | NSR | [11] |
| 51 | 2021 | BmOS | Ophiobolin F | Type B | NSR | [11] |
| 52 | 2021 | DvVS | Variediene | Type B | SR | [14] |
| 53 | 2021 | AcSS | Spiroviolene | Type B | SR | [11] |
| 54 | 2021 | SmVS | Variediene | Type B | SR | [11] |
| 55 | 2021 | TtPS | Preasperterpeniod A | Type B | SR | [11] |
| 56 | 2021 | PhPS | Penichrysol | Type B | SR | [11] |
| 57 | 2021 | TvPS | Preasperterpeniod A | Type B | SR | [11] |
| 58 | 2021 | BpVS | Variediene | Type B | SR | [11] |
| 59 | 2021 | AbVS-2 | Variediene | Type B | SR | [11] |
| 60 | 2021 | AaGS | Variediene | Type B | SR | [11] |
| 61 | 2021 | PaCS | Penichrysol | Type B | SR | [11] |
| 62 | 2022 | CgCS | Colleterpenol | Other | NSR | [44] |
| 63 | 2022 | MpMS | Macrophomene | Other | NSR | [44] |
| 64 | 2022 | AcAS | Aspergildiene A | Type A | NSR | [45] |
| 65 | 2022 | CcCS | Schultriene | Type A | NSR | [46] |
| 66 | 2022 | PstA | Penisentene | Type A | NSR | [47] |
| 67 | 2022 | NnNS | Nigtetraene | Type B | NSR | [46] |
| 68 | 2022 | TvTS | Talaropentaene | Type B | NSR | [44] |
| 69 | 2022 | TlnA | Talarodiene | Type B | SR | [48] |
| 70 | 2024 | EmES | Emerindanols A | Type A | SR | [49] |
| 71 | 2024 | PfNS | Nigtetraene | Type B | NSR | [18] |
| 72 | 2024 | AfAS | Asperfumene | Type B | SR | [50] |
| 73 | 2024 | TndC | Talarodiene | Type B | SR | [51] |
| 74 | 2025 | CcBS | Colcerepene | Other | NSR | [52] |
| 75 | 2025 | FoMS | Mangicdiene | Type B | SR | [9] |

**Table S2.** Mean validation performance across the five folds during training

| Fold | Mean accuracy | | Mean precision | | Mean recall | | Mean F1-score | |
|---|---|---|---|---|---|---|---|---|
| | training | validation | training | validation | training | validation | training | validation |
| 1 | 0.9519 | 0.9388 | 0.9354 | 0.9041 | 0.9239 | 0.9183 | 0.9186 | 0.9042 |
| 2 | 0.9358 | 0.7906 | 0.9060 | 0.7541 | 0.8975 | 0.7208 | 0.8896 | 0.7039 |
| 3 | 0.9249 | 0.8137 | 0.9155 | 0.7483 | 0.9012 | 0.8166 | 0.8943 | 0.7480 |
| 4 | 0.9510 | 0.9649 | 0.9341 | 0.9232 | 0.9276 | 0.9406 | 0.921 | 0.9302 |
| 5 | 0.9447 | 0.8436 | 0.9263 | 0.7563 | 0.9165 | 0.8306 | 0.9092 | 0.7755 |

**Table S3.** Parameter Configuration Summary for the Multimodal Protein Classification Model

| Parameter Category | Parameter Name | Default Value | Description |
|---|---|---|---|
| Data preprocessing | max_seq_len | 350 | Maximum length of protein sequences |
| | max_smiles_len | 128 | Maximum length of SMILES strings |
| Feature extraction | embed_dim | 1152 | Protein feature dimension from ESM model |
| | chemberta_dim | 384 | SMILES feature dimension from ChemBERTa model |
| | num_filters | [64, 128, 256] | Number of filters in multi-scale CNN |
| | filter_lens | [3, 5, 7] | Convolution kernel sizes in multi-scale CNN |
| Model architecture | fusion_dim | 1024 | Feature fusion dimension |
| | attention_heads | 8 | Number of heads in co-attention mechanism |
| | h_dim | 1024 | Hidden dimension in co-attention |
| | dropout | 0.4 | Dropout rate in fully connected layers |
| Training optimization | batch_size | 32 | Training batch size |
| | lr | 1.00E-05 | Initial learning rate |
| | weight_decay | 1.00E-04 | Weight decay coefficient |
| | minor_epochs | 500 | Training epochs per fold |
| | n_splits | 5 | Number of cross-validation folds |
| | patience | 100 | Early stopping patience |
| Loss function | lambda_smiles | 0.5 | Weight for small molecule classification loss |
| | lambda_label | 0.2 | Weight for protein classification loss |
| | lambda_supcon | 1 | Weight for contrastive learning loss |
| | temperature | 0.5 | Temperature parameter for contrastive learning |
| Prediction inference | W_centroid_sim | 0.3 | Weight for class centroid similarity |
| | W_onehot_label | 0.3 | Weight for label embedding |
| | W_onehot_smiles | 0.2 | Weight for SMILES embedding |
| | jitter_strength | 0.01 | Jitter intensity for visualization |
| Visual analysis | perplexity | 30 | t-SNE perplexity |
| | early_exaggeration | 8 | t-SNE early exaggeration factor |
| | learning_rate | 200 | t-SNE learning rate |
| | n_iter | 3000 | t-SNE iteration count |
| | bandwidth | 0.5 | Kernel density estimation bandwidth |
| | density_threshold | 0.05 | Density threshold for boundary points |
| | grid_size | 100 | Density grid size |

**Table S4.** BLAST-based prediction of SR/NSR results for characterized BFTPSs

| BFTPS | Closest homolog | similarity (%) | Actual type | BLAST-predicted type |
|---|---|---|---|---|
| AbFS | Fusicoccadiene synthase | 58.0% | NSR | NSR |
| AcldOS | Ophiobolin F synthase oblA | 88.5% | NSR | NSR |
| AcOS | Ophiobolin F synthase oblA | 65.2% | NSR | NSR |
| Au8003 | Ophiobolin F synthase oblA | 88.5% | NSR | NSR |
| CgDS | Cyclopiane-type diterpene synthase | 28.3% | NSR | SR |
| NnNS | Conidiogenone synthase | 34.3% | NSR | SR |
| PaFS | Fusicoccadiene synthase | 58.0% | NSR | NSR |
| PfNS | Conidiogenone synthase | 32.4% | NSR | SR |
| TvTS | Fusicoccadiene synthase | 43.6% | NSR | NSR |
| AuOS | Ophiobolin F synthase oblA | 88.5% | NSR | NSR |
| PgFS | Fusicoccadiene synthase | 71.9% | NSR | NSR |
| BmOS | Ophiobolin F synthase oblA | 65.2% | NSR | NSR |
| AbVS | Variediene synthase | 72.6% | SR | SR |
| AfAS | Fusicoccadiene synthase | 60.5% | SR | NSR |
| AstC | Preasperterpenoid A synthase PvPS | 72.8% | SR | SR |
| DvVS | Phomopsene synthase | 52.3% | SR | SR |
| EvAS | Stellatatriene synthase | 53.7% | SR | SR |
| EvSS | Bifunctional type I diterpene synthase tndC | 56.3% | SR | SR |
| EvVS | Variediene synthase | 72.6% | SR | SR |
| FgGS | Terpene cyclase ATR13 | 38.6% | SR | NSR |
| FgMS | Bifunctional type I diterpene synthase tndC | 43.5% | SR | SR |
| FoMS | Bifunctional type I diterpene synthase tndC | 43.0% | SR | SR |
| PaPS | Fusicoccadiene synthase | 36.6% | SR | NSR |
| PcCS | Conidiogenone synthase | 93.6% | SR | SR |
| PchDS | Cyclopiane-type diterpene synthase | 93.6% | SR | SR |
| PrDS | Cyclopiane-type diterpene synthase | 89.6% | SR | SR |
| PvPS | Bifunctional sesterterpene synthase astC | 72.6% | SR | SR |
| TlnA | Bifunctional type I diterpene synthase tndC | 91.1% | SR | SR |
| TndC | Stellatatriene synthase | 56.3% | SR | SR |
| AcSS | Conidiogenone synthase | 35.4% | SR | SR |
| SmVS | Variediene synthase | 46.7% | SR | SR |
| TtPS | Bifunctional sesterterpene synthase astC | 89.7% | SR | SR |
| PhPS | Phomopsene synthase | 56.3% | SR | SR |
| TvPS | Preasperterpenoid A synthase PvPS | 94.4% | SR | SR |
| BpVS | Variediene synthase | 43.2% | SR | SR |
| AbVS | Variediene synthase | 72.6% | SR | SR |
| AaGS | Variediene synthase | 69.8% | SR | SR |
| PaCS | Cyclopiane-type diterpene synthase | 81.6% | SR | SR |

**Table S5.** Accession number of putative BFTPS retrieved from NCBI in this study

| | | | | | | | |
|---|---|---|---|---|---|---|---|
| Type A BFTPSs, 103 proteins | | | | | | | |
| CCF32910 | PGH12319 | KIA75942 | TIC90236 | EPS41436 | RDA93524 | OBT63401 | XP_018154066 |
| CEJ53924 | PGH26795 | KIM93250 | TLD13740 | EQB55184 | RDH16203 | OHW89757 | XP_020051309 |
| CEL09395 | PHH80665 | KIM96055 | TLD16733 | GCB22886 | RDW62483 | OHW99200 | XP_020128736 |
| CEL10096 | POS72147 | KIN03517 | TLD27232 | KDN71386 | RLL97216 | OJJ67547 | XP_022394045 |
| CEN61919 | PSN71887 | KJX98376 | TLD33685 | KDN72248 | RYN29235 | OKP11162 | XP_022491462 |
| CRG86078 | PVH95144 | KKY29562 | TLS26487 | KFX91271 | RYP30765 | OLN89755 | XP_024687659 |
| EAU32979 | PYH87634 | KLO81457 | XP_001258098 | KFY54425 | TDZ21940 | OLN97949 | XP_024707785 |
| EAW14129 | PYH89575 | KMK63297 | XP_001260721 | KFY59930 | TDZ61748 | OLN98000 | XP_025401689 |
| EFQ25311 | PYI04682 | KXG54038 | XP_001262975 | KFY96141 | TEA17279 | OQD80556 | XP_025537260 |
| EKG16032 | QBZ56539 | KZL72782 | XP_006696713 | KFZ21954 | TGJ74928 | OQE05967 | XP_025564310 |
| ELA23316 | QBZ58073 | KZL82634 | XP_007701934 | KFZ23023 | THC99267 | OQE42161 | XP_025576901 |
| ELA37212 | QBZ60712 | KZL88287 | XP_009220267 | XP_028511096 | XP_028500364 | OSS53060 | XP_026603673 |
| ELQ36838 | RDA89093 | OBT40079 | XP_018151889 | XP_750866 | OXN24836 | XP_026604799 | |
| Type B BFTPSs, 95 proteins | | | | | | | |
| APA12276 | POS74342 | EYE97880 | XP_014550177 | CZT49095 | RMD43624 | KUL83692 | XP_025461812 |
| APY21859 | PQE19349 | GAO85920 | XP_016596824 | EAQ85668 | RMZ70974 | KUL85185 | XP_025486510 |
| AZQ56744 | PQE22675 | GAQ04021 | XP_018153114 | EED11493 | RWA09438 | KZL77275 | XP_025487035 |
| BAX09282 | PYI01416 | GAT28430 | XP_018187504 | EFQ93661 | RYO55048 | KZN86668 | XP_025501272 |
| BBF45518 | PYI14929 | GCB25423 | XP_018390852 | EJP62241 | SPQ20835 | OCK92653 | XP_025525081 |
| CAP86415 | PZD43626 | KIK55682 | XP_022486475 | EKG20455 | THV50721 | OCL09522 | XP_025555922 |
| CDM32996 | RAQ98630 | KIK55691 | XP_023093164 | EME87121 | TLD03666 | OJJ72250 | XP_025558733 |
| CEL04094 | RAR08378 | KIK55704 | XP_024677291 | EPE34883 | XP_001266466 | OLN96964 | XP_025589290 |
| CEL11848 | RBR18644 | KIN08340 | XP_024705680 | ETS85684 | XP_003067066 | OOF96881 | XP_026604901 |
| CRL20985 | RJE25610 | KOC10771 | XP_025395331 | EUC28897 | XP_003343918 | OQD85661 | XP_026614492 |
| CZR58890 | RLL93374 | KOS48568 | XP_025426141 | EUC38927 | XP_007677283 | OQE14697 | XP_026622213 |
| EUC50502 | PGH21525 | OSS43380 | XP_028489435 | XP_007694490 | XP_028488251 | XP_026631727 | |

**Table S6.** The amino acid sequences of NCBI BFTPSs after correction.

| No | Accession number | Sequence after correction | Length before correction | Length after correction |
|---|---|---|---|---|
| 1 | CAP86415 | MADKITDEYAVGIDPEIYANNPAYSSLFNPYIHKQTIIADH<br>VSVQCHIDLNGIDAVGSKFGNLNAHAGNFTSLCAPNCLP<br>ERFALVAYTVEYAFLHDDETDNAADQEALLLENKMLHQ<br>ALNQSGMTSVSTRVSDKAQRKSEVQAKIAAEYLRLDPV<br>FGEWFLNKWQTFTACVKDVRSLEFPSLDDYLEFRIVDA<br>AADWTLYNFRWGSGITLTPEEEKIADPMSYVAYAELCLV<br>NDLFSWDKEYASHIKSNGDVPLVNAVHIVAVTQGLTHC<br>AAKAVVQAEVRAHEERFCQLKEQYEATDKPSHEVLRW<br>LRLLEHSMAGNWVWSLCVPRYCKVDRNPYKDHLEKYG<br>SDAVRVLTPLDRLCWPKQEIKDMKQSELKDPSSSTYKS<br>HFSPLEPNPGPEQMRLTISQTQQQRPVLNPYTYINSLPS<br>KNVRQTLIAALNSWYKVPVKSLLIIEGAVNFLHNSSLLLD<br>DIQDGSFLRRGRPVAHQIFGVGQTINTATYLMNEALYLI<br>QMLSPSAVSVYTDEMRNLQLGQGRDLYWSYHTHVPTP<br>AQYISMVDGKTGGLFRLISRLMRSEATKNSDLDISQFAT<br>LLGRHFQIRDDYQNLQSEDYTKNKGFCDDLDEGKLSFPI<br>ILSMQSPGFSNTALSSVFKGSQKGQTLSLEMKQYMLEEI<br>TARGAFSETKAVLRKLHTELLRLLIETEKKAGGVENWAL<br>RLLIMKLDIAEEKKVAPPKSDSHWGVNQRRAWKGCQK<br>NGRPIDKACFLRAMEETLQK | 732 | 751 |
| 2 | EED11493 | MEFKYSTVIDPTTYETHGLCDGIDLRCHESPELEEIETLR<br>CQENWREWVGPLGFYKGGLGPRWNFMAITVPECLPER<br>LGVLGYANELAFLHDDVTDVADYGDIHNDDLKAAFEEAA<br>STGRIEGSTSGKRAIQAHIANEMMKIHKQHAITTLEAWA<br>KFAELGSGRQHTTHFKTEKEYIKYRMIDIGTMFWYGMV<br>TFGMGISVPEEELEMCHELANTAYLNLGLTNDLYSWQK<br>EYETAVAMGRDYVANIIGVLMEERKVSEEVAKEICREKIK<br>VTIVDFRKIVADTKARDDVSVDTKRYLEGLLYSLSGNLV<br>WSIDCPRYHRWSSYNERQLDWMKNGIPKSPKDLPKFN<br>GVSNGLANGTEKIVSNGHVNGNGAVHVPATNGVTNGN<br>GATHTPVSNGTNGTTGLKDPFSLDVDTDLVNVFARKEY<br>KAISAPKLHEGEGYPSALKLGPNGDVTIQKSPEIETRVIQ<br>APYDYISSLPSKGVRDQAIDALNVWCRVPAAKVDRIKLV<br>TNMLHNTSLMLDDMEDGSTLRRGKSSTHTIFGAGQTVN<br>AANYQIIRALEEVQKLGDKESVMIFTEELKNLYIGQSLDL<br>HWTNNVICPTVDEYFRMVEHKTGGLFRLFGRLLSLHST<br>NPVKADLTSLLDNFGRYYQTRDDYSNLTSPEYTTQKGF<br>CEDLDEGKFSLPVIHMMLSAPSNTVIRNIWTRRLVNTQA<br>SLSHKQAILELMKKGGSLQFTEDALDVLHAQVEKNISDL<br>EGKFGAQNFQLRLILEM LRKN | 745 | 754 |
| 3 | ETS85684 | MAGTRSSRPGATSFIQHSIPLPRSAYEGVEYFCRFRPRI<br>HRDAILADAGSWQCQVDFFGSSATARADSIRNKNHTSY<br>AVGCINPVVGNFTALCACEAIPDRLALTTYMVEYAYIHD<br>DVIEYAENKDEDRLWETNKELTQGLTLGDEVGAATKDN<br>VRRRQLQAKMAVELMDIDKVKGKECLRLWKEMSDVFV<br>QIRELKFTKLDDYLTFRVIDAGCPWTMSLLCFSMDFTLN<br>DDEVEKTAAITSAAYDGWVLVNDYFSWEKEWKNHQAN<br>GGSGVIANAIFLFMRWYSVDAVEGRRMLRKEILAREEKY<br>CKAKEEFLVSGNVTDKTSQWLELLDHVTAGNFAWSMT<br>TARYQLGGKDAYPALRAANTDNWETSTTDSLSNPISHN<br>ADKIARKINLIFKEQKFLDARGLVNHTEDYPPIVLTAQVS<br>QPDETPEFIHSQVTQARSFTQYEKMILQPQNYLESMPS<br>KGVRNSVIDGLEMWYQVPERSLATIRKIVNLLHSSSLML<br>DDIEDNSPLRRGLPATHTVFGISQTINSANLLMFKALKAA<br>ESLSPAAVRIFIERIIEGHIGQGMELYWTFHTEIPTEEEYF<br>VMVDGKTGGLFILLAELMRSEATRHKDLDTSLLMKLVGR<br>FFQARDDYQNLESAQYTQQKGFAEDIGEGKLSLPLIHAL<br>GSKTPQRGRLMSILQQRKSTVDLPFHIRKLALDDIKATG<br>GLKYAKKMAMSLQDSVNETLTQYEDKVGAKNWILRLVQ | 718 | 741 |

| | | | | |
|---|---|---|---|---|
| | | KRLELEV | | |
| 4 | KDN72248 | METFGKLGEAELYPYSLPIDRNEVVRSGALTSLPVRIHR<br>YNHLADAGALCLTDDWQRAMQDGQHKRSNADQNGLD<br>FCAISSTQAVFMMVDACEEISISAAHQEHLDFADAMEVR<br>DGEAKSRGNDHSHSVRIKNLISTAVLECIKVDRAGAMC<br>MLEAYGKKWLRVMETYKSNEIDNIDDYFEARANNGGM<br>GAYYAMLEFSLGIIITDSEYDLMSEAIRHVERCMLLTNDY<br>WSWPREREQAKHQETGKVFNIVWFLMHTNKSWSEDD<br>AIVKVRKMVYEEEQKWVEAKKSLYQKVPNLGRDAIKFLE<br>NLHTALAGNDYWSSQCYRHNDWKHVPELPGKCAPRV<br>NDLVGLGRALLRNESDLDDADNVAATVHSHKSSPGSPA<br>FHDGQSNHHSSTLSTHSQSSCTTPPSPRSPAINPHLPC<br>SSTSGFAKSPVTGPIDYIRSLPSKKVRTQLIDSLNIWFSV<br>PHSALSVIKEVVDCLHDSSLILDDVEDGSDLRRGFPATH<br>VVYGTGQAVNSATFLYVQAVEAVHRLVKEGGGRLELM<br>DMLLASLKELFQGQSCDLYWTHHRICPTEKEYLYMVDR<br>KTGAMMQLLVGLMQIAATSSDPRYGSLDPTHLEDKDGA<br>RQRLLRFTQLFGRFFQVRDDYLNISSCESSYMEKKGFA<br>EDLDEQKFSYILVHMYARNPEARDKVEGVFRAARQMG<br>KEKHDGIQWKRYILGLLEESGALNATRQVLVEWHKEMM<br>AEIGELEREFGAGNGMLRLLVEMLRI | 771 | 747 |
| 5 | KZL77275 | MSTYSTVVDPEAYETEGLCDGIEVRKSSFTHLEDRGAIR<br>AHQDWNKYVGPCREYRGTLGPKYSFMSLTVPECLPDR<br>LEVISYANEFAFLHDDVTDQVDFERGEVENNEMMKAFL<br>EAAHTGTIDSSVSEIARAGKKKIQSQLFLEMLSIDPECAK<br>TTMKAWARFVEVGSSRKHETRFQTLAEYLPYRIMDVGE<br>MFWYGVVTFGLSLRIPEHETDICRELMAPAWIAVGLQN<br>DLWSWPKEKDAAKKQGKDHVINAIWVLMQEHKTDVRG<br>AKQICRRLIKEYVAKYLQVVKESRNNYSISPDLRKYIEAM<br>KYSISGNVVWSLTCPRYNPQVSFNERQLDWMHQGVPE<br>IEVSSSPSPEAESISPGSPILSESSSKNYSHSDEEGVLG<br>MNSSQSSVSTPPEVDGDEQQIKLGPHLAYEREILEAPY<br>NYIASLPSKGVRDRFIDALNHWLQVPHSVLTKIMEVTRLL<br>HNASLILDDFQDDSPLRRGKAATHTIFGPAQAINTSSYTI<br>VKTINHVAEFASNNCIQETTQKIMALFQGQAMDLFWSYN<br>GRCPTMDEYYRMVDNKTGQLFSIATGLMLDSQKRTSTK<br>APASLDRFTSLLGRYFQVRDDYQNLTSVDYIRQKGFCE<br>DLDEGKYSVPLIYTLQTQPDNIQLINLLSTGRRTGRLTRE<br>QKECVLEILNKFGGFAYTRVVLLRLYSQISDALKELESSF<br>GSSNPEMKVLLLELLRIEN | 663 | 714 |
| 6 | OCK92653 | MDYRNSSVMKIVVAEVENLCADIPLRISNNPQLEVRGIQ<br>RAQDDWRTYFGPLEAFNGAIGDQFNFMAVCLPECLPE<br>RLEILSYANEFGFLHDDIIDEGSDQHVKEDGIKQIQAKML<br>LEMIKIDRERALTAAKAWKEFGDVTAQERKSFGSFDEYL<br>AFRFGDSGERFWFGIATFGMGLTIPAKEMELANRMGRS<br>AYKAGGLTNDLYSWEKELAVSKANGESHVVNAIWVLM<br>QEHSCSVEAATLLLREKIKECVSDFVRIAKEVEGNLELSI<br>DTRKYIESALYGISGNFHWSLQCPRYRPNPLERGTPQR<br>HDTAATGGNPTAEGDADRTSTSEDKPESSSTLNEVIQD<br>ERASKHPKLECNGNLDNKPANGINRNRSAAIEDGAEIFL<br>TEGFPALHDEVVQAPFLYISSLPSKNVRSTAVDALNIWL<br>EVPASSLDIIKSVINKVHATSLMLDDLEDSSPLRRGHPST<br>HTIFGIPQTVNAANFQLVAALEEVQTLNDVRCLRIFTEEL<br>QKINLGQSYDLFWTCNIECPTVSEYLNMIDASKAHNSRR<br>LLLDTDIYSETGGLFRMLVRLMAARSPISEHPAISNVVTL<br>FGRYFQIRDDYQNLVSADYENQKGFCEDLDEGKYSLPLI<br>YALQTEKPNLWLNNILTQRRVAGKLSFEQKRAVLEDMN<br>RKKCLKRTAEMLSILHKAIEQNIESVEKDLGKVNFPLRLL<br>LETLKI | 689 | 706 |
| 7 | APY21859 | MDFTYRYSFEPTDYDTDGLCDGVPVRMHKGADLDEVAI<br>FKAQYDWEKHVGPKLPFRGALGPRHNFICLTLPECLPE<br>RLEIVSYANEFAFLHDDITDVESAETVAAENDEFLDALQ<br>QGVREGDIESRASGKRHLQAWIFKQMAAIDRERAIAAM | 734 | 767 |

| | | | | |
|---|---|---|---|---|
| | | NAWATFINTGAGCAHDTNFKSLDDYLHYRSTDVGYMF<br>WHALIIFGCAITIPEHEVELCHRLALPAIMSVTLTNDIWSY<br>GKEAQAAEKSGKPGEFVNALVVLMREHNCSIEEAERLC<br>RARNKIEVANCLQTTKETRERTDVSQDLKDYLYHMLFG<br>VSGNAIWSTQCRRYHMDAPYNERQQARLNQKKNKLSS<br>TYDPIQASKEALKESTRPEIHRLPTPDSPAKESFAVRPLV<br>NSSGEFDGNGYIDGHEANVRPKPVVNGHSDAEERPAF<br>ERKGSAKRATSANDIDWTAHKKVDHGNGHKKTLSDLML<br>QDLPPMEDDVVMEPYRYLCSLPSKGVRNKTIDALNFWL<br>KVPAERANTIKSITESLHGSSLMLDDIEDHSQLRRGKPS<br>AHAVFGEAQTINSATFQYIQSVSLISKLQSPKALGIFVDEI<br>RQLFIGQAYELQWTSNMICPSLEEYLQMVDGKTGGLFR<br>LLTRLMAAESTTQQDVDFSRLCQLFGRYFQIRDDYANL<br>KLADYTQQKGFCEDLDEGKFSLPLIIAFNEQNKAPKAVA<br>QLRGLLMQRCVNGGLSFEQKVLALSLIDEAGGIEGTEKV<br>LHSLYGEMEAELERLAGIFGEENCQLELILEMLRID | | |
| 8 | OQD85661 | MLPRSQDSPKHFVPDSVPVPREAYEREEYFCKFTPRIH<br>RDAHLADSGCWQCQVDLLGSQGAAHAAAVRSGSNKS<br>YAVGCINPTVGNFTALCAVEAIPERLALISYIVEYAYVHD<br>DLIEYAEEKRDSELYKANKQLTEGLEEYGNEKLSTRSNV<br>KRQMQAKMATELLDLDLEQGQTILRLWKEMSNVFVDIR<br>DLDFQTLEDYFIFRAVDAGCPWTMALMCFSMDFYLTDE<br>ETAQTSSITSAAYDAWVLVNDYFSWEKEYKNYEANGST<br>GQIVSAVFLFMKWHNIDQSTAKKMLRGEIISREERYCQL<br>KADYFARGNVTSRIIKWFQLLDIVTAGNFVWSMTTARYH<br>DDAEDAYPGLRAVNQKKKDSRPFSSLSTPIAPLRINAKR<br>QQSVGLVAPDSPGSLGSSSDSEVDTGLTSPASSKPEQ<br>RKDSIPVPLPAFEEISLEPYVYLKSLPSKGVRNAAIDGLD<br>VWYQAPERSVILIQEIADLLHSSSLMVDDIQDNSPLRRG<br>FPATHMVFGIAQTINSANLLLLKALKAAMTLSASAVNIFTT<br>RLLEGHIGQGMDLNWTNQTHVPTEEEYFTMIDGKTGGL<br>FVLIAQLMRSEATTNKGLDVSTLMKAVGRFFQVRDDFL<br>NLQDADYTKKKGFAEDLSEGKISLPLIHTLAEQSQHRSR<br>LLNILQQRKTGRFLSPELRKLALDDITATGGLDYTKRTLL<br>GLQKDLDTYLTRTEEMAGESNWILRLMKKRLEI | 671 | 729 |
| 9 | OSS53060 | MEALSYQSNLISSDESRKTGCFTTLPVRLHSRDDLADAA<br>SRRFVQDWAREMGDGREQNTYFSFSPVGNWSSLIYPE<br>AIPERLGVLAYLSDLGLIHDDGGEGLSIDDAQAENDNLY<br>AALNPDDKESRTTNLRATKMKKLVSQCILECISLDRELG<br>LKMLAAFRDVWLAISERNSDKEAHTMDEYLRYRSDNGG<br>MLVFWPMLQFSLGISISDEEEALVQPIIDAATEGLLLAND<br>FFSWEREYRELQSGQSKRIVSAVELFIRTEKLTVRDAKE<br>EVRRRIIKSERQFCQRRDELYKTHPDISLKLKRWIDCAG<br>LAVSGNHYWCSACPRQNSWKEDGATNGSSNKRKISDN<br>AIVSTQEVPLKKRKDSSVFDSKRSDGGSESSEVSNYPY<br>YKPSGLALEAPSKYVSSMPSKGVRSTLIEALNTWLHVPH<br>ERLQAIMAVIDSLHNASLILDDLEDNSPLRRGYPSTHILF<br>GQAQSINSANFMFVRAVQQVAQNLSPAALTAVLEELEG<br>LYLGQSWDLYWKHNLACPSEAEYVNMVDHKTGGMFR<br>MLLRLMQAESEETSQLDFHVLTLLFGRFFQIRDDYMNF<br>QDYTSQKGLCEDLDEGKFSYPVVYCLENFPEYRGHFLG<br>VFRQRPTVATVNASPLSKESKLHLTACLKKSGAFEKTIA<br>SLTDMERDLEVEINRLEQQTGEANPMLRLCLAKLSVKGI<br>GKVGEPQPPYAGRLTLRSR | 688 | 711 |
| 10 | TEA17279 | MPSFSQPVPDDVVISSGLRSKFRPCVHQNYDRCVEAAK<br>GMESVYNQVMSTHIESKTLAQVPGLGQVHPMALTIANC<br>LPERLPAITRFADFTILNDDYYDVAKREEIEQVNDGIQSA<br>LQDALASSKGTETTTASDSTFKSKQLQAALVLELMMLD<br>QQFAMEIMSSYSQGLEVATFAPDDLKTLDEYLPVRKINS<br>GLDVTAEMACFGTGLKISRQEKDKLREVVDLATFAITVV<br>NDVYSWPKEIKCHLETPGSNAPFNAVAILMRHGGHSEA<br>EAFQILYDKQAELEEQHLQLLEKLRSQEGGSLPENQELY<br>IANAQQAVSGSELWSIYTTRYPSKEDLQQPRIEFVDGTF | 690 | 758 |

| | | | | |
|---|---|---|---|---|
| | | RYAAEVDTLSEEEVLPSVVKSTSAEVEDQLALLDLTPVS<br>SAEGISTPSSSVSCGCDTEEKKPLSEDTSSHVTHGALQ<br>NLTASTSNPTYKRMKALLPDDPGLKTYASRVAAADDHP<br>AVGPFKYLAMLPSKGVRDTFIDALNWWLKVPEDSLRSIK<br>TIISMLHDSSLILDDIEDDSTLRRGSPAAHMIYGTAQCINA<br>ANYMVVMVLAELQKLHSPAKTMILIEELESLFLGQSEDL<br>HWKYHVECPSTDEYMEMIDNKTGGLFRLCVRLLQAEST<br>RIDVSHLDPRPFVRQLSLFFQIRDDYQNLVSDQYAKQK<br>GFAEDLDEGKISLPMILTLQRARTRPEIMGIMKHKKPGP<br>MAPEMKEFIVREMRKSGALETTRELLRGMQEDLITELRK<br>LEEEFGAKNPTLEL VLRKLWIS | | |
| 11 | PYH89575 | MMPPQVVPIPDDILKKSGMVSIFPTVIHPDYALVNADAAK<br>LIDDFNAREKHRHQHPYHRRHPRAMDYYDKAGIQKANK<br>YNDELRGHLKTDVDDEDAKASMAAKAQQLQAGLLVEM<br>LQIDQTLASDVMATYSRILEAMGQPGKGGDIKTIEEYLPF<br>RLGNSGIEVYQEMCSFGPGVQLSRSEKERLAPITNAAL<br>CSSYLCNDYCSWPKEVKAYFEQGDNPPSAQAEARQNV<br>CEEAVAQQRTHLALIEELEKSEGPLSQEHRRYFEAAQC<br>TASGSELWSVYTARYPTKAELNQPECILVGEILQLVDPA<br>GTTEHPSDPQPVVTEVKKSIVPKQRFFHRGEERPAVGR<br>RQEAPAVGGGAKHHTNGHHTNGHSSHSSGSSNGMHA<br>AGPSKPQNGTGDGFSQASNELQVPRHAVSTIKKIVGLL<br>HRSSLMLDDIEDDSALRRGRPCAHTLYGAAQTINSANY<br>AFVGAFAEVQRLHSPSAAGIFISKLPAQRNLSRIVTHCPP<br>GEVQNMHCGRAMDLSWKYHTHCPTVDEYMTMIDNKT<br>GAMFRLCVRLMEAESTVPCQDIDSSHFVTQLGRYFQVR<br>DDYQNLVSTEYTNQKGFCEDLDEGKISLPLIYTLMNPAG<br>LGAIRYQGHLPAQGEWRIAAPRVKKCILEQMEEAGALD<br>ATRSLVRKMQTDLISELHRVEECFGSKNSLIELVLRRLWI | 678 | 688 |
| 12 | ELQ36838 | MLSPEVLYPYSIPVARETVVKSGSLTSLPVRIHRHDSLA<br>DVGAKCLTGDWGRIMGDGQEKKSNGSPSVAGNWGSFI<br>WPECLPDRLGLLSYLLDAGCLHDADVCEEMPIVAAHAE<br>HLDLYEAMNFSDSRVLDLGSQSTKIKDLVSTVILECMRV<br>DRVCALRMLEAYRKRWQAIIETRDTDKIESLEEYLESRA<br>NNGSMGAYYAMLEFSLGIVITQEEYSLLAAPMRHWKRC<br>MLLTNDYWSWLRARQQTKNQLVGRVVNTIWFLIKQNPT<br>YTDEQAKSKVAEMVVAEEAKWVQSKNRLYAENPKLRD<br>DLVKFLENLHTALAGNDYWSSQRYRHNDWTHVPTQPE<br>PDHPRIHELAGFGRTLVVEESRILARPSSSSSPSHSLGA<br>SASSSSGWASSSSQGDLSSPLSSASSTFDNVDRDRLK<br>HKVTKATTTTAATAEEVISAPIRYIQSMPSKGFRAALVDC<br>YNMWLDVPQREMEIIKTVINCLHDSSLLLDDIEDDSKLRR<br>GFPATHLVYGPAATINSATYLYVQAVQAVHNLGSRKLM<br>DVLLLRLQQLFFGQSLDLHWTFNRECPTKTQYFNMIEQ<br>KTGAFLSLVAELMMVVSAASDYESSTDSPENIFQTFSYQ<br>SGLYFQVRDDYLNITSADYADKKGFAEDLDEQKFSYILV<br>HLAHSRPDLMVQVEGIFKAMRSGKTENAETKKYLVSLL<br>HKSGAVEATKELLVRWQDDIVKEIDVLENHFGKPNPSLR<br>LLMQTLWIDA | 713 | 739 |
| 13 | XP_018187504 | MLTPAPIPMEDTHFRAQGAPAEQSFVYQHSIPVPPQAY<br>EREEYFCKFTPRIHRDAYLADAGSWQCQVDFLDSSDAA<br>RADAVRNQAHKSYAVGCINPIVGNFTALCACEAIPERLA<br>LTTYMVEYAYIHDDGEYFTKYLRETNQELVEGLLDEEVK<br>AGGKDHVRRRQLQAKMAVEMMEMDKEQGLECLRLWK<br>EMSDVFVQIRDLNFTKLDDYLYFRGIDAGCPWTMSLLCF<br>SMDFKLTEDETKSTARITSAAYDAWVLVNDYFSWEKER<br>QNYEANGSVGEIVSAIFIFMKWYSVDEKEAKRMVKKEIM<br>AREEKYCKEKADFVGQGNLTEKTAKWFELLDLVTAGNF<br>AWSMTTARYNLDNEDAYPALRAEHSKKKTATSVDSLTL<br>PISQNVAKSIKQKKEESNEQEYADGSSISSTISSSTPPTT<br>PGDPTDAIDEKIGNVALPVPRLVPSLTQFEELVLEPYAYI<br>ESLPSKGVRNTAIDGLEAWYHVPEKSLEIIRDIANSLHAA<br>SLMLDDIEDESPLRRGYPATHTVFGISQTINSANLVMFK | 700 | 747 |

| | | | | |
|---|---|---|---|---|
| | | ALRAAKSLSPAAVSILTERIIEGHIGQGMDLHWKHHTEIP<br>TEEEYFAMVDGKTGGLFLLLAELMRSEATINKDLDIGPL<br>MEAVGRFFQVRDDYQNLENADYTKQKGFAEDISEGKLS<br>MPLIHALASKSLHRGRLLSILQQRKAGNDLSHEIRKLALD<br>DIKAAGGLEHTKSIALELQKTVDKALTNLEEKTGVKNWIL<br>RLMQKRLEL | | |
| 14 | XP_025395331 | MEFKFSTVVQPETYDAQGLCDGLTVRYHKNSEIEEIDCL<br>RCQEHWRELIGPLGLYKGGLGHPWNGMSIAIPEALPER<br>LGIVSYANELAFLHDDVTDIAQYGDEHNNDLKAAFDQVI<br>RTGTINHSGSGKRALQAYVAKQMLSIDKDRAITSLKAWS<br>TFLDKAGRQEECRFKSEEEYLKYRIHDVGTLFWYGLLTF<br>AQAITIPEDELDMCHQLATTAYLHMALVNDLVSWEKEH<br>QSAVALGKDYCTNFIFVAMEEHGISEGEAQERCRHEIKR<br>ATVDYLQIFEDAKARNDLSHDTMRYLESVLYSMSGNVV<br>WGLHSPRYYTNASFSERQLEWMKNGIPKENKPEGGEF<br>GFGQVEVKMPDRTVNGNASLPASVPEQTANSLEADTSL<br>LSAVIKEHLVGREGFKLGDHDVKVESCNGQDLDSKVGS<br>TAALPGRVLQAPYEYITALPSKGLREQAIDALDVWFRVP<br>KEKLNIIKSITTILHNASLMLDDLEDGSELRRGKPSTHAIF<br>GLGQTINSANYQLIRALQRVHELGDSRSLVVFTEELHNL<br>YVGQSMDLYWTSNLICPSVNEYFQMIEHKTGGLIRLFCR<br>LMALHSTSPVKADLVDFSNRLGRYFQTRDDYQNLVSAE<br>YTKQKGFCEDFEEGKFSLPLIHLLQAMPDSYVLRNVWM<br>QRRVSGTATHAQKKTILDLMKKSGSLKFTSDTLGVLYG<br>HVEKSIDELESRFGAENFQLRLILELLRNG | 716 | 724 |
| 15 | XP_025525081 | MEFKFSTVVDPSTYQTQGLCDGLTVRYHKNTELEEIGC<br>LRCQEHWREQVGPLGLYKGGLGHPWSGMSIAIPEALP<br>DRLGIVSYANELAFLHDDVTDIAKYGDEHNNDLKAAFDQ<br>VISTGTIKHAGSGKRALQAYVAKRMLSIDKDRAITSLKAW<br>PTFLDKAGRQEDYRFKSEDEYLKYRVHDVGMLFWYGL<br>LTFAQAITIPENELEKCHQLATTAYLHMALVNDLVSWDK<br>ERQSATALGKDYVTNFIFVAMEEHGISEDQAQERCRQEI<br>KKATVDYLRVFEDIKARDDLSPDTKRYLESVLYSMSGNV<br>VWSFHSPRYYTDASFSERQLEWMKNGIPKAQTSERGA<br>SGTDKSEVKIPDRTVNGDHTAPNSNGTGIGSHGTINGN<br>GTAKENNSRHVSTPGRNTKGLSNGDTSLLSAVITQHLS<br>GRNGFKLGDHDMKVESFKKGGHDIKEVESTLKGQDLD<br>AKGVREHAIDALNVWFRVPAEKLAIIKSIITILHNASLMLD<br>DLEDGSELRRGKASTHMIFGLGQTVNSANYQLIRALQEI<br>QKLGDSKSLLIFTEELHYLYVGQSMDLYWTSNLICPSINE<br>YFRMVEHKTGGLVRLFGRLMALHSTNPVKVDMIDFSNR<br>LGRYFQTRDDYQNLVSAEYTKQKGFCEDLEEGKFSLPLI<br>HLLQTMPDNHVLRNVWMQRRVRGTATHAQKETILDLM<br>KRNGSLQFTEDTLGILYGHMEKSICELERKFGAENFQLR<br>LILELLRNG | 730 | 740 |
| 16 | XP_025564310 | MDCWQHSRPIDAEAIKKTPSFTTLPIRINERTDVADSATR<br>RALRDWDHCLHDGLADKALISISELGNLGAFAYPEVPPE<br>RLAILTYLTDLGMLHDDGYEAMDMDQARAEHRDFGALF<br>DPDGQPLSSRRSTRAPKLKKLVSQILLEAIRIDRDLGMY<br>MFDMYNKGWLSAAGGKEGKAPQFNSLEEYQAYRRDD<br>FGIRAFWPMVEFGMAMRLSDEDKKLIEPVMEPIDKAIIW<br>TNDYWSFDREYHESVTSGSRLTNVVEVVRRTENISINEA<br>KAKVRQLLVDLEQQYLERKSKFFAQNPSLPFHVRKWVE<br>VAGITVAGTHYWASCAPRHHTWRNHSINVQDPNESNA<br>HRGYSTETDMPNNPSFSNPSSEDAKLTPARLNPEINAQ<br>EFMNLNAQLALDKHDIEQFMRAVLAILSKVVKHCESLFG<br>GRDHEEAKLPQADEATKRVVSFKACSKETQEAEGLWY<br>KPTDTALQAPIQYICSMPSKGVRSRMIEALNYWLEVDEA<br>SLNKIGQLVDILHNASLILDDIEDNSPKRRGKPATHTIFGH<br>SQAINSANFMF<br>VQAVQVARQFRSPNAVEILLEELENLYLGQSWDLCWKY<br>KLKCPSPSEYLNMVDNKTGGLFRLLLRLMQAEGKSTAH<br>DEVNMNGLTILFGRFFQIRDDYMNLRSGLYIEQKGFCED | 763 | 770 |

| | | | | |
|---|---|---|---|---|
| | | LDEGKFSYPIVHCAANHAGFRDLIDGVFRQRPTTITTSG MQPLALEVKQHIVEYLDTTGTFQHCREYLLQLERSIICEI DRIEKVTNEANPMLRLLLEKLSVKYD | | |
| 17 | XP_018390852 | MDRPKQASDDETSYLVSLSHYDTHSLCSDYTLRRHKYE SEANAGCYEARQDWIKYVGPIEEFGGCNPINGNFSAVV LPLTKPERLRMIAYVLECTIPVPYGFPITEVVCLLNADAFL YDNMIELEKPETSRADNQNKREWKNTNPALGKTQIQAKI MIQLASIDYACTERVKSVWTEMIDTTLRDKNKCFTHLEE YVNFRMIDTGAPFVEAMLLFGMGVTLSKEEDIKLEQIRK PCYAALGLANDFFSFDREYADFEKSGKSQTLTNAVWLH MQWYNMDVNAAKSMTLEATKRYEEQFLGSFAEYDPNV GMEDGLTAESLQKTLCLEYDAYIRTEVKEERDESSRRS STDSSLQASEDDSFTTRDDVSISSYTSVSPSTLKDDQIL CNDLLGQENVQAPFDYITSLPSKGVRDTFIDALNIWLNV PETVVSRIKSLGNRLHAASLMLDDIEDGSNLRRGQPATH TVFGIAQTINSGCYEVLQAVSEAQELGAAEVKIVFEELAE LHIGQSYDLFWTYHTQCPSENEYLEMVNKKTGGLFRLL VRLLLESADCELLRVTQGADIEELVSCIGIQYQVRDDYQ NLHSPEYSAQKGFCEDLDEGKMSFPLVHALKSCEDNAE LYELLRQRRDIGYLSDEQKKLVLGQLDRAGSMTYTRDT LRRLQGEIHAGLKRVEDVTGRENWILRALLQRLEV | 688 | 692 |
| 18 | XP_016596824 | MADKIADEYAESIDPEIYANNPAYSSLFDPYIHKQTIIADH VSVQCHIDLNGIDAVGSKFGNLNAHAGNFTSLCAPNCLP ERLALVAYTVEYAFLHDDETDNAADQEALSLENKMLHQ AINQSGMTSVSTRVSPKAQRKAEVQAKIAAEYLRLDPVF GEWFLKAWQTFTASVQDVRSLEFPNLDDYLAFRIVDAA ADWTLYNFRWGSGITLTPEEEKIADPMSYAAYAELCLVN DLFSWDKEYAAYVKSNGDVPLVNAVHIVAVTQGLTHCA AKAVVQAEVRAHEERFCFLKEQYEATSSPSESVLRWLK LLEHSMAGNWVWSLCVPRYFKVDRNPYKDHLAKFGSD AVRVLTPQNELLDSKKETKETEQSVVKAPSPKTTVESDP VLNPYAYINSLPSKNVRQTLIAALNSWYKVPVKSLLIIEGA VNFLHNSSLLLDDIQDGSVLRRGRPVAHQIFGVGQTINT ATYLMNEALSLIQMLSPSAVTVYTDEMRNLQLGQGRDL HWSYHTHVPTPAQYIGMVDGKTGGLFRLISRLMRSEAT TNSDLDISQFATLLGRHFQIRDDYQNLQSEDYTKNKGFC DDLDEGKLSFPIIVSMQSPGFSNTALSSIFKGSQKGETLT LEMKQYILEEITARGAFTETKAVLRKLHTELLRLLVETEKL AGGIENWALRLLIMKLDLGDEKKKVAHKTDSAWKINQR RAWKGSQKNGRPIDKACFLRAMEEATQK | 684 | 728 |
| 19 | XP_024677291 | MEGQKHGVNMAEDGVAKSHPLMYRHSDPALMDAETK SKYISRFQPRLSKYVVTVSHDACIECHIDLFGTGGIGKVVG GMNAHTADFTALCAPEALPERIAFCSYFIEYAFVHDDVS VDAVHDGPSCRLSQDKDGILLKAVGSKRLEDLEDIAEG GLKRKQARAKIWSVLNEIDQDYLGRCQKKFKQWYETG QQMRDKTFASLEDYLAVRALDCGANWVVRMMGWASG VELSPEEEVETGPVTYLAFVVLGVTNDLWSWEKEKRVT RQSGDTLPLINAVKMVMQVQDIGEESAKQTVRALIREHE EQYCLLRDEYLRRPSTSLSVKKWFQVLELSMAGNALWS IHALRYHPDVRNPYICPTEIPSVFKELEVKRSKPESIKKG NAGVGSEDATGPAYKVLEALDETVCLPLHPEATLITAGS METIRVYHLYVIQGFPASPDRCSADLVQCARALGGADF GVASLLHEASLMLDDIQDGSLLRRGKPAVHEIFGVGQTI NSACFQINNSLRLVQQLSPSAVVIFSEQMAHLYTGQAH DISWAGHRIIPTEEDYFRMVDGKTGGLFVLLMRLMQQE ATQNRDLDMRRFMNQLGRYFQIRDDYQNLVSQEYTSQ RGFCQDFDEGKPSFPFIRACHTLEDSTVLMEWLNMLRS GGGASIEAKRYILTRVEESGSLEYTRGVLALLLQDLEGTL RDMEVMTGQENWILRSMLVQLQIKQEKPGDGVEVDEL NIGLAEYLVIRVLRQPSKKEFSAAIPIPPLK | 649 | 756 |
| 20 | KIM96055 | MSTQLVGDPNTTRPVPESLWRDAPEYFSRFIPRIHRDF QVCDDATIQCRIDIVGRGQPFIEAGNLCLRAGNLIALTMP DCIPERLALAAYLIEAGFVYDGVQYVRNLNTSVYSGTVR | 697 | 840 |

| | | | | |
|---|---|---|---|---|
| | | RLGTNLYVPQAELDGEAANADTQITTDKSLYEIKLRNKQ<br>VLSKMSVQVMRFDRQLGENVIRGWTTWKKGEEIILPKT<br>TGKEYQNLNEYLPDRFDDIGWPMVINLLHWGTGVVLDE<br>SSATEIAHILKPIHDVVTLTNDYFSWEKEIAEHIQLNEATP<br>LINSVAFYMKWDSLSATDAKAAVKAKIQQLEEEYSALKT<br>EYIARHGDKTPSSVIRWFSIMEAVVAGNLIWSNFTPRYH<br>VSNIKAHEYEEYYARRIREGASFFDSCTESDAFIPPVSQ<br>ETTSELRWSTRHSRVDCINGGSHESTKAQNLASDKTTA<br>AEDRQDQKIVTQPSTEDSHKRKRRGSVDARQDSATQQ<br>RNSETATLPPESDLGNRFDTKKKQRKVNGVSHDHKEE<br>QNGDAPRKVNGHMNHHSDGAQNGSINGEKATHIVDYR<br>QPRLPDLPEDIINRPFQYIESLPSKGVRSLVLEALNSWYL<br>VPQQQMKQIDSIVRSLHNASLMIDDIQDQTDLRRGKPAT<br>HMIFGIGQTINSAYFQCIEALKRASSNSPSMFEIVIDEFRT<br>GHIGQSLDLYWTFQTVVPSEKEYLQMVDAKTSSLFRLC<br>SRMLRSAATANRDLEIENLATLFGRYFQIRDDYMNLKSQ<br>DYTNQKGFCSDLDEGKYSLPLIHALREPDPKLESTLKQR<br>LQPGGLTVNSKQWILDRLMEKGSLQYTERQIELLYSAIE<br>DCLDNLEKKTGTINWMLRRILYQLQN | | |
| 21 | RYO55048 | MDRPKQASADETSYLVSLSHYDTHSLCSDYTLRRHKYE<br>SEANAGCYEARQDWIKYVGPIEEFGGCNPINGNSSAVV<br>LPLTKPERLRMIAYVLEYAFLYDNMIELEKPETSRADNQ<br>NKREWKNTNPALGKTQIQAKIMIQLAFIDHACTERVKSV<br>WKEMIDTTLRDKNKCFTHLEEYVNFRMIDTGAPFVEAM<br>LLFGMGVTLSKEEDIKLEQIRKPCYAALGLANDFFSFDR<br>EYADFEKSGKSQTLTNAVWLHMQWYNMDVNAAKSMTL<br>EATKRYEEQFLGSCAEFRQNNAPIPEKLDRYLDALAYQI<br>SGNVVWSLNCPRYHREYRYDPNVGMEDGLTAESLQKT<br>LCLEYDAYIRTEVKEERDESSRRSSTDSSLQASEDNNFT<br>TRDDVSISSCTSVSSSTLKDDQILCNDLLGQENVQAPFD<br>YITSLPSKGVRDTFIDALNIWLNVPETVVSRIKSLGNRLH<br>AASLMLDDIEDGSNLRRGQPATHTVFGIAQTINSGCYEIL<br>LAVSEAQELGAAEVKIVLEELAELHIGQSYDLFWTYHTQ<br>CPSENEYLEMVNKKTGGLFRLLVRLLLASADCELLRATQ<br>GADIEKLVSCIGIQYQVRDDHQNLHSPEYRAQKGFCEDL<br>DEGKMSFPLVHALNNCEDNAELYQLLRQRRDIGYLSDE<br>QKKLVLGQLDRAGSMTYTRDTLRRQQGEIHAGLKRVED<br>VTGRENWILRALLQRLEV | 733 | 713 |
| 22 | TLD13740 | MLSQEALYPYSVPVARETVGRSGSLTSLPVRIHRHNSLA<br>DAGAKCLTDDWDRILRDDQDKKSNGSLSAVGHRGSFI<br>WPECIPDRLGLLCYLLDMGCLHDDVCKELSIVAAYAEHL<br>DLGAAMCSSDSRVLSVDSRSAKTKHLVVQHHPRVWLAI<br>TETYETTDKIDNLDEYLESRANNSVMGAYYAMLEFSLGI<br>FITQEEYSLMAAPIKQVGRCMLLTNDHWSWSCDRQQV<br>KTQKATKVLNTVSFLIKQNPTYTEEQAKHEIADMVVAEE<br>AKWVQSKNRLYAENPDLGDDVVKFLENLHTALAGNDY<br>WSSQCYRHNDWTHVPTQPVPCHLHVYELVGLGRAVM<br>AQESRILAGASSPSQDRRKSTSSQEDLSSPPSSVSYTC<br>DNTDGDFLNHKVTKAITTTTITKAEEVISAPIRYIQSLPSK<br>GFRAALVDCYGRWLDVPQREMNIIKTVINSLHDSSLLLD<br>DIEDDSKLRRGFPATHLIYGSAATINSATYLYVQAVQAVH<br>DLGCKRTMDVLLRRLQQLFLGQSLDLHWTFNRTCPTKA<br>EYFGMIEKKTGAFLSLVAELMMAASVSASASAPRDCGS<br>STEGPENVFEAFSYQSGLYFQVRDDYLNITSADYADKK<br>GFAEDLDEQKFSYLLVHLAHARPDLMVQVEGLFKSMRS<br>GRVENAETKKYLVSLLYKSGAVEATRELLVRWQDDIVK<br>EIDILEKHFGTPNPSLRLLMETLWI | 702 | 714 |
| 23 | RAR08378 | MSTSDSSKAYADATSYIAEANAGCLEARSDWTKYIGPT<br>EQFGGCNPINGNLTAVALPLTKPERLRLIAYVFKYAFFYD<br>DGVEQEQPNTSNVDGNQKLEWKNAQTLGKMQMQAKM<br>MTQLSSMDSACTERIKRAWREMLSTTLRDKDKKFDSLE<br>DYMDFRIVDVAGAWTEAILLFGMGMTLSEEEDAQLECIR<br>KSCYAALGLANDYFSFDREYADFQKPGESQTFINSVWL | 688 | 781 |

| | | |
|---|---|---|
| | | HMQWHNVDVAAAKRMTVEVTRRYEGLFLNQCAKYRHE NSPLPEKLDSYLDGLAYLISGNVVWSLNCPRYHPEHRY DPNAGVEDDLTANSLARTLGTDYGFRGTLNAKDASEKD TSVHFTPESEHSRRSSTETLLTAPDDDGSSRDDNISTSS CILTSSSSEKKKQSMYVNLPSKGLLGSVRVQAPFDYIAS LPSKGVRDTFIDALNLWLGVSEPVASRIKSLGGRLFTAS LMIDDIEDGSNLRRGQPAAHTVFGTAQTINSGCYEILQA VNEAQQLGPGAVKIVLEELDELHIGQSYDLYWTHHIMCP SEDEYLEMVMKKTGALFRLLARLLLAHTRTKTLAAPIENL VILVGIQYQIRDDYQNLHSEEYAAQKGSCEDLDEGKFSF PLIHAMSTQSTSGGLLRELLLRRRDLGYLPDEHKKPVLE QFDRAGSMAYTRETLKRLQGEVYDRLEMIEDATKQEN WILRALLQRLELRHESADVPGKLGKVQADLRKSKVELQ VKEEKTRLQKRVARSITTTEENISKAMKDLAAREELAVA RREKKGARSEIX |

**Table S7.** Positions of the 38 previously reported BFTPSs within the predicted clusters.

| Cluster | Published BFTPS | Product | Classfication |
|---|---|---|---|
| 1 | PaFS | Fusicoccadiene | NSR-BFTPS |
| | AbFS | | |
| | PgFS | | |
| 2 | AcOS | Ophiobolin F | |
| | AcldOS | | |
| | BmOS | | |
| | AuOS | | |
| | Au8003 | | |
| 3 | NnNS | Nigtetraene | |
| | PfNS | | |
| 4 | TvTS | Talaropentaene | |
| 5 | CgDS | Dolasta-1(15),8-diene | |
| 7 | PrDS | Penichrysol | SR-BFTPS |
| | PcCS | | |
| | PchDS | | |
| | PhPS | | |
| | PaCS | | |
| 8 | PvPS | Preasperterpeniod A | |
| | AstC | | |
| | TtPS | | |
| | TvPS | | |
| 9 | FgMS | Mangicdiene | |
| | FoMS | | |
| 10 | AfAS | Asperfumene | |
| 11 | EvSS | Stellata-2,6,19-triene | |
| | EvAS | Astellifadiene | |
| | TndC | Talarodiene | |
| | TlnA | | |
| | PaPS | Phomopsene | |
| | FgGS | GJ1012E | |
| 12 | AcSS | Spiroviolene | |
| 13 | EvVS | Variediene | |
| | AbVS-1 | | |
| | DvVS | | |
| | SmVS | | |
| | BpVS | | |
| | AbVS-2 | | |
| | AaGS | | |

**Table S8.** Correspondence between TRACER-derived clusters and SSN components used for the alluvial diagram.

| TRACER-derived cluster | SSN component | Number of BFTPS | Proteins |
|---|---|---|---|
| Cluster 1 | Group 1 | 33 | AbFS; PaFS; PgFS; cd08086_g2062.t1; dq11167_g1997.t1; hf13050_g10624.t1; lf17061_g7219.t1; ni15096_g6987.t1; no11201_g1886.t1; oa01042_g7340.t1; pa17018_g11424.t1; pc17103_g4653.t1; pm13107_g8369.t1; sp17089_g3701.t1; sp17089_g5951.t1; vf16017_g5518.t1; xh12014_g1694.t1; xs14040_g4868.t1; EME87121; KIK55682; KIK55691; KIK55704; XP_014550177; KUL83692; KZL77275; OCL09522; XP_018153114; CZT49095; OLN96964; POS74342; PQE22675; RWA09438; TLD03666 |
| Cluster 2 | Group 1 | 20 | AcldOS; AcOS; Au8003; AuOS; BmOS; bb12017_g7594.t1; bs11134_a63_g4524.t1; he13146_g11738.t1; XP_003067066; EJP62241; EUC28897; XP_007694490; KIN08340; GAO85920; CEL04094; BAX09282; PGH21525; XP_026614492; RLL93374; XP_028488251 |
| Cluster 3 | Group 3 | 16 | NnNS; PfNS; am12567_g12443.t1; ck10162_g6973.t1; dq11167_g11127.t1; fa13326_g3585.t1; fc13317_g660.t1; fp13316_2; hf14026_g5787.t1; hf15128_g9055.t1; hr14028_g7684.t1; sb17049_g11090.t1; ETS85684; XP_018187504; OQD85661; RBR18644 |
| Cluster 4 | Group 1 | 2 | TvTS; KUL85185 |
| Cluster 5 | Group 8 | 1 | mf17066_g4236.t1 |
| | SSN singleton | 4 | CgDS; cl13394_g12660.t1; ht13129_g5464.t1; md13101_g1172.t1 |
| Cluster 6 | Group 7 | 3 | cg11192_g2732.t1; cp16081_g9942.t1; EAQ85668 |
| | Group 8 | 1 | ma17069_g9201.t1 |
| | SSN singleton | 1 | APA12276 |
| Cluster 7 | Group 4 | 15 | PcCS; PrDS; PchDS; PaCS; ac13149_a79_g1877.t1; pe13048_g11157.t1; CAP86415; CDM32996; CRL20985; KOS48568; KZN86668; XP_016596824; XP_022486475; BBF45518; RJE25610 |
| | Group 5 | 1 | PhPS |
| Cluster 8 | Group 1 | 5 | AstC; PvPS; TtPS; TvPS; AZQ56744 |
| Cluster 9 | Group 1 | 9 | FgMS; FoMS; 13316_g9877_t1; cs13151_g15279.t1; fe07089_g5865.t1; fm13297_g11606.t1; fs13303_g8348.t1; APY21859; XP_025589290 |
| Cluster 10 | Group 1 | 1 | AfAS |
| Cluster 11 | Group 1 | 12 | EvAS; EvSS; TlnA; TndC; bs14032_g55_g7540.t1; sp17089_g8741.t1; EED11493; PYI14929; XP_025395331; XP_025487035; XP_025525081; RMD43624 |
| | Group 2 | 1 | FgGS |
| | Group 5 | 2 | PaPS; EYE97880 |
| Cluster 12 | Group 6 | 6 | AcSS; GAQ04021; CEL11848; GAT28430; XP_024677291; XP_026631727 |
| Cluster 13 | Group 2 | 41 | AbVS-1; AaGS; EvVS; SmVS; BpVS; AbVS-2; at12575_g11485.t1; cf16051_g2411.t1; cm16055_g11090.t1; en09116_g4558.t1; fs16075_g8752.t1; pg13102_g5679.t1; pm11196_g3202.t1; XP_001266466; EFQ93661; XP_003343918; EUC38927; XP_007677283; KOC10771; XP_018390852; OJJ72250; OOF96881; OQE14697; OSS43380; XP_023093164; PQE19349; XP_024705680; PYI01416; PZD43626; XP_025426141; XP_025486510; XP_025501272; XP_025555922; XP_025558733; XP_025461812; XP_026604901; XP_026622213; GCB25423; RMZ70974; RYO55048; XP_028489435 |
| | Group 5 | 2 | DvVS; pb17001_g4775.t1 |
| | Group 9 | 3 | cl11194_g22753.t1; co17083_g6831.t1; cp16027_g195.t1 |
| | SSN singleton | 1 | aa12573_v89_g6264.t1 |
| Cluster 14 | Group 2 | 1 | sg17035_g19002.t1 |

| | | | |
|---|---|---|---|
| | Group 10 | 2 | dp12545_g13069.t1; EKG20455 |
| | SSN singleton | 3 | ac13035_a47_g7059.t1; EPE34883; CZR58890 |
| Singleton | Group 1 | 4 | cr15020_g13832.t1; RAQ98630; am17074_g4245.t1; OCK92653 |
| | Group 2 | 2 | RAR08378; EUC50502 |
| | Group 5 | 2 | ec15031_g7858.t1; pa17018_g9084.t1 |
| | Group 7 | 2 | cm13013_g2281.t1; SPQ20835 |
| | Group 8 | 2 | THV50721; be97436_g1673.t1 |
| | SSN singleton | 1 | fs13303_g2078.t1 |

**Table S9.** Detailed information of the 18 BFTPSs selected for machine learning validation

| No. | BFTPS name | Source | Predicted type | Selection rationale* | Enzyme activity | New compound | Actual type |
|---|---|---|---|---|---|---|---|
| 1 | FsPS | *Fusarium sporotrichioides* 13303 | SR | SRC | Yes | Yes | SR |
| 2 | AcPS | *Arthrobotrys conoides* 13035 | SR | Dark | Yes | Yes | SR |
| 3 | FpMS | *Fusarium pseudograminearum* 13316 | SR | PCC | Yes | No | SR |
| 4 | AtVS | *Alternaria tenuissima* 12575 | SR | PCC | Yes | No | SR |
| 5 | AlSS | *Aspergillus lentulus* | SR | PCC | Yes | No | SR |
| 6 | CrNS | *Clonostachys rosea* 15020 | NSR | SRC | Yes | Yes | NSR |
| 7 | BbOS | *Beauveria* sp. 12017 | NSR | PCC | Yes | No | NSR |
| 8 | OaFS | *Ophiosphaerella agrostis* 01042 | NSR | PCC | Yes | No | NSR |
| 9 | DqFS | *Discula quercina* 11167 | NSR | PCC | Yes | No | NSR |
| 10 | XP_023093164 | *Aspergillus oryzae* | SR | PCC | No | — | — |
| 11 | THV50721 | *Botrytis galanthina* | SR | SRC | No | — | — |
| 12 | co17083 | *Cladosporium oxysporum* 17083 | SR | PCC | No | — | — |
| 13 | cm13013 | *Chaetomium murorum* 13013 | SR | SRC | No | — | — |
| 14 | OCK92653 | *Cenococcum geophilum* | SR | SRC | No | — | — |
| 15 | fc13317 | *Fusarium culmorum* 13317 | NSR | PCC | No | — | — |
| 16 | he13146 | *Helicomyces* sp. 13146 | NSR | PCC | No | — | — |
| 17 | xs14040 | *Xylaria* sp. 14040 | NSR | PCC | No | — | — |
| 18 | cg11192 | *Chaetomium globosum* 11192 | NSR | Dark | No | — | — |

*Selection rationale: PCC, product-coherent clustering validation; SRC, singleton-reference-cluster-associated candidates; Dark, candidates from dark clusters without previously characterized representatives.

**Table S10.** Comparison of $^{1}H$ (600 MHz) and $^{13}C$ (150 MHz) NMR (in pyridine-$d_5$) data of **1** to those of fusicoccadiene (400 MHz, in $CD_2Cl_2$) in the literature.[50]

| Pos. | Experimental data of **1** | | Reported data of fusicoccadiene | |
|---|---|---|---|---|
| | $\delta_H$, mult (*J* in Hz) | $\delta_C$ | $\delta_H$, mult (*J* in Hz) | $\delta_C$ |
| 1 | 2.34, d (13.2); 1.97, d (13.8) | 39.9 | 2.31, ABq (13.0, 2.0); 1.95, ABq (13.0, 2.0) | 39.81 |
| 2 | | 137.8 | | 140.31 |
| 3 | | 132.7 | | 132.75 |
| 4 | 2.24, m; 2.15, m | 37.2 | 2.20, m | 37.26 |
| 5 | 1.70, m; 1.46, m | 23.0 | 1.67, m; 1.46, m | 23.11 |
| 6 | 2.52, m | 55.3 | 2.50, m | 55.33 |
| 7 | 2.00, m | 30.4 | 2.04, m | 30.31 |
| 8 | 1.48, m | 32.8 | 1.46, m | 32.89 |
| 9 | 2.10, m; 1.98, m | 22.0 | 2.04, m | 21.86 |
| 10 | | 140.3 | | 141.14 |
| 11 | | 52.2 | | 52.23 |
| 12 | 1.70, m | 39.1 | 1.67, m | 39.17 |
| 13 | 2.27, m; 2.22, m | 27.4 | 2.20, m | 27.27 |
| 14 | | 141.1 | | 137.75 |
| 15 | 2.64, m | 27.7 | 2.64, spt | 27.67 |
| 16 | 1.64, s | 16.0 | 1.64, brs | 15.83 |
| 17 | 0.90, d (6.6) | 22.0 | 0.87, d (7.0) | 21.54 |
| 18 | 1.00, s | 27.3 | 0.94, s | 27.03 |
| 19 | 0.96, d (6.6) | 21.4 | 0.93, d (6.0) | 21.32 |
| 20 | 1.01, d (8.4) | 21.8 | 0.99, d (8.0) | 21.59 |

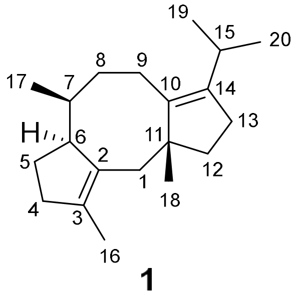

**1**

**Table S11.** Comparison of $^{1}H$ (600 MHz) and $^{13}C$ (150 MHz) NMR (in benzene-$d_6$) data of **2** to those of ophiobolin F (500 MHz, in benzene-$d_6$) in the literature.[7]

| Pos. | Experimental data of **2** | | Reported data of ophiobolin F | |
|---|---|---|---|---|
| | $\delta_H$, mult (*J* in Hz) | $\delta_C$ | $\delta_H$, mult (*J* in Hz) | $\delta_C$ |
| 1 | 1.94, m; 2.00, m | 26.0 | 1.94, m; 2.00, m | 25.70 |
| 2 | 1.61, m | 54.3 | 1.62, m | 54.19 |
| 3 | | 80.3 | | 80.20 |
| 4 | 1.96, m; 1.67, m | 42.0 | 1.96, m; 1.68, m | 41.90 |
| 5 | 1.54, m; 1.43, m | 24.0 | 1.54, m; 1.43, m | 23.83 |
| 6 | 3.15, t (8.3) | 42.6 | 3.15, t (8.3) | 42.52 |
| 7 | | 136.7 | | 136.62 |
| 8 | 5.64, t (8.2) | 129.2 | 5.64, t (8.3) | 129.15 |
| 9 | 2.13, m | 23.8 | 2.13, m | 23.70 |
| 10 | 1.68, m | 55.9 | 1.68, m | 55.85 |
| 11 | | 44.0 | | 43.92 |
| 12 | 1.44, m; 1.34, m | 43.7 | 1.44, m; 1.34, m | 43.66 |
| 13 | 1.44, m; 1.32, m | 36.9 | 1.44, m; 1.32, m | 36.81 |
| 14 | 2.31, m | 46.1 | 2.30, m | 45.97 |
| 15 | 1.68, m | 33.4 | 1.68, m | 33.37 |
| 16 | 1.40, m; 1.27, m | 37.9 | 1.40, m; 1.27, m | 37.77 |
| 17 | 1.95, m | 26.8 | 1.95, m | 26.73 |
| 18 | 5.28, t (7.1) | 125.6 | 5.28, t (6.9) | 125.56 |
| 19 | | 131.0 | | 130.93 |
| 20 | 1.05, s | 29.1 | 1.05, s | 29.00 |
| 21 | 2.02, s | 22.0 | 2.03, s | 21.95 |
| 22 | 0.86, s | 19.1 | 0.85, s | 19.06 |
| 23 | 0.84, d (6.8) | 17.3 | 0.83, d (6.9) | 17.18 |
| 24 | 1.62, s | 17.8 | 1.62, s | 17.76 |
| 25 | 1.71, s | 26.2 | 1.71, s | 26.10 |

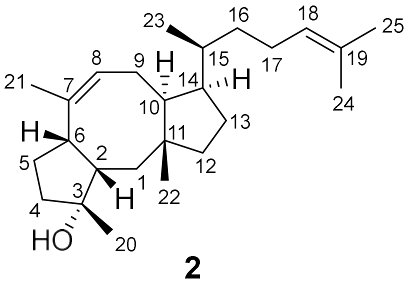

**2**

**Table S12.** Comparison of $^{1}H$ (600 MHz) and $^{13}C$ (150 MHz) NMR (in benzene-$d_6$) data of **8** to those of mangicdiene (400 MHz, in $CDCl_3$) in the literature.[8]

| Pos. | Experimental data of **8** | | Reported data of mangicdiene | |
|---|---|---|---|---|
| | $\delta_H$, mult ($J$ in Hz) | $\delta_C$ | $\delta_H$, mult ($J$ in Hz) | $\delta_C$ |
| 1 | 5.50, d (1.7) | 130.8 | 5.40, d (1.7) | 130.14 |
| 2 | | 144.8 | | 144.37 |
| 3 | 2.54, m | 37.9 | 2.49, m | 37.42 |
| 4 | 1.96, m; 1.25, m | 31.5 | 1.81, m; 1.18, m | 31.14 |
| 5 | 1.60, m; 1.25, m | 44.6 | 1.67, m; 1.17, m | 44.15 |
| 6 | | 57.5 | | 56.37 |
| 7 | 1.81, m | 42.2 | 1.75, td (7.6, 5.1) | 41.58 |
| 8 | 2.10, m; 1.46, m | 33.3 | 1.96, m; 1.36, m | 32.83 |
| 9 | 1.82, m; 1.46, m | 32.9 | 1.82, m; 1.46, m | 32.22 |
| 10 | 1.67, m | 47.2 | 1.59, m | 46.76 |
| 11 | | 44.8 | | 44.54 |
| 12 | 1.46, m; 1.39, m | 40.1 | 1.46, m; 1.37, m | 39.53 |
| 13 | 1.67, m; 1.39, m | 38.8 | 1.60, m; 1.32, m | 38.52 |
| 14 | 1.26, m | 56.7 | 1.27, d (9.4) | 56.71 |
| 15 | | 46.1 | | 45.69 |
| 16 | 1.55, m; 1.39, m | 46.5 | 1.49, m; 1.31, m | 46.24 |
| 17 | 2.15, m | 24.5 | 1.97, m | 23.96 |
| 18 | 5.28, t (7.0) | 126.0 | 5.12, t (7.0) | 125.34 |
| 19 | | 130.7 | | 130.89 |
| 20 | 1.17, d (7.0) | 21.8 | 1.07, d (7.1) | 21.57 |
| 21 | 0.94, d (7.0) | 22.2 | 0.82, d (7.3) | 21.96 |
| 22 | 1.19, s | 30.7 | 1.07, s | 30.23 |
| 23 | 0.92, s | 23.0 | 0.85, s | 22.7 |
| 24 | 1.62, s | 17.7 | 1.60, s | 17.58 |
| 25 | 1.72, s | 25.9 | 1.68, s | 25.75 |

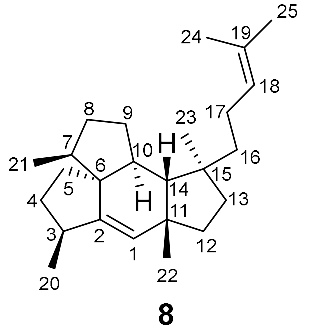

**8**

**Table S13.** Comparison of [1]H (600 MHz) and [13]C (150 MHz) NMR (in benzene-$d_6$) data of **15** to those of spiroviolene (700 MHz, in benzene-$d_6$) in the literature.[13]

| Pos. | Experimental data of **15** | | Reported data of spiroviolene | |
|---|---|---|---|---|
| | $\delta_H$, mult (*J* in Hz) | $\delta_C$ | $\delta_H$, mult (*J* in Hz) | $\delta_C$ |
| 1 | 4.81, d (2.9) | 128.9 | 4.81, d (2.9) | 128.9 |
| 2 | | 148.9 | | 148.9 |
| 3 | 1.60, m | 44.8 | 1.60, m | 44.7 |
| 4 | 1.79, m; 1.38, m | 31.3 | 1.79, m; 1.38, m | 31.3 |
| 5 | 1.74 m; 1.33, m | 30.7 | 1.74 m; 1.33, m | 30.7 |
| 6 | 1.81, m | 46.7 | 1.81, m | 46.6 |
| 7 | | 53.8 | | 53.8 |
| 8 | 1.92, m; 1.69, m | 39.6 | 1.92, ddd (12.7, 6.9, 6.9); 1.69, m | 39.5 |
| 9 | 1.72, m; 1.09, m | 33.1 | 1.72, m; 1.09, dddd (12.2, 12.2, 11.3, 7.6) | 33.1 |
| 10 | 2.77, dddd (12.5, 6.4, 6.4, 2.9) | 59.4 | 2.77, dddd (12.5, 6.4, 6.4, 2.9) | 59.4 |
| 11 | | 63.7 | | 63.7 |
| 12 | 1.73, m; 1.59, m | 38.6 | 1.73, m; 1.59, m | 38.6 |
| 13 | 1.67, m; 1.43, m | 40.8 | 1.67, m; 1.43, dddd (11.8, 6.6, 1.5, 1.5) | 40.8 |
| 14 | 1.58, m | 66.1 | 1.58, m | 66.0 |
| 15 | | 41.3 | | 41.3 |
| 16 | 0.94, d (6.7) | 15.1 | 0.94, d (6.7) | 15.1 |
| 17 | 0.97, d (6.7) | 15.2 | 0.97, d (6.7) | 15.2 |
| 18 | 1.34, s | 32.4 | 1.34, s | 32.4 |
| 19 | 1.04, s | 29.2 | 1.04, s | 29.1 |
| 20 | 1.03, s | 26.1 | 1.03, s | 26.1 |

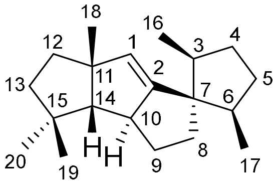

**15**

**Table S14.** Comparison of $^{1}H$ (600 MHz) and $^{13}C$ (150 MHz) NMR (in benzene-$d_6$) data of **16** to those of variediene (700 MHz, in benzene-$d_6$) in the literature.[16]

| Pos. | Experimental data of **16** | | Reported data of variediene | |
|---|---|---|---|---|
| | $\delta_H$, mult (*J* in Hz) | $\delta_C$ | $\delta_H$, mult (*J* in Hz) | $\delta_C$ |
| 1 | 2.19, m; 2.11, m | 45.1 | 2.18, m; 2.11, m | 45.1 |
| 2 | | 144.2 | | 144.2 |
| 3 | | 121.2 | | 121.2 |
| 4 | 1.91, m | 38.3 | 1.94, m; 1.90, m | 38.3 |
| 5 | 2.03, m; 1.89, m | 25.3 | 2.01, m; 1.90, m | 25.3 |
| 6 | 5.26, dd (11.0, 3.8) | 127.4 | 5.26, dd (11.0, 3.8) | 127.4 |
| 7 | | 138.4 | | 138.4 |
| 8 | 2.15, m; 2.06, m | 40.9 | 2.15, m; 2.07, m | 40.9 |
| 9 | 2.19, m; 1.57, m | 41.6 | 2.21, m; 1.55, m | 41.6 |
| 10 | 2.36, m | 45.0 | 2.36, d (10.6) | 44.9 |
| 11 | | 48.9 | | 48.9 |
| 12 | 1.55, m; 1.45, m | 42.0 | 1.54, m; 1.47, m | 42.0 |
| 13 | 1.35, m | 41.6 | 1.35, m | 41.6 |
| 14 | 1.35, m | 68.7 | 1.35, m | 68.7 |
| 15 | | 43.3 | | 43.2 |
| 16 | 1.57, d (1.3) | 21.5 | 1.57, d (1.3) | 21.5 |
| 17 | 1.39, s | 16.3 | 1.39, s | 16.3 |
| 18 | 1.26, s | 31.5 | 1.26, s | 31.5 |
| 19 | 1.03, s | 32.2 | 1.03, s | 32.2 |
| 20 | 0.90, s | 24.7 | 0.90, s | 24.7 |

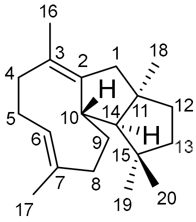

**16**

**Table S15.** 1D and 2D NMR data of **17** in benzene-$d_6$.

| Pos. | $\delta_H$[a], mult (*J* in Hz) | $\delta_C$[b], type | $^1H-^1H$ COSY | HMBC | NOESY |
|---|---|---|---|---|---|
| 1a | 2.24, m | 43.8, $CH_2$ | | | 20 |
| 1b | 1.75, m | | 2 | 2, 3, 10, 11, 22 | |
| 2 | 5.19, dd (12.0, 4.2) | 126.3, CH | 1b | 1, 4, 20 | |
| 3 | | 134.4, C | | | |
| 4a | 2.15, m | 40.2, CH | | 2, 3, 5, 20 | |
| 4b | 2.02, m | | 5a | 2, 3, 5, 20 | |
| 5a | 2.25, m | 24.8, $CH_2$ | 4b, 6 | | 21 |
| 5b | 2.01, m | | 6 | | 21 |
| 6 | 4.87, d (8.4) | 127.9, CH | 5a, 5b | 4, 5, 8, 21 | |
| 7 | | 133.8, C | | | |
| 8a | 2.22, m | 38.3, $CH_2$ | 9a, 9b | | |
| 8b | 2.00, m | | 9a, 9b | 6, 7, 9, 10, 21 | |
| 9a | 1.43, m | 24.6, $CH_2$ | 8a, 8b, 10 | 7, 8, 10, 11, 14 | |
| 9b | 1.22, m | | 8a, 8b, 10 | 7, 8, 10, 11, 14 | |
| 10 | 1.80, m | 42.1, CH | 9a, 9b, 14 | 1, 8, 9, 11, 12, 13, 14 | 2, 6, 12b |
| 11 | | 46.7, C | | | |
| 12a | 1.51, m | 42.5, $CH_2$ | | | |
| 12b | 1.40, m | | 13a | 1, 13, 22 | 10, 14 |
| 13a | 1.70, m | 28.6, $CH_2$ | 12b, 14 | | |
| 13b | 1.58, m | | 14 | | |
| 14 | 2.61, m | 52.8, CH | 10, 13a, 13b | 9, 10, 13, 15, 16, 23 | 10, 12b, 16 |
| 15 | | 136.0, C | | | |
| 16 | 5.31, m | 124.8, CH | 17 | 14, 17, 18, 23 | 14 |
| 17 | 2.85, t (6.6) | 27.6, $CH_2$ | 16, 18 | 15, 16, 18, 19 | 23 |
| 18 | 5.31, m | 124.1, CH | 17 | 17, 24, 25 | |
| 19 | | 131.1, C | | | |
| 20 | 1.50, s | 15.8, $CH_3$ | | 2, 3, 4 | 1a |
| 21 | 1.52, s | 16.6, $CH_3$ | | 6, 7, 8 | 5a, 5b, 22 |
| 22 | 1.06, s | 24.6, $CH_3$ | | 1, 10, 11, 12 | 12a, 21 |
| 23 | 1.67, s | 17.4, $CH_3$ | | 14, 15, 16 | 17 |
| 24 | 1.60, s | 17.8, $CH_3$ | | 18, 19, 25 | |
| 25 | 1.68, s | 25.8, $CH_3$ | | 18, 19, 24 | |

[a]Recorded at 600 MHz in Benzene-$d_6$. [b]Recorded at 150 MHz in Benzene-$d_6$.

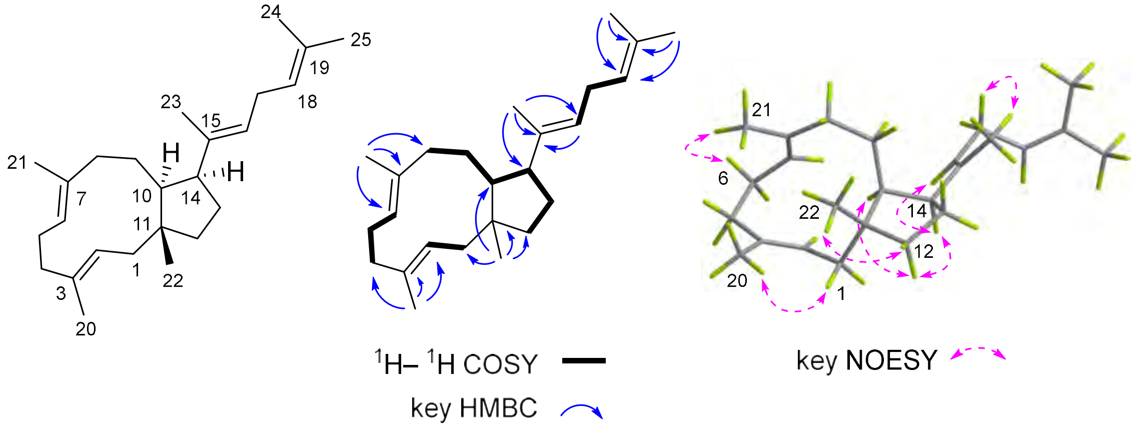

**Table S16.** 1D and 2D NMR data of **18** in benzene-$d_6$.

| Pos. | $\delta_H$[a], mult ($J$ in Hz) | $\delta_C$[b], type | $^1H-^1H$ COSY | HMBC | NOESY |
|---|---|---|---|---|---|
| 1a | 2.23, m | 41.3, $CH_2$ | | 2, 3, 10, 11 | 20, 22 |
| 1b | 1.84, m | | 2 | | 10 |
| 2 | 5.07, dd (9.0, 5.4) | 124.5, CH | 1b | | 4 |
| 3 | | 133.9, C | | | |
| 4 | 2.06, m | 39.9, $CH_2$ | 5 | 2, 3, 20 | 2 |
| 5 | 2.11, m | 24.9, $CH_2$ | 4, 6 | | 21 |
| 6 | 4.96, m | 125.7, CH | 5 | 8, 21 | |
| 7 | | 135.6, C | | | |
| 8a | 2.15, m | 37.5, $CH_2$ | 9b | | |
| 8b | 2.01, m | | 9a | | |
| 9a | 1.50, m | 31.6, $CH_2$ | 8b | 7, 14 | |
| 9b | 1.32, m | | 9a, 10 | | |
| 10 | 1.62, m | 45.8, CH | 9b, 14 | 1 | 1a |
| 11 | | 44.9, C | | | |
| 12a | 1.61, m | 42.2, $CH_2$ | | 13 | |
| 12b | 1.35, m | | 13b | | |
| 13a | 1.68, m | 30.2, $CH_2$ | 14 | | |
| 13b | 1.56, m | | 12a | | |
| 14 | 2.22, m | 57.9, CH | 13a | | 22 |
| 15 | | 152.9, C | | | |
| 16 | 2.17, m | 34.0, $CH_2$ | 17 | 15, 23 | |
| 17 | 2.30, m | 27.3, $CH_2$ | 16, 18 | 19 | |
| 18 | 5.30, m | 125.0, CH | 17 | | |
| 19 | | 131.4, C | | | |
| 20 | 1.51, s | 16.0, $CH_3$ | | 2, 3, 4 | 1a |
| 21 | 1.54, s | 17.6, $CH_3$ | | 6, 7, 8 | 5 |
| 22 | 0.94, s | 23.3, $CH_3$ | | 1, 10, 11, 12 | 1a, 14 |
| 23a | 5.02, m | 109.3, $CH_2$ | | 14, 16 | |
| 23b | 4.90, m | | | | |
| 24 | 1.59, s | 17.8, $CH_3$ | | 25, 18, 19 | |
| 25 | 1.68, s | 25.9, $CH_3$ | | 24, 18, 19 | |

[a]Recorded at 600 MHz in Benzene-$d_6$. [b]Recorded at 150 MHz in Benzene-$d_6$.

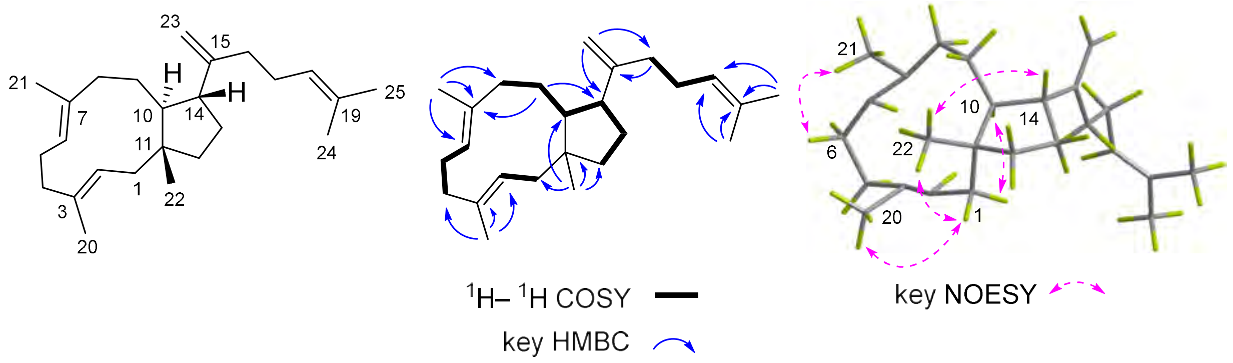

**Table S17**. Comparison of $^{1}H$ (600 MHz) and $^{13}C$ (150 MHz) NMR (in benzene-$d_6$) data of **19** to those of astelifatriene (600 MHz, in benzene-$d_6$) in the literature.[17]

| Pos. | Experimental data of **19** | | Reported data of astelifatriene | |
|---|---|---|---|---|
| | $\delta_H$, mult (*J* in Hz) | $\delta_C$ | $\delta_H$, mult (*J* in Hz) | $\delta_C$ |
| 1 | 2.16, m; 1.74, m | 39.1 | 2.15, m; 1.74, m | 39.1 |
| 2 | 5.04, d (8.2) | 125.0 | 5.04, d (8.2) | 125.0 |
| 3 | | 132.5 | | 132.5 |
| 4 | 2.16, m; 1.95, m | 39.7 | 2.15, m; 1.94, m | 39.7 |
| 5 | 2.26, m; 2.02, m | 26.3 | 2.26, qd (12.8, 4.4); 2.00, m | 26.3 |
| 6 | 4.93, dd (11.6, 4.3) | 124.6 | 4.93, dd (11.4, 3.7) | 124.5 |
| 7 | | 137.7 | | 137.7 |
| 8 | 2.00, m; 1.88, m | 42.0 | 2.00, m; 1.87, m | 42.0 |
| 9 | 1.22, m; 1.01, m | 24.3 | 1.21, m; 1.01, m | 24.3 |
| 10 | 1.63, m | 39.8 | 1.64, m | 39.8 |
| 11 | | 39.1 | | 39.1 |
| 12 | 1.99, m; 1.01, m | 34.8 | 2.00, m; 1.01, m | 34.7 |
| 13 | 1.51, m; 1.44, m | 36.3 | 1.51, m; 1.43, m | 36.3 |
| 14 | 1.58, m | 49.8 | 1.58, t (11.3) | 49.8 |
| 15 | | 43.9 | | 43.9 |
| 16 | 1.41, m; 1.19, m | 40.4 | 1.42, m; 1.21, m | 40.4 |
| 17 | 2.01, m; 1.45, m | 30.8 | 2.00, m; 1.46, m | 30.7 |
| 18 | 2.69, td (11.0, 5.3) | 47.5 | 2.69, td (10.9, 5.2) | 47.5 |
| 19 | | 148.6 | | 148.6 |
| 20 | 1.48, s | 16.7 | 1.47, s | 16.7 |
| 21 | 1.45, s | 16.9 | 1.45, s | 16.9 |
| 22 | 0.85, s | 21.2 | 0.85, s | 21.2 |
| 23 | 0.87, s | 18.0 | 0.87, s | 18.0 |
| 24 | 4.77, d (2.2); 4.70, dd (2.4, 1.3) | 110.5 | 4.77, d (2.4); 4.70, dd (2.5, 1.3) | 110.5 |
| 25 | 1.70, s | 19.1 | 1.70, s | 19.1 |

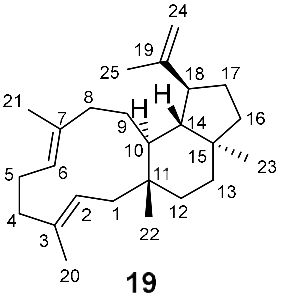

**19**

**Table S18.** 1D and 2D NMR data of **20** in benzene-$d_6$.

| Pos. | $\delta_H$[a], mult (*J* in Hz) | $\delta_C$[b], type | $^1H-^1H$ COSY | HMBC | NOESY |
|---|---|---|---|---|---|
| 1 | 5.09, dd (2.4, 2.4) | 132.1, CH | | | 3 |
| 2 | | 144.7, C | | | |
| 3 | 2.37, m | 40.6, CH | 4a, 4b, 20 | 1, 2 | 1, 5b |
| 4a | 1.64, m | 34.3, $CH_2$ | 3 | 2, 5, 6 | |
| 4b | 1.21, m | | 3, 5a | 2, 5, 6 | |
| 5a | 1.81, m | 26.9, $CH_2$ | 4b | | 21 |
| 5b | 1.61, m | | 6 | | 3 |
| 6 | 2.57, m | 46.1, CH | 5b | 21 | 8b, 10 |
| 7 | | 24.9, C | | | |
| 8a | 0.47, dd (8.0, 3.8 ) | 23.0, $CH_2$ | | 6, 9, 10 | 19, 21 |
| 8b | 0.36, m | | | | 6, 10 |
| 9 | 0.41, m | 26.4, CH | 10 | | 14, 19, 21, 22 |
| 10 | 1.76, m | 43.8, CH | 9, 14 | 8, 9, 11, 14, 22 | 6, 23 |
| 11 | | 42.4, C | | | |
| 12a | 1.81, m | 38.1, $CH_2$ | 13a, 13b | | 23 |
| 12b | 1.30, m | | | | 22 |
| 13a | 1.50, m | 35.9, $CH_2$ | 12a | | |
| 13b | 1.42, m | | 12a | | |
| 14 | 1.44, m | 49.5, CH | 10, 18 | 10, 15, 18, 19, 23 | 9 |
| 15 | | 43.5, C | | | |
| 16a | 1.44, m | 40.7, $CH_2$ | 17a | 14, 15, 18, 23 | |
| 16b | 1.10, m | | 17b | | |
| 17a | 1.68, m | 22.4, $CH_2$ | 16a | | |
| 17b | 1.53, m | | 16b, 17 | | |
| 18 | 1.87, m | 45.1, CH | 14, 17b, 19 | | 23 |
| 19 | 2.50, m | 28.7, CH | 18, 24, 25 | | 8a, 9 |
| 20 | 1.09, d (6.8) | 19.7, $CH_3$ | 3 | 2, 3, 4 | |
| 21 | 0.95, s | 20.7, $CH_3$ | | 6, 7, 8, 9 | 5a, 8a |
| 22 | 1.14, s | 20.7, $CH_3$ | | 1, 10, 11, 12 | 9, 12b |
| 23 | 0.82, s | 18.1, $CH_3$ | | 13, 14, 15, 16 | 10, 12a, 18 |
| 24 | 0.91, d (6.8) | 15.5, $CH_3$ | 19 | 18, 19, 25 | |
| 25 | 1.01, d (6.8) | 23.5, $CH_3$ | 19 | 18, 19, 25 | |

[a]Recorded at 600 MHz in Benzene-$d_6$. [b]Recorded at 150 MHz in Benzene-$d_6$.

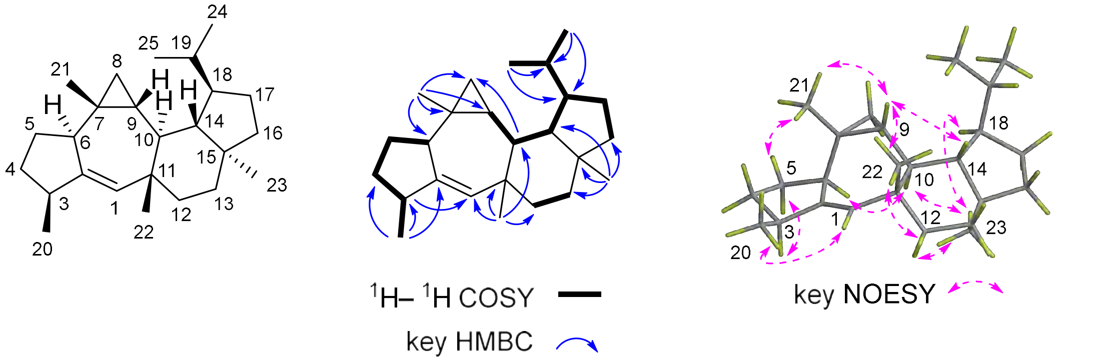

**Table S19.** 1D and 2D NMR data of **21** in benzene-$d_6$.

| Pos. | $\delta_H$[a], mult (*J* in Hz) | $\delta_C$[b], type | $^1H-^1H$ COSY | HMBC | NOESY |
|---|---|---|---|---|---|
| 1 | 5.21, s | 135.3, CH | | 2, 7, 10, 14, 22 | 12a, 21, 22 |
| 2 | | 147.0, C | | | |
| 3 | 2.01, m | 37.3, CH | 20, 4b | 4, 21 | 10 |
| 4a | 1.87, m | 30.2, $CH_2$ | | | |
| 4b | 1.28, m | | 3 | | |
| 5a | 1.93, m | 29.3, $CH_2$ | 6 | 4, 8 | |
| 5b | 1.26, m | | | | |
| 6 | 1.64, m | 48.7, CH | 5a, 8b | 4, 8, 21 | 21 |
| 7 | | 46.8, C | | | |
| 8a | 1.51, m | 31.6, $CH_2$ | 9a | 5 | |
| 8b | 1.19, m | | 6 | | |
| 9a | 1.96, m | 33.9, $CH_2$ | 8a, 10 | | |
| 9b | 0.95, m | | 10 | | |
| 10 | 2.31, m | 53.6, CH | 9a, 9b, 14 | 2 | 3, 18, 23, 25 |
| 11 | | 44.0, C | | | |
| 12a | 1.40, m | 33.1, $CH_2$ | | | |
| 12b | 1.35, m | | 13 | | |
| 13 | 1.45, m | 36.2, $CH_2$ | 12b | 11, 12 | |
| 14 | 1.41, m | 50.5, CH | 10, 18 | 15, 18 | 22, 24 |
| 15 | | 40.0, C | | | |
| 16a | 1.65, m | 41.4, $CH_2$ | 17 | 13, 23 | |
| 16b | 1.52, m | | | | |
| 17 | 1.45, m | 40.4, $CH_2$ | 16a | 15 | |
| 18 | 1.33, s | 59.0, CH | 14 | 10, 11, 13, 14, 15, 23, 24, 25 | 10 |
| 19 | | 43.6, C | | | |
| 20 | 0.87, d (6.9) | 13.8, $CH_3$ | 3 | 3, 4, 7 | 22 |
| 21 | 0.93, s | 18.8, $CH_3$ | | 2, 3, 6, 7 | 6, 22 |
| 22 | 1.21, s | 28.5, $CH_3$ | | 1, 11, 12, 14 | 14, 20, 21, 24 |
| 23 | 1.15, s | 33.1, $CH_3$ | | 13, 15, 16, 18 | 10, 18 |
| 24 | 0.90, s | 25.0, $CH_3$ | | 17, 18, 19, 25 | 14, 22 |
| 25 | 1.00, s | 30.6, $CH_3$ | | 17, 18, 19, 24 | 10 |

[a]Recorded at 600 MHz in Benzene-$d_6$. [b]Recorded at 150 MHz in Benzene-$d_6$.

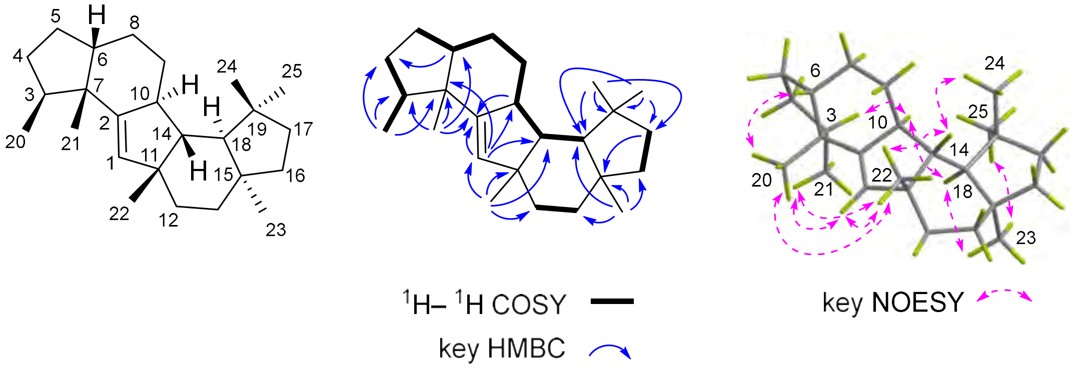

**Table S20.** 1D and 2D NMR data of **22** in benzene-$d_6$.

| Pos. | $\delta_H$[a], mult (*J* in Hz) | $\delta_C$[b], type | $^1$H – $^1$H COSY | HMBC | NOESY |
|---|---|---|---|---|---|
| 1a | 1.48, m | 33.5, $CH_2$ | | | |
| 1b | 1.19, m | | | | |
| 2 | | 139.2, C | | | |
| 3 | 1.92, m | 41.0, CH | 20, 4b | 2, 4, 6, 7, 20 | 12b |
| 4a | 1.76, m | 33.2, $CH_2$ | 5b | 5, 6, 7, 20 | |
| 4b | 1.19, m | | 3, 5b | | |
| 5a | 1.68, m | 27.1, $CH_2$ | 6 | 3, 4, 6, 7 | |
| 5b | 1.41, m | | 4a, 4b | 8 | |
| 6 | 1.83, m | 44.3, CH | 5a, 8 | 2, 8 | 14, 21 |
| 7 | | 44.5, C | | | |
| 8 | 1.64, m | 24.3, $CH_2$ | 6, 9a | 6, 7, 9, 10 | |
| 9a | 2.03, m | 21.7, $CH_2$ | 8 | | |
| 9b | 1.62, m | | | | |
| 10 | | 136.0, C | | | |
| 11 | | 37.7, C | | | |
| 12a | 2.22, m | 48.8, $CH_2$ | 13a | 1, 11, 15, 22 | 22 |
| 12b | 2.08, m | | | | 3, 23 |
| 13a | 1.44, m | 35.5, $CH_2$ | 12a | | |
| 13b | 1.30, m | | | 15, 16, 18, 22, 23 | |
| 14 | 2.21, s | 51.9, CH | 18 | 1, 2, 10, 11, 15, 18, 19, 22 | 6, 21, 22, 24 |
| 15 | | 39.5, C | | | |
| 16a | 1.61, m | 41.0, $CH_2$ | 17 | 13, 15, 23 | |
| 16b | 1.45, m | | | | |
| 17 | 1.45, m | 40.4, $CH_2$ | 16a | | |
| 18 | 1.45, m | 53.7, CH | 14 | | 23 |
| 19 | | 43.4, C | | | |
| 20 | 0.95, d (7.2) | 17.0, $CH_3$ | 3 | 3, 4, 7 | |
| 21 | 0.99, s | 21.2, $CH_3$ | | 2, 3, 6, 7 | 6, 14 |
| 22 | 1.18, s | 29.2, $CH_3$ | | 1, 11, 12, 14 | 12a, 14, 24 |
| 23 | 1.02, s | 32.3, $CH_3$ | | 13, 15, 16, 18 | 12b, 18 |
| 24 | 0.91, s | 24.7, $CH_3$ | | 17, 18, 19, 25 | 14, 22 |
| 25 | 1.00, s | 30.9, $CH_3$ | | 17, 18, 19, 24 | 18 |

[a]Recorded at 600 MHz in Benzene-$d_6$. [b]Recorded at 150 MHz in Benzene-$d_6$.

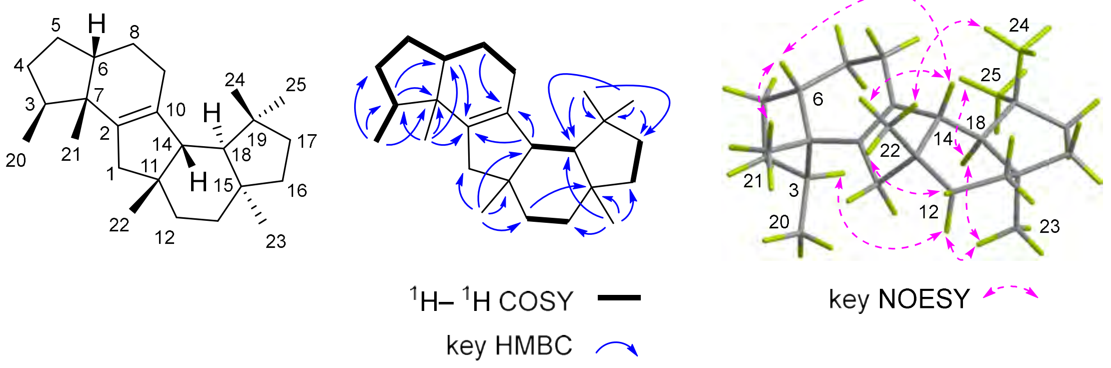

**Table S21.** 1D and 2D NMR data of **23** in benzene-$d_6$.

| Pos. | $\delta_H$[a], mult (*J* in Hz) | $\delta_C$[b], type | $^1H-^1H$ COSY | HMBC | NOESY |
|---|---|---|---|---|---|
| 1 | 1.48, m | 48.1, $CH_2$ | 2 | 2, 7, 10, 11, 12, 14, 22 | |
| 2 | 2.53, m | 48.9, CH | 1, 10 | 1, 6, 7, 8, 9 | 6 |
| 3 | | 142.8, C | | | |
| 4 | 5.44, m | 124.8, CH | 5b | | 20 |
| 5a | 2.64, m | 37.5, $CH_2$ | 6 | | |
| 5b | 1.67, m | | 4 | | |
| 6 | 1.81, m | 43.3, CH | 5a, 21 | 2, 3, 4, 5, 21 | 2, 10 |
| 7 | | 62.4, C | | | |
| 8a | 1.71, m | 26.1, $CH_2$ | | 6, 7, 9 | 21 |
| 8b | 1.50, m | | 9b | | 21, 22 |
| 9a | 1.65, m | 31.5, $CH_2$ | 10 | | 5a, 21 |
| 9b | 1.14, m | | 8b | | |
| 10 | 2.37, m | 55.1, CH | 2, 9a, 14 | | 6, 18, 23 |
| 11 | | 40.8, C | | | |
| 12a | 1.58, m | 29.6, $CH_2$ | | 13 | |
| 12b | 0.94, m | | 13a, 13b | | |
| 13a | 1.47, m | 36.2, $CH_2$ | 12b | 11, 12, 22 | |
| 13b | 1.41, m | | 12b | | |
| 14 | 1.17, m | 52.6, CH | 10, 18 | 9, 10, 12 | 22, 24 |
| 15 | | 39.8, C | | | |
| 16 | 1.59, m | 42.5, $CH_2$ | 17 | 13, 19, 23 | |
| 17 | 1.45, m | 40.5, $CH_2$ | 16 | 15, 18, 19, 24 | |
| 18 | 1.41, m | 59.8, CH | 14 | | 10, 23, 25 |
| 19 | | 43.3, C | | | |
| 20 | 1.76, dt (2.8, 1.2) | 16.5, $CH_3$ | 4, 5a | 3, 4, 7 | 1, 4 |
| 21 | 0.86, d (7.2) | 17.4, $CH_3$ | 6 | 5, 6, 7 | 8a, 8b |
| 22 | 1.00, s | 27.7, $CH_3$ | | 1, 11, 12, 14 | 8b, 14 |
| 23 | 1.18, s | 32.7, $CH_3$ | | 13, 15, 16, 18 | 10 |
| 24 | 0.90, s | 25.3, $CH_3$ | | 17, 18, 19, 25 | 14 |
| 25 | 1.05, s | 31.0, $CH_3$ | | 17, 18, 19, 24 | 18, 23 |

[a]Recorded at 600 MHz in Benzene-$d_6$. [b]Recorded at 150 MHz in Benzene-$d_6$.

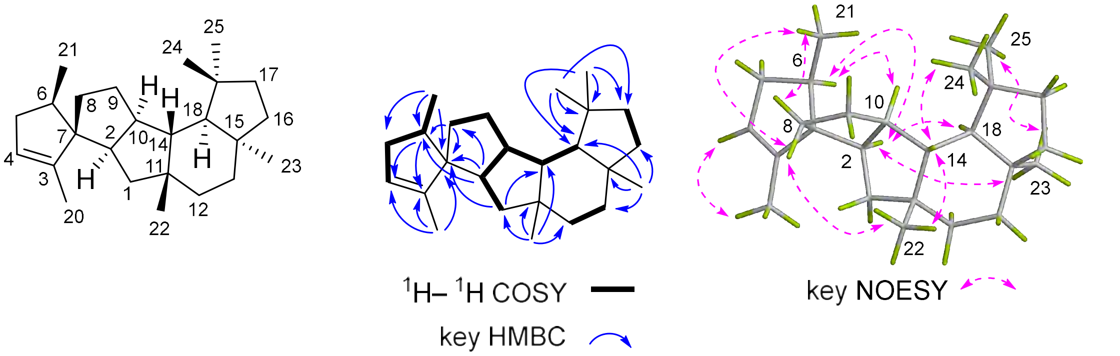

**Table S22.** 1D and 2D NMR data of **24** in benzene-$d_6$.

| Pos. | $\delta_H$[a], mult ($J$ in Hz) | $\delta_C$[b], type | $^1H-^1H$ COSY | HMBC | NOESY |
|---|---|---|---|---|---|
| 1a | 1.71, t (12.9) | 47.0, $CH_2$ | | 2, 3, 11, 12, 22 | |
| 1b | 1.39, m | | 2 | 2, 11, 12, 22 | |
| 2 | 2.48, m | 50.7, CH | 1b, 10 | 3 | 18 |
| 3 | | 149.7, C | | | |
| 4a | 2.45, m | 39.1, $CH_2$ | 5b | 3 | |
| 4b | 1.61, m | | 5b | | |
| 5a | 2.39, m | 28.8, $CH_2$ | 6 | 3, 4, 6, 7 | 21 |
| 5b | 1.95, m | | 4a, 4b, 6 | | |
| 6 | 5.38, dd (12.0, 3.6) | 124.0, CH | 5a, 5b | 5, 8, 21 | |
| 7 | | 135.9, C | | | |
| 8 | 2.06, m | 42.1, $CH_2$ | 9a, 9b | 6, 7, 9, 10, 21 | |
| 9a | 1.60, m | 32.3, $CH_2$ | 8 | | |
| 9b | 1.37, m | | 8 | | |
| 10 | 1.91, m | 52.3, CH | 2, 14 | 1, 2, 8, 9, 11, 14, 18 | 18, 23 |
| 11 | | 38.0, C | | | |
| 12a | 1.33, m | 34.8, $CH_2$ | 13b | 11, 13, 14, 15 | |
| 12b | 1.17, m | | | 11, 13, 14, 15 | |
| 13a | 1.33, m | 36.0, $CH_2$ | 12b | | |
| 13b | 1.28, m | | | | |
| 14 | 1.38, m | 56.8, CH | 10, 18 | | 22, 24 |
| 15 | | 42.3, C | | | |
| 16a | 1.47, m | 41.0, $CH_2$ | | 13, 18, 23 | |
| 16b | 1.41, m | | 17a | 13, 18, 23 | |
| 17a | 1.59, m | 41.5, $CH_2$ | 16b | | |
| 17b | 1.40, m | | | | |
| 18 | 1.13, d (9.0) | 59.7, CH | 14 | 10, 14, 23, 24, 25 | 2, 10 |
| 19 | | 43.2, C | | | |
| 20a | 4.87, s | 112.8, $CH_2$ | | 2, 4 | |
| 20b | 4.83, s | | | | |
| 21 | 1.58, s | 19.1, $CH_3$ | | 6, 7, 8 | 5a |
| 22 | 1.10, s | 31.1, $CH_3$ | | 1, 11, 12, 14 | 14 |
| 23 | 1.06, s | 31.0, $CH_3$ | | 13, 15, 16, 18 | 10, 25 |
| 24 | 0.97, s | 26.2, $CH_3$ | | 17, 18, 19, 25 | 14 |
| 25 | 1.22, s | 33.6, $CH_3$ | | 17, 18, 19, 24 | 23 |

[a]Recorded at 600 MHz in Benzene-$d_6$. [b]Recorded at 150 MHz in Benzene-$d_6$.

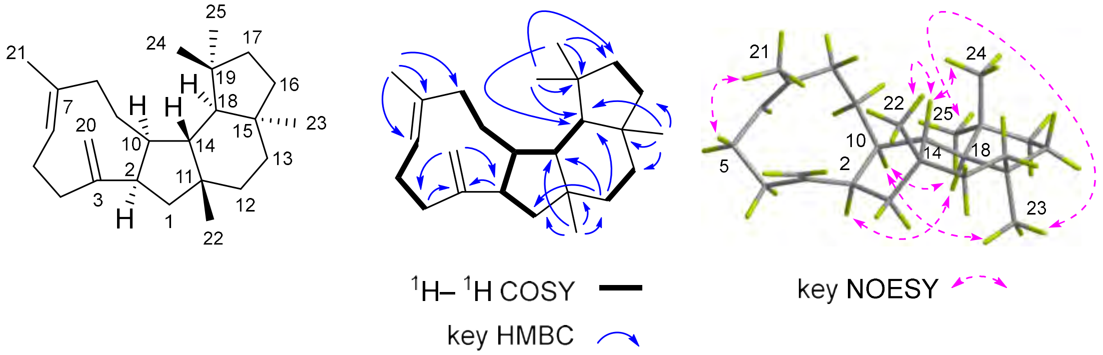

**Table S23.** 1D and 2D NMR data of **25** in benzene-$d_6$.

| Pos. | $\delta_H$[a], mult (*J* in Hz) | $\delta_C$[b], type | $^1H-^1H$ COSY | HMBC | NOESY |
|---|---|---|---|---|---|
| 1a | 2.11, d (14.4) | 42.5, $CH_2$ | | 2, 3, 6, 11, 12, 14 | 12b, 22 |
| 1b | 1.94, m | | | | |
| 2 | | 136.7, C | | | |
| 3 | | 131.2, C | | | |
| 4a | 2.39, m | 36.2, $CH_2$ | 5a, 5b | 2, 3 | 7 |
| 4b | 2.01, m | | 5b | 2, 3 | |
| 5a | 1.89, m | 47.4, $CH_2$ | 4a | 2, 3, 7 | 10 |
| 5b | 1.70, m | | 4a, 4b | 2, 3, 7 | |
| 6 | | 59.1, C | | | |
| 7 | 2.04, m | 44.1, CH | 21, 8a, 8b | 5, 6, 9 | 4a |
| 8a | 1.90, m | 34.0, $CH_2$ | 7, 9b | | |
| 8b | 1.40, m | | 7, 9b | | |
| 9a | 1.96, m | 32.4, $CH_2$ | 10 | 14 | |
| 9b | 1.57, m | | 8a, 8b, 10 | | |
| 10 | 1.79, m | 56.6, CH | 9a, 9b, 14 | | 5a, 23 |
| 11 | | 34.1, C | | | |
| 12a | 1.58, m | 29.6, $CH_2$ | | | |
| 12b | 0.66, m | | 13b | | |
| 13a | 1.57, m | 34.4, $CH_2$ | | | |
| 13b | 1.22, m | | 12b | | |
| 14 | 1.33, m | 46.5, CH | 10 | 1, 9, 10, 11, 12, 18 | 21, 22, 24 |
| 15 | | 40.7, C | | | |
| 16a | 1.64, m | 42.2, $CH_2$ | 17b | | |
| 16b | 1.38, m | | | | |
| 17a | 1.64, m | 39.9, $CH_2$ | 16b | 15, 18, 19 | |
| 17b | 1.33, m | | | | |
| 18 | 1.42, m | 60.5, CH | | 10, 14, 15, 19 | 23, 25 |
| 19 | | 42.9, C | | | |
| 20 | 1.70, s | 14.0, $CH_3$ | | 2, 3, 4 | 1a |
| 21 | 0.98, d (7.2) | 20.0, $CH_3$ | 7 | 6, 7, 8 | 14 |
| 22 | 1.00, s | 31.3, $CH_3$ | | 1, 11, 12, 14 | 14 |
| 23 | 1.18, s | 34.1, $CH_3$ | | 13, 15, 16 ,18 | 10, 18 |
| 24 | 1.04, s | 27.1, $CH_3$ | | 17, 18, 19, 25 | 14 |
| 25 | 1.07, s | 34.4, $CH_3$ | | 17, 18, 19, 24 | 18 |

[a]Recorded at 600 MHz in Benzene-$d_6$. [b]Recorded at 150 MHz in Benzene-$d_6$.

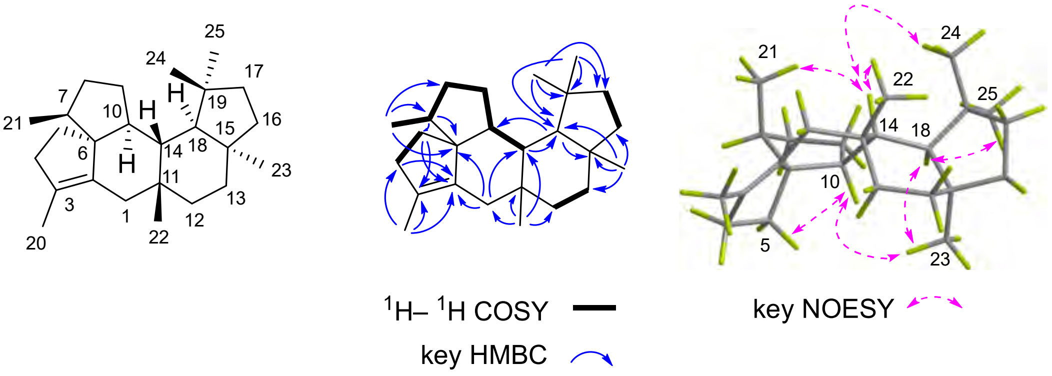

**Table S24.** 1D and 2D NMR data of **26** in benzene-$d_6$.

| Pos. | $\delta_H$[a], mult (*J* in Hz) | $\delta_C$[b], type | $^1H-{}^1H$ COSY | HMBC | NOESY |
|---|---|---|---|---|---|
| 1 | 2.13, m | 48.4, $CH_2$ | | | |
| 2 | | 141.8, C | | | |
| 3 | 1.74, m | 45.6, CH | 4, 20 | 2, 4, 7 | |
| 4 | 1.85, m | 31.7, $CH_2$ | 3, 5 | 7 | |
| 5 | 1.36, m | 31.9, $CH_2$ | 4, 6 | | |
| 6 | 1.74, m | 46.4, CH | 5, 21 | 2, 4, 7 | |
| 7 | | 60.8, C | | | |
| 8a | 2.08, m | 37.7, $CH_2$ | 9a | 2, 6, 7, 9, 10 | 20, 22 |
| 8b | 2.03, m | | | 2, 7, 9, 10 | 21 |
| 9a | 2.25, m | 28.0, $CH_2$ | 8a | 2 | |
| 9b | 2.03, m | | | 14 | |
| 10 | | 152.8, C | | | |
| 11 | | 45.2, C | | | |
| 12a | 1.56, m | 34.1, $CH_2$ | 13a, 13b | 1, 11, 15, 13, 22 | |
| 12b | 1.28, m | | 13a | 11, 13, 15 | |
| 13a | 1.41, m | 34.9, $CH_2$ | 12a, 12b | 18, 23 | |
| 13b | 1.36, m | | 12a | 11, 12, 15, 16, 18, 23 | |
| 14 | 2.21, m | 46.1, CH | 18 | 2, 10, 15, 18, 19 | 21, 22, 24 |
| 15 | | 39.6, C | | | |
| 16a | 1.62, m | 40.8, $CH_2$ | 17a | 13, 15, 23 | |
| 16b | 1.41, m | | | 15, 19, 23 | |
| 17a | 1.48, m | 40.4, $CH_2$ | 16a | 16, 19, 24, 25 | |
| 17b | 1.41, m | | | 15, 19, 24 | |
| 18 | 1.44, m | 55.4, CH | 14 | 10, 11, 14, 13, 23 | 23, 25 |
| 19 | | 43.7, C | | | |
| 20 | 1.00, d (6.6) | 15.3, $CH_3$ | 3 | 3, 4, 7 | 8a, 9b, 22 |
| 21 | 0.95, d (7.2) | 15.9, $CH_3$ | 6 | 5, 6, 7 | 14, 8b |
| 22 | 1.20, s | 30.3, $CH_3$ | | 1, 11, 12, 14 | 8a, 14, 20, 24 |
| 23 | 1.08, s | 32.8, $CH_3$ | | 13, 15, 16, 18 | 18 |
| 24 | 0.90, s | 24.5, $CH_3$ | | 17, 18, 19, 25 | 14, 22 |
| 25 | 1.00, s | 31.0, $CH_3$ | | 17, 18, 19, 24 | 18 |

[a]Recorded at 600 MHz in Benzene-$d_6$. [b]Recorded at 150 MHz in Benzene-$d_6$.

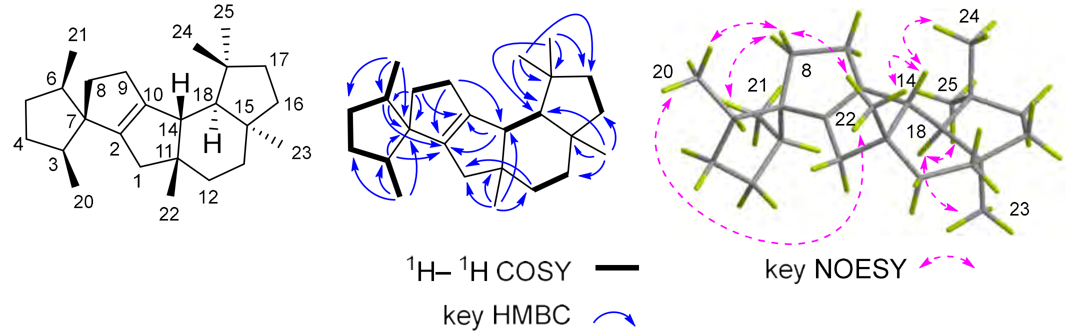

**Table S25.** 1D and 2D NMR data of **27** in benzene-$d_6$.

| Pos. | $\delta_H$[a], mult (*J* in Hz) | $\delta_C$[b], type | $^1H - ^1H$ COSY | HMBC | NOESY |
|---|---|---|---|---|---|
| 1 | 5.17, s | 129.0, CH | | 2, 4, 11, 14, 22 | 20, 22 |
| 2 | 1.66, m | 52.8, CH | 10 | 3, 5, 7, 8 | 10 |
| 3 | | 137.2, C | | | |
| 4 | 1.98, m | 36.5, CH | 20, 5b | 5 | |
| 5a | 1.58, m | 33.4, $CH_2$ | | | |
| 5b | 1.31, m | | 4 | 6 | |
| 6a | 1.63, m | 40.8, $CH_2$ | | | |
| 6b | 1.31, m | | | | |
| 7 | | 43.3, C | | | |
| 8a | 1.59, m | 40.3, $CH_2$ | | | |
| 8b | 1.20, m | | 9a | | |
| 9a | 2.07, m | 31.4, $CH_2$ | 8b | 8 | |
| 9b | 1.50, m | | | | |
| 10 | 2.12, m | 40.1, CH | 2, 9b, 14 | 7 | 2, 18, 23 |
| 11 | | 35.1, C | | | |
| 12a | 1.55, m | 32.0, $CH_2$ | | | |
| 12b | 1.15, m | | | | |
| 13a | 1.54, m | 34.0, $CH_2$ | | | |
| 13b | 1.26, m | | | 12, 15 | |
| 14 | 1.34, m | 43.5, CH | 10, 18 | | 21, 22, 24 |
| 15 | | 40.6, C | | | |
| 16a | 1.65, m | 42.0, $CH_2$ | 17b | | |
| 16b | 1.43, m | | 17a | | |
| 17a | 1.59, m | 40.2, $CH_2$ | 16b | | |
| 17b | 1.36, m | | 16a | | |
| 18 | 1.41, m | 58.2, CH | 14 | 11, 17, 19, 24, 25 | 10, 23 |
| 19 | | 43.0, C | | | |
| 20 | 1.11, d (6.0) | 17.2, $CH_3$ | 4 | 3, 4, 5 | 22 |
| 21 | 0.79, s | 20.8, $CH_3$ | | 2, 6, 7, 8 | 14 |
| 22 | 1.08, s | 29.8, $CH_3$ | | 1, 11, 12, 14 | 14, 20 |
| 23 | 1.21, s | 33.2, $CH_3$ | | 13, 15, 16, 18 | 10, 18, 25 |
| 24 | 1.01, s | 27.1, $CH_3$ | | 17, 18, 19, 25 | 14 |
| 25 | 1.07, s | 33.6, $CH_3$ | | 17,18, 19, 24 | 18 |

[a]Recorded at 600 MHz in Benzene-$d_6$. [b]Recorded at 150 MHz in Benzene-$d_6$.

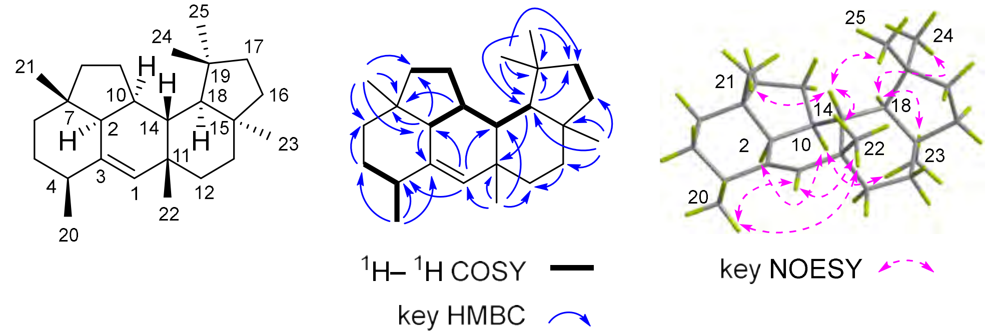

**Table S26.** 1D and 2D NMR data of **28** in benzene-$d_6$.

| Pos. | $\delta_H$[a], mult (*J* in Hz) | $\delta_C$[b], type | $^1H - ^1H$ COSY | HMBC | NOESY |
|---|---|---|---|---|---|
| 1a | 2.09, m | 41.1, $CH_2$ | | 2, 11, 14, 22 | |
| 1b | 1.51, m | | | 2, 3, 14 | 20, 21, 22 |
| 2 | | 145.0, C | | | |
| 3 | | 130.6, C | | | |
| 4 | 2.10, m | 32.1, CH | 5, 20 | 3 | |
| 5 | 1.34, m | 30.3, $CH_2$ | 4, 6a | | |
| 6a | 1.67, m | 42.0, $CH_2$ | 5 | 2, 21 | |
| 6b | 1.29, m | | | 8, 21 | |
| 7 | | 40.2, C | | | |
| 8a | 1.72, m | 37.4, $CH_2$ | | | |
| 8b | 1.33, m | | | | |
| 9 | 1.89, m | 30.3, $CH_2$ | 8a, 8b, 10 | 2, 7, 10 | |
| 10 | 2.00, m | 49.3, CH | 9, 14 | | 4, 23 |
| 11 | | 36.3, C | | | |
| 12a | 1.49, m | 35.0, $CH_2$ | 13a | | |
| 12b | 1.25, m | | | 1, 11, 14, 15 | |
| 13a | 1.69, m | 40.2, $CH_2$ | 12a | 15, 16, 23 | |
| 13b | 1.30, m | | | 18, 23 | |
| 14 | 1.02, m | 46.5, CH | 10, 18 | 10, 22 | 24 |
| 15 | | 39.7, C | | | |
| 16a | 1.49, m | 34.1, $CH_2$ | | | |
| 16b | 1.39, m | | 17a | 15, 18 | |
| 17a | 1.54, m | 40.6, $CH_2$ | 16b | | |
| 17b | 1.39, m | | | 15, 18, 24 | |
| 18 | 1.4 8, m | 59.0, CH | 14 | 10, 11, 14, 15, 19, 25 | 23, 25 |
| 19 | | 43.8, C | | | |
| 20 | 1.09, d (6.0) | 20.7, $CH_3$ | 4 | 3, 4, 5 | 1b |
| 21 | 1.03, s | 26.2, $CH_3$ | | 2, 6, 7, 8 | 1b |
| 22 | 1.01, s | 32.4, $CH_3$ | | 1, 11, 12, 14 | 1b, 24 |
| 23 | 1.10, s | 35.0, $CH_3$ | | 13, 15, 16, 18 | 10, 18 |
| 24 | 0.91, s | 25.7, $CH_3$ | | 17, 18, 19, 25 | 14, 22 |
| 25 | 1.04, s | 31.8, $CH_3$ | | 17, 18, 19, 24 | 18 |

[a]Recorded at 600 MHz in Benzene-$d_6$. [b]Recorded at 150 MHz in Benzene-$d_6$.

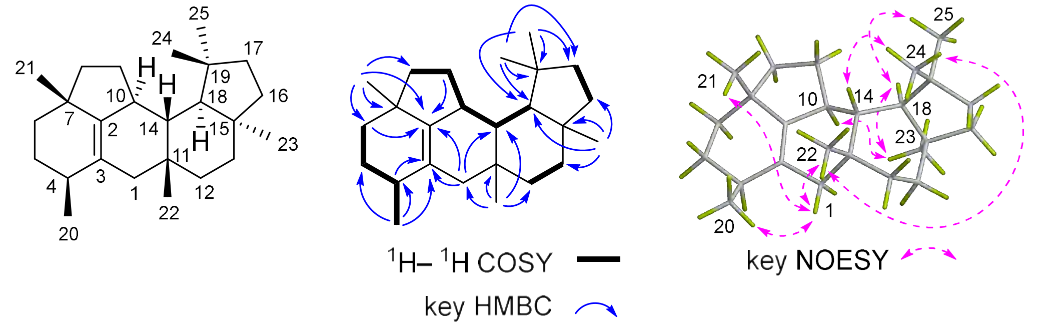

**Table S27.** X-ray crystallographic data for **19**, **22**, **25**, and **27**.

| | **19** | **22** | **25** | **27** |
|---|---|---|---|---|
| Radiation | Cu *Ka* | Cu *Ka* | Cu *Ka* | Cu *Ka* |
| Empirical formula | $C_{25}H_{40}$ | $C_{25}H_{40}$ | $C_{25}H_{40}$ | $C_{25}H_{40}$ |
| Formula weight | 340.57 | 340.57 | 340.57 | 340.57 |
| Temperature/K | 100.0 | 273.15 | 200.00 | 179.0 |
| Crystal system | monoclinic | monoclinic | orthorhombic | monoclinic |
| Space group | $P\ 2_1$ | $P\ 2_1$ | $P\ 2_1\ 2_12_1$ | $P\ 2_1$ |
| *a*/Å | 10.9841(5) | 7.7247(4) | 9.9951(9) | 11.5162(8) |
| *b*/Å | 16.7457(6) | 12.5514(7) | 14.3344(13) | 12.6635(8) |
| *c*/Å | 11.8142(4) | 11.2803(6) | 14.4821(13) | 14.8582(10) |
| *α*/deg | 90 | 90 | 90 | 90 |
| *β*/deg | 95.016(3) | 101.988(2) | 90 | 101.315(4) |
| *γ*/deg | 90 | 90 | 90 | 90 |
| Volume/$Å^3$ | 2164.74(15) | 1069.84(10) | 2074.9(3) | 2124.7(2) |
| *Z* | 4 | 2 | 4 | 4 |
| $\mu/mm^{-1}$ | 0.420 | 0.425 | 0.438 | 0.428 |
| F (000) | 760.0 | 380.0 | 760.0 | 760.0 |
| $\rho_{calc}$ g/$cm^3$ | 1.045 | 1.057 | 1.090 | 1.065 |
| Independent reflections | 9051[$R_{int}$ = 0.0686, $R_{sigma}$ = 0.0209] | 3606[$R_{int}$ = 0.0563, $R_{sigma}$ = 0.0395] | 3740 [$R_{int}$ = 0.0495, $R_{sigma}$ = 0.0326] | 7423 [$R_{int}$ = 0.0769, $R_{sigma}$ = 0.0598] |
| Goodness-of-fit on $F^2$ | 1.062 | 1.063 | 1.053 | 1.291 |
| Final R indexes [I>=2σ (I)] | $R_1$ = 0.0316, $wR_2$ = 0.0759 | $R_1$ = 0.0534, $wR_2$ = 0.1474 | $R_1$ = 0.0405, $wR_2$ = 0.1067 | $R_1$ = 0.1056, $wR_2$ = 0.2872 |
| Final R indexes [all data] | $R_1$ = 0.0343, $wR_2$ = 0.0776 | $R_1$ = 0.0544, $wR_2$ = 0.1485 | $R_1$ = 0.0415, $wR_2$ = 0.1079 | $R_1$ = 0.1093, $wR_2$ = 0.2939 |
| Flack parameter | 0.1(2) | 0.1(4) | -0.1(4) | 0.1(5) |

**Table S28**. Morgan fingerprint-based Tanimoto similarity analysis of new skeleton-containing compounds.

| Query Cpd | Max Tanimoto T | Nearest reference id | nearest_reference_smiles | novelty_class |
|---|---|---|---|---|
| **23** | 0.360656 | TKC207774 | C[C@H]1CC[C@@H]2[C@@H](O)C[C@@H]3C(=C[C@@]4(C)CC[C@]5(C)CCC(C)(C)[C@@H]5[C@@H]34)[C@]12C | Low-similarity / highly novel |
| **24** | 0.461538 | TKC045889 | C/C1=C/CC2(C)CCC3(C)CCC(C)(C)C(O)C3C2CC/C(C)=C\CC1 | Moderately low similarity |
| **25** | 0.372881 | TKC207774 | C[C@H]1CC[C@@H]2[C@@H](O)C[C@@H]3C(=C[C@@]4(C)CC[C@]5(C)CCC(C)(C)[C@@H]5[C@@H]34)[C@]12C | Low-similarity / highly novel |
| **26** | 0.305085 | TKC207774 | C[C@H]1CC[C@@H]2[C@@H](O)C[C@@H]3C(=C[C@@]4(C)CC[C@]5(C)CCC(C)(C)[C@@H]5[C@@H]34)[C@]12C | Low-similarity / highly novel |
| **27** | 0.403509 | TKC207774 | C[C@H]1CC[C@@H]2[C@@H](O)C[C@@H]3C(=C[C@@]4(C)CC[C@]5(C)CCC(C)(C)[C@@H]5[C@@H]34)[C@]12C | Moderately low similarity |
| **28** | 0.322034 | TKC207774 | C[C@H]1CC[C@@H]2[C@@H](O)C[C@@H]3C(=C[C@@]4(C)CC[C@]5(C)CCC(C)(C)[C@@H]5[C@@H]34)[C@]12C | Low-similarity / highly novel |

**Table S29.** 1D and 2D NMR data of **29** in benzene-$d_6$.

| Pos. | $\delta_H$[a], mult (*J* in Hz) | $\delta_C$[b], type | $^1H-^1H$ COSY | HMBC | NOESY |
|---|---|---|---|---|---|
| 1a | 2.34, m | 39.9, $CH_2$ | 2 | 2, 3, 10, 11, 22 | 10, 20 |
| 1b | 1.67, m | | | 10, 11 | 22 |
| 2 | 5.16, m | 125.4, CH | 1a | 4, 20 | |
| 3 | | 133.3, C | | | |
| 4 | 2.10, m | 40.2, CH | 5a | 2, 3, 5, 21 | |
| 5a | 2.21, m | 25.2, $CH_2$ | 4, 6 | | 21 |
| 5b | 2.07, m | | 6 | | 21 |
| 6 | 4.98, m | 127.8, CH | 5a, 5b | 8, 21 | |
| 7 | | 134.6, C | | | |
| 8 | 2.29, m | 39.2, $CH_2$ | 9a, 9b | 6, 7, 9, 10, 21 | 22 |
| 9a | 1.56, m | 32.6, $CH_2$ | 8, 10 | 7, 8, 11 | 23 |
| 9b | 1.52, m | | 8, 10 | | |
| 10 | 1.71, m | 41.0, CH | 9a, 9b, 14 | 1, 11, 14, 15 | 1a |
| 11 | | 47.1, C | | | |
| 12a | 1.45, m | 42.3, $CH_2$ | | | |
| 12b | 1.39, m | | 13a | 13 | |
| 13a | 1.51, m | 26.1, $CH_2$ | 12b, 14 | 14 | 22 |
| 13b | 1.40, m | | 14 | | |
| 14 | 1.63, m | 60.3, CH | 10, 13a, 13b | | 22, 23 |
| 15 | | 74.0, C | | | |
| 16a | 1.59, m | 39.8, $CH_2$ | | 14, 15, 17, 23 | |
| 16b | 1.44, m | | 17a | 14, 15, 18, 23 | |
| 17a | 2.16, m | 23.3, $CH_2$ | 16b, 18 | 18, 19 | |
| 17b | 2.03, m | | 18 | 16, 18, 19 | |
| 18 | 5.27, m | 125.5, CH | 17a, 17b | 17, 24, 25 | |
| 19 | | 131.3, C | | | |
| 20 | 1.62, s | 16.6, $CH_3$ | | | 1a |
| 21 | 1.62, s | 16.6, $CH_3$ | | | 5a, 5b |
| 22 | 1.06, s | 23.7, $CH_3$ | | 1, 10, 11, 12 | 1b, 8, 13a, 14 |
| 23 | 1.09, s | 27.3, $CH_3$ | | 14, 15, 16 | 9a,14 |
| 24 | 1.61, s | 17.7, $CH_3$ | | 18, 19, 25 | |
| 25 | 1.68, s | 25.9, $CH_3$ | | 18, 19, 24 | |

[a]Recorded at 600 MHz in Benzene-$d_6$. [b]Recorded at 150 MHz in Benzene-$d_6$.

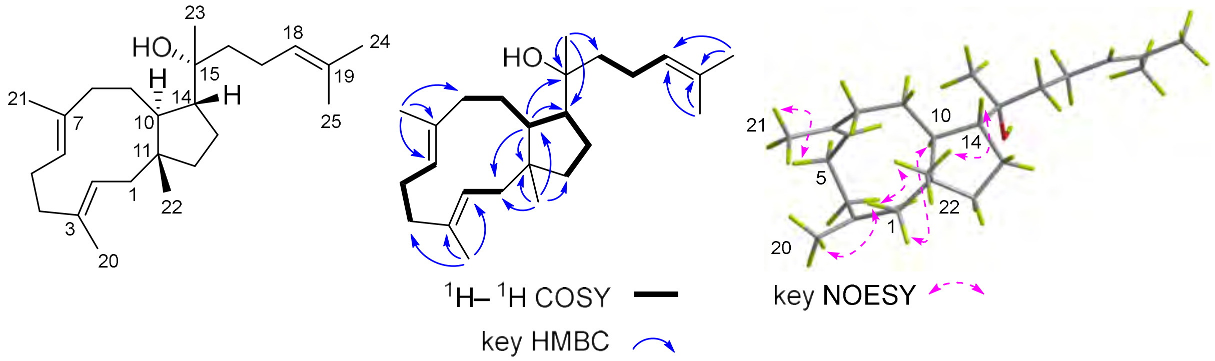

**Table S30.** Oligonucleotides used for construction of expression plasmids.

| Insert | Primer sequence 5'-3' | Vector |
|---|---|---|
| FsPS | F: AAGCTCCGAATTCGAGCTCGATGGCAAATGAGACTTTTTGTGAAG<br>R: GAGCTACTACAGATCCCCGGCTAGGCCTCTAAGCTGCTCA | pUARA4 |
| AcPS | F: AAGCTCCGAATTCGAGCTCGATGAAAGACCTACGCCATTTCAA<br>R: GAGCTACTACAGATCCCCGGTCATTTCATTTCTGTTAACTTGTAGATAAT | pUARA4 |
| FpMS | F: AAGCTCCGAATTCGAGCTCGATGGACTTCACATACCGCTAT<br>R: GAGCTACTACAGATCCCCGGCTAATCTATACGCAGCATCTCC | pUARA4 |
| AtVS | F: AAGCTCCGAATTCGAGCTCGATGGACAGACCAAAGCAAGC<br>R: GAGCTACTACAGATCCCCGGCTACACCTCGAGTCTCTGTAACAAAG | pUARA4 |
| AlSS | F: CTCCGAATTCGAGCTCGATGGAAGGACAGGAGCAC<br>R: CGAGCTACTACAGATCCCCGGTCAATTCATCGACCTCCAA | pUARA4 |
| CrNS | F: AAGCTCCGAATTCGAGCTCGATGGAGTACAAATACTCAG<br>R: GAGCTACTACAGATCCCCGGTTATACCTTGAGCATCTCC | pUARA4 |
| BbOS | F: AAGCTCCGAATTCGAGCTCGATGGACTACAAGTTTTCTACCTTG<br>R: GAGCTACTACAGATCCCCGGCTATACCCTCATCATCTCCAACAT | pUARA4 |
| OaFS | F: AAGCTCCGAATTCGAGCTCGATGAAATATCAATTTTCCACCATTGTAG<br>R: GAGCTACTACAGATCCCCGGTCAGAGCTTGAGCATCACTAG | pUARA4 |
| DqFS | F: AAGCTCCGAATTCGAGCTCGATGGAGTTCAACTACTCGGAA<br>R: GAGCTACTACAGATCCCCGGTTAAACTCGAAGTAGCTCCAATAAG | pUARA4 |
| XP_023093164 | F: GCTCCGAATTCGAGCTCGATGGAACAGACAATCTCGA<br>R: GCTACTACAGATCCCCGGTTAAACTCGCAGTCGTGTC | pUARA4 |
| THV50721 | F: GCTCCGAATTCGAGCTCGATGGCTATCCTACGTCAGTA<br>R: CGAGCTACTACAGATCCCCGGCTATACAGTTTTCAATCGA | pUARA4 |
| co17083 | F: CAAGCTCCGAATTCGAGCTCGATGAAGCAAACGACGTACGAC<br>R: GAGCTACTACAGATCCCCGGCTACAGCTGACTATCTACGTCCTC | pUARA4 |
| cm13013 | F: GCTCCGAATTCGAGCTCGATGGAGTACCCCAACTC<br>R: GCTACTACAGATCCCCGGTCAGACCTTCAGCTCCTTCA | pUARA4 |
| OCK92653 | F: CAAGCTCCGAATTCGAGCTCGATGGATTACAGAAACTCATCTG<br>R: CACGAGCTACTACAGATCCCCGGCTATATCTTCAACGTCTCT | pUARA4 |
| fc13317 | F: AAGCTCCGAATTCGAGCTCGATGACTCAACAGTCCTTTATCCG<br>R: GAGCTACTACAGATCCCCGGCTAGAGTATGCTCAAAAGGCGG | pUARA4 |
| he13146 | F: AAGCTCCGAATTCGAGCTCGATGGACTACTTGTACTCGGTTTTG<br>R: GAGCTACTACAGATCCCCGGTCACACTTTCAGACCCGCTAC | pUARA4 |
| xs14040 | F: AGCTCCGAATTCGAGCTCGATGGCGTACTTTTATTCCT<br>R: CGAGCTACTACAGATCCCCGGCTACAACTTCAACATTTC | pUARA4 |
| cg11192 | F: AGCTCCGAATTCGAGCTCGATGGACCACCCTAACTCT<br>R: AGCTACTACAGATCCCCGGTCAAACCTTCAGCGCCTC | pUARA4 |
| FsPS | F: CTATATCGTAATACCATATGATGGCAAATGAGACTTTTTGT<br>R: TAGTGATGGTGATGGTGATGGGCCTCTAAGCTGCTCAG | pXW55 |
| FsPS-S60A | F: CTGACTATTACGGGGCATGCAACCCCATCAACGGCAATATC<br>R: AAGCCAAATCACCAACATAATGTTGCCAGTCAAGTCG | pXW55 |
| FsPS-I68A | F: CCCATCAACGGCAATGCATTTTCATTGGCACACCCTTTATGTC<br>R: GTTGCAAGACCCGTAATAGTCAGAAGCCAAATCACCAACATA | pXW55 |
| FsPS-F69A | F: CATCAACGGCAATATCGCATCATTGGCACACCCTTTATGTCGACCA<br>R: GGGTTGCAAGACCCGTAATAGTCAGAAGCCAAATCAC | pXW55 |
| FsPS-H73A | F: ATCTTTTCATTGGCAGCACCTTTATGTCGACCAGAACGAGTCA<br>R: ATTGCCGTTGATGGGGTTGCAAGACCCGTAATAGTCAGAAG | pXW55 |
| FsPS-V88A | F: GTTGCCACATACGTGGCAGAGTTGGGATTCTTCCAAGACGATGT<br>R: CTTGACTCGTTCTGGTCGACATAAAGGGTGTGCCAATGAAAAGAT | pXW55 |
| FsPS-W159A | F: CATTGATAAGTGGAAGGAAGCCATGACGTCTCAACGTGAAT<br>R: ACTGGAGCTGAGCATATCGGATCAATAGTTTCAAGTTC | pXW55 |
| FsPS-A185G | F: CAGGGTCATGGATGCGGGAAACGCATTTATCGCTGCG<br>R: CGCATCCATGACCCTGAAGTCAAGATAGTTGTCGAAACTTTCAAATGT | pXW55 |
| FsPS-F188A | F: GATGCGGCAAACGCAGCAATCGCTGCGCTGTTGCTCTTCG<br>R: CATGACCCTGAAGTCAAGATAGTTGTCGAAACTTTCAAATGTCGGACG | pXW55 |
| FsPS-I189A | F: TGCGGCAAACGCATTTGCAGCTGCGCTGTTGCTCTTC<br>R: TCCATGACCCTGAAGTCAAGATAGTTGTCGAAACTTTCAAATGTCGGA | pXW55 |

| | | |
|---|---|---|
| FsPS-Y217A | F: TCATTCATCCATGTGCAGCGGCGCTAGCACTCACGAACG | pXW55 |
| | R: TAGGGGCAAGCATCTCGTGTTCCTCATCTGTCAGTGTA | |
| FsPS-E305A | F: CGATGCACTTTCCTTTGCAATGATGGGAAATCTCATATG | pXW55 |
| | R: AGGTAAAGACATAATCTGTCTAAGTTCTTATGTTGAGTCGTT | |

**Table S31.** Plasmids and transformants constructed in this study.

| Plasmid | Genes to be expressed | Transformant |
|---|---|---|
| pUARA4-*FsPS* | *FsPS* | AO-FsPS |
| pUARA4-*AcPS* | *AcPS* | AO-AcPS |
| pUARA4-*FpMS* | *FpMS* | AO-FpMS |
| pUARA4-*AtVS* | *AtVS* | AO-AtVS |
| pUARA4-*AlSS* | *AlSS* | AO-AlSS |
| pUARA4-*CrNS* | *CrNS* | AO-CrNS |
| pUARA4-*BbOS* | *BbOS* | AO-BbOS |
| pUARA4-*OaFS* | *OaFS* | AO-OaFS |
| pUARA4-*DqFS* | *DqFS* | AO-DqFS |
| pUARA4-*XP_023093164* | *XP_023093164* | AO-XP_023093164 |
| pUARA4-*THV50721* | *THV50721* | AO-THV50721 |
| pUARA4-*co17083* | *co17083* | AO-co17083 |
| pUARA4-*cm13013* | *cm13013* | AO-cm13013 |
| pUARA4-*OCK92653* | *OCK92653* | AO-OCK92653 |
| pUARA4-*fc13317* | *fc13317* | AO-fc13317 |
| pUARA4-*he13146* | *he13146* | AO-he13146 |
| pUARA4-*xs14040* | *xs14040* | AO-xs14040 |
| pUARA4-*cg11192* | *cg11192* | AO-cg11192 |
| pXW55-*FsPS* | *FsPS* | SC-FsPS |
| pXW55-*FsPS*-S60A | *FsPS-S60A* | SC-FsPS-S60A |
| pXW55-*FsPS*-I68A | *FsPS-I68A* | SC-FsPS-I68A |
| pXW55-*FsPS*-F69A | *FsPS-F69A* | SC-FsPS-F69A |
| pXW55-*FsPS*-H73A | *FsPS-H73A* | SC-FsPS-H73A |
| pXW55-*FsPS*-V88A | *FsPS-V88A* | SC-FsPS-V88A |
| pXW55-*FsPS*-W159A | *FsPS-W159A* | SC-FsPS-W159A |
| pXW55-*FsPS*-A185G | *FsPS-A185G* | SC-FsPS-A185G |
| pXW55-*FsPS*-F188A | *FsPS-F188A* | SC-FsPS-F188A |
| pXW55-*FsPS*-I189A | *FsPS-I189A* | SC-FsPS-I189A |
| pXW55-*FsPS*-Y217A | *FsPS-Y217A* | SC-FsPS-Y217A |
| pXW55-*FsPS*-E305A | *FsPS-E305A* | SC-FsPS-E305A |

**Table S32.** Nucleotide sequences of the 18 BFTPSs examined in this study

| Name | Sequence |
|---|---|
| FsPS | atggcaaatgagactttttgtgaagttgaggacatctccaaatatgacacaaagggcttcttcagcggcttcaaactgctcaggagcaagtttgag<br>tggcttgctaaccaggcggtgtacgaatttcgacttgactggcaacattatgttggtgatttggcttctgactattacgggtcttgcaaccccatcaac<br>ggcaatatcttttcattggcacaccctttatgtcgaccagaacgagtcaaggttgccacatacgtggtagagtgtatgataatcaatactgatcctca<br>actttaattataactaaacaattcagtgggattcttccaagacgatgtcatggagcacacggcaaaggcccatccgcaagtatgtctcaattcacc<br>tcaaatcagggacagtgctaacagaccttagtcgcagccatatgctgatgacttctccatgggagaggcggagaaagtcttgggactcagatc<br>aggggtaaagcaactgcttgctaaattgtttctcgaacttgaaactattgatccgatatgctcagctccagtcattgataagtggaaggaatggatg<br>acgtctcaacgtgaattgcgtccgacatttgaaagtttcgacaactatcttgacttcagggtcatggatgcggcaaacgcgtgagtttcaacttttac<br>gtttgaccttaattgggctaatattatttcctaatcgagatttatcgctgcgctgttgctcttcgggctaggttttacactgacagatgaggaacacgag<br>atgcttgcccctatcattcatccatgttatgcggcgctagcactcacgaacgactacttttcatttaataaggagtgcgacgagtcgcaaaagtcca<br>aagattatgaatttaccaacgctacaaaggttctcatggatcttgaaggccttgatgcagatgcggccaaagatcttgtcaagcaaaaggcgaa<br>ctcatatgccgtgcggttctcggatctggtatgcgcttttaaaacgactcaacataagaacttagacagattatgtctttacctcgatgcactttcctttg<br>agatgatgggaaatctcatatggaatctaaactgtccacgataccatccagagaaccgctacaacgatccaaatgctggtcttgagaatgtcat<br>gacagcagaactccgtggcatcgcaccctctgaaggatcaccgtggcacaataagcttctgcacagcgtctggaagcataatgagggaactc<br>atgaatcaccatggtcatcaccaaaagcgtcaagcttgtgcagagtaagcagagacattaacttgtctgacaaagtacgcactctaaaactcac<br>aggcaagatggcattgctaaatcggagtagtacttgaaagcacctttcgaatacatctcttcttgccctggaaagggagttagacgggaatttatc<br>gacgcgcttaacttgtgggctaagttaccgaacgacgcaatattaggagtccaaggagtaatagaccagttacatattgcttcgttaatgtacgtg<br>gctgacccatcatgagctcaaagctgacgatagtccaggatcgatgatatccaagatggatcgaagctcagaaggggatctccttctgcccatg<br>aggttttcggcatacctcaaactatcaacgcaactagcttcgcgattatcgatgcaatcgaaaaggccagacgtctatctattccacatgctgcag<br>atataacaacaagtaagtacaagagcaataaaatttacagcaattgtttagtaacgtaaagcagagcaattgcacgatatctttattgggcaaag<br>ctttgatttgtattggactcgtcattctatcactccatcagagtctgaatatgtgcagatggtgcgaaagagtgagttaccccccaacttcatttatcgc<br>gagcaaacagtctaatgaagttaaagagacaggtggtccgttccagctcataactgagctcatgtgtcacaatcttagcgaagggtaagtttaca<br>tgcttcatgagcagcacacttggcttactattaccgacagtttacgaggaatgttacatgagtttgttaatgttctggccgtgtttttccagatacgagat<br>gattatcaaaatttaacatctactgaagtacgtatttatattcttcctcagtttctcatcgccagatctaacagagctgcagtatacaatgaagaaagg<br>cgagctggaagacttggatgaaggcaaattctcattcatcttggtccatcacttcaattctaagcccgatcgtagcatcgtctacgaacttttacgac<br>agaggcatttgcaaggaggactttctaaggcttcaaaactgcttattatggacgaattgaagcgctcgaagagcatagaatacacacttgatacg<br>ttgaagaagcttgaggagcagtgtgaaaatctattgggaaagattgaagatataactcaagagcggaattgggttctgaggtaccttctacatcg<br>attgagactgagcagcttagaggcctag |
| OaFS | atgaaatatcaattttccaccattgtagacccagcgacctatgacaatgaaggcttgtctaatggcattgatctaagaaagaacaattttacgcact<br>tggaagatcaaggtgcgatccgcgcacagcaggactgggctaagcatgttacgcctatcaaacagtttaagggcacccttggtcacgactaca<br>gtttcatgacggtctgtgtgcccgaatgtataccaatacgtctggaggtcatttcgtatgcaaatgaatttgcatttatgtatgatggtgagtacaggtt<br>gaagaacaagaaagatatgagtctaaaatctaatgaaatgcgcgtagacgatactgagctggatactgagaaaaacgtatgcattcagtagat<br>cttgcacttgcacgtggctaaagatgtggtgctaacctacatagacgtccgcggagaacgataagttgatgggaatctttctagcaggcactcaa<br>ggctggagcccaccgcaggaccaatccagttccggcaaaatacgaattctaaaacaactgttctcggagatggtgaaaatcgacaaagagtg<br>cgcgatagtaaccatgaaagcgtgggccgagttcctcagagtcggttcaagtcgtcagcatggcacagtttttacacggctggaagattacctgc<br>cctatcgtatcatggacgtgggcgaaatgtgagcttggttgcttcaagtctgaccctaaatccgactaacattcatggcattaggttttggtttggcgtt<br>gtaaccttcggcatggccctgcatattccagaccacgaaatggatgcttgccacaagctcctagaacccgcctggatcgctgtcggactggcta<br>acgatgtcttctcatggccgaaagagcgtgatgcaagtcaaaggctcggatgtaatcatgttgtaaatgctgtttgggttgtaatgcaagagcatg<br>gcttttcccagatgcaagctgatcaatactgccgggaacttgccgctcagtatgtcgcccaatacgtggataatattcggaacatcaagaacgac<br>gagtcgatatctccagaccttcgcacttacgttgaagctatgcaatacagcatctcggggaatgtcatatggagcaaattctgccccagatataac<br>cctgagaagtgtttcaaccagactcaactggattggatgcagaatgggcttcccagctccgttgaattggacagcgcttctaatactagtccctcttt<br>cttatcaacaagcacacacagctcgccggtatctgggtcccaaatgacgatcgaatccaaggacggatggactgctgatttttccggcatcgtat<br>ccttgctgctcgattgctcattaccgcctttgtctcataaagtagaactttcacctgcgcccttactgaacattgcatactaacactagcaacagatga<br>tcagcgcaccatggatctatgtcgattcgcttccctctaaaggtacccgagacatgttcttggatgctctcaaccactggctacatgtggacgagcg<br>gagggcaactcaagtgaaaatggcgatccgtatgctacataacgcgtcactaatgtaatgccatttcacgtcacatcccatcaaatcacgactct<br>gctgaccttttcccgcgtgaccaggcttgatgacgttcaagatggctctcgactacgtcggagcaaacctagcgcgcatcgcgttttcggagtcgc<br>tcaaacgaccaattcagctgcattcttggtgaacgaatccatcaaactaatacgcgaacttacgggtgatcaaggggtcgctgtagtgctggaa<br>aaattgaccagtctcttcgttggtcaagctcaagacctgcattcatcgcgaaatctctctattccgtctttgactgaatacatccagacgattgaccaa<br>agtaggttcctagcctcgcagaagaacttcagccctgcgtcatgacaaccgctgaccaagtataccgtagaaactagcgctctcttcgaactgg<br>cctcgaggctcatgtgtctttgctcgacggcgactgtagtgcccaacagctcgctgtcgcgcttctgcattttacttggccgtttctttcagatccgcga<br>cgactatcaaaatctcacatcacctgaagtaagttacgtgttcacgtgtgtacgagctatcaatcattgctaacaaagagacgcgaaaagtatac<br>ctcgcagaaaggattctgcgacgatctgaacagcggaacgtatacgcttcccctggtgtatgcgatttcacaacagtcagagaatctcctgcttc<br>agaatctgctatctacaagactggctgaagggacgctcgacgatgcccaaaaacgtctggctctggatcagatgcagctggtaaagacggac<br>gagtttttgcgaaagatcttagcctcattgtacgacgagctgagagccgaactgcagtatatgtcgagttcatttgcatccgaaaccctcaaatg<br>gagctgatgctagtgatgctcaagctctga |

| | |
|---|---|
| DqFS | atggagttcaactactcggaaatcgtcgatccatccgcctatgatactgaaggactttgccaaggaatcgacgtgcgcatgagcaaattcacga<br>ctcttgaggatcgaggtgccatccgtgctcatgaggactggaacaagcacattggaccctgtcgtaaatacaagggaacacttggacccaggtt<br>cagcttcatctcggtggctataccggagtgcatacctgagagactggaggttgtcgcatatgccaacgagtttgcctttcttcacgatggtatgtgga<br>ccctttgactttgagtcttgaatttcaatcgagacttgctctctctggacgactaaagactcacggtatgttctattgacagatgtcactgatcatgttagt<br>cacgatacggtaagttgatccaaaatcatttttgaacatccccccctgccctttgttccctcccatgagactttgactgatgatcatggaaagggcga<br>gatcgaaaacgatgagatgatgtcggtctttcttgaggctgcccataccggcgcgatcgacagctcaaagaaggccgatgcacggctggaag<br>ggaagaaacgcattcaaacccagttgttcctcgagatgctgtcaattgatcccaaatgcgccaaaaccaccatgaaatcctgggcacggttcgt<br>cgaagttggctccagtcggcaacacgagactcgcttcgtcgagctggccaattacataccgtaccgcattatggatgtgggggagatgtgagtg<br>acgaacctagacgacatgagccttcttggtatttgatgttgacttgcacatgcgcatataggttctggtttgggcttgtaacttttgcccttggccttcac<br>atccccgaccatgaaattgacctttgccgcgagcttatggcaaatgcttggattgctgtgggattgcagaatgatctatggtcttggccgaaggag<br>cgagatgccgccaagcagcacgggaaggaccatgtcgtcaatgccatctgggtcctgatgcaggagcataagacggacgtgagtgaagcc<br>atgcagatttgtcgaaatctaatcgtggagtacgtcgccaagtacctcaaggtaattgaggctacaagaaacgatgagtcaatctcgttggacctt<br>cgtaaatacctcgacgccatgctttacagtatttctgggaatgtggtctggagtcttgaatgcccacgatacaatcccaagatcgcattcaacgag<br>acacagttggaatggatggagcgaggcgttccgtctttggagacctctatttccagggaagaaatatctggaaatgcctcggatagttctgctcctt<br>cgcctttggcagatgagttggtttcgatccgggatagagcaggaagtgctagtaccaaaatttcctctttgagtactgaagggtcttgctcgcctatt<br>cacaccgaggacgacagagccttgaagacattcgacgctagttataaaactctcgaggtcagttcccctccccgggggtccgattgtacgagct<br>ttgtcgaccgaaaggcagggaactgaataatacttctcaaatacagaatgatgtcctcgacgcgccctacgattacattgcatctatgccatctaa<br>aggagtgagagatcagttcatcgatgctttgaacgagtggctgcacgtgcccgaggaaaaggtgacaaagatcaaggatgcaatccgtactttt<br>cacaactcttcactactgtaagtgttgcaatcaacgagaaattggtctctaatcttcgatcaactcaccaattttcctagccttgacgactttcaggac<br>aattctcctctcaggcgcgggaaaccatcggcccataatatctttggaccggcacagaccgtgaattcggctgtttcttcgataatgcggggattc<br>ggtcaagtcatggaattttcttcgagcgaatctgctcagcgggtattgagtacgtactgtttgcatctaccacttgaaaattacaaatgaagggtcga<br>tgtgggttgttaacagaatatgtagagagcattgtggttttgttccaaggtcaggccatggacttgttctggacgtacaacggacgcgtaccgagtg<br>aagaggaatattacacgatgattgatcggagtaggtcttctttgcgcaagtcgatggcttttattgacatccataatagagaccggacaacttttctc<br>aatcgcgaccagcctcctactcgatgccgcagacaccaaggtggaagttcaaaactgtttctacaggctgacgtgtctacttggacggtgtttcca<br>aatacgcgatgactatcagaatctggtatctgacgatgtaagcatccatcccagctattcaaaactatacagaaagacctttcctctgacgagaat<br>gtggctgggtagtaccgaaagcagaagggtttctgtgaggacctcgatgagggaaaatggtcacttgctctgattcacatattgaacaaacagc<br>caaatcacgtaggactgctaaatgcactgtcaacagggaggaagcgtggcggcatgacttcagagcagaagcagtttatcttggacgtcatcg<br>aggaaggtggaggtcttgactatacgaggtccgttttgaaggacctgcattttcagttgtgtgcggaatcacggcggattgaggctttgattgagtct<br>cccaacctcgacatgaggctcttattggagctacttcgagtttaa |
| FpMS | atggacttcacataccgctattcgttcgagcctacggactatgacactgatggtctctgtgatggtgttccggttcgtatgcacaagggtgcagactt<br>ggacgaggttgccatcttcaaagctcagtatgactgggagaagcatgttggtcctaagctgcccttccggggtgcattggggccaagacacaac<br>ttcatctgtcttactcttccagagtgtctgcctgagagactagagattgtgtcttatgcgaatgagtttgccttccttcacgatggtgagatggcacttga<br>accttaactacttttcattgctaacaagtatagatattactgatgttgagtcagctgaaacggtaagactctttacttttcacaccgttgatgcatgctaa<br>cagtcaaaggttgctgctgagaacgacgagttccttgacgcccttcaacaaggtgttagagaaggtgacatccagagccgtgagtctggaaag<br>cgtcatctccaagcttggatcttcaagtccatggtggccattgaccgtgatagagctgtggccgctatgaacgcttgggccacctttatcaacacag<br>gtgcaggatgcgcccacgatacaaacttcaagtcacttgatgagtatcttcattacagagctacagatgtcggctacatgtacgctaatttaactct<br>caacttcatccatcactgacaattcacaggttttggcacgctcttatcatcttcggatgcgccatcaccattcctgaacatgagattgagctatgccat<br>caacttgctcttccagccatcatgtccgtgactttgacaaacgacatctggtcatatggcaaagaagcagaggcagctgagaaatccggcaag<br>cctggtgactttgtcaacgccctcgttgttctgatgagagagcacaactgctccatcgaagaagccgagcgcctctgtagagcacgaaacaag<br>atcgaagtagcgaagtgtctccaagtcacaaaggagacacgagagcgaaaagatgtttcacaagacctcaaggactacctctaccacatgct<br>gtttggtgtcagtggaaatgctatctggagtactcagtgccgaagatatgacatgaccgcgccttacaacgagagacagcaagccagactcaa<br>gcagaccaaggatgagcttacttccacatatgatcctgttcatgctgccaaggaagccatgatggaatctactcgtcctgagattcacagactgc<br>ctactcccgacagtcctactaaggagagctttgctgtccgtcctttggtgaatggcagtggacaatacaatggcaacaatcacgtcaatggagtct<br>ccaatgaagttgacgtgcgtccttccattgaaagacatgcctcaaccaagcgagctacttcagctgatgacatcgactggacggcacacaaga<br>aggttgatagtggggctgaccacaagaagaccttgtctgatatcatgctacaagagttgcctcctatggaagacgatgtaagttcaagatagcctt<br>cttgtcaccaataattactaattggatacaggtcgttatggaaccctaccgatatctatgttctcttccctcaaagggagtcaggaacaagaccattg<br>acgctctcaacttctggctcaaggttcccattgagaatgcaaacacaatcaaggccatcactgaaagcctccacggatcatccctcatgtaaga<br>cttctctagaatatgatggattatcttattaacactttgaaggctggatgacatcgaggatcattcacagctgcgacgtggcaagccttcagcccac<br>gctgtatttggcgaggcacagaccatcaactctgcaacatttcagtacattcagtctgttagcttgatcagccagctcagaagcccaaaggctctg<br>aacatctttgttggtaagtcatggcttcacgtatttgggatgaacatgaactaacagggattgcacagatgagattcgacaacttttcatcggtcagg<br>cttacgagctccagtggacctctaacatgatttgtccacctttggaggagtatttgcgaatggttgacgggagtatgtatcgcaatatcctgttttctta<br>cagtatactgactttgacagaaactggcggtttattccgtctcctcactcgtctcatggctgctgagtccactactgaggtagatgttgacttcagccg<br>tttgtgtcagctgtttggtcggtacttccagatccgagacgattacgccaacctcaagctcgcagacgtaggtctacttattcatttctactcatgcata<br>tcactaaccactcacagtacaccgagcaaaagggcttctgcgaagaccttgatgagggcaagttctcacttcctctcatcatcgccttcaacgag<br>aacaacaaggctcccaaagccgtagctcaactgcgtggtcttatgatgcagcgttgtgttaacggtggtctcacttttgaacagaaggtgctggca<br>ctgaatctcattgaggaggctggtggaatttcaggcacggagaaggtgctgcactcactttatggtgagatggaggctgagttggaaagattggc<br>cagtgtgtttggagccgagaaccaccagcttgagctcattctggagatgctgcgtatagattag |

| | |
|---|---|
| AtVS | atggacagaccaaagcaagcatcagatgatgagacttcttaccttgtctctctttctcactacgatacacacagcctttgctcggactacacattga<br>ggcgccacaaatacgaatccgaggccaacgctggatgttacgaagcgagacaagattggatcaaatacgtcggccaatcgaagaattcg<br>ggggatgtaaccctatcaacggcaacttctcggctgtggtcctaccacttaccaagccggaaaggttgcgcatgatcgcttatgtgcttgagtgta<br>cgatacctgtaccatacggtttccccatcactgaggtagtctgcttactgaatgcagatgcattcctttacgataatatgattgagcttgagaaacca<br>gaaacggtatgatagaagttcaattggtttttcagattgaagatcactgatgatcgtagtcgcgtgctgataatcaaaacaaaagagagtggaaa<br>aacacgaacccggcactgggcaagacacagattcaagcgaagatcatgatacagcttgcttccatcgactatgcgtgtactgaacgggtaaa<br>aagtgtgtggacagagatgattgatacgactctacgagacaagaacaagtgctttacccacctggaagagtacgtgaacttccgcataattgat<br>actggagcgccgtaagttttctctttgctgttgaataagtaatgcgcttgtcaaaactgactgttttaggtttgtggaagccatgctgctgtttggcatgg<br>gtgtgacgctcagcaaagaagaggacattaaactcgagcaaattcgcaagccttgttacgcagctctcggcctggctaatgacttcttctcctttg<br>accgcgaatatgcggacttcgaaaaatctggaaagtcgcaaacactgacgaacgcggtctggcttcacatgcaatggtacaacatggatgta<br>aatgcggcaaaaagcatgactttggaggcgacgaaaagatacgaggagcagtttctggggagctgtgcagagtttcggcagaataatgcgc<br>ccatacctgagaaacttgacagatatttggacgcgctggcttaccagatcagcggaaatgttgtttggagtttgaattgtccgaggtaccaccgag<br>agtacagatatgacccgaatgtgggcatggaagatggactgacagcggagtcgcttcaaaagacactttgtcttgagtacgatgcttatatacga<br>accgaggtaaaggaagagcgggatgaaagctcccggcgttcgtctaccgattcatcccttcaagcctccgaagacaacaacttcacaacgag<br>ggatgatgtttcgatctcttcttgtacatctgtctcttcttcgaccttgaaggacgatcaaatactttgcaatgatctgttaggccaggaaaatgtccag<br>gcacctttcgattacatcacttcactcccctccaaaggtgtacgagacaccttcatcgacgctctcaatatctggctcaacgtacccgagaccgtc<br>gtatcccgtatcaaatcactaggcaaccgtctacacgctgcatctcttatgctcgatgatatcgaagatgggtctaatctccgacgaggtcagccg<br>gcgacacacaccgtcttcggtattgcacaaacgatcaactccggatgttacgaaatcctcctagctgtgagtgaggctcaagaattgggcgccg<br>cagaggtgaagattgtcctcgaagaactcgccgaattgcacatcggccagtcttatgatctcttttggacatatcatacccagtgtccgtccgaga<br>acgagtatctggagatggtgaacaagaagactggaggtctgtttcgattgttggtgcggcttctgcttgcgtctgcagattgcgaattgttgcgagct<br>acgcaaggcgcagatattgagaagcttgtaagctgtattgggatacaataccaggtccgcgatgatcatcagaaccttcattcgccggaataca<br>gagctcaaaagggattttgtgaggatctggatgagggtaaaatgtccttcccactcgttcacgcgttgaacaattgcgaagacaacgctgagctg<br>tatcagcttctccgacaacggcgggatatcggatatttatcagatgagcagaaaaagcttgtgctggggcaactcgaccgagcgggcagcatg<br>acgtatacgcgagatacattgagacgacaacagggggagattcatgctggcttgaagagagttgaggatgttactggaagggaaaattggat<br>attgagggctttgttacagagactcgaggtgtag |
| BbOS | atggactacaagttttctaccttgatcgacgcttctgtccatgacaaggaaggcctctgcccaggcatcgatctgcgcatacatgaggccgccga<br>cctcgaagaggtcggtgcgttcagagcccaagaggactggcgtcgccttgtcgggccgctcaagaacccctacggcggcctcttgggtccga<br>cgtttagtttcatcactgtgacggtacctgaatgcctccccaatagattggagattattgcatacgccctcgaggtcggtttcatccatgatggtatgtg<br>aaaaccctctgtctacttgtggtgaatccttggcgtggctcatcaatttagatgtcatcgacaacgacattcagaacgcagatctagaagaaatgg<br>gggcggcgctgcaagatatgggtgagctcaaggcaaagggcgcttctggaaaggccaaaattgcagctcaaatcatccgagagatgatggc<br>cattgaccacgagagagccacggctgttgccaagagctgggcgtcgggcgtccaacactccagcagacggcaagaggccactgacttcaa<br>gaccctcgaagaatacatcccatatagggccttcgatgttggttacatgtatgtgccgattttatagcccaaaggccacaatgtactcaaactgata<br>cgatgcttcccaggcattggcacgggcttgtcacctttggctgcgccctcaccatcccagacgacgaagccgacgaagtccagcagttgctcat<br>gcctgctctagccactgcctccctcacaaatgacttgttctcctttgagaaggagcgcgacgatgccaacgtgcagaacggagtcctggtcgtca<br>tgaaagagcatagttgcaccgaagaagcagcgagagagattttgaaagagcgtattcgggctgagtgtgccaaatacgtccgcaatgtcaaa<br>gtcgttagcgcgcgcgcagatctctctctcgacacgaagaaatacattgatatcatacagtacactctgtcaggtaaccttgcttggagcacgca<br>gtgtctccgataccacaagaatatcaagcccaatgagtctcaaatgctacgggctacagaaggcgtggagaaacatcccgcaacatgggtac<br>ccaaagacacaggtgactatcctgccgccaatatccaatgcaatagtctcctcagtgaggaacttccgatgatgaataggatcatgggggaaa<br>acggcgatattgtcaacggcgtcggtgacgccgacattgtccacggagtcaatggagtcaaccatggcaatgctaatggactctatggtgctgat<br>cattcaaacggcatcaattgttccaaggactcccctggtatctgtgaggtcaatattctcaacgacaccaacggcagagacaacgctgacagcg<br>tcaatggtctcaactataccaacggcggagatggcgctgccaaagtcaatggtgccgatatctcccatgattgggcaaattctcataaacccgtt<br>cctgatgtatttgattctaagaatctcatggccatggcccttgaacgaaacctgacggacctggacgatagtgtatgttaagtctggacttacccga<br>gcctctggtagcagaattgttccctccgcagcaactgacgatttttataggtcattctgcaaccgtaccactacttaaggtcgctcccatcgaaggg<br>cattcgcgaccaggccatcgatgctctcaataaatggctatctgtccctgccgaatccacagtcagaatcaagagcatcattcaaatgttgcatgg<br>cgcatctctcatgtatgtatgcctttgtcagccccaaccgaatgatgttccgtcgccgctttacaacgcactctgcttggtctcattacaggagctaac<br>gcgtctcaggctcgatgatatgcaagacgcctctccgctccgtcgtggcaagccttccacccacagtatttacggcactgcacagacaatcaac<br>agcgcgacatatcaatatatccaagcaaccactgtggcttcccaacttggcaatcccgcttgtcttggcatttttaccgatgagatgcagcagctcc<br>acgtggggcagagttacgacctgcactggacggagaatacgcgatgcccctccgtgccagagtacctaaagatggtcgacaaaaagacgg<br>gcggcatgtttcgcatgctgacaagactgatgcttgccgaaagcccgaccggctcacagattcctgatactgatctgagccggttcagctgtctac<br>ttggacgcttcttccagatccgcgatgactatcagaacctgatctcggccgactacgcccagcaaaagggatttgcggaagacttggatgaagg<br>aaagtactcatttgcactgattcattgtatccaaacgcttggtgcgcacacaaaatttgcgcaggatgctatgcatctacacgctcttctcatgaagc<br>ggagggtcgagggaaaattgagtgtcgagatgaagaaggagattcttgccatgatgaacaaggcaaaaagcctgaactgcacgcgcagtct<br>tctgagacagatgtttgatgagctggaaagagaggtcggtgttttggaccagagatttggtcgagaaaacttgtctcttagggtcatgttggagatg<br>atgagggtatag |

| | |
|---|---|
| AcPS | atgaaagacctacgccatttcaacattgacccggatgtatcgtcctaccttgacgatatatcatcctataacactctcgggcttgcggaaaactacg<br>aactccgaaggtccaagtatgagagcgagattcagcggggcatcctcgaggcccgccttgactgggagacctatgtcgggtcaaccacagtg<br>tgggcaggcgaatgcccaaagttcggttcctactaccactctgcgcttccaatggcgttgaaggaccggatagcgctaactgcttacatttttgaat<br>gtgtgtctctcatatgctctcgctcctctgtagtgccatcggtggaatatttgatcgagagttactaacttaagctgttcgatgtagatgcatttcgctacg<br>atactgttgctgttgaaaggtctacggaaggatctgtacgtatatacggcggcagtgaaatctgccgcctccggtctcgtcttgtactcgtaggctaa<br>ttgcccccccagctggagcaaaatggctattccgaaaaccttaacgtcgaagaaatacaatatcaaagcaaacagtccgccactggcgcaaa<br>gcagatacaagcaaagatgatcatgcaactaatggaaagcgatcccgactgcggggaggaaatcgtcaatgagtggaaagcaatgttgggt<br>atcactattcagcgcggcaataggaagccattcaaggattttgacgaatactgtaaattccggataatcgacggcgggtgccggtaagttgacta<br>gtttttggcgggggaatcctgcattcctgacctgatcggctatagctaacgctcgaatttagatgggccggtgcattgatcaagtatactatcggtgt<br>acatcttaacgaagaagagcttaagatcgagaacgaggtcttacatcactgttgggtcgctgtaactcttggaaacgacttgctctctttcgaggc<br>agagtatcgtgaggctcttgcccatgatgagcccggaatgaacaacgccgtatgggtcgtcatgcaacttgaaaatctagatcctaacgcagct<br>aaagaagttgttcgtcaaaaggtctacgcggctgagcaagcgtacctacgggttcgggaagagtttactcggaactgtactgagaggtatgctg<br>gattgttgaaatatgtggaggcgtattattatgtggtaccagcgaatacgctatggagcatgacttgccctagatacaatgatccgacacgtcctct<br>gccaccgatatcatcgtaccaaggtgcggccaaagaaatgatgagtttggtcgatcaaggcaaatcaatggatttccagacaggtagtctttaa<br>caaatcccgtgtgagaattgacgaccgcagggctgatatcgtgagaggtagtagaagccaaactaagccacatacgtatcaatggcaccccg<br>agcgaaggaagcagcagcatgtccgatacctccgagggggtggatatcctgacgtctccgacaacgtcgtcaccacctcctagcgagggctt<br>atccagtggagaagaccagggcaactctaaggccgggctaagcgatgatgtaagtacagtatatctcacagttgctctcgtacccaaacgctg<br>cctctctcacaatccaatttgcattattattaatttaattgataacacatttagatattacttgagccttacaattacttgacgtccatgccttccaaaggc<br>gtaagagaagcatttgtggacgcgctcgatctttggtttcaggtacctcgggaactcttgggcgaaatcaaatctattgggacaatactacatacg<br>gcatctattatgttagcaatgccggacttgatatactaattcagagagatgtatgactaacaatcgataatccgcaggctcgacgacatggaagat<br>gattcaccgcttcggcgaggacaggatgctgctcatgttgtttatggccatgctcagacgatgaattcggctaattatcttattgtatgggctatgaac<br>caggtacagaagctaggtaatccatcgtgctggggtgttttcttcggtaagcagagattccttccctaattatctgaaagtgagacgataatttcagtt<br>ttgtgcccccagcgctaatcgtcatcttctaatcttagaggagatccggaattcgttgattggacagagctatgagattcgatggagagataccatg<br>gaatgcccaactgaggagcagtacatcgatatggttgagaagagtatgtggcatgtcaccgttgatattgcaattcgcgattctaacatagctgag<br>caagaaactggtgggctgtttagatttcttgcacggctcatgatcgctacggcgacgacgaacaggtaattccataattcttccccaatcaggttat<br>cttgatattttcagactgaagagagagtatgatccactaaacttaaatataaagacacttatgcctggaccacctttcctatataatcgggcaatact<br>tccaaatccgcgatgatctgatgaatctcacaaatactacggtgtgtataagcgacactaatcccattattcaactctcatataggtaacttgctaaa<br>aacgaaaacccaacattctctctagtacaccgaccacaaggggttttgcgaagacctcgacgaaggcaaattctcatatcttatagttcacgctt<br>ggcataatactgcgccggcggataaccaacgtctccgggaaatattcgaggccagaagcaaagctaacagtatgccgagggagaataaag<br>aggaggttttggatattttacacaccgctgggagtttcgattatatcgagagaaagatgaagtcgctgcaccaagatattatacgtgaaattgatttc<br>ttggataagactactgagaagaaaaattggagtcttaggtatattatctacaagttaacagaaatgaaatga |
| CrNS | atggagtacaaatactcagaaattattcagtatgacacaaagggcctgtgtctggacattcctgtgcgcaagcacttgaacgccagcctagagg<br>accgtggatctcgtgaagcacgtgcggattgggagaagttcgttggccctctctcacagaatgcaggcgcgtactgtcagggtcacagcttcgtg<br>gccatttctgtaccagattgcttgccagatagactggaagttatttcttatgcaaatgagtttgcgtttctacacgatggtaacagtttatctgtcgacaa<br>tgaggaaggaaggcagccaagctaaccgagattcagatgtaactgatgttgtagatcaagatacagtacgttctaacatgttgaaggtttacttct<br>tttatcagaatgctaatttagtaacaggaaactcaagagaacgaccacatgttacaagcactgatggaaggtgttcagcaagggtcgattgagg<br>ccagagagtcaggcaaacgccagatccaagccaagattctcatgactatgatggctctggatcgagaccgagccaaagtagcgactactgc<br>ctgggccaagttcctagagcaagggtcaggaagagaaaaccataatcatttctcaaccctcgaggagtatctgccatacaggtccaaggactc<br>cggtcacatgtttgtctactccttctttttcccgcctcgaataaccagtgctaatactctcacaggttctggcatgccatcagtatatttggatgcggactt<br>actattcctgatcatgagctggccctggttacggaatatgtcatgcctgcttctctcgcatgcagtctccaaaacgatcttctctcattcggcaaggag<br>tatgaggctgcgttagaggctggccagaccgagatctgcaatgccctatgggttctcatgggagagcacaattgcacgttgaaggaggcaaag<br>accctgtgcatccaacgcatccatgaagaggttaggaaatacctgcaatatgcggagcgtgcccaagagatgcatgatttgagcgcagatggc<br>aagaagtacgtggatatgatgcgattcaccatcagtggaaacgccgcatggagtcttcgttgtccacgctataaccgccagcttcaaatcacgat<br>caatgaggatgagcgcctaaccctcgatggcattccaaagaaactcaaaggtagctccgcgttgattgaaaagtcgacgtcccctgaatcgatt<br>gcttccgacgtgtctgatgctggctatgtcactgccaacacagacagcgaatcagatcattcagacgccatgtcccgtgctactattgacggtgttg<br>acaggccagagacacccgcacaagagtcaaaggttaatggggtgaagccagatgatattggatcctccagagagccgttgatccttgagttg<br>gactcagaaagtggtgattggtcttccctgggccccgaagaggcggcgttggccatttatcctcagatgaagtctgatgtaggtcaattatcacat<br>gaatactgtagctcaaactgacatagtcccaggtcgtcactgagccatatcgctacctaagctctcttccatccaaaggagttcgcgacatgacc<br>atcaatgcactcaacttctggctcaaggtccccaaccattctatggatttgatcaagtccgccgtcagaatgttgcatggctcttctctcatgtcggttc<br>gaccacttctacctgtcaacagtcagtattcgctaacacgcccatttaggctcgacgatattgaagacggctctcgcctccgtcgcggcaagccct<br>cgacacacgaaatctttgggcatgcacaaaccatcaattctgcaacctaccaatacattaacgcaactcaagagatcaggaaattgagaaac<br>ccgcattctattgatatttttattggtaagtagttgagctatcgaagtccaattgctattgctcccgactaacgtaaacatacagatgaggtccagcaa<br>ctcttcattggccagagttatgaccttctttggacatctgagttgtcttgcccgacagtcccggagtatttgcagatgattgacggcagtaggtgttctc<br>cttgtcttgttttatactagtgatcccttcaacgtgctgacaaaagcttagaaactggaggcttgttccgtcttctcaccagactcatggtagctgaatct<br>aagtactcaggtgaagtggatttcgcgcgaatgtgctcacttttcgggcgatacttccagattcgagatgactatcaaaacctcaccagtgatgat<br>gtaagaagagaaaagtctgatgttttttttttccgcaggctgacctttggctgcagtacaccaaccaaaaaggattctgtgaggatcttgatgaggg<br>gaaatactcccttcctctgatcctggccctcaccaaatcgcctcgtaaaggacaaattaattctctcctggtacagcggcgtgtaacaggctctttg<br>acatatgagcaaaaatgcttgattctggaacacgttcgggggtctggtgctctcgaggcgaccaagacgatcatggactcgttagctaacgagc<br>tcgagggagaggtggacaggctcgggaagatttttggggaagaaaatctggagctcaagttgattcaggagatgctcaaggtataa |

| | |
|---|---|
| AISS | atggaaggacaggagcacggcctcaacatgccggagaatggtgtcccaaagtctcatccactcatgtacagacactccgaccctgcactcat<br>ggacgcagagagcgaaagcaagtatatctcgcggtttcagccgcgggtcagcaaatatgtcgcggtttctcacgacgcctgcatagagtgcca<br>cattgacctgttcgggacgggcgggatcggaaaagtggttggtgggatgaatgcgcatacggccgactttacggctctctgcgctccagaggcc<br>ttaccggagagaatcgggctttgctcttactttatcgagtatgcgtttgtgcatgatggtgagtgtttacctcttgttggaatgggttggttattgacacgg<br>tatattgctagacgtgtccgtggacgcggttcaagatggccctagctgccgcctttcccaggacaaggatggaatcctaccaaaggcattagggt<br>caaaaaggcttgaagacctccagaatatcgctgaaggcggactgaagagaaagcagtccagagcaaagatctggtcggtcttgaatgaaat<br>cgaccaagactaccttggtcgatgccagaggaccttcaagcaatggtaccaggctgcgcaacagatgagggacaaagcgtttgcaagtctgg<br>aagattacttggctgttcgggccttggactgcggagctaagttcgtgatgtgtcctcttcaacaatacactcccatcgctgacaataataagctgggt<br>cgttcgcatgatgggctgggccagcggggtcgaactctcaccggaagaagaggtggaaaccgggccggtcacctatttggcatttgtggttcttg<br>gtgtgaccaacgatctgtggtcttgggaaaaagagaagagagtgacgcggcagtccggggatactcttcctctgatcaacgccgtccagatgg<br>tcatgcagatgcaggatactgctgaagagtccgcaaaacagatagtccgtggccttatccgcgagcatgaggagcaatactgtcttctccgcga<br>tgaatatctacggagacccagtacctcgctttcggtaaagaaatggttccaagtcctcgagttgtccatggcaggaaacgcactgtggagcatcc<br>atgcccttcgatatcaccttgacgtcaagaatccatatatctgccctcctgaagttccctccgtcttcaaaaaactgcaggtaatgcgatcaaaaca<br>aagcaaaaagaggaaggacgcccggttaggaggtatggaatctattcaatatcaccgaatcagacgctgatcaagtgcgtgcagttgaaggt<br>gcaacagatcctgcatacaaggttcttgaagcattggacgaaacggtatgtcttcctcccattccgagaggccacgctgataactgcaggttctat<br>ggaaaccgtacgagtatatcacctctatgccctccaagggtttccggcatcacctgatcgatgctctgcacacttggttcaatgtaccagagcacg<br>cgttggcgctgatttctggtgtagcatccctgctccatgaagcctccttgatgtgagtttctagaacgcggtcactgtcttgtctttaggttaaccgcac<br>gcaggctggacgatatacaggacggcagccttcttcggcgtgggaggccagccgtccatgaaatatttggtgtcggccagaccatcaactctg<br>catgcttccacatcaataatgccctgcgccttgtccaacagctctcgccctctgctgtgctgatattctccggtcagtcttcttcggatcttcaatcaac<br>gacagtattgtactgaactcacgtagagcaaatggcacatctgtatacaggccaagcccatgatatctcctgggccggccacaggattatccct<br>accgaggaggactatttcaggatggtcgacggcagtatgtacccctgatctgtgctcctcgatctgggctcgtcgccctggctgacagcgaatag<br>aaacaggtggtctctttgtacttctcctgcgcttgatgcagcaggaagccacccaaaacaggttagacgtccctaaccatgccaaaatccattag<br>attgacgacccagggacctcgatatgcgacgcttcatgaaccagctgggacggtacttccaaatccgggacgactatcagaacctcgtatccc<br>aagagtatacgtcccaacgcggcttctgtcaggacctcgacgaaggaaagccttcgttcccgttcatccgcgcctgtcatactctggaggatagc<br>acagtgctcacggaatggctcaacatgctacgtagtggaggcggggtttcgatagaggcgaagcgatacatcctaactcagatagaagaaag<br>cgggagtctagagtatacgcgggatgtcctcagtgccctccttgatgatctagaaggtatgctacgggacatggaatccatgactgggcaggaa<br>aactggattctccgatctatgctggtgcaactgcagatcaaacagacaaagtcggtgcggaaggagagtacggttgctggagtgctgcgcgttt<br>ggggcggatatcgtgagacggcttggaggtcgatgaattga |
| he131<br>46 | atggactacttgtactcggttttggtcgacccatcaacctatgatacgcaagggctttgtgatgagattcccctgaggaaacacattgcggcggatc<br>tggaagaaatcgccaccttcaaggcatatgaagattgggaaaaattcgtcggtcctgtgggcgactttaagggaagccttggaccaacttttagc<br>cttgtctcggttatctatccggtttgccttccggagagagttgaggcaatatcctatctcaaagaaatcactttcttgctagacggtatggtacataacc<br>ctcatggagccatgcactcattatattagatgttattgatggaggtccaccacagctagtaggtatctctaattccgcgttcccatggaagcactgctt<br>acgcttgaataggccgaagctgcaaagaacgacctcctcacatccatcaaagaatacggaatgctgggacttatgcctataataaacaccgg<br>atttggtcaagtaattgccaaaggcttcagacaactgatcgaaatcgaccgcgagaagttccttattaacgcaaggcggttctggcaatcctatg<br>aacgatcgaagacggcgaaacagcttcgaccaaaatttaaggtctacgacgactatatcgcttatcgaattatcgatgtaggctactagtgagttt<br>ccaatacgcctaagcgagaagtttattgctaacggtagggtagtggttggggttatcttatgatctcctttgcctgtgaccttttaattcccgatgagga<br>gacggaccttgcggatgagctaacaaccccagcagtcgtggctgtaagcctaattaacgatttgatctcgtatcaaaatgacatcgacgaagca<br>gagaaggcgggcgatgatcacgtcaaaaacgctgtgtgggtaattatgaaagaaaacaatttcacacttgaagaagcaagccagcgctgcc<br>gcacaaaaataaaagaggagacgaggcgatatctagagattgttagaacatctagcaaccgaacggacatctccctcgatctacggagatat<br>ctggagtacttacaatacgcaatagtcggaagcaccgtgtggagccttaactgccctcgttacaataagggtatcgaatacaatgaacttcagca<br>actacgcagaaagcatgggatcaagaaatatccggccctttggcaattggagctcaactctgagagagctcccggccatatacgaaacgact<br>cgatgttggatatttccaaaaccgcagacctattcacccaaagttctaatgatttagagaaagaccgcttcaatcactcgctgataccgaataagg<br>atgatagctctagctgcgtatctgtgacttccaatgcggaagatgtaatttctcttgcgctcaaaggggatatcgcacagctcgatgaccatgttggt<br>gtttttccctcgaaatcatttattgtgctaacgtcccaggttatcatggaccctatcaactatatcgattcactgccctccaaaggtgtgcgtaataagg<br>tggtagacgcccttaatatgtggttaaaagttcctgcaaaatcagtcgaaacgatcaagacaatcatatcagaactccaccatgcatcattgatgt<br>aagatcctaatttcatggagtataatagctatgctaactggcaacaggcttgacgatatcgaagatggttcgccattgcgacgaggtgaaccagc<br>tgctcatcaaatctatggacattcacaaaccatcaactccgctgcctatctgtttccaaagtcgggtttcaaaattctcaccttgacgaaccaggagt<br>gtatgaagatattccttggtaagtagctaggattcgccaacaatattagagtgatgttgatgttattgtagaggagcttcagcggctttttatcggcca<br>aagttttgatatcaactggactcatggcctcaagtgcccttcgcttagcgaatacctctctatgatcgacgacagtaagcgagagccgggctattat<br>ttgttactagggtactgttaaccgagccatagagaccggcgggcttttccgtatgttggtaagacttatgactgccgaatcgtcggtagagactgcg<br>ccagatcttggaatttttgcgaagctcctcgcacgatatttccaaatccgtgatgactatatgaatctacgatcgtcggatgtgaggctcttaacccct<br>tgttttgttgtctggcctactgatatcaaagtagtatgcgaagcagaaaggatttgccgaagaccttgacgagggaaaatactcgctactcctgatc<br>catctaatacaagctgatcccaataacatgcagctccaaggactactcatgcatcgaagagtctctgggaaattaggagtagaacaaaagaa<br>gcttattttgaagctgatggaaaaggcgaatactttcgaagctacattgtcagttctcaatgcgctttatgatgaaattgatcgggaaatagggaga<br>cttgaaatcgcattcggattagagaacaatcctctacgattgctggtagcgggtctgaaagtgtga |

| | |
|---|---|
| xs140 40 | atggcgtacttttattcctcggtagtcgatccatcgacatatgagactcaaggtctctgtgatggcattgatctccggaagcacgtagcggccgactt<br>ggaagagatcggtacgttaagggcccaagaagactggcgccgcttagttgggccggtagcaagaccttataagggtggcctgaatccgaact<br>acagttttatgactgtgtcggtacccgagtgtatcccggaaaggcttgagataatcagctacgccaatgagttcgcctttctccatgacggtatgtctt<br>taggttacctagcatgtgagcaagcactaacagaacgaagatgtcatagaggttgctagtaaagatgtccgcgatgcttccaatgatgagatgct<br>acaagcatttgaagaaggtgctcagactggcaagatcgaagacaatggtgcctctggcaagcgcaaaattgccgctcagatcctcaaggag<br>atgatggcgattgactcggagagagccatggtcgccattaagagttgggcaacattcgtagaacacgcgagtggacggcagcatcacgttcat<br>ttcaacacactggaagagtatatcccatacagatgtctcgacgtgggatatatgtaagtcttctatcccctccaagcctgccttggtgttttaaccga<br>gttgcgcatcttttaacatcctattaaggttctggcacggcctcgtaactttcggttgcgctatcaccatcccagacgaagaagctgctgaaacaag<br>agagatccttactccagccttgatcgctgcatctctgggtaacgatttattttcctatgagaaggagtacgaagacactatgagagctggcctccct<br>gacgtcgtaaacgctatttgggtcatcatgggggagcatgattgtactgaacaagaggccaaagagatttgcaaagagcgtagtaagtagcttt<br>cgaaatggctctcttgccgttctgttccatctcaggataaagaaaccttcactcaggttaggagatattccagaagcccttgatgctctcgaaatctg<br>aagcatgtttcttaaaacgacgatggctgactacagggtttcagtccgcatggagaatgctaagtatgttaaggtcgtcgaggagacgaggaag<br>cgaatagatctatgcgacgacgtcagaaggtacatcgacgtcatgcagtactccctctcgggaaacctggtctggagtatccaatgcccaaggt<br>ataacaaaggtgctacgttcagcgagttacaaatcttgcgagctaagcatgggcttgagaggtaccctgcgttgtggccaccgaaagacacag<br>ccaatggccttcctgctcccactgctcgcaaggactctgctatcgacctcggtgtcagtgagtttacattagaggaggagcccaaaagggaaga<br>ccaagcaatgtactggggcatcaacggtgtcggcattcgtgtcggccatggtgtcgacgagcacggtgtcggtaactacaacgccgttcagac<br>gaatggccactcctttcctggccaggaaaacggagacaagtatatcaacggcaacagttcagatacttctagtggggctaagaagtcagctca<br>caactcaaagccgagcacagactcacttacactcaatagcgtagtatctttggctcttgatcggaaactaccggagttagacgacggtgtgagtc<br>taccatccttacatatggatatgcaccatgttcattgtgatggctaacgaagaaataatacgaaggtcgttcttcaaccttggcggtacctcaattca<br>cttccatctaaggggttccgtgatcaagccatcgagtctctcaatacttggctcaaggtcccttcgaagacagcccaaaagatcaaaaacgtcgt<br>caagatgcttcataacgcttccctcatgtaaatccccacaagcccaacgaaactacttctctcgctatatctccccgaaagtctgacggtatcttag<br>gcttgacgatcttgaagacggctcaccactccgtcgcggaagaccctccacgcatgatatctatggtcctgcccaaacggtcaacagcgcgact<br>tacttatacattcaagctacacagctagcctctgagctagacagccaatcgtccttccgcattttcacagaagagatgcgccgactttacgtaggc<br>cagagctacgatctgtactggacgcacaacgcactatgcccatctatggcagagtacctgagaatggttgacgataagaccgggggtctgttcc<br>gcatgttgacccgcatgatggtggcagagagtccagtcagcaaggacatgccggacattgatcttgacttgctgagctgtctcattggccggttctt<br>ccagattcgtgacgactaccagaacctggcttcggccgactacgctaagaagaagggtttcgctgaggatcttgatgaaggcaagtactctttca<br>cactcatccactgcatacagactctcgattcggagcccaaattggctggcgaggcgatgcaactgcgggcgctcctcgttaagcgaagattcga<br>tggtaagctaagtaatgaggccaagcgagaggtcttggccatcatgaagaggactaagagcttggaatatacgcttggagtcattagaaagct<br>ccacagtgagttggacagggaagtcgggaacttggaaaagacatttggcggcgagaacttccagttgcggctcttgatggaaatgttgaagttg<br>tag |
| co170 83 | atgaagcaaacgacgtacgacgcggtttctcacgtggtgactatcgaagacgacaccatccaagacttcagtagagacttcaccctccgcaag<br>tccaaagatgaggctgcagccaacgacggaacccacttgttaagaaaagactggcagcgattcataggccccatcgacgcattcggaggca<br>gcaatccttatcaaggcaacttcacctcgcttgcgctaccttacacacaaccggaacgaatcgggatatgcgcatacttcattgagtgtatgtgcgt<br>ctggttcgccgtcggactggtgctgatctccggcgatagattgcatcttgtacgatgacatctgcgaatcggcaacaaaagccctggaggtgagt<br>gcaagcattgggcgtcttttgacagacatttagctaacatggtattctagctggatccacacagtgatggtttgttcttcgagcacgctgactcgatat<br>tgggtgcgaagcagctgcagtcacgcatcattgtggagcttctcaacattgatgaagcttgcggcaatcttatcattgagacttggaagaatgtatt<br>gtcaacagcacaggacctaaagagcaatatacgatatacaaatctagacgagtacatcccgtaccgcattcttgatgttggtgctccgtaagcg<br>acccttttttcatgaatctggtagaataagctgacaaagctcagctgtgttcgagcaatcatgctcttcggcatgagtatccatttgactgccgaagaa<br>gagtctgtcctcaaggacatcattcacccggcttacgctgcctgcgggctcacaaacgactacttttccttcgatcgtgaatggtcagaaagagaa<br>gcgacgcgaaaagactcagtagcacctgagccaatgaagaattcagtatggttgtgtatgggatggtacaacttgtctgccgtggaggcgaaa<br>gaaatggtcaagcgagagaccatcaagtaccaagacgcattcaacaccaagagacgtgctttcgaggaagccaatccggcacactctgac<br>aacattcgaaggtgtctcgacggcgtcgctcagatggtcattggaaacgtggcatggagcttgcaatgcccacgatatcgacccgatcttcgtta<br>cgaccccaacgctgcggttgagaacatgttcctccattctgggacgcctaagttggaggatatcgaagcaacttgcgctagcctcaaatcacag<br>aaacacgcagacaatgccacaatgggagcatatggcgttgtggccgcaccaccagcaaccaggaagcagctccgtaaaagtcttgatgcta<br>ttgccggaagcattctgccgaaaaccgtttccagcaagctattggcggacgtggtaaccgcaccttttgagtggtgtgccagcttgccttcgaaga<br>atgtacgctcgtcgctcatagacgcattcagcatctggtttccggtgtctcgatccgccaaggaccgcatccaaaccattggtcatctactgcataa<br>tgcttcgctgttacttgacgacattggagacggcagccacctcagacgaggacagccgactgcacacaagatcttcggcgtgccagaaacgat<br>taattctgccaacttcgcaatattagaagcaatggagagagtttccgagcttccgccaccctcgttcacagtgtgcgcaacagagatgcgggag<br>ctcttcattggccagggtcatgatcttctttgggctcgccagcacaagtgcccttcagaagcggaatacttcgagatgatcgacggcaagactgg<br>cggactgttccacctcttgatcgggctcttgatatgcaactcgacgacccgggaagctcaaagtcgcgctcagattcgtgtacacaagcttttgac<br>ctcacttggcagactattccaagtccgcgatgactaccagaacctcgtgtgtccagcttatgccagccagaaaggcttctgtgaggacttagacg<br>aagggaagttttcttaccctgtcgtctgtgctctcgggcagactgagaaagacgtttctatgcttcgcaacatcctagcgaatcctcgagaaggtgg<br>gtgcctgtccgacgagctcaagcggcttgcactcaaacagctcgtttcttcgcgcagccttgagaaaccagaacaaccatcatgagacttcat<br>gccgaggctgaggagactatcgccgaggtggaagatatgcttcagacaccaaactggatactcaggctgcttgtcaaaaaattgagggtgga<br>ggacgtagatagtcagctgtag |

| | |
|---|---|
| fc13317 | atgactcaacagtcctttatccgcaactccatccccttacctaggcacacctacgaaggcgaagagtacttttgccgtttcaccccacgcatccac<br>cgagatgcacgtctttcggatgccggttcctggcaatgtcaggtcgacttcctcaaatcgagcaacgatgcgcgggcaggcgcggaccggaac<br>aaagatgttcatagctatgctgtaggctgtatcaatcccgtggtaggcaatgtcactgccctgtgtgcgtgtgaggctctgagcgacaggcttgccc<br>tgacaacgtacatggttgaatacgcctacatacacgacgatggtagggatatctacttgccttctcttcctgactgttgcatgctaatatgactttctag<br>ttatcgaatattccgacaagaaggatgagtctcatgtaacaatcccacccccgagatgttcactaagaattaccgtctaacgattttgaagcttcaa<br>aaagccaacgaacagcttatggagggcctgaccttggcaggtgatactagccgtggtccaaagtcaaaggaccacgtccagcgacgccaa<br>ctgcaggctaagatggtcatggaactcatggaggtcgacaaagaacaggcaaaagaaacactgcgtctctggaaggagatgtctgcagttttt<br>gtccagatcagagacatgaaagttcaccgtcctggaagactatctcaagttccgtgttgtcgatgccggatgcccgtacgtatgaaactccatatt<br>gcatcaatctcattactgacaagcctataccaggtggaccatgagtctcctctgcttttccatggacttcaaactcatacccgaagaagagaagca<br>gacatcggccatcacctcagctgcctatgacagctgggtccttgtcaacgactacttctcttgggagaaggagtggagcaactaccaatccaac<br>ggagccaccggcgtgatagcgaactccatctttctcttcatgaagtggcactcgatcgagaaggaagatggcaagaagatgctccgcaagga<br>gatcatatcacgtgaggagaagtactgcagcttgaaggacaaattccttgccaagggtgctgctaccgagaagacgcgtcaatggttggaact<br>actcgacctcgtgacggcgggcaatttcgcttggagcatgacgacagcacgatatcgagctggcgccccggacgtatatccggctctgtggaa<br>acattatgctgacaagaaaagtgacaccgaaagccttggaagacccatttcagtcaacgccaaagacatggctgatgacattgatgttgtccttc<br>acgatcgtaaatatctcgacctttccactgaagatgtcacgtcggagagtcgcgaatggaagaccagcgaaaaccgacccggtgtcctcaac<br>aacaacaatgacctccagaaatctcagcttggcgaaacgtggtcaattcgccaatacgaggatgtaagtggatttggctgatcatgcacagcg<br>atgacaggtctgacaataactagatgattctccaaccccagaaatacttggaaatgatgccttccaaaggtttccggaacgctgtcatagacgga<br>ttacaagtctggtatcaagttcctgagaaacctctcgcaatcattcgagaccttgtcaatcatttgcacagcgcctccctcatgtgagtgttacaacgt<br>tgaatcgaaccaagataggtctctaatatcactggaaggttcgacgacatcgaggacaattccccctaagacgaggatatccggctactcatg<br>ttgttttcggagtcaatcagaccatcaattcagctagcatactcatgctcaaggccttgaaggccgcggagtctctctcgtcccgagcgtctcgcat<br>gcttctggatttgctcatcgaggggcacatcggtcaaggcatggatctgtactggacttaccatacttctgtgcccacggaagaagagtacttcac<br>catggtggacggcagtaagtttggagtatcttttggtgatcaggttcttgtttactaacacgttggacttagaaactggcagtctctttactcttctggcg<br>gggctgatgcgctccgaagcgactacacatcgcgatctcgatgttagtctcctgatgaaacttgtcggtcgttctttcaagcacgagatgactgcc<br>aaaatctttgttgtgacgaggtatgttcagaaccagccagatacacaaaaccccatttgctgactatacgtctagtatacggacaagaagtgttttg<br>ctgaagatatcggcgagggcaagatatcgcttccacttattcacgtcctcgccaccaagtcacctcagcagggccgccttttgagcatactctag |
| cm13013 | atggagtaccccaactcgtacctcgtcgacccctcctcatacgacaccaccatggacggcctctgcaccaacataccccctccgcgtgcaccgc<br>aacgcctcgctcgccgaccgcggcgctctccgcgcccagaaggactggcagcgtgctttcggctccctgcggcaaggattcgccggaaccat<br>gggtccagagcacaacttgaacttggcctcgacatcgctgccggaaacgctgcctgaccgcctcgagctcatggggtacattgtcgaggtgatg<br>tgtctcgtggacgacatgatggacgcggtcgaatcgcccactgcggctattgggccgtacatggtcgaccttttgcatgcctacaacgtggtgatg<br>ggaggcggggagatggatggctcatcctgctcgccggttgcaggcatggtcattgacgtgggcaggcagatgttcgctgtggacgctgagcgg<br>gcacgggtggcgtgccggtgggctgagaaatgggcaaagctgatgttgagcacctttggcagcgagaaggagccccgtgactttgacgagta<br>ccttgaacacagacgtaccaatgtctcttcacagtacgttttctcttgagcaatctttgcacgggttgactaacgaaacgggcagggcgacgtttgg<br>attgtccatattcgccatgggtctcagcatccccgaggaccaacagcagacgtgcctcgacctttcagagcccatgtggataaccagctgcctcg<br>caaacgactgtcactcctgggaacgcgagcaccaagcagccgccgacaacaaccacgactccgtgtcgaacgccatctgggtgctgatga<br>acaggcactcgatgacgtgcgatgaggcgaaggccgaatgccgtcagcgagctacgcagtacgccgcagagtacctgcgggttgtcgacg<br>cggccaaggccagggacgatctttgtcaggacgccaagttcctgcttgacgtgctgcggtttgcaatcagcgggcatgtcgtgtggggtctgcag<br>tgtctccggtatcatcccgacaaaacgctgagcccgaaacaactcgagatggccaaagccgttgggtcggatctgaccacggggtggtgccat<br>cgcaaccaaaacgccacggacgcgatagctgagcaccccaaggtcgagaccattggggccgtcaccaacggcacgaacagcacccctg<br>gcagtggcttaacagctgtgcgggaggtccctaagctgggcctagaggtgtgcacacccccctttccaactcacctctcttgggatatcccacaag<br>cagctaattcctggaacaggccctcgaagcaccatcgcggtaccttgactcgctgccggccaaaggcatccgcaacaaaaccatcgatgcgtt<br>gaacatctggttcaaggtgccacaaacaacggcctctgcaatcaagcgggccatcaaccttctgcacggatgttccctcatgattgatgatattca<br>agattcatcacagcttcggcgtggaaagccggcaacccatatcgtctttgggacggtgcagacaatcaactctgctgggtaccgcttcctgagtg<br>cactgaacgaagtggtgaagctggacagtgagcggtgtatgggcatattctgtggtgagtcttgctcatctttccatggagacggatgaggcgtgt<br>gttgacagcgaacagaggaagtccaaagcctctacattgggcaaagccacgacgtctcgtggaccagcaaccttgtctgtccaaccgaagac<br>gagtacatggccatggtagatggcagtgagtctgcctgcgtttgcgcacaacccagtgacgctgaccaaaatggcagaaaccgccgggctctt<br>ccgcatgctggcgagactgttggacgcaaagtcagactcgcccaataagccggacatcacgctcctcacccgcttcatgacacttctcggccg<br>gctgttccaaatccgggacgactacatgaacctgacgtcagacgatgtaagtaatgcgcgctgagtaggatgaaggtttcaaacaatgctgac<br>atgccacaaccatcagtacaccaagcaaaagggtttctgcgaagacctcgacgagggaaaatactctctgccagtcattcatgctcttcgggcc<br>tgcgacgcctccgagactgctgcccacagcacagctatactccgcaatttgctctcgcaacgacatctcgccggcaggatgtcgctggaccag<br>aagaagttgtttctggagcacctcaggacgcgagggagtctcgagtacacgcagcaggccatgggcgaattgcaaaccgagctgaagaga<br>atggcggagcagatgggcatgctggacaacaaaagcctgaaggagttgttgaaggagctgaaggtctga |

| | |
|---|---|
| cg111<br>92 | atggaccaccctaactctcacctcgtcgacccctccacctacgacaccaacctcgacggcctcgccaccaacctcccgctccgtataagccac<br>agcgccgcgctcatcgaccgcggcgccttccgcgcccaaaaggactggcagcacgtcgtcgtcggcaccaccgccgccaccgtgccattac<br>caccagggtacgtcggcaccgcagggcccgagtacgggttcctagcggtgtgtgtgcccgacacgctgcccgaccgcgccgagctggtcgg<br>gtacatcgtcgagatggtgttcctgctcgacgacctggtcgaggccgccgagtcgcctgccgcggcggccgcgccgtacctggctgatttgctgt<br>ctgcgaaggacgcgctggagaagggcggggattgtgtgggtgtgggcggcgggcgcgtcgcgcggacgtttgtgggtgttggcagggcgat<br>gctggcgattgatcctgcctgtgcgagggatgctttccgctggctggaggtgtcggtgagggctatgttggtccgtttgggggggagcagggagct<br>tcgtgatttggacgagtacctggactatcgacgggtgaatgtcgcttcgcagtaggtttctctccccaaagatgcacgtggtggagcagggtcag<br>gcctaataaggggatgaaaataggtctatgttcggattggttctctttgccatgggcctcaccatcccggaagaccaacaacaaacgtgcctgga<br>gctttccgaggcgttctggcttcagttcgcgctcacgaacgactgccactcctgggagcgcgagcggaaggcggcgagcgacaacggccagt<br>cgtctgtgacgaacgccatctggatcctgatgaacaagcactcgatgagttgggacgaggcaaaacaggcgtgccgtgacaaggccagaca<br>atacgctgcggagcatgtgcgtgtcgtggaggcagctaaggcgagggatgatctttgccacgacgccaaacgcctgctcgaatggtttaggaa<br>tgcgatcagcggaaatattgtctggggacttcagtgtcctcgctataatgctgatcggacgttaacggcaacacagctcgcgctggcagaagcta<br>tccaggcggacgagacactcggttggatttggggccaagagaagaaggctataaacggtattgccaaacatgcgaaggctgagatcaacgg<br>gggtgttgcaagccgtactgccaccaacggtattatcaccactggtattgttatcaacgatgactcccaccaagagatggaggttgtgcaggacg<br>tcccgagcctgggcacagatgtacgtaacaagagctagaggccgcgttctggtcgttgcaagggctgaccaaatgtctaggttctcgaggcac<br>catcgcgctacattgactctttgcccggaaaaggcatccgtcacaaggctatcgacgcgcttaatgtctggtacagggtgccgccgcagcaggc<br>cgccatcgtttcaaaggccgtccatcttcttcacggggcatccctaatgctggatgacattgaggattcttcccgcttgcgacgcgggaagccggc<br>ggcgcatttggtctttggaaccatgcagaccatcaactcggccgggtttaggtttcttaacgccctcgaggaagtaaggaagctggatagccagc<br>gttgcatggatgttttctgccgtgagtgctaagcccttggctaatattacttgtagttgactaactcgaaactagaggagctccaggatctctacgtcg<br>gacagagctatgatctcgcctggacgtgcaacttgagttgtccgaccgaggaagagtaccttgccatggttgacggcagtgagtcgaatgccctt<br>tttgatgaggcgtaggggttagtagctaatctgggtggtagaaaccgcaggtctctttcggatgctggcgaggatgctcgatgcgcagtcggactc<br>gcccaccaagccggatgtggtcctcatcacgcggttcatgaccttgctaggccgcttcttccaaatccgggatgattacatgaacttgacctcggc<br>agatgtgagtttctctgtgagcgtaacagatagacatgatattgtggctaacccccaaggattagtacaccaagcaaaaagggttctgcgagga<br>cctggacgaaggaaaatactcgcttcccattattcacgccctcagtcgctgtgggaattccaagtcttcggatatcaagtcaaaagccaacagcc<br>acgtagctatccttcaaaacctcctctcccagcgacaaattgcagggaaaatgacattagaccagaagaacctgttccttgaacatctcaaaca<br>gacaggaagtctcgaatacacgaggcaagctatgggagcgctgcaactggagctgaagagcatggcaagggagatgggcatgcataaca<br>acgagaagttggcgggtttgctggaggcgctgaaggtttga |
| XP_02<br>30931<br>64 | atggaacagacaatctcgaattttgaagacgacacatcatatcgcgttgagcgcgggtcgccggatattgaaggcttctgtcgccactatgccct<br>gcgccggcacaagtacgaggagaaggcaaacctaggctccctccagtgccgagcagactgggtgaagtacatcggtcctatcgagagatg<br>gggaagttggaatccttacgaaggtcacttcggttctgtggtcctcccgctgtgcaagcctgacagacttgcaatcattagttacatcttcgagtgtat<br>ggataatctcactataggttcttatagtttagcctatgctaacaacggaactcctttttctctagatgcatttttgtacgataatgttgttgagtcgtctgcc<br>aaagcaacagtaagtaaacacggacaaactgattttgatactgctaggggtagctggagtctaacacgctacaaaatacagttaaacacacat<br>gcggataatatcggcctggatgaaacggaatacagaactatacgatcggtatcaggaacaaaacagattcagtccaagatgatgctcgaact<br>actctccattgatccaacatgcgcggaagtggttcttgactcttggaaaacgatgatcgctacaactgcaaggcatgataaagcaaaaccgttca<br>gtaatttagaggattatgtgaactaccgaatcattgacactggggcgccgtaagtatacggatatctgaaagtttgtccgtatccaaattctaacaa<br>atcttagtttcgttgacacgttgatgcgctttggcatgggcatcctgctgacagaggaagagattgagactgttacccctattgtgaaaccttgttatg<br>cggcgctaggcttagcgaacgattacttttcatttgacgttgagtggaaggaatttcaggagagacagtccgatgaaggtgccatgacaaacgc<br>cgtctggctctttatgcagtggaacaatgttgatgttgccactgcaaagaaactggtgtgggaggtaacaaaccggtatgaggaagaataccag<br>aggcgtgtggaggagttcatcgcgggtgaaggacgcaatgccgccaaattacatacttacttacgagcgttggcctaccagattcctggcaatat<br>agcctggagtttgcgttgtccccggtatcatcccgagctttgcgacgaggcaggccgccttctggaggatgaaatgtgctctgaaacggcaacta<br>atgtgttaggtcaccctgtggagcatgaacgaggctcgaatacaaccggatcggagaagtcgccggtatggactgccgatgataattcctcga<br>aaaggtcctcggtatcttctattgatgcaatcgaggaagaagctgagtcacctaagccggagcaactgggtacggaggtaggaatcttcgcaat<br>tcatcatagtagatgtaattcaaaatgtggttctgaccaaacctggaattcagcaccttttaggtccggccgactagattcgctcccttccgtccaaa<br>ggtgtccgagaagcattcgttgatgcgatgaatgtttggctggtgcttccgaacgatgtcgtgaatcgcgtcaaatcgatcgcagaaacgcttcat<br>aacgcgtctctattgtgagctggcacaatcttcatgtctgatatgactttaacaaagtagcaaacgctgacggttcaccttcaaccctaggttagac<br>gatatcgaagactcatcgccactccgtcggggtcaaccagcagcacatacaattttcggccagggccaaaccatcaactcagcgaattaccta<br>ctaatacaggccatgcatcaagtaagacagctcgataatcagcggtgcatggatgtttttgtcgaagagatgcggaaccttttcatcggtcagag<br>cttcgatctctactggactcgacagggcgagtgtccatcacaggatgaatacctggaaatgattagtcaaagtacctgattctcctctacgtgtctgt<br>ggtcgtgtactgatgattatttcctttaagaaacaggcggtctctttcgtttacttactcggctgatgacggaaaaggcatcgacacaacataacag<br>gtacaactcttttcagatcttggaaatgctataggtcaatgctaatctcttcctcaatagtactatcacacttgactcgctcgtcagcctattgggtcaat<br>acttccagattcgcgacgattataagaatctaacagaagacgtgagaagctaccaaccattggcttcgtctcctatatactgacaattacattaca<br>gtatgcaatccagaaaggcttctgcgaagacctcgacgagggcaaatattcgtttccattagttcatgcgcttaccacgcaaccacgaaacttcc<br>aactacgtggcatcttgcgggaatcacgaaacacgggaggcatgagtctgcccctcaaacaatgtgtccttgaacagctcaagcaggctgga<br>agcatggagtatacacacttgatactgggtaagctaatgaaggatatcatgtgtgagattgggtcccttgagcagaagacggggtgctcgaact<br>gggtgctacgacttttgatgacacgactgcgagtttaa |

| | |
|---|---|
| THV5<br>0721 | ctatacagttttcaatcgatacaacatcagccttaaaatccagttttcccccaacccagtcctttcttcaagcgttccgagttcatgttcaacatccag<br>atgcaagtcattcagtaatgtttttgcgtacgtaaagcttcccatgttttccatctcagtaattgctaatttcttcatttctgaagacattttcctctcattcca<br>ctttcgcgatcttaggatattgagaagtttcgcatcgccattttcggcttcttttccaagagcatgaatgattgggagggtgaaagatccttgatcaag<br>atcattgtagacagtatccagattcttttcatctccctagtgaattaccagttagcaattaggcacgagtttcaaagttcgggtctcggccaaagaaa<br>gttggttaaacatacctttgtactaggaataagatcgttataatcgtctcgaagctgataatacttccccagctttctcatcaatatatccgcgtctatat<br>cacgattcttagtcgcttctgcacgcatcatacctgccgccatgcgaaaaagtcctcctgtttctaaaagtgtaatgttagttttgatttaaaaatgagt<br>atagatattctacacactgtcttcggtctgtttgaaatattcctctagactgggacatttgcaatgaaatgtccaatgtaaatcttgactttgtccaatgtg<br>taattgcgatagagtatctatacaagtaccaagttagcggtccctcgagacaatcgtgggtgtggatgtggtgactgacgcgtaaagcaactgac<br>agattccttgctcagagccatgacaaactcaagacattttacatattgatacgttgccgagttgattgcttggctggtcccaaatacctcgtgtgctga<br>ggggtatcctcggcgtgtgggggagttgtcttcgatatcatctaatctgagaagtgttattatgggaactgaaggaagagcttgatgatagccaag<br>ctaatcattcgtcttacattagagatgtgttatggatgatgctaatgacattccgaatgatttccaagaattcgtgtggaacaaaataccaacagttg<br>agcgcgtcaagtaacgcattccttgagtttgatgctggtaaagatgcaatatatttgtaaggctccaggacaatctgaaagttttagtactgtgtttgg<br>gaggacatatgaagcttgagaattaccttatctgtaccattttgactgaggctagaggaatcaaagtttccgtttatgttcggagttttgacctcgaag<br>ctttctggcgtgtcaacgaagcatgtaccattctcgcttgggagctaagaaaactattagaaccgttttagagaggtgtgggtagagttgagagta<br>cctggtctaattcatccttagaccttttactatcacgctgtccatcagatttgacatcttcgatttgagatgcggggcgctttttatttatctttgtctttgaagt<br>tggggttggcgcccattcgttatatcgtttagaagtagcatgccaaaagcgattgcccgccgccgcgagctcgacaaggcccagaaatcgtctt<br>atatctgggctaggattctctgcttggagatactcgtccctaagcctgcaatattgcatttcatcatccagtattctatctctcaaaagctcccgtgctttg<br>gaatcttctacttcatgaagatgttttaatagccatatggcattaaccattggcatactaccattcgtacgattatattcgtcatattccttttcccatgagt<br>aataatcatttgtcagtaccatggctttcaatagagggagaatcacgggatccaaagacgacctttcattatcgttcaaattcacgtccgcagtgta<br>tataagcatctccctgcagatcctgaaatcgtcagtgtttgatatagaatggtcgaatggaaatgatagcataccaccccccagcatcgaaaaac<br>cgccataccagatattcgtcgagtgttttatacgatttcatatctctttgcggggacttaatttctaaaaatggctgataggcttcgaagaatcgagatg<br>ccccatcaggattcccttcatatatacgaagtgccattcgtgcttttatgctcttcatcatactattacttctgtcgcactcatctttgccaataaatgaatg<br>cattgtttgatgggctgcttcgaactatggatctgttttagttagaccatatcatgggtcatcgtttctcttacctcttcgggggtaaccttttcatccactat<br>atctaaatgtattccaaaccagtgtcaacatgaaatctttgcaatagaaactatccaccatgcatggattggcctaccatcatggatgaatgcatat<br>tcagatagataagtcatgggaataagtcgatctatctttgcttcaggaaatagtgtcgccgaaaagcttccgcttctttcattgatgcagcccacgcc<br>aaatgagcctttccagtcagcttccctaaggtcttcacgacactcccagctcgtattttcgatttcttgatttccctcatgaattcttgggcgaaagttgta<br>tgggaattcagggtcgatttcgaaagcttcaacaggatggggaattgagcgtttgaattcaattggtaggttttgatactgacgtaggatagccat |
| OCK9<br>2653 | ctatatcttcaacgtctctaagagtagacgaagtggaaagtttacttttcccagatccttttccacggattctatattctgctctattgccttgtgaagtata<br>ctcagcatctccgccgtacgcttaaggcatttctttcggttcatatcctctagaacagctcgcttctgttcgaaagacaactttcctgcaacgcgtcgtt<br>gtgtcagaatgttgttcagccacaggttgggcttttcggtctgcaaagcatatataagtggaagagaatattttccttcatcgaggtcctcgcaaaat<br>cctttctggttctcgtactacatgagtcctacgagtcagtttcctgcttcatataactgagatgcgccgatacacttacatcagcagacactaaattctg<br>gtagtcgtcacgtatctggaagtaacggccgaagagagttactacgttgcttatagcaggatgttccgatatgggcgagcgagcagccattagc<br>cggactaacatgcggaataatccaccggtttctgaatatatatcagtatctaaaagcagacgacgagaattgtgagccttgcttgcatcaatcatg<br>ttcagatattctgaaacagtagggcattcaatattgcatgtccagaacaaatcgtaactctgtccaagattgatcttttgcagctcctctatgtgagaa<br>ttgtcagacatgcaaggcgacaggatcaaaggccggcataccagtaaagatgcgcaagcatcggacatcatttagagtttggacctcttctaat<br>gcagcaactagttggaaatttgcagcgttaacggtttgtgggatcccaaagatcgtgtgagttgatgggtgtcctcgtcgtagaggtgaggaatctt<br>ccaagtcgtcaagcctagctgccagtaagtaaaagttaactctcagaagagaagggatgttgatcttacattaatgacgtggcatgtaccttgttg<br>atgacggatttgattatatcgagggaactcgcagggacctccaaccatatgttcaaagcgtctaccgctgttgacctaacgtttttggatggcagtg<br>agctaatgtagagaaaaggagcttgaacaacctttgaatcgtcagtagtcttcccggcttgtaagttttattcagcgctgttacttacttcatcatgca<br>gcgcaggaaagccttctgtcaaaaagatttctgctccatcctcgatggcggcagaacgattccgattgattccattcgctggcttgttatcaagattc<br>ccgttgcattcaagttttggatgtttggaagctctctcatcctgaatcacttcatttagcgtcgatgaactctccggtttatcctcactagtggaggttcga<br>tcagcatctccttccgcggtgggatttcccccggtagctgcagtgtcatgcctttgtggagttcctctctcaaggggatttggccgatagcgtgggcat<br>tggaggctccagtgaaagtttccgctgataccgtacagagcggactctatgtacttcctggtgtcaatcgaaagctccaagtttccctcaacctcttttt<br>gcaatgcgcacgaaatcagagacgcattcctttatcttctctctgagcagaagtgttgcagcctccacggagcacgaatgctcctgcatcaaaac<br>ccaaatagcatttacaacatgtgactcaccgtttgccttcgatacagcaagttctttctcccacgagtataagtcgttagttaaaccgccagccttgta<br>ggcagaccgacccattcggttggccaattccatctcctttgccggtatggtaagtcccatcccgaatgtggcgattccaaaccagaacctgttaga<br>gccgttaatgttaagtggtctactaaatcttctggtgacaccctaccgttcgccagagtcgccaaacctaaatgcaagatattcatcaaatgatcca<br>aagctcttccgttcttgagctgtaacgtcaccgaactctttccatgcctttgcagccgtcaatgctcgctctcgatcgatttttatcatttcaagtagcattt<br>tcgcctggatctgctttatgccgtcttccttcacatgctttacgctcgtttagaaaggacaacatcaccatcagaaattgattgcgcgaacctggtctg<br>agccttcgtctattatgtctaggttacagtgagaacagcatttaatttaatacgcagataactttaccatcgtggagaaacccaaactcattggcata<br>ggagagaatctcgaggcgttcgggaaggcactcggggaggcagacggccataaagttgaattgatctcctatcgctccattaaaggcttccag<br>cggcccgaaatatgttctccagtcgtcctgagcgcgttgaataccccgcacttccaattggggattgttgctgattctcaaaggatatccgcacag<br>aggttttcaacctcagcgacgactatcttcataacagatgagtttctgtaatccat |

### Figures S1 – S78

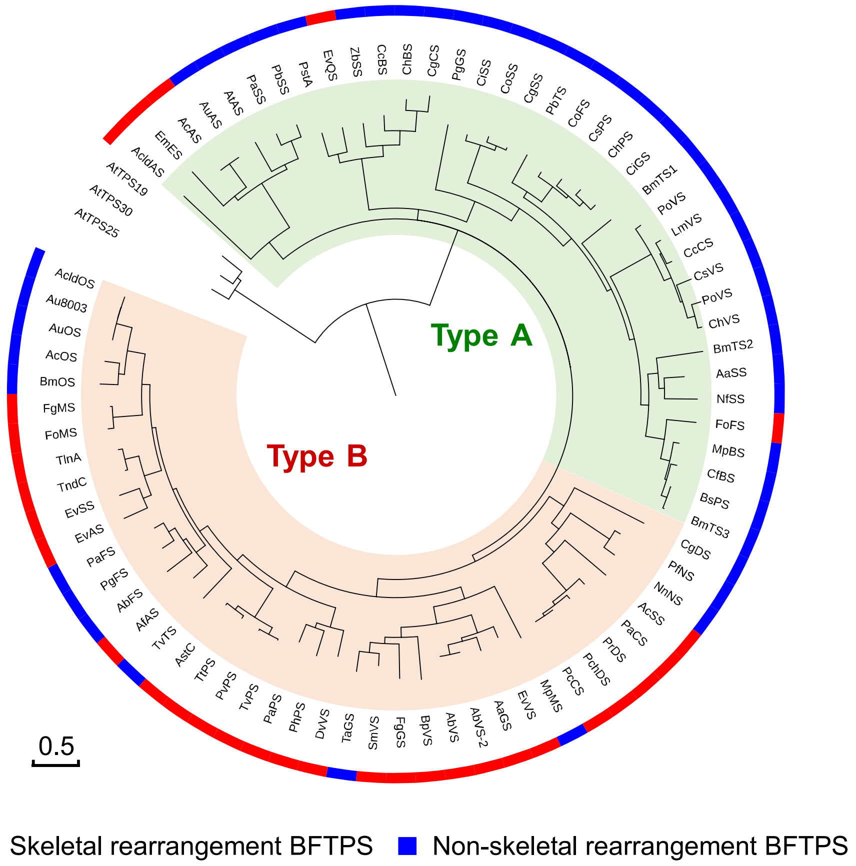

**Figure S1.** Phylogenetic tree of previously characterized bifunctional terpene synthases.

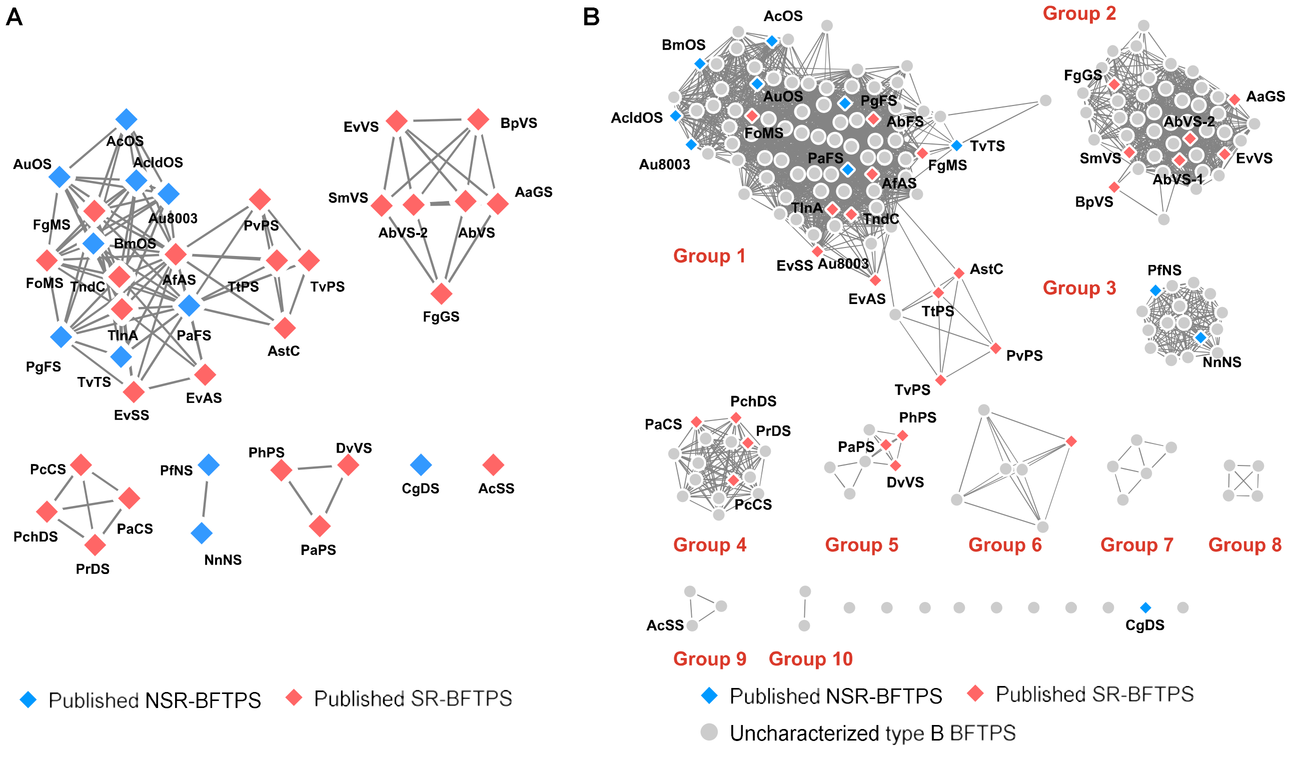

**Figure S2.** System similarity network (SSN) analysis of bifunctional terpene synthases. (A) SSN analysis of published type B BFTPS (E-value: 1e-100). (B) SSN analysis of published type B BFTPS and uncharacterized NCBI and in-house type B BFTPS (E-value: 1e-100).

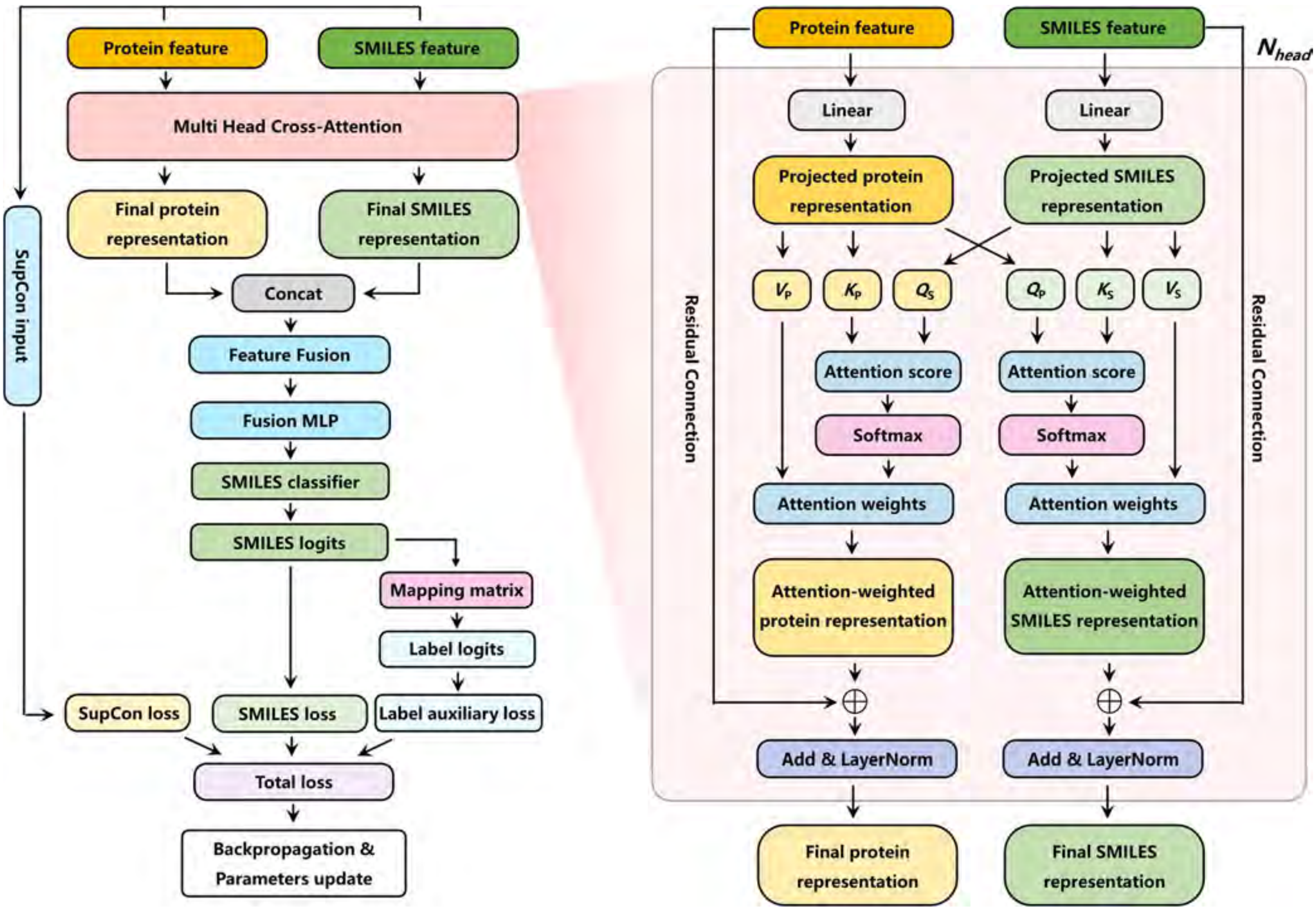

**Figure S3.** Training architecture and cross-attention module of the protein–product representation learning model.

Protein and product SMILES features are processed by a bi-directional multi-head cross-attention module to generate final protein and SMILES representations. The two representations are concatenated and passed through the feature fusion module, fusion MLP, and SMILES classifier to produce SMILES logits. SMILES logits are supervised by SMILES classification loss and are further transformed by a predefined mapping matrix into label-level logits for label auxiliary loss. SupCon loss is calculated from the pre-fusion protein and SMILES features, which are used as contrastive inputs to align experimentally paired protein–product embeddings. The three losses are combined as the total loss for model optimization. In the cross-attention module, $Q_P$, $K_P$, and $V_P$ denote query, key, and value vectors from protein representations, whereas $Q_S$, $K_S$, and $V_S$ denote query, key, and value vectors from SMILES representations. The protein-attended branch uses $Q_P$ with $K_S$, $V_S$ to generate SMILES-informed protein features, while the SMILES-attended branch uses $Q_S$ with $K_P$, $V_P$ to generate protein-informed SMILES features. Each branch is followed by residual connection and LayerNorm to obtain the final modality-specific representations.

**$Q_P$**: query vector from the protein representation. **$K_P$**: key vector from the protein representation. **$V_P$**: value vector from the protein representation. **$Q_S$**: query vector from the SMILES representation. **$K_S$**: key vector from the SMILES representation. **$V_S$**: value vector from the SMILES representation. **Attention score**: similarity score between query and key vectors. **Attention weights**: normalized attention scores after softmax. **Attention-weighted protein representation**: protein representation updated with SMILES information. **Attention-weighted SMILES representation**: SMILES representation updated with protein information. **Residual connection**: addition of the attention output to the corresponding projected input representation. **Add & LayerNorm**: residual addition followed by layer normalization. **$N_{head}$**: number of parallel attention heads. **Concat**: concatenation of final protein and SMILES representations before feature fusion. **Mapping matrix**: predefined SMILES-to-label mapping used to project SMILES logits into label space. **Label logits**: label-level outputs derived from SMILES logits through the mapping matrix. **SupCon input**: pre-fusion protein and SMILES embeddings used for supervised contrastive learning. **SupCon loss**: supervised contrastive loss for aligning paired protein–product embeddings.

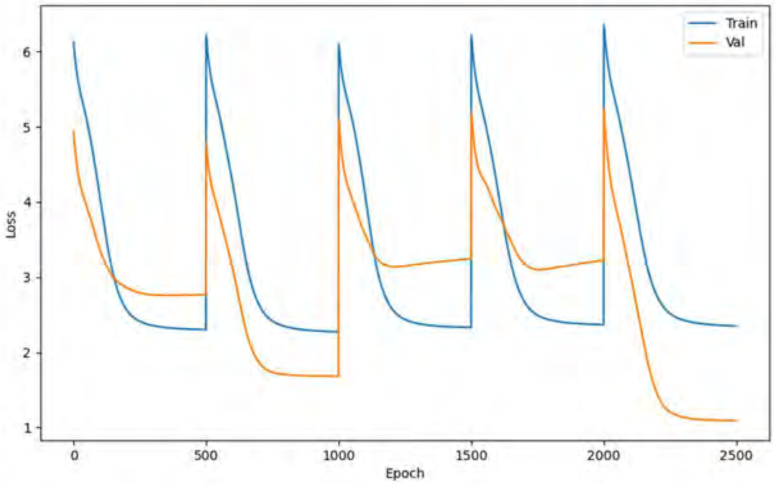

**Figure S4.** Training and validation loss curves of the model after removing overfitting-control strategies. Overfitting was observed in folds 1, 3 and 4.

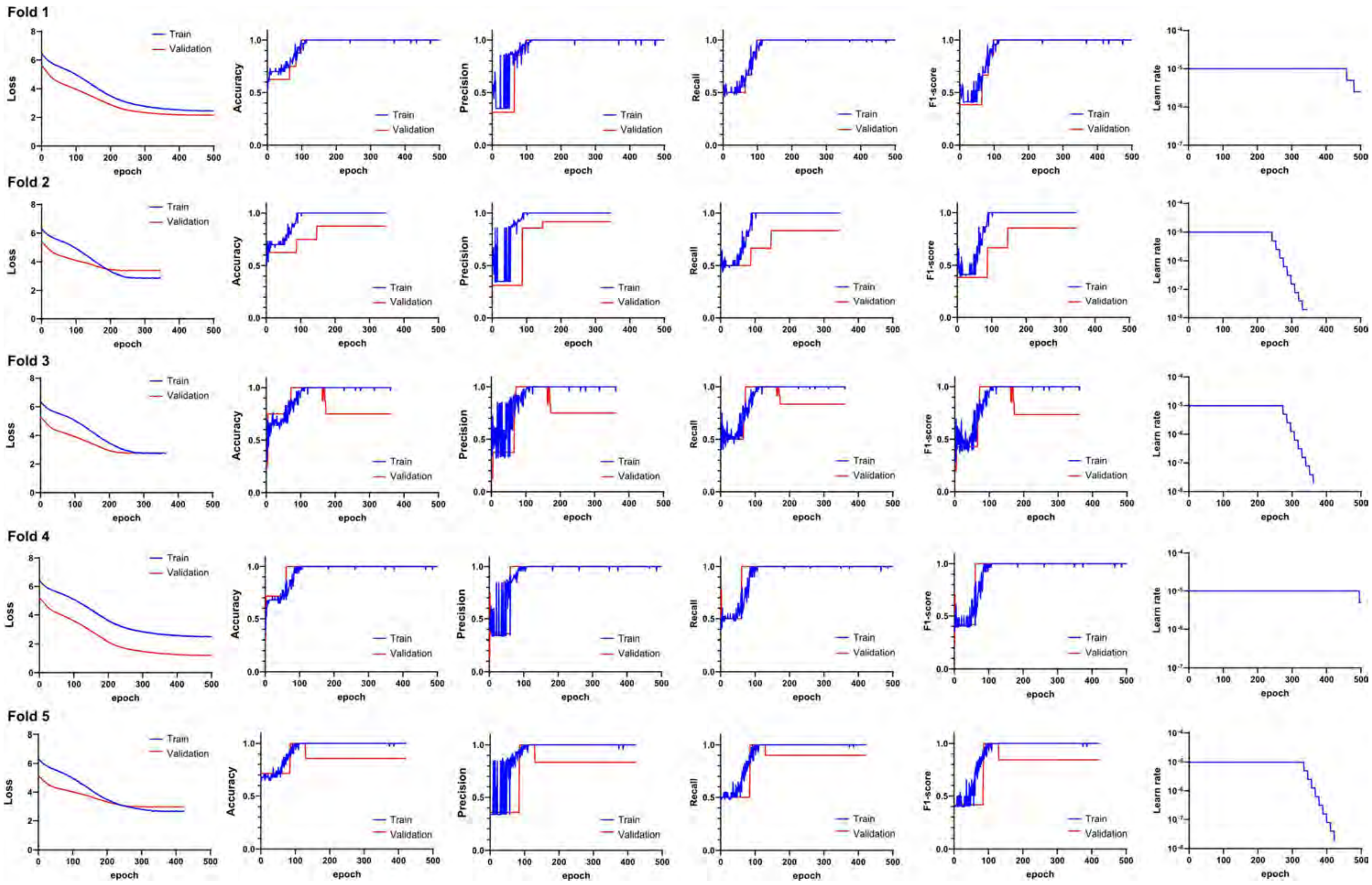

**Figure S5.** Training performance across 5-fold cross-validation.

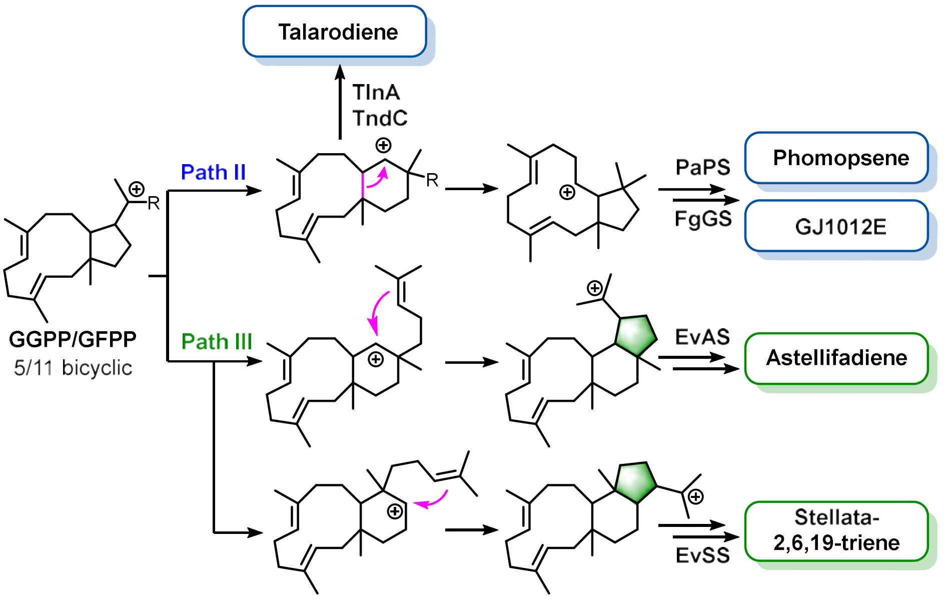

**Figure S6.** Schematic comparison of divergent cyclization pathway and product outcomes among Cluster 11 BFTPSs.

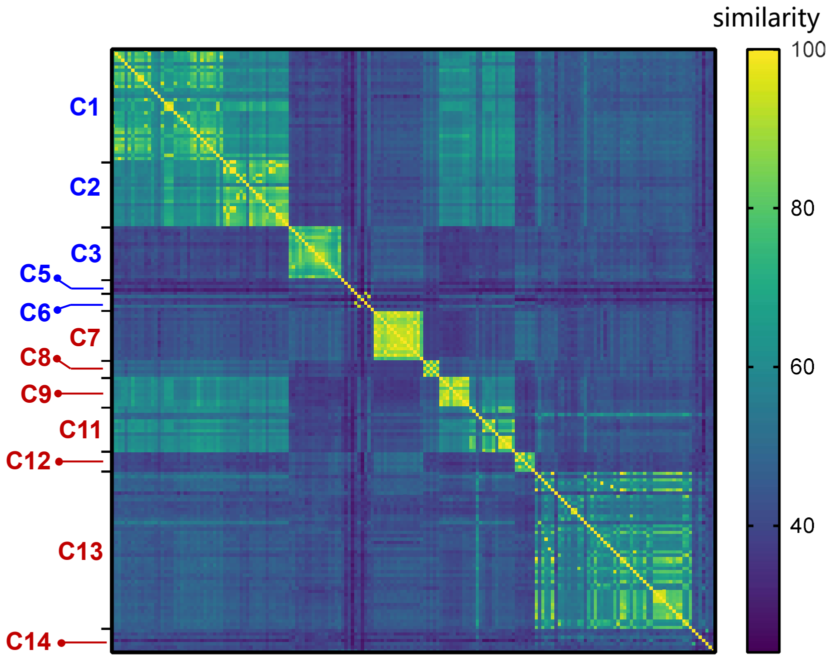

**Figure S7.** Sequence similarity analysis based on machine-learning–derived BFTPS clusters. C4 and C10 were excluded as each contains only a single BFTPS.

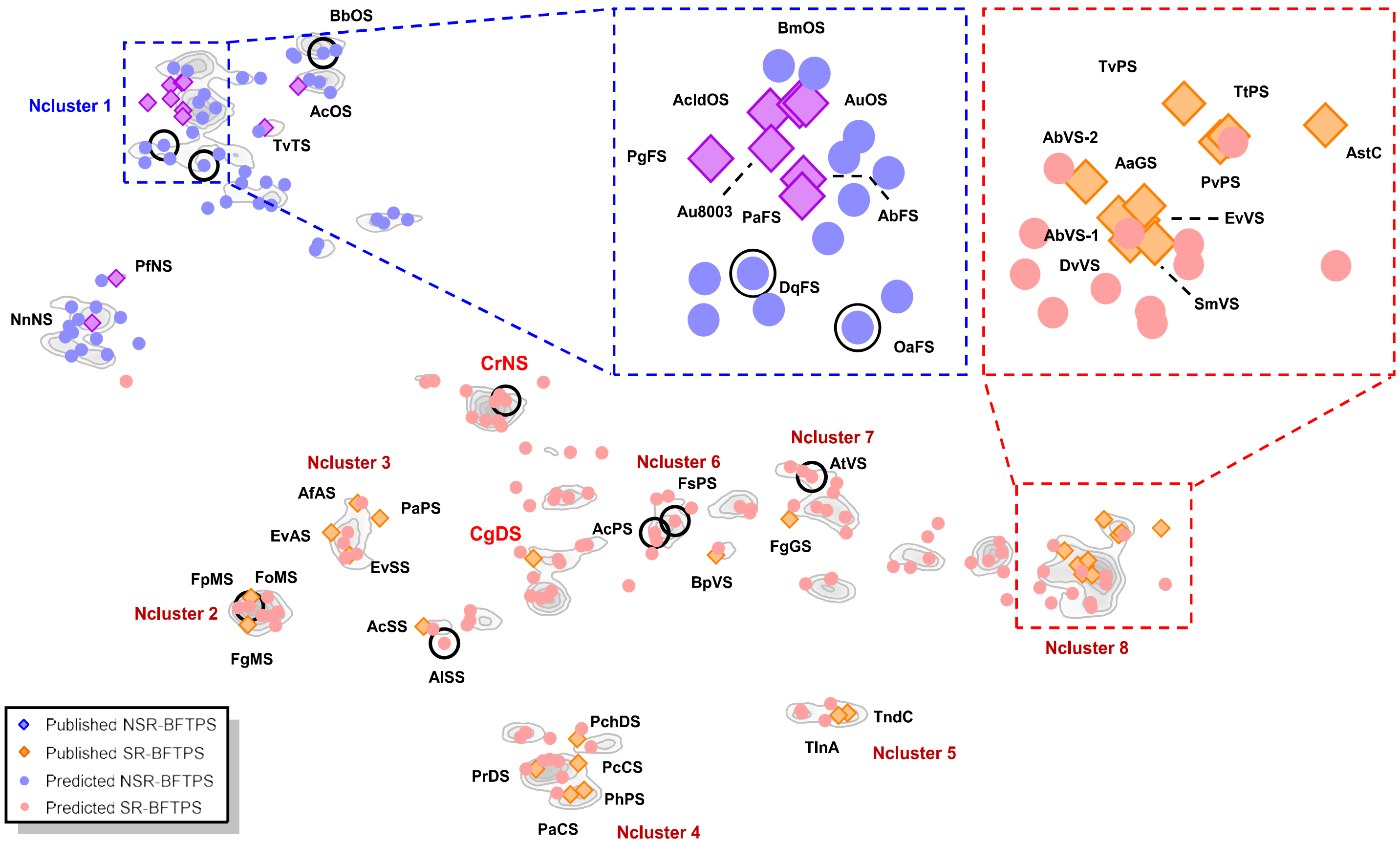

| Cluster | BFTPSs | products | Correctness |
|---|---|---|---|
| Ncluster 1 | BmOS | Ophiobolin F | Yes |
| | AcldOS | | Yes |
| | AuOS | | Yes |
| | Au8003 | Fusicoccadiene | Yes |
| | AbFS | | Yes |
| | PaFS | | Yes |
| | DqFS | | Yes |
| | OaFS | | Yes |
| Ncluster 2 | FoMS | Mangicdiene | Yes |
| | FpMS | | Yes |
| | FgMS | | Yes |
| Ncluster 3 | AfAS | Asperfumene | Yes |
| | EvAS | Astellifadiene | Yes |
| | EvSS | Stellata-2,6,19-triene | Yes |
| Ncluster 4 | PrDS | Penichrysol | Yes |
| | PcCS | | Yes |
| | PhPS | | Yes |
| | PaCS | | Yes |
| Ncluster 5 | TndC | Talarodiene | Yes |
| | TlnA | | Yes |
| Ncluster 6 | FsPS | **18-26, 27** | Yes |
| | AcPS | **18, 19, 22, 24, 26, 28** | Yes |
| Ncluster 7 | AtVS | Variediene | Yes |
| | FgGS | GJ1012E | Yes |
| Ncluster 8 | EvVS | Variediene | Yes |
| | AaGS | | Yes |
| | AbVS-1 | | Yes |
| | AbVS-2 | | Yes |
| | DvVS | | Yes |
| | SmVS | | Yes |
| | PvPS | Preasperterpeniod A | Yes |
| | TtPS | | Yes |
| | TvPS | | Yes |
| singleton | BbOS | Ophiobolin F | Yes |
| singleton | AcOS | | Yes |
| singleton | PgFS | Fusicoccadiene | Yes |
| singleton | TvTS | Talaropentaene | Yes |
| singleton | PfNS | Nigtetraene | Yes |
| singleton | NnNS | | Yes |
| singleton | CrNS | **17** | No |
| singleton | PaPS | Phomopsene | Yes |
| singleton | AcSS | Spiroviolene | Yes |
| singleton | AlSS | | Yes |
| singleton | CgDS | Dolasta-1(15),8-diene | No |
| singleton | BpVS | Variediene | Yes |
| singleton | PchDS | Penichrysol | Yes |
| singleton | AstC | Preasperterpeniod A | Yes |

**Figure S8**. t-SNE projection of the protein-only ablation model, generated without small-molecule information, groups BFTPSs into no-product clusters (Nclusters). Compared with the full model, Nclusters show reduced product coherence and include several misclassified enzymes, highlighted in red. The adjacent table summarizes the product composition and prediction outcome of each Ncluster. Density contours indicate cluster boundaries inferred from kernel density estimation.

**Figure S9.** HR-EI-MS and NMR spectrum of fusicoccadiene (**1**).

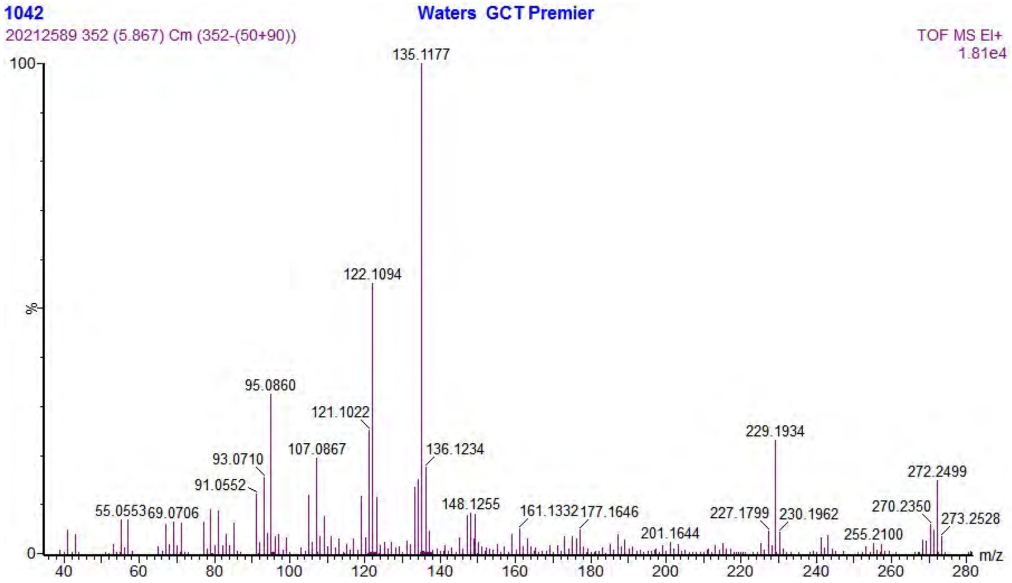

**Figure S9a.** HR-EI-MS spectrum of fusicoccadiene (**1**).

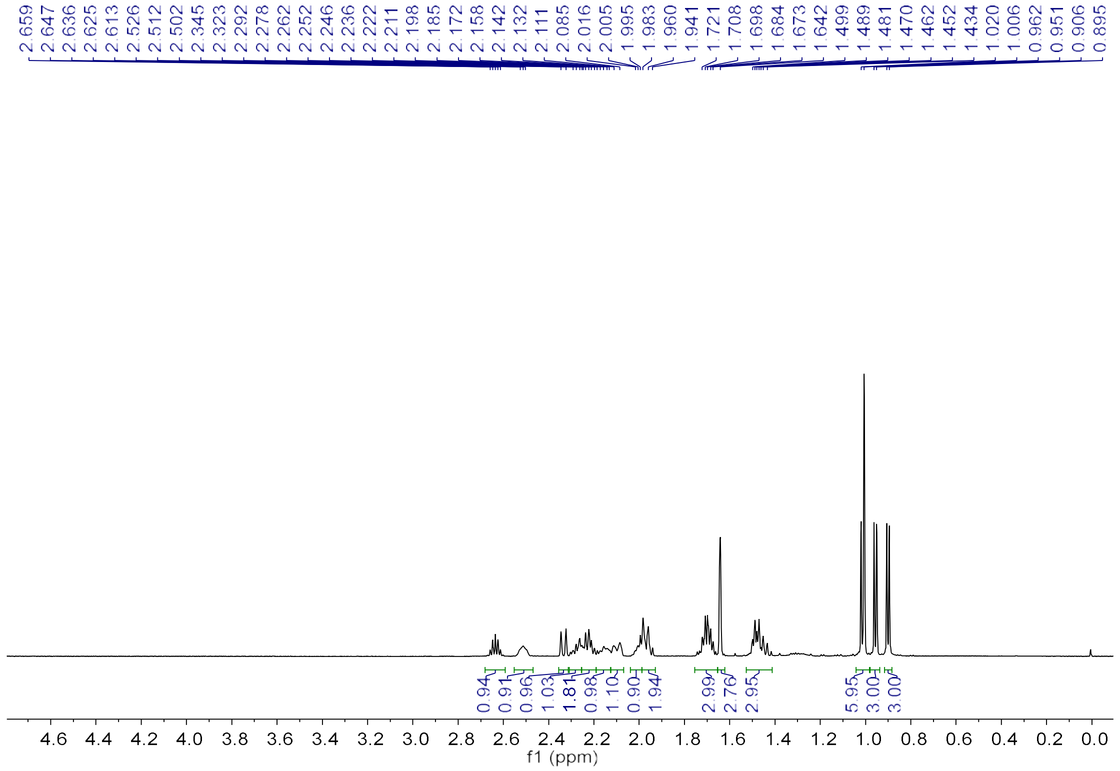

**Figure S9b.** $^{1}H$ NMR spectrum of fusicoccadiene (**1**) (600 MHz, Pyridine-$d_5$).

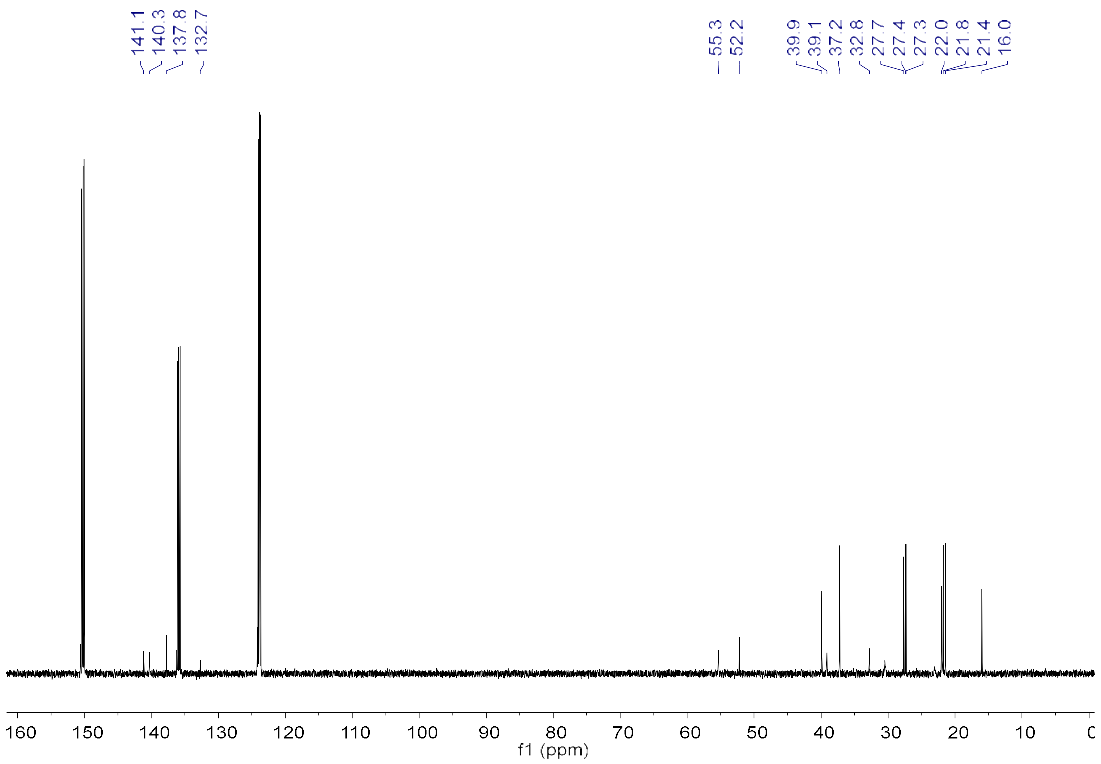

**Figure S9c.** $^{13}C$ NMR spectrum of fusicoccadiene (**1**) (150 MHz, Pyridine-$d_5$).

**Figure S10.** HR-EI-MS and NMR spectrum of ophiobolin F (**2**).

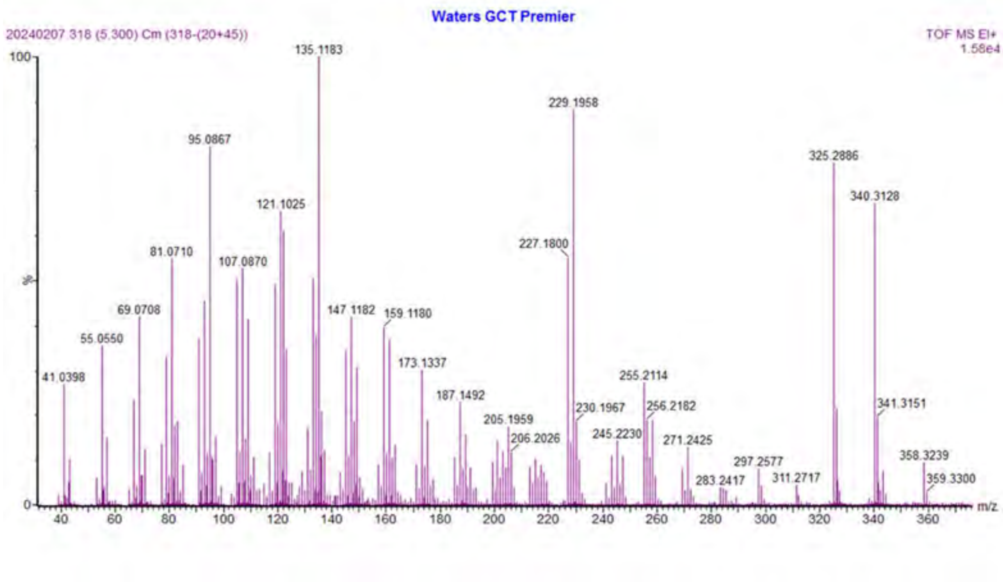

**Figure S10a.** HR-EI-MS spectrum of ophiobolin F (**2**).

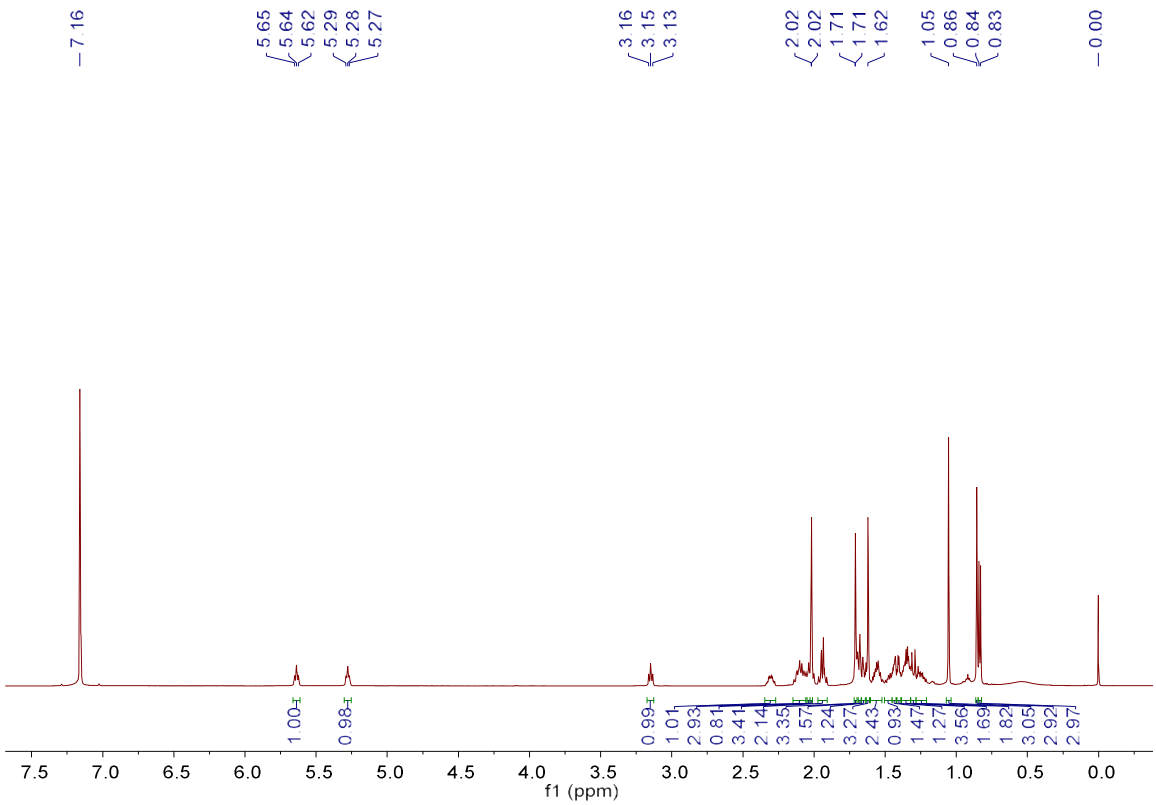

**Figure S10b.** $^{1}$H NMR spectrum of ophiobolin F (**2**) (600 MHz, Benzene-$d_6$).

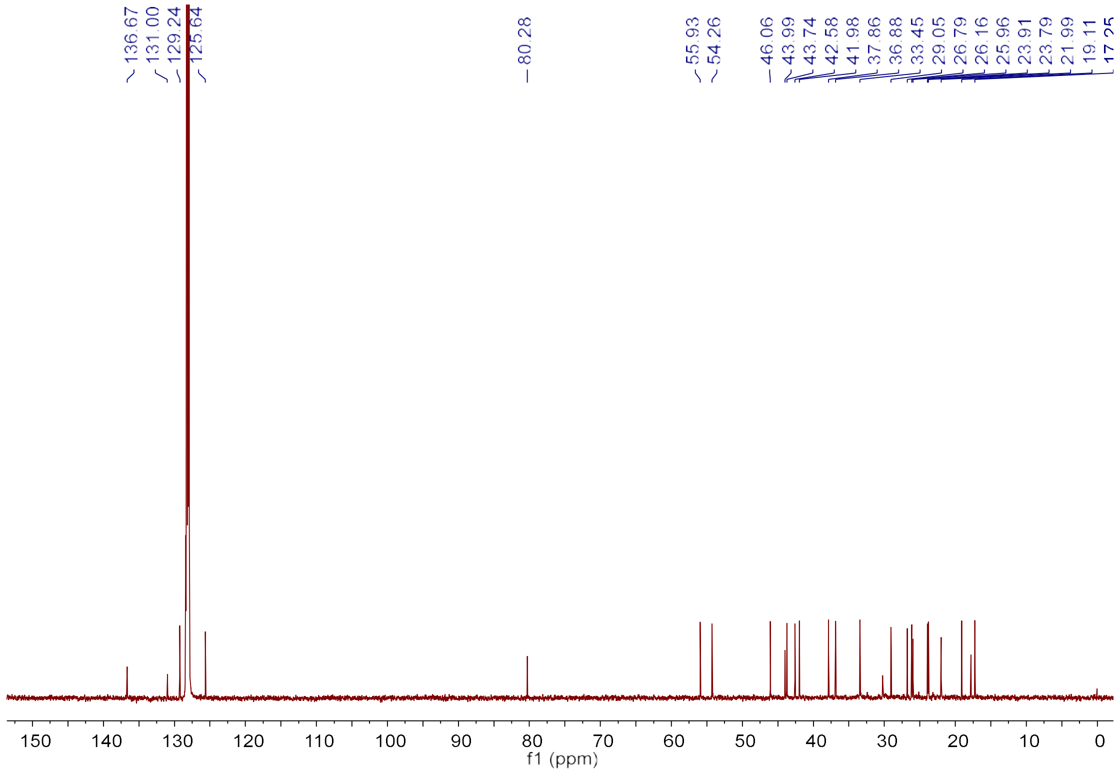

**Figure S10c.** $^{13}C$ NMR spectrum of ophiobolin F (**2**) (150 MHz, Benzene-$d_6$).

**Figure S11.** HR-EI-MS and NMR spectrum of mangicdiene (**8**).

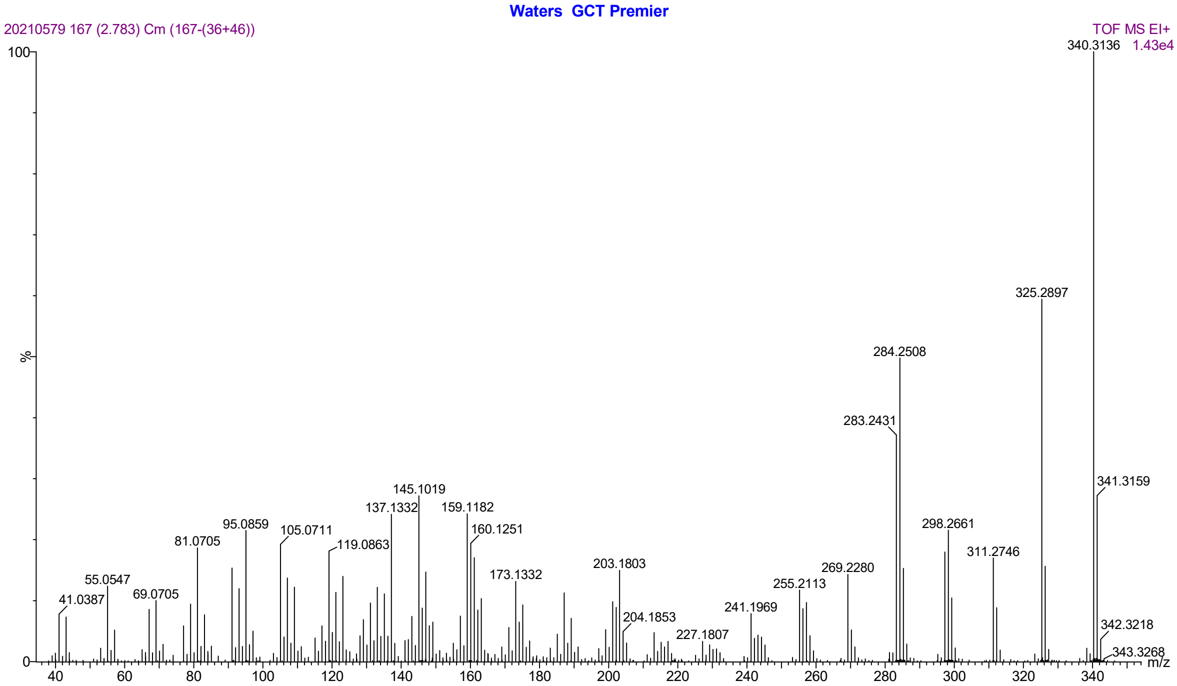

**Figure S11a.** HR-EI-MS spectrum of mangicdiene (**8**).

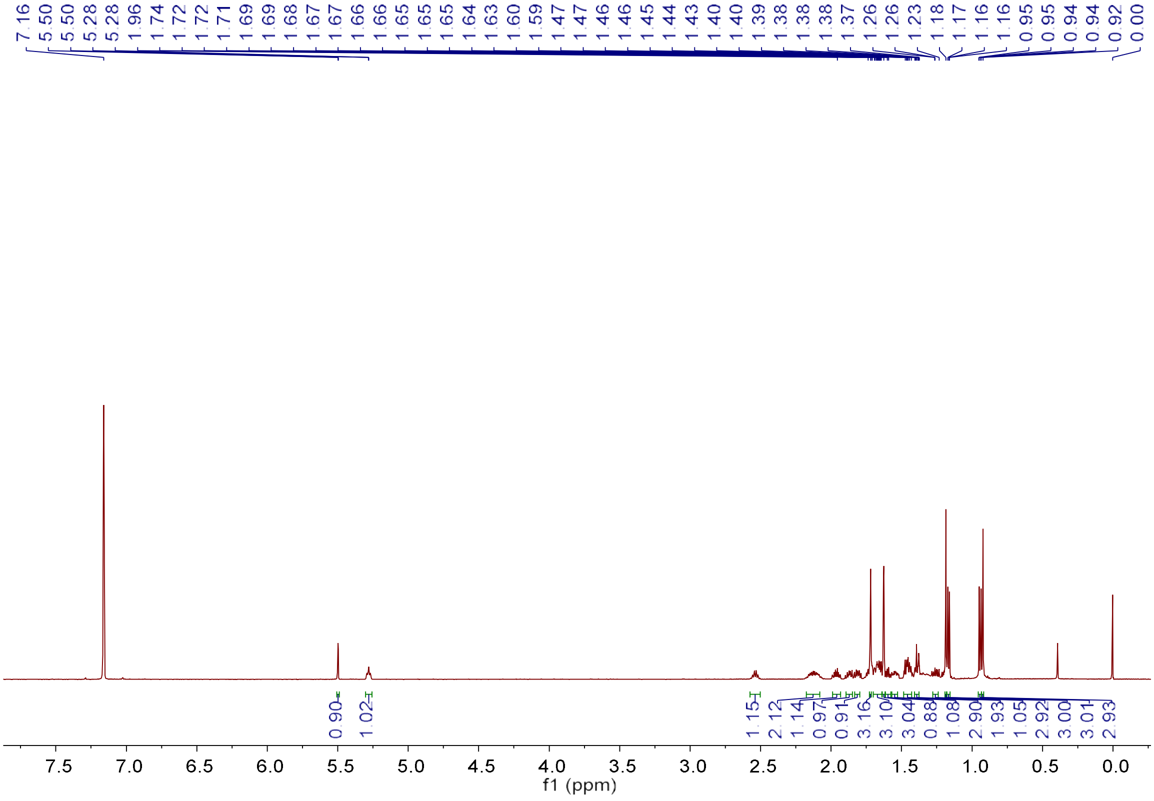

**Figure S11b.** $^{1}$H NMR spectrum of mangicdiene (**8**) (600 MHz, Benzene-$d_6$).

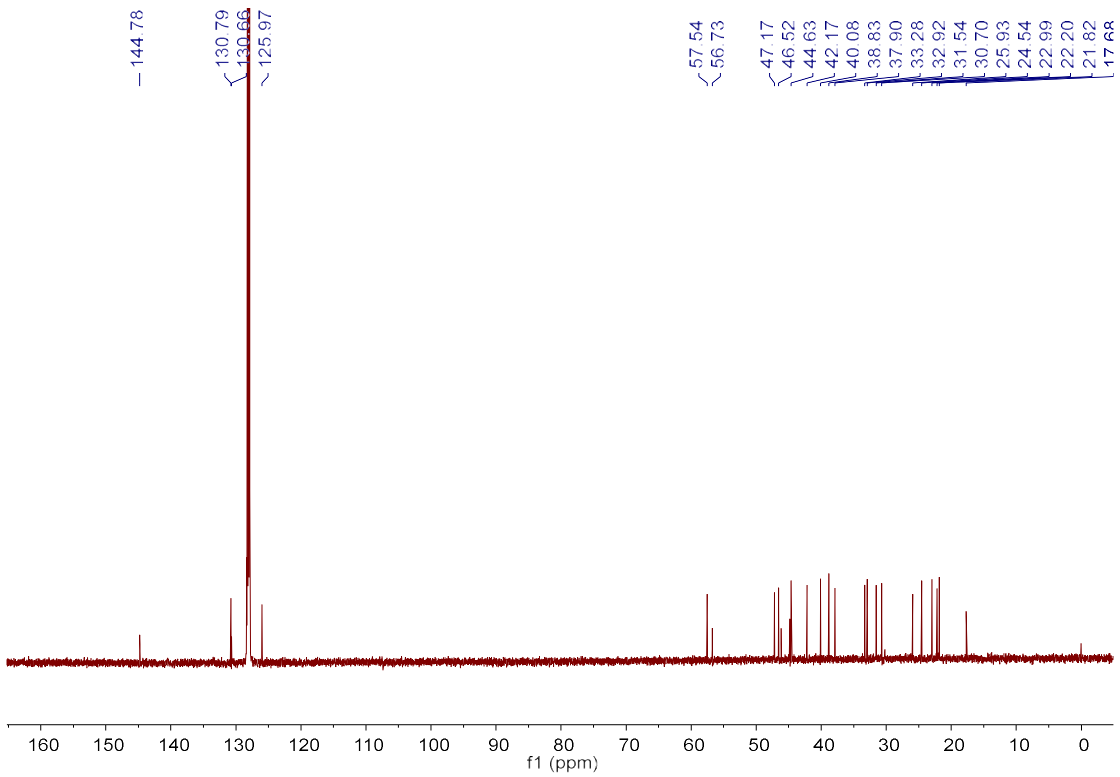

**Figure S11c.** $^{13}$C NMR spectrum of mangicdiene (**8**) (150 MHz, Benzene-$d_6$).

**Figure S12.** HR-EI-MS and NMR spectrum of spiroviolene (**15**).

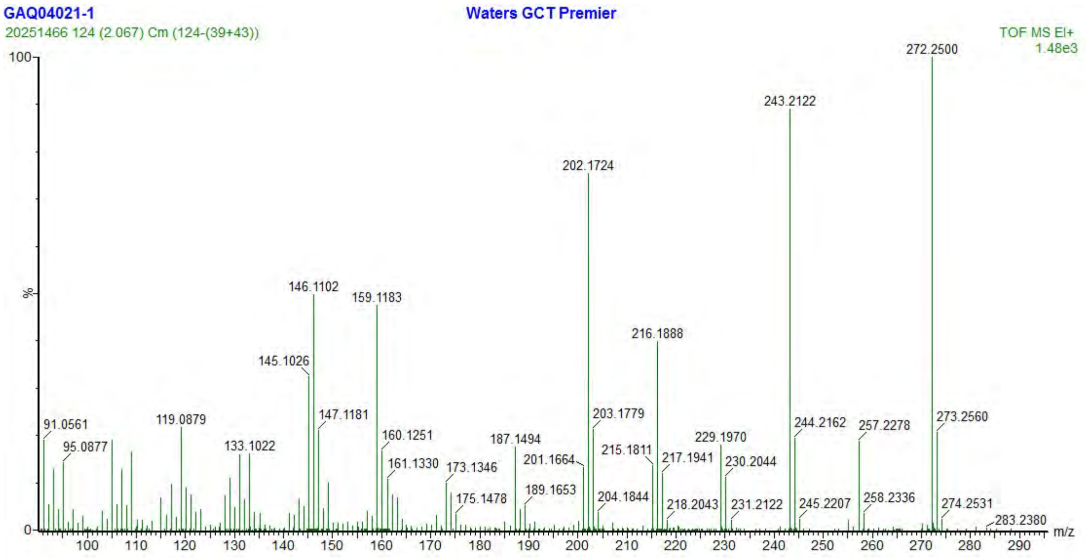

**Figure S12a.** HR-EI-MS spectrum of spiroviolene (**15**).

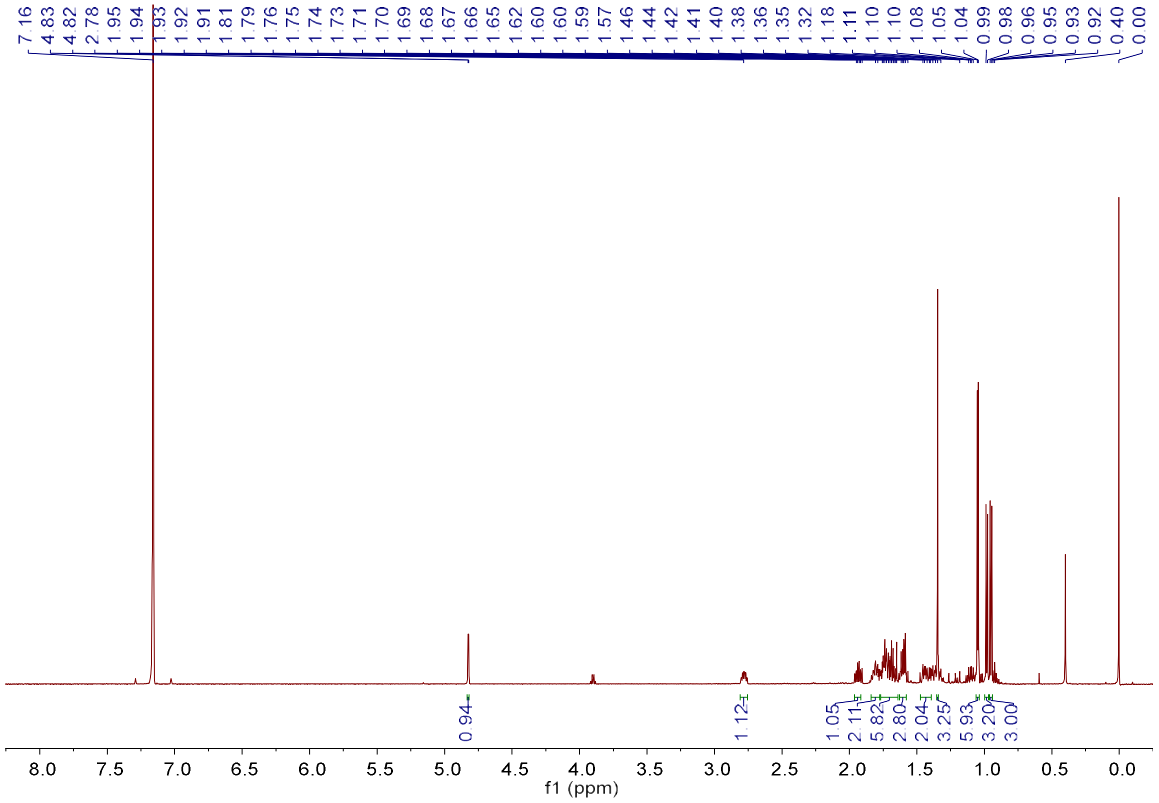

**Figure S12b.** $^{1}H$ NMR spectrum of spiroviolene (**15**) (600 MHz, Benzene-$d_6$).

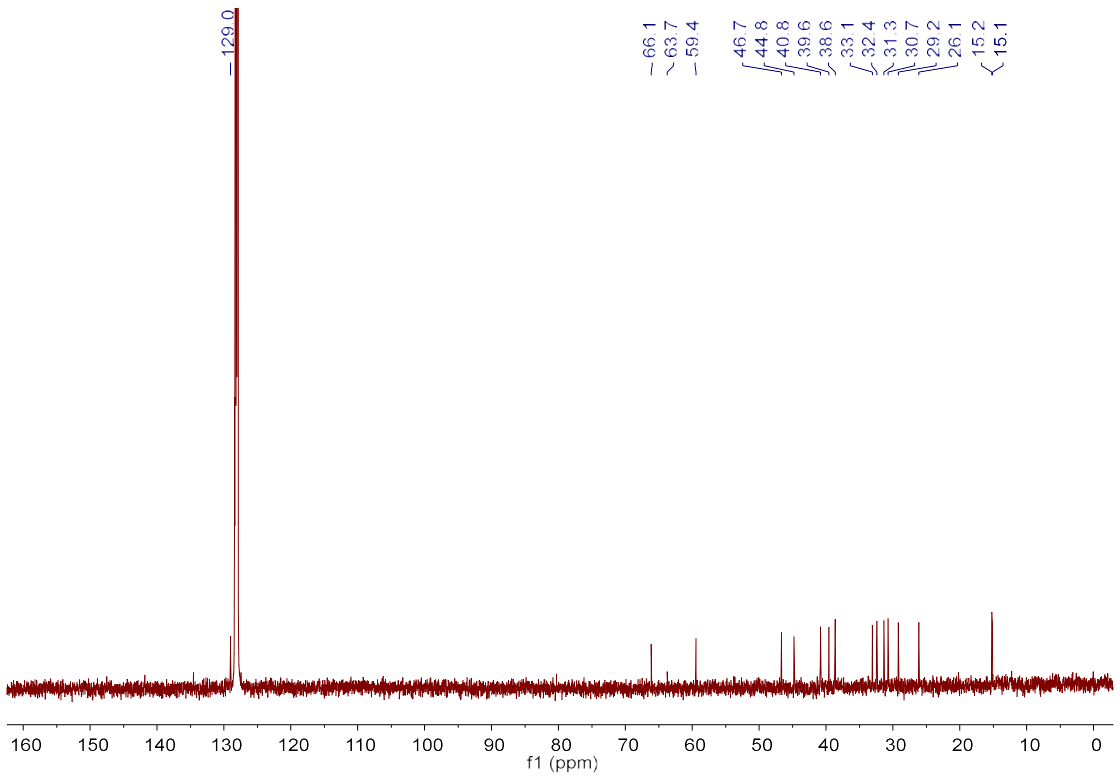

**Figure S12c.** $^{13}C$ NMR spectrum of spiroviolene (**15**) (150 MHz, Benzene-$d_6$).

Figure S13. HR-EI-MS and NMR spectrum of variediene (**16**).

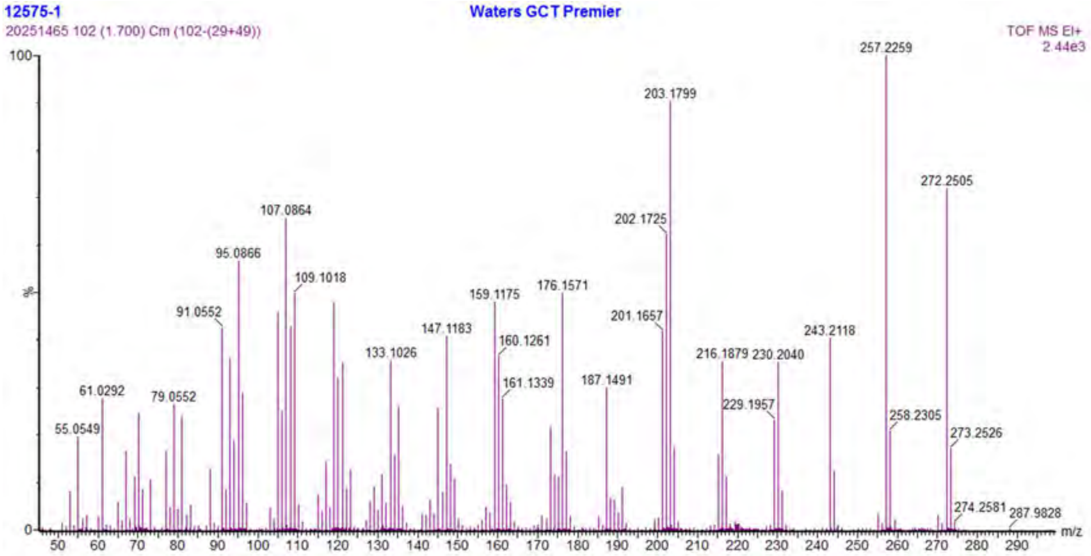

**Figure S13a.** HR-EI-MS spectrum of variediene (**16**).

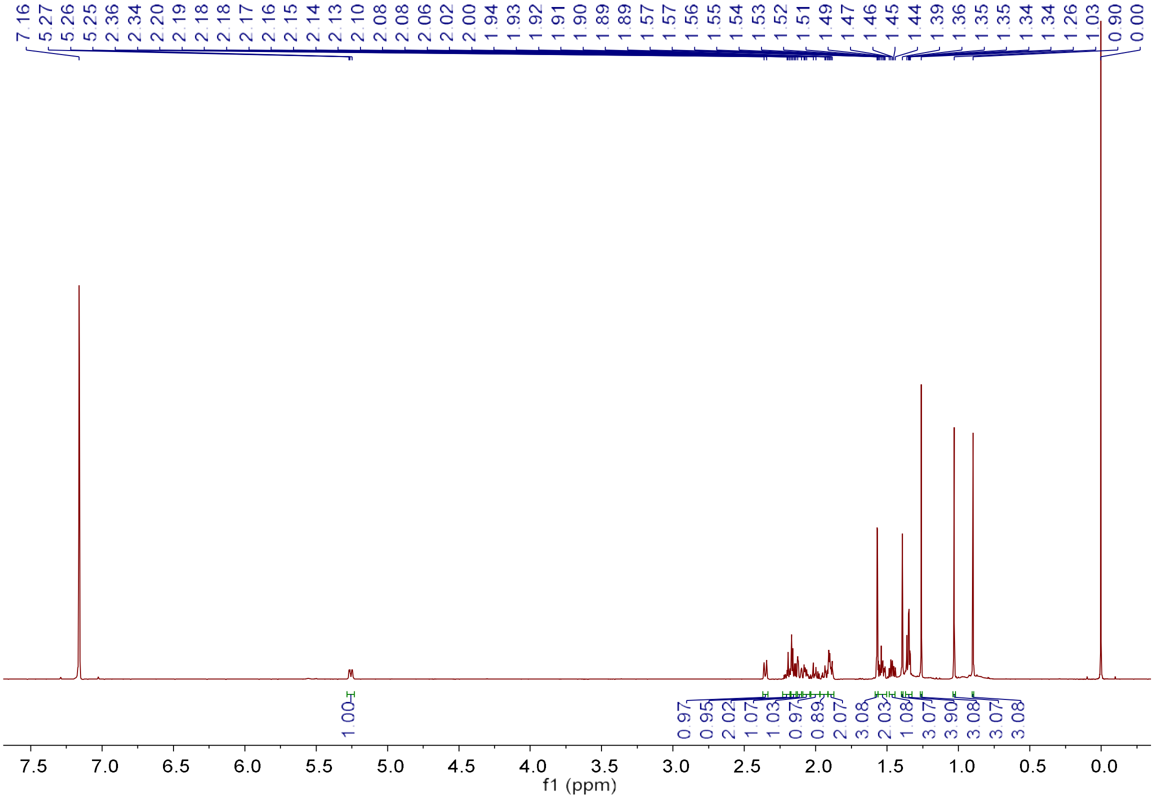

**Figure S13b.** [1]H NMR spectrum of variediene (**16**) (600 MHz, Benzene-$d_6$).

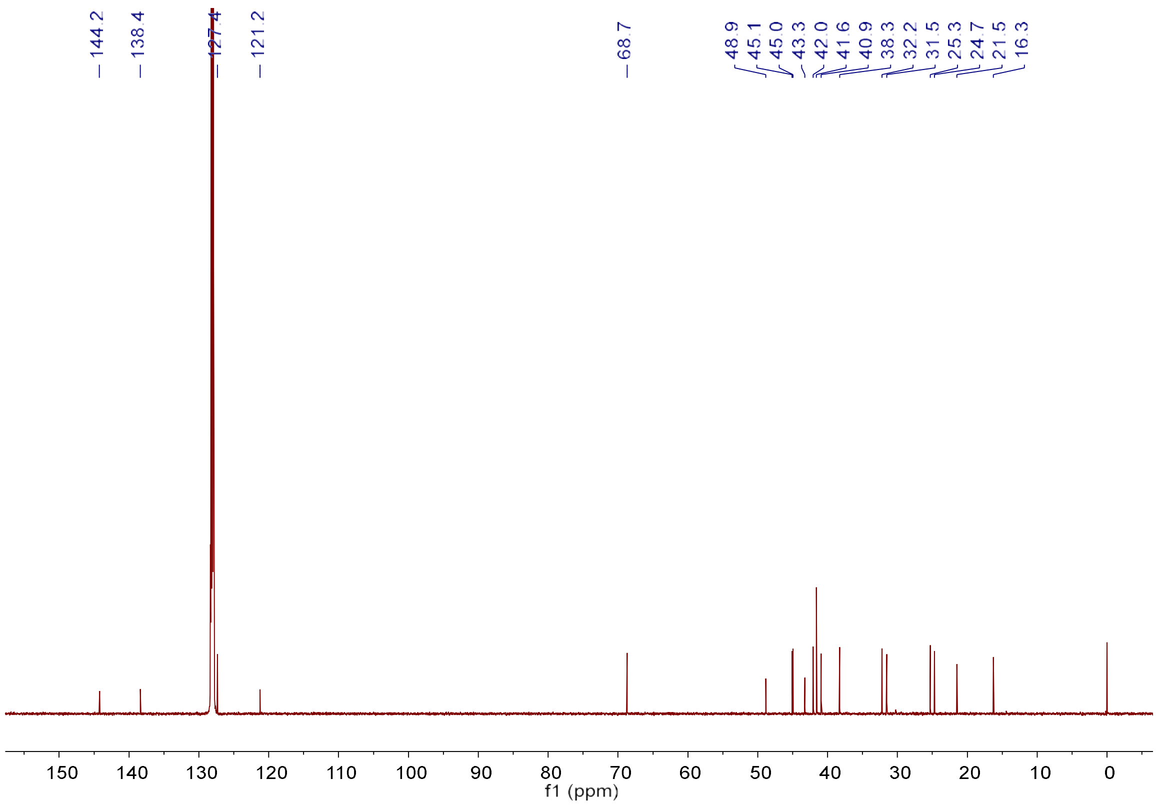

**Figure S13c.** $^{13}$C NMR spectrum of variediene (**16**) (150 MHz, Benzene-$d_6$).

**Figure S14.** HR-EI-MS and NMR spectrum of **17**.

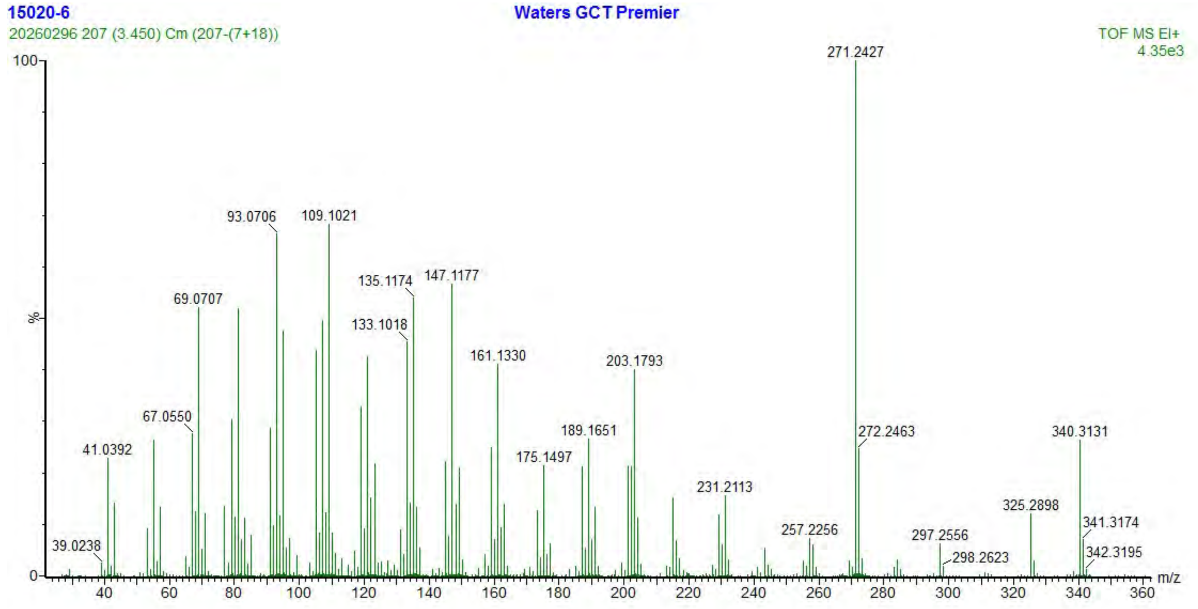

**Figure S14a.** HR-EI-MS spectrum of **17**.

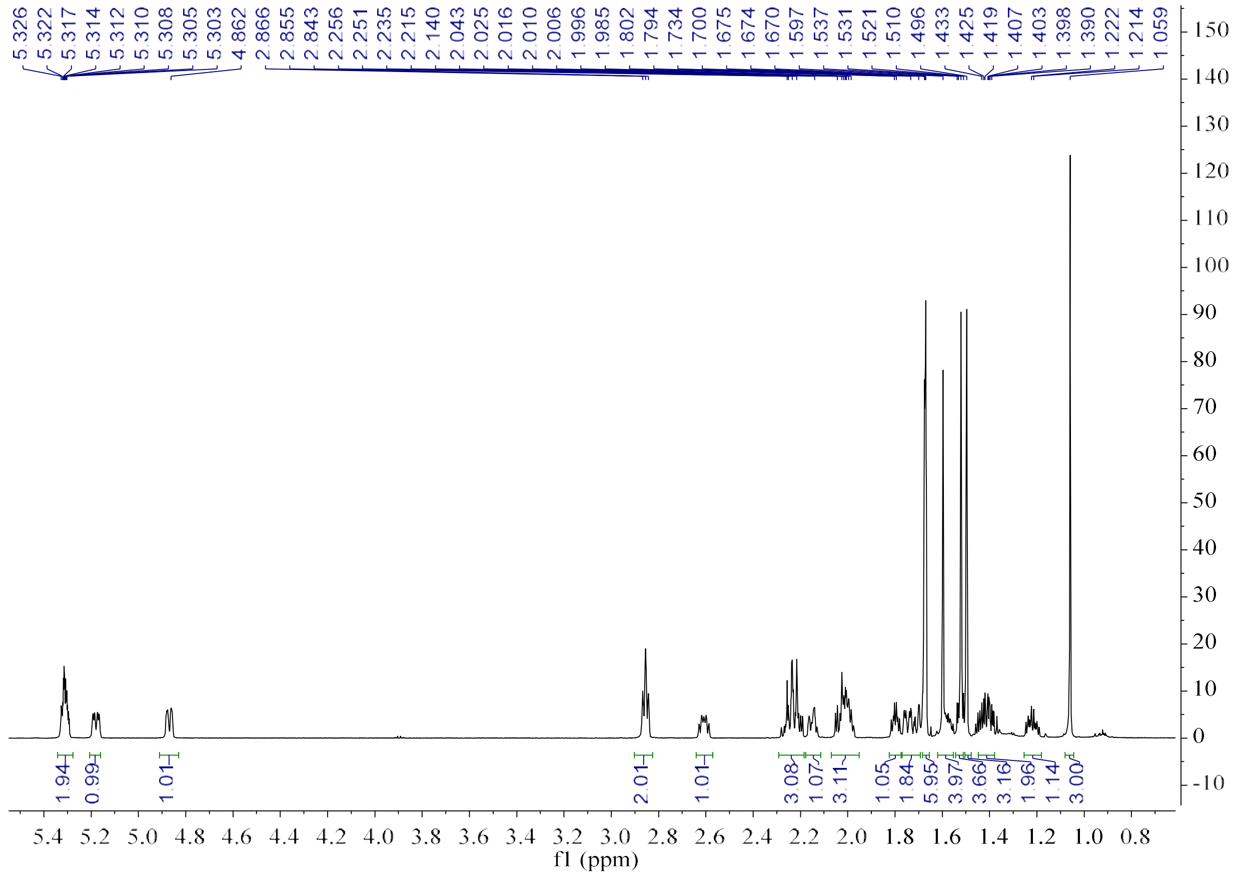

**Figure S14b.** [1]H NMR spectrum of **17** (600 MHz, Benzene-$d_6$).

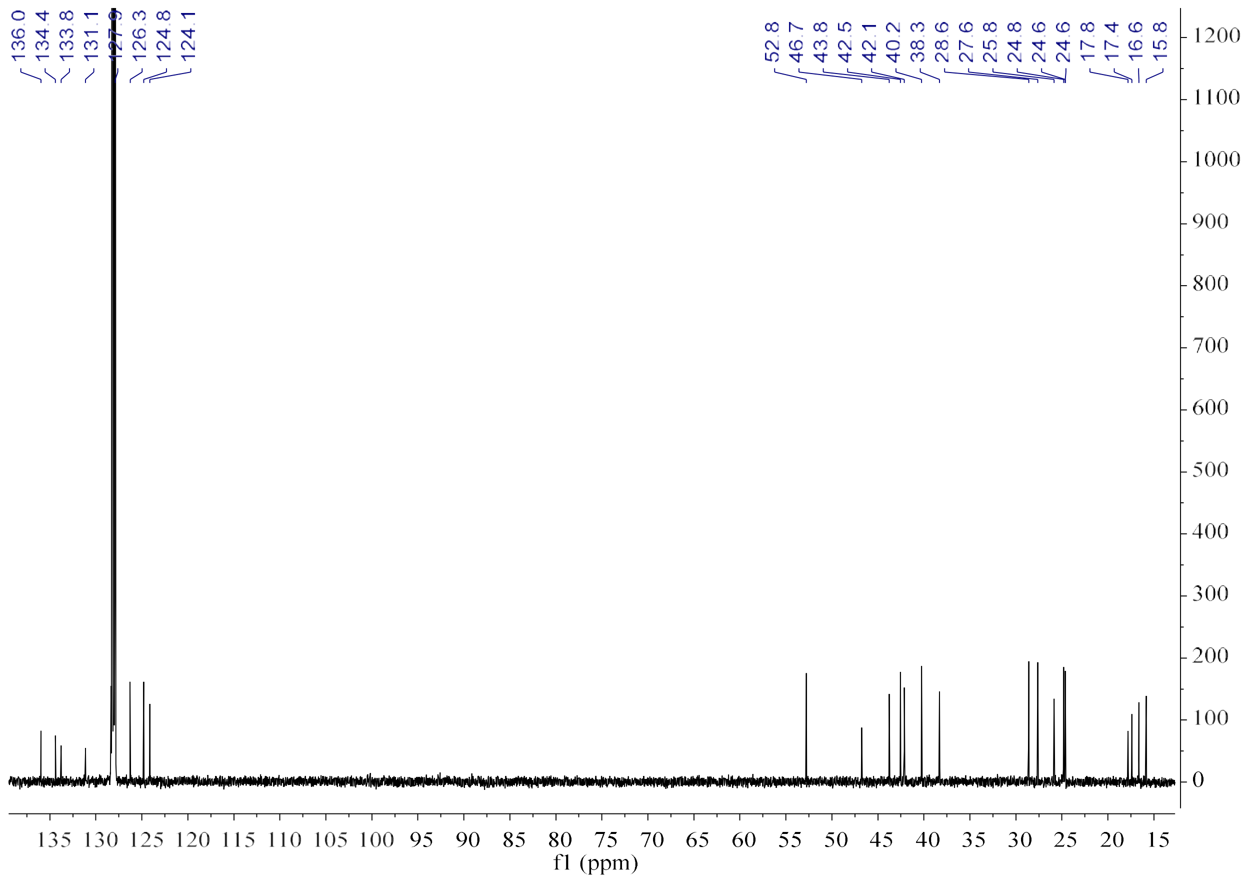

**Figure S14c.** $^{13}$C NMR spectrum of **17** (150 MHz, Benzene-$d_6$).

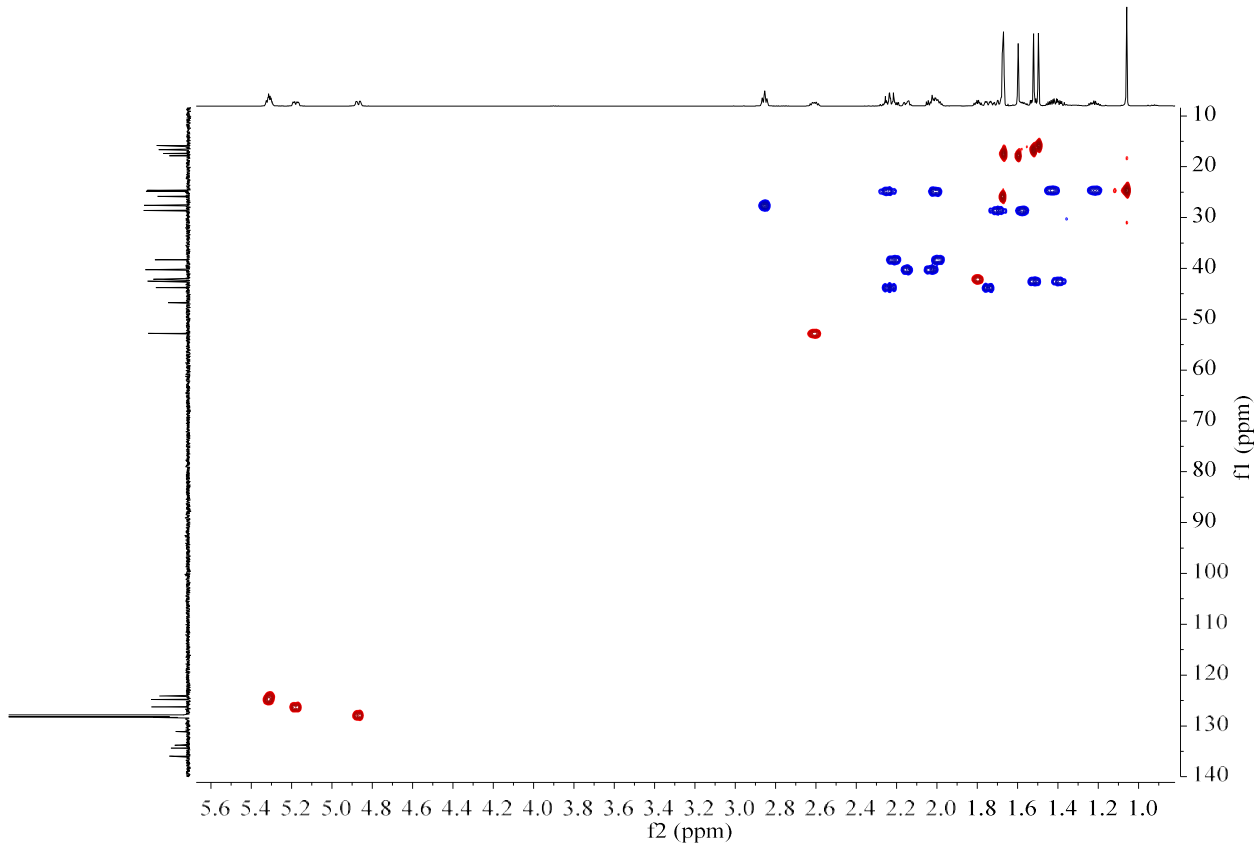

**Figure S14d.** HSQC spectrum of **17** (150 MHz/600 MHz, Benzene-$d_6$).

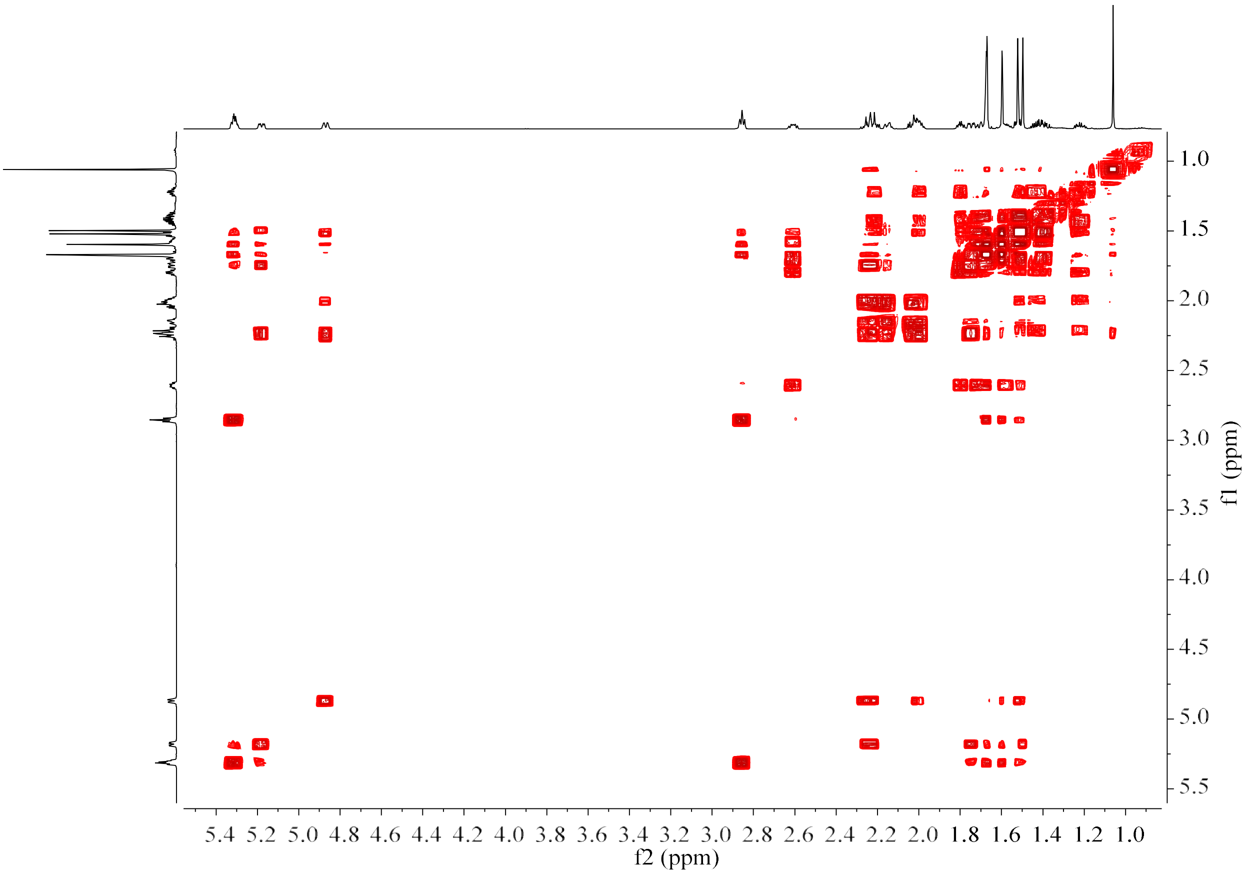

**Figure S14e.** $^{1}H$–$^{1}H$ COSY spectrum of **17** (600 MHz, Benzene-$d_6$).

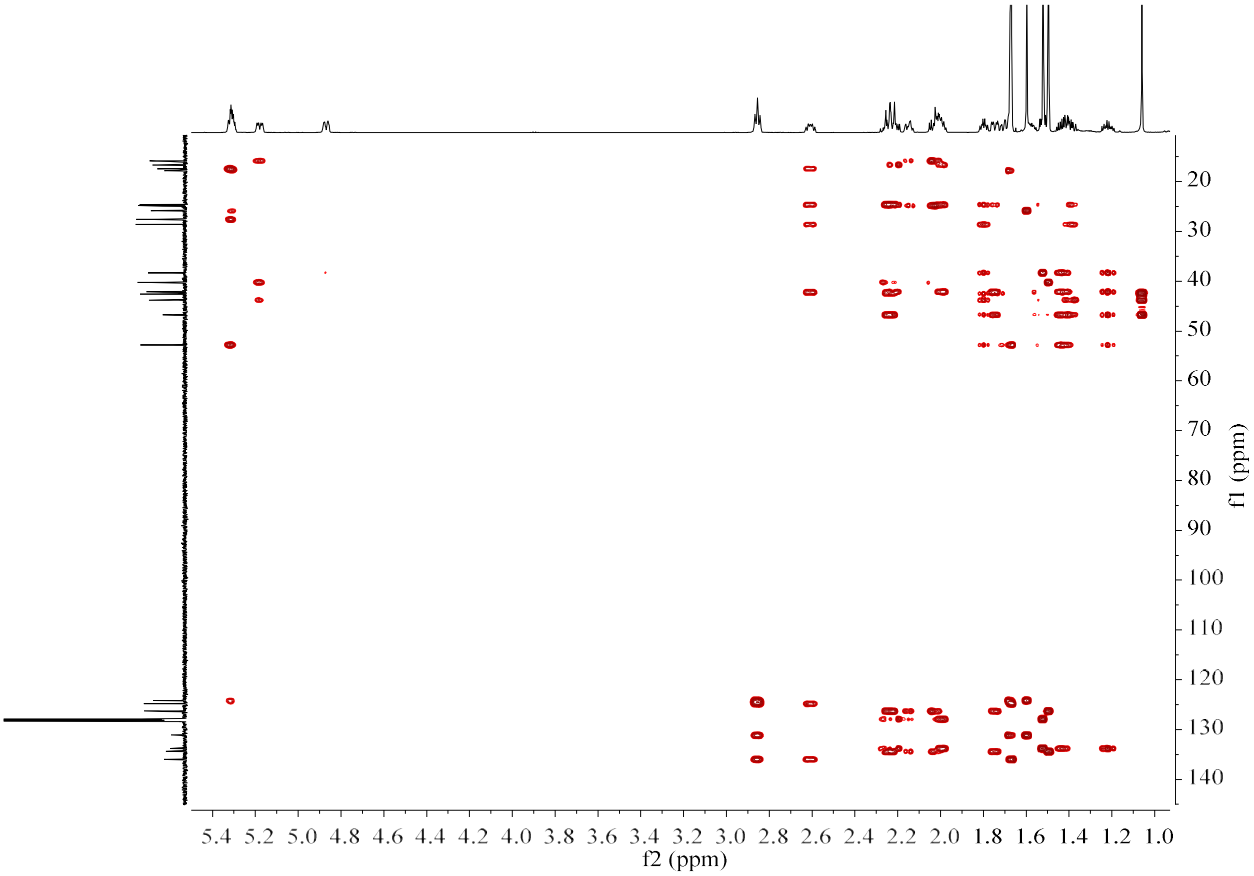

**Figure S14f.** HMBC spectrum of **17** (150 MHz/600 MHz, Benzene-$d_6$).

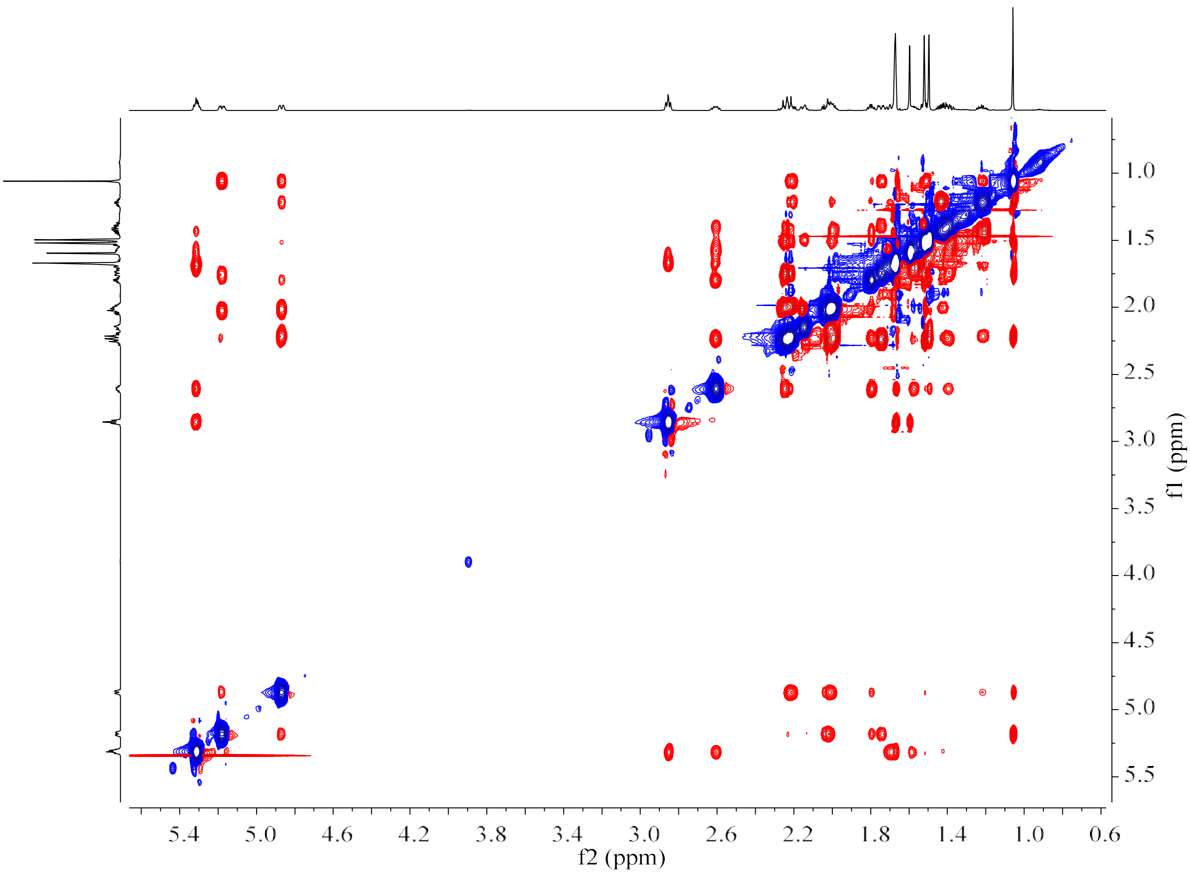

**Figure S14g.** NOESY spectrum of **17** (600 MHz, Benzene-$d_6$).

**Figure S15.** HR-EI-MS and NMR spectrum of **18**.

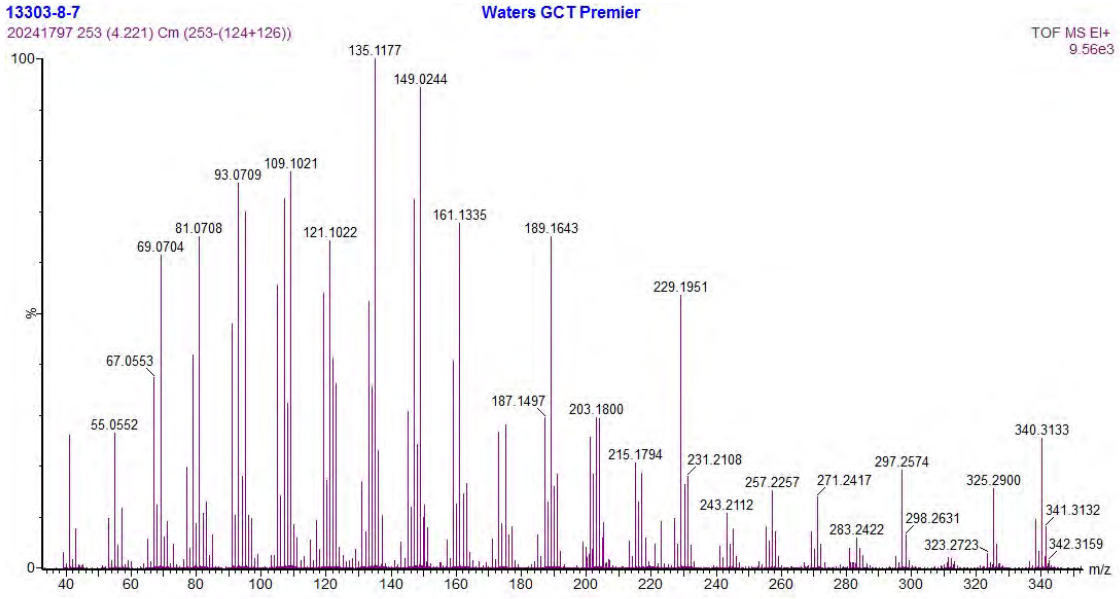

**Figure S15a.** HR-EI-MS spectrum of **18**.

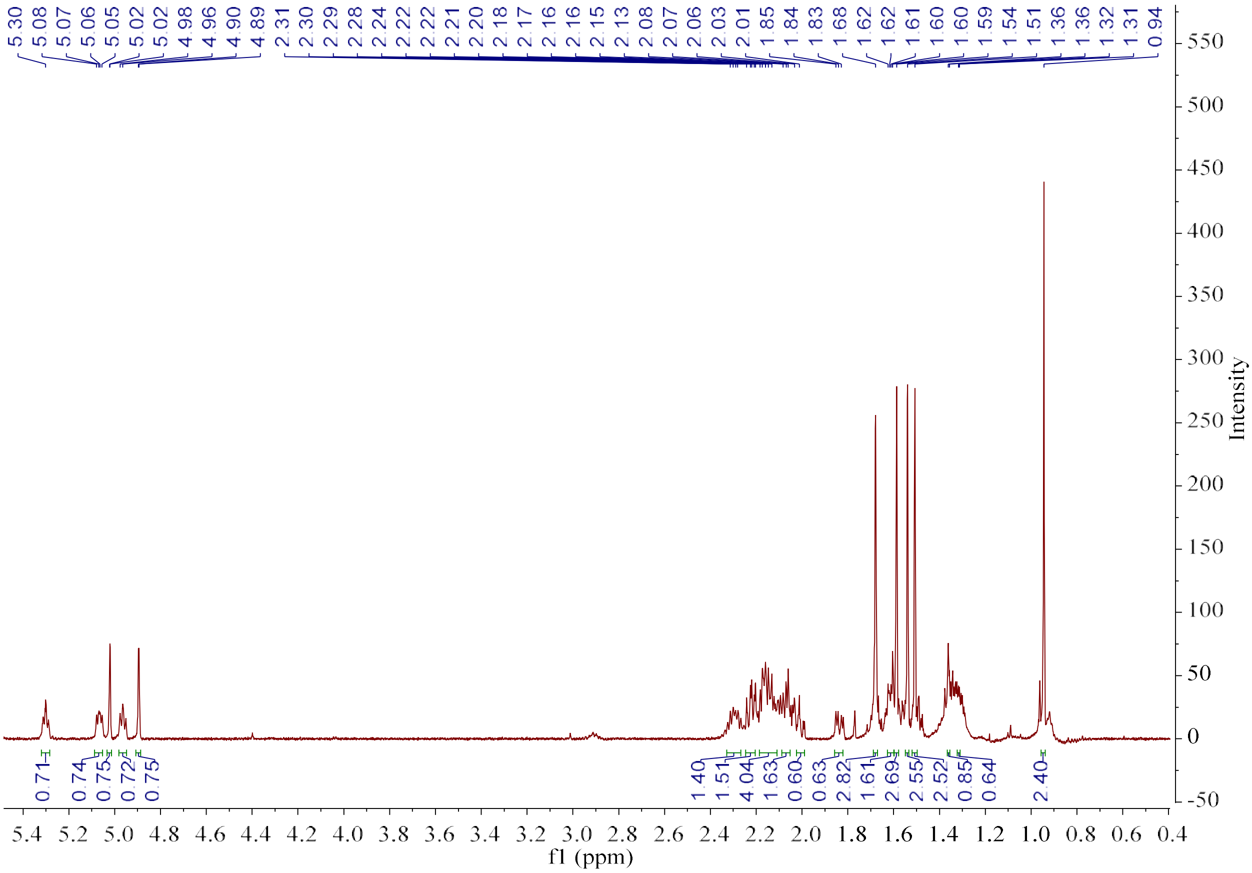

**Figure S15b.** $^{1}$H NMR spectrum of **18** (600 MHz, Benzene-$d_6$).

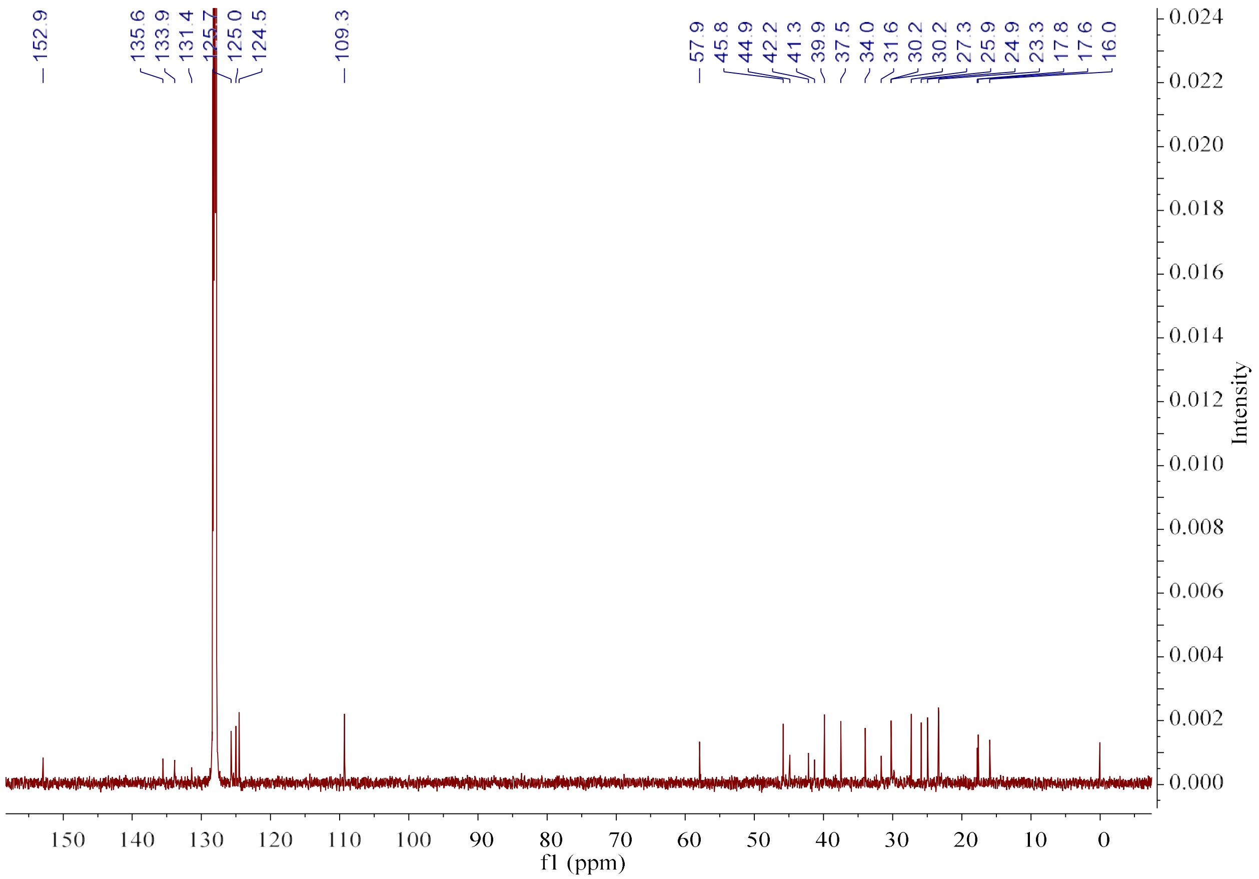

**Figure S15c.** $^{13}$C NMR spectrum of **18** (150 MHz, Benzene-*d*$_6$).

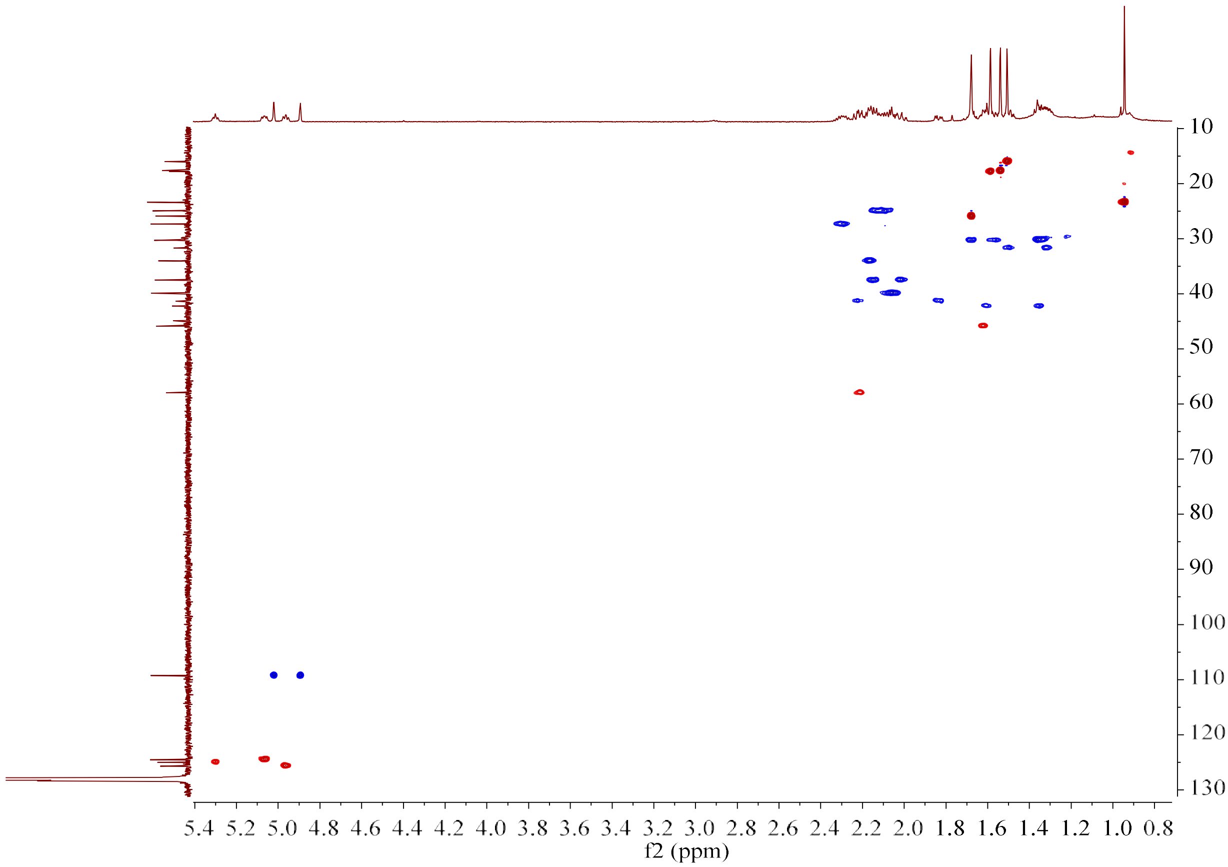

**Figure S15d.** HSQC spectrum of **18** (150 MHz/600 MHz, Benzene-*d*$_6$).

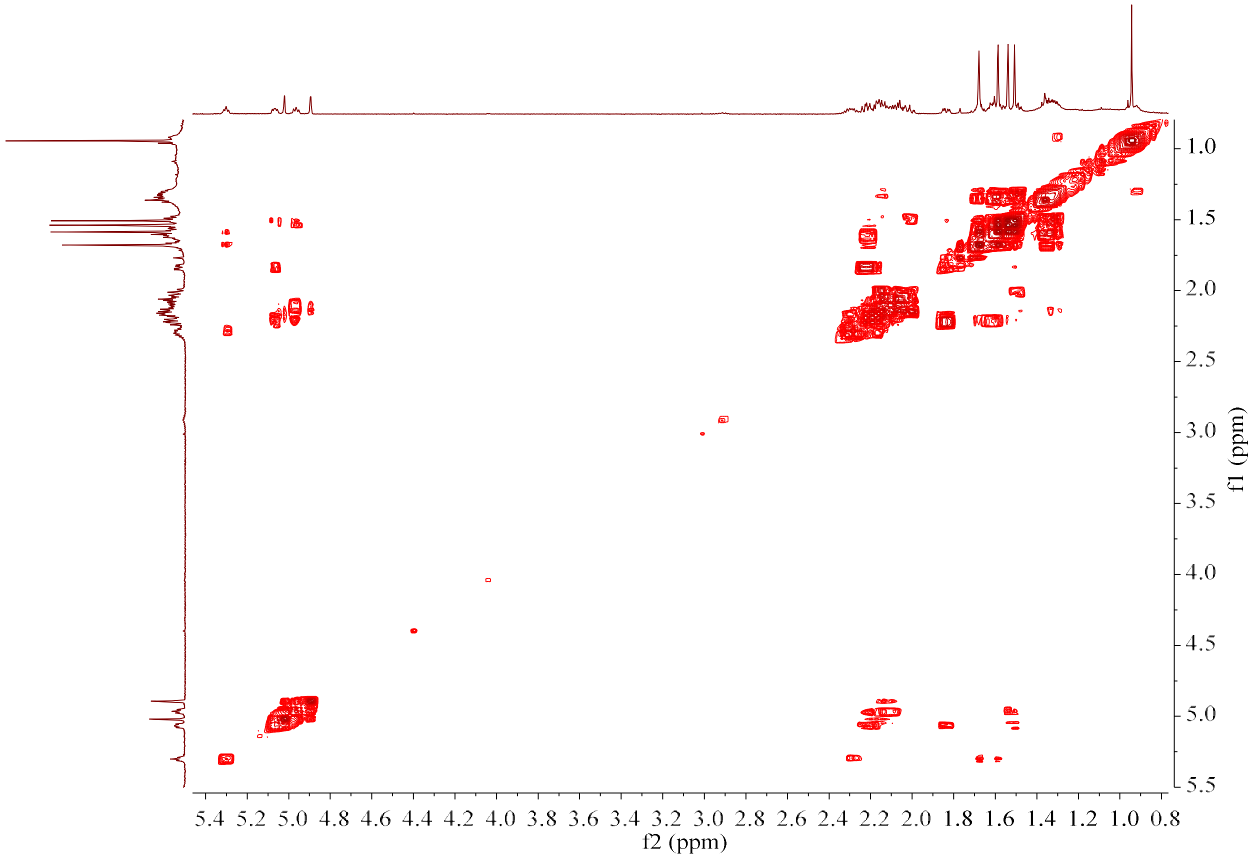

**Figure S15e.** $^{1}H$–$^{1}H$ COSY spectrum of **18** (600 MHz, Benzene-$d_6$).

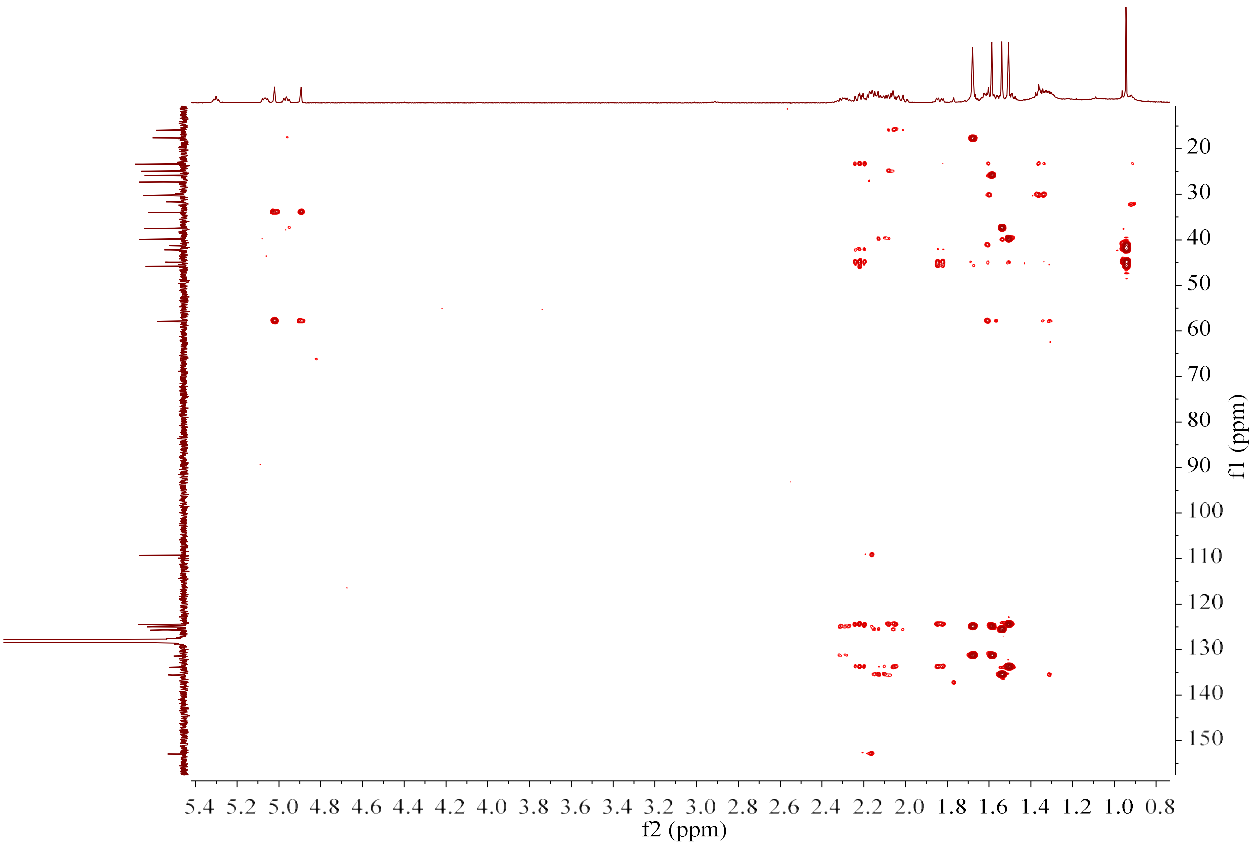

**Figure S15f.** HMBC spectrum of **18** (150 MHz/600 MHz, Benzene-$d_6$).

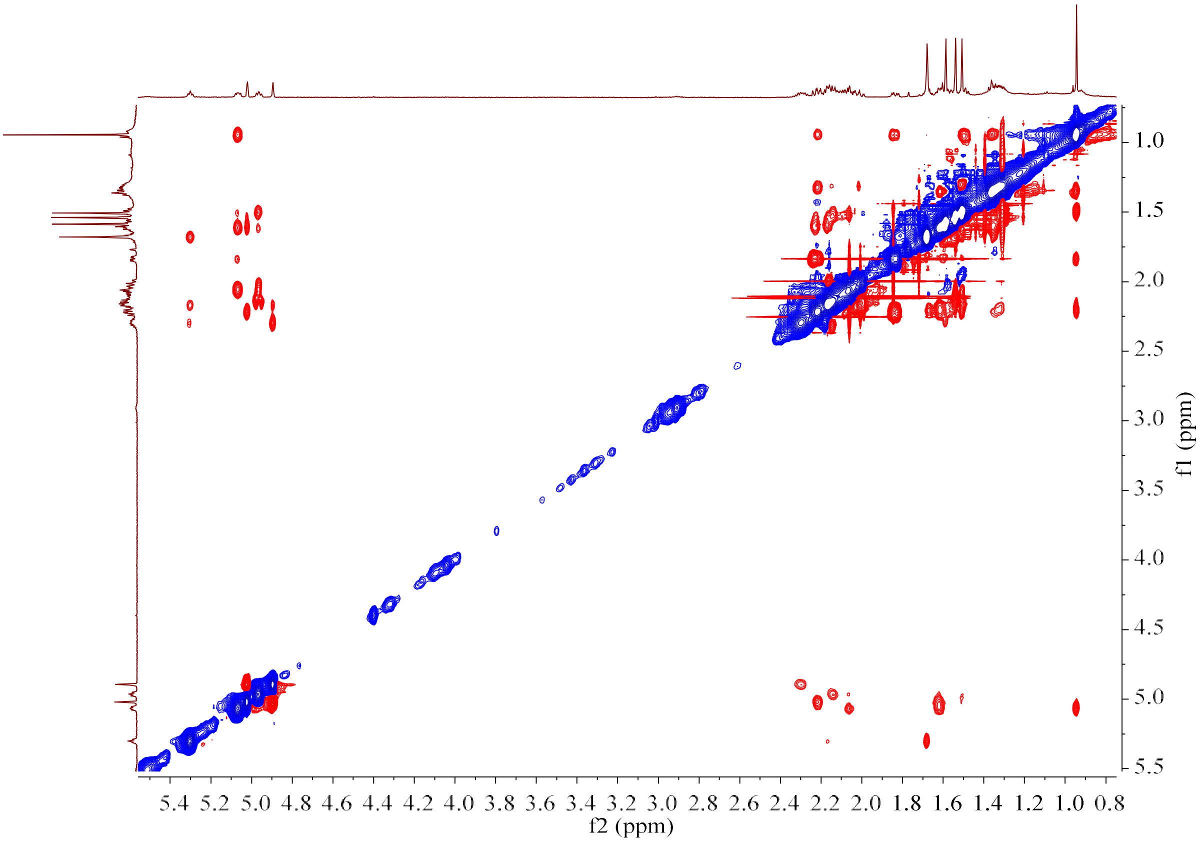

**Figure S15g.** NOESY spectrum of **18** (600 MHz, Benzene-$d_6$).

**Figure S16.** HR-EI-MS and NMR spectrum of astelifatriene (**19**).

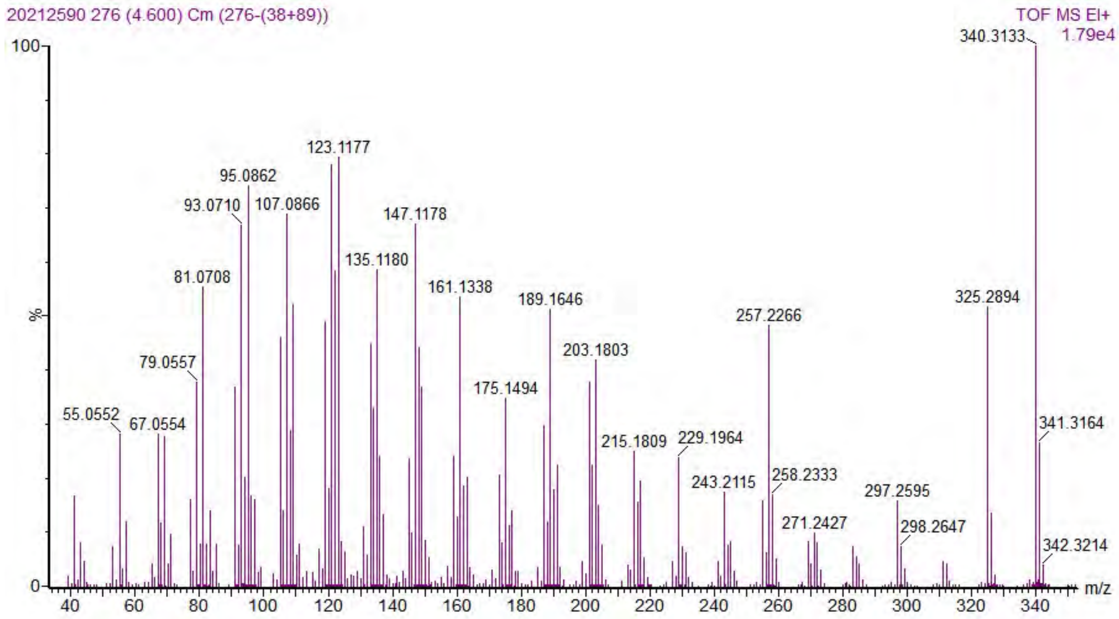

**Figure S16a.** HR-EI-MS spectrum of astelifatriene (**19**).

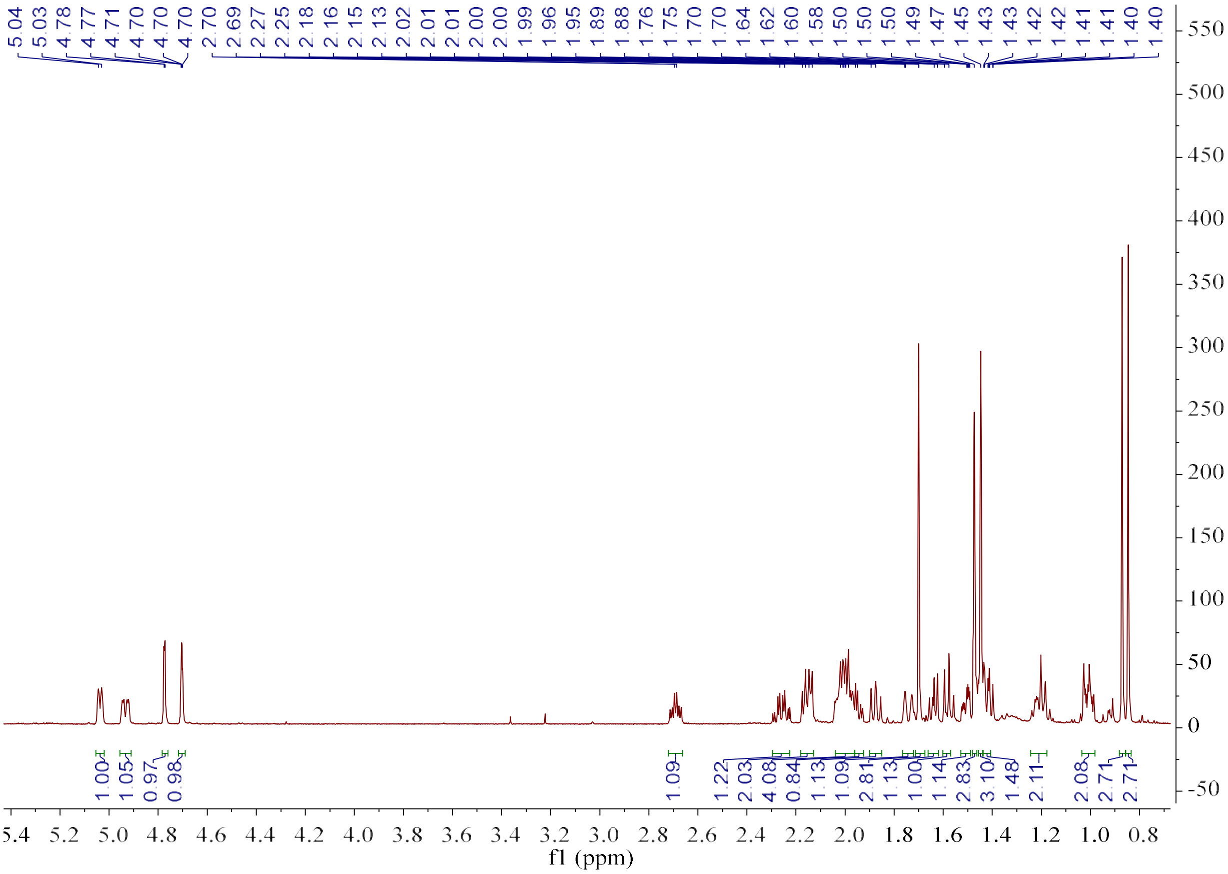

**Figure S16b.** [1]H NMR spectrum of astelifatriene (**19**). (600 MHz, Benzene-$d_6$).

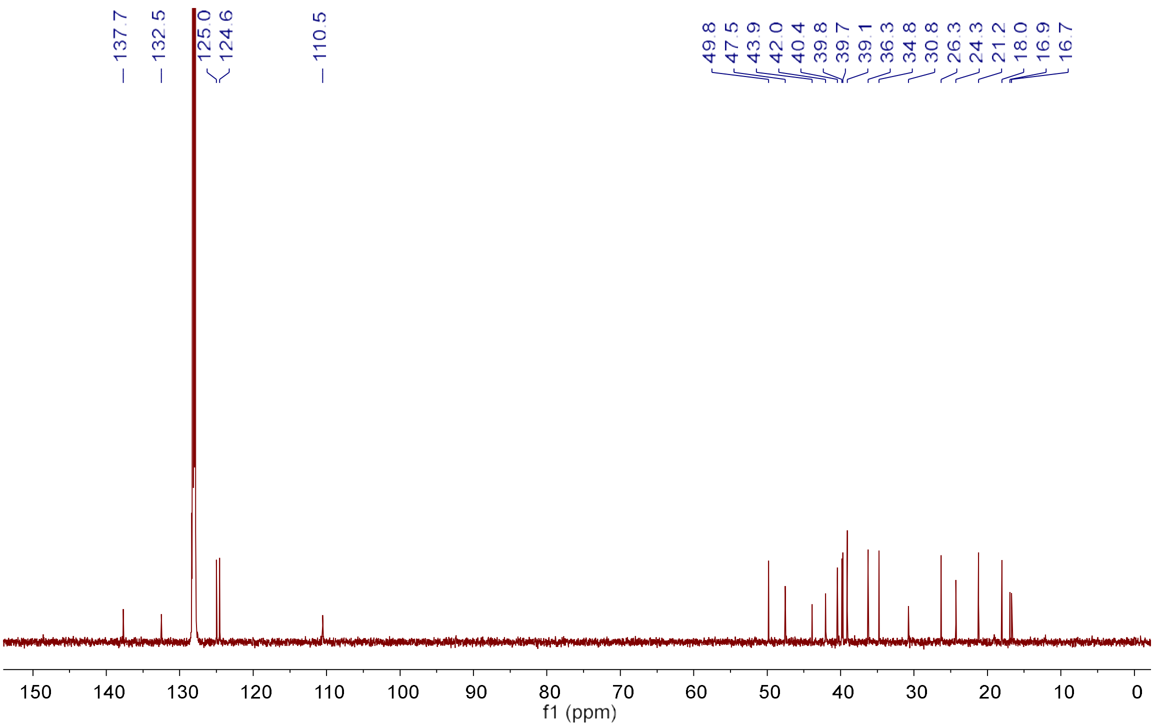

**Figure S16c.** $^{13}C$ NMR spectrum of astelifatriene (**19**) (150 MHz, Benzene-$d_6$).

**Figure S17.** HR-EI-MS and NMR spectrum of **20**.

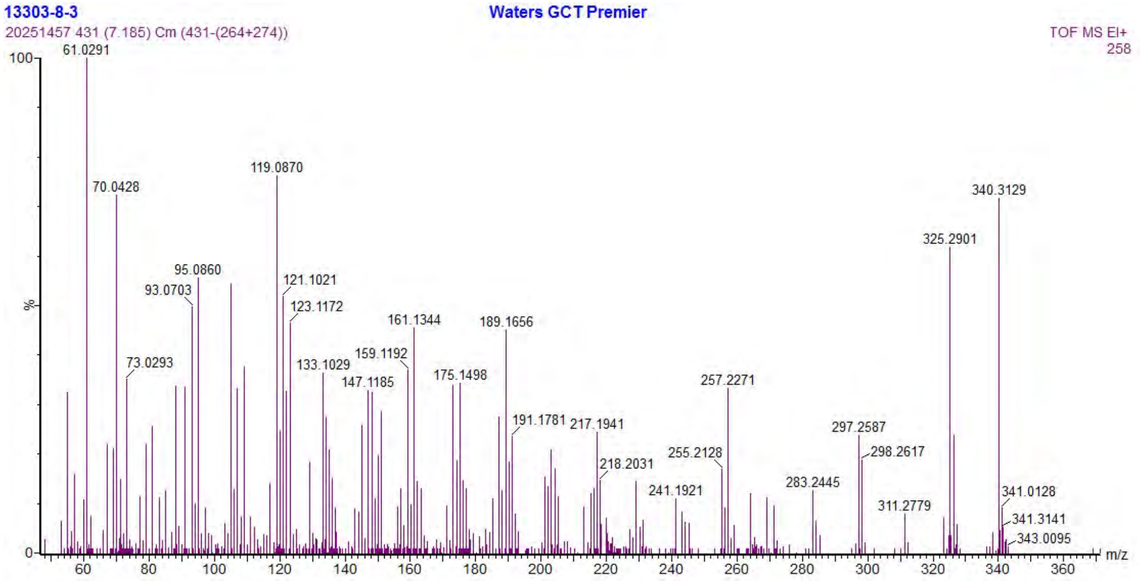

**Figure S17a.** HR-EI-MS spectrum of **20**.

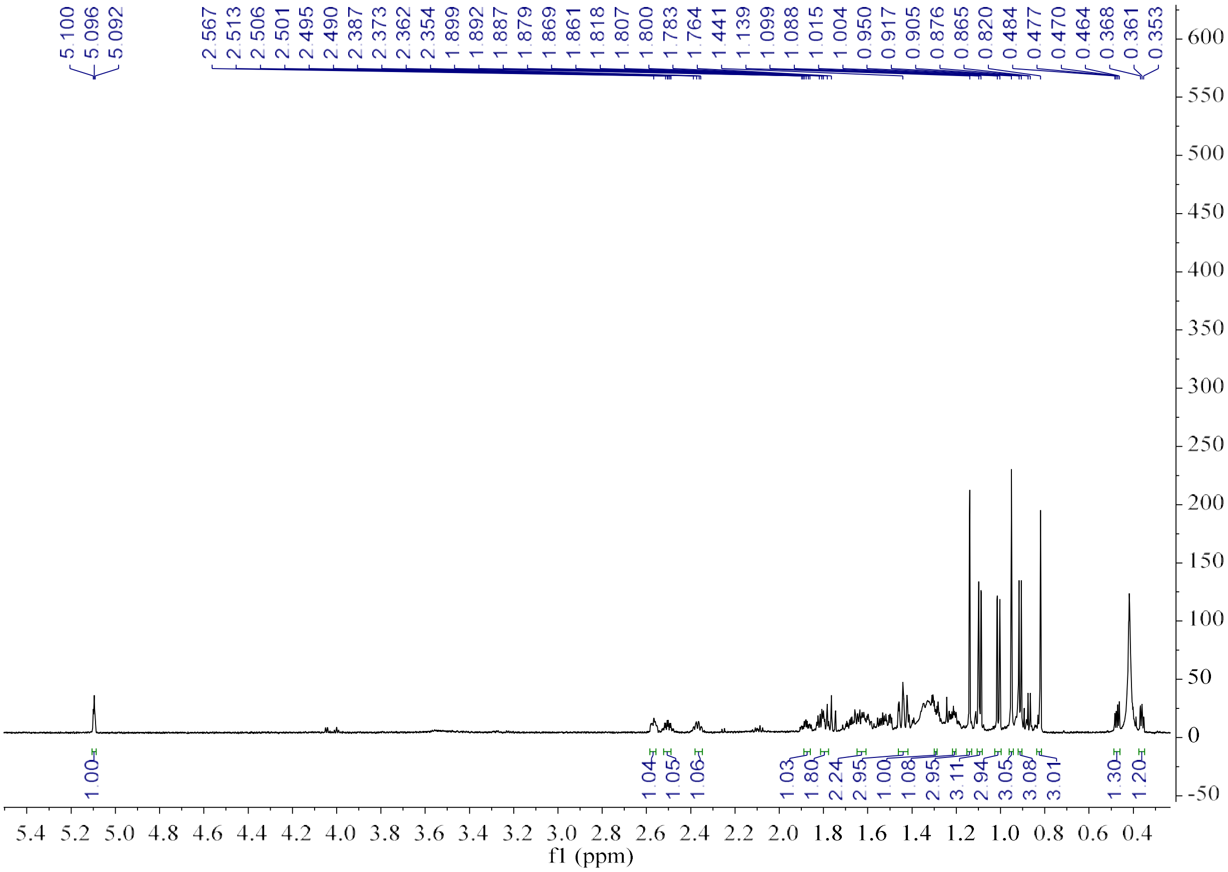

**Figure S17b.** $^{1}$H NMR spectrum of **20** (600 MHz, Benzene-$d_6$).

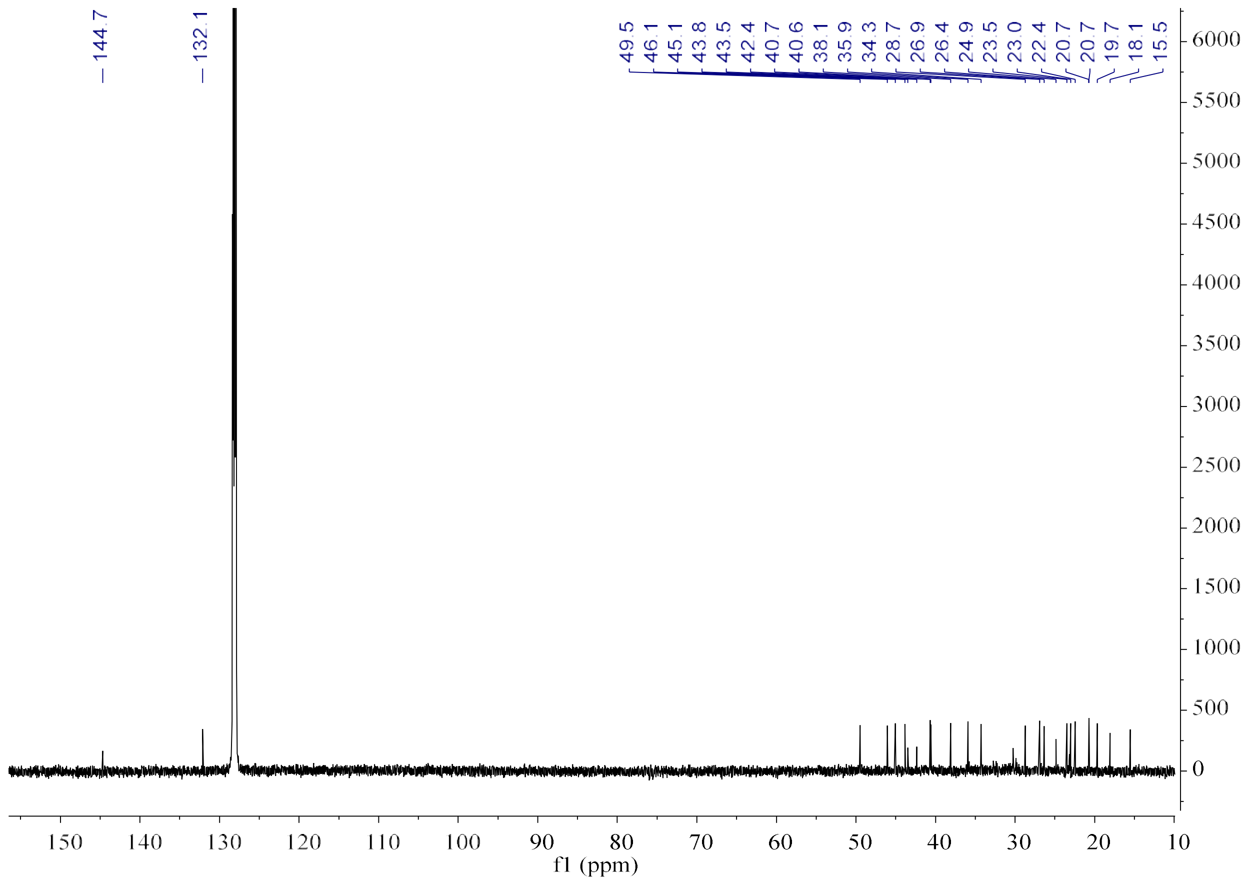

**Figure S17c.** $^{13}C$ NMR spectrum of **20** (150 MHz, Benzene-$d_6$).

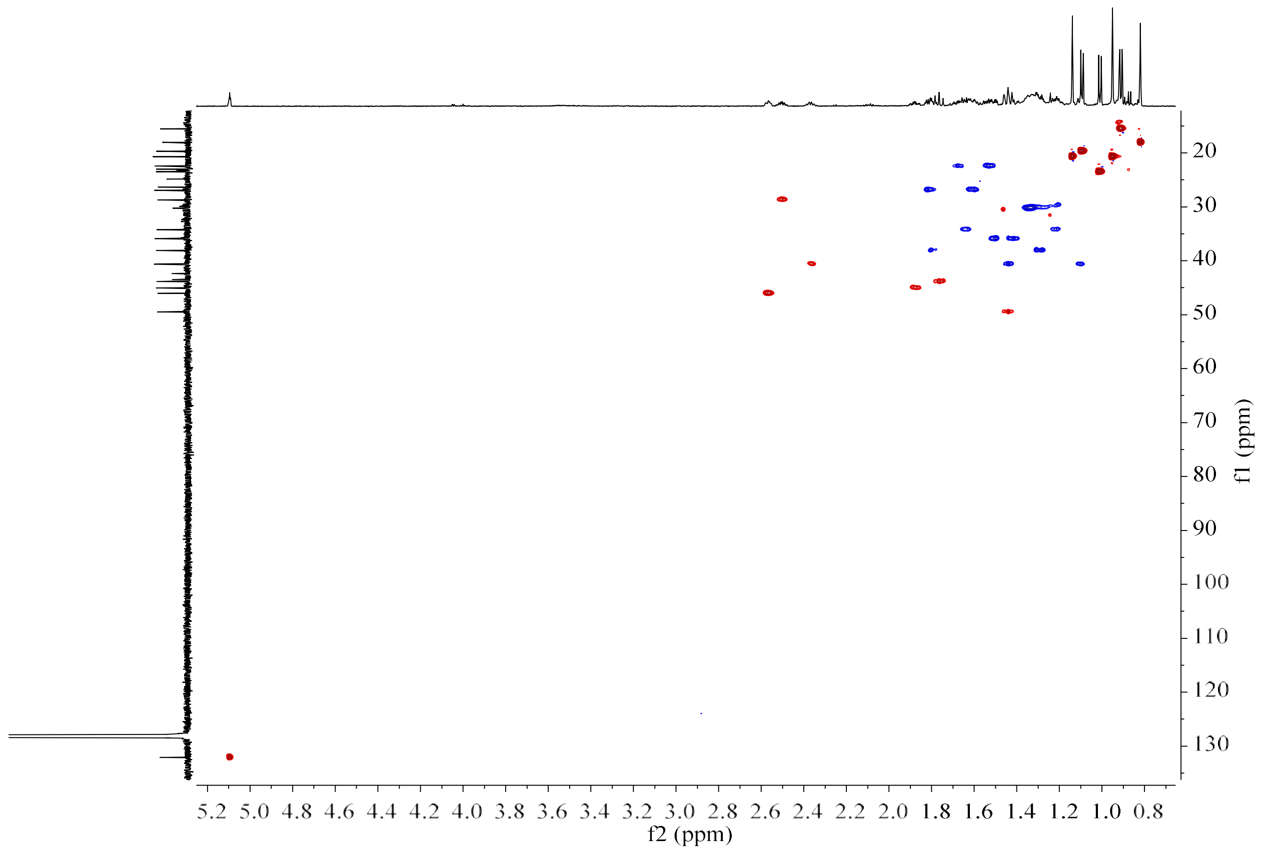

**Figure S17d.** HSQC spectrum of **20** (150 MHz/600 MHz, Benzene-$d_6$).

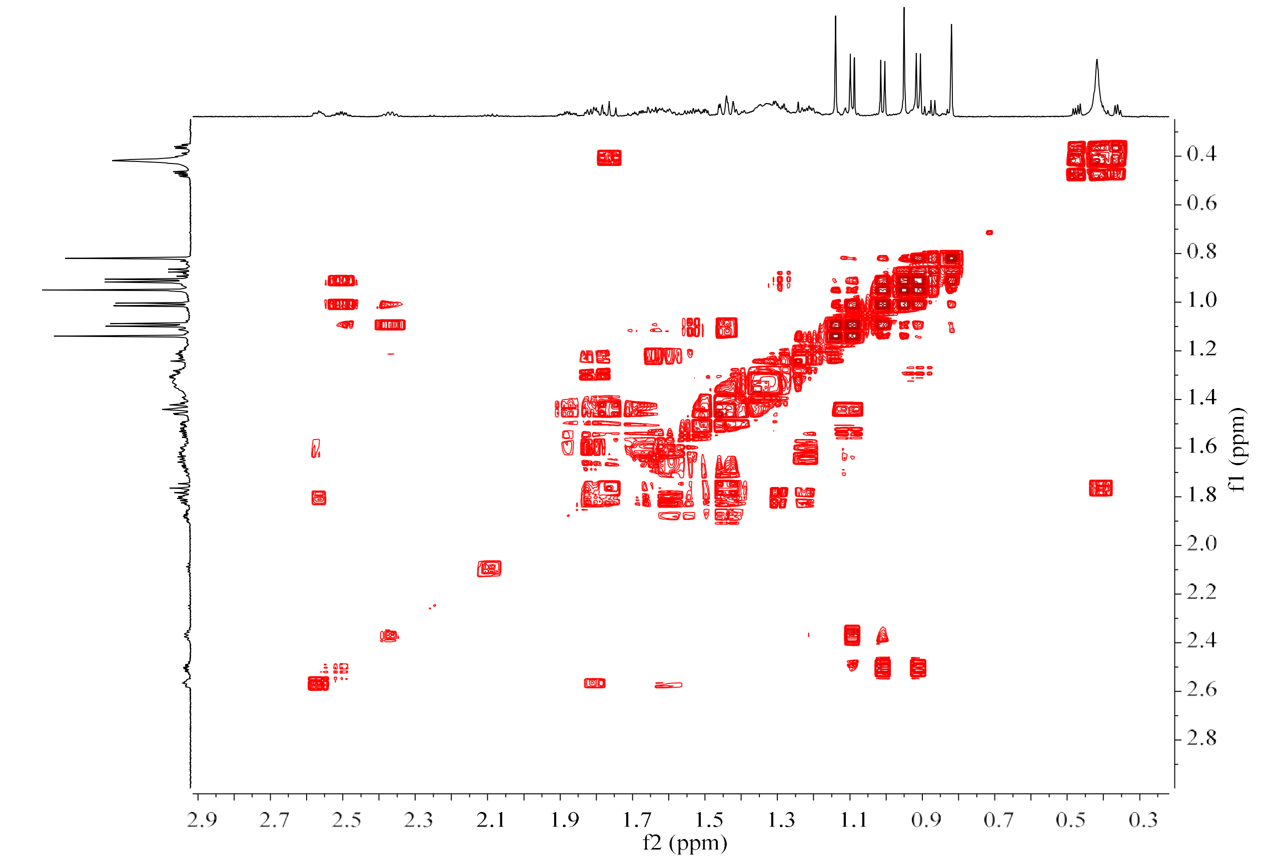

**Figure S17e.** $^1$H–$^1$H COSY spectrum of **20** (600 MHz, Benzene-$d_6$).

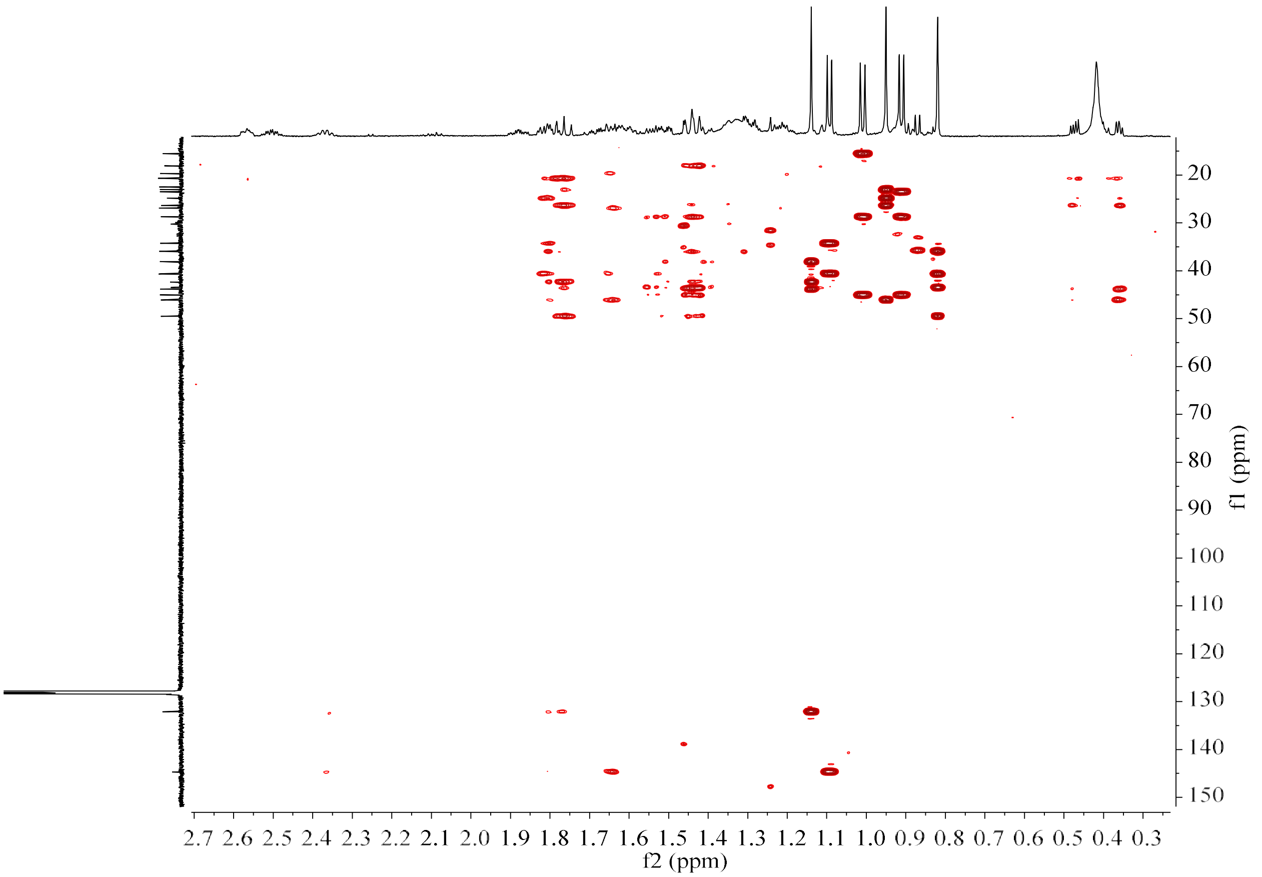

**Figure S17f.** HMBC spectrum of **20** (150 MHz/600 MHz, Benzene-$d_6$).

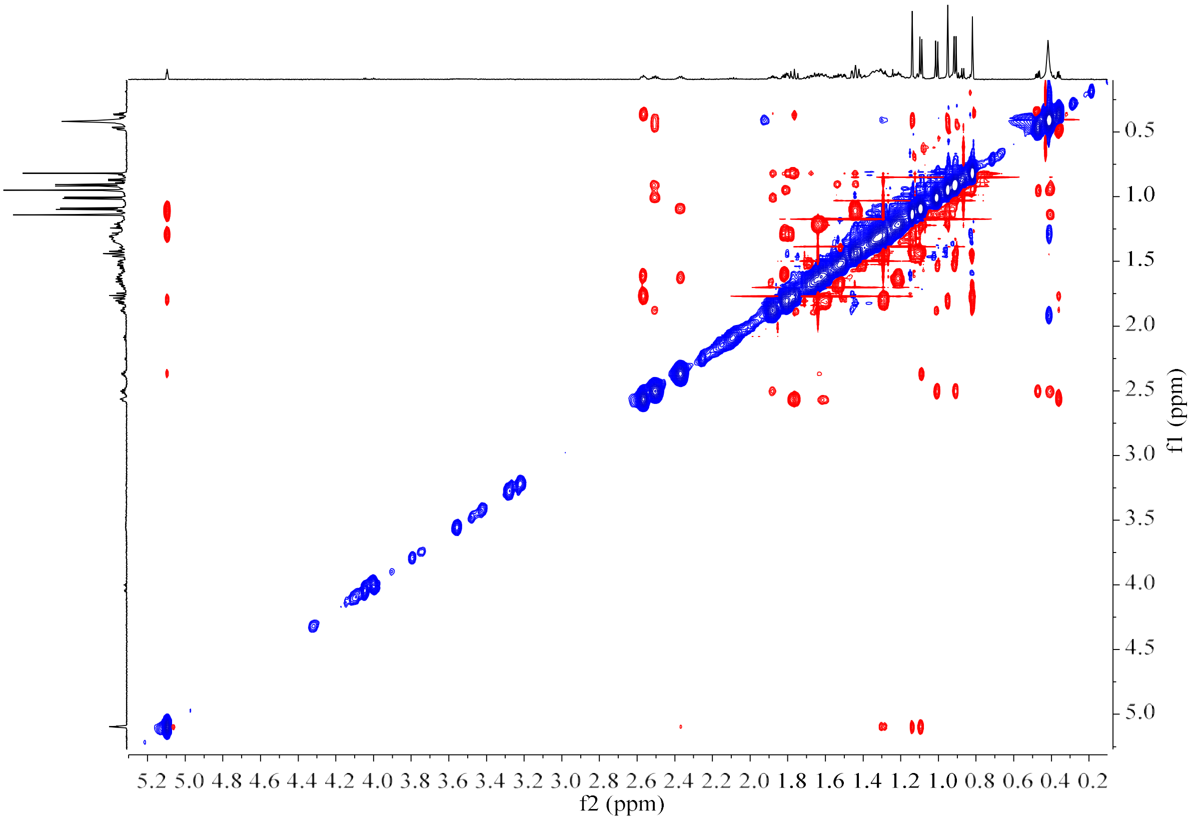

**Figure S17g.** NOESY spectrum of **20** (600 MHz, Benzene-$d_6$).

**Figure S18.** HR-EI-MS and NMR spectrum of **21**.

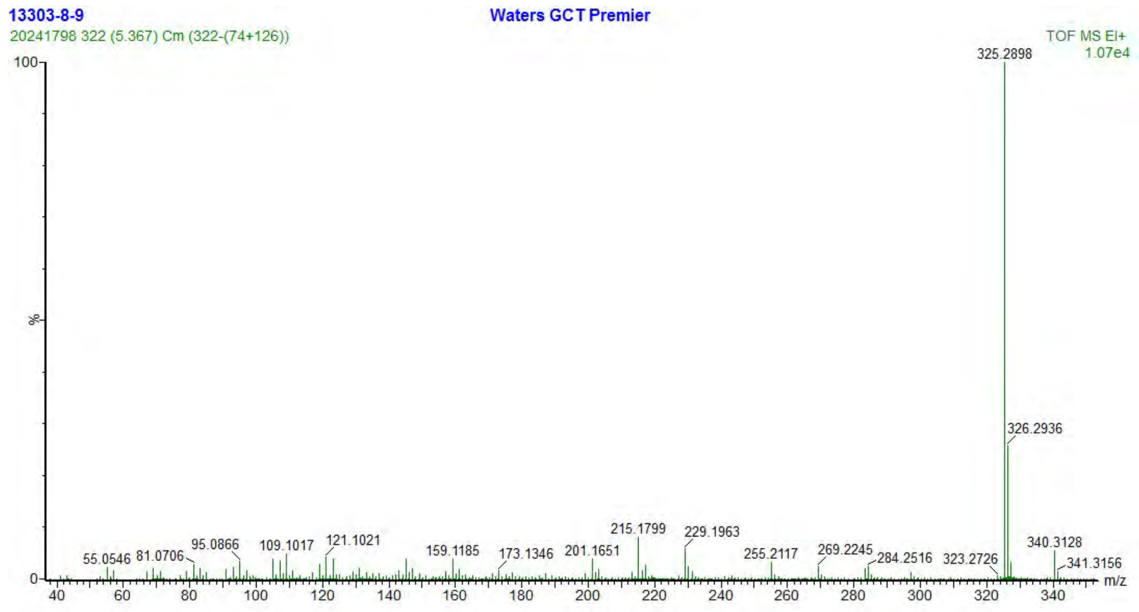

**Figure S18a.** HR-EI-MS spectrum of **21**.

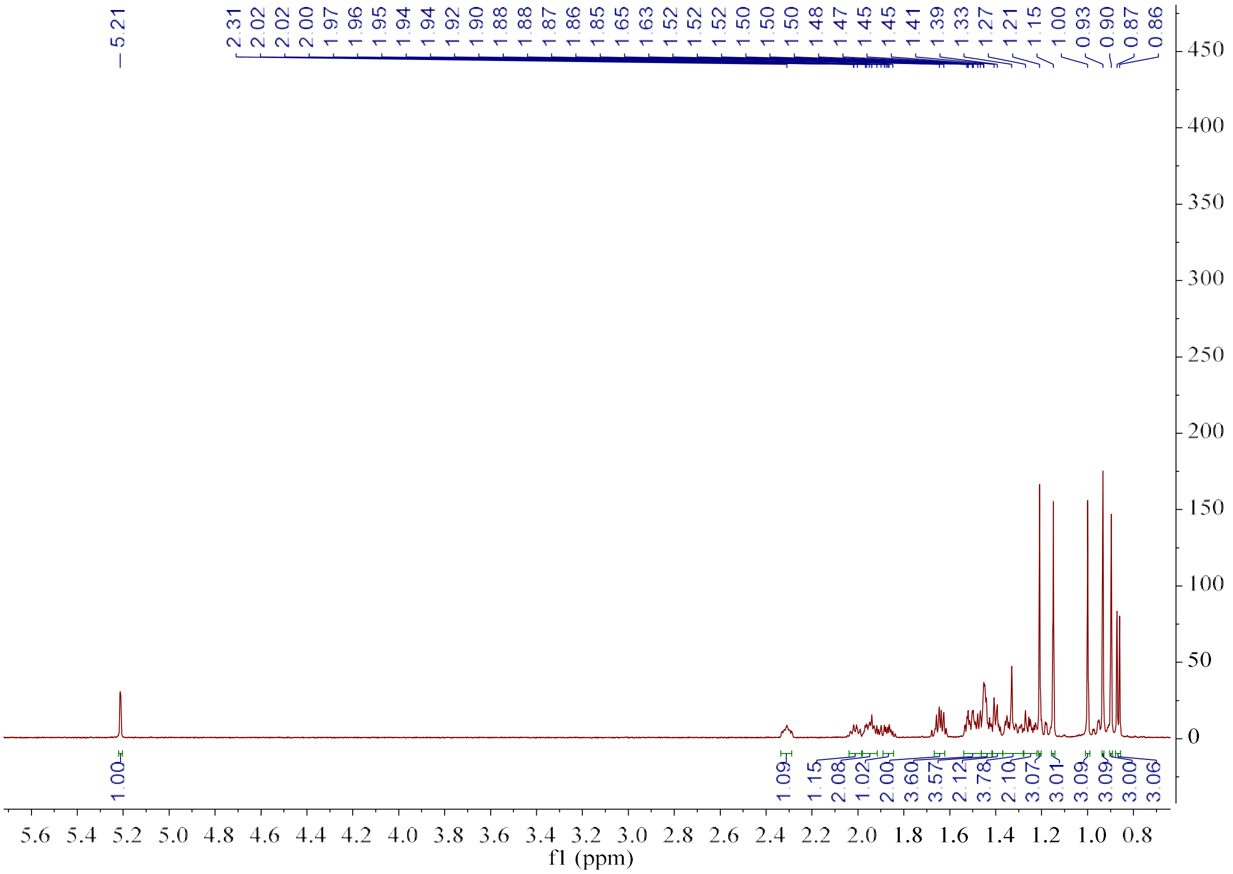

**Figure S18b.** $^1$H NMR spectrum of **21** (600 MHz, Benzene-$d_6$).

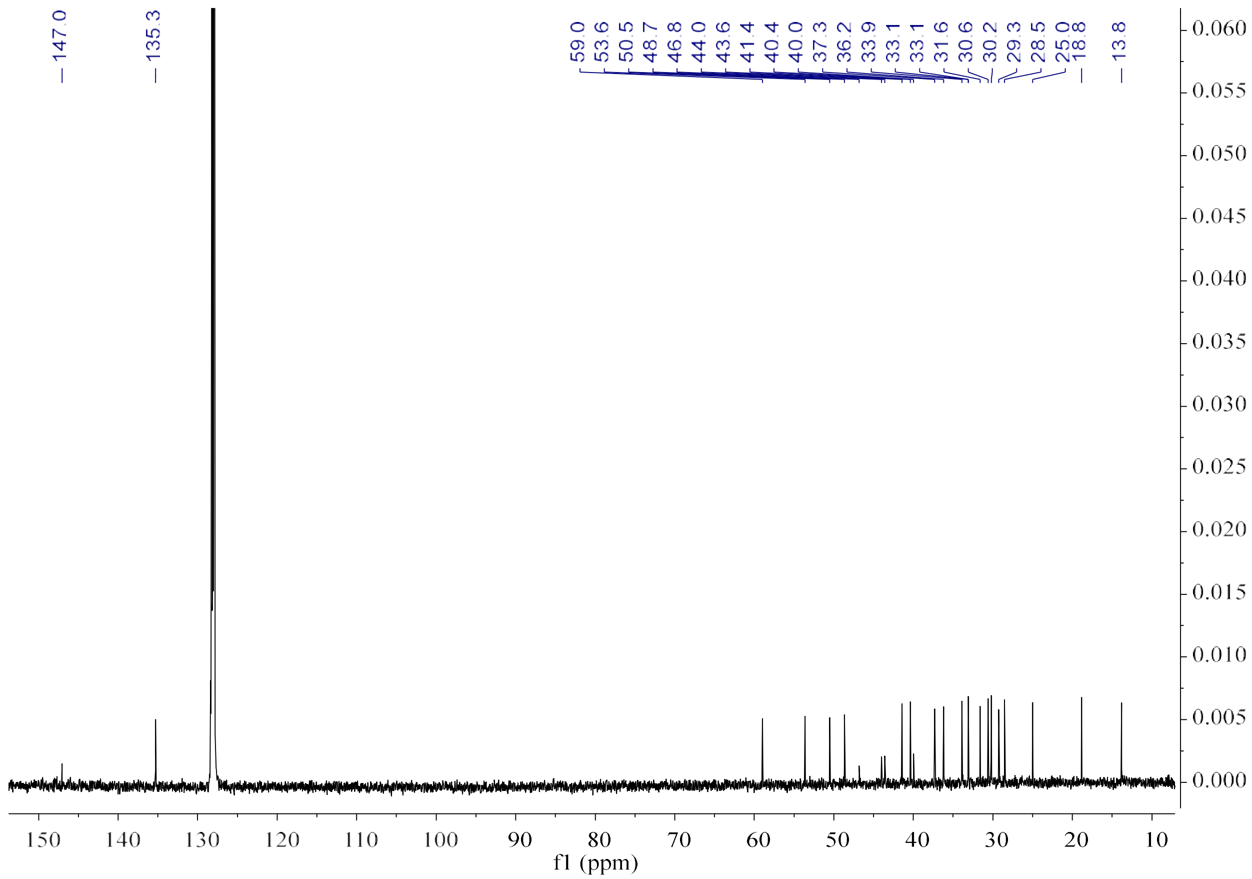

**Figure S18c.** $^{13}C$ NMR spectrum of **21** (150 MHz, Benzene-$d_6$).

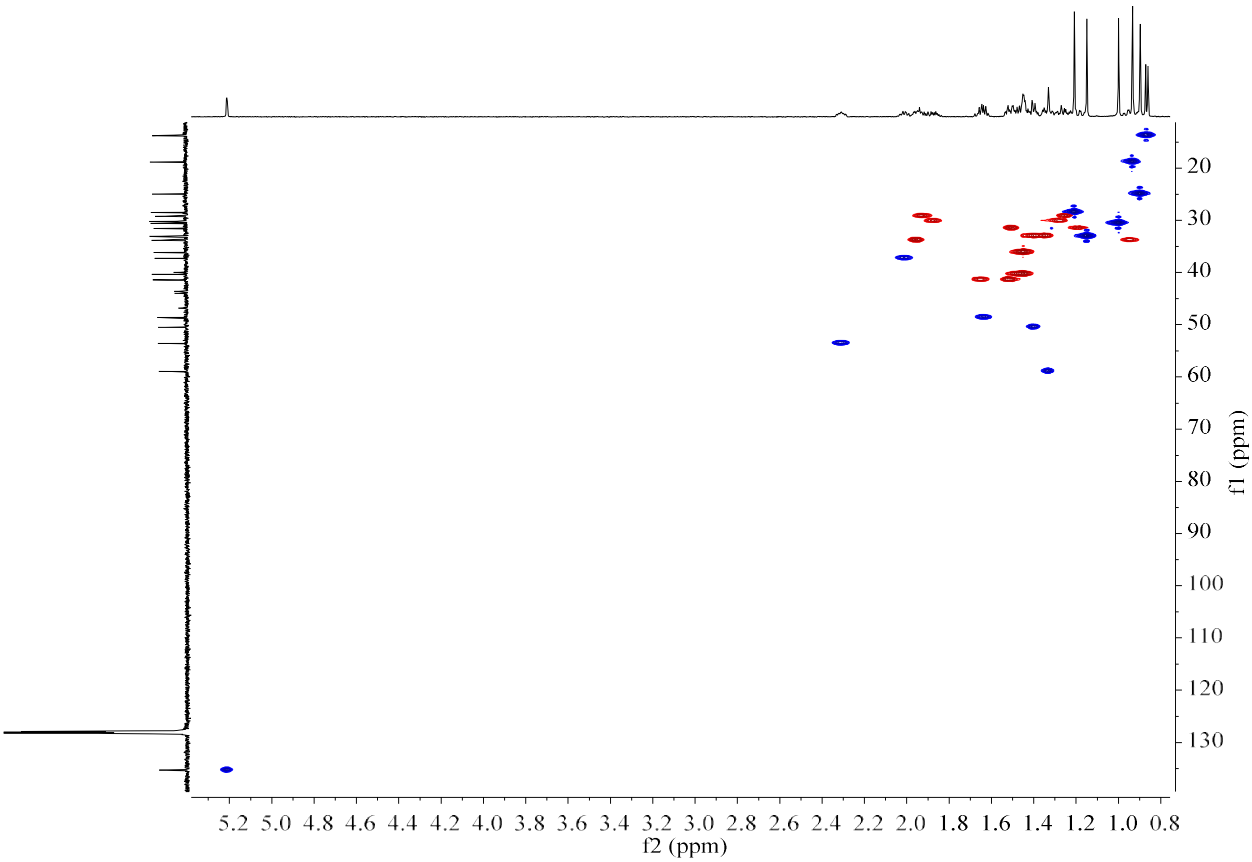

**Figure S18d.** HSQC spectrum of **21** (150 MHz/600 MHz, Benzene-$d_6$).

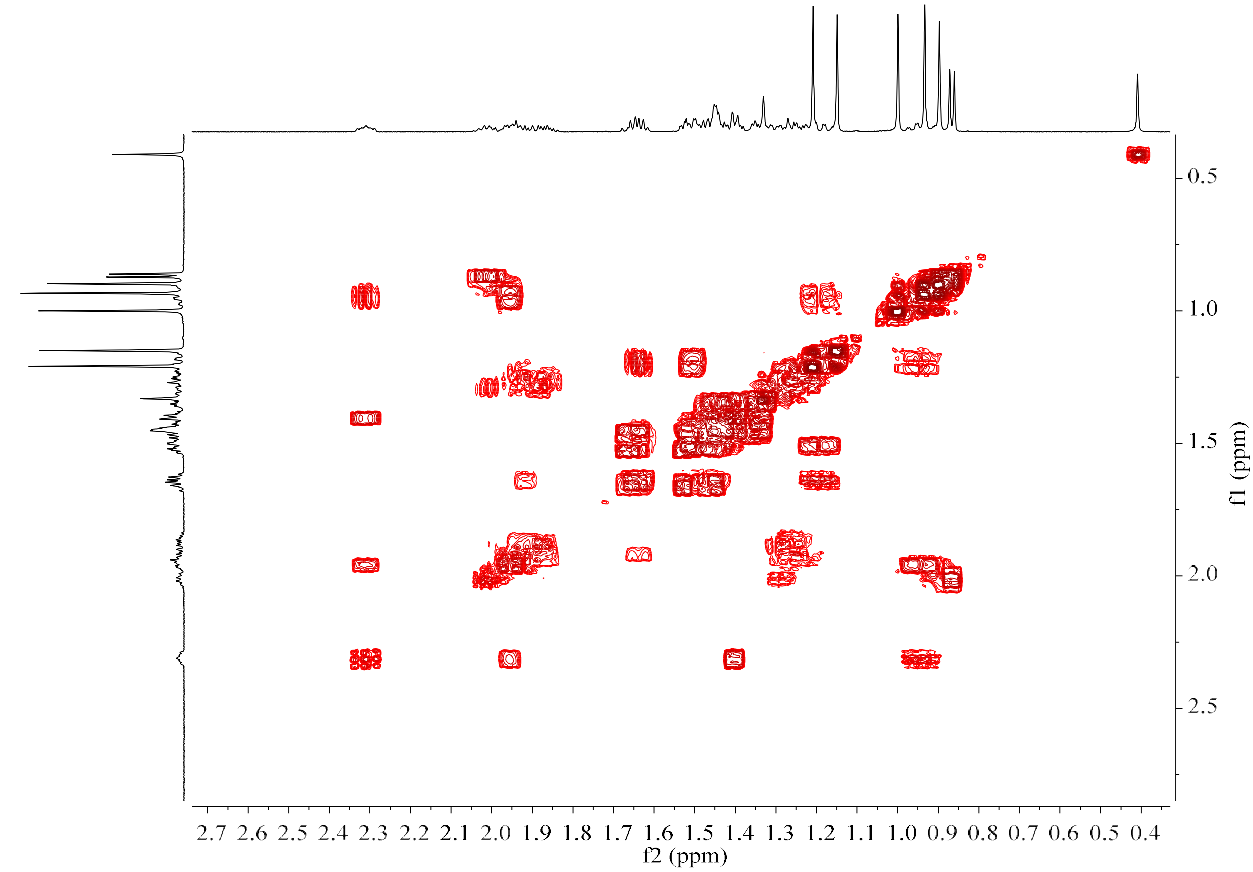

**Figure S18e.** $^1$H–$^1$H COSY spectrum of **21** (600 MHz, Benzene-*d*$_6$).

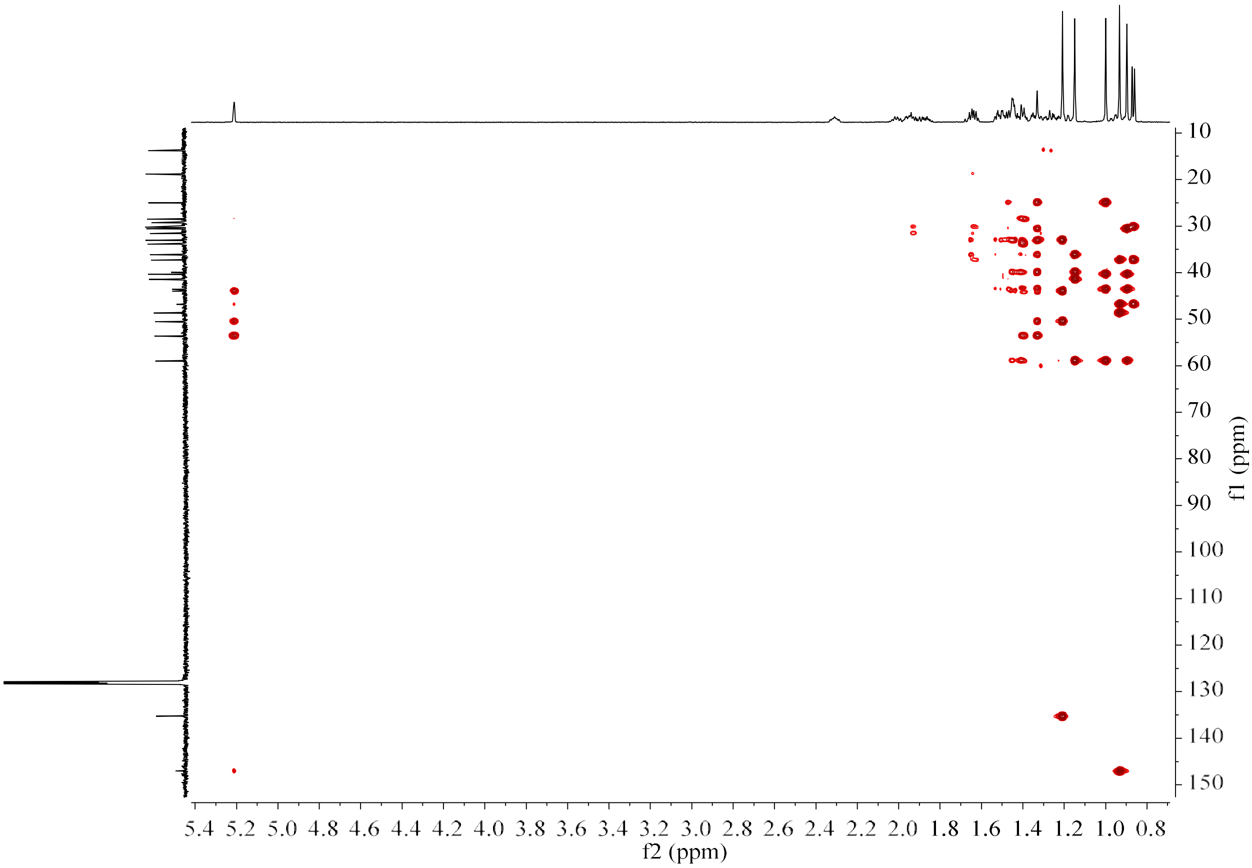

**Figure S18f.** HMBC spectrum of **21** (150 MHz/600 MHz, Benzene-*d*$_6$).

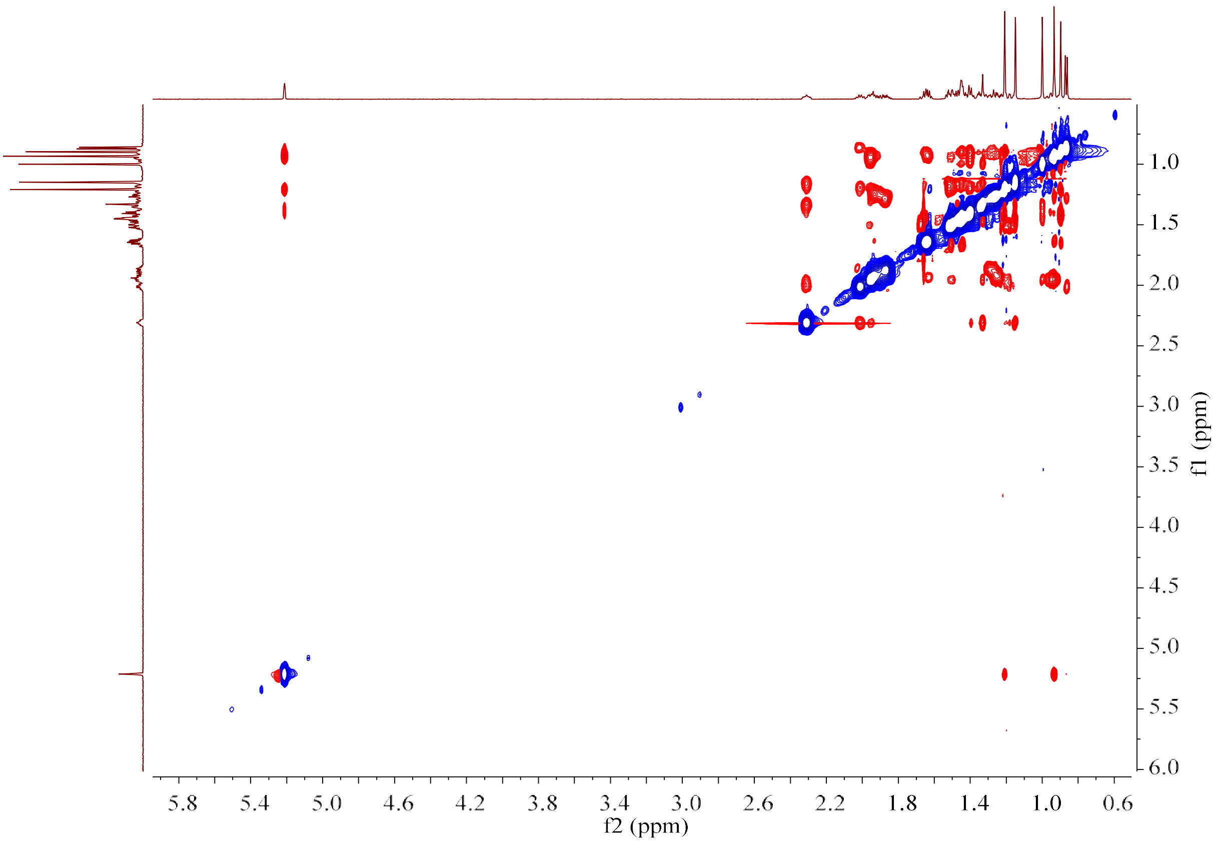

**Figure S18g.** NOESY spectrum of **21** (600 MHz, Benzene-$d_6$).

**Figure S19.** HR-EI-MS and NMR spectrum of **22**.

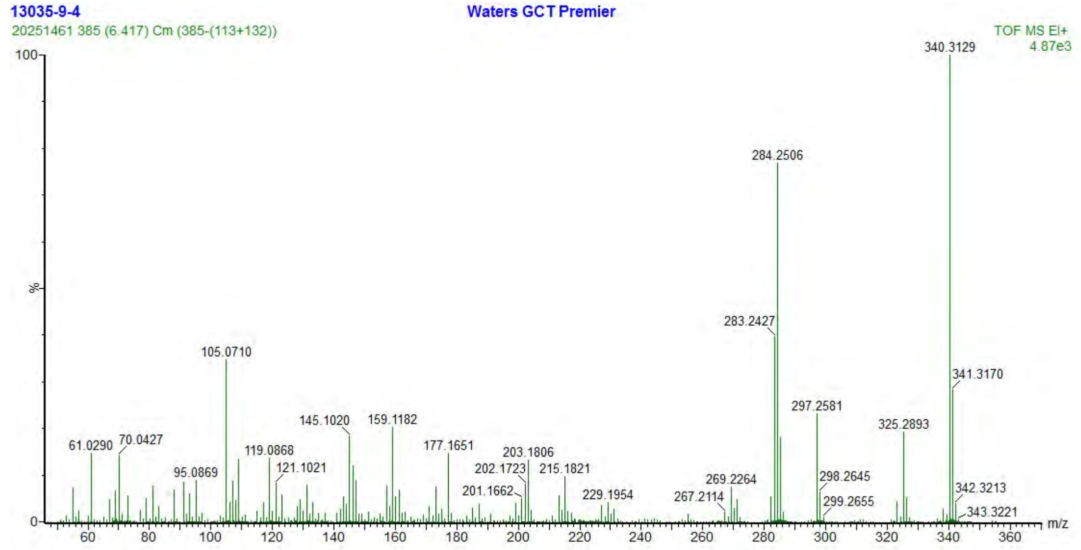

**Figure S19a.** HR-EI-MS spectrum of **22**.

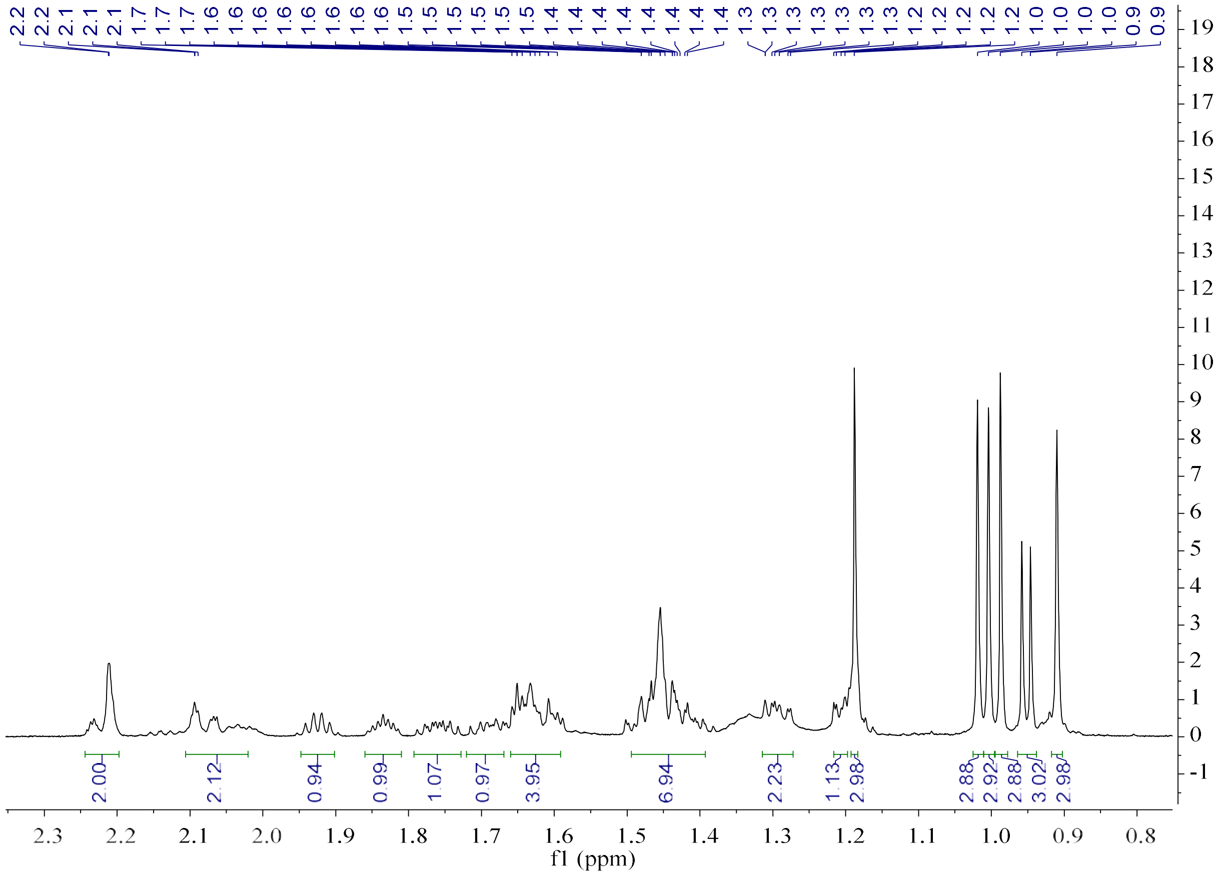

**Figure S19b.** $^{1}H$ NMR spectrum of **22** (600 MHz, Benzene-$d_6$).

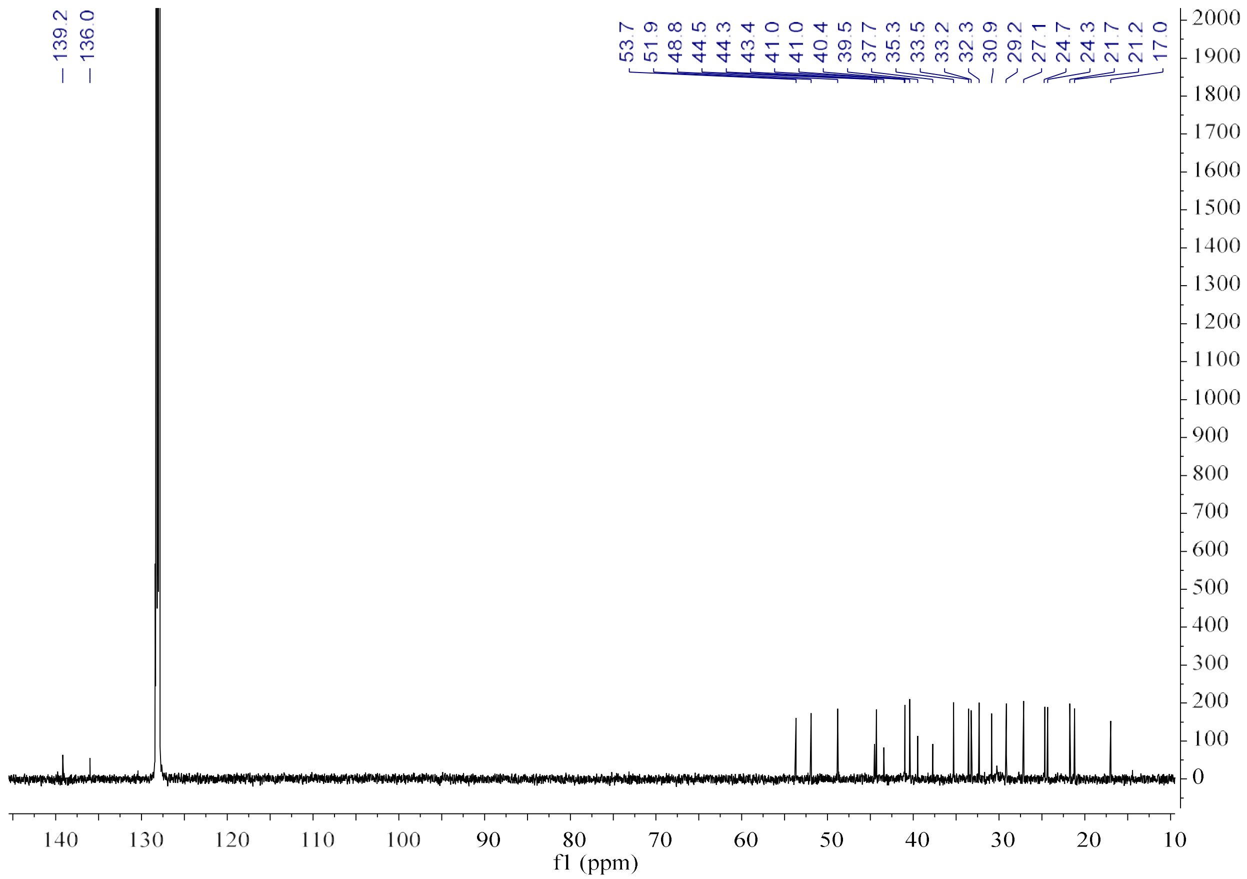

**Figure S19c.** $^{13}$C NMR spectrum of **22** (150 MHz, Benzene-$d_6$).

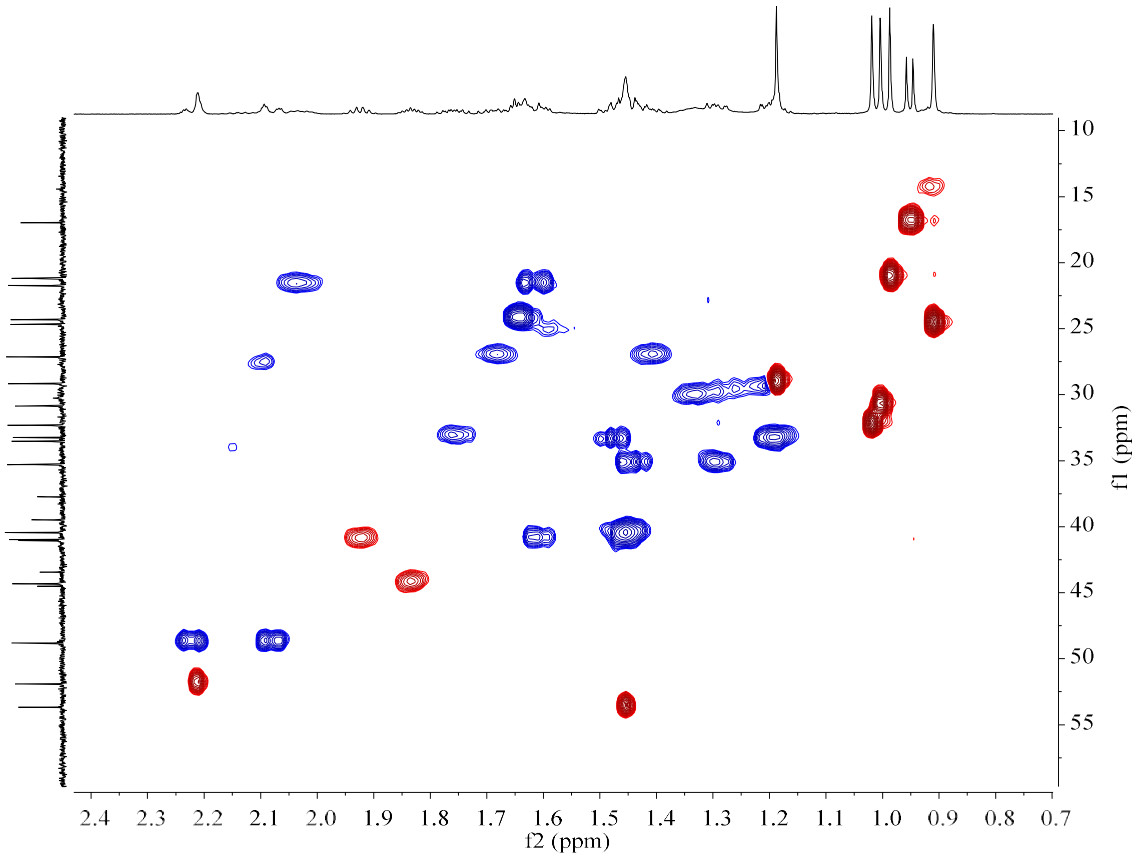

**Figure S19d.** HSQC spectrum of **22** (150 MHz/600 MHz, Benzene-$d_6$).

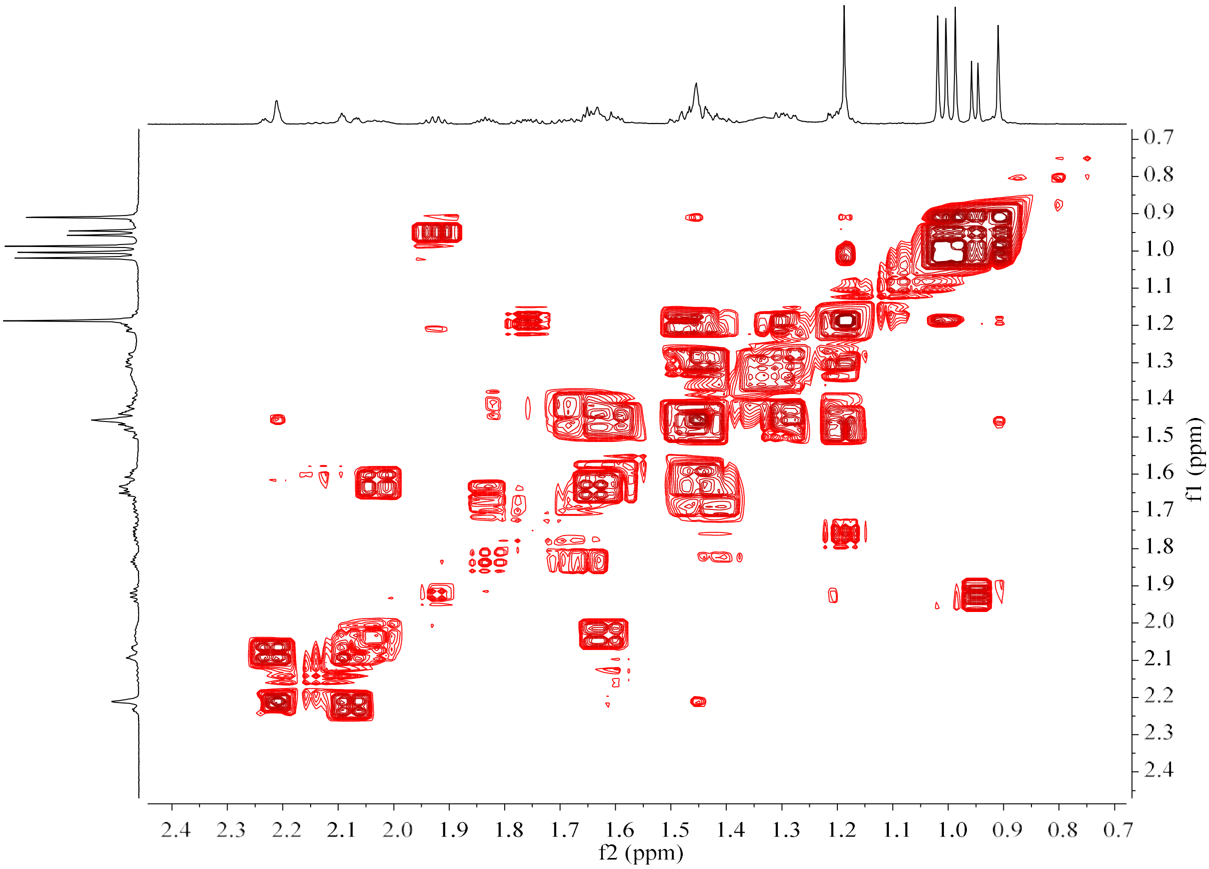

**Figure S19e.** $^{1}H$–$^{1}H$ COSY spectrum of **22** (600 MHz, Benzene-$d_6$).

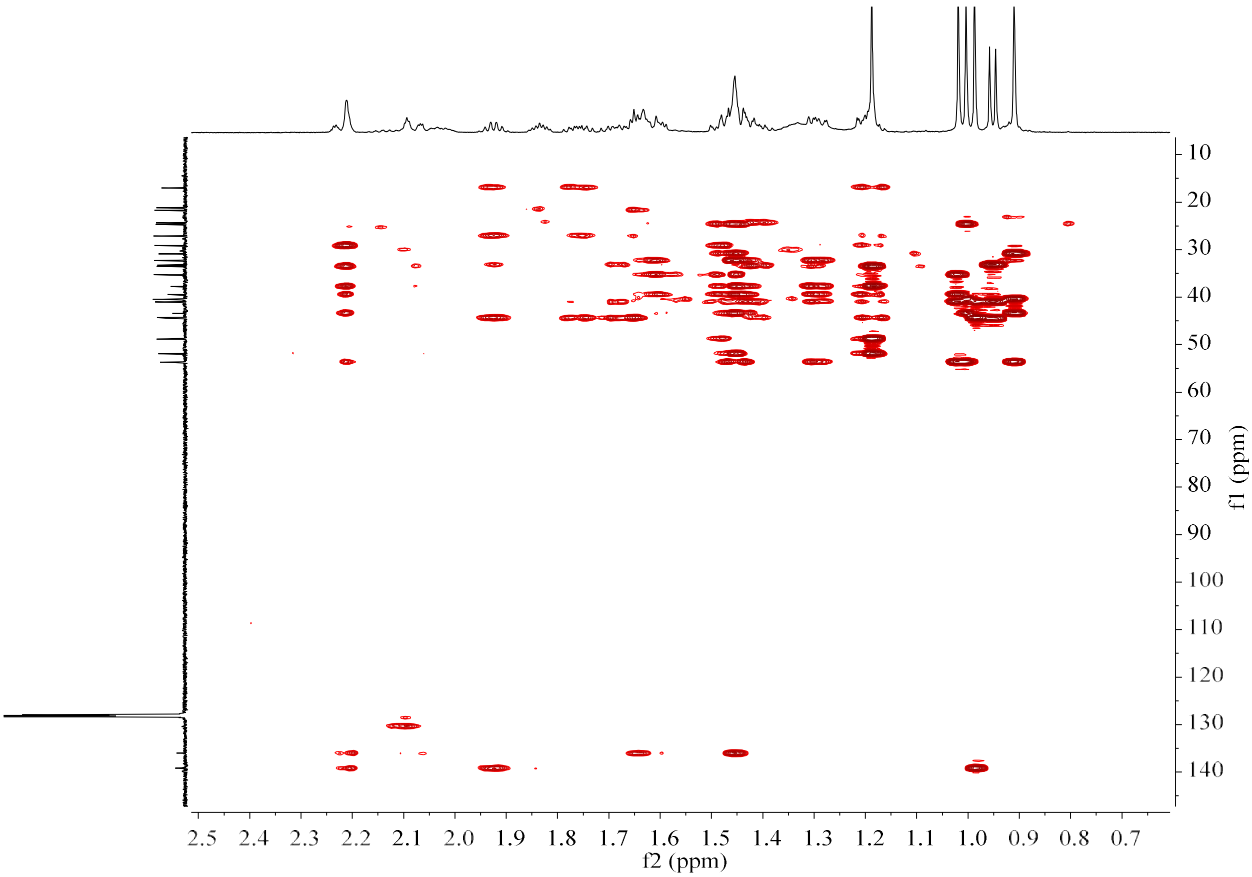

**Figure S19f.** HMBC spectrum of **22** (150 MHz/600 MHz, Benzene-$d_6$).

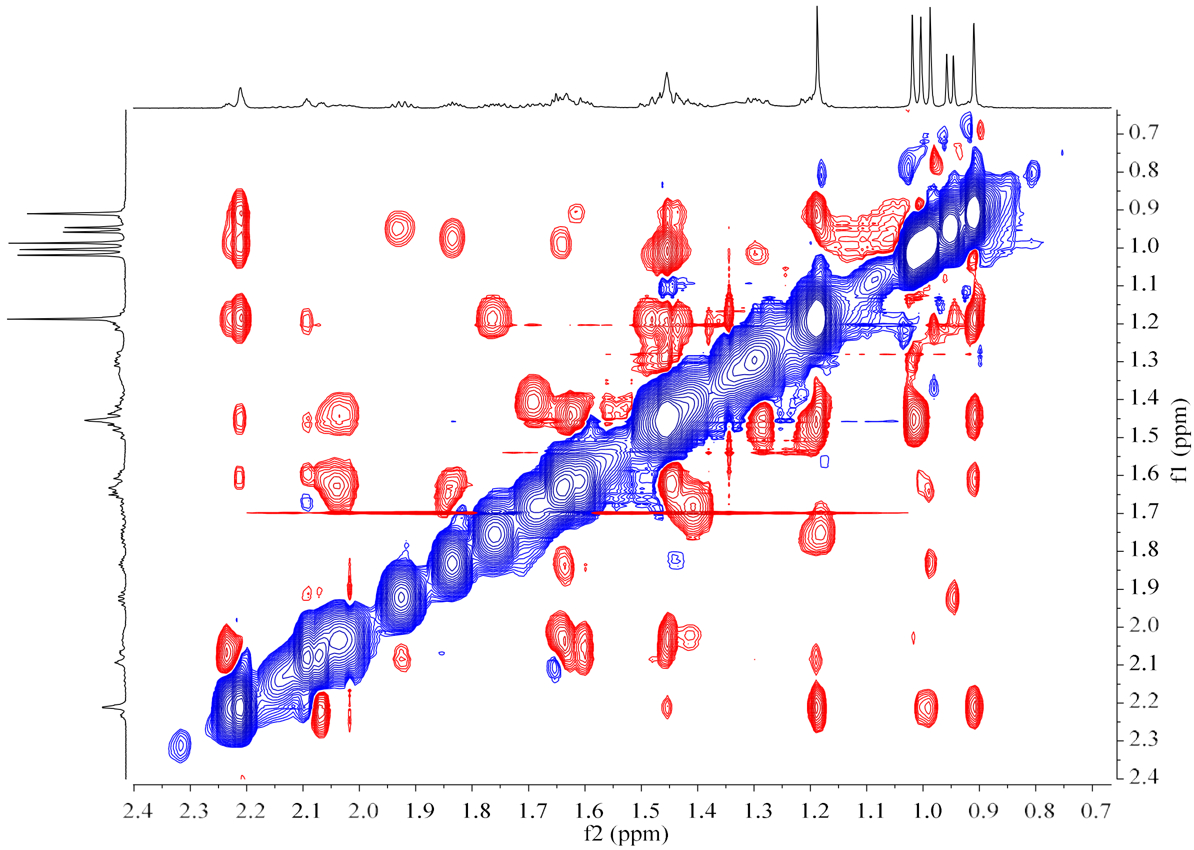

**Figure S19g.** NOESY spectrum of **22** (600 MHz, Benzene-$d_6$).

**Figure S20.** HR-EI-MS and NMR spectrum of **23**.

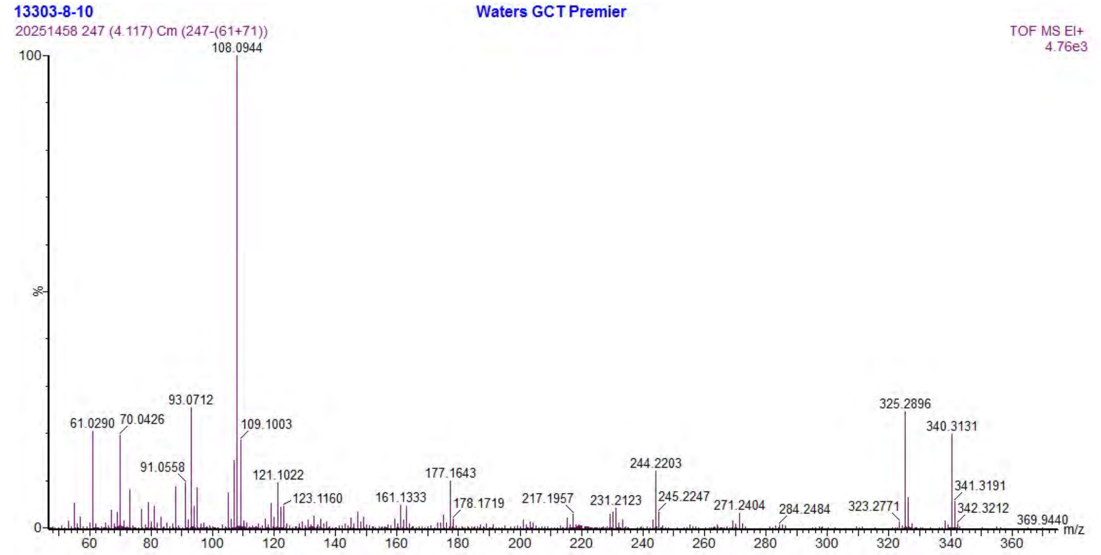

**Figure S20a.** HR-EI-MS spectrum of **23**.

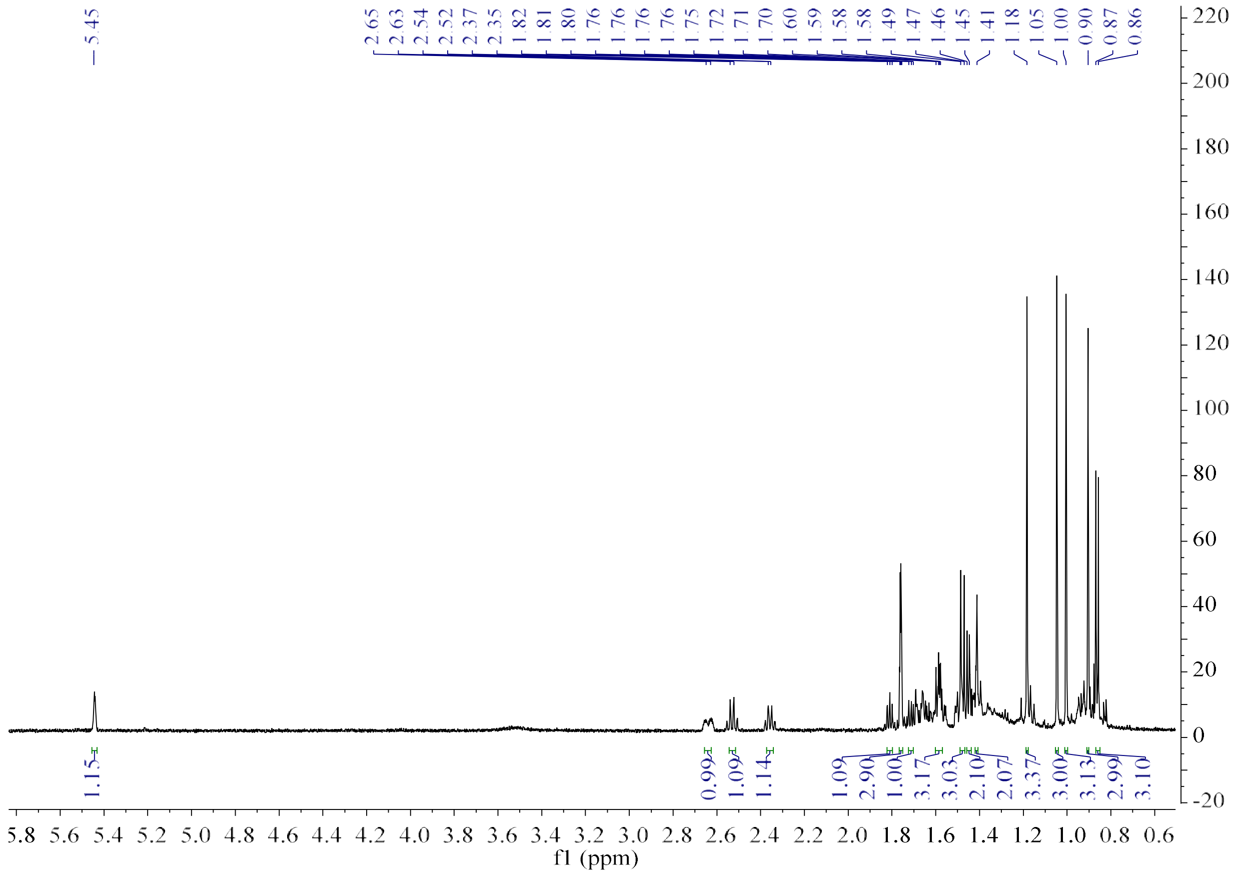

**Figure S20b.** [1]H NMR spectrum of **23** (600 MHz, Benzene-$d_6$).

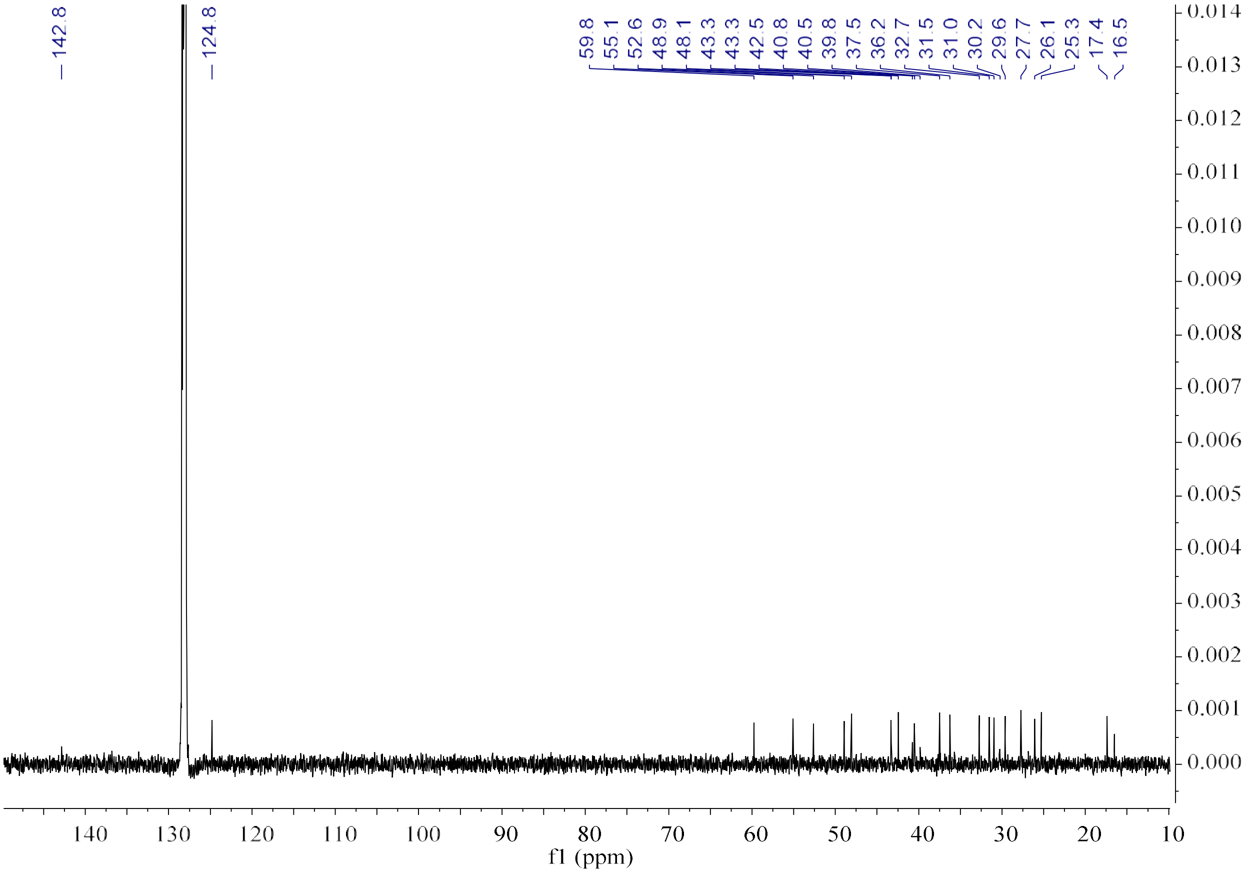

**Figure S20c.** $^{13}C$ NMR spectrum of **23** (150 MHz, Benzene-$d_6$).

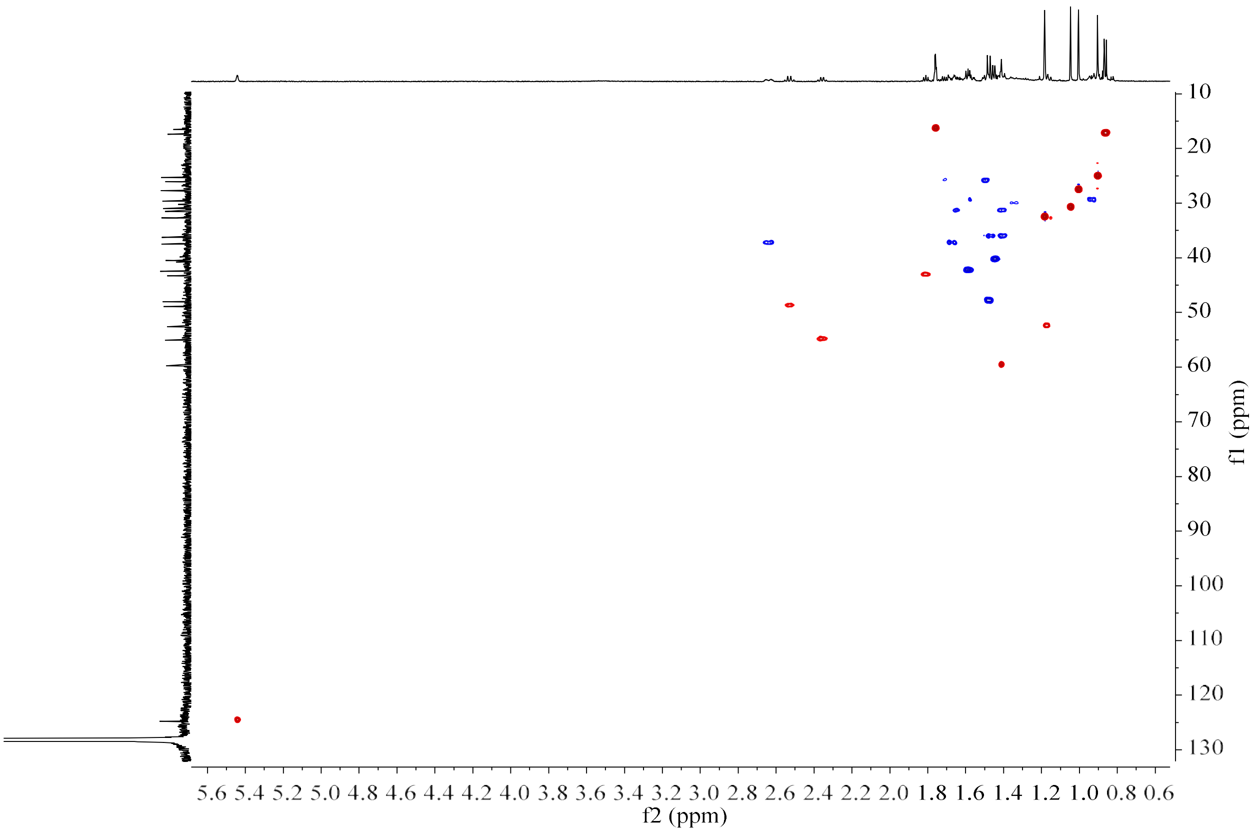

**Figure S20d.** HSQC spectrum of **23** (150 MHz/600 MHz, Benzene-$d_6$).

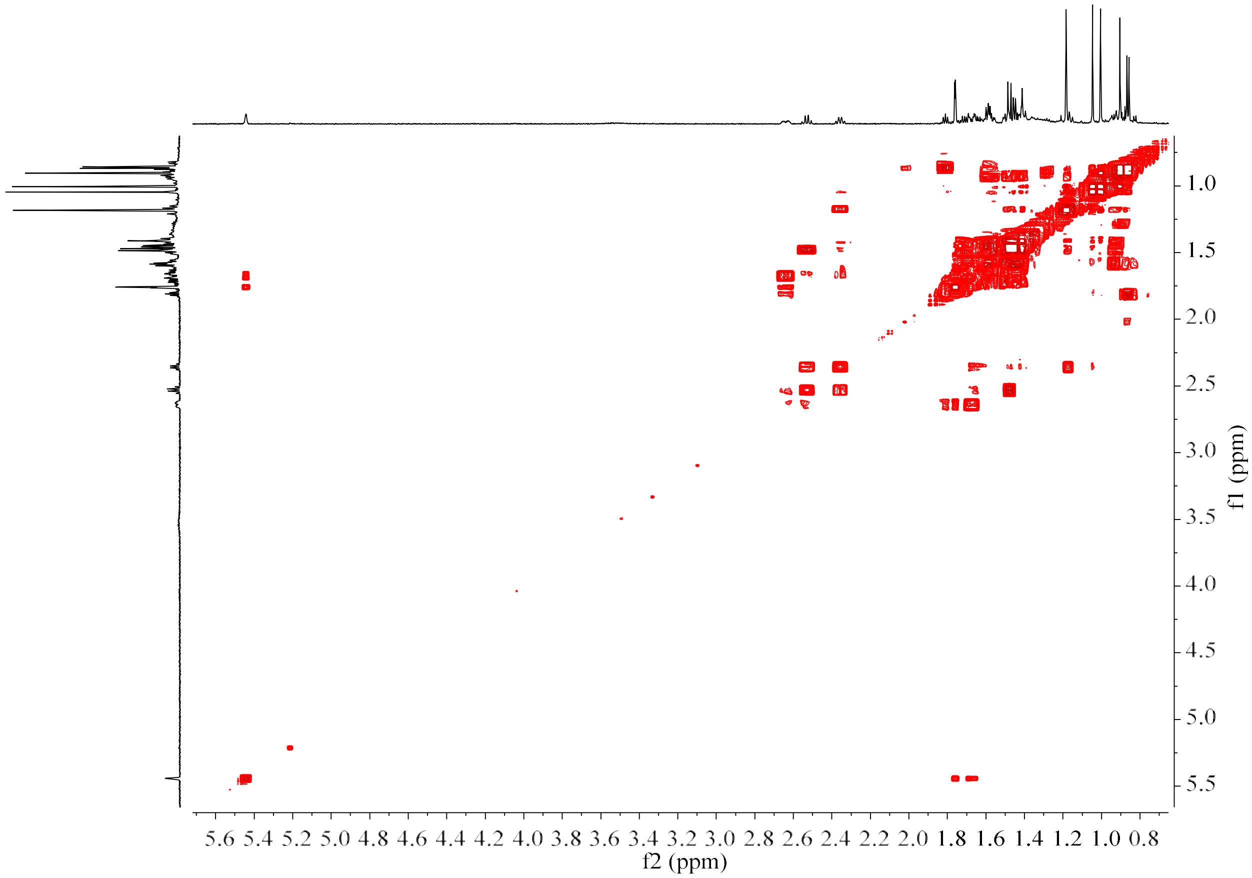

**Figure S20e.** $^1$H–$^1$H COSY spectrum of **23** (600 MHz, Benzene-$d_6$).

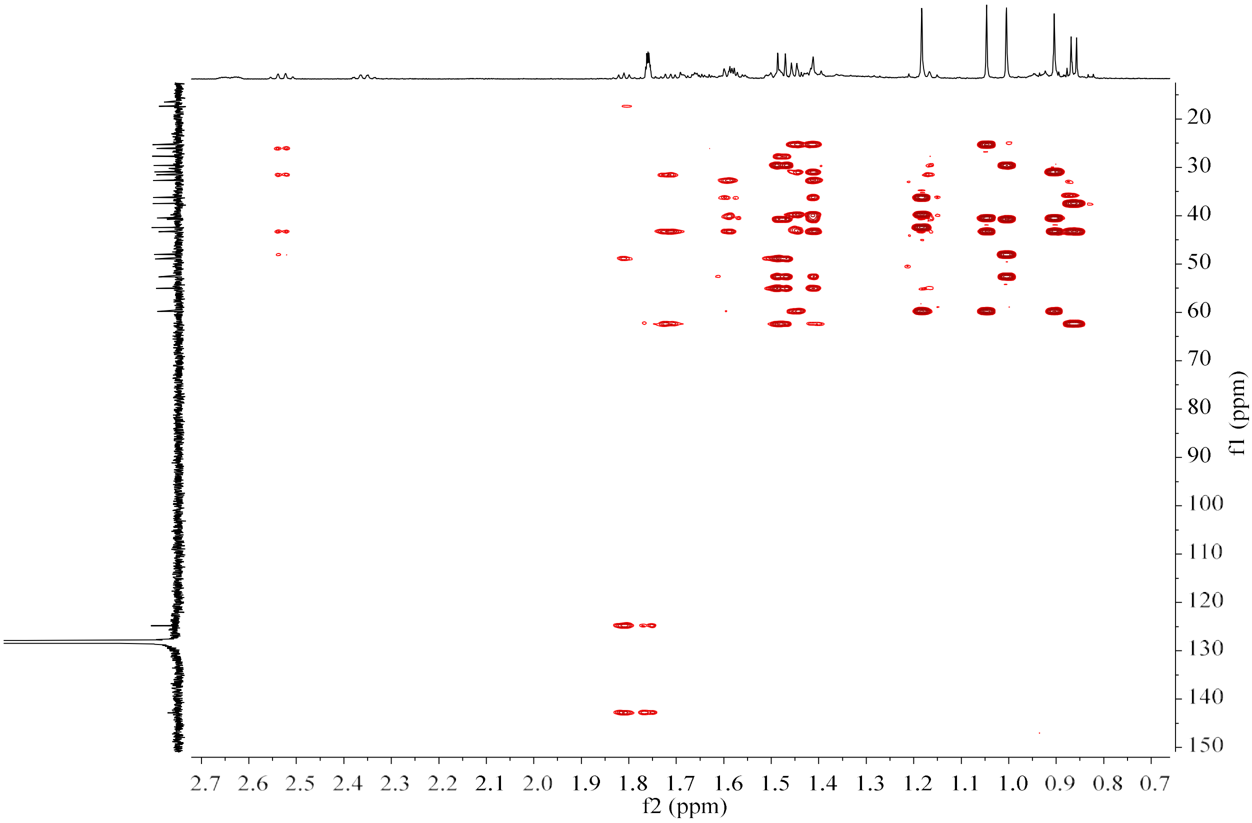

**Figure S20f.** HMBC spectrum of **23** (150 MHz/600 MHz, Benzene-$d_6$).

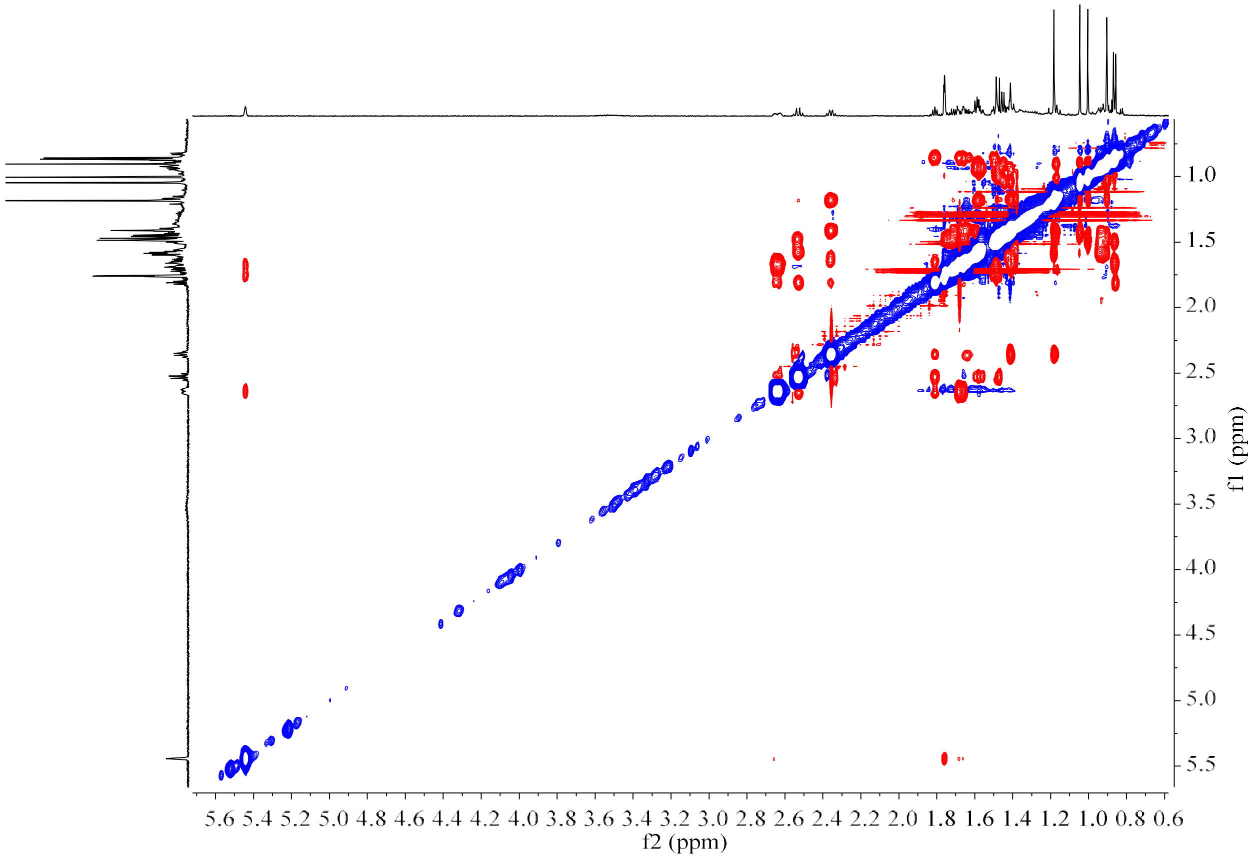

**Figure S20g.** NOESY spectrum of **23** (600 MHz, Benzene-$d_6$).

**Figure S21.** HR-EI-MS and NMR spectrum of **24**.

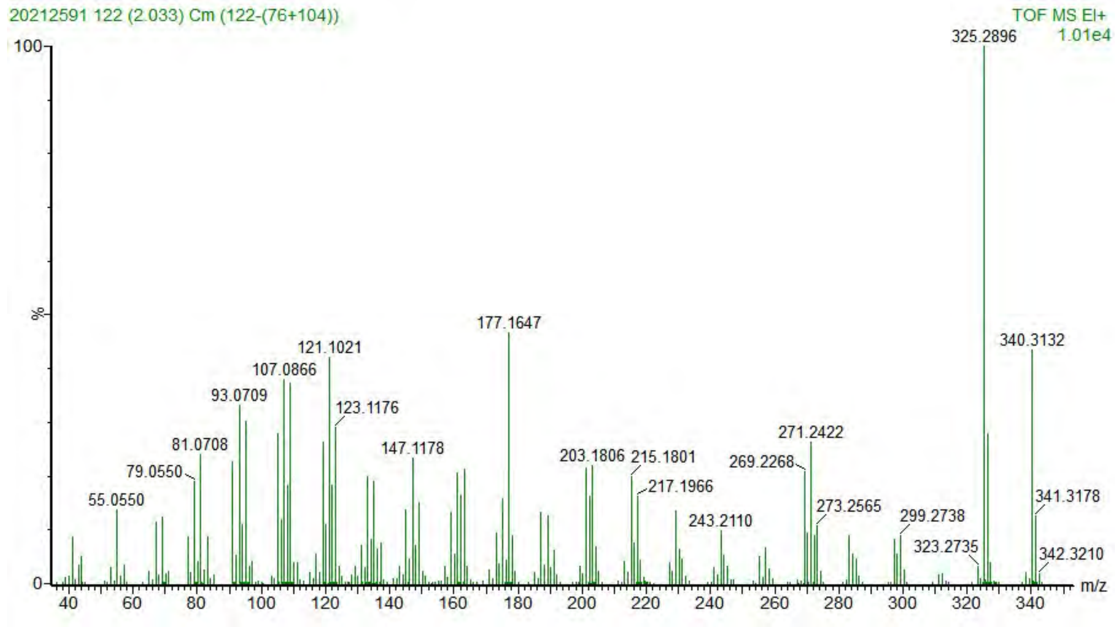

**Figure S21a.** HR-EI-MS spectrum of **24**.

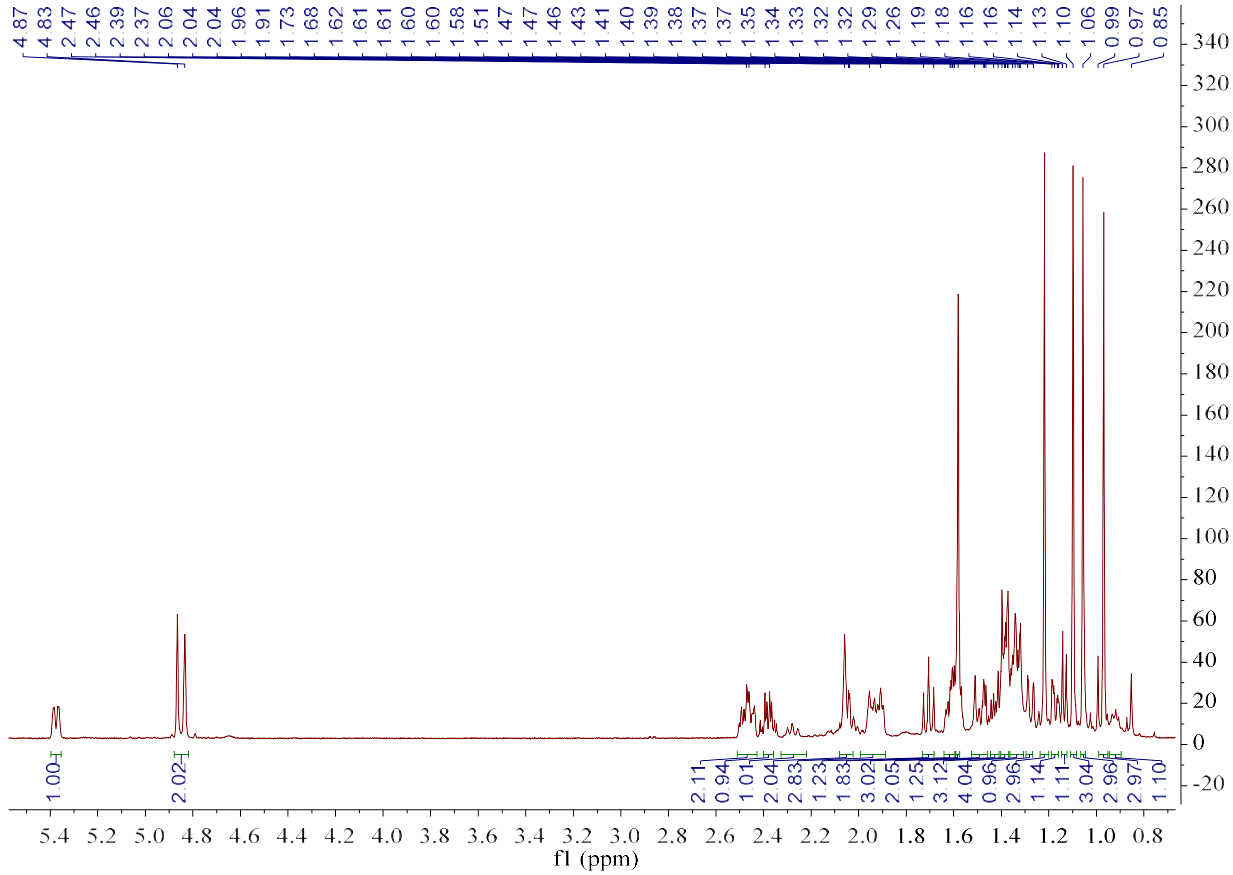

**Figure S21b.** $^{1}$H NMR spectrum of **24** (600 MHz, Benzene-$d_6$).

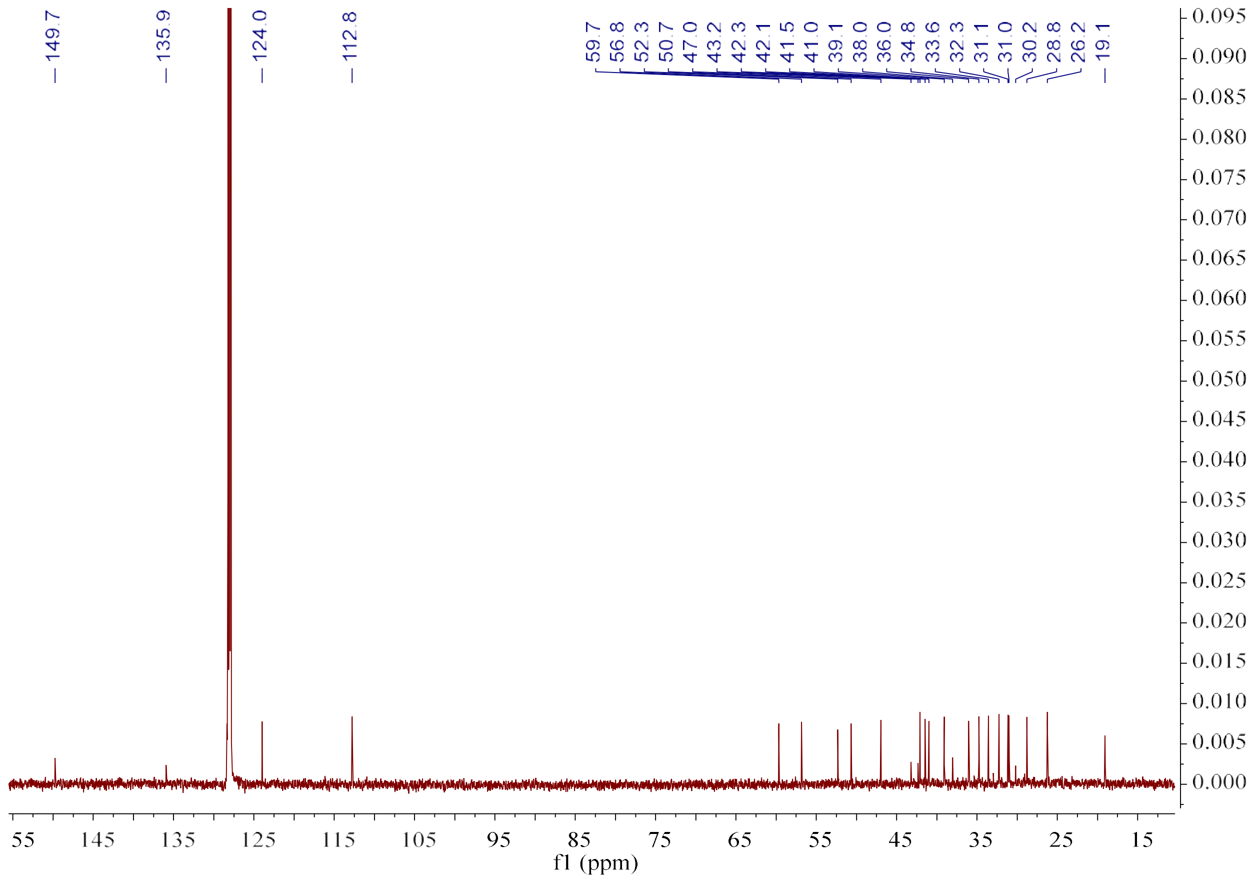

**Figure S21c.** $^{13}$C NMR spectrum of **24** (150 MHz, Benzene-$d_6$).

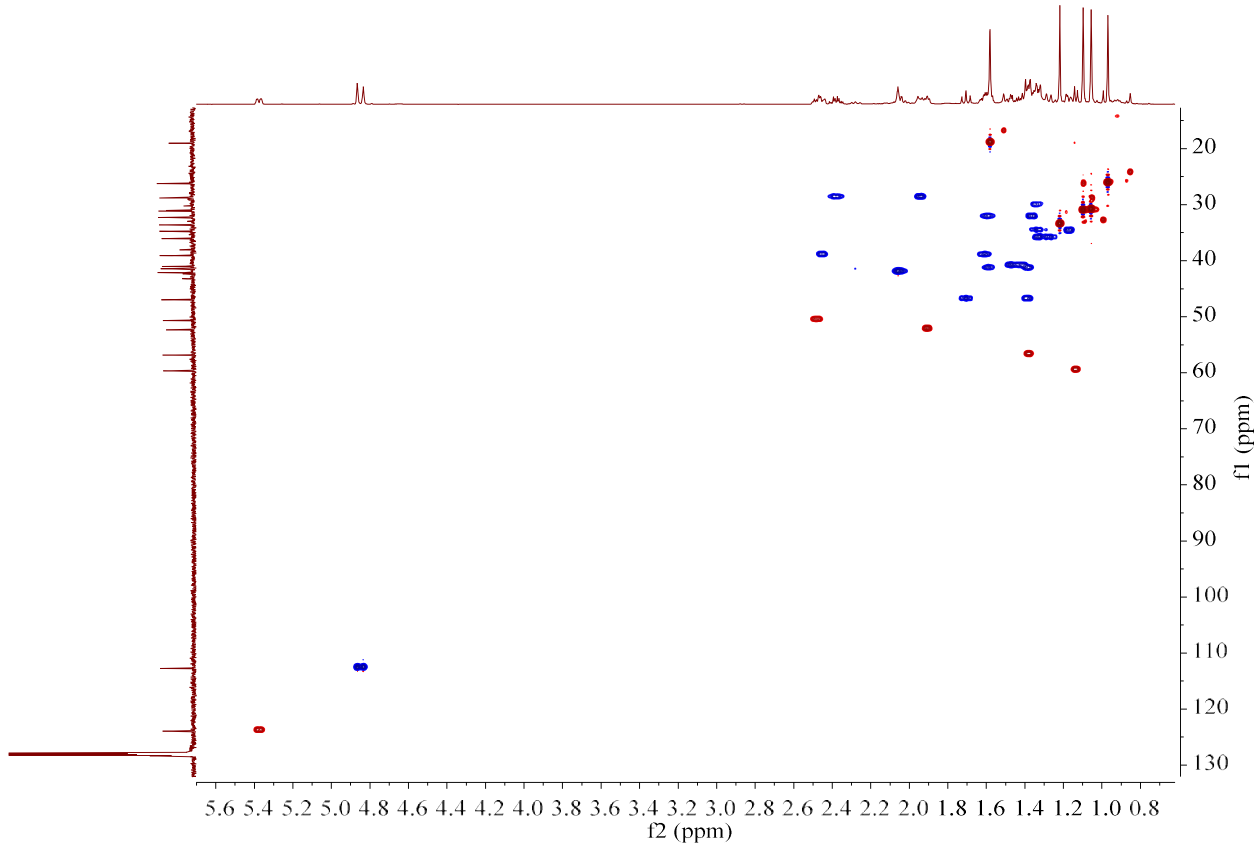

**Figure S21d.** HSQC spectrum of **24** (150 MHz/600 MHz, Benzene-$d_6$).

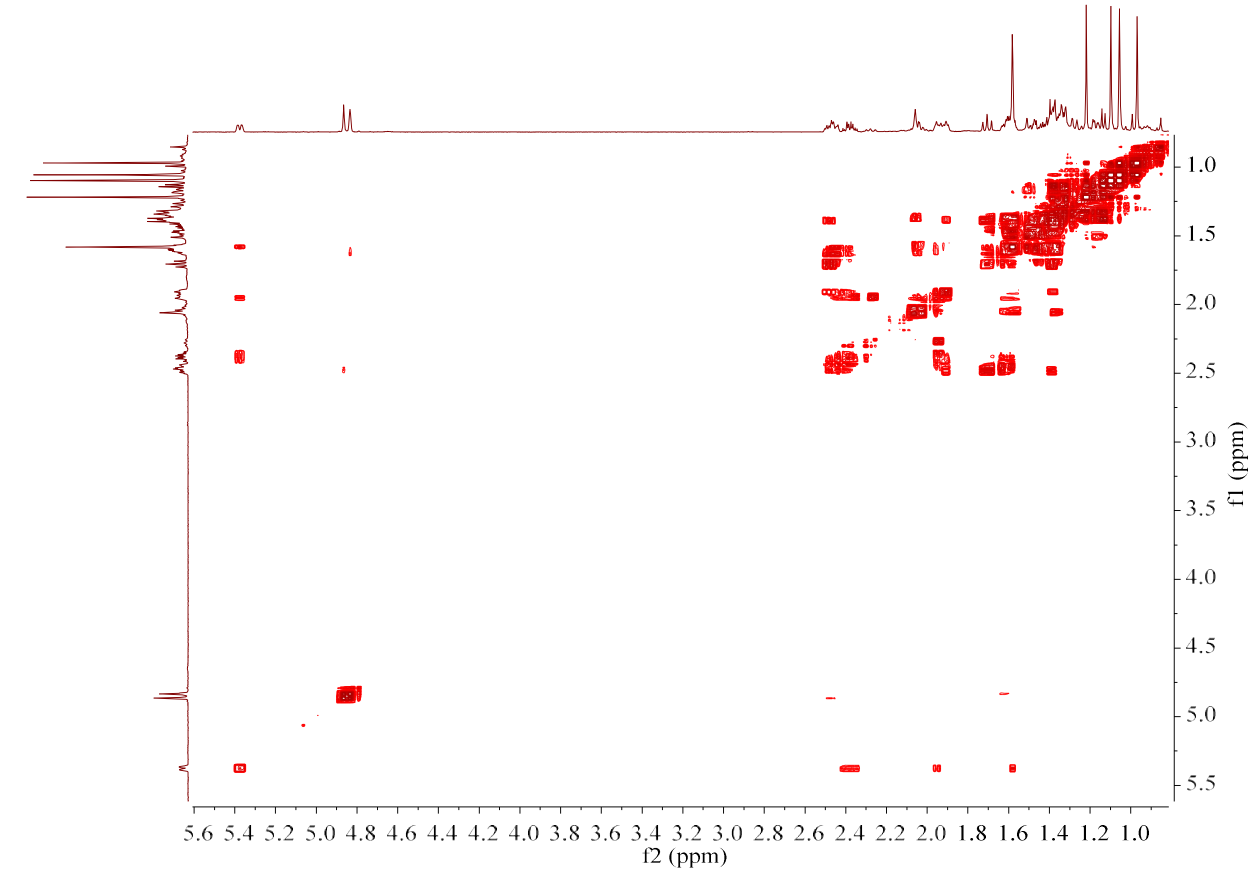

**Figure S21e.** $^{1}H$–$^{1}H$ COSY spectrum of **24** (600 MHz, Benzene-$d_6$).

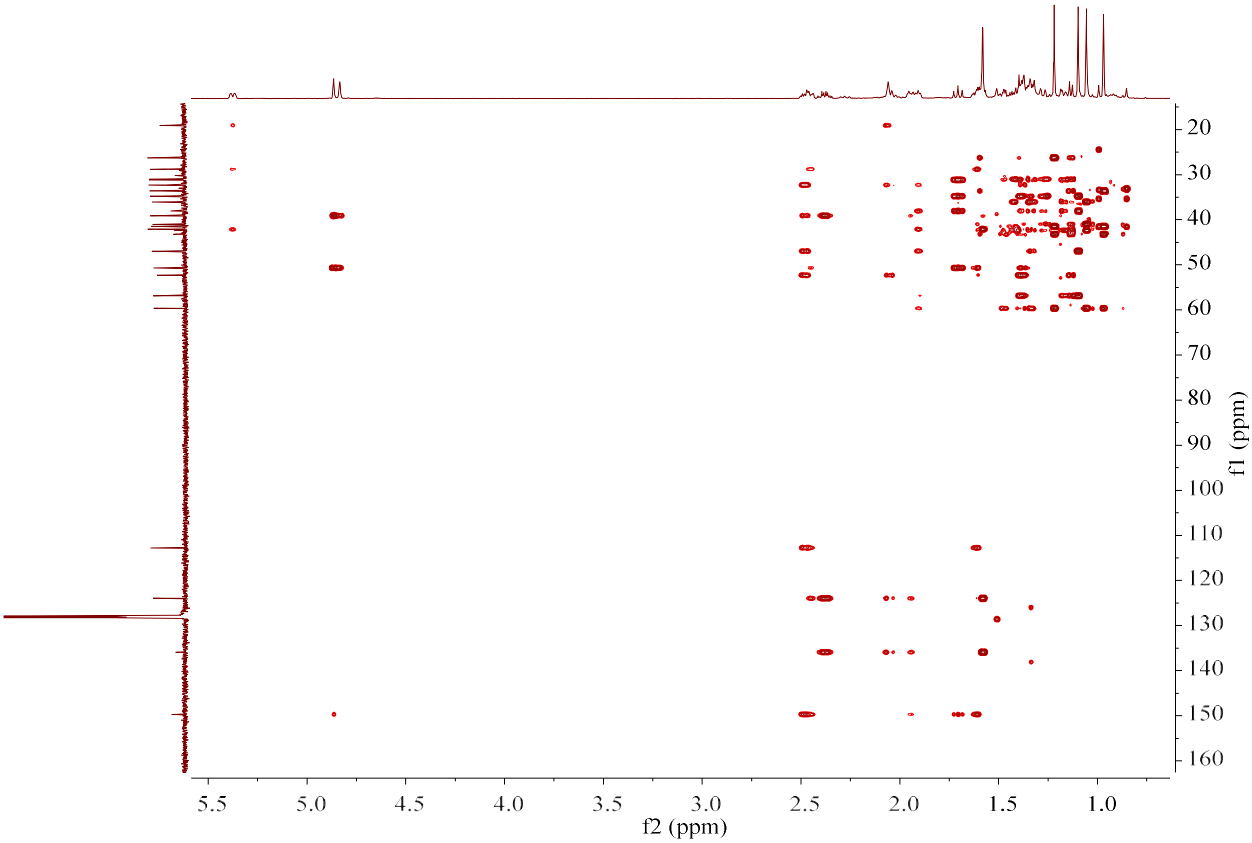

**Figure S21f.** HMBC spectrum of **24** (150 MHz/600 MHz, Benzene-$d_6$).

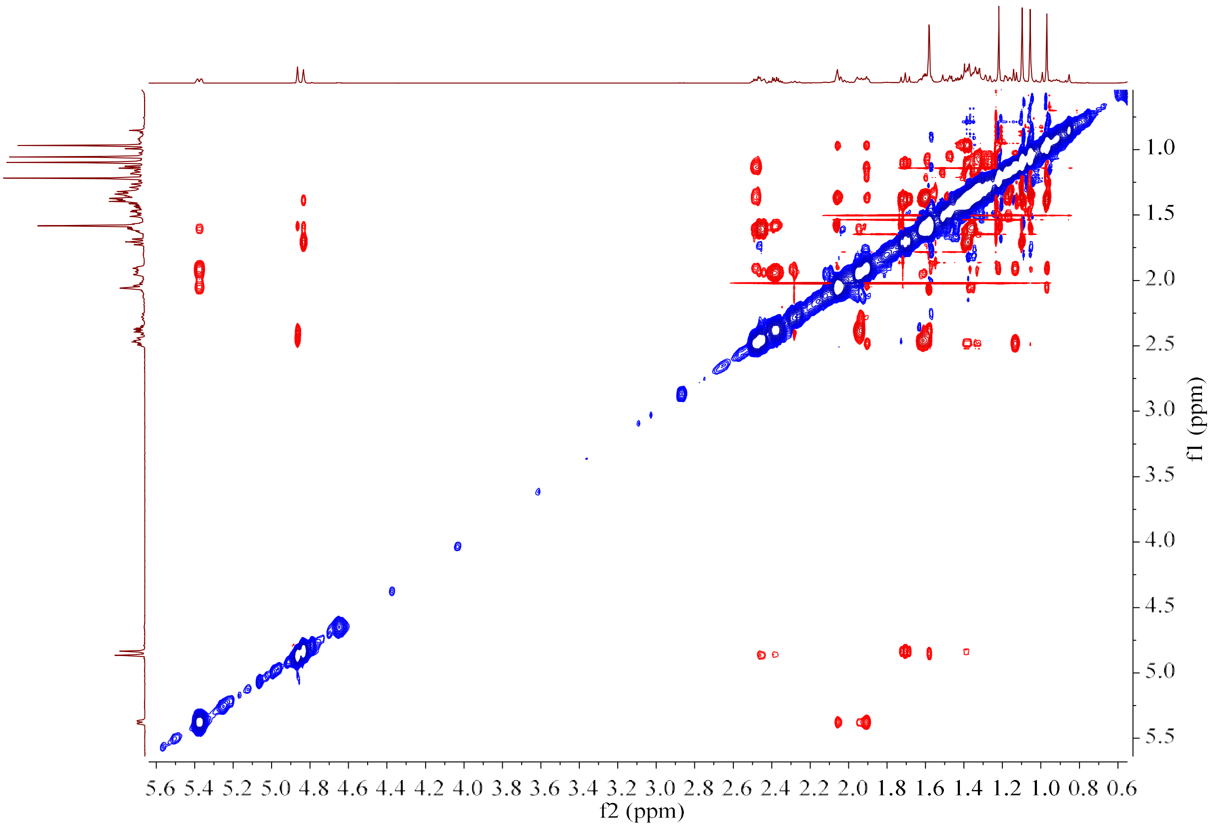

**Figure S21g.** NOESY spectrum of **24** (600 MHz, Benzene-$d_6$).

**Figure S22.** HR-EI-MS and NMR spectrum of **25**.

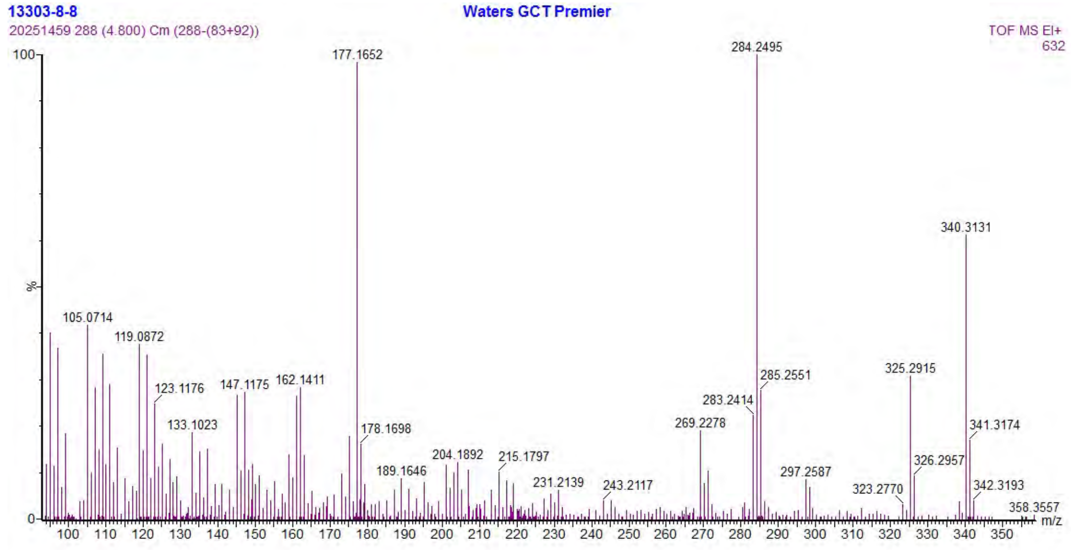

**Figure S22a.** HR-EI-MS spectrum of **25**.

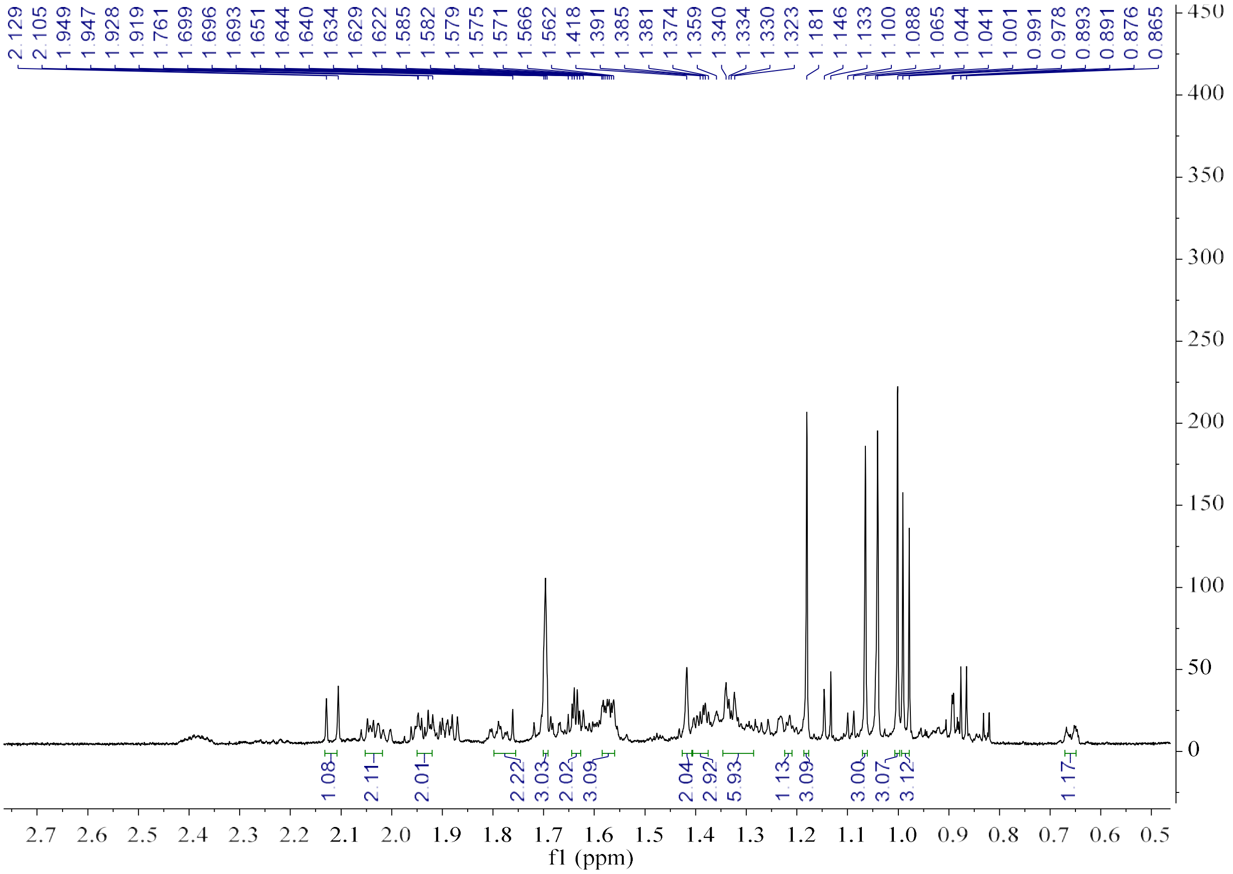

**Figure S22b.** $^1H$ NMR spectrum of **25** (600 MHz, Benzene-$d_6$).

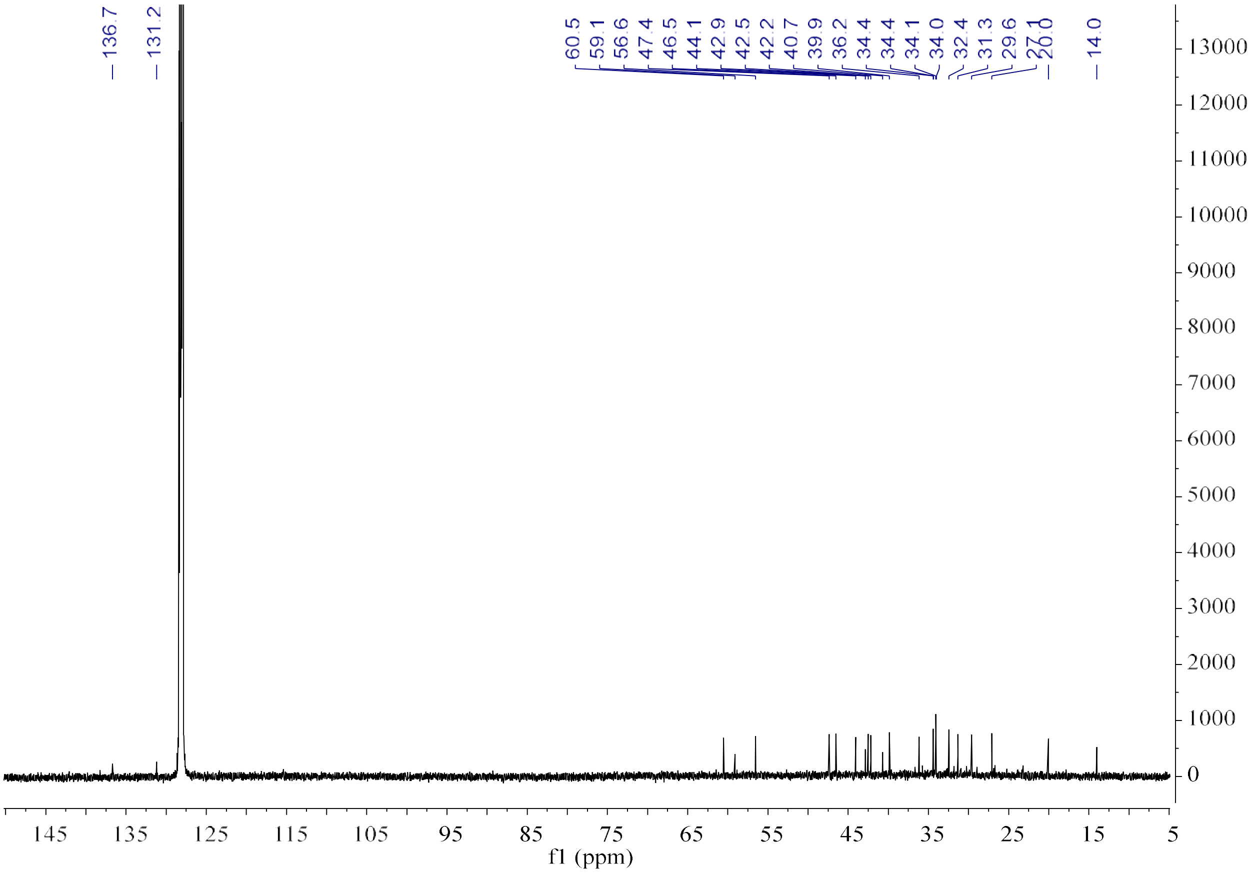

**Figure S22c.** $^{13}$C NMR spectrum of **25** (150 MHz, Benzene-$d_6$).

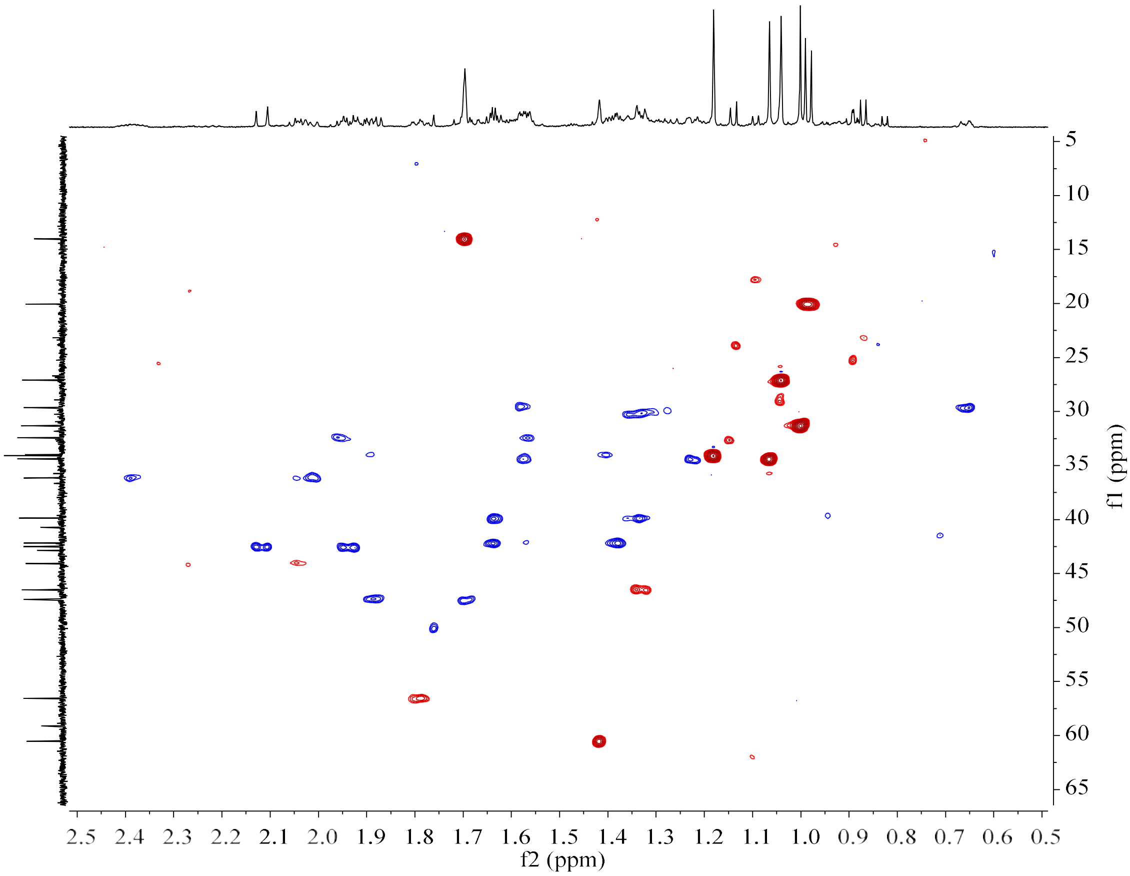

**Figure S22d.** HSQC spectrum of **25** (150 MHz/600 MHz, Benzene-$d_6$).

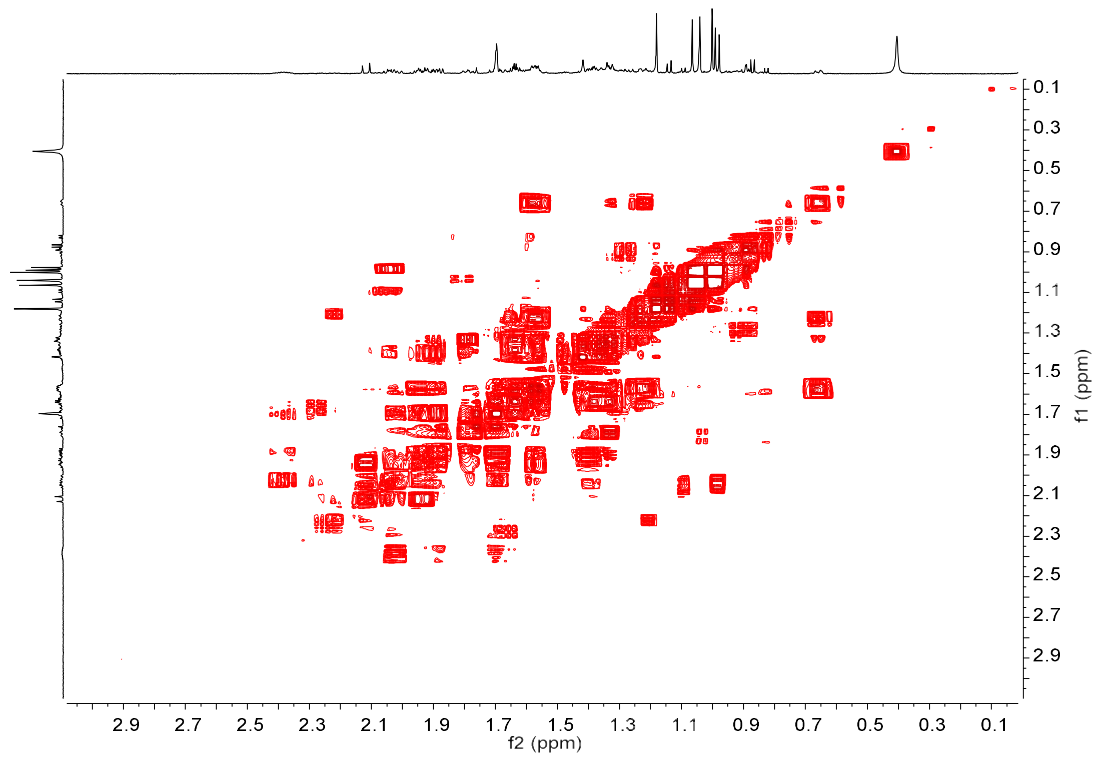

**Figure S22e.** $^1H$–$^1H$ COSY spectrum of **25** (600 MHz, Benzene-$d_6$).

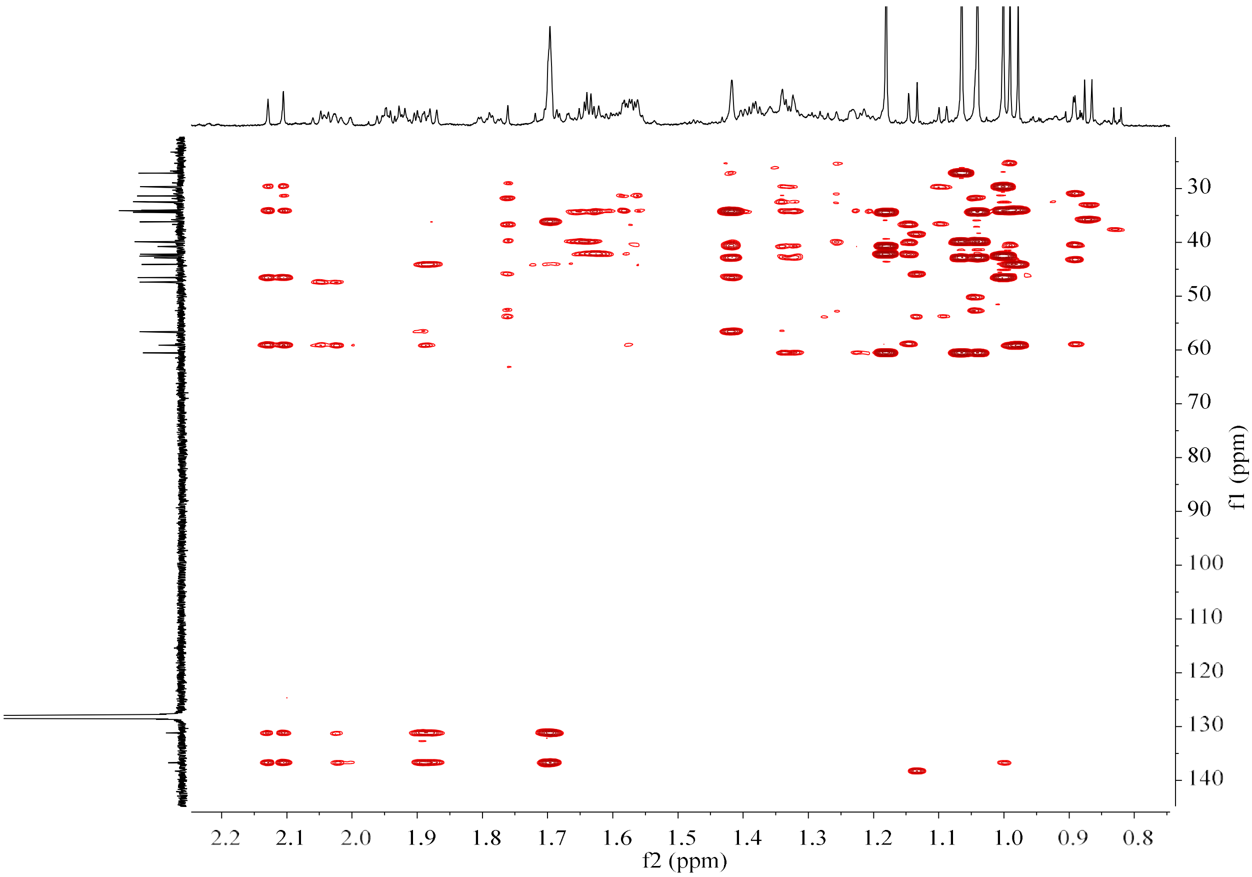

**Figure S22f.** HMBC spectrum of **25** (150 MHz/600 MHz, Benzene-$d_6$).

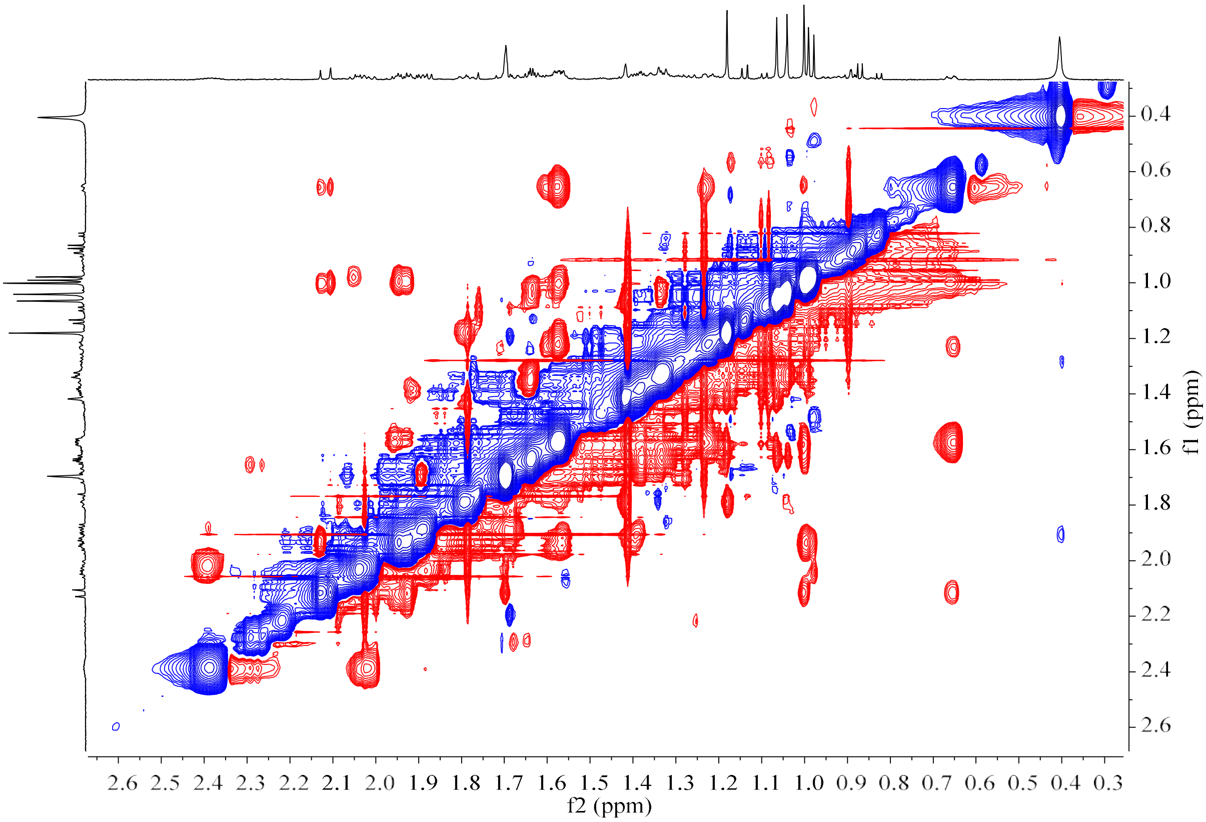

**Figure S22g.** NOESY spectrum of **25** (600 MHz, Benzene-$d_6$).

**Figure S23.** HR-EI-MS and NMR spectrum of **26**.

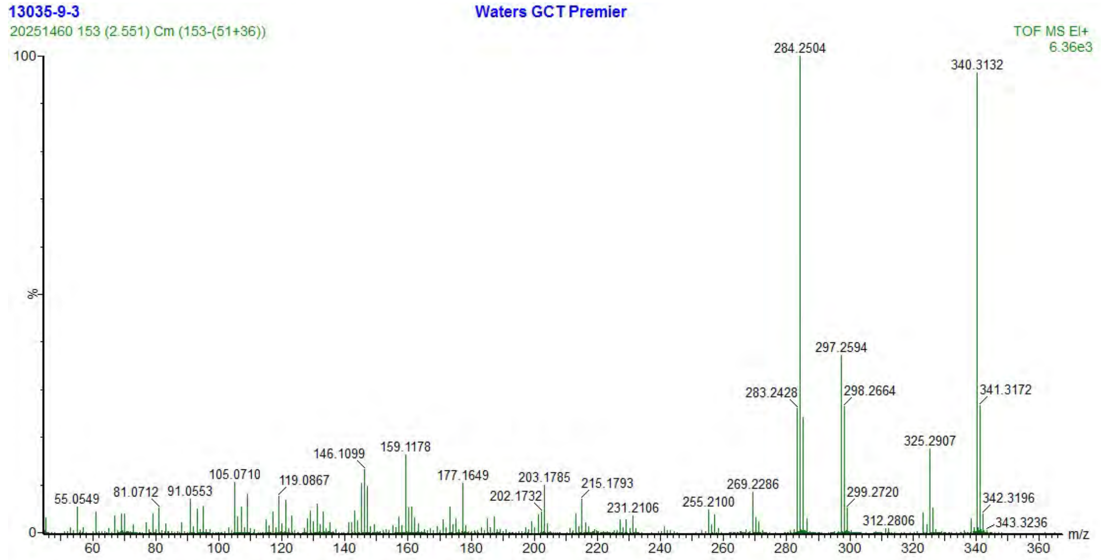

**Figure S23a.** HR-EI-MS spectrum of **26**.

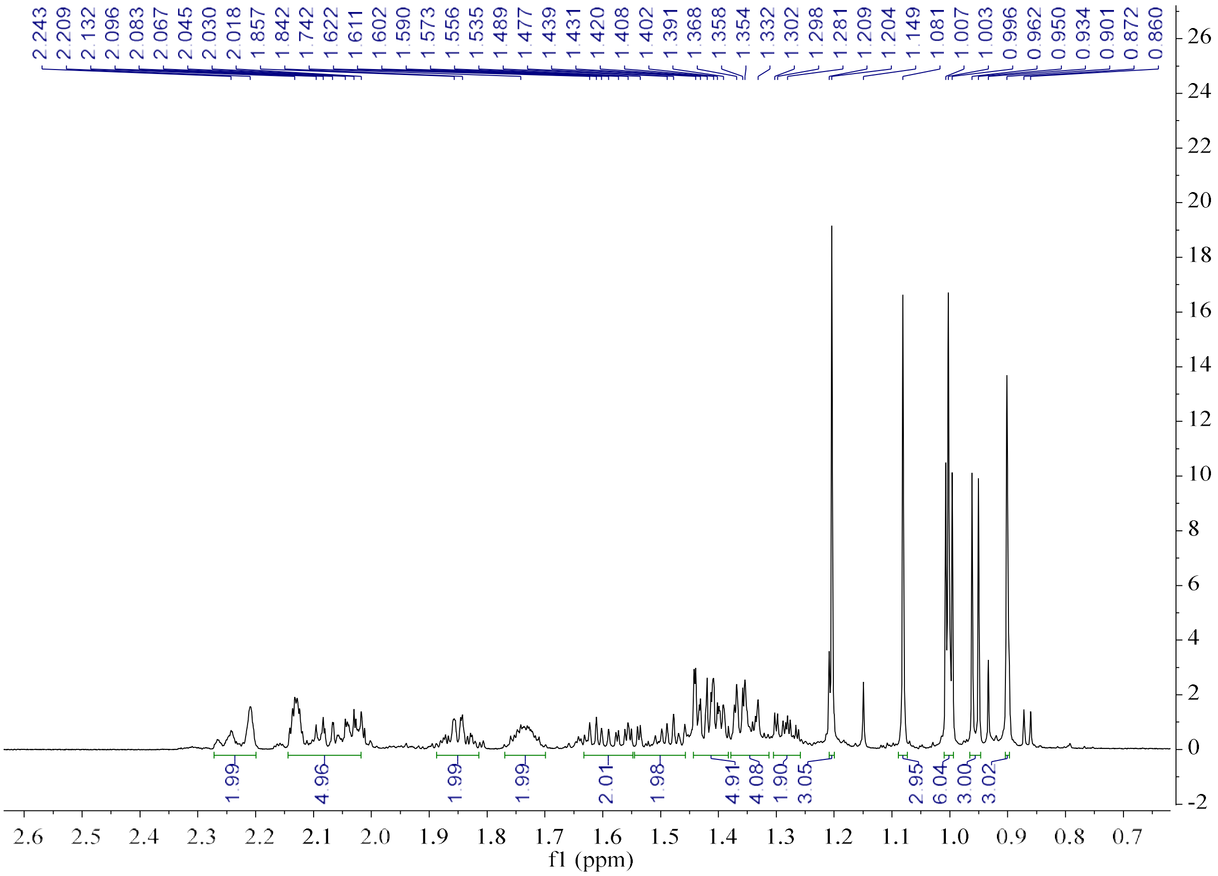

**Figure S23b.** $^{1}H$ NMR spectrum of **26** (600 MHz, Benzene-$d_6$).

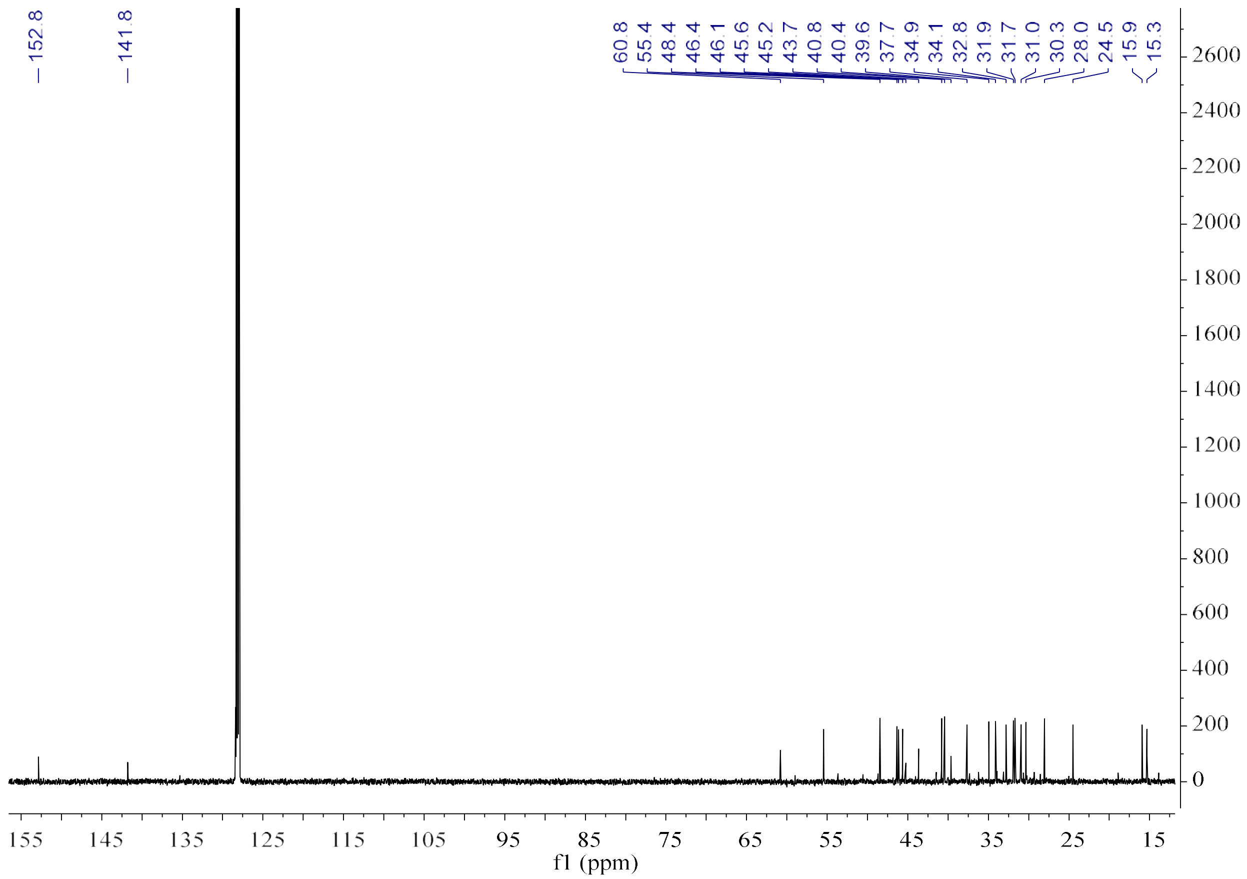

**Figure S23c.** $^{13}$C NMR spectrum of **26** (150 MHz, Benzene-$d_6$).

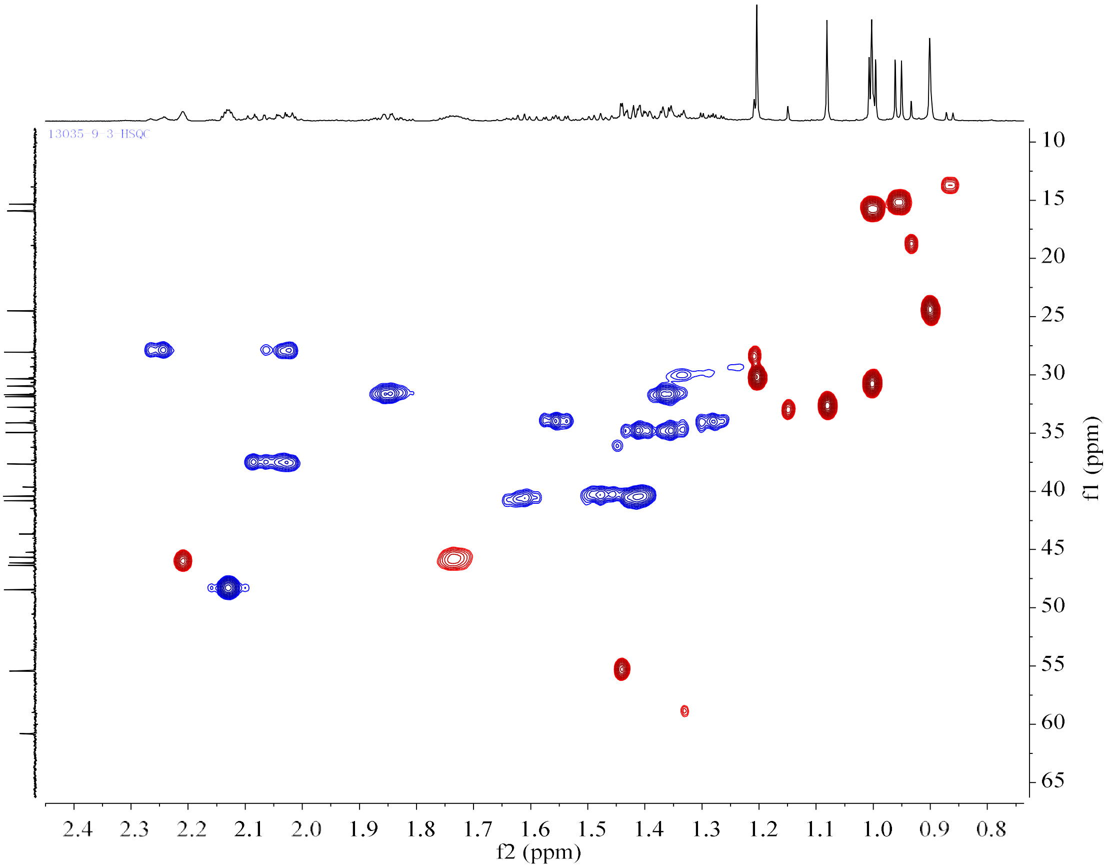

**Figure S23d.** HSQC spectrum of **26** (150 MHz/600 MHz, Benzene-$d_6$).

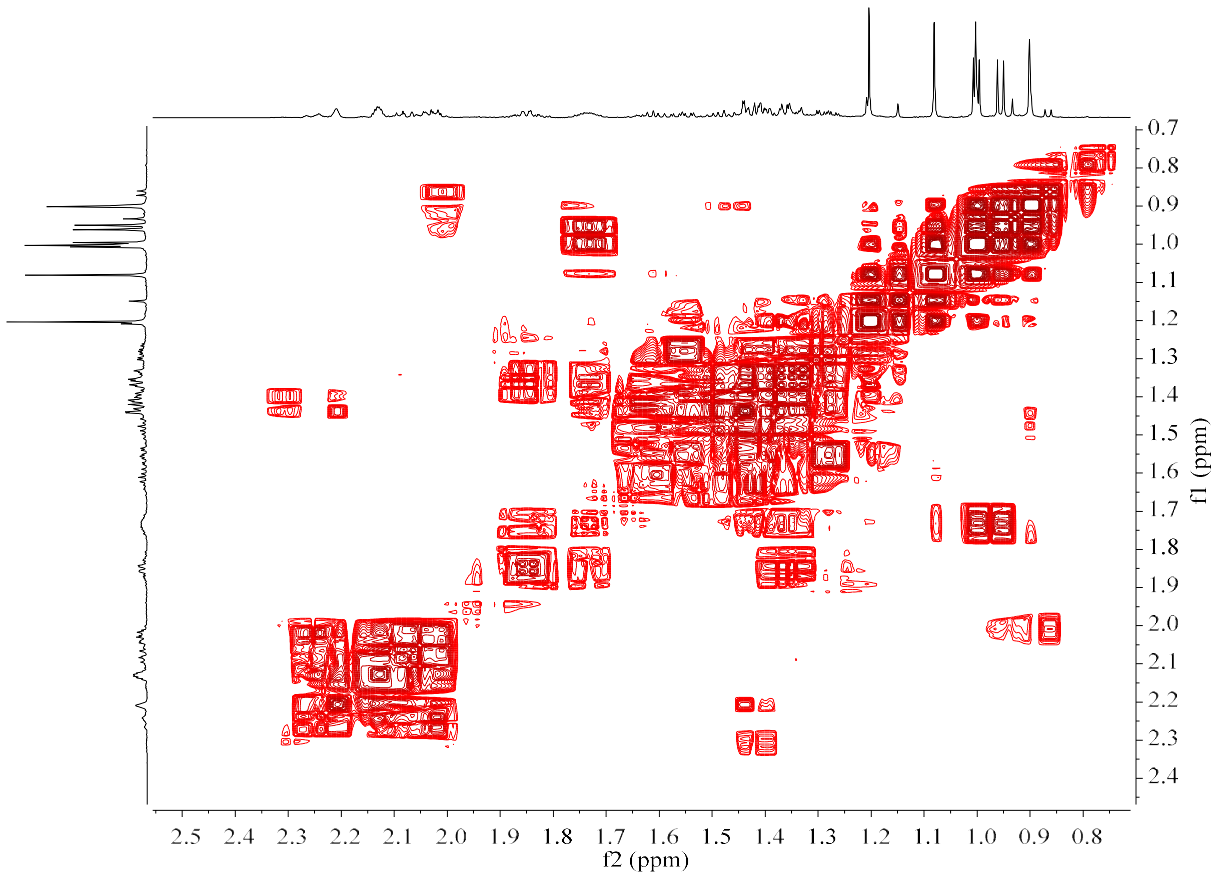

**Figure S23e.** $^1H–^1H$ COSY spectrum of **26** (600 MHz, Benzene-$d_6$).

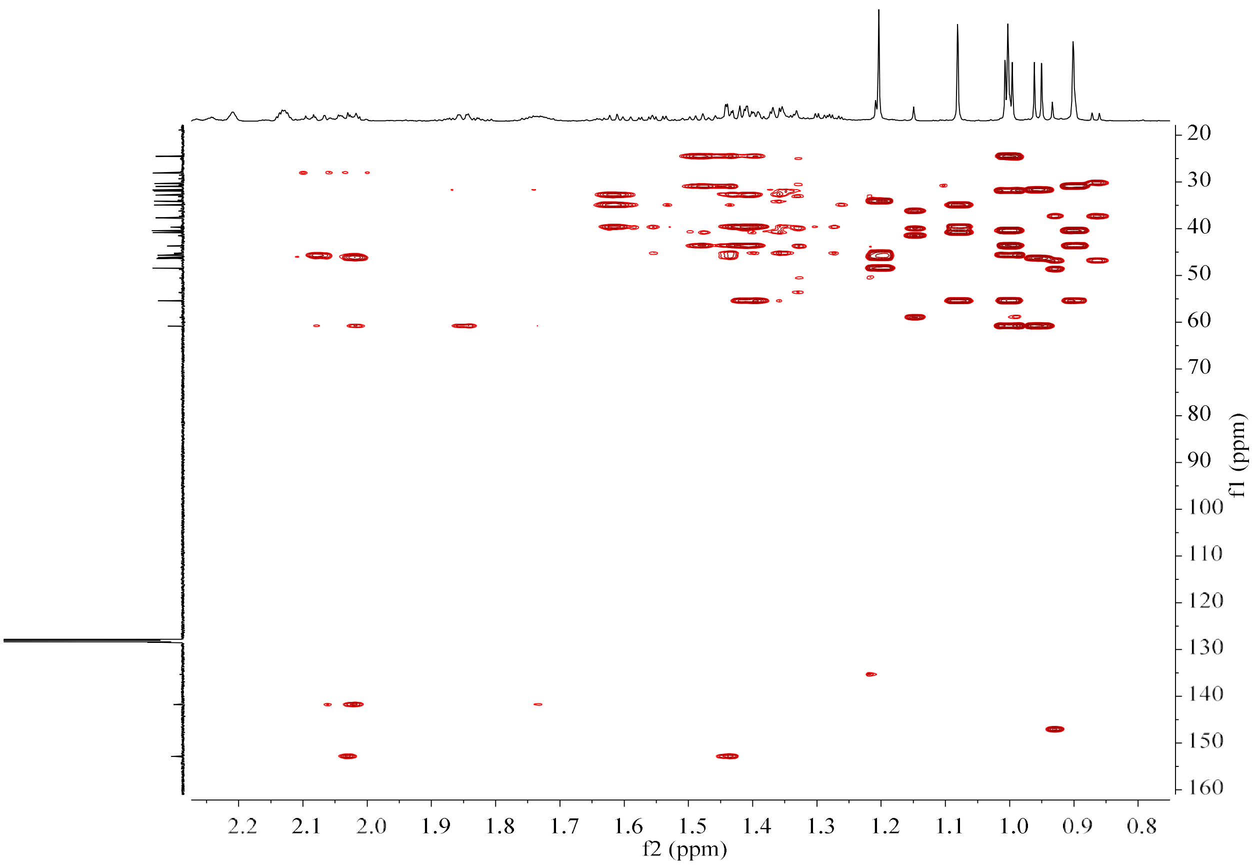

**Figure S23f.** HMBC spectrum of **26** (150 MHz/600 MHz, Benzene-$d_6$).

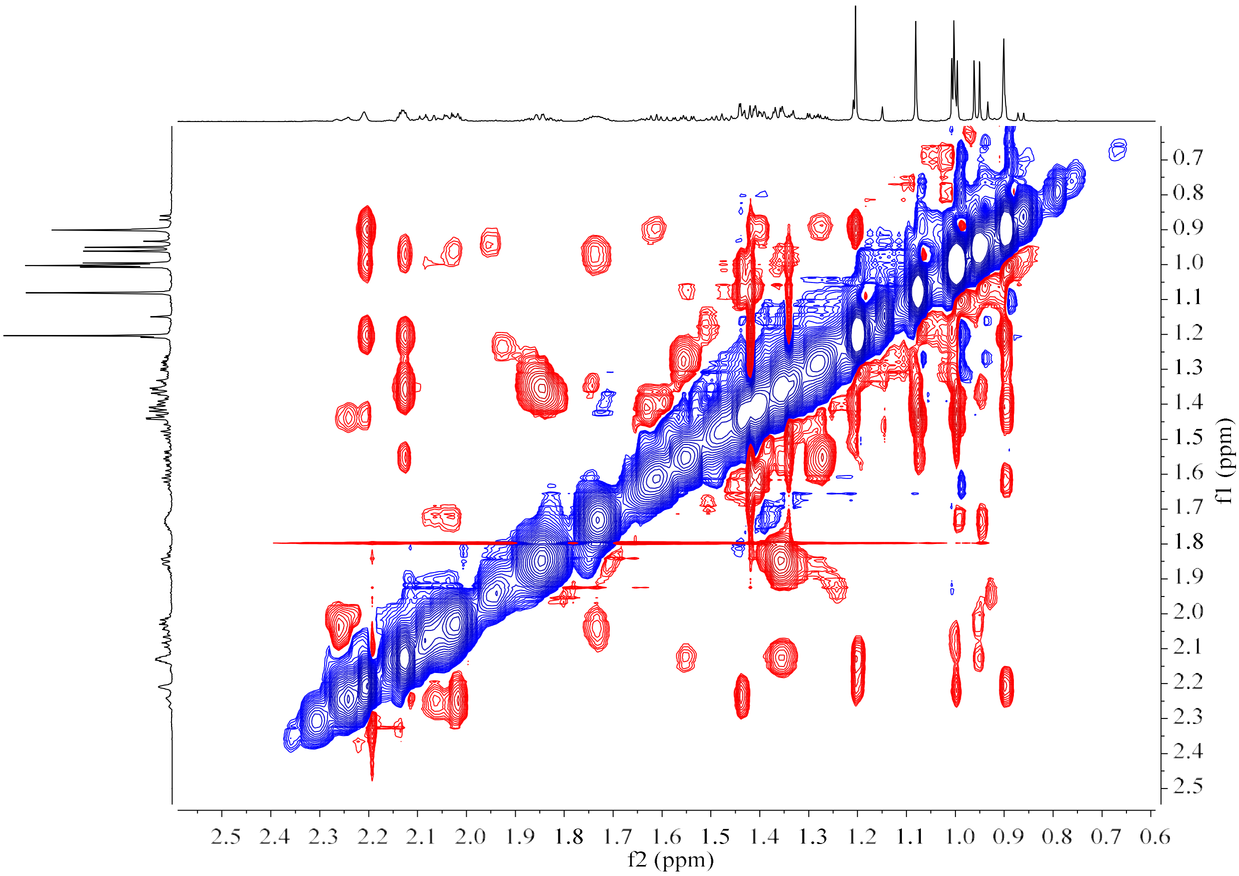

**Figure S23g.** NOESY spectrum of **26** (600 MHz, Benzene-$d_6$).

**Figure S24.** HR-EI-MS and NMR spectrum of **27**.

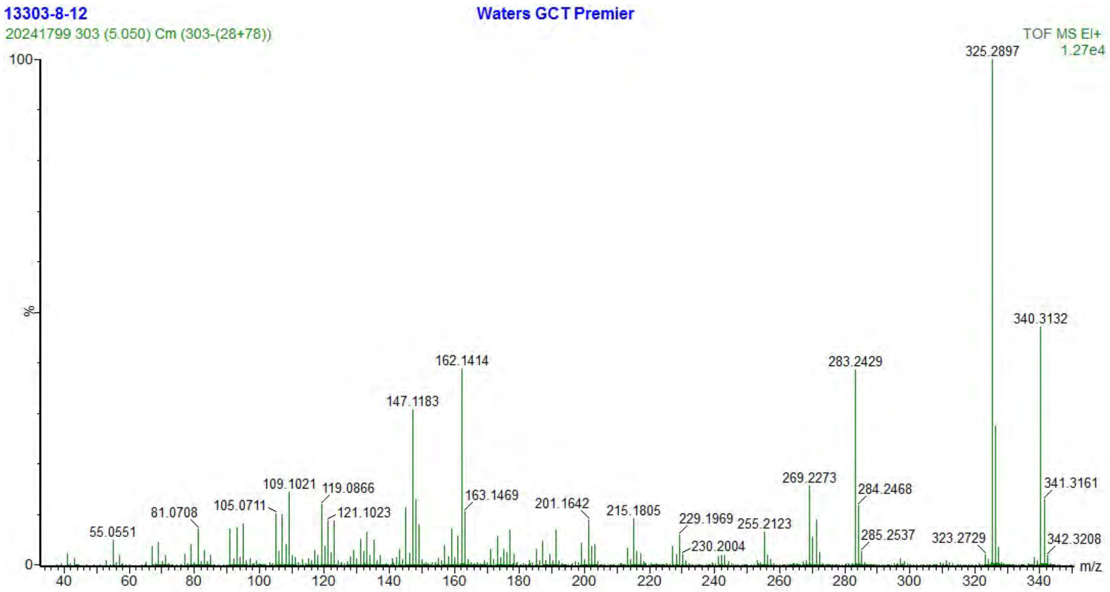

**Figure S24a.** HR-EI-MS spectrum of **27**.

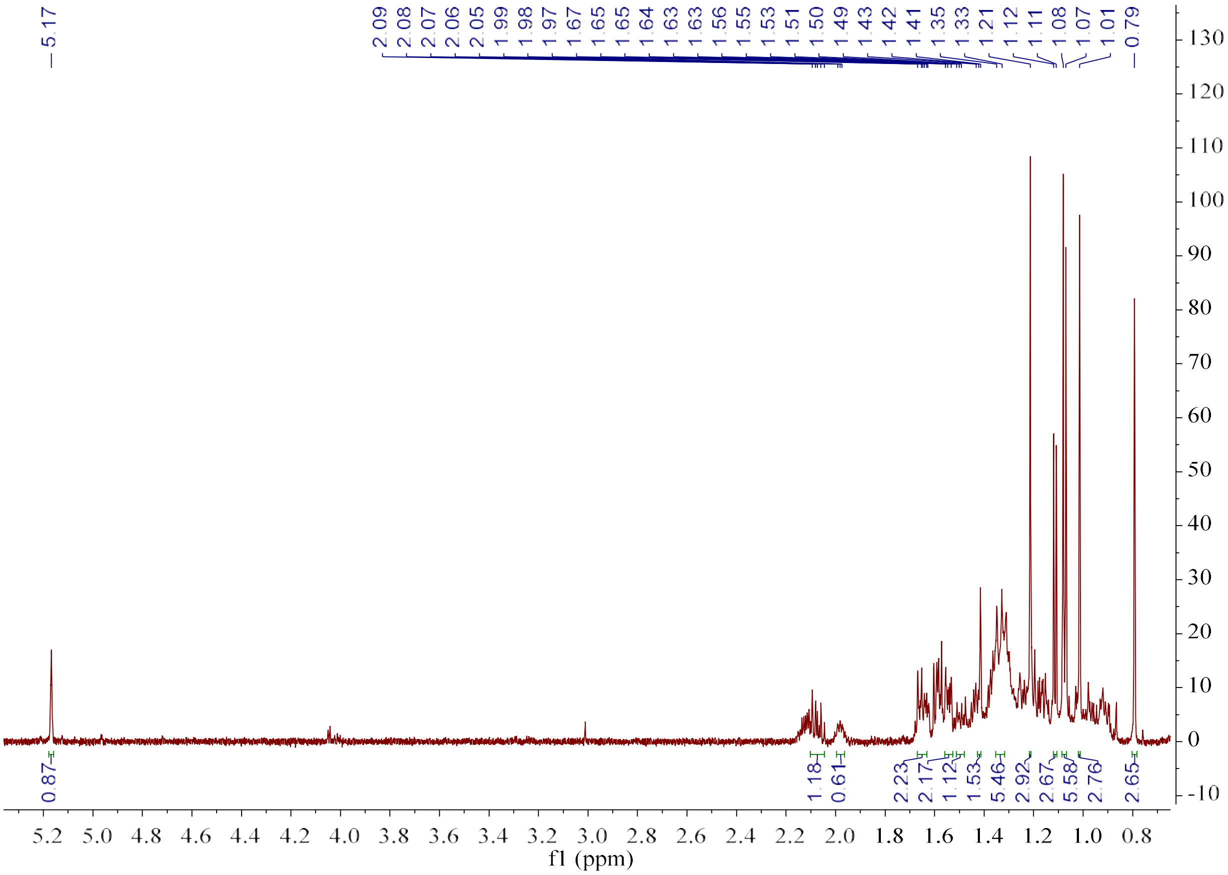

**Figure S24b.** [1]H NMR spectrum of **27** (600 MHz, Benzene-$d_6$).

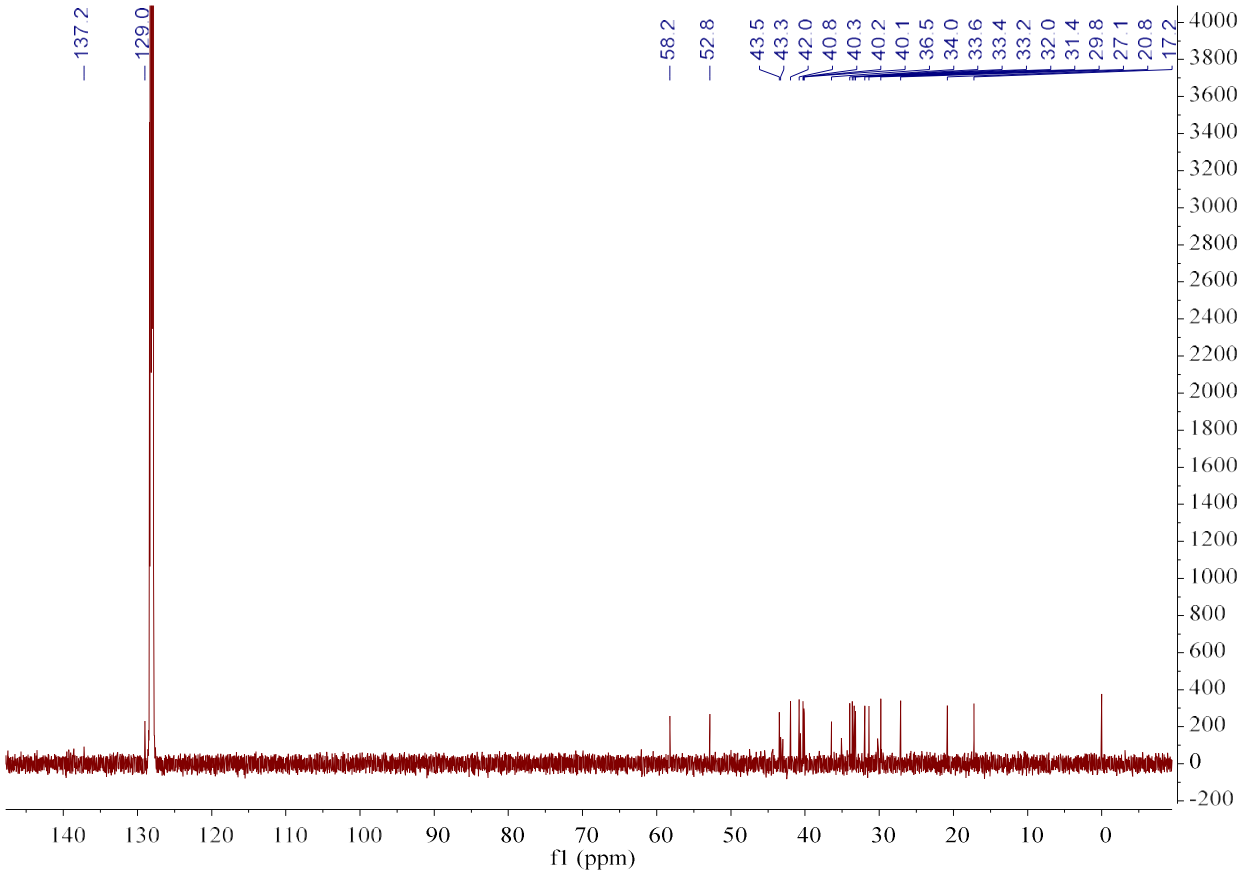

**Figure S24c.** $^{13}$C NMR spectrum of **27** (150 MHz, Benzene-$d_6$).

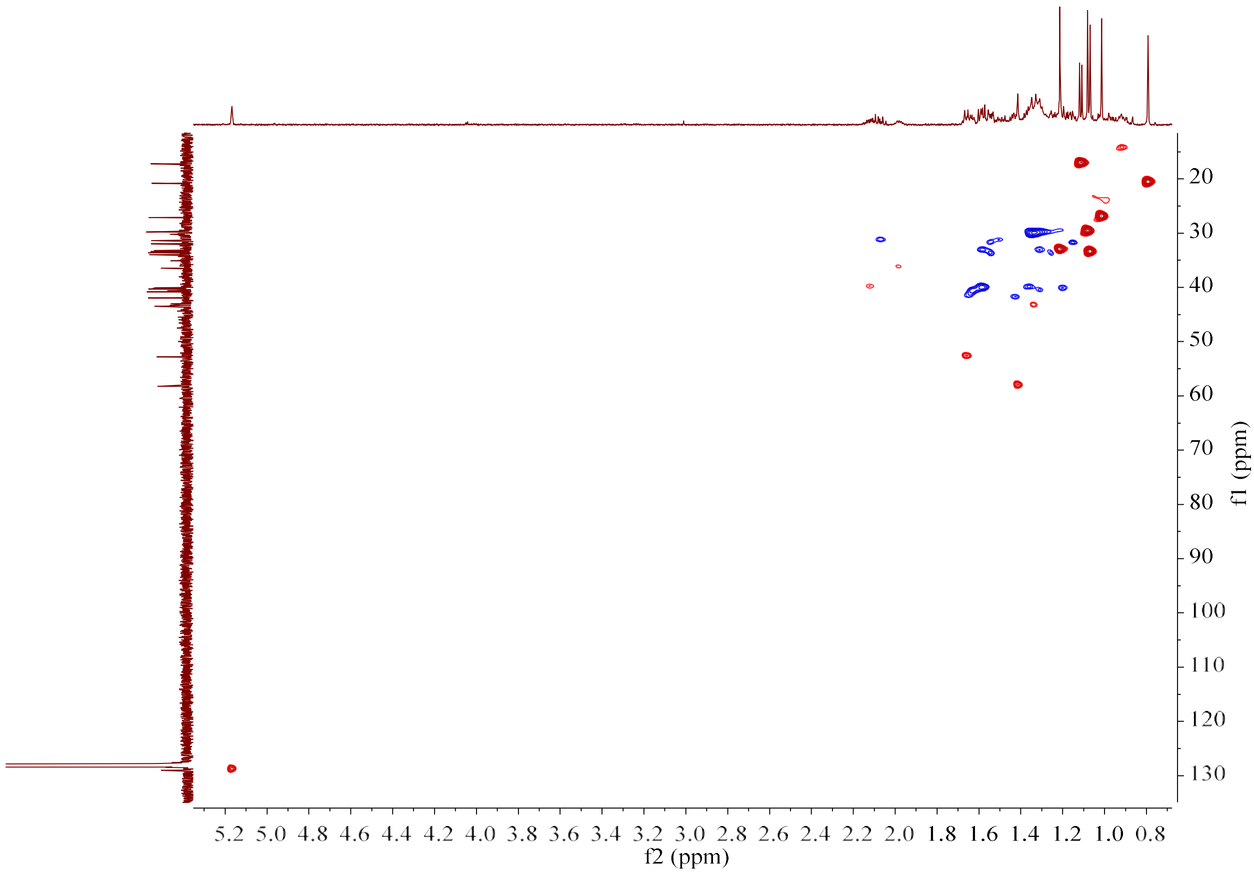

**Figure S24d.** HSQC spectrum of **27** (150 MHz/600 MHz, Benzene-$d_6$).

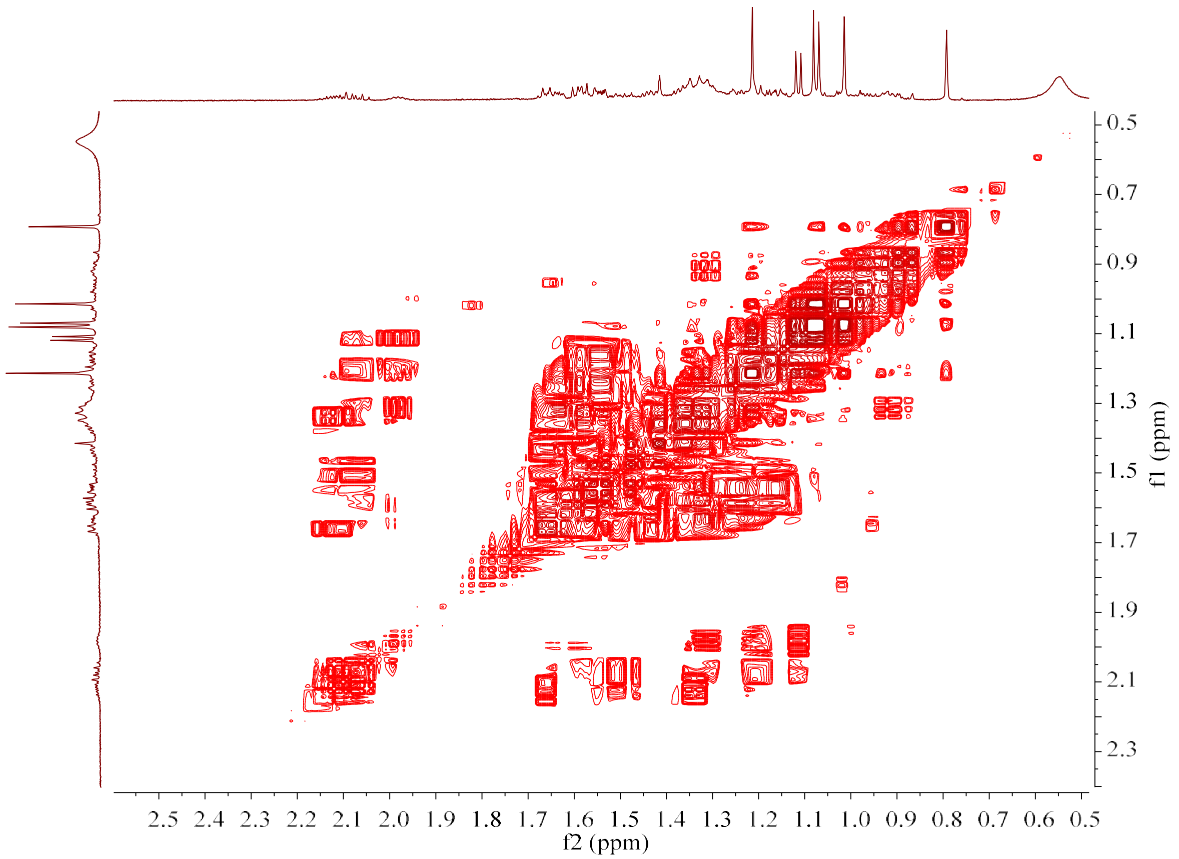

**Figure S24e.** $^1H$–$^1H$ COSY spectrum of **27** (600 MHz, Benzene-$d_6$).

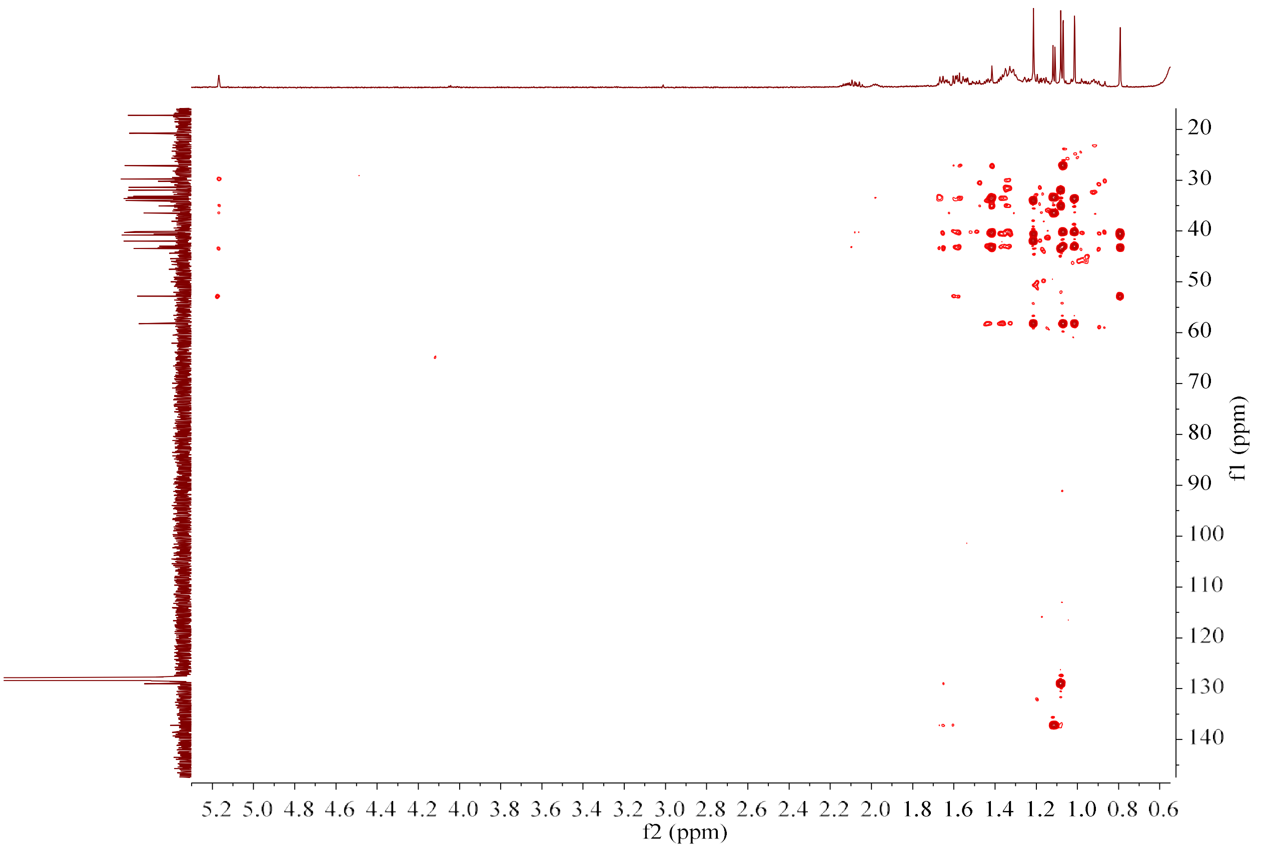

**Figure S24f.** HMBC spectrum of **27** (150 MHz/600 MHz, Benzene-$d_6$).

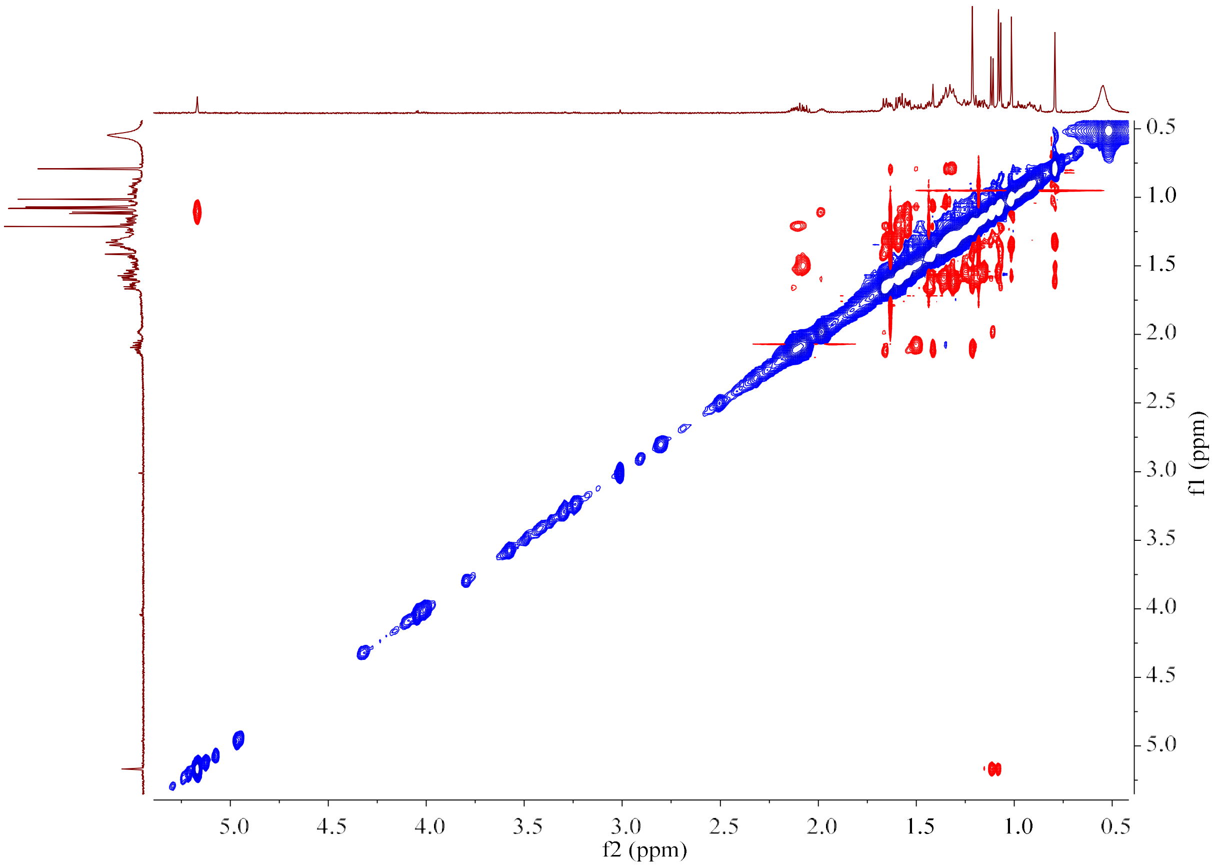

**Figure S24g.** NOESY spectrum of **27** (600 MHz, Benzene-$d_6$).

**Figure S25.** HR-EI-MS and NMR spectrum of **28**.

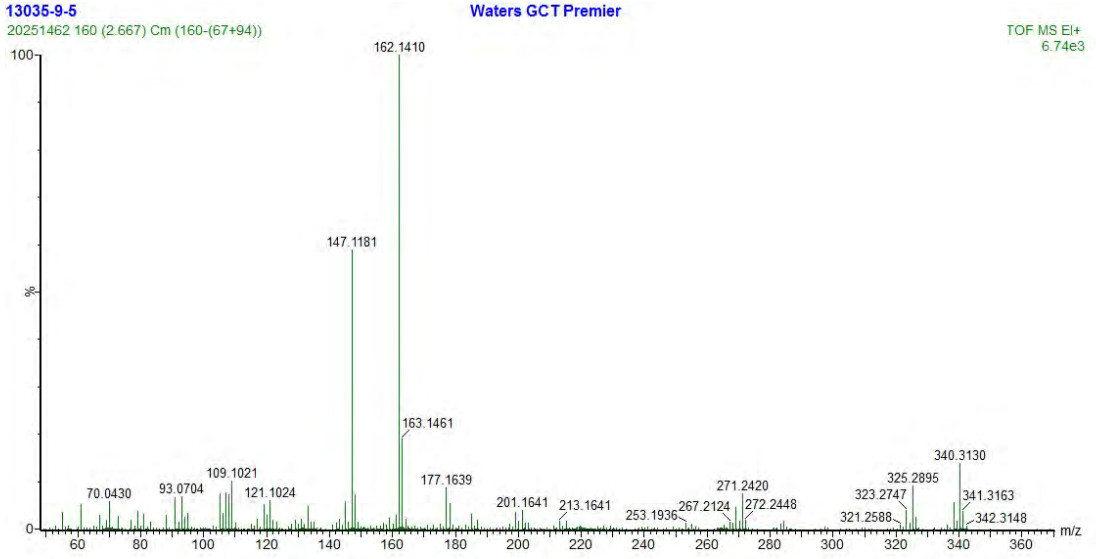

**Figure S25a.** HR-EI-MS spectrum of **28**.

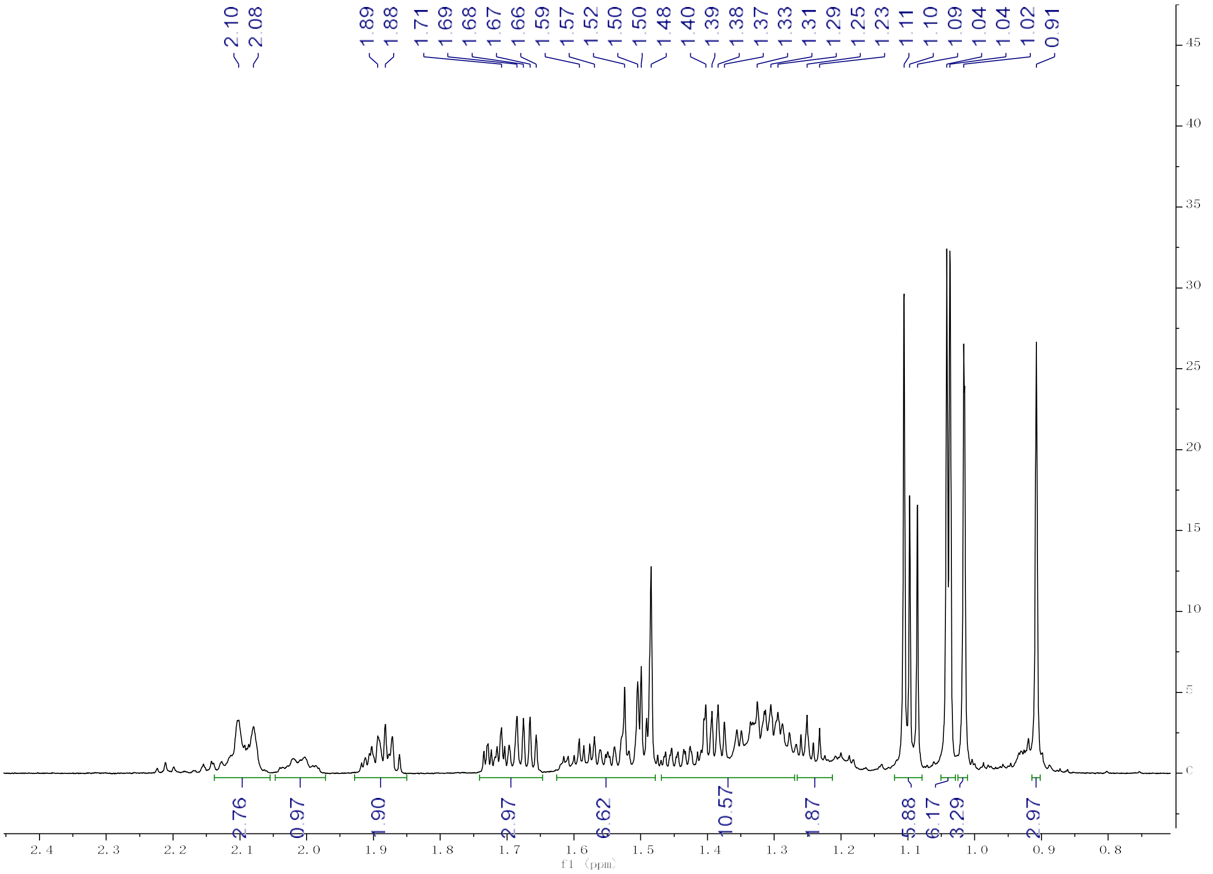

**Figure S25b.** [1]H NMR spectrum of **28** (600 MHz, Benzene-$d_6$).

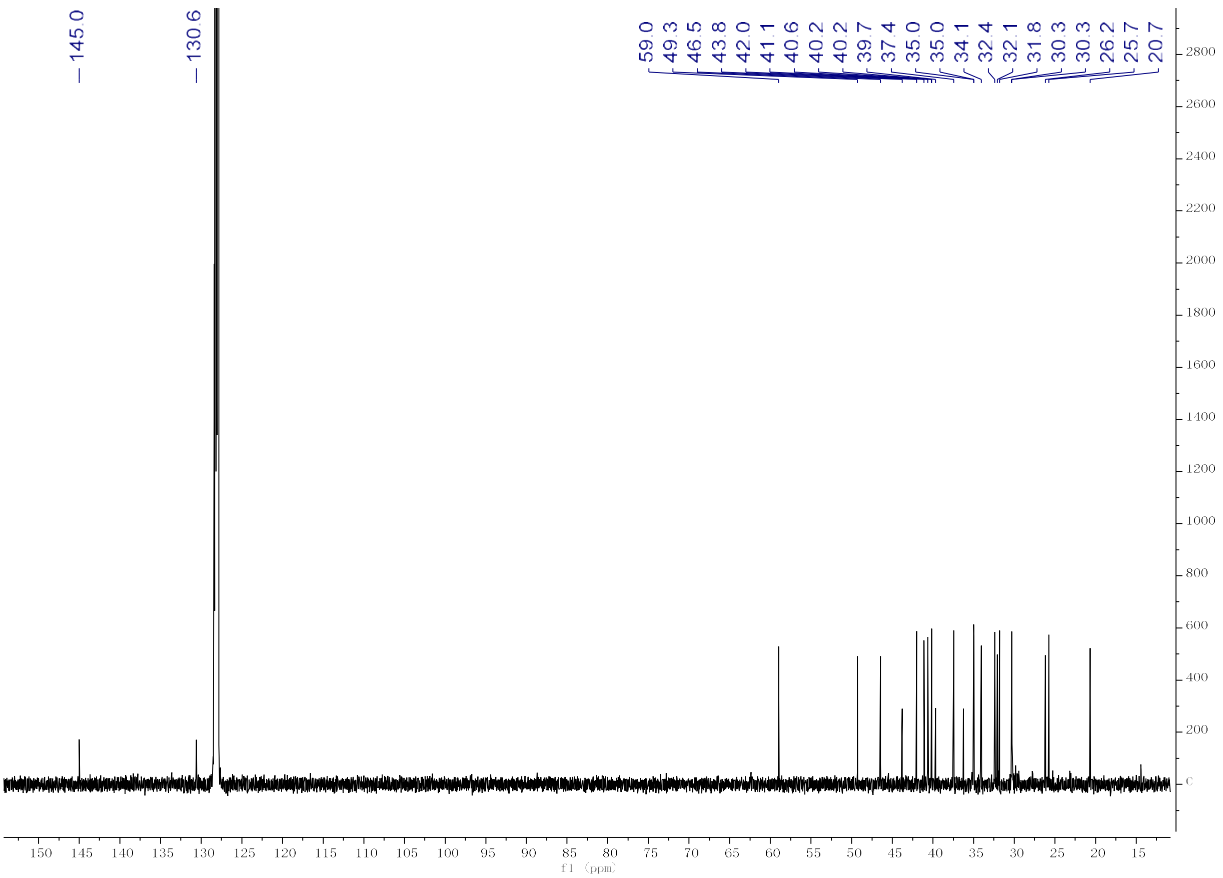

**Figure S25c.** $^{13}C$ NMR spectrum of **28** (150 MHz, Benzene-$d_6$).

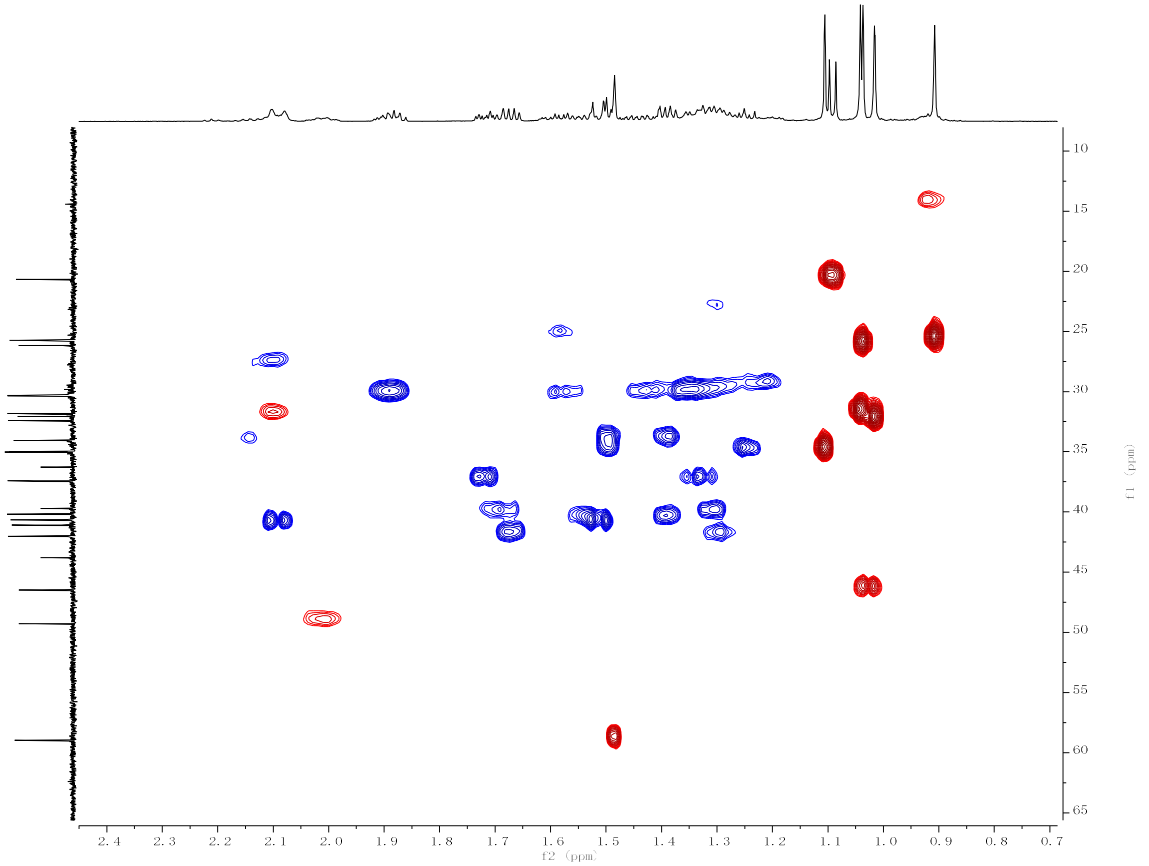

**Figure S25d.** HSQC spectrum of **28** (150 MHz/600 MHz, Benzene-$d_6$).

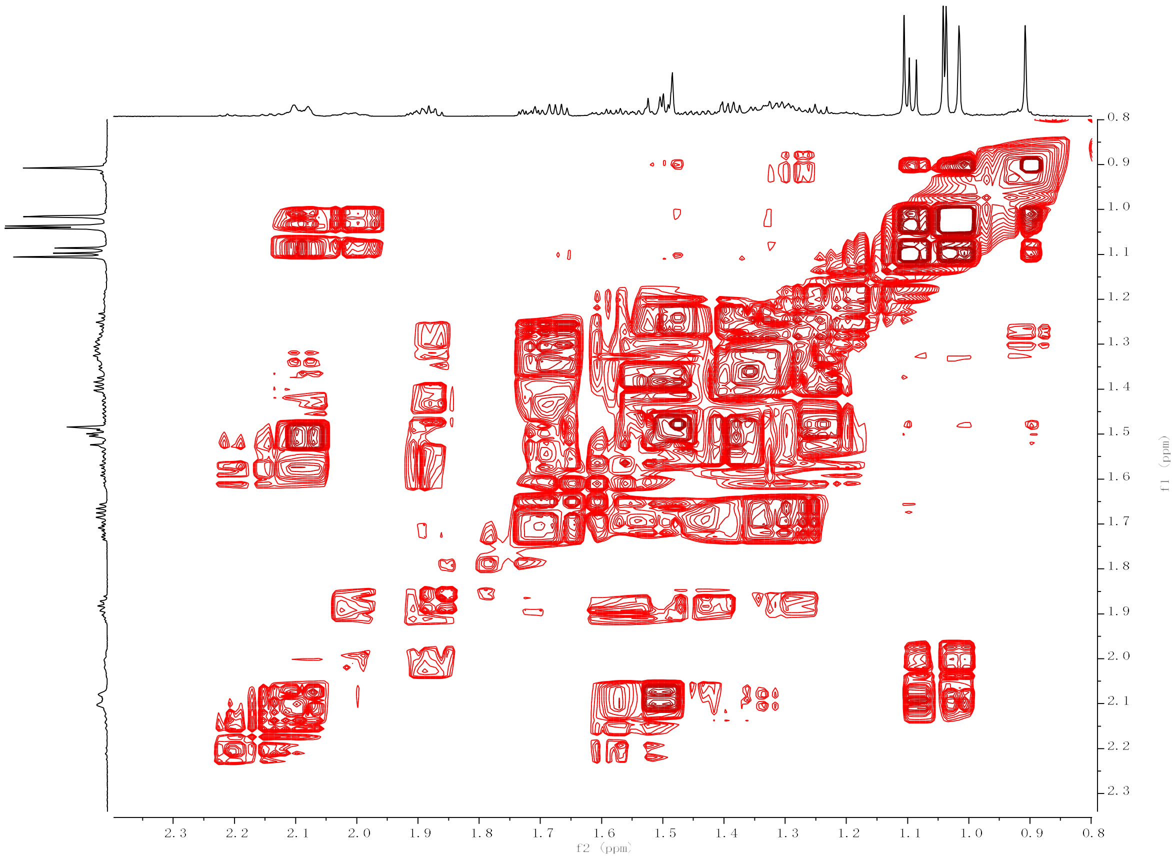

**Figure S25e.** $^{1}H$–$^{1}H$ COSY spectrum of **28** (600 MHz, Benzene-$d_6$).

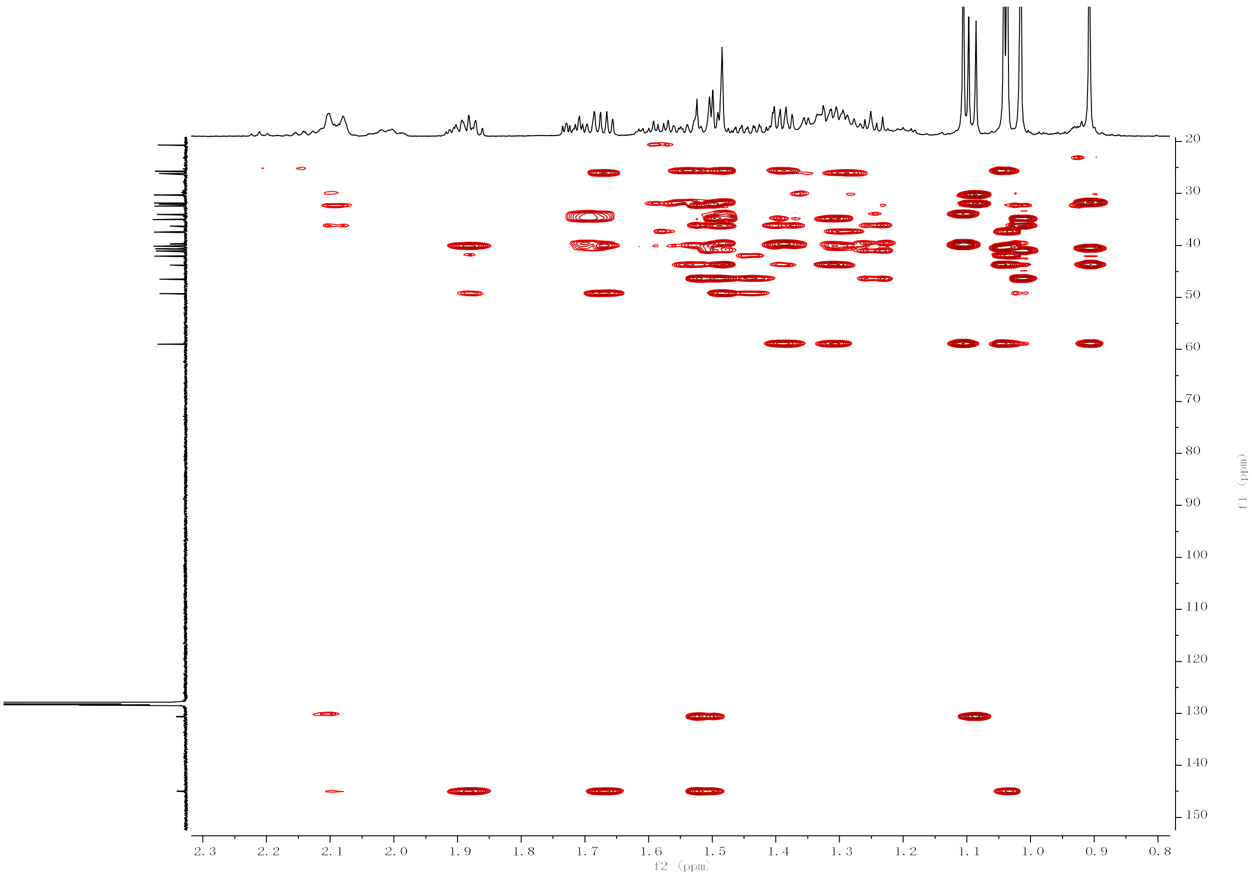

**Figure S25f.** HMBC spectrum of **28** (150 MHz/600 MHz, Benzene-$d_6$).

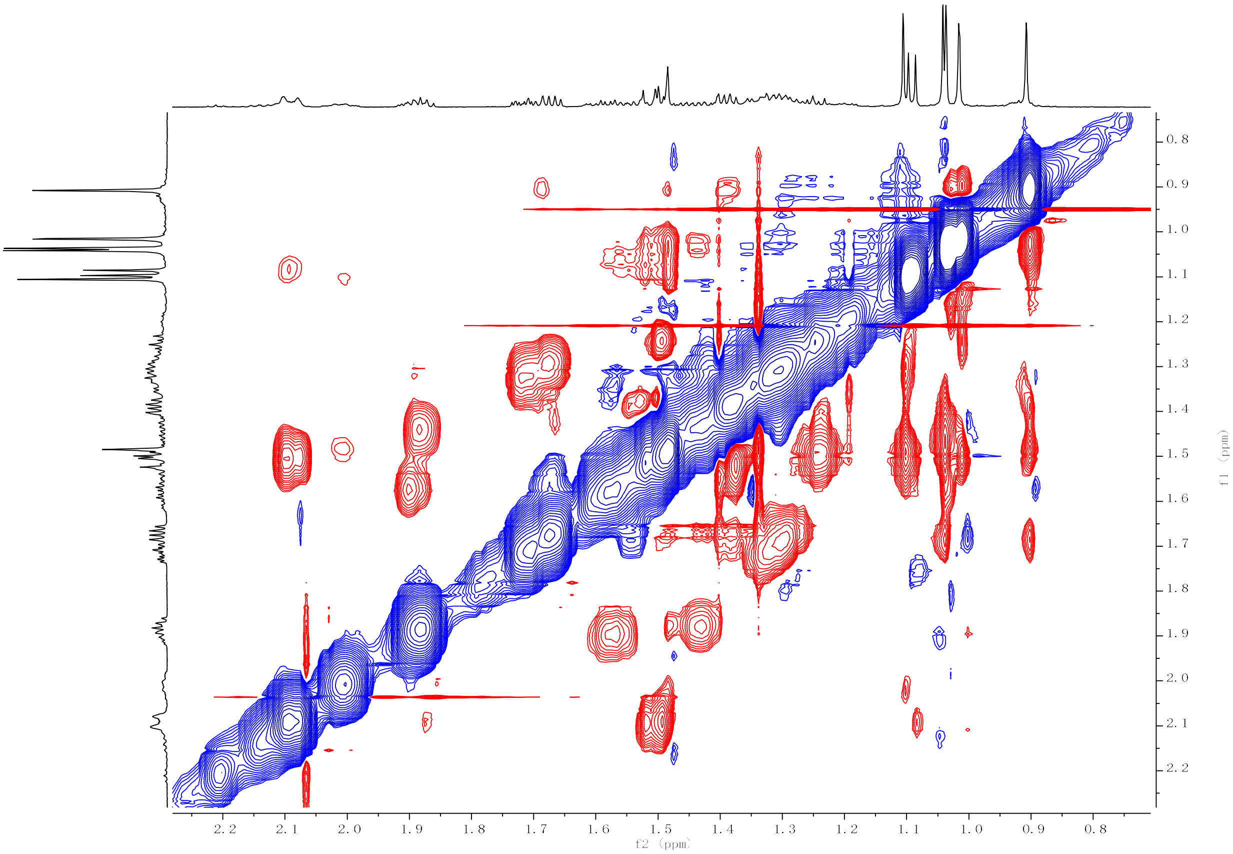

**Figure S25g.** NOESY spectrum of **28** (600 MHz, Benzene-$d_6$).

**Figure S26.** HR-EI-MS and NMR spectrum of **29**.

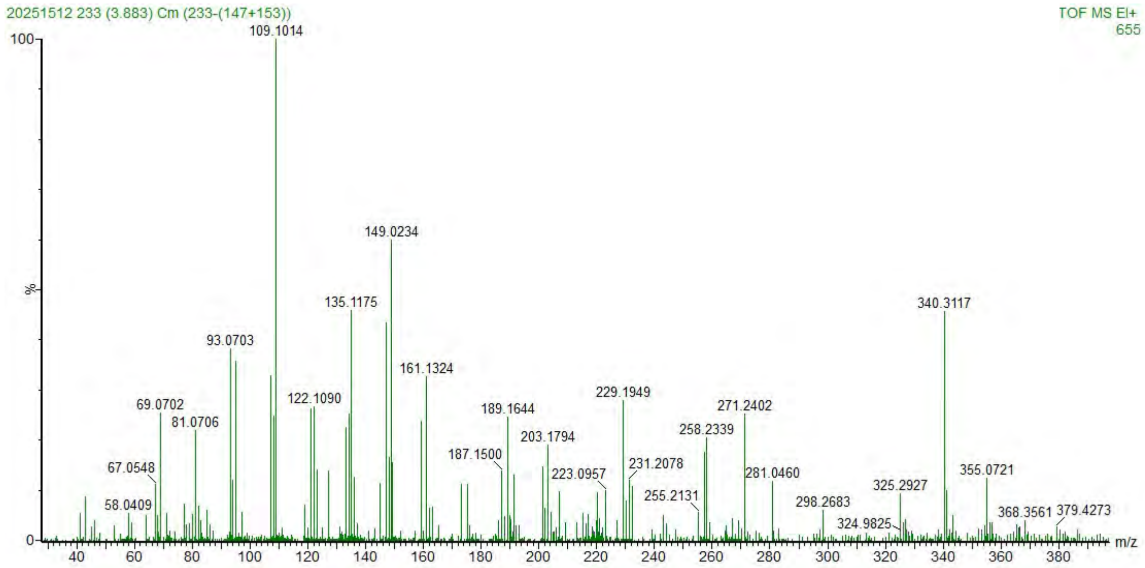

**Figure S26a.** HR-EI-MS spectrum of **29**.

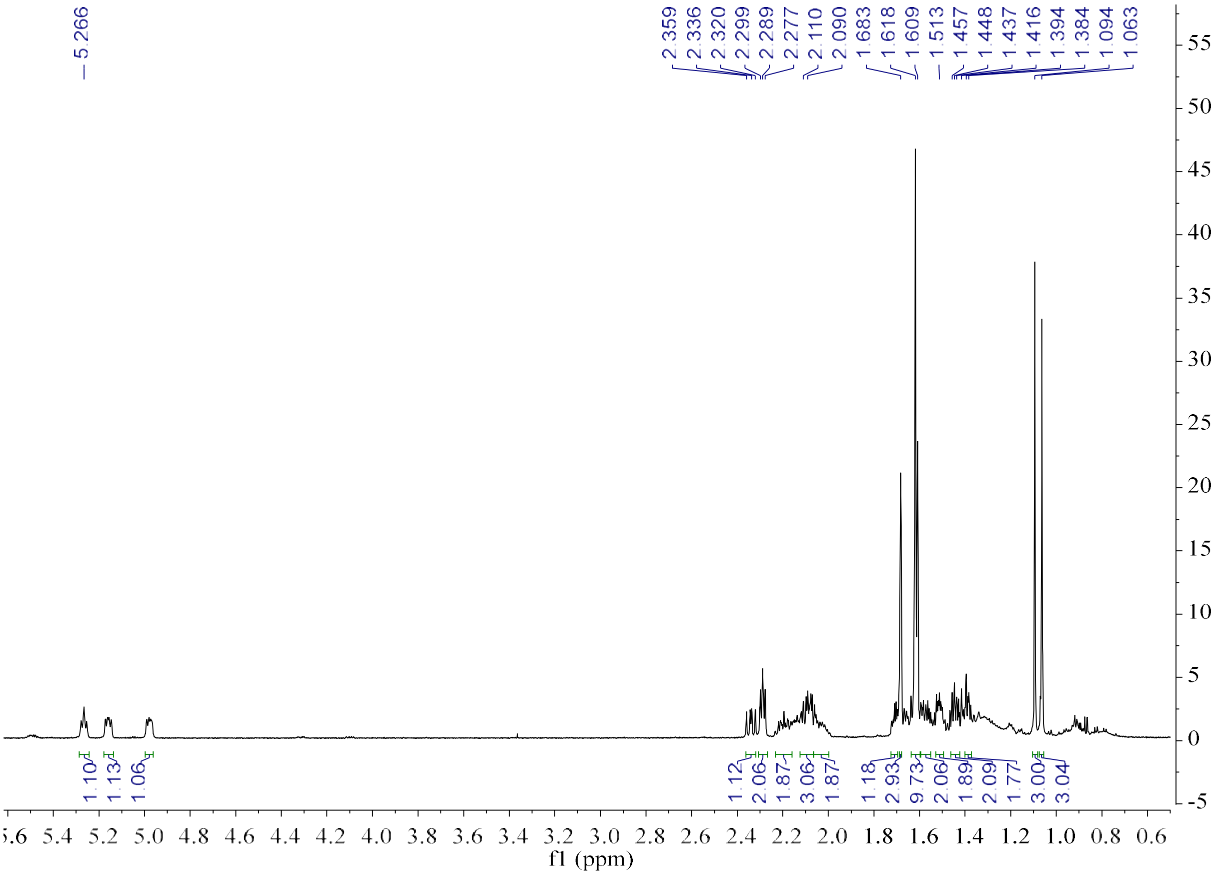

**Figure S26b.** [1]H NMR spectrum of **29** (600 MHz, Benzene-$d_6$).

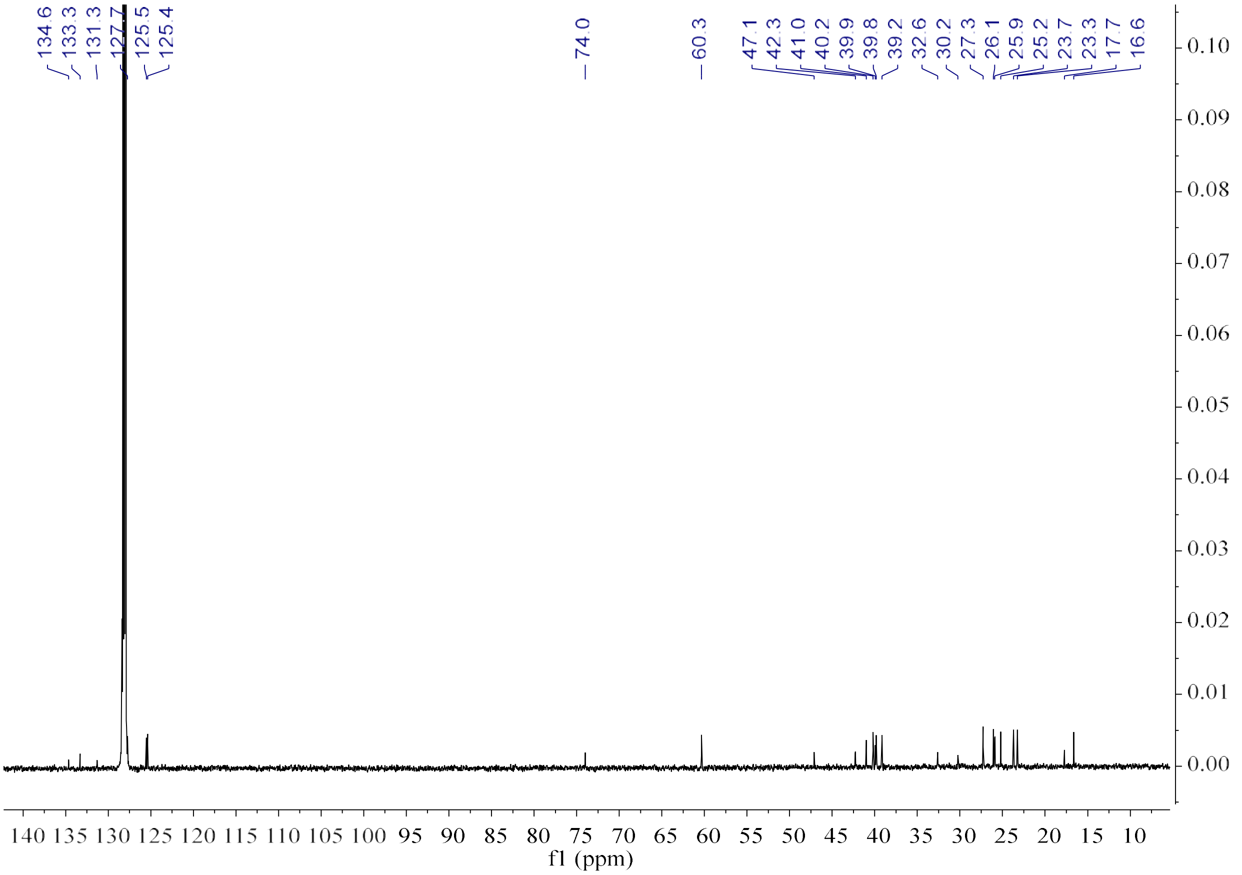

**Figure S26c.** $^{13}$C NMR spectrum of **29** (150 MHz, Benzene-$d_6$).

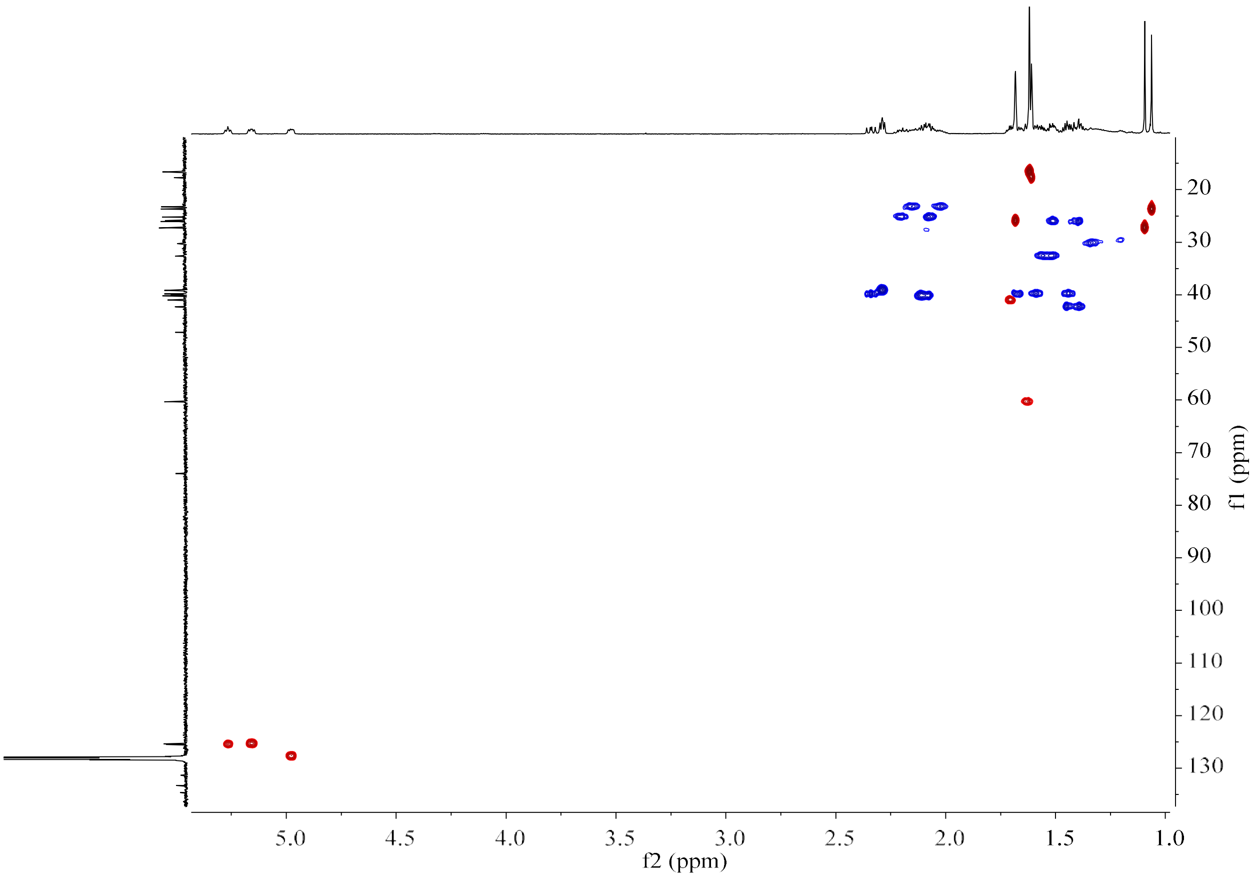

**Figure S26d.** HSQC spectrum of **29** (150 MHz/600 MHz, Benzene-$d_6$).

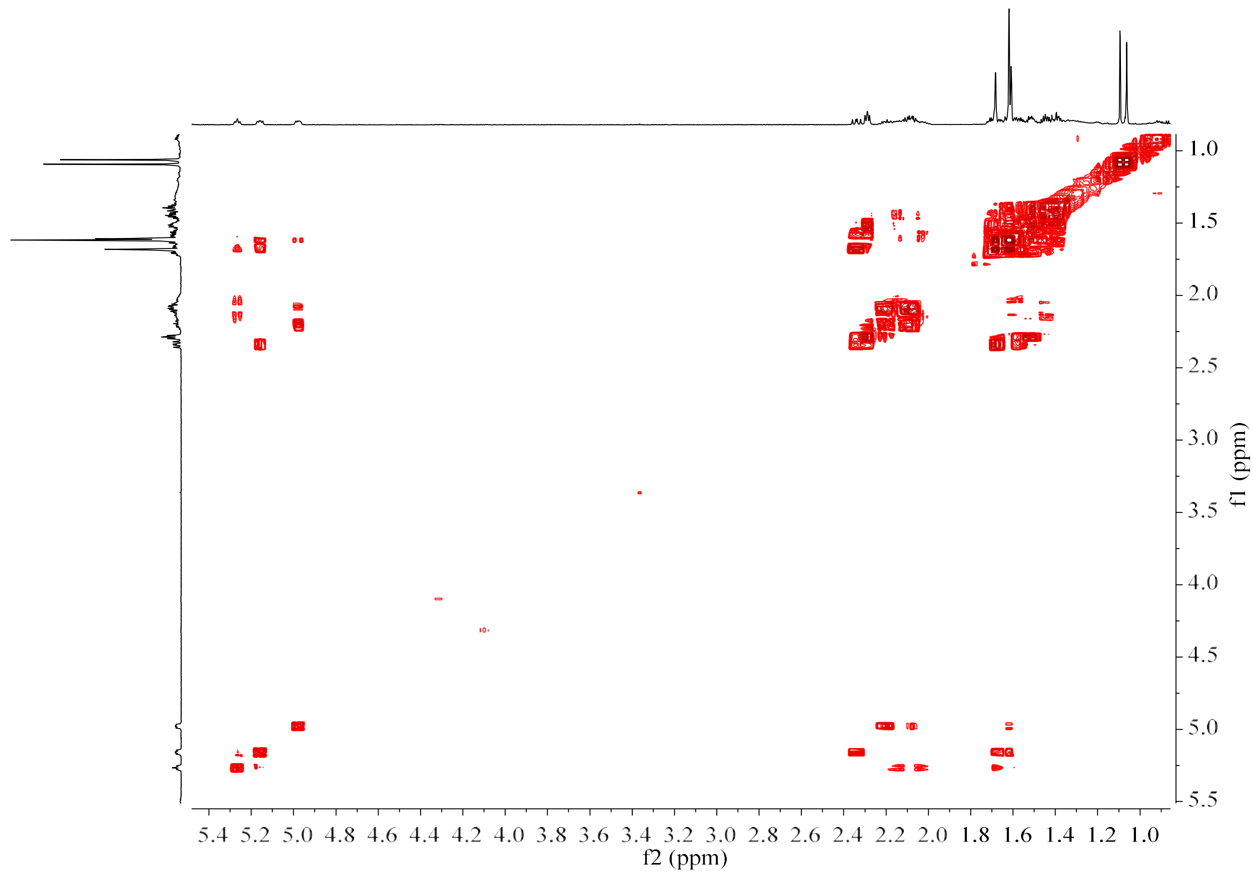

**Figure S26e.** $^{1}H$–$^{1}H$ COSY spectrum of **29** (600 MHz, Benzene-$d_6$).

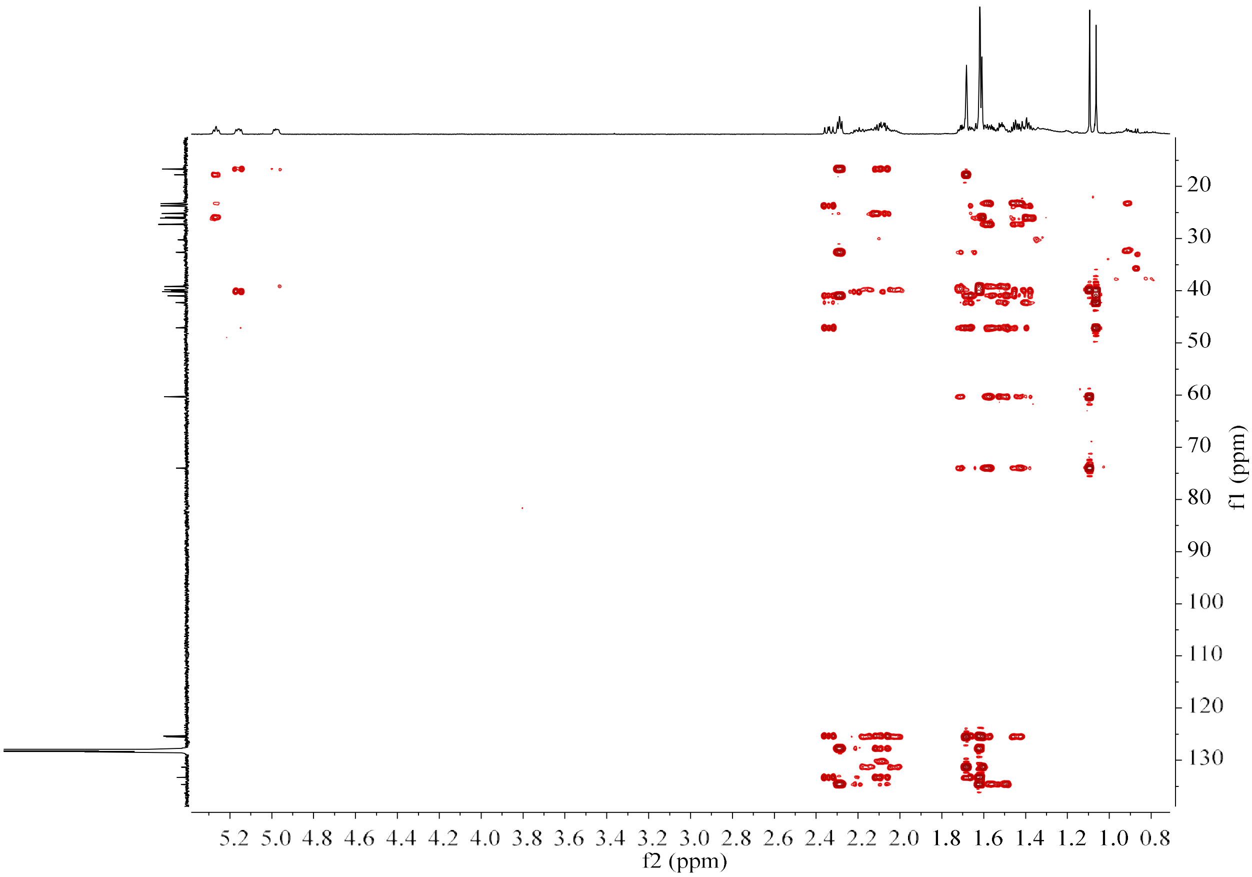

**Figure S26f.** HMBC spectrum of **29** (150 MHz/600 MHz, Benzene-$d_6$).

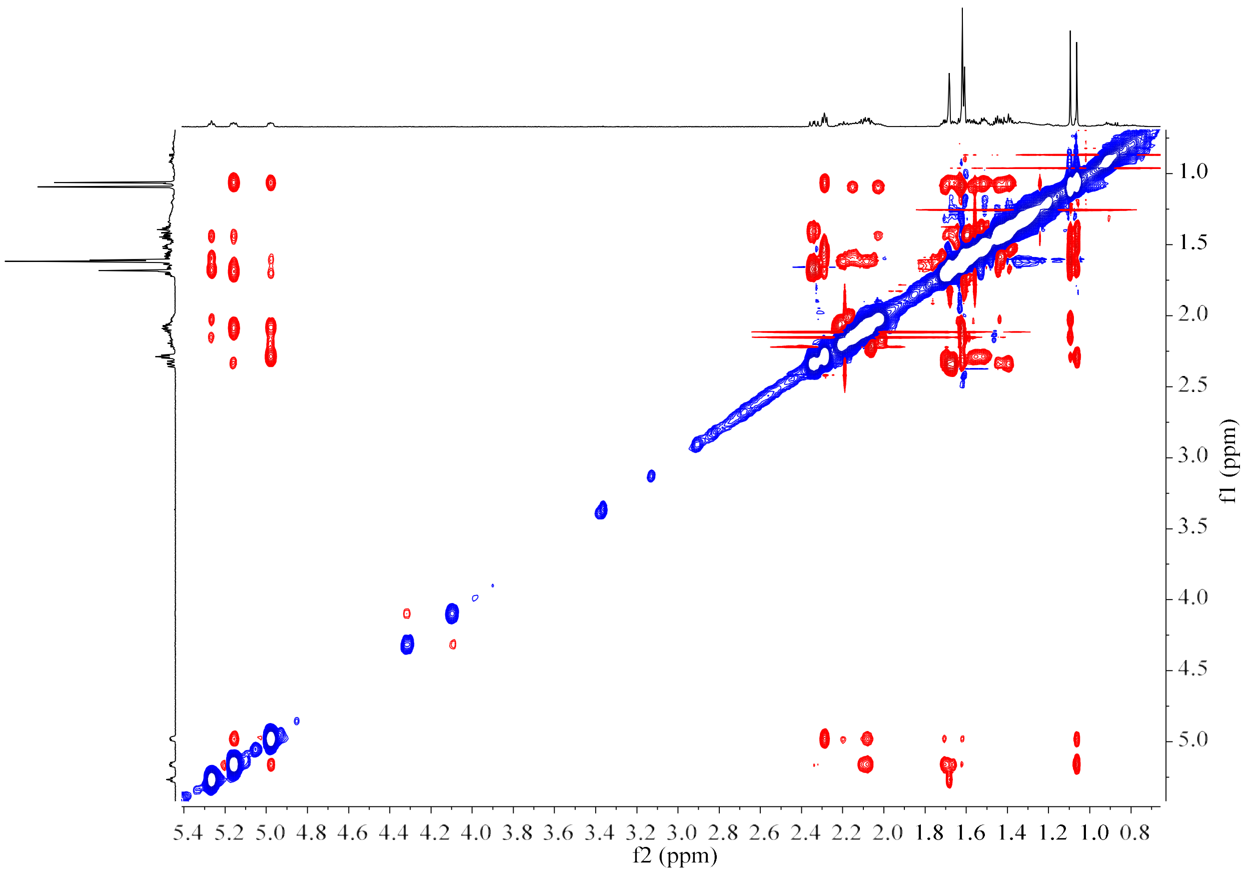

**Figure S26g.** NOESY spectrum of **29** (600 MHz, Benzene-$d_6$).

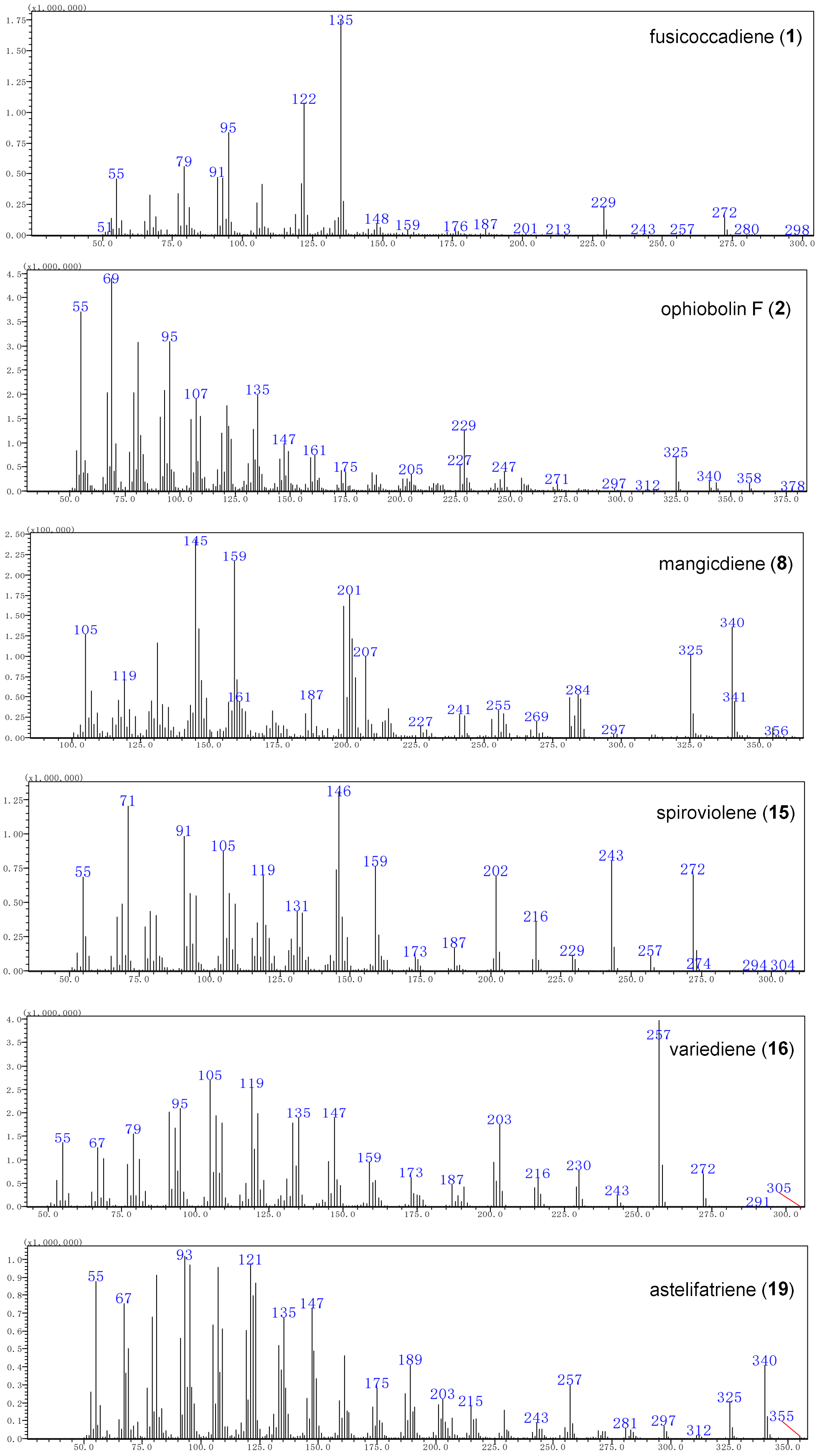

**Figure S27**. GC-MS spectra of compounds **1**, **2**, **8**, **15**, **16**, and **19**.

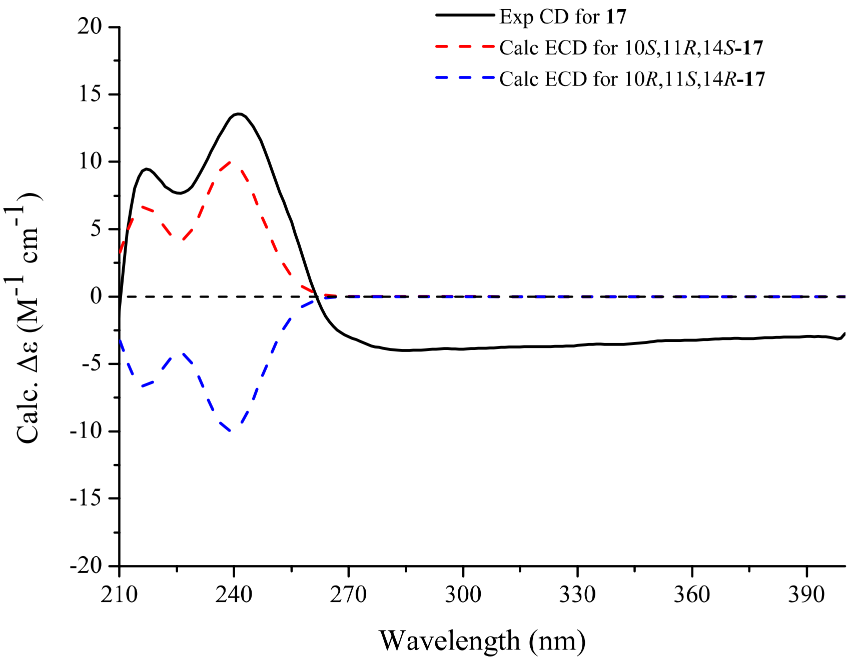

**Figure S28**. Computed ECD and experimental CD spectra of **17**.

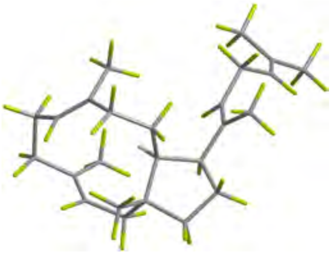

Conf.**17**-A 49.1%

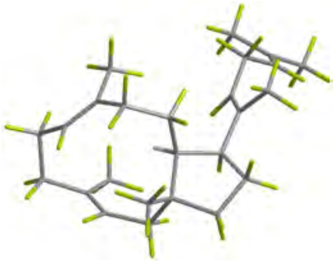

Conf.**17**-B 49%

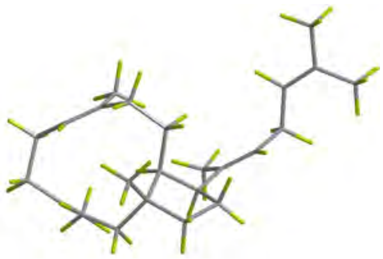

Conf.**17**-C 6.9%

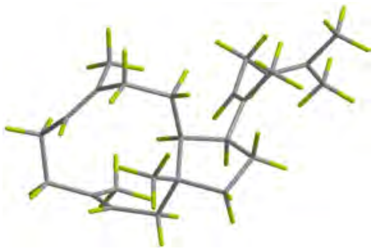

Conf.**17**-D 6.8%

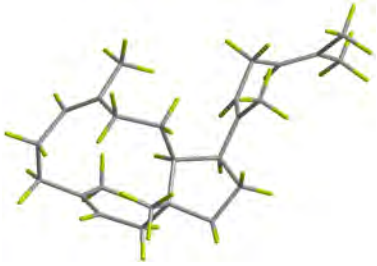

Conf.**17**-E 0.3%

**Figure S29.** DFT-optimized structures for low-energy conformers of 10*S*,11*R*,14*S*-**17** at the B3LYP/6-31G(d) level in acetonitrile (PCM).

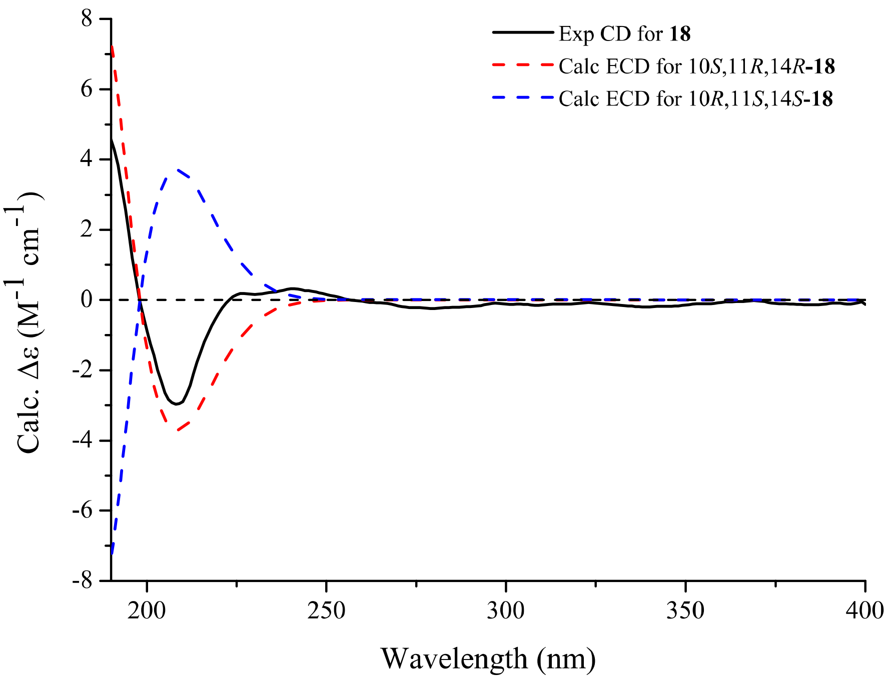

**Figure S30.** Computed ECD and experimental CD spectra of **18**.

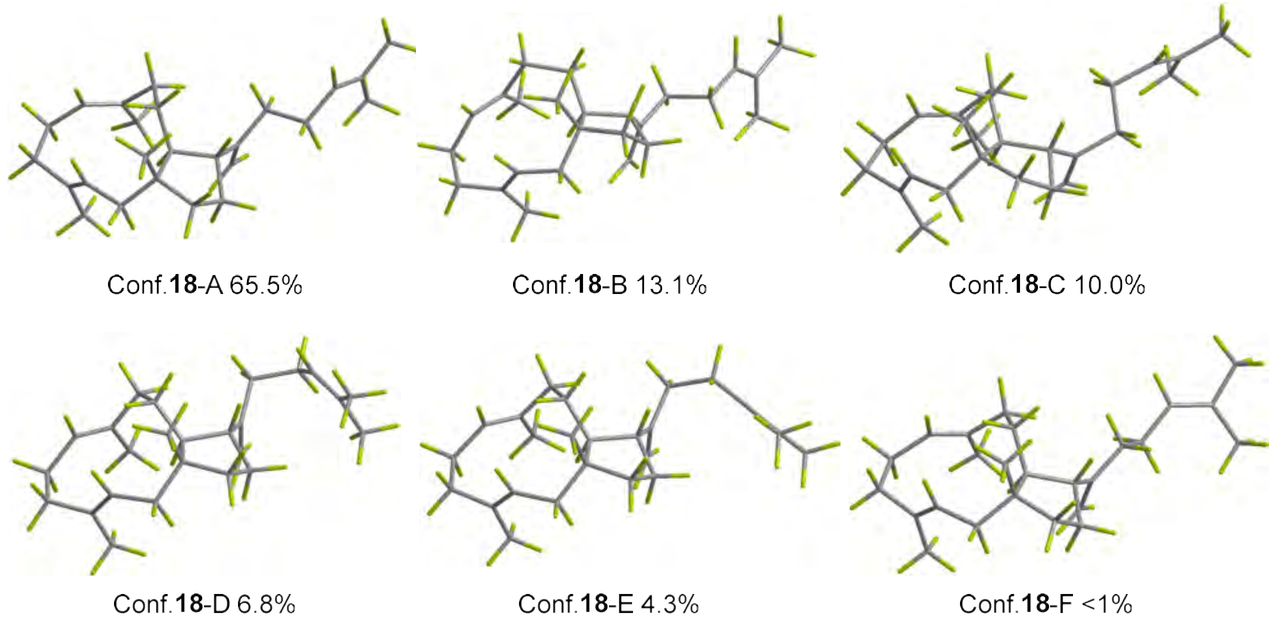

**Figure S31.** DFT-optimized structures for low-energy conformers of 10*S*,11*R*,14*R*-**18** at the B3LYP/6-31G(d) level in acetonitrile (PCM)

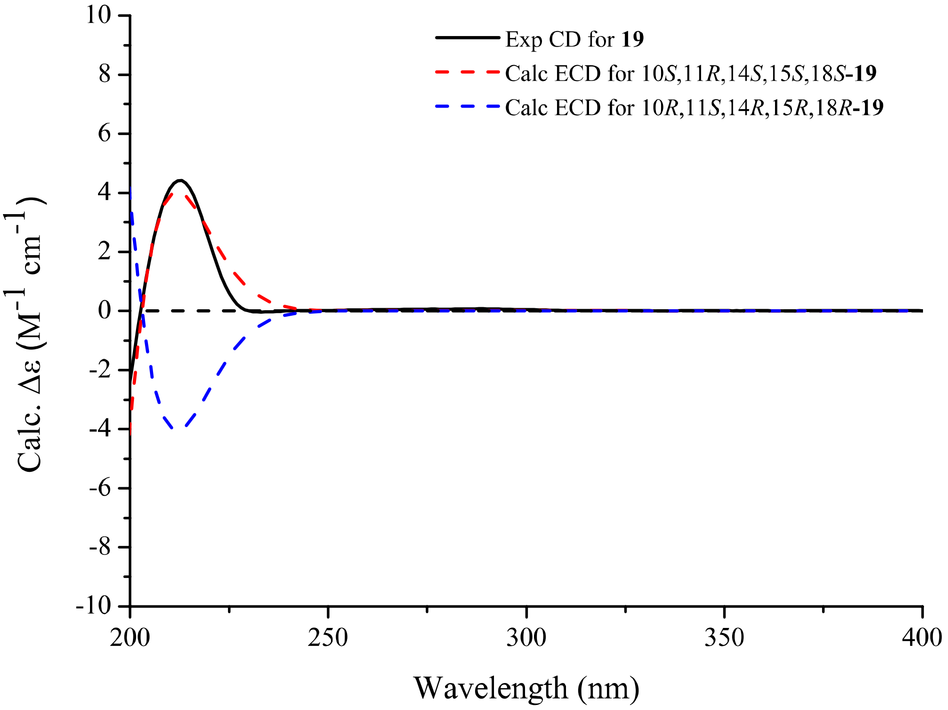

**Figure S32.** Computed ECD and experimental CD spectra of **19**.

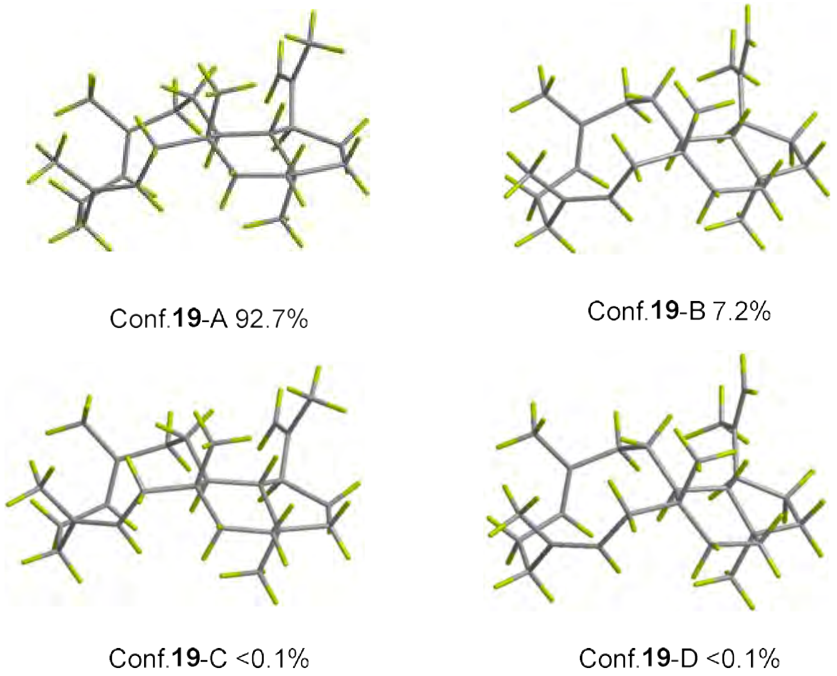

**Figure S33.** DFT-optimized structures for low-energy conformers of 10*S*,11*R*,14*S*,15*S*,18*S*-**19** at the B3LYP/6-31G(d) level in acetonitrile (PCM).

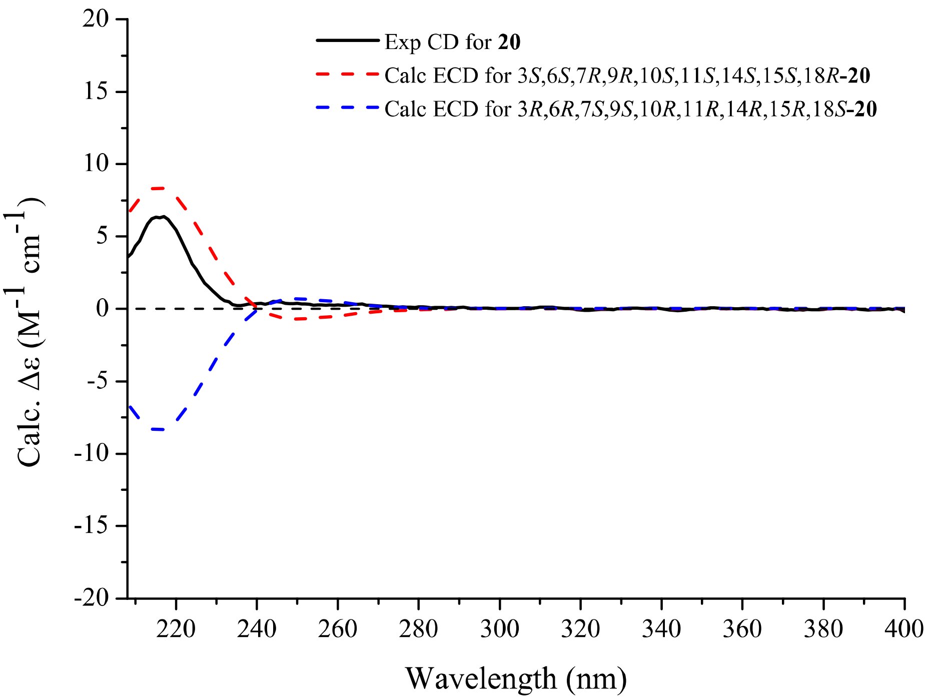

**Figure S34.** Computed ECD and experimental CD spectra of **20**.

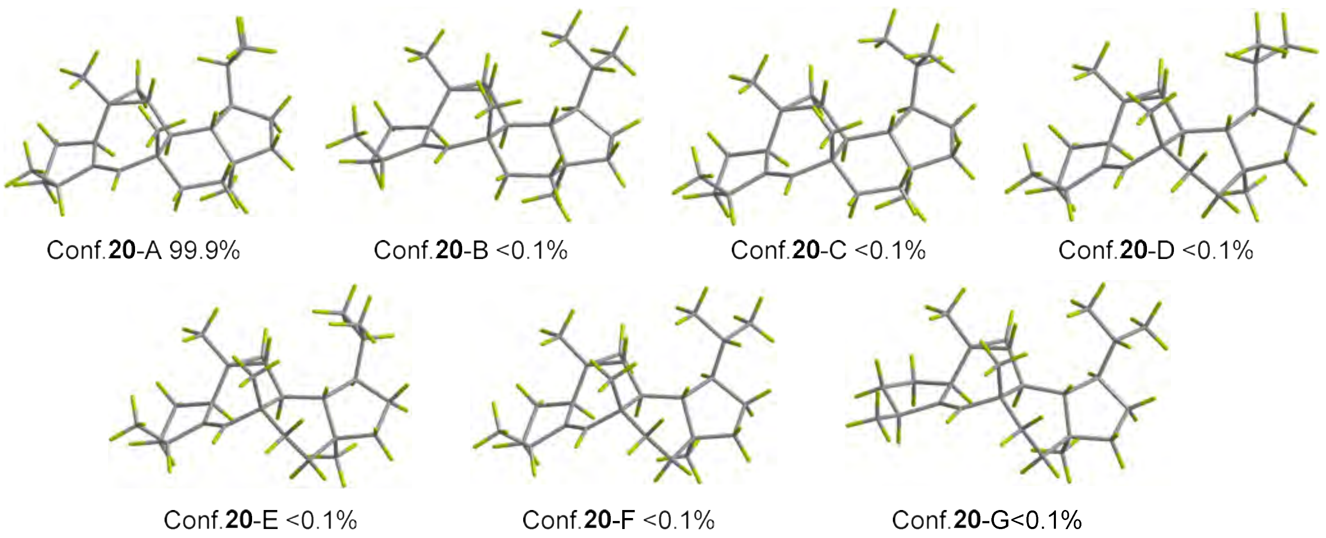

**Figure S35.** DFT-optimized structures for low-energy conformers of 3*S*,6*S*,7*R*,9*R*,10*S*,11*S*,14*S*,15*S*,18*R*-**20** at the B3LYP/6-31G(d) level in acetonitrile (PCM).

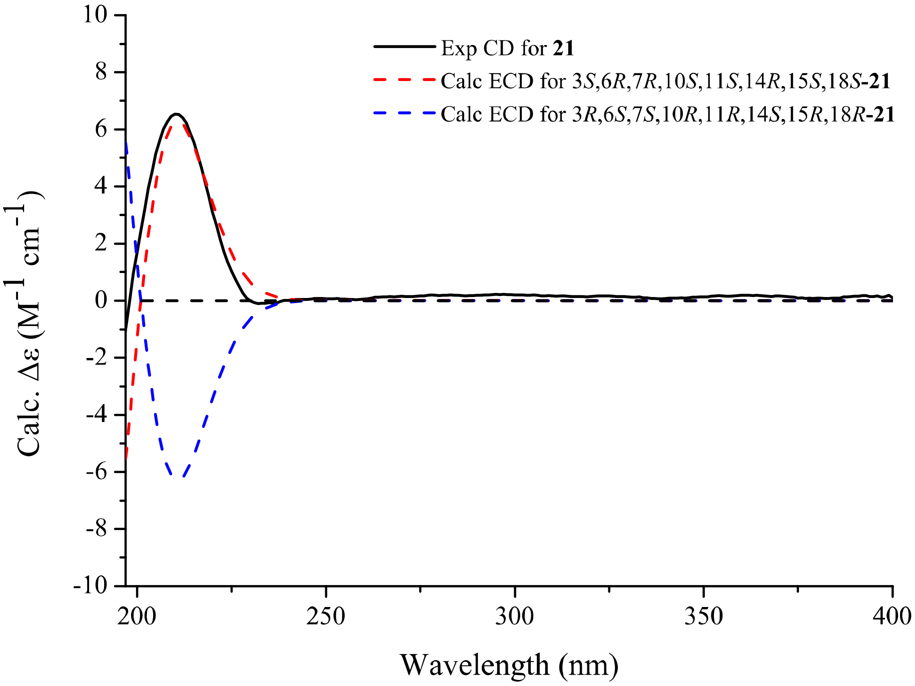

**Figure S36.** Computed ECD and experimental CD spectra of **21**.

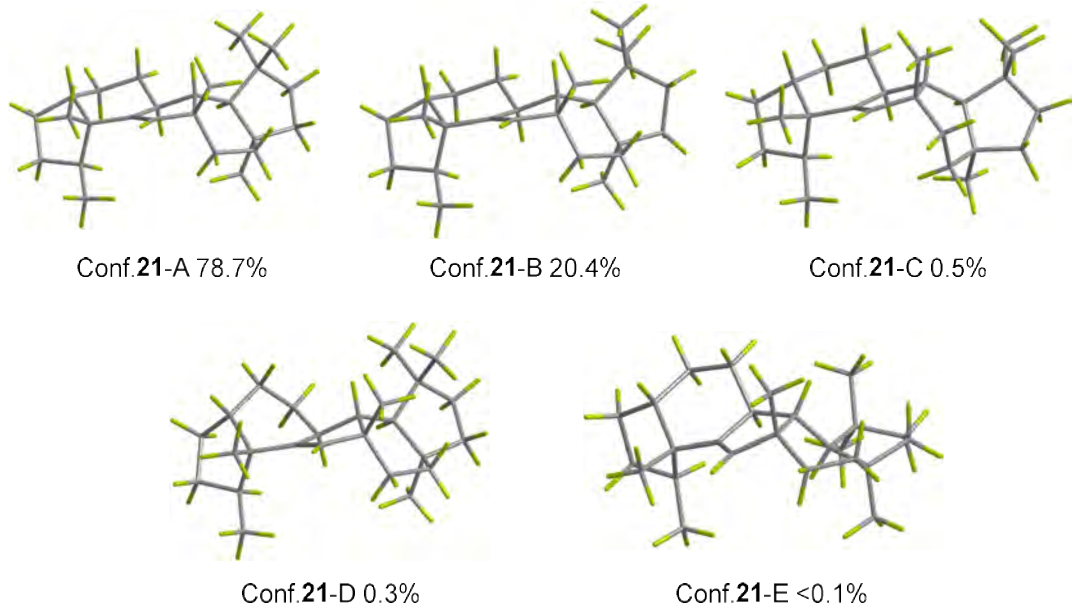

**Figure S37.** DFT-optimized structures for low-energy conformers of 3*S*,6*R*,7*R*,10*S*,11*S*,14*R*,15*S*,18*S*-**21** at the B3LYP/6-31G(d) level in acetonitrile (PCM).

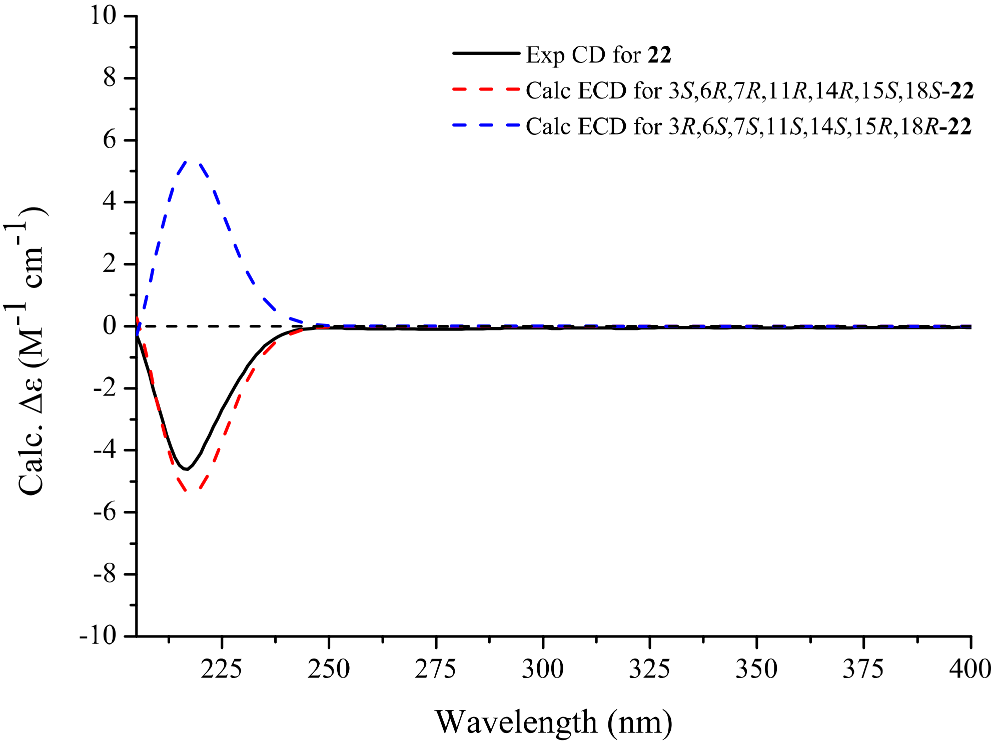

**Figure S38.** Computed ECD and experimental CD spectra of **22**.

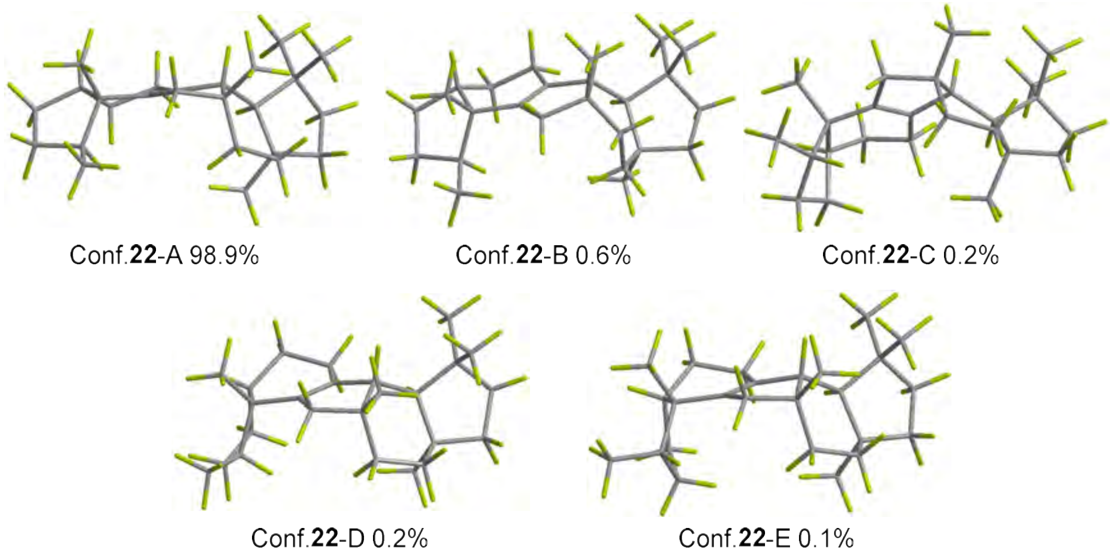

**Figure S39.** DFT-optimized structures for low-energy conformers of 3*S*,6*R*,7*R*,11*R*,14*R*,15*S*,18*S*-**22** at the B3LYP/6-31G(d) level in acetonitrile (PCM).

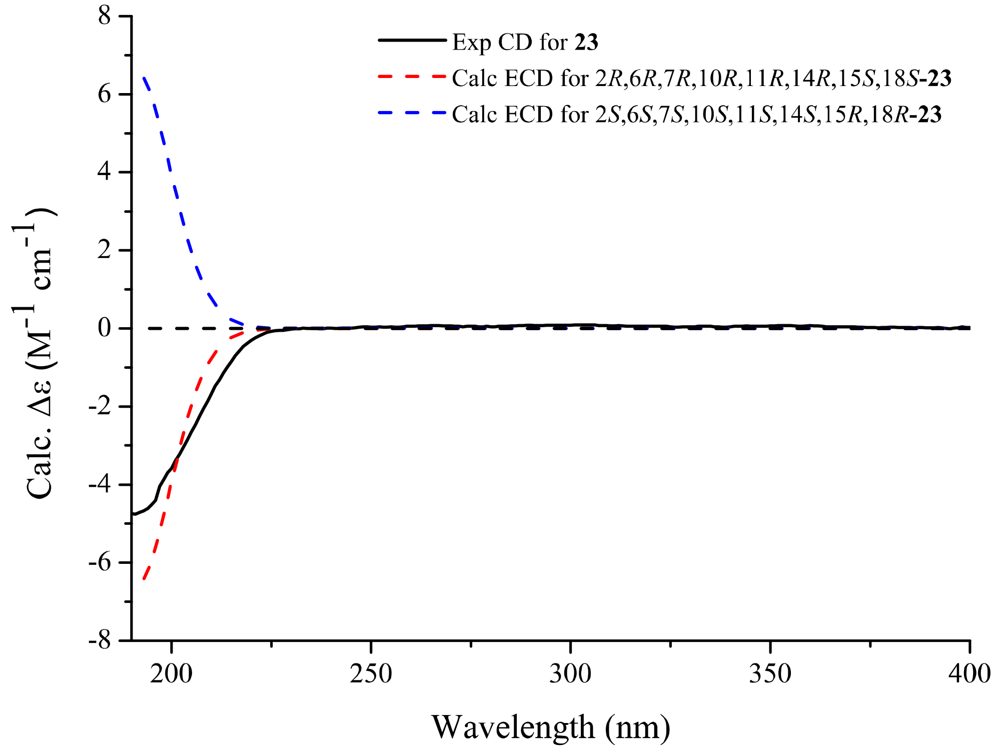

**Figure S40.** Computed ECD and experimental CD spectra of **23**.

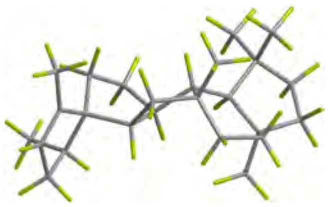

Conf.**23**-A 69%

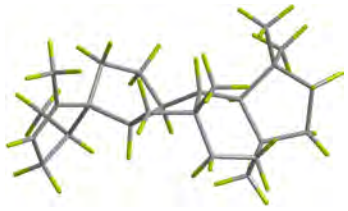

Conf.**23**-B 27.9%

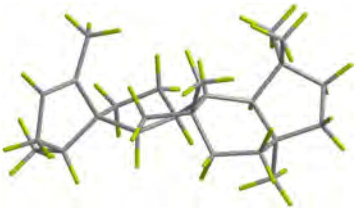

Conf.**23**-C 2.1%

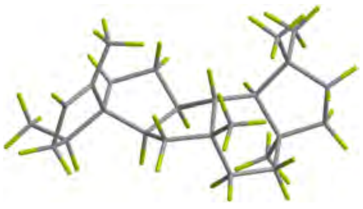

Conf.**23**-D 0.7%

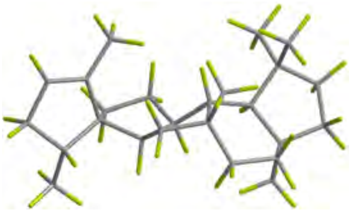

Conf.**23**-E 0.2%

**Figure S41.** DFT-optimized structures for low-energy conformers of 2*R*,6*R*,7*R*,10*R*,11*R*,14*R*,15*S*,18*S*-**23** at the B3LYP/6-31G(d) level in acetonitrile (PCM).

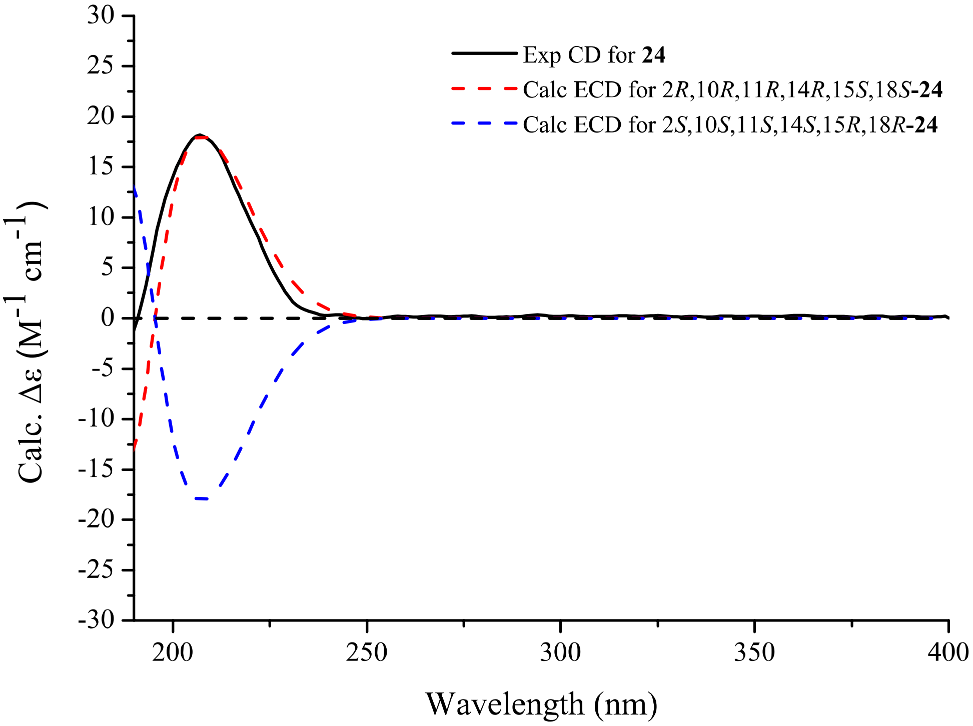

**Figure S42.** Computed ECD and experimental CD spectra of **24**.

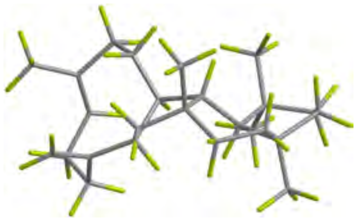

Conf.**24**-A 99.9%

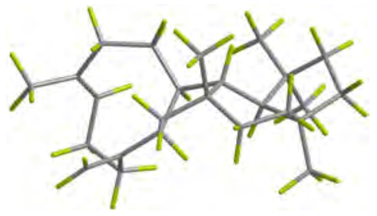

Conf.**24**-B <0.1%

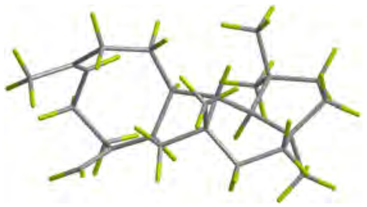

Conf.**24**-C <0.1%

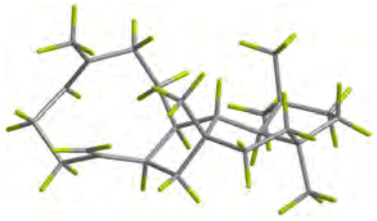

Conf.**24**-D<0.1%

**Figure S43.** DFT-optimized structures for low-energy conformers of 2*R*,10*R*,11*R*,14*R*,15*S*,18*S*-**24** at the B3LYP/6-31G(d) level in acetonitrile (PCM).

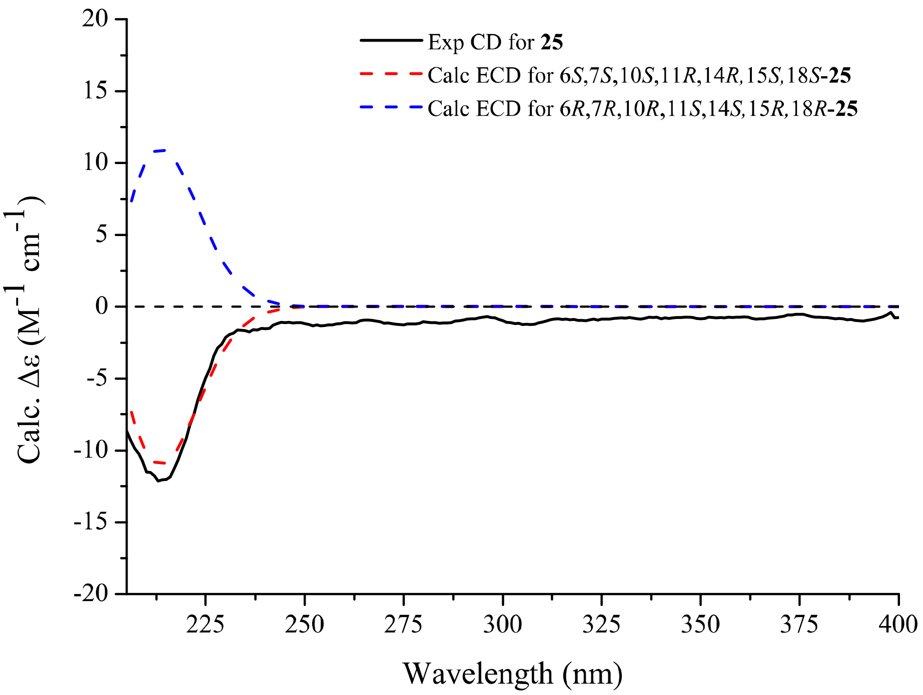

**Figure S44.** Computed ECD and experimental CD spectra of **25**.

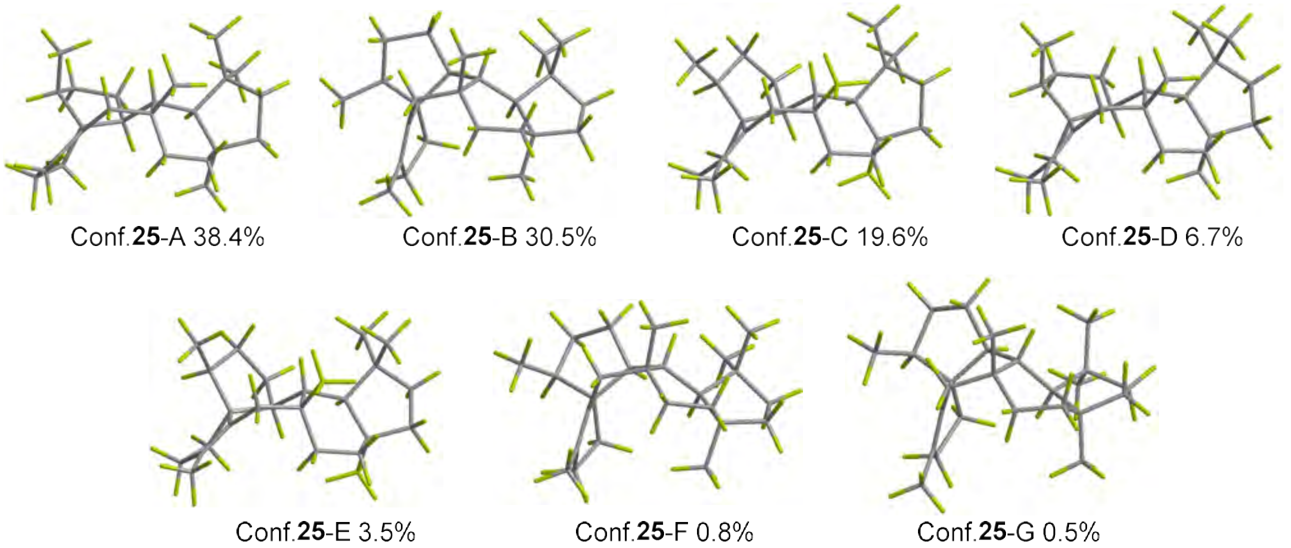

**Figure S45.** DFT-optimized structures for low-energy conformers of 6*S*,7S,10*S*,11*R*,14*R*,15*S*,18*S*-**25** at the B3LYP/6-31G(d) level in acetonitrile (PCM).

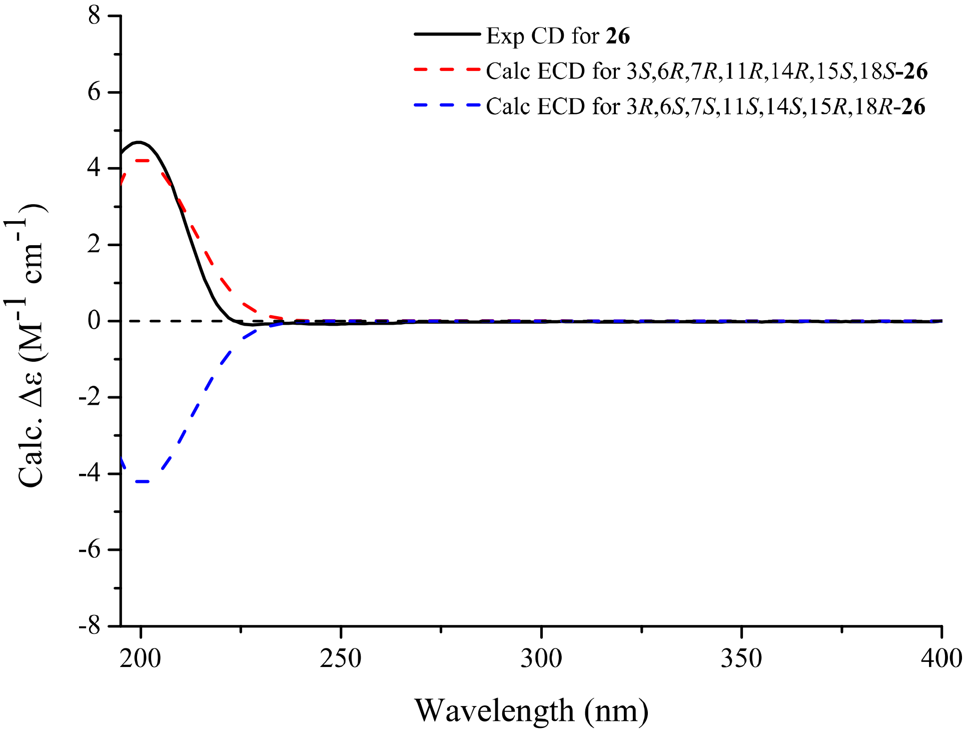

**Figure S46.** Computed ECD and experimental CD spectra of **26**.

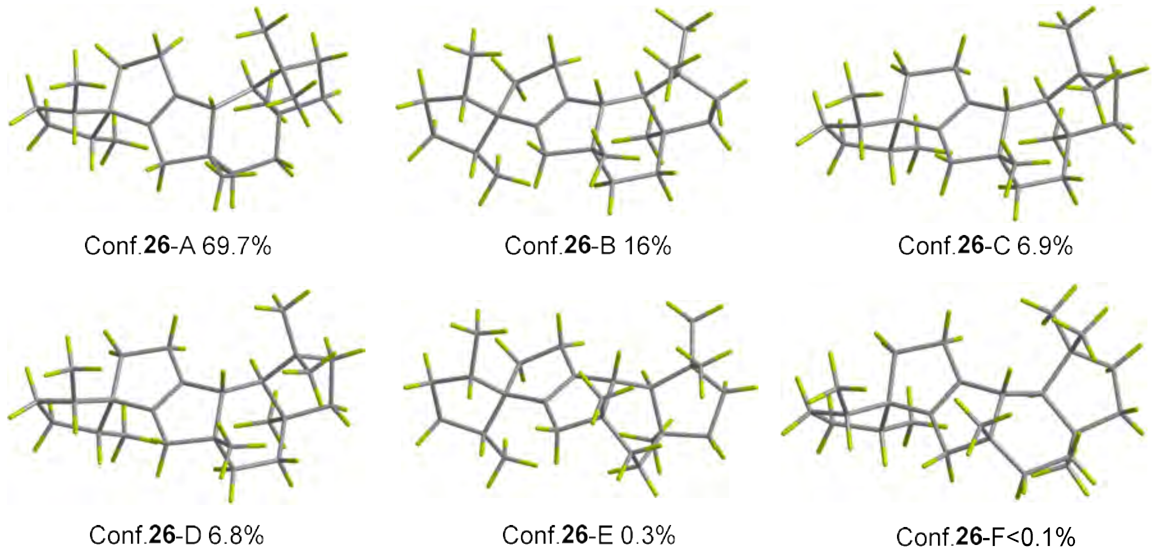

**Figure S47.** DFT-optimized structures for low-energy conformers of 3*S*,6*R*,7*R*,11*R*,14*R*,15*S*,18*S*-**26** at the B3LYP/6-31G(d) level in acetonitrile (PCM).

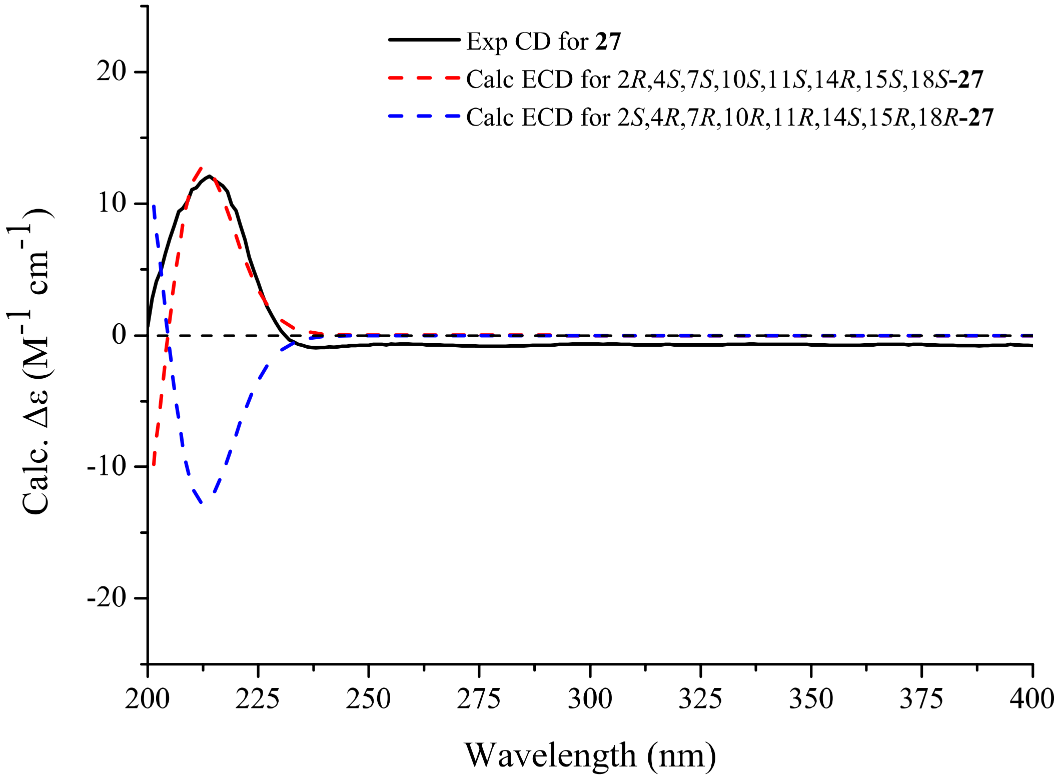

**Figure S48.** Computed ECD and experimental CD spectra of **27**.

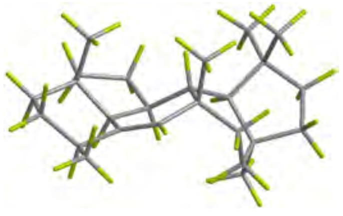

Conf.**27**-A 97.2%

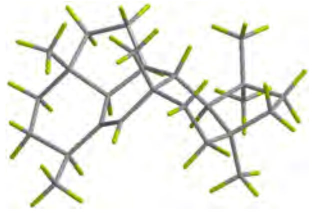

Conf.**27**-B 0.9%

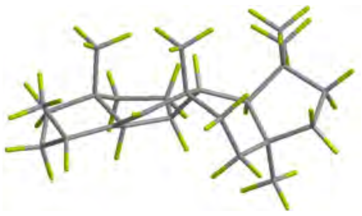

Conf.**27**-C 0.9%

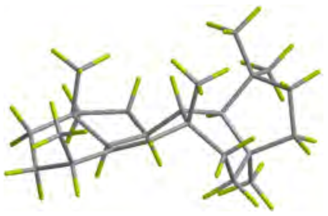

Conf.**27**-D 0.6%

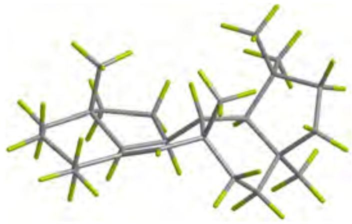

Conf.**27**-E 0.3%

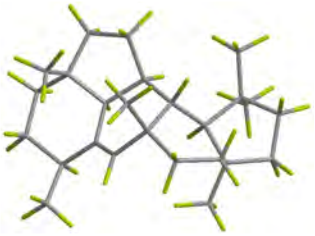

Conf.**27**-F 0.1%

**Figure S49.** DFT-optimized structures for low-energy conformers of 2*R*,4*S*,7*S*,10*S*,11*S*,14*R*,15*S*,18*S*-**27** at the B3LYP/6-31G(d) level in acetonitrile (PCM).

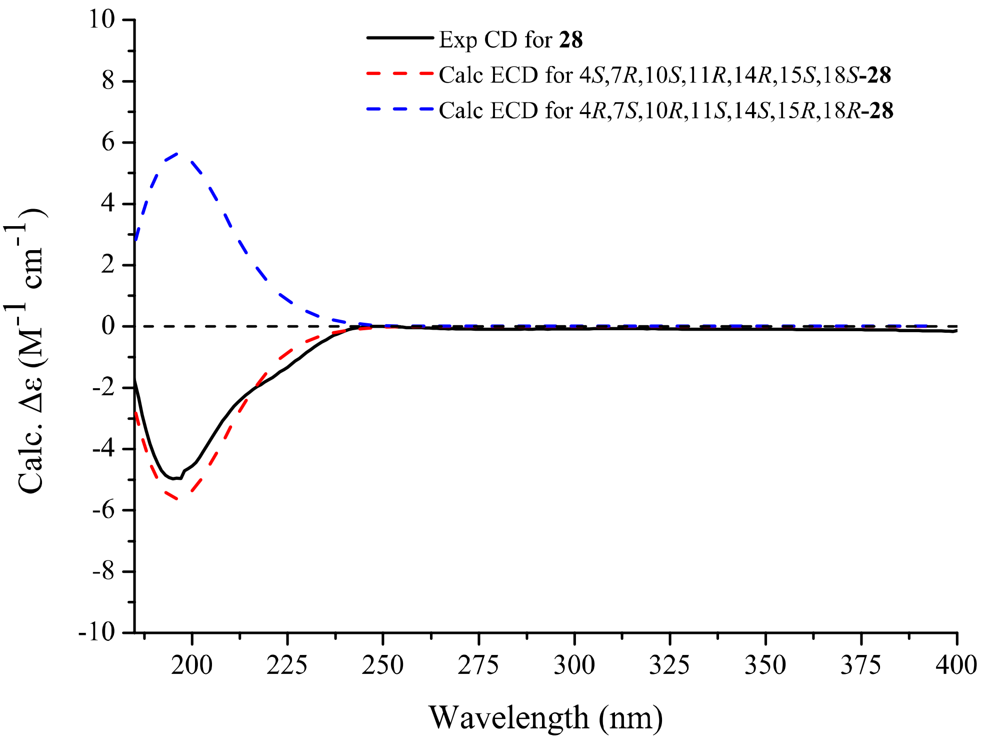

**Figure S50.** Computed ECD and experimental CD spectra of **28**.

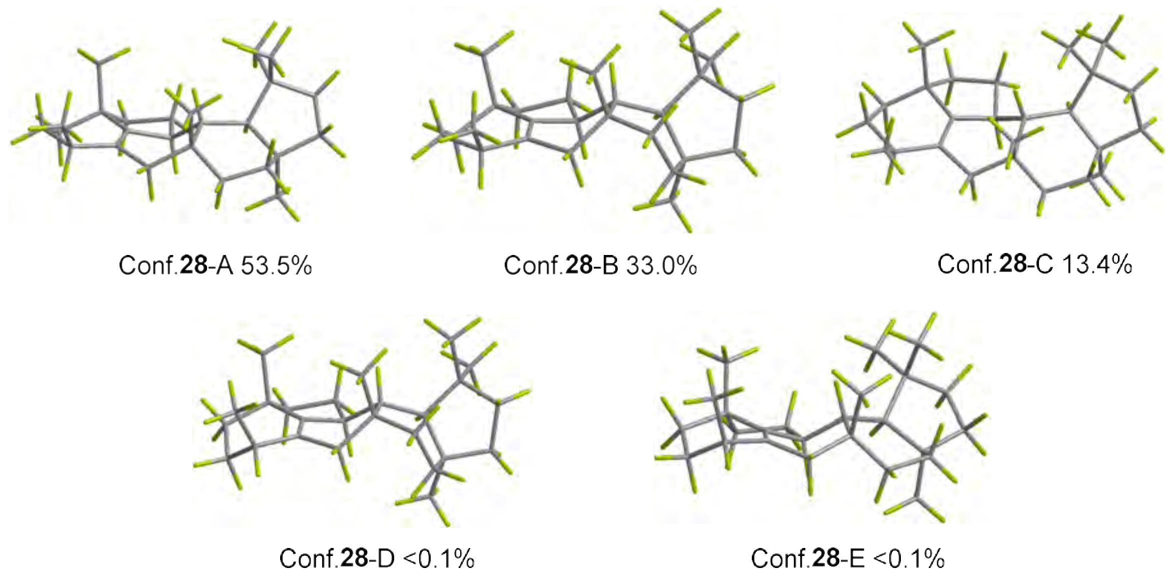

**Figure S51.** DFT-optimized structures for low-energy conformers of 4*S*,7*R*,10*S*,11*R*,14*R*,15*S*,18*S*-**28** at the B3LYP/6-31G(d) level in acetonitrile (PCM).

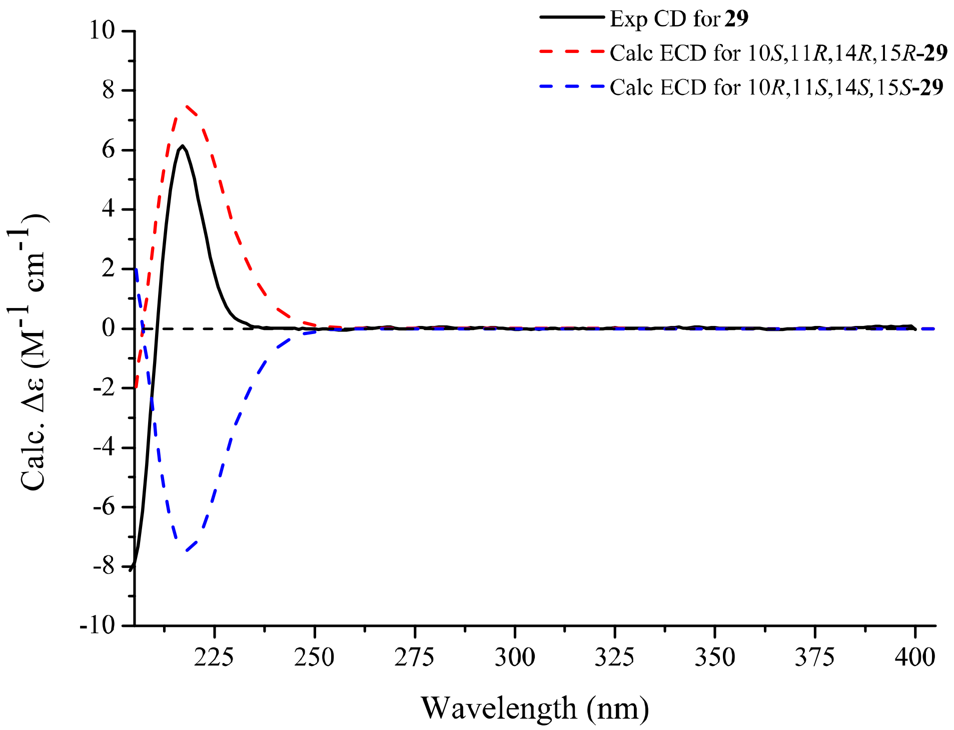

**Figure S52.** Computed ECD and experimental CD spectra of **29**.

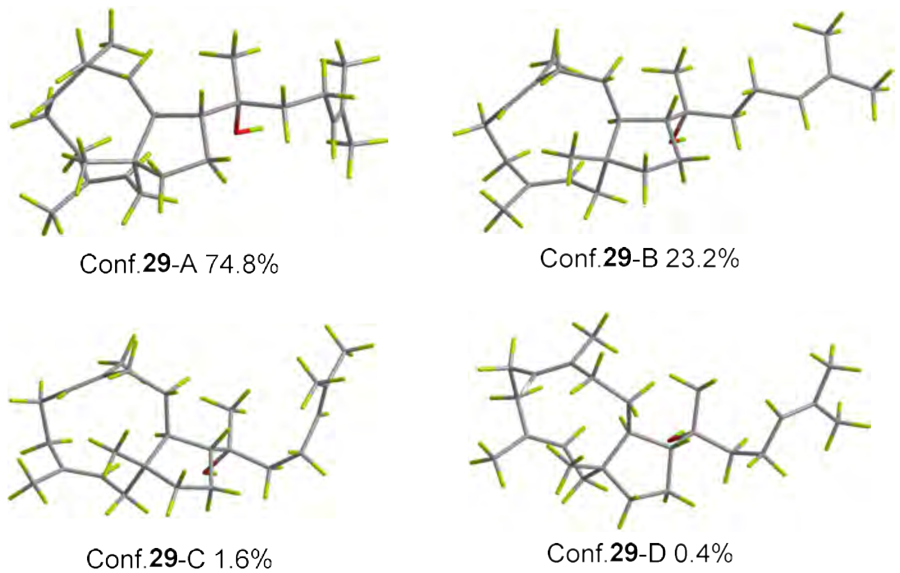

**Figure S53.** DFT-optimized structures for low-energy conformers of 10*S*,11*R*,14*R*,15*R*-**29** at the B3LYP/6-31G(d) level in acetonitrile (PCM).

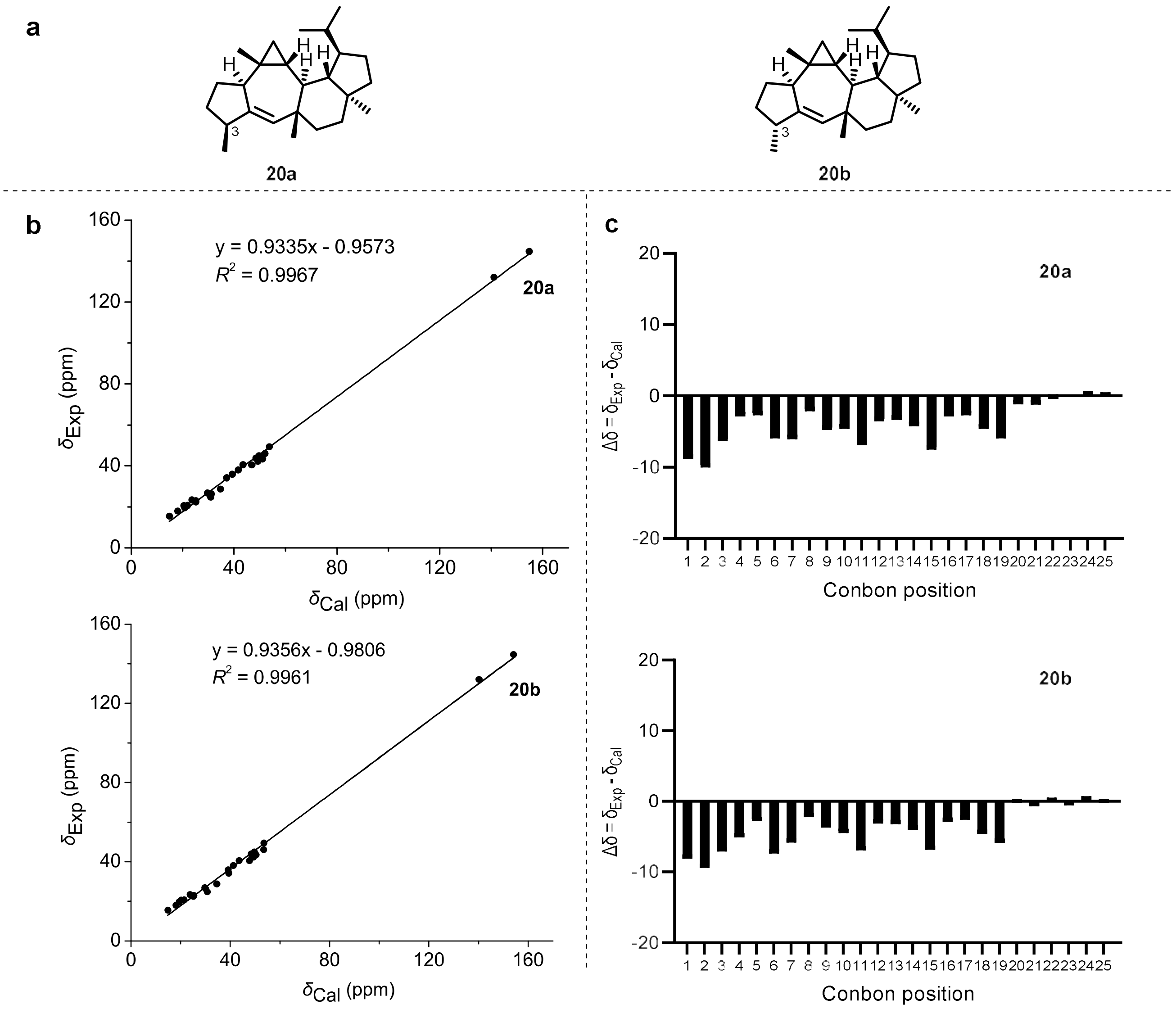

**Figure S54.** $^{13}$C NMR calculation results of two possible isomers of **20a** and **20b**. (a) Two possible isomers of **20a** and **20b**. (b) Linear correlation plots of calculated versus experimental $^{13}$C NMR chemical shift. (c) Relative errors between the experimental $\delta_C$ and calculated $\delta_C$ of two potential structures.

| A | B | C | D | E | F | G | H |
|---|---|---|---|---|---|---|---|
| Functional | | Solvent? | | Basis Set | | Type of Data | |
| B3LYP | | PCM | | 6-311+G(d,p) | | Shielding Tensors | |
| | | DP4+ | 99.52% | 0.48% | - | - | - |
| Nuclei | sp2? | Experimental | Isomer 1 | Isomer 2 | Isomer 3 | Isomer 4 | Isomer 5 |
| C | x | 132.1 | 41.566 | 42.266 | | | |
| C | x | 144.7 | 27.766 | 28.366 | | | |
| C | | 40.6 | 135.566 | 134.766 | | | |
| C | | 34.3 | 145.366 | 143.166 | | | |
| C | | 26.9 | 152.866 | 152.766 | | | |
| C | | 46.1 | 130.466 | 128.966 | | | |
| C | | 24.9 | 151.566 | 151.766 | | | |
| C | | 23 | 157.266 | 157.166 | | | |
| C | | 26.4 | 151.366 | 152.366 | | | |
| C | | 43.8 | 134.066 | 134.166 | | | |
| C | | 42.4 | 133.266 | 133.166 | | | |
| C | | 38.1 | 140.866 | 141.266 | | | |
| C | | 35.9 | 143.166 | 143.366 | | | |
| C | | 49.5 | 128.766 | 128.966 | | | |
| C | | 43.5 | 131.466 | 132.066 | | | |
| C | | 40.7 | 138.966 | 138.966 | | | |
| C | | 22.4 | 157.366 | 157.366 | | | |
| C | | 45.1 | 132.766 | 132.866 | | | |
| C | | 28.7 | 147.766 | 147.866 | | | |
| C | | 19.7 | 161.666 | 163.166 | | | |
| C | | 20.7 | 160.566 | 161.066 | | | |
| C | | 20.7 | 161.966 | 162.266 | | | |
| C | | 18.1 | 164.466 | 164.466 | | | |
| C | | 15.5 | 167.566 | 167.666 | | | |
| C | | 23.5 | 158.866 | 158.766 | | | |

| A | B | C | D | E | F | G | H |
|---|---|---|---|---|---|---|---|
| Functional | | Solvent? | | Basis Set | | Type of Data | |
| B3LYP | | PCM | | 6-311+G(d,p) | | Shielding Tensors | |
| | | Isomer 1 | Isomer 2 | Isomer 3 | Isomer 4 | Isomer 5 | Isomer 6 |
| sDP4+ (H data) | | - | - | - | - | - | - |
| sDP4+ (C data) | | 96.53% | 3.47% | - | - | - | - |
| sDP4+ (all data) | | 96.53% | 3.47% | - | - | - | - |
| uDP4+ (H data) | | - | - | - | - | - | - |
| uDP4+ (C data) | | 88.27% | 11.73% | - | - | - | - |
| uDP4+ (all data) | | 88.27% | 11.73% | - | - | - | - |
| DP4+ (H data) | | - | - | - | - | - | - |
| DP4+ (C data) | | 99.52% | 0.48% | - | - | - | - |
| DP4+ (all data) | | 99.52% | 0.48% | - | - | - | - |

**Figure S55. DP4⁺** analysis results of two possible isomers of **20a** (Isomer 1) and **20b** (Isomer 2).

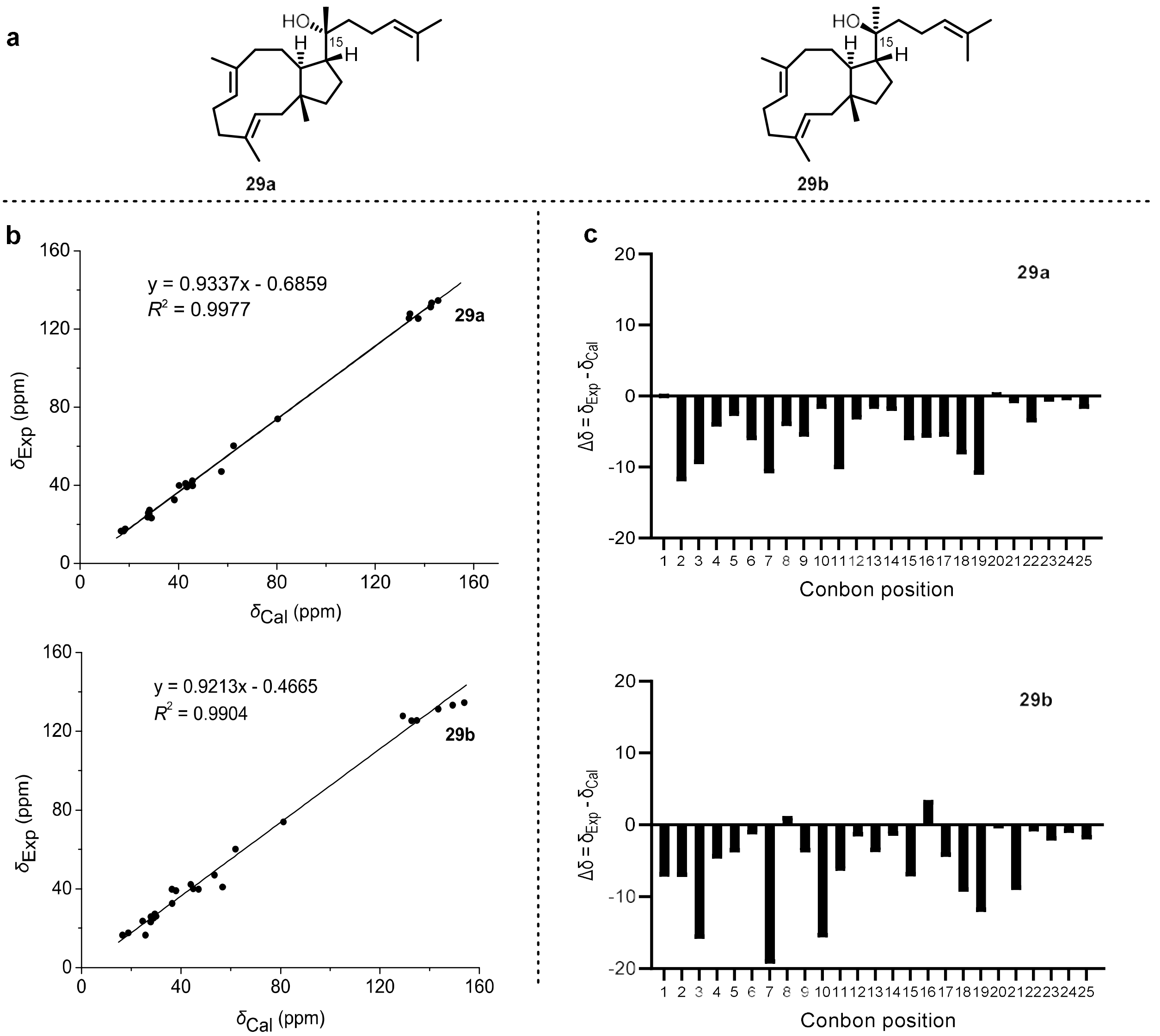

**Figure S56.** $^{13}$C NMR calculation results of two possible isomers of **29a** and **29b**. (a) Two possible isomers of **29a** and **29b**. (b) Linear correlation plots of calculated versus experimental $^{13}$C NMR chemical shift. (c) Relative errors between the experimental $\delta_C$ and calculated $\delta_C$ of two potential structures.

| A | B | C | D | E | F | G | H |
|---|---|---|---|---|---|---|---|
| Functional | | Solvent? | | Basis Set | | Type of Data | |
| B3LYP | | PCM | | 6-311+G(d,p) | | Shielding Tensors | |
| | | DP4+ | 100.00% | 0.00% | - | - | - |
| Nuclei | sp2? | xperimenta | Isomer 1 | Isomer 2 | Isomer 3 | Isomer 4 | Isomer 5 |
| C | | 39.9 | 142.266 | 135.366 | | | |
| C | x | 125.4 | 45.066 | 49.766 | | | |
| C | x | 133.3 | 39.566 | 33.266 | | | |
| C | | 40.2 | 137.966 | 137.566 | | | |
| C | | 25.2 | 154.466 | 153.466 | | | |
| C | x | 127.8 | 48.466 | 53.366 | | | |
| C | x | 134.6 | 36.966 | 28.566 | | | |
| C | | 39.2 | 139.066 | 144.466 | | | |
| C | | 32.6 | 144.166 | 146.066 | | | |
| C | | 41 | 139.666 | 125.766 | | | |
| C | | 47.1 | 125.066 | 128.966 | | | |
| C | | 42.3 | 136.866 | 138.566 | | | |
| C | | 26.1 | 154.566 | 152.566 | | | |
| C | | 60.3 | 120.066 | 120.666 | | | |
| C | | 74 | 102.266 | 101.366 | | | |
| C | | 39.8 | 136.766 | 146.166 | | | |
| C | | 23.3 | 153.466 | 154.666 | | | |
| C | x | 125.5 | 48.766 | 47.666 | | | |
| C | x | 131.3 | 40.066 | 39.066 | | | |
| C | | 16.6 | 165.766 | 165.966 | | | |
| C | | 16.6 | 164.866 | 156.766 | | | |
| C | | 23.7 | 155.066 | 157.866 | | | |
| C | | 27.3 | 154.366 | 152.966 | | | |
| C | | 17.7 | 164.166 | 163.666 | | | |
| C | | 25.9 | 154.766 | 154.566 | | | |

| A | B | C | D | E | F | G | H |
|---|---|---|---|---|---|---|---|
| Functional | | Solvent? | | Basis Set | | Type of Data | |
| B3LYP | | PCM | | 6-311+G(d,p) | | Shielding Tensors | |
| | | Isomer 1 | Isomer 2 | Isomer 3 | Isomer 4 | Isomer 5 | Isomer 6 |
| sDP4+ (H data) | | - | - | - | - | - | - |
| sDP4+ (C data) | | 100.00% | 0.00% | - | - | - | - |
| sDP4+ (all data) | | 100.00% | 0.00% | - | - | - | - |
| uDP4+ (H data) | | - | - | - | - | - | - |
| uDP4+ (C data) | | 100.00% | 0.00% | - | - | - | - |
| uDP4+ (all data) | | 100.00% | 0.00% | - | - | - | - |
| DP4+ (H data) | | - | - | - | - | - | - |
| DP4+ (C data) | | 100.00% | 0.00% | - | - | - | - |
| DP4+ (all data) | | 100.00% | 0.00% | - | - | - | - |

**Figure S57. DP4$^+$** analysis results of two possible isomers of **29a** (Isomer 1) and **29b** (Isomer 2).

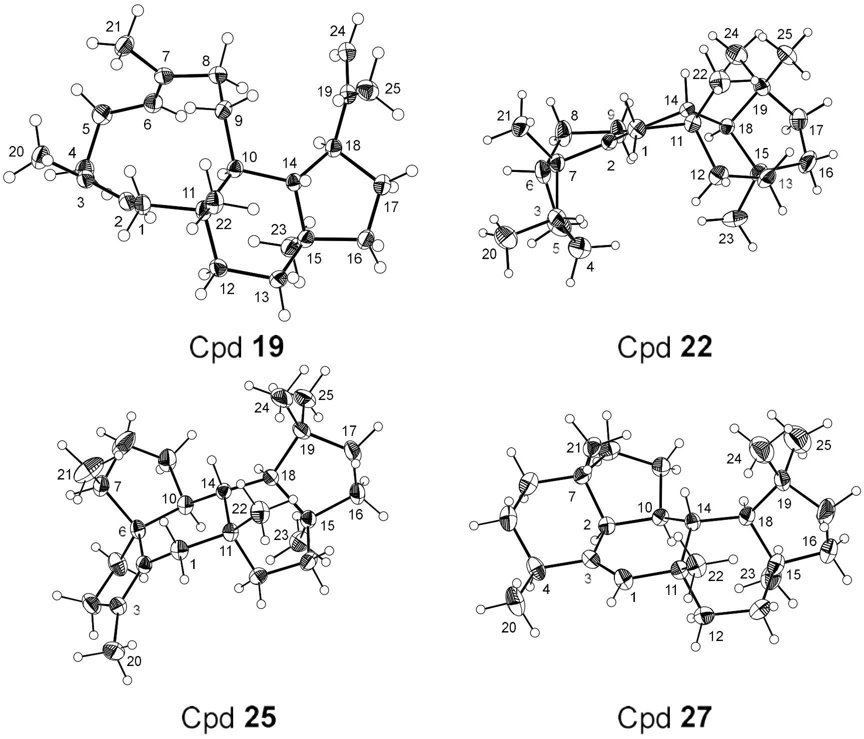

**Figure S58.** ORTEP (Oak Ridge Thermal-Ellipsoid Plot) drawing for X-ray structure of compound **19**, **22**, **25**, and **27**.

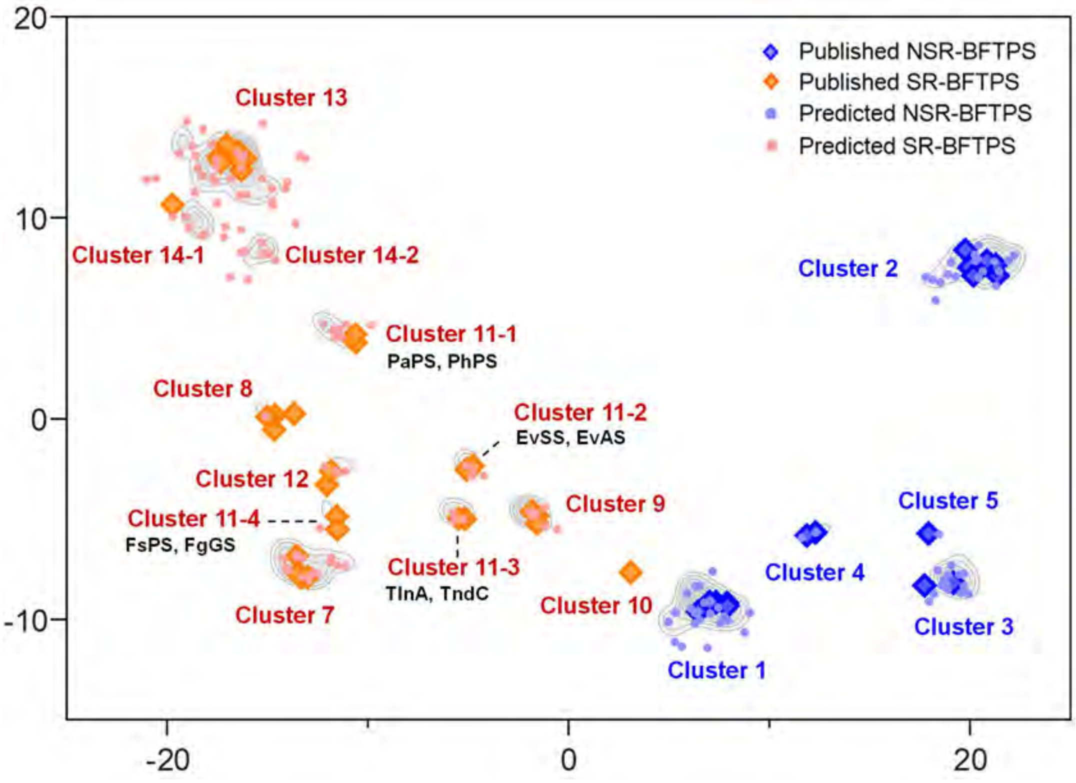

**Figure S59.** Iterative prediction of the updated model incorporating newly characterized BFTPSs. Cluster 11 was redistributed into four smaller product-related clusters (Cluster 11-1–11-4).

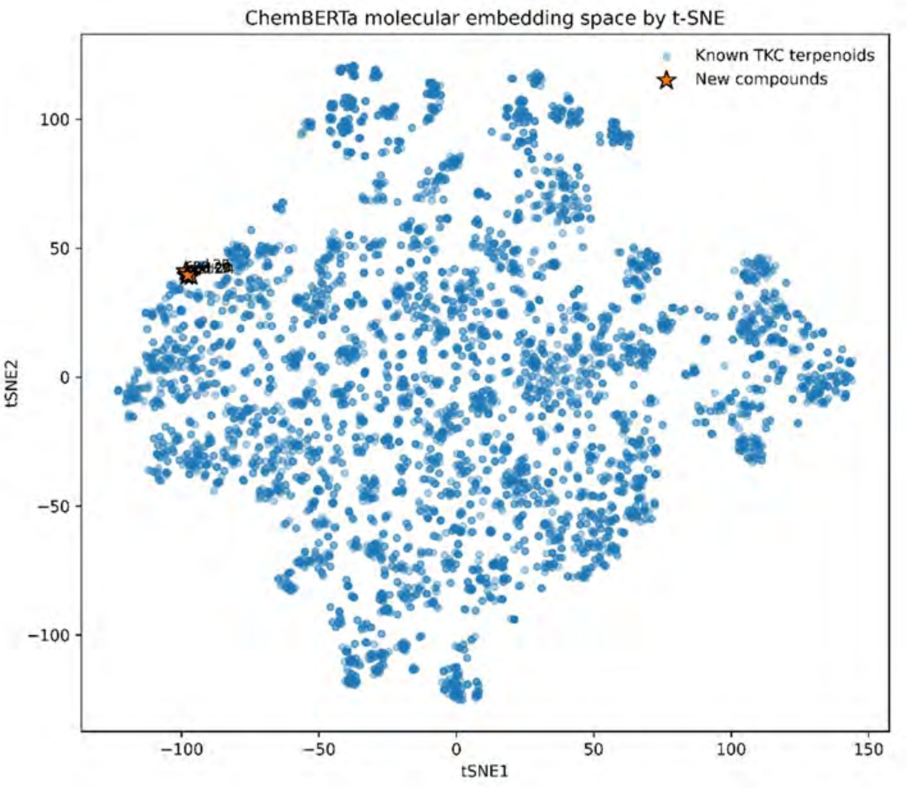

**Figure S60.** t-SNE projection of ChemBERTa molecular embeddings showing the distribution of known sesterterpenoids and newly identified compounds **23–28**.

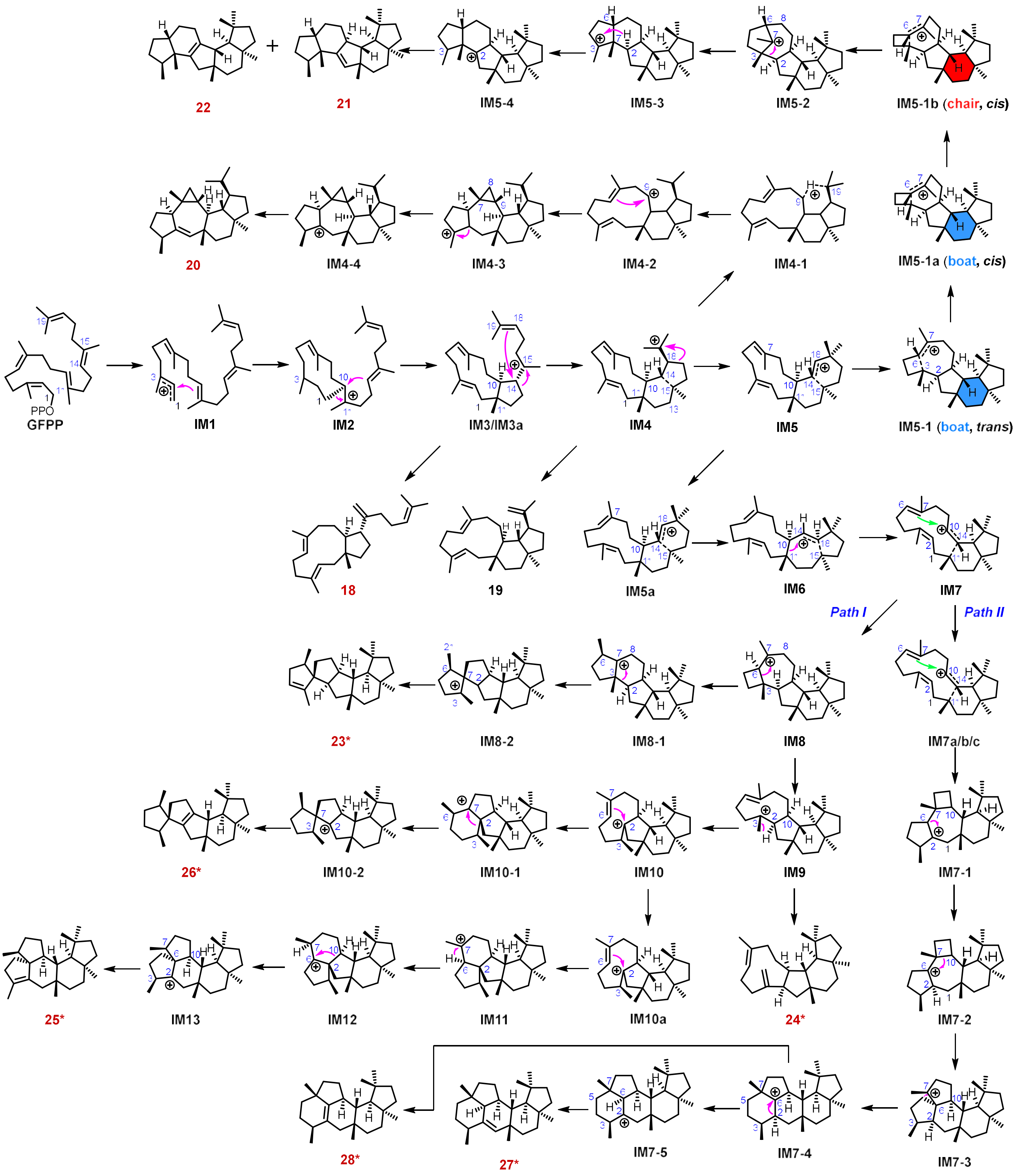

**Figure S61.** Computed cyclization pathways of FsPS and AcPS-derived compounds **18–28**.

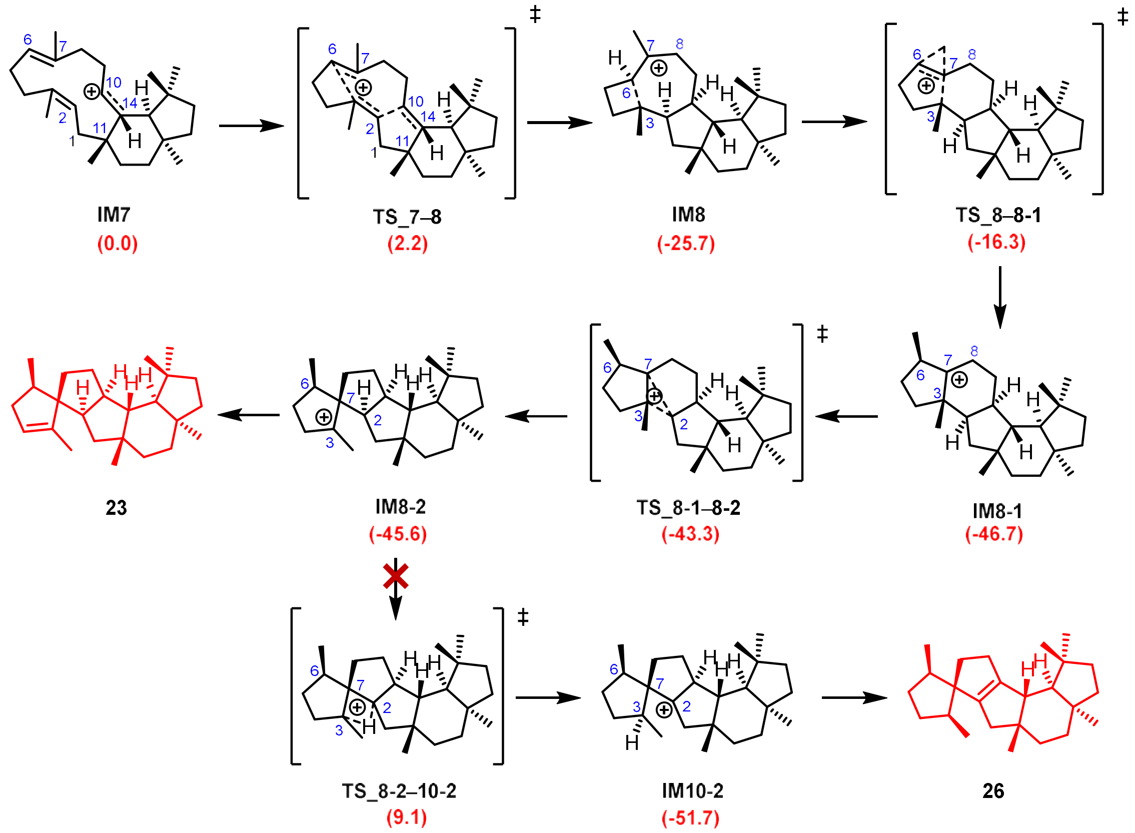

**Figure S62.** DFT-computed reaction pathways for the conversion of **IM7** into compound **23**, with the free energies of key Intermediates (IMs) and transition states (TSs) indicated in parentheses (in red).

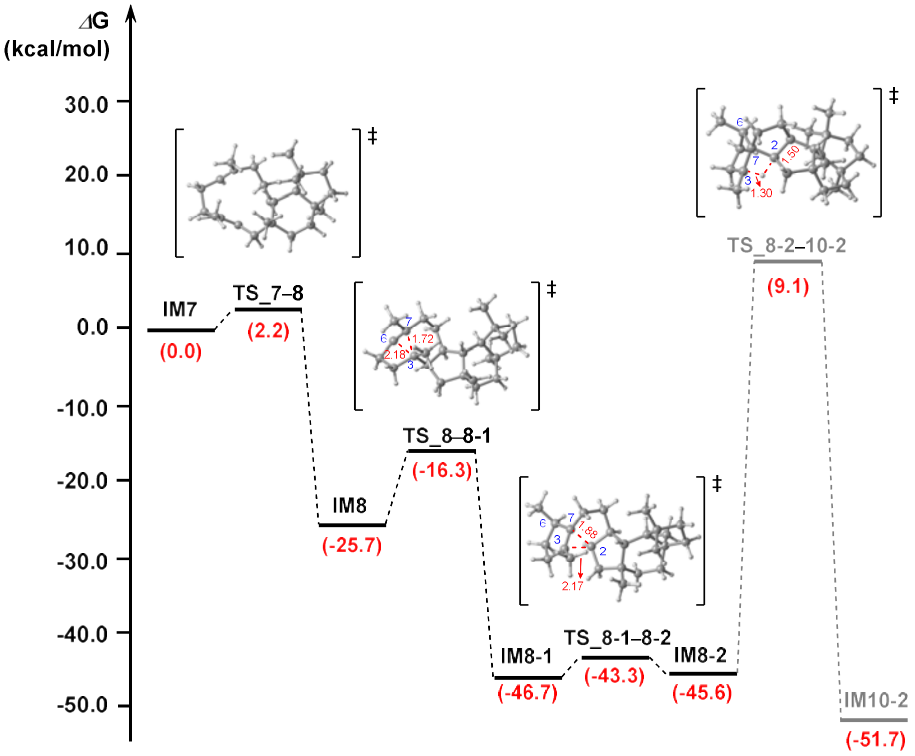

**Figure S63.** The Gibbs free energy profile and the optimized structures for the **IM7** cyclized into **23**. Gibbs free energies (kcal/mol) were calculated at the mPW1PW91/6-311+G(d,p) level based on M06-2X/6-31+G(d,p) geometries. The potential energy profiles relative to **IM7** are shown in red. **IM** means intermediate, **TS** means transition state.

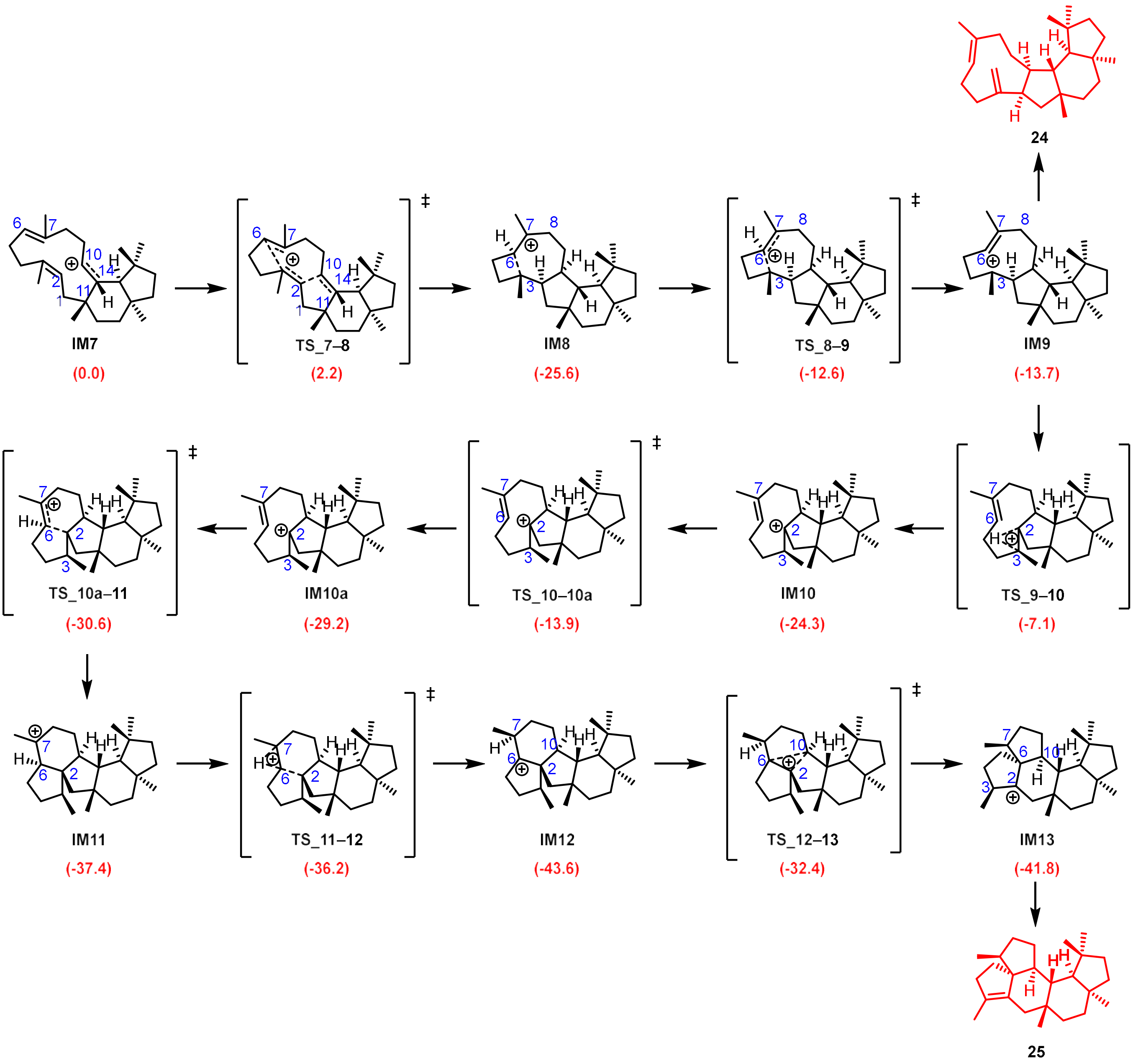

**Figure S64.** DFT-computed reaction pathways for the conversion of **IM7** into compounds **24** and **25**, with the free energies of key Intermediates (IMs) and transition states (TSs) indicated in parentheses (in red).

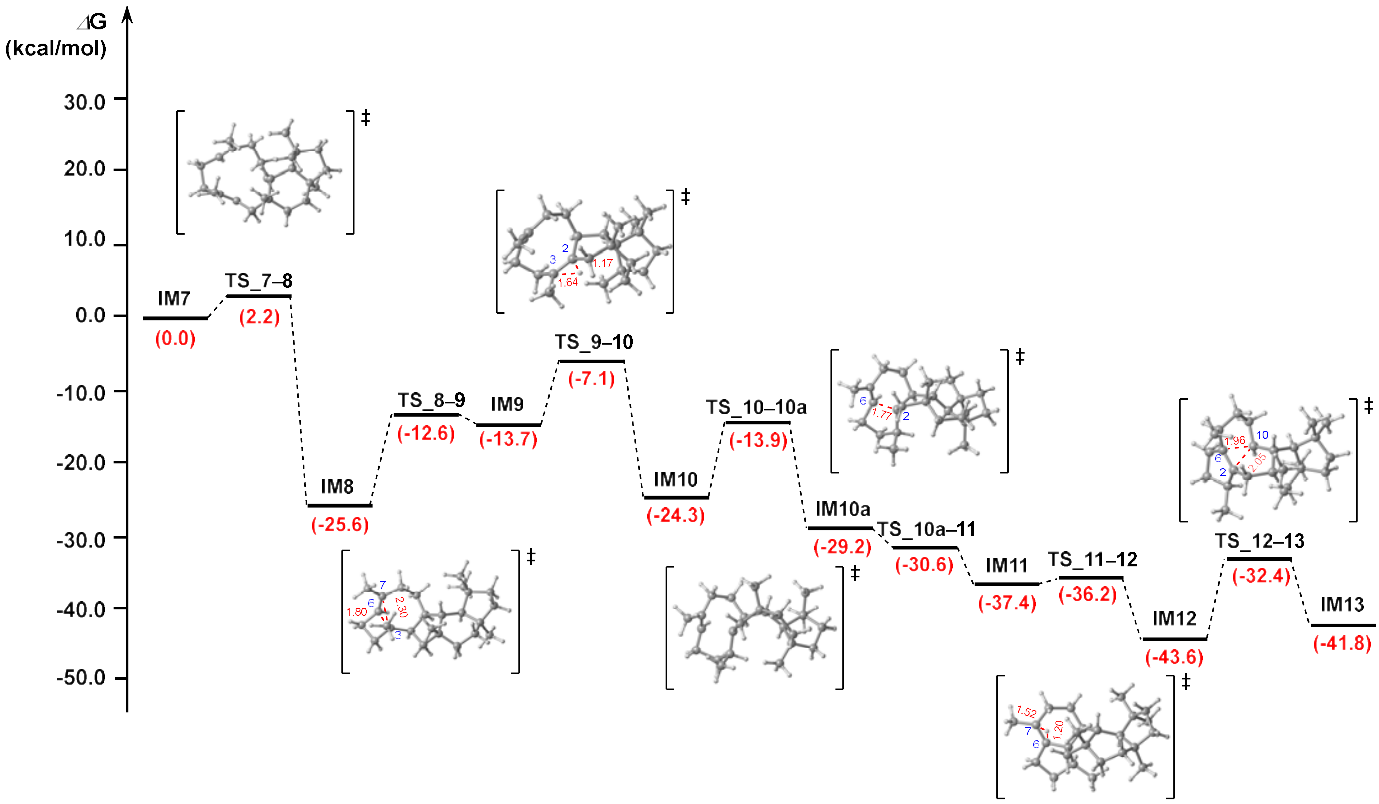

**Figure S65.** The Gibbs free energy profile and the optimized structures for the **IM7** cyclized into **24** and **25**. Gibbs free energies (kcal/mol) were calculated at the mPW1PW91/6-311+G(d,p) level based on M06-2X/6-31+G(d,p) geometries. The potential energy profiles relative to **IM7** are shown in red. **IM** means intermediate, **TS** means transition state.

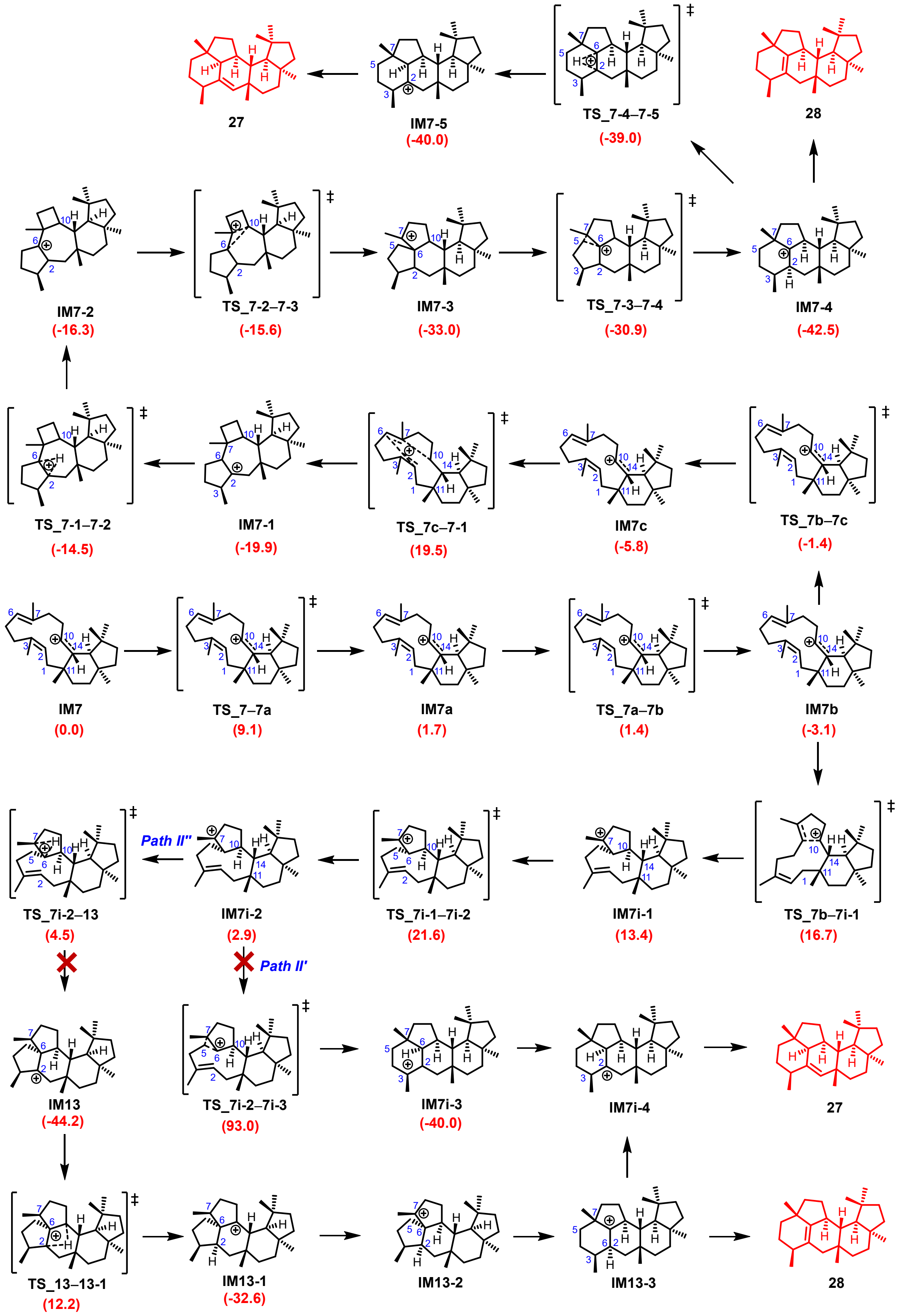

**Figure S66.** DFT-computed reaction pathways for the conversion of **IM7** into compound **27** and **28**, with the free energies of key Intermediates (**IMs**) and transition states (**TSs**) indicated in parentheses (in red).

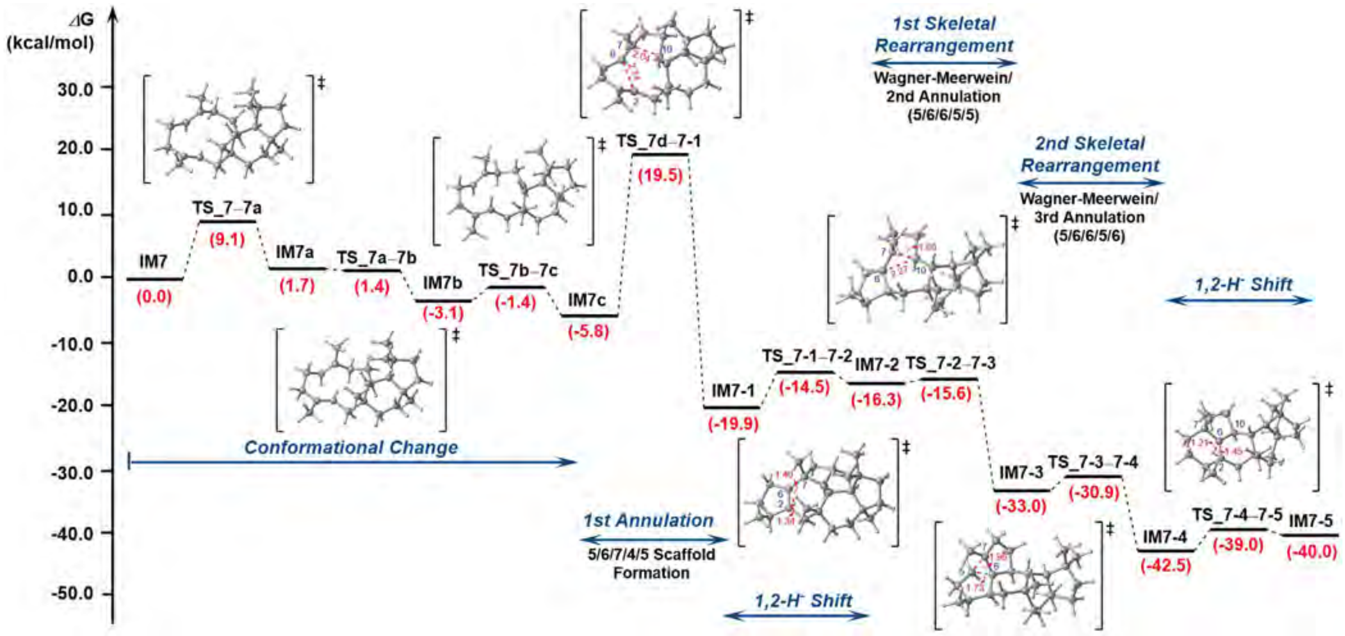

**Figure S67.** The Gibbs free energy profile and the optimized structures for the **IM7** cyclized into **27** and **28**. Gibbs free energies (kcal/mol) were calculated at the mPW1PW91/6-311+G(d,p) level based on M06-2X/6-31+G(d,p) geometries. The potential energy profiles relative to **IM7** are shown in red. **IM** means intermediate, **TS** means transition state.

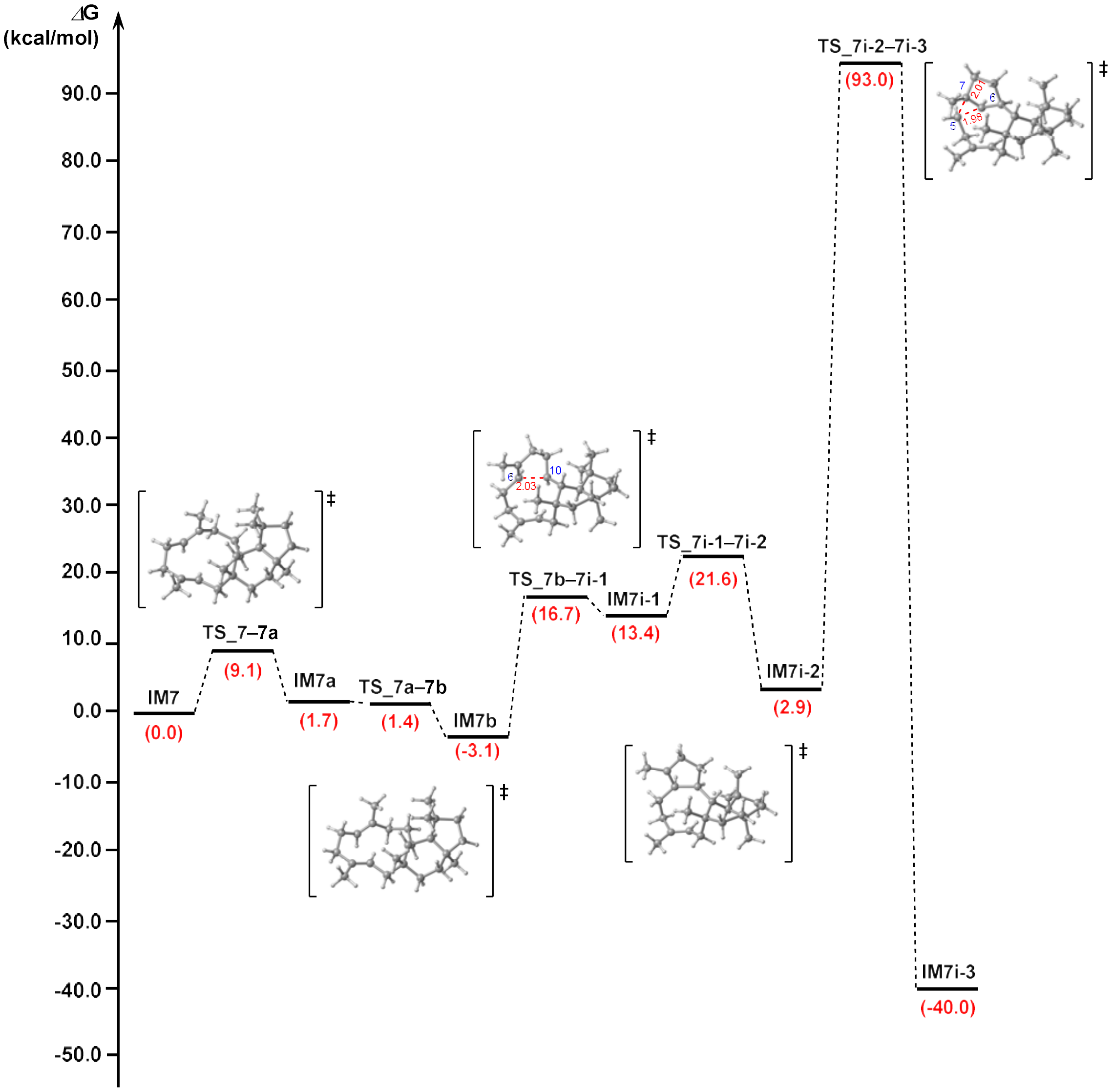

**Figure S68.** The Gibbs free energy profile and the optimized structures of energetically implausible ***Path II'*** for the **IM7** cyclized into **27** and **28**. Gibbs free energies (kcal/mol) were calculated at the mPW1PW91/6-311+G(d,p) level based on M06-2X/6-31+G(d,p) geometries. The potential energy profile relative to **IM7** are shown in red. **IM** means intermediate, **TS** means transition state.

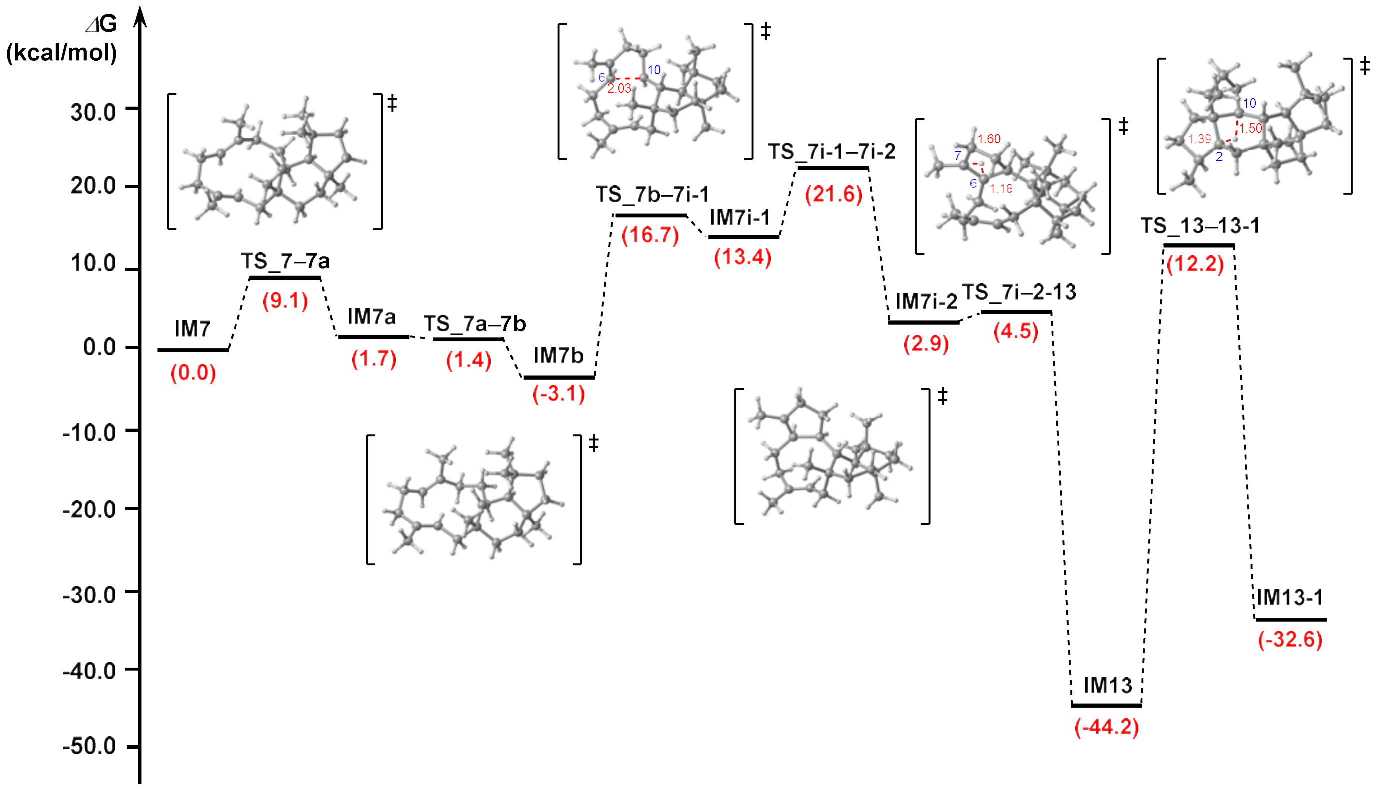

**Figure S69.** The Gibbs free energy profile and the optimized structures of energetically implausible ***Path II''*** for the **IM7** cyclized into **27** and **28**. Gibbs free energies (kcal/mol) were calculated at the mPW1PW91/6-311+G(d,p) level based on M06-2X/6-31+G(d,p) geometries. The potential energy profiles relative to **IM7** are shown in red. **IM** means intermediate, **TS** means transition state.

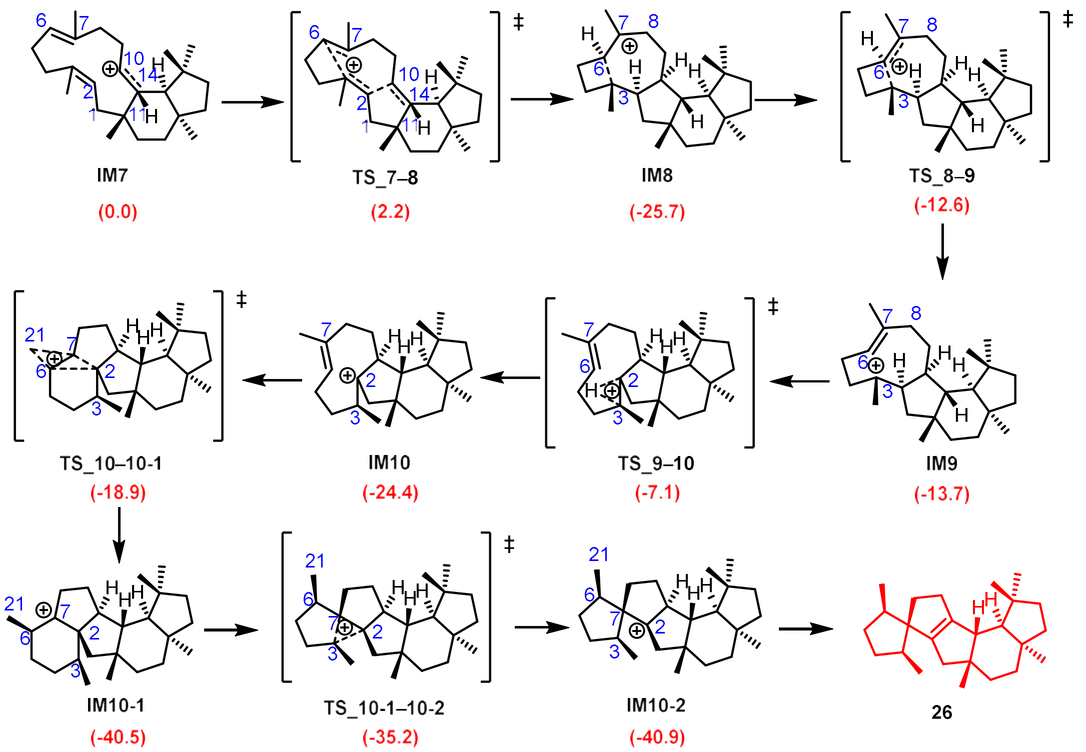

**Figure S70.** DFT-computed reaction pathways for the conversion of **IM7** into compound **26**, with the free energies of key Intermediates (IMs) and transition states (TSs) indicated in parentheses (in red).

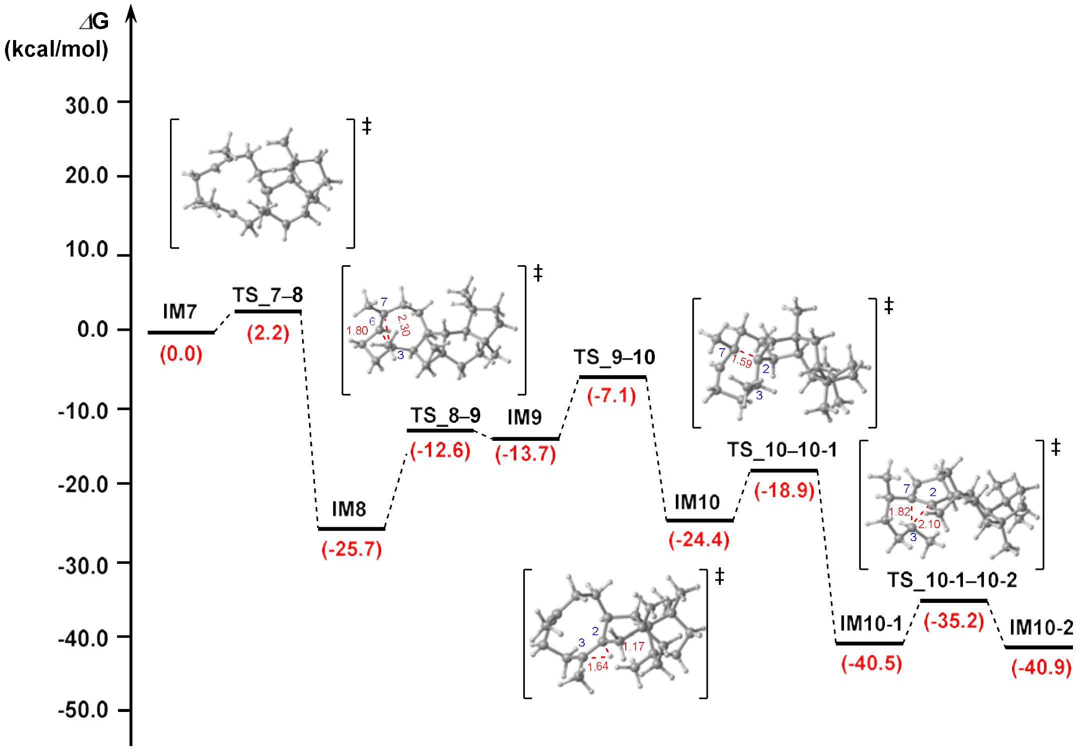

**Figure S71.** The Gibbs free energy profile and the optimized structures for the **IM7** cyclized into **26**. Gibbs free energies (kcal/mol) were calculated at the mPW1PW91/6-311+G(d,p) level based on M06-2X/6-31+G(d,p) geometries. The potential energy profiles relative to **IM7** are shown in red. **IM** means intermediate, **TS** means transition state.

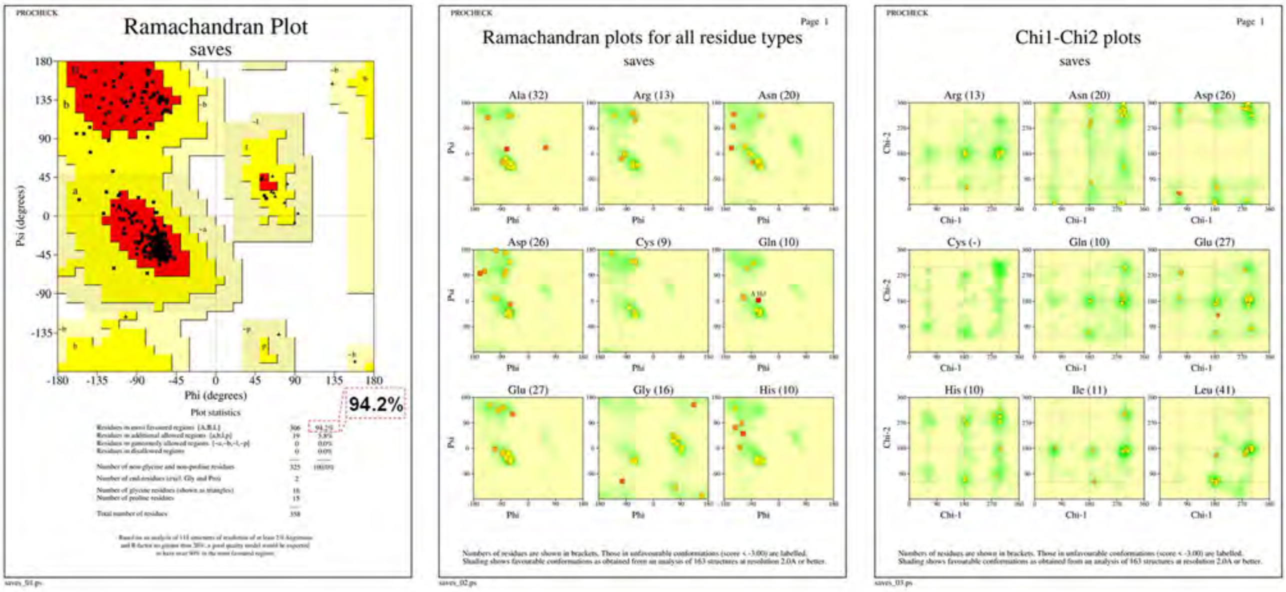

**Figure S72.** Ramachandran plot of the FsPS, as predicted using AlphaFold3.

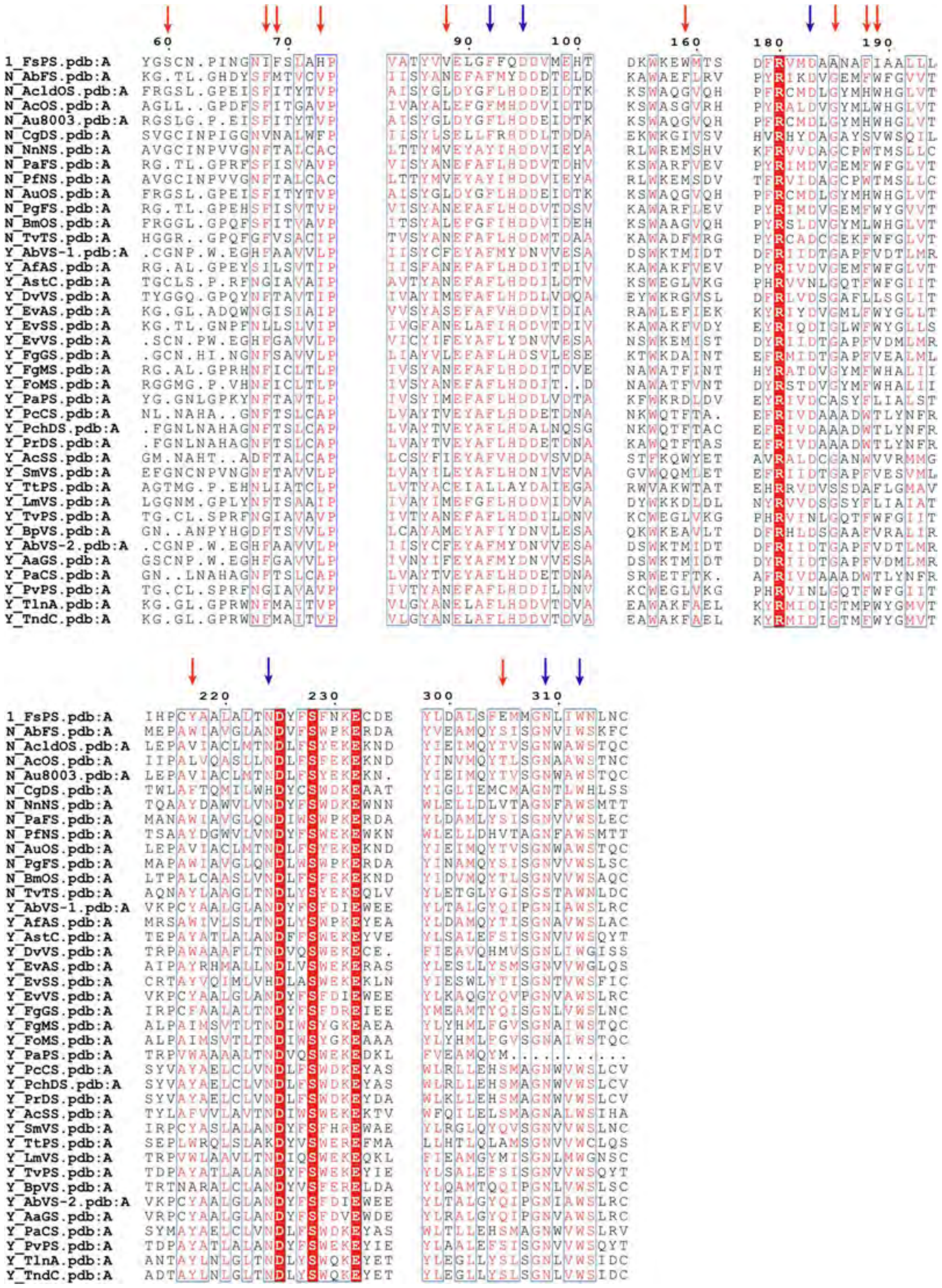

**Figure S73.** Structural alignment of FsPS and 38 previously reported type B BFTPSs. Key pocket residues are indicated by arrows above, with conserved residues shown in blue and non-conserved residues in red. Conservative amino acids were shown with red background.

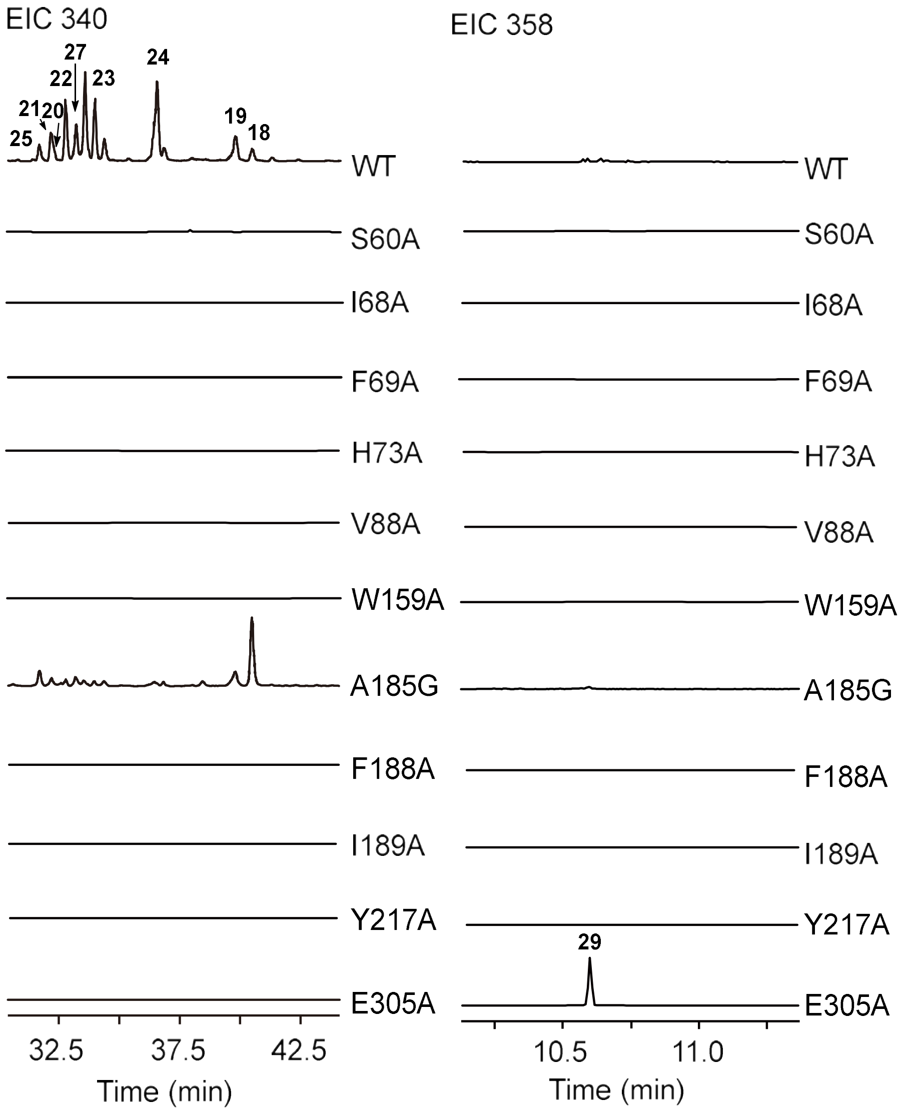

**Figure S74.** GC-MS chromatograms of key-residue mutants in the FsPS active site.

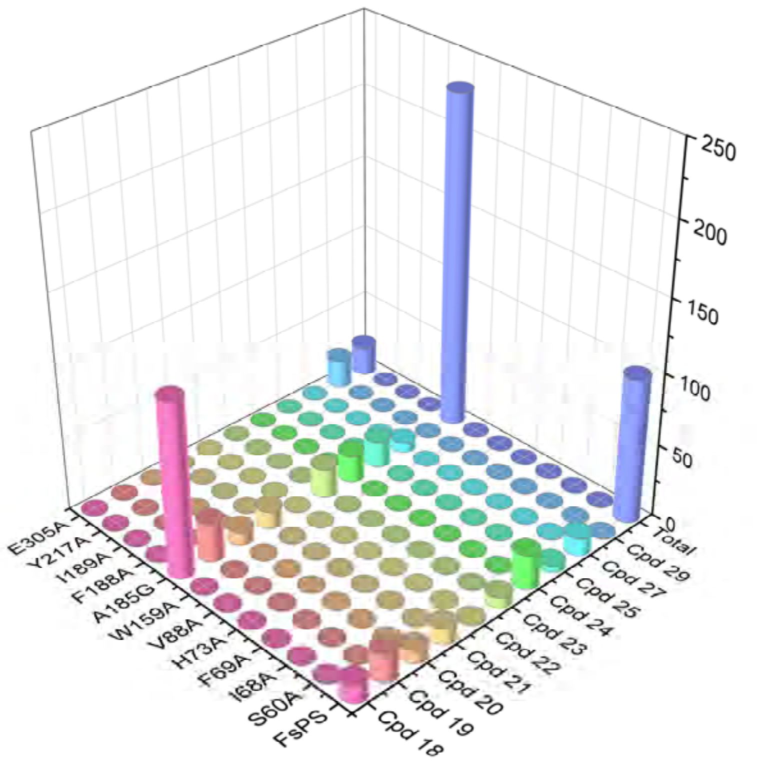

**Figure S75.** Product profile comparisons of FsPS active-site mutants.

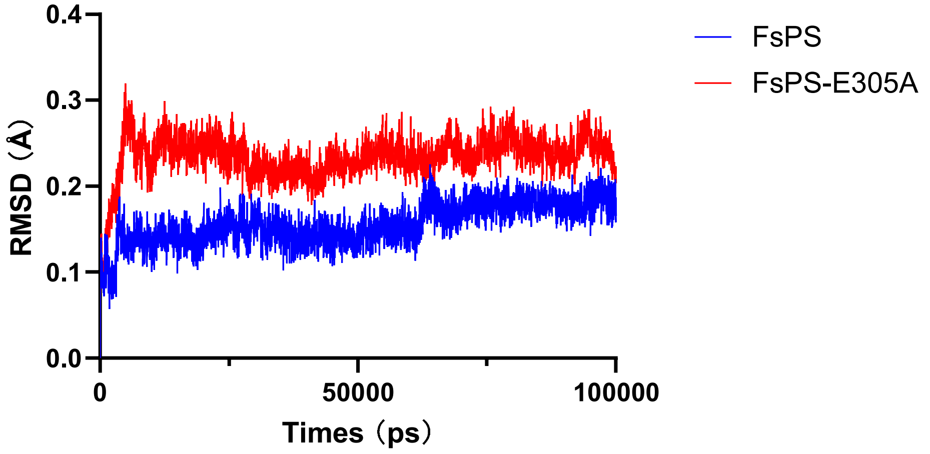

**Figure S76.** RMSD of backbone atoms (Cα, C, N, and O) for FsPS and FsPS-E305A complexed with **IM3**.

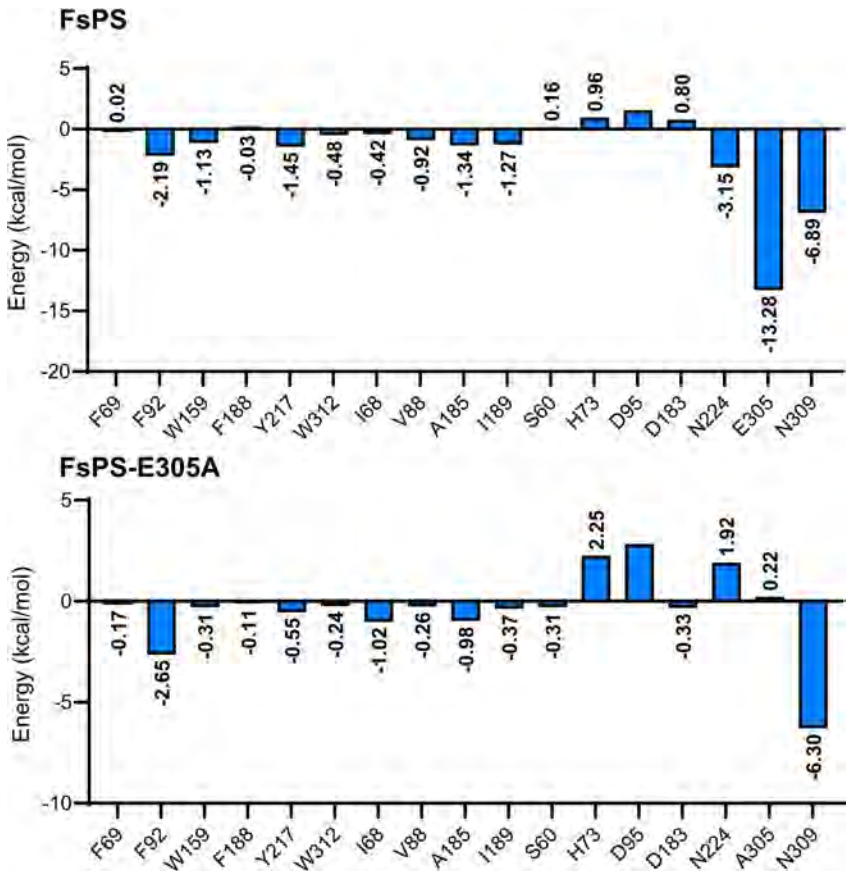

**Figure S77.** MM/PBSA energy decomposition analysis of key pocket residues of FsPS and FsPS-E305A interacting with **IM3**.

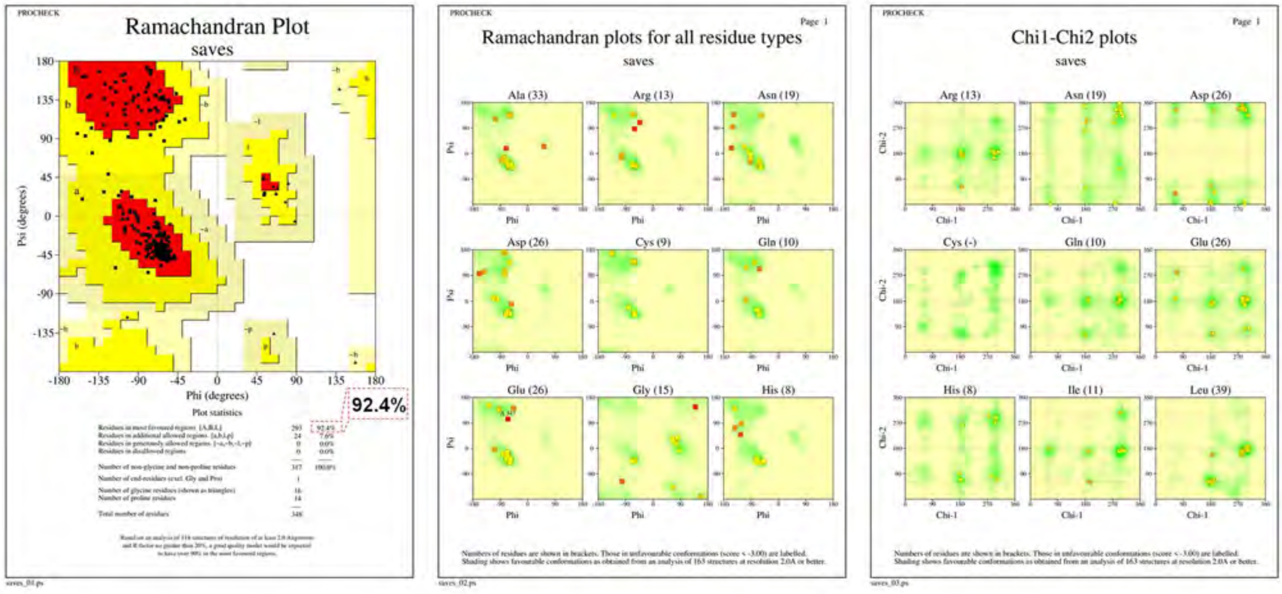

**Figure S78**. Ramachandran plot of the FsPS-E305A, as predicted using AlphaFold3.

### Cartesian Coordinates and Energies

**IM7**
Sum of electronic and thermal Free Energies
= -976.501352

| | | | |
|---|---|---|---|
| C | 1.89794700 | 0.29294400 | -0.26763800 |
| C | 3.00477200 | -0.81015300 | -0.47515700 |
| C | 2.61985300 | -2.14358600 | 0.17623500 |
| C | 1.26399600 | -2.63991500 | -0.28848200 |
| C | 0.09587500 | -1.73778300 | 0.14531500 |
| C | 0.36233900 | -0.22738600 | -0.12260100 |
| H | 3.39812000 | -2.87679300 | -0.06410400 |
| H | 1.98744400 | 0.92262200 | -1.19763500 |
| H | 1.26606700 | -2.74570300 | -1.38080900 |
| H | 1.06913200 | -3.64432600 | 0.10314400 |
| H | -0.15999800 | 0.41332400 | 0.58718800 |
| H | 2.62454300 | -2.04059100 | 1.26681400 |
| C | 2.40242000 | 1.30184200 | 0.83011900 |
| C | 4.24168600 | -0.19572300 | 0.24107000 |
| H | 4.42056600 | -0.72026000 | 1.18467300 |
| H | 5.14513400 | -0.31128300 | -0.36307300 |
| C | 3.90727400 | 1.26996100 | 0.50674300 |
| H | 4.49710400 | 1.70314200 | 1.31984200 |
| H | 4.09290000 | 1.87669600 | -0.38984200 |
| C | -1.15027300 | -2.23798400 | -0.64523600 |
| H | -1.40589200 | -3.21688300 | -0.22338400 |
| C | 0.32941500 | 0.29484900 | -1.45069100 |
| H | 0.68545000 | -0.38487700 | -2.22921500 |
| C | -2.36533600 | -1.35651500 | -0.71890100 |
| C | -2.04969800 | 2.16745900 | -0.14706000 |
| C | -0.32930800 | 1.51638400 | -2.00286000 |
| H | 0.32793100 | 1.96134000 | -2.76017100 |
| H | -1.11870100 | 1.02407900 | -2.60215600 |
| C | -3.18699400 | 1.63354700 | -0.60566900 |
| H | -3.30826700 | 1.56240000 | -1.69124400 |
| C | -0.98271600 | 2.61173000 | -1.14272000 |
| H | -0.21232400 | 3.19847400 | -0.64465700 |
| H | -1.44107300 | 3.29421000 | -1.86747900 |
| C | -4.32313200 | 1.04323800 | 0.17598600 |
| H | -4.12453800 | 1.06768500 | 1.25103700 |
| H | -5.23356000 | 1.62936300 | 0.00178900 |
| C | -3.45885500 | -1.36007500 | 0.05678800 |
| H | -2.38422900 | -0.67609800 | -1.57036400 |
| C | -4.60039600 | -0.41320000 | -0.26076500 |
| H | -5.51473900 | -0.75779800 | 0.23198100 |
| H | -4.80065200 | -0.42372500 | -1.33839100 |
| C | 3.33128400 | -1.00416800 | -1.96351700 |
| H | 2.49377300 | -1.39208200 | -2.55155700 |
| H | 3.65844300 | -0.06150500 | -2.41759900 |
| H | 4.14986100 | -1.72155700 | -2.06823900 |
| C | 2.13590700 | 0.78889800 | 2.24818500 |
| H | 2.59859000 | 1.47295400 | 2.96596600 |
| H | 1.06475800 | 0.75635100 | 2.47113600 |
| H | 2.55307200 | -0.20542400 | 2.42335400 |
| C | 1.86972000 | 2.72332100 | 0.70641300 |
| H | 2.45681900 | 3.37660400 | 1.35921300 |
| H | 1.96616400 | 3.10871400 | -0.31507800 |
| H | 0.82995600 | 2.80061300 | 1.02809800 |
| C | -0.17231000 | -1.88174100 | 1.64871000 |
| H | 0.68074600 | -1.56700200 | 2.25423800 |
| H | -1.03881200 | -1.27810300 | 1.94268900 |
| H | -0.38850200 | -2.92624200 | 1.89214200 |
| C | -1.79031400 | 2.45080800 | 1.31237900 |
| H | -2.68913800 | 2.34180400 | 1.91810000 |
| H | -1.02272800 | 1.79987100 | 1.75106500 |
| H | -1.43730700 | 3.48094000 | 1.43658000 |
| C | -3.66130600 | -2.25001100 | 1.25198700 |
| H | -2.82322000 | -2.92374500 | 1.43551400 |
| H | -3.81475800 | -1.64577800 | 2.15489500 |
| H | -4.56582600 | -2.85348400 | 1.12113300 |
| H | -0.83340200 | -2.43837700 | -1.67740100 |

**TS_7−8**
Sum of electronic and thermal Free Energies
= -976.497832

| | | | |
|---|---|---|---|
| C | 1.84543100 | 0.44593100 | -0.30422700 |
| C | 3.02175100 | -0.51049100 | -0.68658600 |
| C | 2.90347000 | -1.89961700 | -0.04524300 |
| C | 1.55083100 | -2.53818500 | -0.29936200 |
| C | 0.36810400 | -1.78275100 | 0.33843400 |
| C | 0.37898800 | -0.26588700 | -0.05483700 |
| H | 3.70062600 | -2.53403900 | -0.44957100 |
| H | 1.73906600 | 1.12961900 | -1.15913700 |
| H | 1.38935100 | -2.62822000 | -1.38215800 |
| H | 1.53057400 | -3.56532500 | 0.08366300 |
| H | -0.11762900 | 0.36194600 | 0.69363100 |
| H | 3.08977200 | -1.83473300 | 1.03134000 |
| C | 2.35433900 | 1.40644700 | 0.82841000 |
| C | 4.27412200 | 0.23164600 | -0.13410400 |
| H | 4.68526200 | -0.31781200 | 0.71859300 |
| H | 5.06722700 | 0.29113000 | -0.88427300 |
| C | 3.79190700 | 1.61472900 | 0.30816300 |
| H | 4.43051400 | 2.06895200 | 1.07186200 |
| H | 3.76678100 | 2.30347500 | -0.54632500 |
| C | -0.93089900 | -2.51448800 | -0.17686100 |
| H | -1.21789100 | -3.23385600 | 0.59678200 |
| C | -0.16761000 | -0.03471700 | -1.36642100 |
| H | 0.02312900 | -0.83539200 | -2.08879800 |
| C | -2.15416000 | -1.73897800 | -0.60331500 |
| C | -2.40875900 | 1.96213100 | -0.21839800 |

| | | | |
|---|---|---|---|
| C | -0.75914200 | 1.16523400 | -1.98340500 |
| H | 0.00103300 | 1.48625500 | -2.72144900 |
| H | -1.56255500 | 0.78737900 | -2.63220100 |
| C | -3.50302200 | 1.43389900 | -0.77358400 |
| H | -3.52920700 | 1.35369600 | -1.86366200 |
| C | -1.24043900 | 2.33364700 | -1.12040900 |
| H | -0.41172100 | 2.72005000 | -0.52796900 |
| H | -1.52716700 | 3.14092300 | -1.80327200 |
| C | -4.63177900 | 0.72161900 | -0.09832100 |
| H | -4.65223400 | 0.90037000 | 0.98020100 |
| H | -5.59858400 | 1.03970400 | -0.50009600 |
| C | -3.17132200 | -1.34177700 | 0.19121500 |
| H | -2.32898700 | -1.67034000 | -1.67929900 |
| C | -4.44885100 | -0.78822700 | -0.38844100 |
| H | -5.29510000 | -1.34317100 | 0.03384200 |
| H | -4.46742400 | -0.94653700 | -1.47224700 |
| C | 3.12233200 | -0.62919200 | -2.21409200 |
| H | 2.21449600 | -1.03708200 | -2.67419000 |
| H | 3.31140200 | 0.35193300 | -2.66388800 |
| H | 3.94876300 | -1.28929100 | -2.49383600 |
| C | 2.37219900 | 0.80451600 | 2.23386200 |
| H | 2.82496200 | 1.52594300 | 2.92084100 |
| H | 1.35980200 | 0.60298000 | 2.59814600 |
| H | 2.95680300 | -0.11572800 | 2.29616500 |
| C | 1.62084800 | 2.74278700 | 0.89283800 |
| H | 2.15550100 | 3.41419400 | 1.57195000 |
| H | 1.58214300 | 3.23181300 | -0.08655600 |
| H | 0.60399900 | 2.63816000 | 1.28531500 |
| C | 0.43144600 | -1.90901600 | 1.86699600 |
| H | 1.38568800 | -1.58049200 | 2.27499400 |
| H | -0.35750300 | -1.32226400 | 2.34867900 |
| H | 0.29546300 | -2.95583100 | 2.15393300 |
| C | -2.23775700 | 2.23729500 | 1.25256400 |
| H | -3.15637400 | 2.07617800 | 1.81776700 |
| H | -1.45179500 | 1.61782500 | 1.70521600 |
| H | -1.93748000 | 3.28081200 | 1.40280300 |
| C | -3.12496200 | -1.42759400 | 1.69157800 |
| H | -2.44402200 | -2.20432600 | 2.04560500 |
| H | -2.78696700 | -0.47259500 | 2.11523800 |
| H | -4.11922500 | -1.62390200 | 2.10229100 |
| H | -0.64288800 | -3.12615400 | -1.03865900 |

**IM8**

Sum of electronic and thermal Free Energies
= -976.542241

| | | | |
|---|---|---|---|
| C | 1.87804200 | 0.47143000 | -0.32831800 |
| C | 3.09205600 | -0.42852900 | -0.74350200 |
| C | 3.08264500 | -1.79279200 | -0.03853600 |
| C | 1.72556000 | -2.47930800 | -0.13442700 |
| C | 0.61036400 | -1.70040000 | 0.59147600 |
| C | 0.55120300 | -0.21998000 | 0.08112600 |
| H | 3.85793600 | -2.42858000 | -0.48274100 |
| H | 1.64866000 | 1.04868500 | -1.23499500 |
| H | 1.45551000 | -2.60930700 | -1.19086900 |
| H | 1.77362500 | -3.48965100 | 0.29083000 |
| H | 0.11267000 | 0.38715600 | 0.88611500 |
| H | 3.36029400 | -1.67279700 | 1.01462400 |
| C | 2.43664000 | 1.55551000 | 0.63697700 |
| C | 4.34000200 | 0.40709400 | -0.33688700 |
| H | 4.82374900 | -0.03486600 | 0.54104000 |
| H | 5.09082500 | 0.42517500 | -1.13239300 |
| C | 3.81415900 | 1.80716000 | -0.00031400 |
| H | 4.48338700 | 2.37265500 | 0.65628600 |
| H | 3.68356600 | 2.39497900 | -0.91911600 |
| C | -0.76575700 | -2.33008300 | 0.19679500 |
| H | -1.31594000 | -2.68992100 | 1.07238300 |
| C | -0.46711000 | -0.29625700 | -1.08368300 |
| H | 0.04878000 | -0.83330700 | -1.89253000 |
| C | -1.56352600 | -1.27938500 | -0.61398200 |
| C | -3.10658900 | 1.51769500 | -0.37656800 |
| C | -0.94983400 | 1.01074200 | -1.72670500 |
| H | -0.07707300 | 1.57056000 | -2.07491500 |
| H | -1.51356800 | 0.77811000 | -2.63867900 |
| C | -3.69022500 | 0.36320700 | -0.88846800 |
| H | -3.32606200 | 0.02750300 | -1.85703400 |
| C | -1.75643600 | 1.96246400 | -0.84310900 |
| H | -1.17609800 | 2.25140900 | 0.04324100 |
| H | -1.92868200 | 2.91125200 | -1.37495100 |
| C | -4.92417600 | -0.36438900 | -0.38607300 |
| H | -5.43908500 | 0.15548000 | 0.42217000 |
| H | -5.64579500 | -0.61148200 | -1.16412500 |
| C | -2.73856800 | -0.72783400 | 0.19237400 |
| H | -2.01228700 | -1.75084700 | -1.49599300 |
| C | -4.06086100 | -1.53716900 | 0.11947000 |
| H | -4.35486500 | -1.97008700 | 1.07885900 |
| H | -3.98135900 | -2.33039400 | -0.62762500 |
| C | 3.09283400 | -0.63169100 | -2.26745600 |
| H | 2.17210100 | -1.10184400 | -2.63015000 |
| H | 3.19459500 | 0.33136500 | -2.78021700 |
| H | 3.93232600 | -1.26452700 | -2.57346900 |
| C | 2.60275300 | 1.08778800 | 2.08695700 |
| H | 3.04582800 | 1.89157300 | 2.68426100 |
| H | 1.63855400 | 0.83550900 | 2.54189100 |
| H | 3.25713800 | 0.21646800 | 2.17261000 |
| C | 1.57480200 | 2.81914600 | 0.63819200 |
| H | 2.08107200 | 3.63094900 | 1.17066100 |
| H | 1.36707100 | 3.16560100 | -0.38183700 |
| H | 0.62045200 | 2.64987400 | 1.15492200 |
| C | 0.81024400 | -1.81650500 | 2.10766300 |
| H | 1.79739300 | -1.48153900 | 2.42695500 |
| H | 0.07276300 | -1.22077100 | 2.65752600 |
| H | 0.69985900 | -2.86098500 | 2.41834400 |

| | | | |
|---|---|---|---|
| C | -3.74311900 | 2.36603700 | 0.66547100 |
| H | -4.68351700 | 1.97578200 | 1.04873600 |
| H | -3.05479800 | 2.57140800 | 1.49085500 |
| H | -3.94677600 | 3.33461500 | 0.18766300 |
| C | -2.48955700 | -0.24511400 | 1.61104400 |
| H | -2.16609300 | -1.11397300 | 2.19126100 |
| H | -1.71401200 | 0.51808300 | 1.70121200 |
| H | -3.41015200 | 0.11768800 | 2.07580200 |
| H | -0.60527400 | -3.21002600 | -0.43373100 |

**TS_8−8-1**
Sum of electronic and thermal Free Energies
= -976.527339

| | | | |
|---|---|---|---|
| C | 2.14560300 | 0.21289000 | -0.41872100 |
| C | 3.34446200 | -0.75558600 | -0.06644200 |
| C | 2.92135300 | -1.98179100 | 0.74207400 |
| C | 1.65950300 | -2.60017600 | 0.15681000 |
| C | 0.44188600 | -1.68445400 | 0.37564800 |
| C | 0.73899600 | -0.15068200 | 0.16375100 |
| H | 3.74031900 | -2.71007900 | 0.75352000 |
| H | 2.03331300 | 0.15243600 | -1.51055700 |
| H | 1.81166200 | -2.78140300 | -0.91309700 |
| H | 1.45126700 | -3.57851800 | 0.60683800 |
| H | 0.65074300 | 0.32068100 | 1.14992500 |
| H | 2.75201900 | -1.69489300 | 1.78791300 |
| C | 2.67504400 | 1.67412400 | -0.10967300 |
| C | 4.35162800 | 0.11748800 | 0.70010700 |
| H | 4.08329300 | 0.16378600 | 1.76298100 |
| H | 5.36894800 | -0.28275200 | 0.64172800 |
| C | 4.20269700 | 1.48774400 | 0.05555800 |
| H | 4.64490400 | 2.29885300 | 0.64285200 |
| H | 4.69264900 | 1.50144100 | -0.92446200 |
| C | -0.65574500 | -2.03665400 | -0.65756700 |
| H | -1.26579800 | -2.89386600 | -0.35034300 |
| C | -0.40843000 | 0.37333600 | -0.77050900 |
| H | 0.05798100 | 0.44877600 | -1.76048700 |
| C | -1.46736700 | -0.76377900 | -0.97493300 |
| C | -3.33280100 | 1.38715200 | 0.42849300 |
| C | -0.95972600 | 1.78261100 | -0.49662800 |
| H | -0.12158400 | 2.44004100 | -0.27560800 |
| H | -1.40158400 | 2.19801800 | -1.41053400 |
| C | -3.67732600 | 0.80517400 | -0.78676400 |
| H | -3.07670600 | 1.07923800 | -1.65068900 |
| C | -1.95079800 | 1.91616800 | 0.65436900 |
| H | -1.55981300 | 1.43067700 | 1.56353200 |
| H | -2.07325700 | 2.97093900 | 0.94077700 |
| C | -4.90043800 | -0.02977200 | -1.11371800 |
| H | -5.62856700 | -0.07894600 | -0.30377800 |
| H | -5.41012800 | 0.25906500 | -2.03218700 |
| C | -2.86224300 | -0.73627200 | -0.32764600 |
| H | -1.69981900 | -0.76026100 | -2.04532400 |
| C | -4.02237900 | -1.29321700 | -1.20663100 |
| H | -4.45690300 | -2.21351600 | -0.80845900 |
| H | -3.68137100 | -1.46519300 | -2.22962300 |
| C | 4.00190400 | -1.21880600 | -1.38025900 |
| H | 3.30963800 | -1.79382100 | -2.00335900 |
| H | 4.33896100 | -0.36805200 | -1.98024400 |
| H | 4.87260000 | -1.84886300 | -1.17119300 |
| C | 2.15475900 | 2.25974200 | 1.21492400 |
| H | 2.64595200 | 3.21985100 | 1.40055300 |
| H | 1.07719500 | 2.44430200 | 1.22765100 |
| H | 2.38020200 | 1.60328300 | 2.06254900 |
| C | 2.41587300 | 2.64725100 | -1.26695500 |
| H | 2.86617800 | 3.62280600 | -1.05307000 |
| H | 2.86949700 | 2.26719200 | -2.18855400 |
| H | 1.35472200 | 2.81402800 | -1.47263700 |
| C | -0.01770400 | -1.90639800 | 1.82813200 |
| H | 0.84224800 | -1.87525000 | 2.50257400 |
| H | -0.70330000 | -1.13401400 | 2.17834700 |
| H | -0.49591100 | -2.88454100 | 1.95103700 |
| C | -4.27160500 | 1.50261000 | 1.57590500 |
| H | -5.18760700 | 0.92579700 | 1.46638700 |
| H | -3.78638700 | 1.26831100 | 2.52713200 |
| H | -4.55284300 | 2.56491000 | 1.61917700 |
| C | -3.06373700 | -1.20815000 | 1.10129800 |
| H | -2.72065300 | -2.24545900 | 1.14778300 |
| H | -2.51154700 | -0.64577700 | 1.85204900 |
| H | -4.12396200 | -1.20821100 | 1.36708600 |
| H | -0.16700400 | -2.33433600 | -1.59193100 |

**IM8-1**
Sum of electronic and thermal Free Energies
= -976.575728

| | | | |
|---|---|---|---|
| C | 2.15513100 | 0.22566600 | -0.41699700 |
| C | 3.30645800 | -0.81724400 | -0.16681800 |
| C | 2.94041400 | -1.80147700 | 0.94555400 |
| C | 1.63830000 | -2.52370200 | 0.61683600 |
| C | 0.44603000 | -1.53913600 | 0.56363700 |
| C | 0.82340100 | -0.04060600 | 0.33632900 |
| H | 3.75512200 | -2.52210000 | 1.08311300 |
| H | 1.92649200 | 0.17205100 | -1.49255100 |
| H | 1.74947800 | -3.05076200 | -0.33623800 |
| H | 1.43154200 | -3.29863100 | 1.36311300 |
| H | 0.89856700 | 0.42319900 | 1.32685800 |
| H | 2.83670600 | -1.26000800 | 1.89785100 |
| C | 2.79103600 | 1.64188000 | -0.18014100 |
| C | 4.55234700 | 0.02562800 | 0.18303300 |
| H | 4.64975600 | 0.14699100 | 1.26805400 |
| H | 5.47314500 | -0.44954500 | -0.16949200 |
| C | 4.27846300 | 1.37251700 | -0.48014400 |
| H | 4.92403100 | 2.18095900 | -0.12211500 |
| H | 4.42445300 | 1.29640100 | -1.56580000 |

| | | | |
|---|---|---|---|
| C | -0.47697200 | -1.85362400 | -0.62810500 |
| H | -1.06461700 | -2.76670300 | -0.48585200 |
| C | -0.38448000 | 0.59263700 | -0.44422600 |
| H | 0.05073200 | 0.97339200 | -1.37275600 |
| C | -1.31408100 | -0.58934000 | -0.93873300 |
| C | -3.47872700 | 1.18647800 | 0.57027100 |
| C | -1.02199600 | 1.84719900 | 0.19444400 |
| H | -0.23415400 | 2.37116200 | 0.74041600 |
| H | -1.33019600 | 2.55048700 | -0.58893700 |
| C | -3.57911500 | 0.92147700 | -0.79055500 |
| H | -2.84084800 | 1.38693700 | -1.44003000 |
| C | -2.18864100 | 1.66394100 | 1.15804900 |
| H | -1.93164400 | 0.98612400 | 1.98475800 |
| H | -2.41976900 | 2.61843500 | 1.65316900 |
| C | -4.69528100 | 0.20028900 | -1.51937800 |
| H | -5.55818300 | -0.02194000 | -0.89142000 |
| H | -5.03532200 | 0.70036100 | -2.42548500 |
| C | -2.80493300 | -0.69902300 | -0.55989000 |
| H | -1.35863700 | -0.47825800 | -2.02593600 |
| C | -3.78025300 | -1.01937000 | -1.73526800 |
| H | -4.25365700 | -1.99975800 | -1.63961300 |
| H | -3.26079600 | -0.95966800 | -2.69357300 |
| C | 3.59600200 | -1.57544100 | -1.47484300 |
| H | 2.69437700 | -2.01425900 | -1.91421300 |
| H | 4.01782300 | -0.89659300 | -2.22337100 |
| H | 4.31919800 | -2.38078400 | -1.30609700 |
| C | 2.66823100 | 2.12736500 | 1.27397000 |
| H | 3.33313200 | 2.98263600 | 1.43096600 |
| H | 1.65589700 | 2.46283900 | 1.51942300 |
| H | 2.94296000 | 1.35435000 | 1.99887400 |
| C | 2.24315800 | 2.72117800 | -1.11825900 |
| H | 2.81714900 | 3.64737700 | -1.00542500 |
| H | 2.31991200 | 2.40666600 | -2.16503000 |
| H | 1.19577800 | 2.96647100 | -0.91075400 |
| C | -0.25826500 | -1.63164600 | 1.92518700 |
| H | 0.48939600 | -1.54943000 | 2.72141300 |
| H | -0.96903500 | -0.81723100 | 2.08408900 |
| H | -0.77918000 | -2.58699600 | 2.05122600 |
| C | -4.60533100 | 1.01342300 | 1.52642600 |
| H | -5.46627000 | 0.48907500 | 1.11749100 |
| H | -4.28046400 | 0.53960400 | 2.45676800 |
| H | -4.92974400 | 2.03071900 | 1.78818800 |
| C | -3.23556600 | -1.50635800 | 0.65444000 |
| H | -2.90144700 | -2.53382600 | 0.48405000 |
| H | -2.81676400 | -1.17974500 | 1.60373900 |
| H | -4.32479100 | -1.54016300 | 0.73537800 |
| H | 0.15789200 | -2.03611100 | -1.50317600 |

**TS_8-1−8-2**
Sum of electronic and thermal Free Energies
= -976.570287

| | | | |
|---|---|---|---|
| C | 2.04747000 | 0.52920000 | -0.28998000 |
| C | 3.07213000 | -0.43031000 | -0.99078000 |
| C | 2.87346000 | -1.90438000 | -0.60421000 |
| C | 1.41409000 | -2.34062000 | -0.68643000 |
| C | 0.52904000 | -1.58690000 | 0.32450000 |
| C | 0.66837000 | -0.04879000 | 0.11486000 |
| H | 3.48988000 | -2.53002000 | -1.26039000 |
| H | 1.83139000 | 1.30178000 | -1.04344000 |
| H | 1.03587000 | -2.17874000 | -1.70556000 |
| H | 1.32470000 | -3.41765000 | -0.49641000 |
| H | 0.36223000 | 0.44571000 | 1.05081000 |
| H | 3.24698000 | -2.07793000 | 0.41123000 |
| C | 2.82789000 | 1.31183000 | 0.80199000 |
| C | 4.45909000 | 0.09258000 | -0.52053000 |
| H | 4.88987000 | -0.58972000 | 0.22046000 |
| H | 5.17185000 | 0.14724000 | -1.34884000 |
| C | 4.19584000 | 1.46256000 | 0.11380000 |
| H | 4.98320000 | 1.77419000 | 0.80783000 |
| H | 4.12380000 | 2.23317000 | -0.66548000 |
| C | -0.97059000 | -1.89429000 | 0.01970000 |
| H | -1.49098000 | -2.24109000 | 0.91472000 |
| C | -0.39701000 | 0.27774000 | -0.95237000 |
| H | -0.01113000 | -0.05510000 | -1.92727000 |
| C | -1.58277000 | -0.64786000 | -0.65414000 |
| C | -2.78555000 | 0.66010000 | -0.04585000 |
| C | -0.85214000 | 1.73253000 | -1.03529000 |
| H | -0.02441000 | 2.43405000 | -0.90327000 |
| H | -1.30333000 | 1.93862000 | -2.01185000 |
| C | -3.96611000 | 0.61398000 | -1.02461000 |
| H | -3.62032000 | 0.73650000 | -2.05758000 |
| C | -1.90621000 | 1.87940000 | 0.06794000 |
| H | -1.42797000 | 1.92721000 | 1.05075000 |
| H | -2.50384000 | 2.78695000 | -0.04406000 |
| C | -4.58200000 | -0.78757000 | -0.78952000 |
| H | -5.66859000 | -0.76652000 | -0.89099000 |
| H | -4.21164000 | -1.51568000 | -1.51535000 |
| C | -2.98559000 | -0.28792000 | 0.95602000 |
| H | -2.18458000 | -0.91393000 | -1.51911000 |
| C | -4.14314000 | -1.18141000 | 0.63311000 |
| H | -4.91422000 | -0.95786000 | 1.38518000 |
| H | -3.89572000 | -2.24115000 | 0.76145000 |
| C | 2.96451000 | -0.28171000 | -2.51740000 |
| H | 1.96658000 | -0.53273000 | -2.89510000 |
| H | 3.18018000 | 0.74914000 | -2.81955000 |
| H | 3.68305000 | -0.93648000 | -3.02119000 |
| C | 2.98238000 | 0.55626000 | 2.12607000 |
| H | 3.59048000 | 1.14820000 | 2.81772000 |
| H | 2.01356000 | 0.38519000 | 2.60915000 |
| H | 3.47680000 | -0.41136000 | 2.00144000 |
| C | 2.17820000 | 2.66765000 | 1.08401000 |

| | | | |
|---|---|---|---|
| H | 2.76501000 | 3.23773000 | 1.81142000 |
| H | 2.10450000 | 3.26821000 | 0.17007000 |
| H | 1.16860000 | 2.55272000 | 1.50237000 |
| C | 0.85168000 | -2.06028000 | 1.74651000 |
| H | 1.90585000 | -1.94685000 | 1.99737000 |
| H | 0.27891000 | -1.50291000 | 2.49566000 |
| H | 0.60012000 | -3.12170000 | 1.85043000 |
| C | -4.96291000 | 1.73291000 | -0.69905000 |
| H | -5.83464000 | 1.64623000 | -1.35183000 |
| H | -5.31170000 | 1.66549000 | 0.33802000 |
| H | -4.53507000 | 2.72600000 | -0.85206000 |
| C | -2.35106000 | -0.31838000 | 2.29707000 |
| H | -3.01932000 | 0.24929000 | 2.95990000 |
| H | -2.28668000 | -1.33347000 | 2.69577000 |
| H | -1.37098000 | 0.15712000 | 2.32424000 |
| H | -1.05287000 | -2.71775000 | -0.69905000 |

**IM8-2**
Sum of electronic and thermal Free Energies
= -976.573703

| | | | |
|---|---|---|---|
| C | 0.36061300 | -0.36403100 | 0.86595000 |
| C | 1.24052300 | 0.37764300 | -0.17042100 |
| C | 0.92080600 | 1.84716600 | 0.10567100 |
| C | -0.60328300 | 1.87371000 | 0.41056300 |
| C | -0.98510100 | 0.38921100 | 0.77416500 |
| C | -1.39823900 | 2.36946000 | -0.80824300 |
| C | -2.87559000 | 2.01221500 | -0.66871600 |
| C | -3.08229900 | 0.49780300 | -0.81623800 |
| C | -1.89382600 | -0.34035700 | -0.24585900 |
| C | -4.29074300 | -0.02967900 | -0.02994300 |
| C | -4.05641600 | -1.53630300 | 0.01794400 |
| C | -2.53490900 | -1.69032500 | 0.29223300 |
| H | -1.46847600 | 0.39292100 | 1.75712500 |
| C | -3.24516900 | 0.17381700 | -2.31146100 |
| H | -1.24421200 | -0.60175500 | -1.09668300 |
| C | -2.35455600 | -1.92162600 | 1.80196700 |
| C | -1.97577000 | -2.90578400 | -0.45742500 |
| H | 0.23430300 | -1.40883200 | 0.57334200 |
| C | 1.14643400 | -0.28808700 | 2.18711900 |
| C | 2.64407000 | -0.15593400 | 1.80281300 |
| C | 2.71751400 | -0.23982400 | 0.27333500 |
| C | -0.85321400 | 2.81449500 | 1.59850900 |
| C | 2.89409100 | -1.62738800 | -0.40598300 |
| C | 3.09473800 | -1.27422000 | -1.89625600 |
| C | 3.75393600 | 0.11449200 | -1.86840800 |
| C | 3.47105400 | 0.67673900 | -0.52256200 |
| C | 4.10663600 | -2.37514800 | 0.15569300 |
| C | 4.00283700 | 1.99010600 | -0.12342000 |
| H | 1.00728700 | 0.09810700 | -1.19991300 |
| H | 1.18439100 | 2.51754200 | -0.72039200 |
| H | 1.46302800 | 2.18777700 | 0.99485800 |
| H | -1.25750000 | 3.45234100 | -0.90874800 |
| H | -1.00019300 | 1.91530400 | -1.72773200 |
| H | -3.47237800 | 2.53876700 | -1.42150300 |
| H | -3.24520700 | 2.35322000 | 0.30563900 |
| H | -5.24629100 | 0.25336600 | -0.48350900 |
| H | -4.27433500 | 0.38618500 | 0.98739900 |
| H | -4.65256900 | -2.03949100 | 0.78542600 |
| H | -4.32430200 | -1.99312300 | -0.94081000 |
| H | -4.17469100 | 0.60448600 | -2.69776800 |
| H | -2.41484600 | 0.59161300 | -2.89260000 |
| H | -3.26630300 | -0.90244500 | -2.50382500 |
| H | -1.30401000 | -1.96465100 | 2.10835600 |
| H | -2.80868300 | -2.88052600 | 2.06948700 |
| H | -2.85147400 | -1.14956900 | 2.39887300 |
| H | -2.07098200 | -2.77887200 | -1.54120300 |
| H | -0.91577500 | -3.07579000 | -0.22899700 |
| H | -2.51974000 | -3.81408300 | -0.17721100 |
| H | 0.96267900 | -1.17352600 | 2.79982800 |
| H | 0.83126000 | 0.57339700 | 2.78122100 |
| H | 3.05273000 | 0.79308200 | 2.16033300 |
| H | 3.26454100 | -0.93844800 | 2.24279600 |
| H | -0.33109100 | 2.46783700 | 2.49645000 |
| H | -0.49726300 | 3.82509800 | 1.36966500 |
| H | -1.91567000 | 2.88191900 | 1.84545200 |
| H | 2.00023200 | -2.24057900 | -0.26401400 |
| H | 2.14388900 | -1.23509100 | -2.43213400 |
| H | 3.71898300 | -2.01031700 | -2.40574000 |
| H | 4.85299900 | 0.05315600 | -1.93182900 |
| H | 3.45926300 | 0.80370400 | -2.66868600 |
| H | 4.26465300 | -3.29630700 | -0.41015700 |
| H | 3.97428500 | -2.65543600 | 1.20250800 |
| H | 5.02258700 | -1.77590100 | 0.07913700 |
| H | 5.06942300 | 1.80677900 | 0.08766600 |
| H | 3.96533600 | 2.71376100 | -0.94163200 |
| H | 3.55930700 | 2.39546900 | 0.78334800 |

**TS_8-2−12**
Sum of electronic and thermal Free Energies
= -976.486910

| | | | |
|---|---|---|---|
| C | 0.40115000 | -0.27199000 | 0.88689000 |
| C | 1.34193000 | 0.52973000 | 0.04833000 |
| C | 0.83540000 | 1.97391000 | 0.10165000 |
| C | -0.67200000 | 1.90210000 | 0.48850000 |
| C | -0.97121000 | 0.39453000 | 0.81776000 |
| C | -1.55458000 | 2.39792000 | -0.67102000 |
| C | -2.99030000 | 1.90622000 | -0.50446000 |
| C | -3.06772000 | 0.39507000 | -0.76296000 |
| C | -1.81119000 | -0.36775000 | -0.23306000 |
| C | -4.23014000 | -0.29110000 | -0.03097000 |
| C | -3.85038000 | -1.76850000 | -0.06400000 |
| C | -2.32942000 | -1.78941000 | 0.24807000 |

| | | | |
|---|---|---|---|
| H | -1.46686000 | 0.34334000 | 1.79322000 |
| C | -3.19494000 | 0.16010000 | -2.27801000 |
| H | -1.13817000 | -0.52972000 | -1.09273000 |
| C | -2.17436000 | -2.04545000 | 1.75784000 |
| C | -1.63842000 | -2.92386000 | -0.51806000 |
| H | 0.37663000 | -1.34186000 | 0.69526000 |
| C | 1.32752000 | 0.01423000 | 2.19281000 |
| C | 2.78706000 | 0.49304000 | 1.76483000 |
| C | 2.72955000 | 0.06351000 | 0.29264000 |
| C | -0.91573000 | 2.79243000 | 1.71664000 |
| C | 2.96269000 | -1.48221000 | 0.15945000 |
| C | 2.76523000 | -1.79276000 | -1.32927000 |
| C | 3.40763000 | -0.58951000 | -2.01934000 |
| C | 2.99759000 | 0.57475000 | -1.13808000 |
| C | 4.38859000 | -1.82877000 | 0.61038000 |
| C | 3.51682000 | 1.95073000 | -1.44300000 |
| H | 1.73550000 | 0.53455000 | -1.40066000 |
| H | 0.97623000 | 2.50176000 | -0.84446000 |
| H | 1.40348000 | 2.53192000 | 0.85502000 |
| H | -1.50619000 | 3.49239000 | -0.71427000 |
| H | -1.15814000 | 2.02950000 | -1.62767000 |
| H | -3.66119000 | 2.43212000 | -1.19219000 |
| H | -3.34543000 | 2.13782000 | 0.50732000 |
| H | -5.20113000 | -0.07668000 | -0.48871000 |
| H | -4.27226000 | 0.06416000 | 1.00811000 |
| H | -4.41242000 | -2.37326000 | 0.65417000 |
| H | -4.04705000 | -2.19005000 | -1.05549000 |
| H | -4.15594000 | 0.53579000 | -2.64367000 |
| H | -2.40135000 | 0.68033000 | -2.82595000 |
| H | -3.12599000 | -0.89967000 | -2.53905000 |
| H | -1.13634000 | -2.00323000 | 2.10578000 |
| H | -2.54624000 | -3.04900000 | 1.98471000 |
| H | -2.75987000 | -1.33974000 | 2.35637000 |
| H | -1.71350000 | -2.77073000 | -1.60011000 |
| H | -0.57378000 | -3.00823000 | -0.26257000 |
| H | -2.10477000 | -3.88614000 | -0.28239000 |
| H | 1.38357000 | -0.92630000 | 2.74498000 |
| H | 0.83540000 | 0.75077000 | 2.82759000 |
| H | 2.90564000 | 1.56847000 | 1.90396000 |
| H | 3.55886000 | -0.00545000 | 2.34845000 |
| H | -0.32934000 | 2.46749000 | 2.58145000 |
| H | -0.64211000 | 3.83007000 | 1.49789000 |
| H | -1.96775000 | 2.78028000 | 2.01303000 |
| H | 2.26515000 | -2.05130000 | 0.77968000 |
| H | 1.69770000 | -1.83621000 | -1.58383000 |
| H | 3.21219000 | -2.74431000 | -1.62227000 |
| H | 4.50281000 | -0.65318000 | -1.98871000 |
| H | 3.11292000 | -0.44422000 | -3.06137000 |
| H | 4.62827000 | -2.85398000 | 0.31708000 |
| H | 4.50549000 | -1.76801000 | 1.69378000 |
| H | 5.13619000 | -1.16744000 | 0.15972000 |
| H | 4.61054000 | 1.87707000 | -1.40483000 |
| H | 3.23943000 | 2.28023000 | -2.44730000 |
| H | 3.21520000 | 2.69881000 | -0.71062000 |

**IM12**
Sum of electronic and thermal Free Energies
= -976.583765

| | | | |
|---|---|---|---|
| C | -0.33022500 | -0.79303000 | -0.47619300 |
| C | -1.46037400 | -0.12709500 | 0.35621400 |
| C | -0.83851700 | 1.14743200 | 0.96560900 |
| C | 0.18458400 | 1.59699600 | -0.08590100 |
| C | 0.82340300 | 0.25336100 | -0.56502700 |
| C | 1.20921700 | 2.58935300 | 0.50108600 |
| C | 2.59574100 | 2.33133300 | -0.08261900 |
| C | 3.13133900 | 0.96722400 | 0.37378200 |
| C | 2.04769300 | -0.16330600 | 0.27520200 |
| C | 4.27242100 | 0.45519100 | -0.51930000 |
| C | 4.30029500 | -1.04406100 | -0.24336000 |
| C | 2.80796700 | -1.46245800 | -0.22012200 |
| H | 1.13736000 | 0.36906800 | -1.60982500 |
| C | 3.62855900 | 1.07425700 | 1.82521100 |
| H | 1.67432300 | -0.36525900 | 1.29272400 |
| C | 2.43044700 | -1.90757800 | -1.64298400 |
| C | 2.60045600 | -2.64060300 | 0.73813400 |
| H | 0.04449600 | -1.66985800 | 0.05621100 |
| C | -0.99171200 | -1.23476900 | -1.81228400 |
| C | -2.44331800 | -0.71892400 | -1.78200900 |
| C | -2.58531400 | 0.07151500 | -0.54877400 |
| C | -0.53128600 | 2.26179200 | -1.26914400 |
| C | -2.10486300 | -1.06938500 | 1.49366600 |
| C | -3.31919800 | -0.35891500 | 2.09634300 |
| C | -4.37691500 | -0.04722700 | 1.04306800 |
| C | -3.78906000 | 0.80520500 | -0.15540100 |
| C | -2.42557300 | -2.49165900 | 1.03231200 |
| C | -4.82301100 | 1.03734700 | -1.25049200 |
| H | -3.47386200 | 1.75448500 | 0.29710400 |
| H | -0.32842000 | 0.86450800 | 1.89430700 |
| H | -1.56330300 | 1.92893000 | 1.21365200 |
| H | 0.88374600 | 3.61620300 | 0.30161400 |
| H | 1.25107700 | 2.48933000 | 1.59164700 |
| H | 3.29394300 | 3.11998800 | 0.21901100 |
| H | 2.54263700 | 2.36765200 | -1.17974900 |
| H | 5.22254100 | 0.95782800 | -0.31158200 |
| H | 4.02977800 | 0.63919700 | -1.57483800 |
| H | 4.86274300 | -1.61072700 | -0.99187200 |
| H | 4.77058200 | -1.24452900 | 0.72541400 |
| H | 4.46315900 | 1.77940900 | 1.89532400 |
| H | 2.83772400 | 1.41980400 | 2.49911900 |
| H | 3.97177700 | 0.10820300 | 2.20658500 |
| H | 1.38531300 | -2.21720400 | -1.72923700 |
| H | 3.04158700 | -2.77217400 | -1.91905200 |

| | | | |
|---|---|---|---|
| H | 2.61849800 | -1.12517700 | -2.38605100 |
| H | 2.85371500 | -2.35806600 | 1.76574100 |
| H | 1.56477300 | -3.00287300 | 0.73297100 |
| H | 3.24083000 | -3.48271000 | 0.45462700 |
| H | -0.97568800 | -2.32096700 | -1.92837700 |
| H | -0.45849200 | -0.81630800 | -2.67020300 |
| H | -2.75464200 | -0.12461500 | -2.65136200 |
| H | -3.19944000 | -1.52333600 | -1.73092300 |
| H | -1.18869000 | 1.57131900 | -1.81726000 |
| H | -1.13308600 | 3.11632100 | -0.94040000 |
| H | 0.19853600 | 2.62480300 | -2.00017800 |
| H | -1.29549500 | -1.12847600 | 2.23112800 |
| H | -3.00718300 | 0.56491800 | 2.59494000 |
| H | -3.76012400 | -0.99943700 | 2.86813700 |
| H | -4.81769500 | -0.96132600 | 0.63053700 |
| H | -5.19794700 | 0.54366300 | 1.45745500 |
| H | -2.71097300 | -3.08987600 | 1.90186700 |
| H | -1.56142600 | -2.97920400 | 0.57479400 |
| H | -3.25709800 | -2.54102400 | 0.32299700 |
| H | -5.18437000 | 0.09249200 | -1.66757400 |
| H | -5.68320000 | 1.56869200 | -0.83811100 |
| H | -4.41455100 | 1.64436400 | -2.06340700 |

**TS_8−9**
Sum of electronic and thermal Free Energies
= -976.521486

| | | | |
|---|---|---|---|
| C | 2.14560000 | 0.21289000 | -0.41872000 |
| C | 3.34446000 | -0.75559000 | -0.06644000 |
| C | 2.92135000 | -1.98179000 | 0.74207000 |
| C | 1.65950000 | -2.60018000 | 0.15681000 |
| C | 0.44189000 | -1.68445000 | 0.37565000 |
| C | 0.73900000 | -0.15068000 | 0.16375000 |
| H | 3.74032000 | -2.71008000 | 0.75352000 |
| H | 2.03331000 | 0.15244000 | -1.51056000 |
| H | 1.81166000 | -2.78140000 | -0.91310000 |
| H | 1.45127000 | -3.57852000 | 0.60684000 |
| H | 0.65074000 | 0.32068000 | 1.14993000 |
| H | 2.75202000 | -1.69489000 | 1.78791000 |
| C | 2.67504000 | 1.67412000 | -0.10967000 |
| C | 4.35163000 | 0.11749000 | 0.70011000 |
| H | 4.08329000 | 0.16379000 | 1.76298000 |
| H | 5.36895000 | -0.28275000 | 0.64173000 |
| C | 4.20270000 | 1.48774000 | 0.05556000 |
| H | 4.64490000 | 2.29885000 | 0.64285000 |
| H | 4.69265000 | 1.50144000 | -0.92446000 |
| C | -0.65575000 | -2.03665000 | -0.65757000 |
| H | -1.26580000 | -2.89387000 | -0.35034000 |
| C | -0.40843000 | 0.37334000 | -0.77051000 |
| H | 0.05798000 | 0.44878000 | -1.76049000 |
| C | -1.46737000 | -0.76378000 | -0.97493000 |
| C | -3.33280000 | 1.38715000 | 0.42849000 |
| C | -0.95973000 | 1.78261000 | -0.49663000 |
| H | -0.12158000 | 2.44004000 | -0.27561000 |
| H | -1.40158000 | 2.19802000 | -1.41053000 |
| C | -3.67733000 | 0.80517000 | -0.78676000 |
| H | -3.07671000 | 1.07924000 | -1.65069000 |
| C | -1.95080000 | 1.91617000 | 0.65437000 |
| H | -1.55981000 | 1.43068000 | 1.56353000 |
| H | -2.07326000 | 2.97094000 | 0.94078000 |
| C | -4.90044000 | -0.02977000 | -1.11372000 |
| H | -5.62857000 | -0.07895000 | -0.30378000 |
| H | -5.41013000 | 0.25907000 | -2.03219000 |
| C | -2.86224000 | -0.73627000 | -0.32765000 |
| H | -1.69982000 | -0.76026000 | -2.04532000 |
| C | -4.02238000 | -1.29322000 | -1.20663000 |
| H | -4.45690000 | -2.21352000 | -0.80846000 |
| H | -3.68137000 | -1.46519000 | -2.22962000 |
| C | 4.00190000 | -1.21881000 | -1.38026000 |
| H | 3.30964000 | -1.79382000 | -2.00336000 |
| H | 4.33896000 | -0.36805000 | -1.98024000 |
| H | 4.87260000 | -1.84886000 | -1.17119000 |
| C | 2.15476000 | 2.25974000 | 1.21492000 |
| H | 2.64595000 | 3.21985000 | 1.40055000 |
| H | 1.07720000 | 2.44430000 | 1.22765000 |
| H | 2.38020000 | 1.60328000 | 2.06255000 |
| C | 2.41587000 | 2.64725000 | -1.26696000 |
| H | 2.86618000 | 3.62281000 | -1.05307000 |
| H | 2.86950000 | 2.26719000 | -2.18855000 |
| H | 1.35472000 | 2.81403000 | -1.47264000 |
| C | -0.01770000 | -1.90640000 | 1.82813000 |
| H | 0.84225000 | -1.87525000 | 2.50257000 |
| H | -0.70330000 | -1.13401000 | 2.17835000 |
| H | -0.49591000 | -2.88454000 | 1.95104000 |
| C | -4.27161000 | 1.50261000 | 1.57591000 |
| H | -5.18761000 | 0.92580000 | 1.46639000 |
| H | -3.78639000 | 1.26831000 | 2.52713000 |
| H | -4.55284000 | 2.56491000 | 1.61918000 |
| C | -3.06374000 | -1.20815000 | 1.10130000 |
| H | -2.72065000 | -2.24546000 | 1.14778000 |
| H | -2.51155000 | -0.64578000 | 1.85205000 |
| H | -4.12396000 | -1.20821000 | 1.36709000 |
| H | -0.16700000 | -2.33434000 | -1.59193000 |

**IM9**
Sum of electronic and thermal Free Energies
= -976.523131

| | | | |
|---|---|---|---|
| C | 2.15513100 | 0.22566600 | -0.41699700 |
| C | 3.30645800 | -0.81724400 | -0.16681800 |
| C | 2.94041400 | -1.80147700 | 0.94555400 |
| C | 1.63830000 | -2.52370200 | 0.61683600 |
| C | 0.44603000 | -1.53913600 | 0.56363700 |
| C | 0.82340100 | -0.04060600 | 0.33632900 |

| | | | |
|---|---|---|---|
| H | 3.75512200 | -2.52210000 | 1.08311300 |
| H | 1.92649200 | 0.17205100 | -1.49255100 |
| H | 1.74947800 | -3.05076200 | -0.33623800 |
| H | 1.43154200 | -3.29863100 | 1.36311300 |
| H | 0.89856700 | 0.42319900 | 1.32685800 |
| H | 2.83670600 | -1.26000800 | 1.89785100 |
| C | 2.79103600 | 1.64188000 | -0.18014100 |
| C | 4.55234700 | 0.02562800 | 0.18303300 |
| H | 4.64975600 | 0.14699100 | 1.26805400 |
| H | 5.47314500 | -0.44954500 | -0.16949200 |
| C | 4.27846300 | 1.37251700 | -0.48014400 |
| H | 4.92403100 | 2.18095900 | -0.12211500 |
| H | 4.42445300 | 1.29640100 | -1.56580000 |
| C | -0.47697200 | -1.85362400 | -0.62810500 |
| H | -1.06461700 | -2.76670300 | -0.48585200 |
| C | -0.38448000 | 0.59263700 | -0.44422600 |
| H | 0.05073200 | 0.97339200 | -1.37275600 |
| C | -1.31408100 | -0.58934000 | -0.93873300 |
| C | -3.47872700 | 1.18647800 | 0.57027100 |
| C | -1.02199600 | 1.84719900 | 0.19444400 |
| H | -0.23415400 | 2.37116200 | 0.74041600 |
| H | -1.33019600 | 2.55048700 | -0.58893700 |
| C | -3.57911500 | 0.92147700 | -0.79055500 |
| H | -2.84084800 | 1.38693700 | -1.44003000 |
| C | -2.18864100 | 1.66394100 | 1.15804900 |
| H | -1.93164400 | 0.98612400 | 1.98475800 |
| H | -2.41976900 | 2.61843500 | 1.65316900 |
| C | -4.69528100 | 0.20028900 | -1.51937800 |
| H | -5.55818300 | -0.02194000 | -0.89142000 |
| H | -5.03532200 | 0.70036100 | -2.42548500 |
| C | -2.80493300 | -0.69902300 | -0.55989000 |
| H | -1.35863700 | -0.47825800 | -2.02593600 |
| C | -3.78025300 | -1.01937000 | -1.73526800 |
| H | -4.25365700 | -1.99975800 | -1.63961300 |
| H | -3.26079600 | -0.95966800 | -2.69357300 |
| C | 3.59600200 | -1.57544100 | -1.47484300 |
| H | 2.69437700 | -2.01425900 | -1.91421300 |
| H | 4.01782300 | -0.89659300 | -2.22337100 |
| H | 4.31919800 | -2.38078400 | -1.30609700 |
| C | 2.66823100 | 2.12736500 | 1.27397000 |
| H | 3.33313200 | 2.98263600 | 1.43096600 |
| H | 1.65589700 | 2.46283900 | 1.51942300 |
| H | 2.94296000 | 1.35435000 | 1.99887400 |
| C | 2.24315800 | 2.72117800 | -1.11825900 |
| H | 2.81714900 | 3.64737700 | -1.00542500 |
| H | 2.31991200 | 2.40666600 | -2.16503000 |
| H | 1.19577800 | 2.96647100 | -0.91075400 |
| C | -0.25826500 | -1.63164600 | 1.92518700 |
| H | 0.48939600 | -1.54943000 | 2.72141300 |
| H | -0.96903500 | -0.81723100 | 2.08408900 |
| H | -0.77918000 | -2.58699600 | 2.05122600 |
| C | -4.60533100 | 1.01342300 | 1.52642600 |
| H | -5.46627000 | 0.48907500 | 1.11749100 |
| H | -4.28046400 | 0.53960400 | 2.45676800 |
| H | -4.92974400 | 2.03071900 | 1.78818800 |
| C | -3.23556600 | -1.50635800 | 0.65444000 |
| H | -2.90144700 | -2.53382600 | 0.48405000 |
| H | -2.81676400 | -1.17974500 | 1.60373900 |
| H | -4.32479100 | -1.54016300 | 0.73537800 |
| H | 0.15789200 | -2.03611100 | -1.50317600 |

**TS_9−10**
Sum of electronic and thermal Free Energies
= -976.512624

| | | | |
|---|---|---|---|
| C | 0.59394000 | -0.49509000 | -0.76798000 |
| C | 1.37090000 | -1.80555000 | -1.18395000 |
| C | 2.76321000 | -2.12964000 | -0.61292000 |
| C | 3.73770000 | -0.97257000 | -0.64060000 |
| C | 2.22552000 | 1.28665000 | 0.37253000 |
| C | 1.19356000 | 0.30451000 | 0.40257000 |
| C | 3.59243000 | -0.01955000 | -1.56880000 |
| C | 3.94657000 | 1.42846000 | -1.39743000 |
| C | 2.61832000 | 2.05392000 | -0.85456000 |
| C | 0.78943000 | -0.43839000 | 1.66381000 |
| C | -0.62163000 | -1.02101000 | 1.34584000 |
| C | -0.81849000 | -0.84695000 | -0.20962000 |
| C | -1.69280000 | -0.23324000 | 2.11491000 |
| C | -1.78122000 | 1.22967000 | 1.67747000 |
| C | -2.16342000 | 1.43615000 | 0.18631000 |
| C | -1.93678000 | 0.13592000 | -0.66382000 |
| C | -3.68253000 | 1.69492000 | 0.04737000 |
| C | -4.32307000 | 0.31662000 | -0.00006000 |
| C | -3.37033000 | -0.51940000 | -0.87982000 |
| C | -3.51285000 | -2.00457000 | -0.53944000 |
| C | -3.75120000 | -0.34165000 | -2.35962000 |
| C | -1.43791000 | 2.66499000 | -0.38105000 |
| C | -0.65021000 | -2.49207000 | 1.77467000 |
| H | -1.62514000 | 0.47210000 | -1.66171000 |
| C | 4.73380000 | -0.93716000 | 0.48770000 |
| C | 2.85233000 | 1.72968000 | 1.64817000 |
| H | -1.08976000 | -1.81654000 | -0.63194000 |
| H | 0.50341000 | 0.14057000 | -1.65427000 |
| H | 1.43360000 | -1.79053000 | -2.27630000 |
| H | 0.72932000 | -2.65902000 | -0.94992000 |
| H | 2.66923000 | -2.45808000 | 0.43026000 |
| H | 3.15844000 | -3.00292000 | -1.14699000 |
| H | 2.86858000 | -0.19064000 | -2.36458000 |
| H | 0.58498000 | 1.30181000 | 0.42269000 |
| H | 4.76291000 | 1.59456000 | -0.69052000 |
| H | 4.20070000 | 1.93638000 | -2.33039000 |
| H | 1.84959000 | 1.97013000 | -1.62770000 |
| H | 2.75909000 | 3.10891000 | -0.59449000 |

| | | | |
|---|---|---|---|
| H | 0.77915000 | 0.19582000 | 2.55552000 |
| H | 1.51180000 | -1.24377000 | 1.84508000 |
| H | -2.66109000 | -0.72447000 | 1.97701000 |
| H | -1.47301000 | -0.28911000 | 3.18832000 |
| H | -0.81660000 | 1.72102000 | 1.88972000 |
| H | -2.50102000 | 1.75836000 | 2.31172000 |
| H | -3.87545000 | 2.22126000 | -0.89525000 |
| H | -4.06156000 | 2.33381000 | 0.85119000 |
| H | -4.39616000 | -0.11023000 | 1.00718000 |
| H | -5.33756000 | 0.33166000 | -0.41113000 |
| H | -2.92451000 | -2.64916000 | -1.20130000 |
| H | -4.56170000 | -2.29457000 | -0.65888000 |
| H | -3.23519000 | -2.22286000 | 0.49523000 |
| H | -3.75504000 | 0.71387000 | -2.65240000 |
| H | -3.03864000 | -0.86513000 | -3.00750000 |
| H | -4.74731000 | -0.75041000 | -2.55615000 |
| H | -1.69004000 | 3.55822000 | 0.20158000 |
| H | -0.34164000 | 2.58549000 | -0.37627000 |
| H | -1.73353000 | 2.84499000 | -1.41937000 |
| H | -0.44677000 | -2.57976000 | 2.84770000 |
| H | -1.62220000 | -2.95148000 | 1.58159000 |
| H | 0.10845000 | -3.07635000 | 1.24208000 |
| H | 5.40955000 | -0.08102000 | 0.44717000 |
| H | 5.35061000 | -1.84211000 | 0.44888000 |
| H | 4.23533000 | -0.95330000 | 1.46510000 |
| H | 3.78442000 | 2.26450000 | 1.45500000 |
| H | 2.16965000 | 2.44516000 | 2.13122000 |
| H | 3.01349000 | 0.90494000 | 2.34278000 |

**IM10**
Sum of electronic and thermal Free Energies
= -976.540187

| | | | |
|---|---|---|---|
| C | 0.64456300 | -0.73242800 | -0.87750700 |
| C | 1.42307800 | -2.06220600 | -1.15105100 |
| C | 2.77385200 | -2.13904000 | -0.40399300 |
| C | 3.38576800 | -0.76709800 | -0.24944100 |
| C | 1.44480900 | 1.71521300 | -0.14251700 |
| C | 1.49180500 | 0.18579100 | 0.00802400 |
| C | 3.17973600 | 0.10910600 | -1.29035400 |
| C | 3.50883200 | 1.54418500 | -1.45660200 |
| C | 2.09953200 | 2.17834800 | -1.44796900 |
| C | 1.23949900 | -0.32377900 | 1.44198700 |
| C | -0.17698700 | -0.99514700 | 1.42244600 |
| C | -0.66189200 | -0.89850800 | -0.05980600 |
| C | -1.13392300 | -0.31600800 | 2.40622500 |
| C | -1.53334600 | 1.07144200 | 1.95045800 |
| C | -2.35503600 | 1.06359100 | 0.65178500 |
| C | -1.72407400 | 0.17802300 | -0.48373600 |
| C | -3.76689300 | 0.43730900 | 0.89580300 |
| C | -3.84079000 | -0.82190500 | 0.02254000 |
| C | -2.95485800 | -0.49636700 | -1.19035600 |
| C | -2.63665100 | -1.73046100 | -2.04849500 |
| C | -3.67573200 | 0.49891500 | -2.11879800 |
| C | -2.47121200 | 2.53640100 | 0.22128500 |
| C | -0.05279500 | -2.46565700 | 1.85257900 |
| H | -1.24009300 | 0.84870400 | -1.20862800 |
| C | 4.37102300 | -0.52738900 | 0.85282600 |
| C | 2.05055000 | 2.48271100 | 1.04613400 |
| H | -1.08198000 | -1.87361600 | -0.30052000 |
| H | 0.42446700 | -0.25529600 | -1.83773700 |
| H | 1.59935900 | -2.14423200 | -2.22714300 |
| H | 0.80986400 | -2.92752300 | -0.89276500 |
| H | 2.67119000 | -2.59910200 | 0.58331500 |
| H | 3.47752000 | -2.77200600 | -0.95815700 |
| H | 2.65296100 | -0.29290600 | -2.15345900 |
| H | 0.36924300 | 1.93955700 | -0.19087100 |
| H | 4.14010200 | 1.95030600 | -0.66426200 |
| H | 4.00344000 | 1.70405700 | -2.41910000 |
| H | 1.53624200 | 1.82695500 | -2.31970300 |
| H | 2.14380700 | 3.26805400 | -1.50558500 |
| H | 1.30068900 | 0.49471000 | 2.15843900 |
| H | 1.96384000 | -1.06758900 | 1.77256700 |
| H | -2.03068000 | -0.94148600 | 2.50937500 |
| H | -0.66607100 | -0.28750300 | 3.39810200 |
| H | -0.63327300 | 1.68780000 | 1.81363100 |
| H | -2.12094800 | 1.57511200 | 2.72661900 |
| H | -4.54520400 | 1.15046000 | 0.60540400 |
| H | -3.93495600 | 0.21508300 | 1.95445900 |
| H | -3.41524000 | -1.68320100 | 0.55907300 |
| H | -4.86269800 | -1.09239300 | -0.26328700 |
| H | -1.83673400 | -1.52204100 | -2.77079600 |
| H | -3.52339000 | -2.01234400 | -2.62294200 |
| H | -2.35546000 | -2.61196100 | -1.46638600 |
| H | -4.12225200 | 1.33864000 | -1.58720600 |
| H | -2.99087400 | 0.89851800 | -2.87563200 |
| H | -4.48929900 | -0.01091400 | -2.64314100 |
| H | -3.00916400 | 3.10496200 | 0.98761800 |
| H | -1.48091300 | 2.99647600 | 0.11420000 |
| H | -2.99669400 | 2.67071400 | -0.72438000 |
| H | 0.30265300 | -2.52860900 | 2.88707300 |
| H | -1.02909700 | -2.95803600 | 1.80464800 |
| H | 0.63742100 | -3.03785200 | 1.22740100 |
| H | 4.72014900 | 0.50434800 | 0.89787400 |
| H | 5.23810800 | -1.16852100 | 0.65481100 |
| H | 3.97724400 | -0.82136900 | 1.82845800 |
| H | 2.23667400 | 3.51862600 | 0.75293100 |
| H | 1.38543900 | 2.51180400 | 1.91040100 |
| H | 3.00145900 | 2.05337200 | 1.37649900 |

**TS_10−10a**
Sum of electronic and thermal Free Energies
= -976.523603

| | | | |
|---|---|---|---|
| C | -0.64369000 | 0.61405000 | -0.95351000 |
| C | -1.44516000 | -0.22665000 | 0.02604000 |
| C | -0.96800000 | 0.20140000 | 1.42098000 |
| C | 0.35959000 | 1.01895000 | 1.25867000 |
| C | 0.72787000 | 0.83427000 | -0.24231000 |
| C | 1.46928000 | 0.56658000 | 2.23883000 |
| C | 2.30069000 | -0.70074000 | 1.94986000 |
| C | 2.22310000 | -1.28017000 | 0.51781000 |
| C | 1.83243000 | -0.21395000 | -0.55514000 |
| C | 3.61551000 | -1.71264000 | 0.01933000 |
| C | 4.30418000 | -0.42256000 | -0.39295000 |
| C | 3.19628000 | 0.45671000 | -1.02331000 |
| C | 0.15098000 | 2.50698000 | 1.60393000 |
| H | 1.09803000 | 1.78752000 | -0.62085000 |
| C | 1.35255000 | -2.54077000 | 0.56786000 |
| H | 1.48537000 | -0.79547000 | -1.42311000 |
| C | 3.42001000 | 1.90864000 | -0.57696000 |
| C | 3.26411000 | 0.40886000 | -2.55501000 |
| H | -0.51172000 | 0.08770000 | -1.90579000 |
| C | -1.37473000 | 1.98128000 | -1.23142000 |
| C | -2.61687000 | 2.21914000 | -0.35086000 |
| C | -3.37974000 | 0.93790000 | -0.13611000 |
| C | -3.46780000 | 0.10159000 | -1.21637000 |
| C | -3.93028000 | -1.29176000 | -1.31169000 |
| C | -2.57529000 | -2.03956000 | -1.41412000 |
| C | -4.24441000 | 0.81302000 | 1.08441000 |
| C | -1.75324000 | -1.69901000 | -0.17142000 |
| C | -2.35711000 | -2.37315000 | 1.07224000 |
| H | -0.80272000 | -0.69938000 | 2.01846000 |
| H | -1.71460000 | 0.78025000 | 1.97473000 |
| H | 2.02282000 | -1.49592000 | 2.65016000 |
| H | 3.34160000 | -0.46051000 | 2.18942000 |
| H | 3.49128000 | -2.37514000 | -0.84838000 |
| H | 4.16114000 | -2.27716000 | 0.78348000 |
| H | 4.71599000 | 0.07858000 | 0.49131000 |
| H | 5.13818000 | -0.58137000 | -1.08352000 |
| H | -0.60396000 | 3.00832000 | 1.00414000 |
| H | -0.12957000 | 2.61421000 | 2.65784000 |
| H | 1.08721000 | 3.05248000 | 1.45382000 |
| H | 1.90424000 | -3.32607000 | 1.09574000 |
| H | 0.42826000 | -2.39250000 | 1.13168000 |
| H | 1.12248000 | -2.92358000 | -0.43372000 |
| H | 2.80991000 | 2.63097000 | -1.12961000 |
| H | 4.46600000 | 2.17331000 | -0.76169000 |
| H | 3.23559000 | 2.04291000 | 0.49508000 |
| H | 3.26655000 | -0.62462000 | -2.91889000 |
| H | 2.40218000 | 0.92259000 | -3.00021000 |
| H | 4.17093000 | 0.89996000 | -2.92230000 |
| H | -1.66964000 | 1.99892000 | -2.28389000 |
| H | -0.67447000 | 2.81067000 | -1.11547000 |
| H | -2.36162000 | 2.63030000 | 0.62811000 |
| H | -3.27192000 | 2.96046000 | -0.82463000 |
| H | -2.98923000 | 0.44351000 | -2.13321000 |
| H | -4.50780000 | -1.45090000 | -2.22674000 |
| H | -4.51297000 | -1.63375000 | -0.45462000 |
| H | -2.72956000 | -3.11952000 | -1.48021000 |
| H | -2.05762000 | -1.72856000 | -2.32796000 |
| H | -4.77865000 | -0.13615000 | 1.13674000 |
| H | -4.98992000 | 1.61555000 | 1.04480000 |
| H | -3.68067000 | 0.95970000 | 2.00959000 |
| H | -0.75470000 | -2.11807000 | -0.35165000 |
| H | -3.15553000 | -1.77063000 | 1.51538000 |
| H | -1.61749000 | -2.56110000 | 1.85253000 |
| H | -2.78123000 | -3.34166000 | 0.79708000 |
| H | 2.16597000 | 1.40749000 | 2.30633000 |
| H | 1.02051000 | 0.49606000 | 3.23733000 |

**IM10a**
Sum of electronic and thermal Free Energies
= -976.548019

| | | | |
|---|---|---|---|
| C | 0.61052500 | 1.00899000 | -0.25975300 |
| C | 1.55828300 | -0.13053000 | 0.13155300 |
| C | 1.42439900 | -1.10356200 | -1.04752000 |
| C | -0.10095800 | -1.08161700 | -1.37891900 |
| C | -0.64826000 | 0.26462400 | -0.78377300 |
| C | -0.83841900 | -2.29482600 | -0.80006000 |
| C | -2.33996300 | -2.03047700 | -0.77749300 |
| C | -2.72613500 | -1.00316500 | 0.30147200 |
| C | -1.73077200 | 0.21749400 | 0.33257200 |
| C | -4.11803500 | -0.39561400 | -0.02692500 |
| C | -3.82503500 | 1.00856400 | -0.55384700 |
| C | -2.64707900 | 1.49455600 | 0.30756500 |
| C | -0.23820200 | -1.08184800 | -2.90990100 |
| H | -1.07215200 | 0.82679600 | -1.62029200 |
| C | -2.75486700 | -1.71797500 | 1.66106600 |
| H | -1.20837100 | 0.19928500 | 1.29923400 |
| C | -2.03716500 | 2.78482500 | -0.25606800 |
| C | -3.12142400 | 1.81649600 | 1.73452000 |
| H | 0.34656000 | 1.60617200 | 0.61774800 |
| C | 1.31315900 | 1.92753200 | -1.31061600 |
| C | 2.76680200 | 1.49799100 | -1.65175900 |
| C | 3.38407600 | 0.68591300 | -0.54066200 |
| C | 3.07801000 | 1.09215600 | 0.74006800 |
| C | 3.43526700 | 0.53596500 | 2.07139200 |
| C | 2.05525100 | 0.10102800 | 2.60403900 |
| C | 4.42853000 | -0.33160300 | -0.87290400 |
| C | 1.47331200 | -0.80987800 | 1.51689600 |
| C | 2.13262300 | -2.19682600 | 1.57901200 |
| H | 1.79966200 | -2.10377800 | -0.83040300 |
| H | 1.95102400 | -0.75147300 | -1.93330300 |
| H | -2.89459800 | -2.95732700 | -0.59335600 |

| | | | |
|---|---|---|---|
| H | -2.66469400 | -1.67277200 | -1.76261300 |
| H | -4.73724100 | -0.34033300 | 0.87431700 |
| H | -4.66543800 | -1.01425700 | -0.74485800 |
| H | -3.50979000 | 0.96153600 | -1.60684800 |
| H | -4.68552200 | 1.68392600 | -0.50360800 |
| H | 0.24046700 | -0.19708100 | -3.34566200 |
| H | 0.23597600 | -1.97048700 | -3.34044900 |
| H | -1.28556100 | -1.07433900 | -3.22219200 |
| H | -3.54107900 | -2.48004200 | 1.67058300 |
| H | -1.80550900 | -2.22086000 | 1.88189200 |
| H | -2.95438300 | -1.02141200 | 2.48009000 |
| H | -1.20651500 | 3.15226300 | 0.35806100 |
| H | -2.80255900 | 3.56730800 | -0.25162900 |
| H | -1.69033200 | 2.69041200 | -1.28924600 |
| H | -3.61104500 | 0.97125400 | 2.22132600 |
| H | -2.27288100 | 2.11553400 | 2.36121200 |
| H | -3.83473800 | 2.64691600 | 1.71985200 |
| H | 1.31139600 | 2.95350400 | -0.93279200 |
| H | 0.73277900 | 1.95513300 | -2.23580600 |
| H | 2.81689600 | 0.93637500 | -2.58847600 |
| H | 3.39315500 | 2.38594200 | -1.80100100 |
| H | 2.49749400 | 2.01026500 | 0.81548600 |
| H | 3.87496400 | 1.32141600 | 2.69274200 |
| H | 4.13352000 | -0.30175200 | 2.02382000 |
| H | 2.12693000 | -0.41372100 | 3.56447900 |
| H | 1.42615200 | 0.98640800 | 2.75194600 |
| H | 4.77096500 | -0.89411800 | -0.00437600 |
| H | 5.28689500 | 0.21145800 | -1.28607400 |
| H | 4.08628100 | -1.02245500 | -1.64705200 |
| H | 0.39665400 | -0.93876500 | 1.69410300 |
| H | 3.14419900 | -2.19840700 | 1.16086200 |
| H | 1.55548100 | -2.95581300 | 1.04886900 |
| H | 2.20337700 | -2.51482800 | 2.62190300 |
| H | -0.59829200 | -3.18158100 | -1.39876400 |
| H | -0.49367700 | -2.50558100 | 0.22054600 |

**TS_10a−11**
Sum of electronic and thermal Free Energies
= -976.550250

| | | | |
|---|---|---|---|
| C | -0.60185000 | 0.96768000 | -0.34719000 |
| C | -1.58002000 | -0.21862000 | -0.23383000 |
| C | -1.44297000 | -0.64063000 | 1.25273000 |
| C | 0.03979000 | -0.34869000 | 1.64654000 |
| C | 0.62195000 | 0.56843000 | 0.51759000 |
| C | 0.86793000 | -1.62766000 | 1.82075000 |
| C | 2.35645000 | -1.31330000 | 1.72886000 |
| C | 2.77661000 | -0.97439000 | 0.28822000 |
| C | 1.76581000 | 0.02848000 | -0.38696000 |
| C | 4.14539000 | -0.24165000 | 0.30535000 |
| C | 3.80966000 | 1.22797000 | 0.06425000 |
| C | 2.66136000 | 1.18955000 | -0.95795000 |
| C | 0.02904000 | 0.42175000 | 2.97750000 |
| H | 1.01275000 | 1.46195000 | 1.01492000 |
| C | 2.87296000 | -2.27695000 | -0.51922000 |
| H | 1.30292000 | -0.48879000 | -1.23676000 |
| C | 2.01768000 | 2.57383000 | -1.10851000 |
| C | 3.18874000 | 0.79028000 | -2.34534000 |
| H | -0.30251000 | 1.07808000 | -1.39588000 |
| C | -1.29335000 | 2.30214000 | 0.08616000 |
| C | -2.64209000 | 2.12612000 | 0.81921000 |
| C | -3.42485000 | 0.96114000 | 0.32590000 |
| C | -3.05862000 | 0.46493000 | -0.92931000 |
| C | -3.66841000 | -0.63471000 | -1.75224000 |
| C | -2.40564000 | -1.18639000 | -2.42456000 |
| C | -4.48249000 | 0.35274000 | 1.16400000 |
| C | -1.42848000 | -1.41542000 | -1.25515000 |
| C | -1.67752000 | -2.81353000 | -0.67811000 |
| H | -1.72317000 | -1.68103000 | 1.42285000 |
| H | -2.07471000 | -0.05515000 | 1.92180000 |
| H | 0.61509000 | -2.09541000 | 2.78006000 |
| H | 0.61495000 | -2.35613000 | 1.04308000 |
| H | 2.95441000 | -2.16284000 | 2.07762000 |
| H | 2.60235000 | -0.47359000 | 2.39215000 |
| H | 4.79482000 | -0.61695000 | -0.49251000 |
| H | 4.67702000 | -0.40885000 | 1.24758000 |
| H | 3.45216000 | 1.69420000 | 0.99463000 |
| H | 4.66099000 | 1.82044000 | -0.28748000 |
| H | -0.48022000 | 1.38737000 | 2.86914000 |
| H | -0.48860000 | -0.15158000 | 3.75438000 |
| H | 1.04237000 | 0.62960000 | 3.33185000 |
| H | 3.64219000 | -2.93131000 | -0.09548000 |
| H | 1.92745000 | -2.83211000 | -0.52397000 |
| H | 3.13913000 | -2.08087000 | -1.56183000 |
| H | 1.20874000 | 2.58082000 | -1.84818000 |
| H | 2.77702000 | 3.27731000 | -1.46500000 |
| H | 1.63379000 | 2.97663000 | -0.16663000 |
| H | 3.68746000 | -0.18090000 | -2.34533000 |
| H | 2.36438000 | 0.74017000 | -3.06644000 |
| H | 3.90750000 | 1.53163000 | -2.70971000 |
| H | -1.45037000 | 2.92086000 | -0.80241000 |
| H | -0.63100000 | 2.88279000 | 0.73244000 |
| H | -2.53218000 | 2.05244000 | 1.90638000 |
| H | -3.27099000 | 3.01521000 | 0.64939000 |
| H | -2.60425000 | 1.21783000 | -1.57475000 |
| H | -4.39792000 | -0.23778000 | -2.46303000 |
| H | -4.16109000 | -1.39795000 | -1.14309000 |
| H | -2.58317000 | -2.10476000 | -2.98754000 |
| H | -2.01555000 | -0.44058000 | -3.12656000 |
| H | -4.61822000 | -0.71041000 | 0.95485000 |
| H | -5.42191000 | 0.85357000 | 0.88426000 |
| H | -4.31939000 | 0.51886000 | 2.23016000 |
| H | -0.39547000 | -1.37616000 | -1.61629000 |

| | | | |
|---|---|---|---|
| H | -1.58068000 | -3.54269000 | -1.48626000 |
| H | -2.68501000 | -2.91637000 | -0.25993000 |
| H | -0.96360000 | -3.09617000 | 0.09533000 |

**IM11**

Sum of electronic and thermal Free Energies
= -976.560973

| | | | |
|---|---|---|---|
| C | 0.61086200 | -1.00531200 | 0.27704400 |
| C | 1.55584500 | 0.12884300 | -0.13239000 |
| C | 1.42366700 | 1.11832200 | 1.03296200 |
| C | -0.10086700 | 1.09935600 | 1.36816300 |
| C | -0.64827200 | -0.25391600 | 0.79001900 |
| C | -0.84056100 | 2.30537600 | 0.77817100 |
| C | -2.34151100 | 2.03829700 | 0.75361900 |
| C | -2.72339200 | 0.99993500 | -0.31601600 |
| C | -1.72989800 | -0.22157900 | -0.32779400 |
| C | -4.11727100 | 0.39785500 | 0.01176700 |
| C | -3.82924500 | -0.99905700 | 0.55967500 |
| C | -2.64684000 | -1.49840300 | -0.28737700 |
| C | -0.23623300 | 1.11585400 | 2.89903500 |
| H | -1.07362700 | -0.80446600 | 1.63368000 |
| C | -2.74415600 | 1.69845800 | -1.68385400 |
| H | -1.20571200 | -0.21755400 | -1.29366900 |
| C | -2.03966300 | -2.77979200 | 0.29838300 |
| C | -3.11320700 | -1.84237800 | -1.71172900 |
| H | 0.34610800 | -1.61874600 | -0.58911600 |
| C | 1.31641200 | -1.90497600 | 1.34293700 |
| C | 2.77207900 | -1.47091800 | 1.67136800 |
| C | 3.38391500 | -0.67470500 | 0.54548400 |
| C | 3.07778100 | -1.10551500 | -0.72707800 |
| C | 3.42868800 | -0.56845000 | -2.06783100 |
| C | 2.04643800 | -0.14184700 | -2.60125100 |
| C | 4.42728900 | 0.35058800 | 0.85838300 |
| C | 1.46975000 | 0.78673900 | -1.52678400 |
| C | 2.13058100 | 2.17172300 | -1.61030300 |
| H | 1.79797700 | 2.11561700 | 0.80022100 |
| H | 1.95336400 | 0.77877500 | 1.92212300 |
| H | -0.60395300 | 3.19751300 | 1.37044800 |
| H | -0.49383400 | 2.50994000 | -0.24314300 |
| H | -2.89628900 | 2.96291100 | 0.55818800 |
| H | -2.66974600 | 1.69021800 | 1.74119200 |
| H | -4.73023600 | 0.33058100 | -0.89301400 |
| H | -4.66951500 | 1.02679700 | 0.71717400 |
| H | -3.52063600 | -0.93892200 | 1.61403600 |
| H | -4.69050400 | -1.67394400 | 0.51332000 |
| H | 0.24413000 | 0.23656100 | 3.34422600 |
| H | 0.23604400 | 2.01030300 | 3.31959200 |
| H | -1.28374300 | 1.10951900 | 3.21141200 |
| H | -3.52972000 | 2.46112500 | -1.70666600 |
| H | -1.79315000 | 2.19807200 | -1.90586700 |
| H | -2.94034700 | 0.99185500 | -2.49529800 |
| H | -1.21238200 | -3.15976900 | -0.31264600 |
| H | -2.80728800 | -3.56019500 | 0.31102800 |
| H | -1.68737600 | -2.66749300 | 1.32790300 |
| H | -3.60324800 | -1.00631700 | -2.21377500 |
| H | -2.26056900 | -2.14804200 | -2.32976300 |
| H | -3.82436100 | -2.67458700 | -1.68794900 |
| H | 1.31271000 | -2.93773800 | 0.98385100 |
| H | 0.73850000 | -1.91469200 | 2.27015800 |
| H | 2.82687000 | -0.89675000 | 2.60024000 |
| H | 3.39890900 | -2.35717500 | 1.82965000 |
| H | 2.49641200 | -2.02483800 | -0.78376000 |
| H | 3.86695800 | -1.36174200 | -2.68035800 |
| H | 4.12523800 | 0.27143200 | -2.03407500 |
| H | 2.11418700 | 0.35689800 | -3.57054600 |
| H | 1.41639100 | -1.02959200 | -2.73093800 |
| H | 4.77400700 | 0.89212500 | -0.02194600 |
| H | 5.28361700 | -0.18166900 | 1.28956400 |
| H | 4.08036300 | 1.06066100 | 1.61307700 |
| H | 0.39334100 | 0.91460000 | -1.70554900 |
| H | 2.20246000 | 2.47395000 | -2.65791800 |
| H | 3.14138100 | 2.17814100 | -1.19058500 |
| H | 1.55412600 | 2.93967200 | -1.09211900 |

**TS_11−12**

Sum of electronic and thermal Free Energies
= -976.559103

| | | | |
|---|---|---|---|
| C | 0.63570000 | -0.75042000 | -0.53294000 |
| C | 1.64842000 | 0.41885000 | -0.33429000 |
| C | 1.35288000 | 0.98209000 | 1.08791000 |
| C | -0.04302000 | 0.47892000 | 1.54816000 |
| C | -0.56782000 | -0.48913000 | 0.42986000 |
| C | -1.00138000 | 1.65162000 | 1.80593000 |
| C | -2.44789000 | 1.17944000 | 1.78965000 |
| C | -2.90484000 | 0.82060000 | 0.36752000 |
| C | -1.83967000 | -0.07589000 | -0.37905000 |
| C | -4.20009000 | -0.03194000 | 0.44370000 |
| C | -3.74520000 | -1.46127000 | 0.17152000 |
| C | -2.66230000 | -1.30893000 | -0.90878000 |
| C | 0.11306000 | -0.29674000 | 2.86814000 |
| H | -0.81617000 | -1.43022000 | 0.93523000 |
| C | -3.17532000 | 2.12076000 | -0.40527000 |
| H | -1.49001000 | 0.48587000 | -1.25146000 |
| C | -1.91621000 | -2.63126000 | -1.10822000 |
| C | -3.29574000 | -0.94510000 | -2.26223000 |
| H | 0.26970000 | -0.70866000 | -1.56575000 |
| C | 1.30209000 | -2.13845000 | -0.36726000 |
| C | 2.49566000 | -2.18981000 | 0.59903000 |
| C | 3.55051000 | -1.20989000 | 0.22747000 |
| C | 3.11871000 | -0.00215000 | -0.35422000 |
| C | 4.00310000 | 1.20053000 | -0.66861000 |
| C | 3.00672000 | 2.25346000 | -1.19563000 |

| | | | |
|---|---|---|---|
| C | 4.98387000 | -1.52693000 | 0.45205000 |
| C | 1.68320000 | 1.50750000 | -1.44733000 |
| C | 0.51029000 | 2.46978000 | -1.56262000 |
| H | 1.40417000 | 2.07603000 | 1.09384000 |
| H | 2.12030000 | 0.64556000 | 1.79874000 |
| H | -0.74454000 | 2.11758000 | 2.76490000 |
| H | -0.86826000 | 2.42519000 | 1.04190000 |
| H | -3.11368000 | 1.95393000 | 2.18717000 |
| H | -2.56593000 | 0.30790000 | 2.44730000 |
| H | -4.91875000 | 0.28789000 | -0.31807000 |
| H | -4.69720000 | 0.08261000 | 1.41223000 |
| H | -3.29328000 | -1.89457000 | 1.07639000 |
| H | -4.55454000 | -2.13045000 | -0.13952000 |
| H | 0.72439000 | -1.19754000 | 2.74448000 |
| H | 0.58538000 | 0.32968000 | 3.63206000 |
| H | -0.85526000 | -0.62662000 | 3.25415000 |
| H | -4.00699000 | 2.66407000 | 0.05596000 |
| H | -2.30801000 | 2.78895000 | -0.41737000 |
| H | -3.44631000 | 1.91922000 | -1.44587000 |
| H | -1.08950000 | -2.54430000 | -1.82278000 |
| H | -2.61261000 | -3.37145000 | -1.51524000 |
| H | -1.53615000 | -3.04420000 | -0.16922000 |
| H | -3.85579000 | -0.00868000 | -2.22944000 |
| H | -2.51944000 | -0.83968000 | -3.02922000 |
| H | -3.98414000 | -1.73212000 | -2.58734000 |
| H | 1.66277000 | -2.49487000 | -1.34201000 |
| H | 0.56652000 | -2.87148000 | -0.03581000 |
| H | 2.17823000 | -1.89209000 | 1.61116000 |
| H | 2.91142000 | -3.19753000 | 0.67621000 |
| H | 3.27674000 | -0.82791000 | -1.21896000 |
| H | 4.77580000 | 0.95424000 | -1.40176000 |
| H | 4.51339000 | 1.52801000 | 0.24282000 |
| H | 2.85148000 | 3.03404000 | -0.44386000 |
| H | 3.36712000 | 2.74729000 | -2.10027000 |
| H | 5.12532000 | -1.49815000 | 1.54248000 |
| H | 5.66428000 | -0.80409000 | 0.00298000 |
| H | 5.22530000 | -2.54315000 | 0.13170000 |
| H | 1.77276000 | 0.96111000 | -2.40016000 |
| H | 0.72182000 | 3.20667000 | -2.34289000 |
| H | 0.35106000 | 3.02010000 | -0.63036000 |
| H | -0.41994000 | 1.96795000 | -1.83130000 |

**IM12**
Sum of electronic and thermal Free Energies
= -976.570899

| | | | |
|---|---|---|---|
| C | 0.18660100 | -0.40891900 | 1.37696200 |
| C | 0.61018400 | 0.64581000 | 0.30586300 |
| C | 0.55689300 | -1.85313200 | 0.97251600 |
| C | 0.86011400 | -1.98084000 | -0.51415200 |
| C | 2.07353900 | -1.12815000 | -0.94052400 |
| C | 1.95978700 | 0.36016900 | -0.43290900 |
| C | 3.37706100 | -1.67714600 | -0.28046500 |
| C | 3.79708300 | -0.67248700 | 0.80032000 |
| C | 3.31768600 | 0.68835600 | 0.26612700 |
| H | 0.71605600 | 1.58999700 | 0.85640400 |
| C | 2.17879800 | -1.21094600 | -2.46965800 |
| H | 1.92095500 | 1.02049800 | -1.30894400 |
| C | 3.26300600 | 1.79743200 | 1.31784200 |
| C | 4.30737700 | 1.15933600 | -0.81881500 |
| C | 0.76139000 | -0.10923500 | 2.76943800 |
| C | -0.40705900 | 0.93891300 | -0.86177400 |
| C | -1.92104371 | 0.49749228 | -0.71052008 |
| C | -2.09806100 | -0.46660800 | 0.38628300 |
| C | -1.37885300 | -0.28844200 | 1.62727700 |
| C | -2.45859500 | -0.33378500 | -1.94117400 |
| C | -3.57728000 | -1.19055700 | -1.33618300 |
| C | -3.01121400 | -1.57420200 | 0.04552200 |
| C | -0.55049800 | 2.45749100 | -1.07010100 |
| C | -1.68901100 | 2.82696300 | -0.12653200 |
| C | -2.75571100 | 1.76895500 | -0.40516000 |
| C | -3.99425800 | -2.04808900 | 1.11602500 |
| C | -3.88449700 | 1.72140600 | 0.62484300 |
| H | -0.02095500 | 0.46563300 | -1.76735800 |
| H | 1.44443700 | -2.16119900 | 1.53645000 |
| H | -0.23340400 | -2.55221800 | 1.27887800 |
| H | 1.06261100 | -3.02813500 | -0.76603000 |
| H | -0.02007700 | -1.69480000 | -1.10973800 |
| H | 4.16116400 | -1.75852500 | -1.03997900 |
| H | 3.23473300 | -2.68695600 | 0.11959000 |
| H | 3.32001000 | -0.91007800 | 1.75645600 |
| H | 4.87678300 | -0.67926600 | 0.98208900 |
| H | 1.25893200 | -0.86784500 | -2.95836200 |
| H | 3.00267100 | -0.60202900 | -2.85101800 |
| H | 2.35431700 | -2.24582200 | -2.78374600 |
| H | 2.90277100 | 2.73402200 | 0.87457900 |
| H | 4.26846900 | 1.98937100 | 1.70537400 |
| H | 2.62455400 | 1.56284300 | 2.17023100 |
| H | 4.43670300 | 0.42418900 | -1.61690400 |
| H | 3.96552900 | 2.09417300 | -1.27609100 |
| H | 5.29155100 | 1.34152600 | -0.37612800 |
| H | 0.51222300 | 0.90502700 | 3.09946800 |
| H | 0.35706700 | -0.81821100 | 3.49973600 |
| H | 1.84405300 | -0.21891300 | 2.78360700 |
| H | -1.53003400 | 0.74441100 | 1.96975400 |
| H | -1.68986900 | -0.98131500 | 2.41166600 |
| H | -2.79740500 | 0.34134200 | -2.73031600 |
| H | -1.68446600 | -0.98059700 | -2.37132100 |
| H | -3.82343200 | -2.06765600 | -1.93685900 |
| H | -4.49843100 | -0.61176000 | -1.20505100 |
| H | -2.25496400 | -2.37920600 | -0.11647300 |
| H | -0.84205900 | 2.67720800 | -2.10359500 |
| H | 0.38318400 | 2.99085500 | -0.86980500 |

| | | | |
|---|---|---|---|
| H | -1.35951700 | 2.77577000 | 0.92128900 |
| H | -2.09205400 | 3.83200600 | -0.28514900 |
| H | -3.20565600 | 1.97502200 | -1.38303400 |
| H | -3.48736600 | -2.38238400 | 2.02375200 |
| H | -4.57244000 | -2.88784900 | 0.72586900 |
| H | -4.69265200 | -1.24916400 | 1.37849500 |
| H | -4.64388600 | 0.97255400 | 0.38413100 |
| H | -4.38418500 | 2.69393200 | 0.63100600 |
| H | -3.52772600 | 1.54315800 | 1.64363500 |

**TS_12−13**
Sum of electronic and thermal Free Energies
= -976.553018

| | | | |
|---|---|---|---|
| C | 0.16913000 | -0.32386000 | 1.50689000 |
| C | 0.43154000 | 0.51301000 | 0.21766000 |
| C | 0.81574000 | -1.72857000 | 1.44929000 |
| C | 1.14324000 | -2.20088000 | 0.04114000 |
| C | 2.15097000 | -1.28833000 | -0.67720000 |
| C | 1.75458000 | 0.22987000 | -0.57566000 |
| C | 3.55333000 | -1.38441000 | -0.00390000 |
| C | 3.77640000 | -0.06814000 | 0.74741000 |
| C | 3.03791000 | 0.98826000 | -0.09093000 |
| H | 0.45438000 | 1.56027000 | 0.53340000 |
| C | 2.22295000 | -1.74610000 | -2.14033000 |
| H | 1.56722000 | 0.58581000 | -1.59648000 |
| C | 2.79075000 | 2.29331000 | 0.66637000 |
| C | 3.88717000 | 1.33434000 | -1.32826000 |
| C | 0.66434000 | 0.38600000 | 2.77564000 |
| C | -0.68482000 | 0.43323000 | -0.84903000 |
| C | -2.59991000 | 0.54290000 | -0.42524000 |
| C | -2.06875000 | -0.50267000 | 0.33450000 |
| C | -1.38078000 | -0.37929000 | 1.65101000 |
| C | -3.35556000 | -0.04933000 | -1.61536000 |
| C | -3.63153000 | -1.49566000 | -1.19501000 |
| C | -2.44871000 | -1.85434000 | -0.28208000 |
| C | -0.82419000 | 1.77131000 | -1.59760000 |
| C | -1.54593000 | 2.76744000 | -0.69783000 |
| C | -2.78568000 | 2.03410000 | -0.13538000 |
| C | -2.76749000 | -2.94386000 | 0.73905000 |
| C | -3.11483000 | 2.44163000 | 1.30438000 |
| H | -0.59522000 | -0.41857000 | -1.52667000 |
| H | 1.74926000 | -1.70645000 | 2.02445000 |
| H | 0.17431000 | -2.45193000 | 1.96499000 |
| H | 1.55815000 | -3.21469000 | 0.07780000 |
| H | 0.23027000 | -2.27925000 | -0.55902000 |
| H | 4.32143000 | -1.51380000 | -0.77315000 |
| H | 3.62604000 | -2.25723000 | 0.65358000 |
| H | 3.34665000 | -0.12302000 | 1.75352000 |
| H | 4.83661000 | 0.17726000 | 0.86893000 |
| H | 1.24031000 | -1.68801000 | -2.62561000 |
| H | 2.91745000 | -1.13658000 | -2.72519000 |
| H | 2.56019000 | -2.78667000 | -2.19840000 |
| H | 2.22381000 | 3.00768000 | 0.05502000 |
| H | 3.74882000 | 2.76507000 | 0.90585000 |
| H | 2.25580000 | 2.14836000 | 1.60731000 |
| H | 4.14117000 | 0.45284000 | -1.92195000 |
| H | 3.35557000 | 2.03668000 | -1.98060000 |
| H | 4.82548000 | 1.80632000 | -1.02082000 |
| H | 0.26194000 | 1.40288000 | 2.85201000 |
| H | 0.33654000 | -0.17447000 | 3.65822000 |
| H | 1.75235000 | 0.44434000 | 2.81759000 |
| H | -1.71273000 | 0.53842000 | 2.13632000 |
| H | -1.66540000 | -1.21288000 | 2.29867000 |
| H | -4.26268000 | 0.52380000 | -1.81830000 |
| H | -2.74480000 | -0.00665000 | -2.52594000 |
| H | -3.73039000 | -2.18071000 | -2.03918000 |
| H | -4.56067000 | -1.54300000 | -0.61488000 |
| H | -1.61784000 | -2.20575000 | -0.90465000 |
| H | -1.36312000 | 1.63681000 | -2.53895000 |
| H | 0.18592000 | 2.10968000 | -1.85371000 |
| H | -0.90055000 | 3.07847000 | 0.13104000 |
| H | -1.82032000 | 3.67463000 | -1.24070000 |
| H | -3.64910000 | 2.31158000 | -0.74771000 |
| H | -1.88032000 | -3.26617000 | 1.28981000 |
| H | -3.16812000 | -3.81623000 | 0.21643000 |
| H | -3.52412000 | -2.60800000 | 1.45559000 |
| H | -3.87897000 | 1.80044000 | 1.75238000 |
| H | -3.50309000 | 3.46343000 | 1.29388000 |
| H | -2.23149000 | 2.44564000 | 1.94781000 |

**IM13**
Sum of electronic and thermal Free Energies
= -976.568013

| | | | |
|---|---|---|---|
| C | 0.08695400 | -0.13846200 | 1.35612000 |
| C | 0.48731100 | 0.47329700 | -0.01622700 |
| C | 0.83449300 | -1.46497100 | 1.63471700 |
| C | 1.40405400 | -2.10873500 | 0.37820700 |
| C | 2.44548700 | -1.24084800 | -0.34548000 |
| C | 1.91264700 | 0.21965200 | -0.57450600 |
| C | 3.73121000 | -1.07669500 | 0.51809600 |
| C | 3.73225300 | 0.36572300 | 1.03367300 |
| C | 3.05046100 | 1.17383500 | -0.08225100 |
| H | 0.39251900 | 1.55532300 | 0.09948500 |
| C | 2.76950100 | -1.93681000 | -1.67392300 |
| H | 1.81176200 | 0.38660800 | -1.65722900 |
| C | 2.59642200 | 2.56028000 | 0.37741100 |
| C | 4.04795800 | 1.38145700 | -1.23643100 |
| C | 0.30425200 | 0.81881800 | 2.53640400 |
| C | -0.58837700 | 0.04721900 | -1.05995000 |
| C | -2.94148200 | 0.40552500 | -0.14718500 |
| C | -1.82525000 | -0.49304200 | -0.31493500 |
| C | -1.45501500 | -0.36079900 | 1.25669500 |

| | | | |
|---|---|---|---|
| C | -4.18175400 | -0.33557500 | 0.23399900 |
| C | -3.81131800 | -1.82254800 | 0.12303300 |
| C | -2.50091400 | -1.84466400 | -0.68900600 |
| C | -0.96821000 | 1.24179200 | -1.95242800 |
| C | -1.66720200 | 2.34400800 | -1.15026100 |
| C | -2.95865500 | 1.84056500 | -0.50042200 |
| C | -1.73178200 | -3.14399900 | -0.50620100 |
| C | -3.50777400 | 2.76096600 | 0.60257400 |
| H | -0.20192500 | -0.76237500 | -1.69108500 |
| H | 1.65949600 | -1.25678900 | 2.32712100 |
| H | 0.17938900 | -2.16539800 | 2.16781900 |
| H | 1.86125000 | -3.07295600 | 0.63136800 |
| H | 0.59603600 | -2.33031200 | -0.32117600 |
| H | 4.61683100 | -1.25538100 | -0.10097700 |
| H | 3.77044900 | -1.81133700 | 1.32988300 |
| H | 3.15539500 | 0.44291600 | 1.96267000 |
| H | 4.73782100 | 0.73927800 | 1.25467200 |
| H | 1.86596700 | -2.06825900 | -2.28241000 |
| H | 3.48822200 | -1.36503700 | -2.26720300 |
| H | 3.19806700 | -2.92877200 | -1.49268700 |
| H | 2.06250300 | 3.08867200 | -0.42335500 |
| H | 3.46982400 | 3.16622200 | 0.63816100 |
| H | 1.94910800 | 2.52890400 | 1.25776200 |
| H | 4.44583300 | 0.43833500 | -1.61871000 |
| H | 3.57054900 | 1.90586800 | -2.07170600 |
| H | 4.89674100 | 1.98525500 | -0.89997700 |
| H | -0.18847800 | 1.78404300 | 2.36402600 |
| H | -0.10380600 | 0.38518000 | 3.45667000 |
| H | 1.36393200 | 1.00930100 | 2.71077200 |
| H | -1.94917800 | 0.50001300 | 1.73965900 |
| H | -1.82178700 | -1.24744400 | 1.78008400 |
| H | -4.55556000 | -0.02170800 | 1.21608200 |
| H | -4.95928900 | -0.03851500 | -0.48300900 |
| H | -4.59587400 | -2.40790900 | -0.35977200 |
| H | -3.64883400 | -2.26041800 | 1.11279100 |
| H | -2.75195800 | -1.74743900 | -1.75623600 |
| H | -1.62310000 | 0.91001000 | -2.76981500 |
| H | -0.06285000 | 1.64270500 | -2.42056600 |
| H | -1.00725900 | 2.75005200 | -0.37586400 |
| H | -1.92059400 | 3.18748100 | -1.79927300 |
| H | -3.73198200 | 1.80256900 | -1.29461900 |
| H | -0.92029800 | -3.25008400 | -1.22830200 |
| H | -2.41683500 | -3.98214100 | -0.66486200 |
| H | -1.31994700 | -3.23933500 | 0.50418500 |
| H | -4.48528300 | 2.43682500 | 0.96644400 |
| H | -3.60979000 | 3.77262500 | 0.20500700 |
| H | -2.80869700 | 2.79747800 | 1.44382800 |

**TS_10−10-1**
Sum of electronic and thermal Free Energies
= -976.531544

| | | | |
|---|---|---|---|
| C | 0.66347000 | -0.72315000 | -0.80313000 |
| C | 1.39870000 | -2.02476000 | -1.23245000 |
| C | 2.72549000 | -2.06173000 | -0.46324000 |
| C | 3.03009000 | -0.58506000 | -0.17517000 |
| C | 1.53842000 | 1.57410000 | -0.03252000 |
| C | 1.60904000 | 0.03962000 | 0.18673000 |
| C | 3.65668000 | 0.09218000 | -1.28352000 |
| C | 3.67225000 | 1.53347000 | -1.42877000 |
| C | 2.15432000 | 1.96846000 | -1.37595000 |
| C | 1.12151000 | -0.41140000 | 1.58440000 |
| C | -0.28410000 | -1.06834000 | 1.43722000 |
| C | -0.68053000 | -0.88900000 | -0.06098000 |
| C | -1.30077000 | -0.44619000 | 2.39784000 |
| C | -1.64369000 | 0.97560000 | 2.00186000 |
| C | -2.38470000 | 1.06198000 | 0.65868000 |
| C | -1.70488000 | 0.22565000 | -0.48501000 |
| C | -3.82284000 | 0.46171000 | 0.78256000 |
| C | -3.86889000 | -0.75063000 | -0.15706000 |
| C | -2.90400000 | -0.38553000 | -1.29572000 |
| C | -2.55799000 | -1.58194000 | -2.19523000 |
| C | -3.54752000 | 0.67065000 | -2.21338000 |
| C | -2.43507000 | 2.55851000 | 0.30464000 |
| C | -0.18208000 | -2.56597000 | 1.76989000 |
| H | -1.17566000 | 0.92573000 | -1.14681000 |
| C | 4.17480000 | -0.44906000 | 0.91408000 |
| C | 2.10090000 | 2.44547000 | 1.09952000 |
| H | -1.11460000 | -1.84099000 | -0.36396000 |
| H | 0.49808000 | -0.12722000 | -1.70692000 |
| H | 1.59792000 | -1.99208000 | -2.31003000 |
| H | 0.79777000 | -2.92005000 | -1.05879000 |
| H | 2.61735000 | -2.58712000 | 0.49062000 |
| H | 3.53797000 | -2.55332000 | -1.00887000 |
| H | 4.11246000 | -0.52161000 | -2.06867000 |
| H | 0.46819000 | 1.80969000 | -0.10895000 |
| H | 4.13746000 | 2.00468000 | -0.55236000 |
| H | 4.18259000 | 1.86410000 | -2.33217000 |
| H | 1.62141000 | 1.52917000 | -2.22265000 |
| H | 2.14710000 | 3.05209000 | -1.51985000 |
| H | 1.10474000 | 0.42556000 | 2.28549000 |
| H | 1.80192000 | -1.14719000 | 2.02551000 |
| H | -2.21220000 | -1.05916000 | 2.40211000 |
| H | -0.89979000 | -0.48563000 | 3.41840000 |
| H | -0.72215000 | 1.57526000 | 1.94875000 |
| H | -2.26462000 | 1.45213000 | 2.76940000 |
| H | -4.56587000 | 1.20767000 | 0.48154000 |
| H | -4.06118000 | 0.19081000 | 1.81611000 |
| H | -3.49361000 | -1.64660000 | 0.35936000 |
| H | -4.87684000 | -0.98293000 | -0.51709000 |
| H | -1.70737000 | -1.35665000 | -2.85179000 |
| H | -3.41127000 | -1.81290000 | -2.83928000 |

| | | | |
|---|---|---|---|
| H | -2.33211000 | -2.49710000 | -1.64201000 |
| H | -3.99646000 | 1.49981000 | -1.66726000 |
| H | -2.81201000 | 1.08207000 | -2.91456000 |
| H | -4.34651000 | 0.20986000 | -2.80199000 |
| H | -2.99356000 | 3.10473000 | 1.07273000 |
| H | -1.42466000 | 2.98595000 | 0.27233000 |
| H | -2.90669000 | 2.75975000 | -0.65774000 |
| H | 0.11971000 | -2.70398000 | 2.81415000 |
| H | -1.15024000 | -3.05811000 | 1.63268000 |
| H | 0.54699000 | -3.08650000 | 1.14071000 |
| H | 4.41993000 | 0.58117000 | 1.16436000 |
| H | 5.08027000 | -0.97356000 | 0.59846000 |
| H | 3.78649000 | -0.95269000 | 1.80187000 |
| H | 2.13863000 | 3.49178000 | 0.78205000 |
| H | 1.47095000 | 2.41014000 | 1.98973000 |
| H | 3.10780000 | 2.15532000 | 1.41359000 |

**IM10-1**
Sum of electronic and thermal Free Energies
= -976.565933

| | | | |
|---|---|---|---|
| C | 0.37609000 | -1.20554000 | -0.06096300 |
| C | 0.99931100 | -2.34304100 | 0.79415000 |
| C | 2.51645600 | -2.11215800 | 0.74560900 |
| C | 2.71109400 | -0.72012500 | 0.30946800 |
| C | 2.06994900 | -0.26625700 | -1.90079300 |
| C | 1.53631200 | -0.19682200 | -0.36317100 |
| C | 4.01447400 | -0.03248700 | 0.32784900 |
| C | 4.11299700 | 0.92242400 | -0.86960600 |
| C | 3.52612300 | 0.23207600 | -2.10375200 |
| C | 1.01914700 | 1.19997300 | 0.09537200 |
| C | 0.00661200 | 0.85672600 | 1.20143200 |
| C | -0.73746900 | -0.37344400 | 0.62969900 |
| C | -0.96173500 | 1.99218800 | 1.54582200 |
| C | -1.84044000 | 2.38658400 | 0.37546700 |
| C | -2.68666500 | 1.23695200 | -0.20297400 |
| C | -1.91727800 | -0.11632200 | -0.35848900 |
| C | -3.89355200 | 0.88262000 | 0.72472900 |
| C | -3.81077400 | -0.62714800 | 1.01882900 |
| C | -3.01461700 | -1.22121300 | -0.15840000 |
| C | -2.48196600 | -2.63143200 | 0.13683900 |
| C | -3.94036000 | -1.36424700 | -1.38316400 |
| C | -3.16205500 | 1.75805200 | -1.57311700 |
| C | 0.75901100 | 0.49177000 | 2.48845800 |
| H | -1.52169300 | -0.17209300 | -1.38119400 |
| C | 4.23212400 | 0.67412100 | 1.69462800 |
| C | 1.09939000 | 0.46124300 | -2.83095500 |
| H | -1.13262300 | -0.92173600 | 1.48974600 |
| H | 0.00132200 | -1.62682800 | -0.99896900 |
| H | 0.73106600 | -3.33181600 | 0.41829000 |
| H | 0.64869800 | -2.28831100 | 1.82842900 |
| H | 3.09608900 | -2.36222400 | 1.64218800 |
| H | 2.99300500 | -2.69798100 | -0.06579100 |
| H | 4.78680100 | -0.81166300 | 0.25384200 |
| H | 2.02285400 | -1.33420500 | -2.14332200 |
| H | 3.58377600 | 1.85675500 | -0.66054000 |
| H | 5.16251000 | 1.18354200 | -1.03056100 |
| H | 4.16898200 | -0.61750400 | -2.36252200 |
| H | 3.55751700 | 0.91310700 | -2.95829900 |
| H | 0.51609800 | 1.67295800 | -0.75010700 |
| H | 1.80403000 | 1.87666500 | 0.44168500 |
| H | -1.58726500 | 1.65187700 | 2.38368500 |
| H | -0.39974800 | 2.86006600 | 1.91303500 |
| H | -1.21787900 | 2.81542900 | -0.42064500 |
| H | -2.52081200 | 3.19131300 | 0.67759100 |
| H | -4.83301700 | 1.13211400 | 0.22005600 |
| H | -3.87721900 | 1.46941100 | 1.64897100 |
| H | -3.27455100 | -0.80313300 | 1.96098400 |
| H | -4.79382300 | -1.09833900 | 1.12256600 |
| H | -1.93721400 | -3.03837600 | -0.72447900 |
| H | -3.32610900 | -3.30088400 | 0.32745100 |
| H | -1.83246300 | -2.68796300 | 1.01420100 |
| H | -4.65953800 | -0.54912600 | -1.46854100 |
| H | -3.36965200 | -1.42067300 | -2.31703700 |
| H | -4.52309900 | -2.28599800 | -1.29370900 |
| H | -3.70148100 | 2.70222200 | -1.43945100 |
| H | -2.30223000 | 1.95931500 | -2.22361900 |
| H | -3.82228500 | 1.07306500 | -2.10173300 |
| H | 1.23885600 | 1.38033300 | 2.91302100 |
| H | 0.07258200 | 0.09039800 | 3.24065100 |
| H | 1.54481400 | -0.26268000 | 2.34412200 |
| H | 5.23295500 | 1.11023100 | 1.68182400 |
| H | 4.16950700 | -0.02727400 | 2.52967700 |
| H | 3.50516500 | 1.47363100 | 1.84604100 |
| H | 1.34909100 | 0.19015700 | -3.86073200 |
| H | 0.05820600 | 0.18257600 | -2.65288800 |
| H | 1.19570700 | 1.54669000 | -2.74173500 |

**TS_10-1−10-2**
Sum of electronic and thermal Free Energies
= -976.557432

| | | | |
|---|---|---|---|
| C | 0.35288000 | 1.06385000 | -0.35525000 |
| C | 1.05143000 | 2.00873000 | -1.35527000 |
| C | 2.51546000 | 2.04629000 | -0.85499000 |
| C | 2.72579000 | 0.68996000 | -0.17000000 |
| C | 2.65672000 | 0.76553000 | 1.64440000 |
| C | 1.47003000 | 0.12166000 | 0.03910000 |
| C | 4.05500000 | -0.03934000 | -0.29002000 |
| C | 4.18854000 | -1.04001000 | 0.87230000 |
| C | 3.35670000 | -0.51817000 | 2.06104000 |
| C | 0.96171000 | -1.26402000 | 0.22153000 |
| C | -0.11476000 | -1.27593000 | -0.91093000 |

| | | | |
|---|---|---|---|
| C | -0.80752000 | 0.10613000 | -0.74650000 |
| C | -1.10840000 | -2.43104000 | -0.82495000 |
| C | -1.95209000 | -2.38967000 | 0.43172000 |
| C | -2.75752000 | -1.09001000 | 0.61707000 |
| C | -1.95147000 | 0.21456000 | 0.31369000 |
| C | -3.97934000 | -1.01129000 | -0.35402000 |
| C | -3.87871000 | 0.33392000 | -1.10062000 |
| C | -3.02828000 | 1.23913000 | -0.18825000 |
| C | -2.46647000 | 2.46656000 | -0.92073000 |
| C | -3.90686000 | 1.78818000 | 0.95379000 |
| C | -3.21023000 | -1.12739000 | 2.09030000 |
| C | 0.59583000 | -1.36122000 | -2.27072000 |
| H | -1.50534000 | 0.56685000 | 1.25612000 |
| C | 4.22465000 | -0.69204000 | -1.66308000 |
| C | 1.51358000 | 1.22946000 | 2.52931000 |
| H | -1.20684000 | 0.37177000 | -1.72914000 |
| H | 0.04495000 | 1.66135000 | 0.50951000 |
| H | 0.59056000 | 2.99767000 | -1.38096000 |
| H | 1.00560000 | 1.59750000 | -2.36844000 |
| H | 3.23981000 | 2.17589000 | -1.66371000 |
| H | 2.67749000 | 2.87066000 | -0.15044000 |
| H | 4.82273000 | 0.73876000 | -0.18020000 |
| H | 3.37369000 | 1.59003000 | 1.57039000 |
| H | 3.83028000 | -2.02451000 | 0.55819000 |
| H | 5.24098000 | -1.16779000 | 1.13437000 |
| H | 3.98278000 | -0.28347000 | 2.92794000 |
| H | 2.62348000 | -1.25653000 | 2.40041000 |
| H | 0.46369000 | -1.35113000 | 1.19812000 |
| H | 1.71456000 | -2.04861000 | 0.12559000 |
| H | -1.75584000 | -2.36504000 | -1.71139000 |
| H | -0.57541000 | -3.38644000 | -0.90423000 |
| H | -1.30934000 | -2.54700000 | 1.30894000 |
| H | -2.65531000 | -3.23054000 | 0.43478000 |
| H | -4.91061000 | -1.07346000 | 0.21886000 |
| H | -3.99369000 | -1.85333000 | -1.05362000 |
| H | -3.37616000 | 0.19688000 | -2.06725000 |
| H | -4.85607000 | 0.77884000 | -1.31432000 |
| H | -1.87239000 | 3.09337000 | -0.24284000 |
| H | -3.29603000 | 3.08212000 | -1.28070000 |
| H | -1.85358000 | 2.22438000 | -1.79255000 |
| H | -4.65518000 | 1.07184000 | 1.29383000 |
| H | -3.30556000 | 2.09621000 | 1.81665000 |
| H | -4.45513000 | 2.66656000 | 0.60064000 |
| H | -3.76852000 | -2.05132000 | 2.27651000 |
| H | -2.34010000 | -1.12889000 | 2.75783000 |
| H | -3.84711000 | -0.29453000 | 2.38074000 |
| H | 1.11319000 | -2.32214000 | -2.36851000 |
| H | -0.13546000 | -1.29224000 | -3.08149000 |
| H | 1.33700000 | -0.57008000 | -2.42569000 |
| H | 5.22561000 | -1.12310000 | -1.74442000 |
| H | 4.10423000 | 0.02720000 | -2.47856000 |
| H | 3.49757000 | -1.49789000 | -1.80161000 |
| H | 1.93107000 | 1.35369000 | 3.53392000 |
| H | 1.11614000 | 2.19970000 | 2.22511000 |
| H | 0.69810000 | 0.50373000 | 2.59804000 |

**IM10-2**
Sum of electronic and thermal Free Energies
= -976.566627

| | | | |
|---|---|---|---|
| C | 0.34438200 | 0.87049000 | -0.62361400 |
| C | 1.11479100 | 1.63993000 | -1.72930700 |
| C | 2.46918800 | 1.91262500 | -1.03352000 |
| C | 2.71102800 | 0.70459600 | -0.08530400 |
| C | 3.03541300 | 1.09571800 | 1.43855100 |
| C | 1.41883100 | 0.04276800 | -0.05092100 |
| C | 3.96470500 | -0.16774800 | -0.39632400 |
| C | 4.26352100 | -0.96659200 | 0.88611600 |
| C | 3.60470900 | -0.19034400 | 2.04617200 |
| C | 0.92542700 | -1.28112700 | 0.32761100 |
| C | -0.23458000 | -1.48932300 | -0.70525000 |
| C | -0.87156900 | -0.07091100 | -0.78429900 |
| C | -1.24614900 | -2.56127300 | -0.31831500 |
| C | -2.00764300 | -2.21987300 | 0.94477900 |
| C | -2.77476400 | -0.88583300 | 0.88845800 |
| C | -1.95544800 | 0.29895000 | 0.28272500 |
| C | -4.04101900 | -0.96913200 | -0.02392600 |
| C | -3.94280400 | 0.18534800 | -1.04188300 |
| C | -3.02979600 | 1.23272700 | -0.37486700 |
| C | -2.46826100 | 2.26434800 | -1.36457800 |
| C | -3.83999000 | 2.03424200 | 0.66401100 |
| C | -3.15346500 | -0.59998500 | 2.35521100 |
| C | 0.38118500 | -1.86838300 | -2.05879400 |
| H | -1.45226500 | 0.82410500 | 1.10975500 |
| C | 3.89101600 | -1.00820600 | -1.66434100 |
| C | 1.92911600 | 1.75219900 | 2.25057000 |
| H | -1.30604000 | 0.01929600 | -1.78274400 |
| H | 0.13732400 | 1.64226900 | 0.14376900 |
| H | 0.58933400 | 2.54932400 | -2.02498500 |
| H | 1.24090900 | 1.01803700 | -2.62064800 |
| H | 3.28987700 | 2.02704000 | -1.74600100 |
| H | 2.41117800 | 2.84070400 | -0.45384800 |
| H | 4.75642400 | 0.58146700 | -0.52788500 |
| H | 3.84423800 | 1.82458900 | 1.29469700 |
| H | 3.86169000 | -1.98260400 | 0.81152200 |
| H | 5.34261300 | -1.07666800 | 1.01823500 |
| H | 4.30426400 | 0.04963300 | 2.85099400 |
| H | 2.79749800 | -0.77518700 | 2.50407800 |
| H | 0.47715300 | -1.14487500 | 1.33307400 |
| H | 1.67737500 | -2.07086100 | 0.39281800 |
| H | -1.94585300 | -2.66163000 | -1.16045900 |
| H | -0.74248300 | -3.53075300 | -0.22157700 |
| H | -1.31323000 | -2.20322200 | 1.79770700 |

| | | | |
|---|---|---|---|
| H | -2.72353600 | -3.01732100 | 1.17462300 |
| H | -4.94345400 | -0.87514300 | 0.58904200 |
| H | -4.11363300 | -1.93833300 | -0.52795200 |
| H | -3.48923800 | -0.16851100 | -1.97725700 |
| H | -4.91815300 | 0.60709000 | -1.30522700 |
| H | -1.81785400 | 2.98627900 | -0.85274000 |
| H | -3.29384900 | 2.83311800 | -1.80171400 |
| H | -1.91178200 | 1.82899300 | -2.19836800 |
| H | -4.58737300 | 1.42848300 | 1.17698700 |
| H | -3.19146500 | 2.49526000 | 1.41744000 |
| H | -4.38421700 | 2.83718600 | 0.15840700 |
| H | -3.73047400 | -1.44174700 | 2.75298400 |
| H | -2.25202900 | -0.49834000 | 2.97114500 |
| H | -3.74860200 | 0.30014000 | 2.49196100 |
| H | 0.90408100 | -2.82856100 | -1.98935200 |
| H | -0.40614200 | -1.96940600 | -2.81133200 |
| H | 1.09929200 | -1.12732600 | -2.42570100 |
| H | 4.86557600 | -1.45980700 | -1.86746300 |
| H | 3.61846100 | -0.40710800 | -2.53876200 |
| H | 3.16838800 | -1.82403400 | -1.56391800 |
| H | 2.33053200 | 2.06582600 | 3.21785100 |
| H | 1.52935000 | 2.64715000 | 1.76256700 |
| H | 1.09918600 | 1.06419700 | 2.45450900 |

**TS_7−7a**
Sum of electronic and thermal Free Energies
= -976.486917

| | | | |
|---|---|---|---|
| C | 1.95911000 | 0.26592000 | -0.27015000 |
| C | 3.01210000 | -0.89935000 | -0.40690000 |
| C | 2.51563000 | -2.22909000 | 0.16185000 |
| C | 1.17071000 | -2.62110000 | -0.41746000 |
| C | 0.02473000 | -1.68416000 | 0.00009000 |
| C | 0.35772000 | -0.17077000 | -0.18304000 |
| H | 3.26663000 | -2.99623000 | -0.05772000 |
| H | 2.17094000 | 0.85821000 | -1.18680000 |
| H | 1.23526000 | -2.66585000 | -1.51236000 |
| H | 0.89730000 | -3.63194000 | -0.09661000 |
| H | -0.09852000 | 0.44654000 | 0.58702000 |
| H | 2.44891000 | -2.17069000 | 1.25375000 |
| C | 2.41972000 | 1.24096000 | 0.87268000 |
| C | 4.21302000 | -0.35862000 | 0.42077000 |
| H | 4.27781000 | -0.89434000 | 1.37264000 |
| H | 5.15734000 | -0.52567000 | -0.10388000 |
| C | 3.94281000 | 1.12633000 | 0.66614000 |
| H | 4.49432000 | 1.52525000 | 1.52237000 |
| H | 4.23143000 | 1.71808000 | -0.21263000 |
| C | -1.20086000 | -2.11403000 | -0.85810000 |
| H | -1.41858000 | -3.14797000 | -0.56618000 |
| C | 0.38702000 | 0.45983000 | -1.45504000 |
| H | 0.64853000 | -0.20001000 | -2.28712000 |
| C | -2.43738000 | -1.26716000 | -0.78842000 |
| C | -2.07903000 | 2.18274000 | -0.19935000 |
| C | 0.06160000 | 1.88329000 | -1.84345000 |
| H | 0.92346000 | 2.48842000 | -1.56061000 |
| H | 0.06339000 | 1.89979000 | -2.93623000 |
| C | -3.26146000 | 1.61996000 | -0.48113000 |
| H | -3.53395000 | 1.53459000 | -1.53539000 |
| C | -1.23097000 | 2.64911000 | -1.36645000 |
| H | -0.90357000 | 3.67940000 | -1.18560000 |
| H | -1.88474000 | 2.68941000 | -2.24030000 |
| C | -4.26065000 | 1.00303000 | 0.44981000 |
| H | -3.87648000 | 0.94483000 | 1.47306000 |
| H | -5.17165000 | 1.61395000 | 0.47480000 |
| C | -3.48706000 | -1.39609000 | 0.03752000 |
| H | -2.48589000 | -0.47441000 | -1.53248000 |
| C | -4.64145000 | -0.41294000 | -0.03967000 |
| H | -5.47824000 | -0.77971000 | 0.56196000 |
| H | -5.00533000 | -0.34190000 | -1.07149000 |
| C | 3.43305000 | -1.05877000 | -1.87533000 |
| H | 2.60019000 | -1.31692000 | -2.53775000 |
| H | 3.88830000 | -0.13520000 | -2.25113000 |
| H | 4.17690000 | -1.85516000 | -1.96602000 |
| C | 2.03375000 | 0.73486000 | 2.26607000 |
| H | 2.51681000 | 1.36469000 | 3.01885000 |
| H | 0.95517000 | 0.78834000 | 2.44226000 |
| H | 2.35682000 | -0.29429000 | 2.44017000 |
| C | 1.96120000 | 2.68951000 | 0.71147000 |
| H | 2.34386000 | 3.27868000 | 1.55014000 |
| H | 2.35754000 | 3.13979000 | -0.20538000 |
| H | 0.87274000 | 2.79004000 | 0.71712000 |
| C | -0.30852000 | -1.87508000 | 1.48580000 |
| H | 0.52055000 | -1.58641000 | 2.13546000 |
| H | -1.18139000 | -1.27308000 | 1.76221000 |
| H | -0.54165000 | -2.92457000 | 1.68720000 |
| C | -1.65849000 | 2.52512000 | 1.21146000 |
| H | -2.51211000 | 2.51638000 | 1.88996000 |
| H | -0.90734000 | 1.85019000 | 1.63797000 |
| H | -1.23233000 | 3.53385000 | 1.24261000 |
| C | -3.63024000 | -2.46896000 | 1.08270000 |
| H | -2.81080000 | -3.18833000 | 1.08315000 |
| H | -3.69230000 | -2.02308000 | 2.08355000 |
| H | -4.56566000 | -3.01637000 | 0.92527000 |
| H | -0.88012000 | -2.16545000 | -1.90711000 |

**IM7a**
Sum of electronic and thermal Free Energies
= -976.498582

| | | | |
|---|---|---|---|
| C | 1.98253900 | 0.33865400 | -0.17764500 |
| C | 3.04274800 | -0.74175600 | -0.62175800 |
| C | 2.63629900 | -2.16338000 | -0.25107900 |
| C | 1.24493900 | -2.50384500 | -0.75047500 |
| C | 0.12548900 | -1.69186200 | -0.07545300 |

| | | | |
|---|---|---|---|
| C | 0.40129900 | -0.15705600 | -0.05311100 |
| H | 3.37576600 | -2.85841500 | -0.67748300 |
| H | 2.10262200 | 1.12014700 | -0.96631800 |
| H | 1.19335900 | -2.35653500 | -1.84494200 |
| H | 1.02934200 | -3.56596400 | -0.59225600 |
| H | -0.01003000 | 0.33479100 | 0.82547400 |
| H | 2.68158100 | -2.30204500 | 0.83219300 |
| C | 2.52355000 | 1.09580300 | 1.09308600 |
| C | 4.31226900 | -0.29685200 | 0.16624200 |
| H | 4.49997700 | -1.01053300 | 0.98281500 |
| H | 5.19704400 | -0.31040200 | -0.46530900 |
| C | 4.01706900 | 1.08782600 | 0.72617600 |
| H | 4.64447100 | 1.33736000 | 1.59672000 |
| H | 4.20310000 | 1.86469000 | -0.02997300 |
| C | -1.17308600 | -2.03923900 | -0.84819100 |
| H | -1.37986000 | -3.09210400 | -0.64761700 |
| C | 0.35273200 | 0.65340800 | -1.22138500 |
| H | 0.53996900 | 0.13751500 | -2.17433700 |
| C | -2.37168700 | -1.16533700 | -0.58269000 |
| C | -2.38699700 | 2.24339800 | -0.43819900 |
| C | 0.01718700 | 2.11390900 | -1.32836200 |
| H | 0.27939800 | 2.60770300 | -0.38776900 |
| H | 0.62879900 | 2.54647300 | -2.12402700 |
| C | -3.53388100 | 1.57697800 | -0.60928700 |
| H | -3.81439800 | 1.29535000 | -1.63051800 |
| C | -1.46818800 | 2.47289100 | -1.62453900 |
| H | -1.43093300 | 3.53801200 | -1.88280600 |
| H | -1.83246900 | 1.93929900 | -2.50390200 |
| C | -4.41111500 | 1.01254100 | 0.46466600 |
| H | -3.95044600 | 1.16876100 | 1.44868100 |
| H | -5.38903400 | 1.49766700 | 0.47339600 |
| C | -3.44442800 | -1.43698400 | 0.20729600 |
| H | -2.37289500 | -0.23924700 | -1.17302100 |
| C | -4.65223600 | -0.50449500 | 0.25400000 |
| H | -5.31928900 | -0.84116400 | 1.06577500 |
| H | -5.22730700 | -0.64272200 | -0.67140100 |
| C | 3.32415100 | -0.63040200 | -2.12865600 |
| H | 2.44396200 | -0.82776500 | -2.74575500 |
| H | 3.69570400 | 0.37347400 | -2.38051900 |
| H | 4.09833400 | -1.34370000 | -2.41420100 |
| C | 2.28753800 | 0.30585800 | 2.38510400 |
| H | 2.81213800 | 0.80071700 | 3.20171400 |
| H | 1.22865600 | 0.26112300 | 2.64766600 |
| H | 2.66240600 | -0.72454200 | 2.32743200 |
| C | 2.02387900 | 2.52116100 | 1.29244200 |
| H | 2.58077700 | 2.96622900 | 2.12689100 |
| H | 2.19543800 | 3.14459400 | 0.41688400 |
| H | 0.96497600 | 2.56253600 | 1.57057300 |
| C | -0.06105300 | -2.14717200 | 1.38320300 |
| H | 0.81668700 | -1.94192500 | 2.00790600 |
| H | -0.92848400 | -1.63818700 | 1.83239300 |
| H | -0.24532900 | -3.21976500 | 1.42020600 |
| C | -1.92914000 | 2.76627300 | 0.89641000 |
| H | -2.76980000 | 3.00360700 | 1.54426600 |
| H | -1.29289000 | 2.04986200 | 1.44112700 |
| H | -1.34643800 | 3.68645100 | 0.78079600 |
| C | -3.61616200 | -2.73189200 | 1.01806400 |
| H | -2.78180900 | -3.42852500 | 0.91305200 |
| H | -3.72227400 | -2.49354400 | 2.09916600 |
| H | -4.54143500 | -3.25512600 | 0.70199400 |
| H | -0.96166000 | -1.97471600 | -1.92010900 |

**TS_7a−7b**
Sum of electronic and thermal Free Energies
= -976.499166

| | | | |
|---|---|---|---|
| C | 1.99254000 | 0.33865000 | -0.17765000 |
| C | 3.05275000 | -0.73176000 | -0.63176000 |
| C | 2.64630000 | -2.16338000 | -0.27108000 |
| C | 1.24494000 | -2.49385000 | -0.75048000 |
| C | 0.14549000 | -1.69186000 | -0.03545000 |
| C | 0.42130000 | -0.15706000 | -0.03311000 |
| H | 3.37577000 | -2.84842000 | -0.71748000 |
| H | 2.10262000 | 1.12015000 | -0.96632000 |
| H | 1.17336000 | -2.33654000 | -1.83494000 |
| H | 1.02934000 | -3.55596000 | -0.59226000 |
| H | 0.00997000 | 0.34479000 | 0.84547000 |
| H | 2.71158000 | -2.31205000 | 0.81219000 |
| C | 2.54355000 | 1.09580000 | 1.09309000 |
| C | 4.32227000 | -0.29685000 | 0.15624000 |
| H | 4.50998000 | -1.00053000 | 0.97282000 |
| H | 5.20704000 | -0.31040000 | -0.48531000 |
| C | 4.03707000 | 1.09783000 | 0.71618000 |
| H | 4.66447000 | 1.34736000 | 1.57672000 |
| H | 4.21310000 | 1.86469000 | -0.04997000 |
| C | -1.18309000 | -2.01924000 | -0.76819000 |
| H | -1.37986000 | -3.08210000 | -0.59762000 |
| C | 0.36273000 | 0.64341000 | -1.21139000 |
| H | 0.53997000 | 0.11752000 | -2.15434000 |
| C | -2.35169000 | -1.15534000 | -0.41269000 |
| C | -2.39700000 | 2.26340000 | -0.47820000 |
| C | 0.01719000 | 2.10391000 | -1.32836000 |
| H | 0.26940000 | 2.60770000 | -0.39777000 |
| H | 0.63880000 | 2.52647000 | -2.12403000 |
| C | -3.52388000 | 1.55698000 | -0.61929000 |
| H | -3.79440000 | 1.21535000 | -1.62052000 |
| C | -1.46819000 | 2.45289000 | -1.65454000 |
| H | -1.43093000 | 3.50801000 | -1.95281000 |
| H | -1.82247000 | 1.88930000 | -2.52390000 |
| C | -4.37112000 | 1.01254000 | 0.49467000 |
| H | -3.88045000 | 1.17876000 | 1.45868000 |
| H | -5.33903000 | 1.52767000 | 0.53340000 |
| C | -3.49443000 | -1.46698000 | 0.21730000 |

| | | | |
|---|---|---|---|
| H | -2.25290000 | -0.14925000 | -0.79302000 |
| C | -4.66224000 | -0.49450000 | 0.33400000 |
| H | -5.27929000 | -0.80116000 | 1.18578000 |
| H | -5.29731000 | -0.64272000 | -0.55140000 |
| C | 3.32415000 | -0.61040000 | -2.13866000 |
| H | 2.44396000 | -0.81777000 | -2.75576000 |
| H | 3.68570000 | 0.39347000 | -2.39052000 |
| H | 4.09833000 | -1.32370000 | -2.43420000 |
| C | 2.31754000 | 0.30586000 | 2.38510000 |
| H | 2.84214000 | 0.81072000 | 3.20171000 |
| H | 1.25866000 | 0.26112000 | 2.65767000 |
| H | 2.70241000 | -0.71454000 | 2.32743000 |
| C | 2.03388000 | 2.52116000 | 1.29244000 |
| H | 2.59078000 | 2.97623000 | 2.11689000 |
| H | 2.19544000 | 3.14459000 | 0.40688000 |
| H | 0.97498000 | 2.55254000 | 1.57057000 |
| C | -0.00105000 | -2.14717000 | 1.42320000 |
| H | 0.89669000 | -1.97193000 | 2.01791000 |
| H | -0.83848000 | -1.62819000 | 1.90239000 |
| H | -0.21533000 | -3.21977000 | 1.45021000 |
| C | -1.94914000 | 2.83627000 | 0.84641000 |
| H | -2.79980000 | 3.10361000 | 1.47427000 |
| H | -1.33289000 | 2.12986000 | 1.42113000 |
| H | -1.35644000 | 3.74645000 | 0.70080000 |
| C | -3.79616000 | -2.83189000 | 0.77806000 |
| H | -3.00181000 | -3.55853000 | 0.60305000 |
| H | -3.96227000 | -2.76354000 | 1.85917000 |
| H | -4.72144000 | -3.22513000 | 0.34199000 |
| H | -1.01166000 | -1.91472000 | -1.85011000 |

**IM7b**
Sum of electronic and thermal Free Energies
= -976.506263

| | | | |
|---|---|---|---|
| C | 1.98253900 | 0.33865400 | -0.17764500 |
| C | 3.04274800 | -0.74175600 | -0.62175800 |
| C | 2.63629900 | -2.16338000 | -0.25107900 |
| C | 1.24493900 | -2.50384500 | -0.75047500 |
| C | 0.12548900 | -1.69186200 | -0.07545300 |
| C | 0.40129900 | -0.15705600 | -0.05311100 |
| H | 3.37576600 | -2.85841500 | -0.67748300 |
| H | 2.10262200 | 1.12014700 | -0.96631800 |
| H | 1.19335900 | -2.35653500 | -1.84494200 |
| H | 1.02934200 | -3.56596400 | -0.59225600 |
| H | -0.01003000 | 0.33479100 | 0.82547400 |
| H | 2.68158100 | -2.30204500 | 0.83219300 |
| C | 2.52355000 | 1.09580300 | 1.09308600 |
| C | 4.31226900 | -0.29685200 | 0.16624200 |
| H | 4.49997700 | -1.01053300 | 0.98281500 |
| H | 5.19704400 | -0.31040200 | -0.46530900 |
| C | 4.01706900 | 1.08782600 | 0.72617600 |
| H | 4.64447100 | 1.33736000 | 1.59672000 |
| H | 4.20310000 | 1.86469000 | -0.02997300 |
| C | -1.17308600 | -2.03923900 | -0.84819100 |
| H | -1.37986000 | -3.09210400 | -0.64761700 |
| C | 0.35273200 | 0.65340800 | -1.22138500 |
| H | 0.53996900 | 0.13751500 | -2.17433700 |
| C | -2.37168700 | -1.16533700 | -0.58269000 |
| C | -2.38699700 | 2.24339800 | -0.43819900 |
| C | 0.01718700 | 2.11390900 | -1.32836200 |
| H | 0.27939800 | 2.60770300 | -0.38776900 |
| H | 0.62879900 | 2.54647300 | -2.12402700 |
| C | -3.53388100 | 1.57697800 | -0.60928700 |
| H | -3.81439800 | 1.29535000 | -1.63051800 |
| C | -1.46818800 | 2.47289100 | -1.62453900 |
| H | -1.43093300 | 3.53801200 | -1.88280600 |
| H | -1.83246900 | 1.93929900 | -2.50390200 |
| C | -4.41111500 | 1.01254100 | 0.46466600 |
| H | -3.95044600 | 1.16876100 | 1.44868100 |
| H | -5.38903400 | 1.49766700 | 0.47339600 |
| C | -3.44442800 | -1.43698400 | 0.20729600 |
| H | -2.37289500 | -0.23924700 | -1.17302100 |
| C | -4.65223600 | -0.50449500 | 0.25400000 |
| H | -5.31928900 | -0.84116400 | 1.06577500 |
| H | -5.22730700 | -0.64272200 | -0.67140100 |
| C | 3.32415100 | -0.63040200 | -2.12865600 |
| H | 2.44396200 | -0.82776500 | -2.74575500 |
| H | 3.69570400 | 0.37347400 | -2.38051900 |
| H | 4.09833400 | -1.34370000 | -2.41420100 |
| C | 2.28753800 | 0.30585800 | 2.38510400 |
| H | 2.81213800 | 0.80071700 | 3.20171400 |
| H | 1.22865600 | 0.26112300 | 2.64766600 |
| H | 2.66240600 | -0.72454200 | 2.32743200 |
| C | 2.02387900 | 2.52116100 | 1.29244200 |
| H | 2.58077700 | 2.96622900 | 2.12689100 |
| H | 2.19543800 | 3.14459400 | 0.41688400 |
| H | 0.96497600 | 2.56253600 | 1.57057300 |
| C | -0.06105300 | -2.14717200 | 1.38320300 |
| H | 0.81668700 | -1.94192500 | 2.00790600 |
| H | -0.92848400 | -1.63818700 | 1.83239300 |
| H | -0.24532900 | -3.21976500 | 1.42020600 |
| C | -1.92914000 | 2.76627300 | 0.89641000 |
| H | -2.76980000 | 3.00360700 | 1.54426600 |
| H | -1.29289000 | 2.04986200 | 1.44112700 |
| H | -1.34643800 | 3.68645100 | 0.78079600 |
| C | -3.61616200 | -2.73189200 | 1.01806400 |
| H | -2.78180900 | -3.42852500 | 0.91305200 |
| H | -3.72227400 | -2.49354400 | 2.09916600 |
| H | -4.54143500 | -3.25512600 | 0.70199400 |
| H | -0.96166000 | -1.97471600 | -1.92010900 |

**TS_7b−7i-1**
Sum of electronic and thermal Free Energies

= -976.474776

| | | | |
|---|---|---|---|
| C | 1.76458000 | -0.06436000 | -0.39659000 |
| C | 2.76103000 | -1.07328000 | 0.33973000 |
| C | 2.32046000 | -1.27532000 | 1.78617000 |
| C | 0.89735000 | -1.83351000 | 1.88978000 |
| C | -0.14379000 | -1.21965000 | 0.89836000 |
| C | 0.41468000 | 0.16053000 | 0.40471000 |
| H | 3.01777000 | -1.94930000 | 2.29759000 |
| H | 1.46841000 | -0.52994000 | -1.34874000 |
| H | 0.90758000 | -2.91915000 | 1.74136000 |
| H | 0.54567000 | -1.67917000 | 2.91570000 |
| H | 0.65140000 | 0.70358000 | 1.33087000 |
| H | 2.38585000 | -0.31591000 | 2.31668000 |
| C | 2.66106000 | 1.17211000 | -0.80192000 |
| C | 4.14118000 | -0.38187000 | 0.23588000 |
| H | 4.32319000 | 0.25432000 | 1.10991000 |
| H | 4.95718000 | -1.10659000 | 0.19110000 |
| C | 4.01459000 | 0.47010000 | -1.01711000 |
| H | 4.82623000 | 1.19047000 | -1.15728000 |
| H | 3.97695000 | -0.16564000 | -1.91150000 |
| C | -0.42358000 | -2.21969000 | -0.26896000 |
| H | -0.82533000 | -3.13487000 | 0.17880000 |
| C | -0.45217000 | 1.12272000 | -0.43075000 |
| H | -0.38508000 | 0.91916000 | -1.49629000 |
| C | -1.35555000 | -1.62813000 | -1.27178000 |
| C | -2.46875000 | 1.94215000 | 0.80928000 |
| C | -0.32456000 | 2.59152000 | -0.04288000 |
| H | 0.69215000 | 2.82755000 | 0.24981000 |
| H | -0.59220000 | 3.24526000 | -0.87902000 |
| C | -2.46831000 | 1.32731000 | -0.45042000 |
| H | -2.15683000 | 2.01189000 | -1.24123000 |
| C | -1.28378000 | 2.77625000 | 1.12459000 |
| H | -0.84145000 | 2.36976000 | 2.04671000 |
| H | -1.57003000 | 3.80845000 | 1.36165000 |
| C | -3.67326000 | 0.50342000 | -0.88950000 |
| H | -4.13945000 | 0.02999000 | -0.02012000 |
| H | -4.40000000 | 1.23944000 | -1.26131000 |
| C | -2.69095000 | -1.68940000 | -1.27698000 |
| H | -0.89576000 | -0.95056000 | -1.99062000 |
| C | -3.44132000 | -0.57951000 | -1.96300000 |
| H | -4.41483000 | -0.90292000 | -2.34466000 |
| H | -2.87185000 | -0.16878000 | -2.80418000 |
| C | 2.91043000 | -2.44065000 | -0.36133000 |
| H | 2.09310000 | -3.13379000 | -0.16854000 |
| H | 3.01580000 | -2.33058000 | -1.44610000 |
| H | 3.81572000 | -2.92968000 | 0.00971000 |
| C | 2.88415000 | 2.22985000 | 0.31795000 |
| H | 3.95089000 | 2.35440000 | 0.52075000 |
| H | 2.52242000 | 3.21823000 | 0.02119000 |
| H | 2.41938000 | 1.96585000 | 1.27420000 |
| C | 2.22595000 | 1.85060000 | -2.10602000 |
| H | 3.00398000 | 2.54798000 | -2.43416000 |
| H | 2.08683000 | 1.11203000 | -2.90319000 |
| H | 1.30384000 | 2.43230000 | -2.01392000 |
| C | -1.43648000 | -1.01897000 | 1.69784000 |
| H | -1.30283000 | -0.24110000 | 2.46164000 |
| H | -2.28279000 | -0.76032000 | 1.06108000 |
| H | -1.69275000 | -1.94618000 | 2.22161000 |
| C | -3.53673000 | 1.76684000 | 1.82327000 |
| H | -3.31620000 | 2.30413000 | 2.74594000 |
| H | -4.49270000 | 2.11395000 | 1.41136000 |
| H | -3.67358000 | 0.70360000 | 2.05328000 |
| C | -3.52111000 | -2.65121000 | -0.47077000 |
| H | -4.14348000 | -3.24557000 | -1.14882000 |
| H | -2.91668000 | -3.34071000 | 0.11924000 |
| H | -4.20761000 | -2.13676000 | 0.21285000 |
| H | 0.50632000 | -2.49191000 | -0.76060000 |

**IM7i-1**
Sum of electronic and thermal Free Energies
= -976.480010

| | | | |
|---|---|---|---|
| C | -0.61313100 | -1.21044300 | 0.31729400 |
| C | -2.24626100 | -1.17220200 | 0.39634200 |
| C | -0.22707500 | 2.28713100 | -0.02984300 |
| C | -0.13056400 | 1.13986600 | -1.09011100 |
| C | 0.89259000 | 1.54588500 | -2.20065500 |
| C | 2.24612600 | 0.82701900 | -2.12681700 |
| C | 2.77470200 | 0.69232500 | -0.69770600 |
| C | 3.94864000 | -0.29444800 | -0.60233000 |
| C | 3.96343600 | -0.66377500 | 0.87381000 |
| C | 2.47995400 | -0.97610400 | 1.19309400 |
| C | 1.69735700 | 0.07725700 | 0.29305500 |
| C | 0.35357600 | -0.22501400 | -0.47527300 |
| C | -0.95608500 | 1.75675400 | 1.15555600 |
| C | -2.25896800 | 1.85931100 | 1.42477300 |
| C | -2.87871500 | 0.71467700 | 2.18214300 |
| C | -3.31467200 | -0.29417000 | 1.10438000 |
| C | -2.68009800 | -1.79471800 | -0.83584400 |
| C | -1.60156900 | -2.60037000 | -1.41863000 |
| C | -0.50725600 | -2.61140100 | -0.34968500 |
| C | -4.01994100 | -1.68880100 | -1.42228600 |
| C | -1.50034400 | 1.01467500 | -1.77010100 |
| C | 3.26730700 | 2.07280000 | -0.21595300 |
| H | -0.28240500 | -1.24909400 | 1.35644500 |
| H | 0.62189900 | -0.81001500 | -1.36422900 |
| H | 1.44724800 | 0.86732200 | 1.00366300 |
| C | 2.19031300 | -0.72978300 | 2.67958600 |
| C | 2.27945200 | -2.47350200 | 0.91620600 |
| C | -3.20312500 | 2.87063000 | 0.83549900 |
| H | -2.32622000 | -2.15350300 | 0.94756300 |
| H | -0.74516800 | 3.13083100 | -0.49640800 |
| H | 0.45583000 | 1.34962200 | -3.18681200 |

| | | | |
|---|---|---|---|
| H | 1.03645800 | 2.63152400 | -2.15299100 |
| H | 2.16381400 | -0.16857100 | -2.58046700 |
| H | 2.97977100 | 1.36761700 | -2.73629600 |
| H | 4.88742700 | 0.13870000 | -0.96040400 |
| H | 3.74584300 | -1.18522700 | -1.21221200 |
| H | 4.61274100 | -1.51196100 | 1.10948600 |
| H | 4.31603400 | 0.18471900 | 1.47072700 |
| H | -2.16894600 | 0.26050000 | 2.88158800 |
| H | -3.76723400 | 1.00935600 | 2.75019500 |
| H | -4.04238500 | -0.99419900 | 1.53412100 |
| H | -3.86230800 | 0.26837200 | 0.33995900 |
| H | -1.26706200 | -2.03086500 | -2.30482800 |
| H | -1.95120100 | -3.56592900 | -1.79868600 |
| H | -0.71409400 | -3.41330400 | 0.37031600 |
| H | 0.46728000 | -2.80252900 | -0.78582600 |
| H | -4.79352400 | -1.82254600 | -0.65757300 |
| H | -4.13334600 | -0.64727200 | -1.76549300 |
| H | -4.17464700 | -2.36790300 | -2.25942400 |
| H | -2.31680500 | 0.91092700 | -1.04969300 |
| H | -1.70544700 | 1.91797800 | -2.35423600 |
| H | -1.51747000 | 0.17544500 | -2.47779300 |
| H | 3.43551900 | 2.09873100 | 0.86488400 |
| H | 4.21343600 | 2.32017100 | -0.70806700 |
| H | 2.56727700 | 2.87534900 | -0.45824500 |
| H | 2.39066500 | 0.31185500 | 2.95218100 |
| H | 1.14692400 | -0.94873900 | 2.94101700 |
| H | 2.82286200 | -1.36812300 | 3.30566000 |
| H | 1.30817600 | -2.84126200 | 1.25171300 |
| H | 2.41922800 | -2.73045400 | -0.13872300 |
| H | 3.02963200 | -3.03231100 | 1.48378300 |
| H | -4.02909800 | 2.40325100 | 0.28455200 |
| H | -3.65972800 | 3.45382700 | 1.64283500 |
| H | -2.70301500 | 3.56889800 | 0.16338600 |
| H | 0.76695900 | 2.64296500 | 0.23932900 |
| H | -0.40880700 | 1.01426200 | 1.73597400 |

**TS_7i-1−7i-2**
Sum of electronic and thermal Free Energies
= -976.466966

| | | | |
|---|---|---|---|
| C | -0.70147000 | -1.17022000 | 0.00760000 |
| C | -2.30661000 | -0.97946000 | 0.41288000 |
| C | 0.03241000 | 2.35572000 | -0.17831000 |
| C | 0.11726000 | 1.17644000 | -1.19749000 |
| C | 1.25954000 | 1.47397000 | -2.21863000 |
| C | 2.56340000 | 0.69460000 | -1.96703000 |
| C | 2.92976000 | 0.55614000 | -0.48371000 |
| C | 4.04958000 | -0.48261000 | -0.25947000 |
| C | 3.80785000 | -0.94987000 | 1.16948000 |
| C | 2.29046000 | -1.22857000 | 1.19900000 |
| C | 1.71618000 | -0.00090000 | 0.39452000 |
| C | 0.45251000 | -0.18193000 | -0.50683000 |
| C | -0.76971000 | 1.91526000 | 0.99089000 |
| C | -2.04530000 | 2.17294000 | 1.27179000 |
| C | -2.74671000 | 1.11582000 | 2.07893000 |
| C | -3.25250000 | 0.08816000 | 1.05848000 |
| C | -3.05455000 | -1.80763000 | -0.52062000 |
| C | -2.22355000 | -2.80895000 | -1.18969000 |
| C | -0.79307000 | -2.31061000 | -1.06785000 |
| C | -4.49452000 | -1.73552000 | -0.79226000 |
| C | -1.21338000 | 1.08875000 | -1.95607000 |
| C | 3.45356000 | 1.91205000 | 0.03376000 |
| H | -0.29306000 | -1.58827000 | 0.92515000 |
| H | 0.83120000 | -0.78729000 | -1.33506000 |
| H | 1.51165000 | 0.75900000 | 1.15341000 |
| C | 1.78333000 | -1.25614000 | 2.64479000 |
| C | 2.06249000 | -2.61132000 | 0.55556000 |
| C | -2.89487000 | 3.25190000 | 0.66315000 |
| H | -2.32751000 | -1.80276000 | 1.18293000 |
| H | -0.41036000 | 3.21716000 | -0.68894000 |
| H | 0.91300000 | 1.23830000 | -3.23192000 |
| H | 1.45445000 | 2.55276000 | -2.21394000 |
| H | 2.48520000 | -0.30459000 | -2.41422000 |
| H | 3.38843000 | 1.18568000 | -2.49633000 |
| H | 5.04376000 | -0.05633000 | -0.41998000 |
| H | 3.94354000 | -1.32524000 | -0.95405000 |
| H | 4.39143000 | -1.82945000 | 1.45752000 |
| H | 4.05827000 | -0.14827000 | 1.87515000 |
| H | -2.07609000 | 0.65175000 | 2.80967000 |
| H | -3.61617000 | 1.50173000 | 2.62198000 |
| H | -4.04629000 | -0.49750000 | 1.53584000 |
| H | -3.73232000 | 0.64552000 | 0.24494000 |
| H | -2.56981000 | -2.99283000 | -2.21324000 |
| H | -2.42884000 | -3.75394000 | -0.65366000 |
| H | -0.09459000 | -3.11698000 | -0.84949000 |
| H | -0.50450000 | -1.90423000 | -2.03947000 |
| H | -5.10206000 | -1.50740000 | 0.08439000 |
| H | -4.60011000 | -0.86021000 | -1.46055000 |
| H | -4.86333000 | -2.61424000 | -1.32107000 |
| H | -2.06664000 | 1.04485000 | -1.26944000 |
| H | -1.34854000 | 1.97310000 | -2.58747000 |
| H | -1.23823000 | 0.21280000 | -2.61741000 |
| H | 3.52886000 | 1.93394000 | 1.12560000 |
| H | 4.45266000 | 2.09612000 | -0.37328000 |
| H | 2.83510000 | 2.75685000 | -0.27406000 |
| H | 2.01278000 | -0.31500000 | 3.15541000 |
| H | 0.69884000 | -1.41161000 | 2.70973000 |
| H | 2.26199000 | -2.06745000 | 3.20412000 |
| H | 1.10401000 | -3.05134000 | 0.84557000 |
| H | 2.11578000 | -2.58657000 | -0.53782000 |
| H | 2.83340000 | -3.30595000 | 0.90253000 |
| H | -3.76883000 | 2.84343000 | 0.13982000 |
| H | -3.28156000 | 3.90598000 | 1.45240000 |

| | | | |
|---|---|---|---|
| H | -2.33943000 | 3.87136000 | -0.04210000 |
| H | 1.03257000 | 2.65374000 | 0.13122000 |
| H | -0.32112000 | 1.11540000 | 1.57882000 |

**IM7i-2**
Sum of electronic and thermal Free Energies
= -976.498610

| | | | |
|---|---|---|---|
| C | -0.70081500 | -0.79612300 | -0.67050000 |
| C | -1.82745100 | -0.98607700 | 0.38160900 |
| C | -0.00621700 | 2.42992000 | 0.62078800 |
| C | 0.37040800 | 1.73253200 | -0.72943900 |
| C | 1.73859700 | 2.33054500 | -1.19956300 |
| C | 2.93738500 | 1.37985000 | -1.09048000 |
| C | 2.96248700 | 0.57002400 | 0.20837800 |
| C | 4.07313900 | -0.50890500 | 0.17962400 |
| C | 3.47005400 | -1.66924000 | 0.95581200 |
| C | 2.03049500 | -1.76310600 | 0.41468600 |
| C | 1.60485000 | -0.24282200 | 0.38784200 |
| C | 0.52590300 | 0.17863100 | -0.65669100 |
| C | -1.14543600 | 1.75561100 | 1.30087300 |
| C | -2.45311400 | 1.96838700 | 1.12158600 |
| C | -3.38789100 | 0.82548700 | 1.43549800 |
| C | -3.22886600 | -0.14356900 | 0.23137000 |
| C | -2.63136600 | -2.05240900 | -0.12556500 |
| C | -2.45511300 | -2.15546100 | -1.60750000 |
| C | -1.46396400 | -1.03755800 | -1.99279700 |
| C | -3.36786000 | -3.05097300 | 0.68225000 |
| C | -0.68448000 | 2.10868100 | -1.77649000 |
| C | 3.27910200 | 1.48362600 | 1.40611900 |
| H | -0.18479300 | -1.74580100 | -0.53302900 |
| H | 0.95991900 | -0.07291400 | -1.63692700 |
| H | 1.21403100 | -0.01579600 | 1.39278000 |
| C | 1.20656600 | -2.64012200 | 1.36827500 |
| C | 2.10414500 | -2.42180700 | -0.98023200 |
| C | -3.07126000 | 3.14100300 | 0.41152200 |
| H | -1.55696500 | -1.03078100 | 1.43463900 |
| H | -0.22175400 | 3.47879100 | 0.39009600 |
| H | 1.65277700 | 2.65172200 | -2.24361600 |
| H | 1.93545400 | 3.24250900 | -0.62404900 |
| H | 2.93407000 | 0.68571100 | -1.94092500 |
| H | 3.86769000 | 1.95316500 | -1.18110700 |
| H | 5.00530300 | -0.14038500 | 0.61614600 |
| H | 4.30201500 | -0.81350600 | -0.84752800 |
| H | 4.01300200 | -2.61291900 | 0.84151500 |
| H | 3.44455700 | -1.43384500 | 2.02814200 |
| H | -3.14850700 | 0.31867400 | 2.37502800 |
| H | -4.43357700 | 1.14376300 | 1.48070100 |
| H | -4.13540100 | -0.76870500 | 0.16080700 |
| H | -3.17123200 | 0.39958000 | -0.71223300 |
| H | -3.40409600 | -2.18065600 | -2.15290700 |
| H | -2.00035700 | -3.15108300 | -1.74781100 |
| H | -0.78442800 | -1.35721200 | -2.78640500 |
| H | -1.99431400 | -0.15689600 | -2.35743200 |
| H | -3.58062300 | -2.69458600 | 1.69161900 |
| H | -4.27631700 | -3.40336900 | 0.18900100 |
| H | -2.68705800 | -3.91033300 | 0.77174700 |
| H | -1.69478900 | 1.82744400 | -1.46557600 |
| H | -0.68147200 | 3.19138500 | -1.94172600 |
| H | -0.46812100 | 1.62829600 | -2.73746400 |
| H | 3.24938600 | 0.92304600 | 2.34685800 |
| H | 4.28890200 | 1.89059100 | 1.29390800 |
| H | 2.60948500 | 2.33697900 | 1.50559300 |
| H | 1.11501200 | -2.16533800 | 2.35166800 |
| H | 0.19586300 | -2.86584800 | 1.00588600 |
| H | 1.70761200 | -3.60352700 | 1.51245900 |
| H | 1.13005800 | -2.71294800 | -1.38723700 |
| H | 2.58228500 | -1.77440800 | -1.72034000 |
| H | 2.69806400 | -3.33921300 | -0.91560300 |
| H | -3.65100300 | 2.82675400 | -0.46640100 |
| H | -3.76952900 | 3.65148400 | 1.08386200 |
| H | -2.32836200 | 3.86680100 | 0.07972400 |
| H | 0.85206600 | 2.42915600 | 1.28867000 |
| H | -0.86062300 | 0.88747800 | 1.89561400 |

**TS_7i-2−7i-3**
Sum of electronic and thermal Free Energies
= -976.353218

| | | | |
|---|---|---|---|
| C | 0.49773000 | 1.11523000 | -0.20380000 |
| C | 1.96427000 | 1.12402000 | 0.35047000 |
| C | 0.26594000 | -2.18107000 | 0.47811000 |
| C | -0.11416000 | -1.43119000 | -0.84080000 |
| C | -1.27604000 | -2.20912000 | -1.53840000 |
| C | -2.61620000 | -1.47277000 | -1.54966000 |
| C | -2.97004000 | -0.86168000 | -0.19281000 |
| C | -4.18156000 | 0.08515000 | -0.30115000 |
| C | -3.97100000 | 1.05963000 | 0.84631000 |
| C | -2.47525000 | 1.43175000 | 0.75109000 |
| C | -1.80067000 | 0.05531000 | 0.36146000 |
| C | -0.57127000 | 0.04611000 | -0.60363000 |
| C | 1.20288000 | -1.40532000 | 1.34958000 |
| C | 2.54056000 | -1.46386000 | 1.40149000 |
| C | 3.25933000 | -0.32814000 | 2.09707000 |
| C | 3.76097000 | 0.73067000 | 1.08295000 |
| C | 2.93107000 | 1.42667000 | -0.61361000 |
| C | 2.36620000 | 2.77703000 | -1.05305000 |
| C | 0.91451000 | 2.16177000 | -1.36583000 |
| C | 3.58063000 | 0.61997000 | -1.69831000 |
| C | 1.09940000 | -1.47078000 | -1.76235000 |
| C | -3.32839000 | -1.97779000 | 0.80383000 |
| H | 0.01793000 | 1.67775000 | 0.59368000 |
| H | -0.92112000 | 0.39376000 | -1.58627000 |
| H | -1.45725000 | -0.38324000 | 1.30905000 |

| | | | |
|---|---|---|---|
| C | -2.01180000 | 1.96746000 | 2.11245000 |
| C | -2.32345000 | 2.53701000 | -0.30934000 |
| C | 3.40435000 | -2.49802000 | 0.73338000 |
| H | 1.92540000 | 1.93023000 | 1.08651000 |
| H | -0.63738000 | -2.40388000 | 1.04812000 |
| H | -0.99279000 | -2.42859000 | -2.57343000 |
| H | -1.38936000 | -3.18292000 | -1.04778000 |
| H | -2.59656000 | -0.68119000 | -2.30958000 |
| H | -3.41160000 | -2.16074000 | -1.85926000 |
| H | -5.13039000 | -0.45692000 | -0.24710000 |
| H | -4.16729000 | 0.62016000 | -1.25899000 |
| H | -4.61178000 | 1.94542000 | 0.80189000 |
| H | -4.17257000 | 0.56437000 | 1.80379000 |
| H | 2.60627000 | 0.14585000 | 2.83496000 |
| H | 4.16064000 | -0.67162000 | 2.62208000 |
| H | 4.06015000 | 1.66396000 | 1.56158000 |
| H | 4.56205000 | 0.31328000 | 0.47445000 |
| H | 2.85477000 | 3.19851000 | -1.93463000 |
| H | 2.30651000 | 3.53080000 | -0.26472000 |
| H | 0.16564000 | 2.94303000 | -1.50564000 |
| H | 0.96933000 | 1.59713000 | -2.30230000 |
| H | 4.54561000 | 1.09105000 | -1.91980000 |
| H | 3.74949000 | -0.42352000 | -1.44348000 |
| H | 2.98866000 | 0.66594000 | -2.61721000 |
| H | 1.98308000 | -1.11021000 | -1.24121000 |
| H | 1.30282000 | -2.49838000 | -2.08202000 |
| H | 0.93885000 | -0.86847000 | -2.66499000 |
| H | -2.53694000 | -2.72255000 | 0.91229000 |
| H | -3.53715000 | -1.57382000 | 1.79953000 |
| H | -4.22360000 | -2.50772000 | 0.46287000 |
| H | -0.96646000 | 2.30039000 | 2.12205000 |
| H | -2.62009000 | 2.83131000 | 2.40210000 |
| H | -2.12579000 | 1.20404000 | 2.88931000 |
| H | -3.09624000 | 3.29728000 | -0.16130000 |
| H | -1.36564000 | 3.05668000 | -0.22502000 |
| H | -2.42343000 | 2.16258000 | -1.33372000 |
| H | 3.95821000 | -3.06281000 | 1.49182000 |
| H | 2.82617000 | -3.20724000 | 0.14057000 |
| H | 4.15653000 | -2.04596000 | 0.07467000 |
| H | 0.69112000 | -3.14727000 | 0.18584000 |
| H | 0.72603000 | -0.62473000 | 1.94853000 |

**IM7i-3**

Sum of electronic and thermal Free Energies
= -976.565100

| | | | |
|---|---|---|---|
| C | 0.31614100 | 1.26906100 | -0.38341200 |
| C | 1.45801500 | 0.66363300 | 0.44030800 |
| C | 0.81448200 | -1.67598900 | -0.21460800 |
| C | -0.17958600 | -1.12826300 | -1.26414700 |
| C | -1.29257300 | -2.20678400 | -1.47218100 |
| C | -2.69788700 | -1.65195700 | -1.28504200 |
| C | -2.87407700 | -1.02192000 | 0.09657200 |
| C | -4.23584000 | -0.30875000 | 0.21345000 |
| C | -3.96930500 | 0.83341700 | 1.18208800 |
| C | -2.59550000 | 1.39818900 | 0.76091500 |
| C | -1.79559800 | 0.09583600 | 0.35629200 |
| C | -0.77072300 | 0.24674800 | -0.80357900 |
| C | 1.64454800 | -0.74864500 | 0.55330500 |
| C | 2.67525400 | -1.19491300 | 1.53545800 |
| C | 4.01685100 | -0.38306200 | 1.39108200 |
| C | 3.85959500 | 1.07688400 | 0.94356200 |
| C | 2.87964900 | 1.14352300 | -0.22558800 |
| C | 2.38810100 | 2.49954800 | -0.75365400 |
| C | 1.02563800 | 2.18862000 | -1.40962300 |
| C | 3.45822200 | 0.31854200 | -1.39689600 |
| C | 0.53132600 | -0.97992200 | -2.61713200 |
| C | -2.79036800 | -2.09681900 | 1.19216300 |
| H | -0.17828000 | 1.95375400 | 0.30230800 |
| H | -1.29328900 | 0.66085300 | -1.67457200 |
| H | -1.22137800 | -0.20140600 | 1.25223700 |
| C | -2.00669100 | 2.16160600 | 1.95744600 |
| C | -2.79773400 | 2.36431900 | -0.41702600 |
| C | 2.95147400 | -2.69496900 | 1.57907300 |
| H | 1.52304400 | 1.12353900 | 1.43124900 |
| H | 0.25058800 | -2.12893400 | 0.63397100 |
| H | -1.19790400 | -2.63134500 | -2.47600200 |
| H | -1.14526600 | -3.04735200 | -0.78367100 |
| H | -2.90775900 | -0.90278700 | -2.05987000 |
| H | -3.43539600 | -2.45099000 | -1.42407000 |
| H | -5.02363200 | -0.98928600 | 0.55161500 |
| H | -4.54649700 | 0.08275800 | -0.76225900 |
| H | -4.74246100 | 1.60833400 | 1.17239500 |
| H | -3.90953000 | 0.45208400 | 2.20965900 |
| H | 4.53578600 | -0.44354600 | 2.35038100 |
| H | 4.63603300 | -0.92434500 | 0.66795700 |
| H | 3.51979000 | 1.71294100 | 1.76822300 |
| H | 4.84014700 | 1.45085700 | 0.62766500 |
| H | 3.10922200 | 2.93667000 | -1.45178400 |
| H | 2.25481200 | 3.19726500 | 0.08265000 |
| H | 0.44822000 | 3.09828000 | -1.59268000 |
| H | 1.16369400 | 1.70754700 | -2.38047800 |
| H | 4.52644000 | 0.54897900 | -1.46296700 |
| H | 3.36970000 | -0.76770200 | -1.29374700 |
| H | 3.01909900 | 0.59774600 | -2.35314100 |
| H | 1.34964000 | -0.26764700 | -2.59867300 |
| H | 0.93420400 | -1.94450000 | -2.94425100 |
| H | -0.18802700 | -0.64761700 | -3.37250700 |
| H | -1.79347900 | -2.54695600 | 1.26529800 |
| H | -3.01970400 | -1.66869900 | 2.17342600 |
| H | -3.50446000 | -2.90467300 | 1.00104300 |
| H | -1.11999000 | 2.75660200 | 1.71530700 |
| H | -2.75275700 | 2.86319300 | 2.34580500 |

| | | | |
|---|---|---|---|
| H | -1.74878400 | 1.47409000 | 2.77112300 |
| H | -3.44181700 | 3.19164700 | -0.10334500 |
| H | -1.85798300 | 2.80227100 | -0.76924900 |
| H | -3.27770100 | 1.87774400 | -1.27241900 |
| H | 2.05255500 | -3.27816000 | 1.79513200 |
| H | 3.37719300 | -3.04697400 | 0.63386400 |
| H | 3.67902300 | -2.90721000 | 2.36546200 |
| H | 1.39966500 | -2.52691100 | -0.59205300 |
| H | 2.23675400 | -0.87771800 | 2.49801500 |

**TS_7i-2−13**
Sum of electronic and thermal Free Energies
= -976.494214

| | | | |
|---|---|---|---|
| C | 0.71689000 | -0.54662000 | 0.95204000 |
| C | 1.66041000 | -1.10993000 | -0.19712000 |
| C | 0.76985000 | 2.57057000 | 0.13789000 |
| C | -0.48306000 | 1.91148000 | 0.78234000 |
| C | -1.71256000 | 2.53406000 | 0.07704000 |
| C | -2.96663000 | 1.68267000 | 0.17871000 |
| C | -2.81372000 | 0.41232000 | -0.65617000 |
| C | -3.99383000 | -0.55948000 | -0.44635000 |
| C | -3.35968000 | -1.93645000 | -0.58260000 |
| C | -2.02840000 | -1.82417000 | 0.18968000 |
| C | -1.52211000 | -0.39196000 | -0.25031000 |
| C | -0.57342000 | 0.31564000 | 0.77024000 |
| C | 1.20976000 | 1.77650000 | -1.04330000 |
| C | 2.43408000 | 1.30608000 | -1.29603000 |
| C | 2.55275000 | 0.12314000 | -2.23940000 |
| C | 1.62091000 | -1.00274000 | -1.73922000 |
| C | 2.90713000 | -1.41010000 | 0.42162000 |
| C | 2.88601000 | -1.08124000 | 1.87742000 |
| C | 1.73769000 | -0.07477000 | 2.00602000 |
| C | 4.08309000 | -2.03997000 | -0.22070000 |
| C | -0.53131000 | 2.37049000 | 2.25234000 |
| C | -2.74730000 | 0.75619000 | -2.15550000 |
| H | 0.30153000 | -1.46784000 | 1.37496000 |
| H | -1.06372000 | 0.13809000 | 1.73593000 |
| H | -0.97334000 | -0.55171000 | -1.18225000 |
| C | -1.12211000 | -2.99050000 | -0.23573000 |
| C | -2.32863000 | -1.94034000 | 1.69319000 |
| C | 3.68708000 | 1.74967000 | -0.59038000 |
| H | 1.63159000 | -2.27231000 | 0.00057000 |
| H | 0.49072000 | 3.59746000 | -0.13238000 |
| H | -1.87563000 | 3.52620000 | 0.51364000 |
| H | -1.48774000 | 2.70981000 | -0.98130000 |
| H | -3.16432000 | 1.41732000 | 1.22741000 |
| H | -3.84269000 | 2.24313000 | -0.16799000 |
| H | -4.79598000 | -0.38160000 | -1.16910000 |
| H | -4.42901000 | -0.43514000 | 0.55155000 |
| H | -3.98222000 | -2.75187000 | -0.20033000 |
| H | -3.15136000 | -2.15525000 | -1.63861000 |
| H | 2.27649000 | 0.38083000 | -3.26700000 |
| H | 3.59009000 | -0.22733000 | -2.27853000 |
| H | 0.59795000 | -0.80402000 | -2.04766000 |
| H | 1.88825000 | -1.96539000 | -2.18951000 |
| H | 3.87184000 | -0.72540000 | 2.19589000 |
| H | 2.70147000 | -2.00865000 | 2.44232000 |
| H | 1.30677000 | -0.06358000 | 3.00904000 |
| H | 2.10779000 | 0.92872000 | 1.77832000 |
| H | 3.87700000 | -2.44186000 | -1.21264000 |
| H | 4.84533000 | -1.25469000 | -0.31944000 |
| H | 4.51107000 | -2.80793000 | 0.43005000 |
| H | 0.25989000 | 1.91909000 | 2.85738000 |
| H | -0.41066000 | 3.45703000 | 2.31008000 |
| H | -1.49180000 | 2.11678000 | 2.71394000 |
| H | -1.82032000 | 1.26790000 | -2.43296000 |
| H | -2.80416000 | -0.15658000 | -2.75863000 |
| H | -3.58557000 | 1.39950000 | -2.44312000 |
| H | -0.26905000 | -3.14703000 | 0.43665000 |
| H | -1.69052000 | -3.92646000 | -0.21822000 |
| H | -0.75016000 | -2.85600000 | -1.25905000 |
| H | -2.84692000 | -2.88458000 | 1.88847000 |
| H | -1.42498000 | -1.94094000 | 2.31289000 |
| H | -2.97091000 | -1.13006000 | 2.05113000 |
| H | 4.46936000 | 1.97326000 | -1.32409000 |
| H | 3.53587000 | 2.64398000 | 0.01535000 |
| H | 4.09667000 | 0.96518000 | 0.06484000 |
| H | 1.58153000 | 2.66313000 | 0.86564000 |
| H | 0.39261000 | 1.40567000 | -1.66755000 |

**TS_13−13-1**
Sum of electronic and thermal Free Energies
= -976.481918

| | | | |
|---|---|---|---|
| C | -0.70289000 | 0.65943000 | -0.59500000 |
| C | -2.16405000 | 0.77176000 | -0.24546000 |
| C | -1.49877000 | -1.22332000 | 1.22844000 |
| C | 0.04623000 | -1.09298000 | 1.15821000 |
| C | 0.55645000 | -2.38306000 | 0.48312000 |
| C | 2.04561000 | -2.34817000 | 0.18482000 |
| C | 2.39887000 | -1.24804000 | -0.82032000 |
| C | 3.93254000 | -0.99536000 | -0.84716000 |
| C | 4.10945000 | 0.52445000 | -0.76498000 |
| C | 2.95289000 | 1.00727000 | 0.12879000 |
| C | 1.78863000 | 0.14483000 | -0.44510000 |
| C | 0.42611000 | 0.18684000 | 0.33250000 |
| C | -2.17029000 | -0.71896000 | -0.00372000 |
| C | -3.37090000 | -1.39634000 | -0.66904000 |
| C | -4.15022000 | -0.19623000 | -1.23655000 |
| C | -3.06101000 | 0.83499000 | -1.52507000 |
| C | -2.24402000 | 2.09851000 | 0.52167000 |
| C | -1.17536000 | 2.89533000 | -0.33604000 |
| C | -0.48693000 | 1.92434000 | -1.39273000 |

| | | | |
|---|---|---|---|
| C | -1.95592000 | 2.10614000 | 2.02105000 |
| C | 0.55922000 | -1.03143000 | 2.60161000 |
| C | 1.90831000 | -1.64742000 | -2.22157000 |
| H | -1.15772000 | -0.72678000 | -0.95940000 |
| H | 0.50555000 | 1.02078000 | 1.05122000 |
| H | 1.59630000 | 0.60120000 | -1.42147000 |
| C | 2.70868000 | 2.50755000 | -0.04065000 |
| C | 3.27179000 | 0.73546000 | 1.60272000 |
| C | -4.18734000 | -2.24193000 | 0.31468000 |
| H | -3.23270000 | 2.54219000 | 0.37624000 |
| H | -1.74852000 | -2.27044000 | 1.41578000 |
| H | 0.32344000 | -3.21826000 | 1.15405000 |
| H | -0.00325000 | -2.57216000 | -0.44396000 |
| H | 2.60227000 | -2.20443000 | 1.11714000 |
| H | 2.37118000 | -3.31584000 | -0.21417000 |
| H | 4.39557000 | -1.42015000 | -1.74251000 |
| H | 4.40979000 | -1.47791000 | 0.01258000 |
| H | 5.08839000 | 0.82272000 | -0.37708000 |
| H | 4.00381000 | 0.97323000 | -1.76165000 |
| H | -4.74191000 | -0.45208000 | -2.11786000 |
| H | -4.83901000 | 0.17952000 | -0.46873000 |
| H | -2.50317000 | 0.50392000 | -2.40997000 |
| H | -3.44141000 | 1.83908000 | -1.72437000 |
| H | -0.40235000 | 3.30054000 | 0.32317000 |
| H | -1.62744000 | 3.73247000 | -0.87157000 |
| H | -1.02541000 | 1.95539000 | -2.33833000 |
| H | 0.55320000 | 2.18098000 | -1.57104000 |
| H | -2.72183000 | 1.55018000 | 2.56975000 |
| H | -0.98028000 | 1.68400000 | 2.27448000 |
| H | -1.97085000 | 3.13530000 | 2.39026000 |
| H | 0.23994000 | -0.11005000 | 3.10019000 |
| H | 0.15219000 | -1.87617000 | 3.16708000 |
| H | 1.64514000 | -1.08583000 | 2.66313000 |
| H | 0.81340000 | -1.66281000 | -2.29160000 |
| H | 2.27181000 | -0.93451000 | -2.96963000 |
| H | 2.27602000 | -2.64089000 | -2.49765000 |
| H | 1.83276000 | 2.84714000 | 0.52787000 |
| H | 3.56660000 | 3.07639000 | 0.33073000 |
| H | 2.57081000 | 2.77762000 | -1.09359000 |
| H | 4.14523000 | 1.32366000 | 1.90076000 |
| H | 2.44584000 | 1.03110000 | 2.25953000 |
| H | 3.50511000 | -0.31513000 | 1.79441000 |
| H | -5.11771000 | -2.54513000 | -0.17279000 |
| H | -3.66085000 | -3.14775000 | 0.62207000 |
| H | -4.45061000 | -1.66617000 | 1.20782000 |
| H | -1.89071000 | -0.63066000 | 2.06464000 |
| H | -3.02121000 | -2.04627000 | -1.48079000 |

**IM13-1**
Sum of electronic and thermal Free Energies
= -976.553240

| | | | |
|---|---|---|---|
| C | -0.52158800 | 0.89585700 | -0.25729000 |
| C | -1.88108800 | 0.41685500 | -0.47804700 |
| C | -1.42144100 | -1.36157700 | 1.27240400 |
| C | 0.12115300 | -1.19481900 | 1.16519100 |
| C | 0.68354300 | -2.41009700 | 0.39578900 |
| C | 2.13426900 | -2.24826600 | -0.02732300 |
| C | 2.35335900 | -1.05423800 | -0.96552100 |
| C | 3.87721600 | -0.80962900 | -1.17579600 |
| C | 4.13932100 | 0.65985800 | -0.81275600 |
| C | 3.09423900 | 1.01262000 | 0.26656400 |
| C | 1.86566400 | 0.29316400 | -0.35873000 |
| C | 0.51233900 | 0.18280500 | 0.50116200 |
| C | -2.07644900 | -1.03456800 | -0.05447100 |
| C | -3.53223700 | -1.36940800 | -0.41750400 |
| C | -3.74617000 | -0.51421300 | -1.70707500 |
| C | -2.47000900 | 0.34509700 | -1.92721600 |
| C | -2.61329000 | 1.65706300 | 0.22414300 |
| C | -1.85130600 | 2.86475300 | -0.38605400 |
| C | -0.41517500 | 2.33731200 | -0.61726900 |
| C | -2.58167200 | 1.71041600 | 1.75032100 |
| C | 0.67579900 | -1.22832500 | 2.59849500 |
| C | 1.62925000 | -1.28431400 | -2.29772200 |
| H | -1.49774300 | -1.58514800 | -0.81000400 |
| H | 0.70359400 | 0.90676100 | 1.32095100 |
| H | 1.63105500 | 0.91875000 | -1.23006500 |
| C | 2.90428500 | 2.52588000 | 0.37133200 |
| C | 3.53186000 | 0.49471300 | 1.63929200 |
| C | -4.57196100 | -1.09956400 | 0.66954700 |
| H | -3.65430300 | 1.63472700 | -0.10970300 |
| H | -1.61904200 | -2.40399400 | 1.55101600 |
| H | 0.58511500 | -3.28225300 | 1.05231800 |
| H | 0.07598600 | -2.63186200 | -0.48748000 |
| H | 2.76959200 | -2.15031000 | 0.85797300 |
| H | 2.47373700 | -3.15519900 | -0.54144000 |
| H | 4.18480800 | -1.04043400 | -2.19951100 |
| H | 4.44975500 | -1.47142000 | -0.51772900 |
| H | 5.16122500 | 0.83776800 | -0.46487800 |
| H | 3.97852500 | 1.30256800 | -1.68788700 |
| H | -3.93526600 | -1.13433700 | -2.58596400 |
| H | -4.61981200 | 0.13431100 | -1.58688900 |
| H | -1.75683500 | -0.16709100 | -2.57999300 |
| H | -2.68711700 | 1.32293400 | -2.36024900 |
| H | -1.85006500 | 3.71327100 | 0.30103800 |
| H | -2.30421400 | 3.20411300 | -1.31836400 |
| H | -0.11687800 | 2.39371400 | -1.67574300 |
| H | 0.35826000 | 2.85345400 | -0.04761900 |
| H | -3.11022600 | 0.87353100 | 2.20901100 |
| H | -1.55871300 | 1.72730000 | 2.14806800 |
| H | -3.07267600 | 2.63115500 | 2.07821500 |
| H | 0.36060900 | -0.34719000 | 3.16983200 |

| | | | |
|---|---|---|---|
| H | 0.28848500 | -2.11196100 | 3.11456800 |
| H | 1.76510000 | -1.28042900 | 2.62692600 |
| H | 0.53661800 | -1.23349300 | -2.18247900 |
| H | 1.91941500 | -0.52314900 | -3.02990900 |
| H | 1.86843100 | -2.26668900 | -2.71711800 |
| H | 2.17091800 | 2.79365500 | 1.14393000 |
| H | 3.84684700 | 3.00159200 | 0.65846800 |
| H | 2.59167100 | 2.96270900 | -0.58378900 |
| H | 4.42922200 | 1.03530700 | 1.95535100 |
| H | 2.76568400 | 0.67449400 | 2.40132100 |
| H | 3.77705500 | -0.56934000 | 1.63807400 |
| H | -5.55980500 | -1.42511500 | 0.33182300 |
| H | -4.33824800 | -1.65038900 | 1.58635300 |
| H | -4.65028800 | -0.03784700 | 0.91963900 |
| H | -1.81324200 | -0.74342000 | 2.08540800 |
| H | -3.58669000 | -2.43528600 | -0.66227800 |

**TS_7b−7c**
Sum of electronic and thermal Free Energies
= -976.503496

| | | | |
|---|---|---|---|
| C | 2.04458000 | 0.30133000 | -0.21977000 |
| C | 3.03028000 | -0.83101000 | -0.68533000 |
| C | 2.61759000 | -2.21702000 | -0.17942000 |
| C | 1.14758000 | -2.51527000 | -0.43695000 |
| C | 0.18893000 | -1.61130000 | 0.35897000 |
| C | 0.50835000 | -0.10798000 | 0.15049000 |
| H | 3.24805000 | -2.96529000 | -0.67222000 |
| H | 2.06654000 | 0.99773000 | -1.09887000 |
| H | 0.92643000 | -2.44070000 | -1.51092000 |
| H | 0.92399000 | -3.55148000 | -0.16255000 |
| H | 0.18649000 | 0.52661000 | 0.97920000 |
| H | 2.82825000 | -2.30177000 | 0.89183000 |
| C | 2.74585000 | 1.18771000 | 0.87738000 |
| C | 4.37916000 | -0.36597000 | -0.06338000 |
| H | 4.61733000 | -0.97917000 | 0.81081000 |
| H | 5.20194000 | -0.48947000 | -0.77202000 |
| C | 4.19020000 | 1.09579000 | 0.35182000 |
| H | 4.91260000 | 1.42002000 | 1.10633000 |
| H | 4.30191000 | 1.76032000 | -0.51551000 |
| C | -1.24057000 | -1.90896000 | -0.14193000 |
| H | -1.37680000 | -2.99531000 | -0.06289000 |
| C | 0.38111000 | 0.52028000 | -1.12527000 |
| H | 0.47469000 | -0.13877000 | -1.99239000 |
| C | -2.35607000 | -1.18828000 | 0.57292000 |
| C | -2.44976000 | 2.01574000 | -0.56716000 |
| C | -0.02510000 | 1.93195000 | -1.42813000 |
| H | 0.19881000 | 2.58094000 | -0.58236000 |
| H | 0.55223000 | 2.27921000 | -2.29071000 |
| C | -3.55380000 | 1.26086000 | -0.59512000 |
| H | -3.74173000 | 0.68311000 | -1.50189000 |
| C | -1.53448000 | 2.06018000 | -1.77040000 |
| H | -1.62421000 | 3.03667000 | -2.26334000 |
| H | -1.83027000 | 1.30727000 | -2.50921000 |
| C | -4.54616000 | 1.05405000 | 0.51285000 |
| H | -4.26910000 | 1.64946000 | 1.38437000 |
| H | -5.52584000 | 1.42081000 | 0.18814000 |
| C | -3.65806000 | -1.37184000 | 0.33057000 |
| H | -2.07967000 | -0.39838000 | 1.27359000 |
| C | -4.69326000 | -0.44838000 | 0.94187000 |
| H | -4.63725000 | -0.50434000 | 2.03313000 |
| H | -5.69031000 | -0.80505000 | 0.67021000 |
| C | 3.16140000 | -0.84153000 | -2.21650000 |
| H | 2.23326000 | -1.11392000 | -2.72926000 |
| H | 3.48441000 | 0.13667000 | -2.59231000 |
| H | 3.91619000 | -1.57393000 | -2.51456000 |
| C | 2.64281000 | 0.55838000 | 2.26964000 |
| H | 3.25121000 | 1.14284000 | 2.96616000 |
| H | 1.61640000 | 0.56370000 | 2.64859000 |
| H | 3.01091000 | -0.46983000 | 2.29162000 |
| C | 2.25922000 | 2.63279000 | 0.94068000 |
| H | 2.89005000 | 3.18787000 | 1.64103000 |
| H | 2.32826000 | 3.13251000 | -0.03146000 |
| H | 1.23051000 | 2.70822000 | 1.31127000 |
| C | 0.26575000 | -1.94803000 | 1.85540000 |
| H | 1.27742000 | -1.87077000 | 2.25589000 |
| H | -0.37897000 | -1.29332000 | 2.44872000 |
| H | -0.07989000 | -2.97435000 | 2.01075000 |
| C | -2.04602000 | 2.88749000 | 0.59786000 |
| H | -2.88942000 | 3.13812000 | 1.24014000 |
| H | -1.28894000 | 2.40270000 | 1.23163000 |
| H | -1.61447000 | 3.83226000 | 0.24705000 |
| C | -4.18233000 | -2.38751000 | -0.64703000 |
| H | -3.40766000 | -3.03989000 | -1.05340000 |
| H | -4.94343000 | -3.01579000 | -0.17205000 |
| H | -4.67729000 | -1.88140000 | -1.48637000 |
| H | -1.28584000 | -1.68502000 | -1.22057000 |

**IM7c**
Sum of electronic and thermal Free Energies
= -976.506263

| | | | |
|---|---|---|---|
| C | 2.06038400 | 0.36816100 | -0.18838300 |
| C | 3.04203000 | -0.67334200 | -0.83948200 |
| C | 2.71293400 | -2.11694600 | -0.44473600 |
| C | 1.23969500 | -2.44695600 | -0.63889900 |
| C | 0.31140500 | -1.66377800 | 0.30434300 |
| C | 0.57449600 | -0.13615100 | 0.24035400 |
| H | 3.33564800 | -2.78841400 | -1.04605300 |
| H | 1.99319900 | 1.14724000 | -0.99239200 |
| H | 0.94642500 | -2.26996200 | -1.68288800 |
| H | 1.06681200 | -3.51354400 | -0.46090200 |
| H | 0.30387000 | 0.38744900 | 1.15985000 |
| H | 2.99583200 | -2.29281700 | 0.59863000 |

| | | | |
|---|---|---|---|
| C | 2.81424900 | 1.16840000 | 0.93688500 |
| C | 4.41904000 | -0.21686600 | -0.27530700 |
| H | 4.74411700 | -0.90256300 | 0.51274900 |
| H | 5.18776800 | -0.23747000 | -1.05188400 |
| C | 4.21504400 | 1.18882200 | 0.29786500 |
| H | 4.98371500 | 1.46710600 | 1.02440600 |
| H | 4.23562900 | 1.93891900 | -0.50433500 |
| C | -1.14331200 | -1.94892600 | -0.12353600 |
| H | -1.29204900 | -3.03420400 | -0.05113400 |
| C | 0.32354500 | 0.63918400 | -0.93370200 |
| H | 0.36952500 | 0.10451500 | -1.88655500 |
| C | -2.19283500 | -1.20377000 | 0.65680400 |
| C | -2.70049900 | 1.89135000 | -0.57693200 |
| C | -0.15238600 | 2.06169300 | -1.01444800 |
| H | -0.08862700 | 2.54271300 | -0.03778900 |
| H | 0.50213100 | 2.60431100 | -1.70472200 |
| C | -3.72113300 | 1.09621100 | -0.91624000 |
| H | -3.69107200 | 0.63605200 | -1.90476300 |
| C | -1.59799600 | 2.18226600 | -1.56876300 |
| H | -1.68296200 | 3.22638600 | -1.89617300 |
| H | -1.71526500 | 1.56424600 | -2.46590200 |
| C | -4.87181400 | 0.69164000 | -0.02855900 |
| H | -5.33385000 | 1.56610900 | 0.44256200 |
| H | -5.64856800 | 0.22736400 | -0.64273100 |
| C | -3.51428800 | -1.34510800 | 0.50989300 |
| H | -1.85217500 | -0.40267700 | 1.31403900 |
| C | -4.43357200 | -0.30240400 | 1.08613500 |
| H | -3.90964700 | 0.25003800 | 1.87341900 |
| H | -5.32721500 | -0.75025500 | 1.53397600 |
| C | 3.05327200 | -0.52766400 | -2.36915000 |
| H | 2.09448900 | -0.77567700 | -2.83567200 |
| H | 3.32338200 | 0.49278200 | -2.66605400 |
| H | 3.79940800 | -1.20277600 | -2.79636400 |
| C | 2.84400500 | 0.40272200 | 2.26297000 |
| H | 3.48421800 | 0.94265800 | 2.96689200 |
| H | 1.85127400 | 0.32691300 | 2.71651800 |
| H | 3.24932100 | -0.60592100 | 2.15454800 |
| C | 2.28062300 | 2.57716900 | 1.18118900 |
| H | 2.95428200 | 3.09692400 | 1.86861900 |
| H | 2.23214700 | 3.16559000 | 0.25875600 |
| H | 1.29138700 | 2.56839300 | 1.65232700 |
| C | 0.49332700 | -2.14087100 | 1.75270100 |
| H | 1.51536400 | -2.02043700 | 2.11474800 |
| H | -0.17148400 | -1.60123400 | 2.43426200 |
| H | 0.23658000 | -3.20221700 | 1.81765000 |
| C | -2.59782300 | 2.58739800 | 0.75544100 |
| H | -3.55496900 | 2.60245600 | 1.27953800 |
| H | -1.87534400 | 2.08961000 | 1.41862400 |
| H | -2.26452900 | 3.62497100 | 0.63541200 |
| C | -4.13999600 | -2.36459000 | -0.40213100 |
| H | -4.42915800 | -1.90200600 | -1.35559000 |
| H | -3.46860000 | -3.19506100 | -0.62892800 |
| H | -5.05249200 | -2.77382300 | 0.04181300 |
| H | -1.24888100 | -1.70754500 | -1.19437500 |

**TS_7c−7-1**

Sum of electronic and thermal Free Energies
= -976.470290

| | | | |
|---|---|---|---|
| C | 1.78067000 | 0.40413000 | 0.07370000 |
| C | 2.86773000 | -0.21860000 | -0.86418000 |
| C | 2.61368000 | -1.68865000 | -1.20863000 |
| C | 1.18502000 | -1.91439000 | -1.67965000 |
| C | 0.11972000 | -1.56254000 | -0.62036000 |
| C | 0.27311000 | -0.05901000 | -0.13851000 |
| H | 3.31849000 | -1.99935000 | -1.98892000 |
| H | 1.81156000 | 1.45093000 | -0.22145000 |
| H | 0.99491000 | -1.32960000 | -2.59043000 |
| H | 1.04313000 | -2.96447000 | -1.96339000 |
| H | -0.19096000 | -0.00151000 | 0.86159000 |
| H | 2.82783000 | -2.32420000 | -0.34289000 |
| C | 2.39285000 | 0.52078000 | 1.50813000 |
| C | 4.18156000 | -0.03864000 | -0.05378000 |
| H | 4.51900000 | -1.00694000 | 0.33107000 |
| H | 4.99153000 | 0.35291000 | -0.67614000 |
| C | 3.83556000 | 0.89772000 | 1.11085000 |
| H | 4.53159000 | 0.81041000 | 1.95079000 |
| H | 3.85798000 | 1.94403000 | 0.77718000 |
| C | -1.24270000 | -1.94845000 | -1.27684000 |
| H | -1.31636000 | -3.04093000 | -1.21758000 |
| C | -0.43013000 | 0.99639000 | -0.98126000 |
| H | -0.86628000 | 0.64552000 | -1.91944000 |
| C | -2.47677000 | -1.29717000 | -0.72819000 |
| C | -2.08482000 | 1.92767000 | -0.21312000 |
| C | -0.00039000 | 2.45726000 | -1.03965000 |
| H | 0.52755000 | 2.74888000 | -0.13191000 |
| H | 0.63876000 | 2.68809000 | -1.89799000 |
| C | -3.20326000 | 1.34434000 | -0.79287000 |
| H | -3.30668000 | 1.51998000 | -1.86648000 |
| C | -1.39829000 | 3.04443000 | -1.01897000 |
| H | -1.48259000 | 4.01539000 | -0.52374000 |
| H | -1.83213000 | 3.12658000 | -2.01880000 |
| C | -4.39180000 | 0.68754000 | -0.16745000 |
| H | -4.94382000 | 1.50359000 | 0.32810000 |
| H | -5.03483000 | 0.32592000 | -0.97367000 |
| C | -3.12673000 | -1.44992000 | 0.43131000 |
| H | -2.86928000 | -0.54891000 | -1.41362000 |
| C | -4.17086000 | -0.44020000 | 0.85093000 |
| H | -3.87533000 | -0.03099000 | 1.82448000 |
| H | -5.13542000 | -0.93170000 | 1.02351000 |
| C | 2.93038000 | 0.60625000 | -2.15980000 |
| H | 1.96013000 | 0.64926000 | -2.67228000 |
| H | 3.23788000 | 1.63638000 | -1.94568000 |

| | | | |
|---|---|---|---|
| H | 3.65272000 | 0.17892000 | -2.86229000 |
| C | 2.41720000 | -0.73390000 | 2.38558000 |
| H | 2.95606000 | -0.50391000 | 3.31043000 |
| H | 1.41067000 | -1.05510000 | 2.66923000 |
| H | 2.93263000 | -1.57330000 | 1.91111000 |
| C | 1.72479000 | 1.63736000 | 2.31301000 |
| H | 2.27312000 | 1.81645000 | 3.24319000 |
| H | 1.70601000 | 2.58390000 | 1.75932000 |
| H | 0.69849000 | 1.37111000 | 2.59310000 |
| C | 0.31009000 | -2.47072000 | 0.60269000 |
| H | 1.31278000 | -2.40495000 | 1.01168000 |
| H | -0.38751000 | -2.21102000 | 1.40277000 |
| H | 0.14099000 | -3.51567000 | 0.32392000 |
| C | -1.93390000 | 1.95471000 | 1.28861000 |
| H | -2.77722000 | 2.48692000 | 1.74356000 |
| H | -1.90437000 | 0.94493000 | 1.70940000 |
| H | -1.01896000 | 2.47421000 | 1.58217000 |
| C | -2.92616000 | -2.57490000 | 1.41138000 |
| H | -3.89754000 | -3.02475000 | 1.64322000 |
| H | -2.27777000 | -3.35714000 | 1.01637000 |
| H | -2.50866000 | -2.21915000 | 2.36066000 |
| H | -1.19731000 | -1.71970000 | -2.34767000 |

**IM7-1**
Sum of electronic and thermal Free Energies
= -976.533136

| | | | |
|---|---|---|---|
| C | 2.06893500 | 0.31491800 | -0.19619600 |
| C | 2.89247600 | -0.83123700 | -0.86432700 |
| C | 2.32880800 | -2.21668700 | -0.53150900 |
| C | 0.84603800 | -2.30737100 | -0.84646200 |
| C | -0.01701900 | -1.36404300 | 0.02330200 |
| C | 0.53518400 | 0.11356100 | 0.05299300 |
| H | 2.87303300 | -2.97561700 | -1.10624800 |
| H | 2.13573600 | 1.15314400 | -0.90286600 |
| H | 0.69388200 | -2.06945200 | -1.90717700 |
| H | 0.48952000 | -3.33576000 | -0.70631300 |
| H | 0.36695500 | 0.46206000 | 1.07883800 |
| H | 2.50163400 | -2.45590400 | 0.52373800 |
| C | 2.91115800 | 0.80827300 | 1.02101700 |
| C | 4.32662900 | -0.67303300 | -0.28597100 |
| H | 4.54373700 | -1.48410500 | 0.41811800 |
| H | 5.08716800 | -0.72703400 | -1.07052100 |
| C | 4.33105200 | 0.67427100 | 0.44255100 |
| H | 5.10589700 | 0.74583200 | 1.21257000 |
| H | 4.50412100 | 1.49034400 | -0.27143000 |
| C | -1.45734400 | -1.56879500 | -0.61248900 |
| H | -1.69781100 | -2.62903700 | -0.48671200 |
| C | -0.30036300 | 1.03897700 | -0.83726300 |
| H | -0.47010400 | 0.56774100 | -1.81873700 |
| C | -2.66641600 | -0.78447500 | -0.34756000 |
| C | -1.66702300 | 1.55558000 | -0.28809200 |
| C | 0.06737500 | 2.51428100 | -1.12964900 |
| H | 0.57636600 | 2.99021300 | -0.28923100 |
| H | 0.62962100 | 2.71620800 | -2.04286000 |
| C | -2.85538800 | 0.65435800 | -0.60270600 |
| H | -2.92790300 | 0.57232700 | -1.71715400 |
| C | -1.44770500 | 2.83252900 | -1.13546000 |
| H | -1.77349900 | 3.77924100 | -0.69800700 |
| H | -1.87247800 | 2.73464000 | -2.13991200 |
| C | -4.27740900 | 0.98285100 | -0.10799200 |
| H | -4.24924600 | 1.28317200 | 0.94305400 |
| H | -4.72446700 | 1.80208900 | -0.67337400 |
| C | -3.94104800 | -1.39867200 | 0.05292100 |
| H | -4.09095500 | -2.39115300 | -0.38237800 |
| C | -5.01870200 | -0.35109100 | -0.27526600 |
| H | -5.89394800 | -0.44538100 | 0.37038500 |
| H | -5.35070400 | -0.47855900 | -1.31100500 |
| C | 2.91508500 | -0.61926700 | -2.38629500 |
| H | 1.90831500 | -0.51479800 | -2.80828800 |
| H | 3.46112600 | 0.29862300 | -2.62857000 |
| H | 3.41303500 | -1.45145300 | -2.89541300 |
| C | 2.77084900 | -0.05397100 | 2.28189100 |
| H | 3.45771600 | 0.31145500 | 3.05200700 |
| H | 1.76031300 | 0.00389300 | 2.70195200 |
| H | 3.01180700 | -1.10586200 | 2.10492300 |
| C | 2.60274400 | 2.25790200 | 1.39675500 |
| H | 3.27918300 | 2.59857200 | 2.18734400 |
| H | 2.72831600 | 2.92631300 | 0.53815800 |
| H | 1.57990500 | 2.36976000 | 1.77875600 |
| C | -0.09847100 | -1.91534500 | 1.45424300 |
| H | 0.88955900 | -1.93367300 | 1.91611400 |
| H | -0.73861200 | -1.28846200 | 2.08621400 |
| H | -0.48871200 | -2.93930300 | 1.46584400 |
| C | -1.64373600 | 1.88803500 | 1.20524400 |
| H | -2.49436200 | 2.52320900 | 1.46787000 |
| H | -1.67970500 | 0.98912200 | 1.83415100 |
| H | -0.73777400 | 2.43770600 | 1.47185100 |
| C | -3.75059900 | -1.58616300 | 1.59857000 |
| H | -4.70757300 | -1.94221700 | 1.98642800 |
| H | -2.98005800 | -2.32585700 | 1.82090800 |
| H | -3.50439700 | -0.64827600 | 2.10384100 |
| H | -1.34530800 | -1.40607800 | -1.70036800 |

**TS_7-1−7-2**
Sum of electronic and thermal Free Energies
= -976.524491

| | | | |
|---|---|---|---|
| C | 2.11577000 | 0.24791000 | -0.27394000 |
| C | 2.88877000 | -1.02474000 | -0.78090000 |
| C | 2.30058000 | -2.31607000 | -0.18623000 |
| C | 0.81213000 | -2.45122000 | -0.47166000 |
| C | -0.07232000 | -1.34012000 | 0.16727000 |
| C | 0.58032000 | 0.12423000 | 0.07212000 |

| | | | |
|---|---|---|---|
| H | 2.82822000 | -3.18179000 | -0.60456000 |
| H | 2.17389000 | 0.96240000 | -1.10412000 |
| H | 0.66682000 | -2.45620000 | -1.55834000 |
| H | 0.44866000 | -3.42447000 | -0.11875000 |
| H | 0.49273000 | 0.52576000 | 1.08496000 |
| H | 2.47851000 | -2.35049000 | 0.89404000 |
| C | 3.02692000 | 0.89552000 | 0.83936000 |
| C | 4.35739000 | -0.81973000 | -0.28764000 |
| H | 4.59337000 | -1.53479000 | 0.50787000 |
| H | 5.08219000 | -0.99937000 | -1.08720000 |
| C | 4.42791000 | 0.61602000 | 0.24794000 |
| H | 5.22760000 | 0.76182000 | 0.98134000 |
| H | 4.61751000 | 1.31822000 | -0.57394000 |
| C | -1.43759000 | -1.54017000 | -0.59527000 |
| H | -1.74451000 | -2.57219000 | -0.39690000 |
| C | -0.28820000 | 1.03999000 | -0.80725000 |
| H | -0.55603000 | 0.50706000 | -1.72988000 |
| C | -2.68514000 | -0.70008000 | -0.37395000 |
| C | -1.61081000 | 1.62196000 | -0.20008000 |
| C | 0.04689000 | 2.48476000 | -1.28225000 |
| H | 0.61961000 | 3.05925000 | -0.55430000 |
| H | 0.52461000 | 2.58228000 | -2.25844000 |
| C | -2.78868000 | 0.71233000 | -0.23245000 |
| H | -2.85378000 | 0.05624000 | -1.46718000 |
| C | -1.47176000 | 2.81192000 | -1.20430000 |
| H | -1.76687000 | 3.79870000 | -0.84038000 |
| H | -1.97893000 | 2.61601000 | -2.15452000 |
| C | -4.22652000 | 1.10134000 | 0.02998000 |
| H | -4.30039000 | 1.31508000 | 1.10419000 |
| H | -4.52064000 | 2.01395000 | -0.49426000 |
| C | -4.05822000 | -1.33572000 | -0.15131000 |
| H | -4.22818000 | -2.13937000 | -0.87475000 |
| C | -5.03707000 | -0.15340000 | -0.35906000 |
| H | -5.95262000 | -0.26106000 | 0.22486000 |
| H | -5.32752000 | -0.08973000 | -1.41404000 |
| C | 2.86225000 | -1.08022000 | -2.32276000 |
| H | 1.84606000 | -1.05682000 | -2.73269000 |
| H | 3.39751000 | -0.22350000 | -2.74539000 |
| H | 3.34825000 | -1.98933000 | -2.69348000 |
| C | 2.90209000 | 0.24170000 | 2.23075000 |
| H | 3.61155000 | 0.70991000 | 2.92088000 |
| H | 1.90566000 | 0.37827000 | 2.66484000 |
| H | 3.12166000 | -0.82842000 | 2.22212000 |
| C | 2.80031000 | 2.40629000 | 1.00163000 |
| H | 3.54601000 | 2.83186000 | 1.68144000 |
| H | 2.88993000 | 2.93307000 | 0.04567000 |
| H | 1.81768000 | 2.63417000 | 1.43207000 |
| C | -0.30215000 | -1.68172000 | 1.65478000 |
| H | 0.63654000 | -1.65385000 | 2.21259000 |
| H | -0.98342000 | -0.97232000 | 2.13889000 |
| H | -0.72423000 | -2.68574000 | 1.77261000 |
| C | -1.49826000 | 2.13613000 | 1.25668000 |
| H | -2.34506000 | 2.78563000 | 1.49647000 |
| H | -1.46529000 | 1.32697000 | 1.99144000 |
| H | -0.58844000 | 2.72790000 | 1.37283000 |
| C | -4.11982000 | -1.93461000 | 1.26943000 |
| H | -5.10385000 | -2.38416000 | 1.42377000 |
| H | -3.37035000 | -2.71607000 | 1.41682000 |
| H | -3.97258000 | -1.17239000 | 2.04122000 |
| H | -1.24067000 | -1.52622000 | -1.67321000 |

**IM7-2**
Sum of electronic and thermal Free Energies
= -976.527280

| | | | |
|---|---|---|---|
| C | 2.10218500 | 0.19945300 | -0.32714600 |
| C | 2.83780500 | -1.13286000 | -0.66034200 |
| C | 2.24539600 | -2.30495900 | 0.12905700 |
| C | 0.77416600 | -2.49888000 | -0.18088000 |
| C | -0.12765600 | -1.29479800 | 0.20328500 |
| C | 0.59270800 | 0.13012300 | 0.06732000 |
| H | 2.79649000 | -3.22012200 | -0.11987800 |
| H | 2.15714300 | 0.80443200 | -1.24543500 |
| H | 0.67218900 | -2.71234200 | -1.25088500 |
| H | 0.39189900 | -3.38648900 | 0.33784000 |
| H | 0.54816400 | 0.55352200 | 1.07705500 |
| H | 2.38295700 | -2.14653100 | 1.20448300 |
| C | 3.03018100 | 0.95111600 | 0.68289200 |
| C | 4.31491800 | -0.89054100 | -0.24002100 |
| H | 4.56215200 | -1.49556100 | 0.63945900 |
| H | 5.01335900 | -1.18543100 | -1.02865200 |
| C | 4.40929300 | 0.60022100 | 0.09788000 |
| H | 5.22736100 | 0.83671300 | 0.78602800 |
| H | 4.56994600 | 1.18860800 | -0.81533000 |
| C | -1.37410700 | -1.51646400 | -0.71136400 |
| H | -1.69714500 | -2.54800300 | -0.52560400 |
| C | -0.27145200 | 1.04584300 | -0.79772900 |
| H | -0.55080600 | 0.51454700 | -1.71705200 |
| C | -2.66177600 | -0.67045300 | -0.61855600 |
| C | -1.57195900 | 1.59787300 | -0.16719300 |
| C | 0.03074000 | 2.48789900 | -1.28110500 |
| H | 0.58060300 | 3.08765000 | -0.55764200 |
| H | 0.50079100 | 2.58931300 | -2.25995800 |
| C | -2.72199600 | 0.70373800 | -0.17388600 |
| H | -2.85612700 | -0.26280000 | -1.66724700 |
| C | -1.49238300 | 2.77514400 | -1.20473100 |
| H | -1.82506800 | 3.75093400 | -0.84533000 |
| H | -1.99987200 | 2.52879900 | -2.14202700 |
| C | -4.10758300 | 1.06425900 | 0.19647100 |
| H | -4.08371300 | 1.08213300 | 1.30185900 |
| H | -4.39873100 | 2.07110400 | -0.11738100 |
| C | -4.01965400 | -1.31712000 | -0.24052400 |
| H | -4.28821400 | -2.06979400 | -0.98696500 |

| | | | |
|---|---|---|---|
| C | -4.97381100 | -0.10885500 | -0.29768300 |
| H | -5.87066100 | -0.24610800 | 0.30823600 |
| H | -5.29097500 | 0.08273300 | -1.32854500 |
| C | 2.75389500 | -1.39940100 | -2.17051200 |
| H | 1.72530700 | -1.33964000 | -2.54667300 |
| H | 3.33956800 | -0.65114800 | -2.71496400 |
| H | 3.15096600 | -2.38864100 | -2.42371700 |
| C | 2.93448600 | 0.44661700 | 2.13136700 |
| H | 3.63537200 | 1.01041300 | 2.75549000 |
| H | 1.93673100 | 0.59961400 | 2.55895500 |
| H | 3.18919700 | -0.60985300 | 2.23450300 |
| C | 2.81008000 | 2.46313200 | 0.69764700 |
| H | 3.58909500 | 2.95143100 | 1.29217600 |
| H | 2.84651200 | 2.88418000 | -0.31303000 |
| H | 1.84821100 | 2.72626000 | 1.15609100 |
| C | -0.54758000 | -1.46470700 | 1.67064800 |
| H | 0.32308400 | -1.38706000 | 2.33033100 |
| H | -1.26194900 | -0.69398300 | 1.98642200 |
| H | -1.01282000 | -2.44040800 | 1.84215000 |
| C | -1.43474100 | 2.18148400 | 1.26169400 |
| H | -2.28017100 | 2.83980100 | 1.47925700 |
| H | -1.37591100 | 1.41034300 | 2.03288800 |
| H | -0.51930400 | 2.77467900 | 1.31454000 |
| C | -3.98085100 | -1.97289100 | 1.13980000 |
| H | -4.96239200 | -2.38952600 | 1.37809200 |
| H | -3.25681600 | -2.78994300 | 1.17739700 |
| H | -3.71775300 | -1.25832100 | 1.92816000 |
| H | -1.04049100 | -1.51535200 | -1.75343800 |

**TS_7-2−7-3**
Sum of electronic and thermal Free Energies
= -976.526140

| | | | |
|---|---|---|---|
| C | -0.10131000 | -1.19176000 | -0.85569000 |
| C | -0.59932000 | 0.28126000 | -0.55119000 |
| C | -0.59258000 | -2.18505000 | 0.23754000 |
| C | -0.99517000 | -1.53763000 | 1.55464000 |
| C | -2.17383000 | -0.56193000 | 1.39877000 |
| C | -1.97504000 | 0.42534000 | 0.19116000 |
| C | -3.48191000 | -1.34539000 | 1.06665000 |
| C | -3.81903000 | -1.05816000 | -0.40196000 |
| C | -3.29937000 | 0.36900000 | -0.64406000 |
| H | -0.69824000 | 0.79170000 | -1.51794000 |
| C | -2.31831000 | 0.20065000 | 2.72266000 |
| H | -1.95156000 | 1.44162000 | 0.60244000 |
| C | -3.17083000 | 0.75019000 | -2.11955000 |
| C | -4.29689000 | 1.36509000 | -0.01911000 |
| C | -0.65836000 | -1.65587000 | -2.21345000 |
| C | 0.34150000 | 1.16193000 | 0.28145000 |
| C | 2.53724000 | 0.57714000 | 0.28059000 |
| C | 2.33611000 | -0.88143000 | 0.09586000 |
| C | 1.44192000 | -1.36914000 | -1.03104000 |
| C | 3.59627000 | 0.78489000 | 1.31089000 |
| C | 4.28670000 | -0.59248000 | 1.45346000 |
| C | 3.78766000 | -1.40826000 | 0.23845000 |
| C | 0.11864000 | 2.69022000 | 0.42432000 |
| C | 1.63420000 | 2.97406000 | 0.32167000 |
| C | 1.79504000 | 1.60444000 | -0.36461000 |
| C | 4.61085000 | -1.15545000 | -1.02330000 |
| C | 1.75353000 | 1.73645000 | -1.88841000 |
| H | 0.48385000 | 0.75625000 | 1.28963000 |
| H | -1.46114000 | -2.73327000 | -0.14649000 |
| H | 0.17596000 | -2.94730000 | 0.41554000 |
| H | -1.28390000 | -2.31445000 | 2.27198000 |
| H | -0.13831000 | -1.02527000 | 2.01551000 |
| H | -4.29248000 | -0.99475000 | 1.71381000 |
| H | -3.37241000 | -2.41674000 | 1.26517000 |
| H | -3.31519000 | -1.77316000 | -1.06017000 |
| H | -4.89075000 | -1.13987000 | -0.61132000 |
| H | -1.40325000 | 0.75907000 | 2.95954000 |
| H | -3.14457000 | 0.91599000 | 2.69308000 |
| H | -2.50571000 | -0.49473000 | 3.54822000 |
| H | -2.78803000 | 1.77326000 | -2.22520000 |
| H | -4.15789000 | 0.72618000 | -2.59183000 |
| H | -2.52057000 | 0.08625000 | -2.69003000 |
| H | -4.48186000 | 1.16706000 | 1.03936000 |
| H | -3.92706000 | 2.39272000 | -0.10904000 |
| H | -5.25907000 | 1.30822000 | -0.53751000 |
| H | -0.35186000 | -0.98521000 | -3.02501000 |
| H | -0.28020000 | -2.65763000 | -2.44397000 |
| H | -1.74510000 | -1.71585000 | -2.21435000 |
| H | 1.78182000 | -0.96136000 | -1.98526000 |
| H | 1.60534000 | -2.44986000 | -1.11903000 |
| H | 4.25862000 | 1.58391000 | 0.95040000 |
| H | 3.17011000 | 1.16317000 | 2.24925000 |
| H | 3.98009000 | -1.07632000 | 2.38470000 |
| H | 5.37410000 | -0.49620000 | 1.47408000 |
| H | 3.76929000 | -2.47914000 | 0.45950000 |
| H | -0.37774000 | 3.00489000 | 1.34361000 |
| H | -0.43173000 | 3.08181000 | -0.43596000 |
| H | 1.93689000 | 3.83692000 | -0.27283000 |
| H | 2.10637000 | 3.02877000 | 1.30471000 |
| H | 4.19151000 | -1.67853000 | -1.88712000 |
| H | 5.63780000 | -1.50306000 | -0.88804000 |
| H | 4.65301000 | -0.08526000 | -1.26986000 |
| H | 2.72341000 | 2.11483000 | -2.22279000 |
| H | 0.99598000 | 2.47629000 | -2.15654000 |
| H | 1.53087000 | 0.81695000 | -2.42073000 |
| H | 1.85744000 | -1.16645000 | 1.05602000 |

**IM7-3**
Sum of electronic and thermal Free Energies
= -976.553956

| | | | |
|---|---|---|---|
| C | 0.18628300 | -1.05575000 | 1.03508800 |
| C | 0.54031600 | 0.31475000 | 0.33039600 |
| C | 0.85668000 | -2.25567400 | 0.30642200 |
| C | 1.34022900 | -1.94985300 | -1.10445200 |
| C | 2.39778500 | -0.83463400 | -1.14605200 |
| C | 1.96251600 | 0.42756900 | -0.31221700 |
| C | 3.72960400 | -1.31432800 | -0.49067500 |
| C | 3.87902200 | -0.55911000 | 0.83566700 |
| C | 3.18590700 | 0.79285400 | 0.59227900 |
| H | 0.47143000 | 1.07688800 | 1.12254800 |
| C | 2.62952700 | -0.47904400 | -2.62178500 |
| H | 1.87168300 | 1.25927100 | -1.02130900 |
| C | 2.85423900 | 1.56042400 | 1.87323000 |
| C | 4.13763100 | 1.68868200 | -0.22516700 |
| C | 0.65664800 | -1.05401100 | 2.49972900 |
| C | -0.44242700 | 0.80376600 | -0.79340800 |
| C | -1.89222600 | 0.46217300 | -0.45151100 |
| C | -2.07387500 | -1.01960100 | -0.16184000 |
| C | -1.34786300 | -1.30546300 | 1.14072800 |
| C | -3.00538100 | 0.71110100 | -1.62341600 |
| C | -4.11263300 | -0.31719600 | -1.32955000 |
| C | -3.54813600 | -1.42150100 | -0.38823400 |
| C | -0.45620700 | 2.34060100 | -0.90605900 |
| C | -1.51991400 | 2.75908800 | 0.11494300 |
| C | -2.40087500 | 1.57886600 | 0.30176000 |
| C | -4.38167800 | -1.66696200 | 0.87115600 |
| C | -3.58062000 | 1.67643500 | 1.18126500 |
| H | -0.17333300 | 0.33736500 | -1.74660600 |
| H | 1.71962100 | -2.59066300 | 0.89522000 |
| H | 0.16730800 | -3.10821700 | 0.28611900 |
| H | 1.77475400 | -2.85390100 | -1.54691100 |
| H | 0.49369100 | -1.68498000 | -1.75214700 |
| H | 4.56845500 | -1.07951900 | -1.15424900 |
| H | 3.74282600 | -2.40056800 | -0.35211500 |
| H | 3.38895200 | -1.10790900 | 1.64531400 |
| H | 4.92544000 | -0.43042700 | 1.13208000 |
| H | 1.70152000 | -0.14587300 | -3.10242700 |
| H | 3.36830700 | 0.31845700 | -2.73857400 |
| H | 2.99197000 | -1.35438500 | -3.17199200 |
| H | 2.35400800 | 2.50998000 | 1.64059500 |
| H | 3.77845200 | 1.80699900 | 2.40535300 |
| H | 2.21889600 | 1.00310000 | 2.56278700 |
| H | 4.46828900 | 1.21250600 | -1.15125100 |
| H | 3.65130000 | 2.63483600 | -0.48967000 |
| H | 5.03127900 | 1.92354100 | 0.36150900 |
| H | 0.22407700 | -0.21628200 | 3.05957000 |
| H | 0.33380000 | -1.98226800 | 2.98436100 |
| H | 1.73962600 | -1.00317400 | 2.59452600 |
| H | -1.77491900 | -0.68248500 | 1.94153600 |
| H | -1.49687800 | -2.34428800 | 1.45518700 |
| H | -3.38635200 | 1.73055000 | -1.75826300 |
| H | -2.41431700 | 0.48317300 | -2.51726600 |
| H | -4.45911000 | -0.73181000 | -2.28035400 |
| H | -4.98964900 | 0.16483700 | -0.88624300 |
| H | -3.54203200 | -2.36991700 | -0.93398400 |
| H | -0.76220700 | 2.65421000 | -1.90928800 |
| H | 0.51246900 | 2.79951400 | -0.69977500 |
| H | -1.08511600 | 2.91245100 | 1.12010100 |
| H | -2.07923600 | 3.67542100 | -0.10126000 |
| H | -4.00324700 | -2.53472500 | 1.41770000 |
| H | -5.42001400 | -1.87845200 | 0.60037600 |
| H | -4.38326200 | -0.82477200 | 1.56684100 |
| H | -3.85238500 | 2.71311800 | 1.38268400 |
| H | -3.30115100 | 1.19437900 | 2.13055500 |
| H | -4.43884500 | 1.12087900 | 0.80412100 |
| H | -1.50340100 | -1.50594300 | -0.96518400 |

**TS_7-3−7-4**
Sum of electronic and thermal Free Energies
= -976.550546

| | | | |
|---|---|---|---|
| C | 0.19181000 | -1.11630000 | 0.96162000 |
| C | 0.54964000 | 0.29427000 | 0.33564000 |
| C | 0.88020000 | -2.26996000 | 0.17710000 |
| C | 1.37336000 | -1.88756000 | -1.21151000 |
| C | 2.42360000 | -0.76459000 | -1.18508000 |
| C | 1.97546000 | 0.44787000 | -0.28735000 |
| C | 3.75316000 | -1.27204000 | -0.54601000 |
| C | 3.88651000 | -0.59195000 | 0.82198000 |
| C | 3.18855000 | 0.76828000 | 0.64781000 |
| H | 0.46027000 | 1.01885000 | 1.15988000 |
| C | 2.66355000 | -0.32784000 | -2.63743000 |
| H | 1.88556000 | 1.31798000 | -0.94889000 |
| C | 2.84025000 | 1.46301000 | 1.96531000 |
| C | 4.14234000 | 1.71260000 | -0.11059000 |
| C | 0.64723000 | -1.18940000 | 2.42954000 |
| C | -0.42894000 | 0.83060000 | -0.77494000 |
| C | -1.83655000 | 0.45659000 | -0.37955000 |
| C | -2.06867000 | -1.02346000 | -0.24924000 |
| C | -1.34331000 | -1.38840000 | 1.04324000 |
| C | -3.12887000 | 0.93623000 | -1.42342000 |
| C | -4.19528000 | -0.13467000 | -1.17802000 |
| C | -3.55801000 | -1.38099000 | -0.49333000 |
| C | -0.46689000 | 2.37039000 | -0.84611000 |
| C | -1.56565000 | 2.76474000 | 0.15257000 |
| C | -2.42618000 | 1.53638000 | 0.30885000 |
| C | -4.33854000 | -1.90299000 | 0.71540000 |
| C | -3.55727000 | 1.56820000 | 1.27579000 |
| H | -0.16834000 | 0.38677000 | -1.74104000 |
| H | 1.74149000 | -2.62623000 | 0.75544000 |
| H | 0.19989000 | -3.12698000 | 0.10674000 |
| H | 1.81860000 | -2.76439000 | -1.69600000 |

| | | | |
|---|---|---|---|
| H | 0.53052000 | -1.59648000 | -1.85270000 |
| H | 4.59591000 | -0.99471000 | -1.18783000 |
| H | 3.77267000 | -2.36427000 | -0.46897000 |
| H | 3.39289000 | -1.18752000 | 1.59589000 |
| H | 4.92961000 | -0.47457000 | 1.13429000 |
| H | 1.73698000 | 0.02671000 | -3.10559000 |
| H | 3.39871000 | 0.47865000 | -2.70501000 |
| H | 3.03442000 | -1.16980000 | -3.23229000 |
| H | 2.33765000 | 2.42139000 | 1.77973000 |
| H | 3.75788000 | 1.68375000 | 2.51969000 |
| H | 2.20046000 | 0.86629000 | 2.61669000 |
| H | 4.48300000 | 1.29067000 | -1.05914000 |
| H | 3.65331000 | 2.66977000 | -0.32551000 |
| H | 5.02989000 | 1.91832000 | 0.49593000 |
| H | 0.20436000 | -0.38468000 | 3.02787000 |
| H | 0.32557000 | -2.14378000 | 2.86118000 |
| H | 1.72904000 | -1.13825000 | 2.53669000 |
| H | -1.78338000 | -0.81713000 | 1.87394000 |
| H | -1.48398000 | -2.44666000 | 1.28715000 |
| H | -3.51911000 | 1.95849000 | -1.46703000 |
| H | -2.53878000 | 0.80859000 | -2.33693000 |
| H | -4.63983000 | -0.39449000 | -2.14282000 |
| H | -5.00890000 | 0.27478000 | -0.57317000 |
| H | -3.56308000 | -2.19662000 | -1.22220000 |
| H | -0.73632000 | 2.69812000 | -1.85517000 |
| H | 0.49088000 | 2.83390000 | -0.60155000 |
| H | -1.15284000 | 2.95206000 | 1.15387000 |
| H | -2.13989000 | 3.65648000 | -0.11630000 |
| H | -3.93959000 | -2.86803000 | 1.03811000 |
| H | -5.38740000 | -2.05701000 | 0.44698000 |
| H | -4.30754000 | -1.23951000 | 1.58334000 |
| H | -4.23202000 | 2.40432000 | 1.06915000 |
| H | -3.10310000 | 1.75961000 | 2.25606000 |
| H | -4.11528000 | 0.64071000 | 1.34053000 |
| H | -1.49921000 | -1.46927000 | -1.07484000 |

**IM7-4**

Sum of electronic and thermal Free Energies
= -976.569163

| | | | |
|---|---|---|---|
| C | 0.16636700 | -1.43945600 | 0.24028700 |
| C | 0.48066000 | 0.09123600 | 0.06549100 |
| C | 1.29562600 | -2.35018700 | -0.27803000 |
| C | 2.00872100 | -1.72760200 | -1.46579600 |
| C | 2.81987300 | -0.49091500 | -1.04534200 |
| C | 1.96941700 | 0.49845800 | -0.16729300 |
| C | 4.01046200 | -0.91947800 | -0.13252200 |
| C | 3.65755000 | -0.48087400 | 1.29279400 |
| C | 2.83248800 | 0.80131900 | 1.10645000 |
| H | 0.16470400 | 0.55822000 | 1.00147500 |
| C | 3.32753100 | 0.19117600 | -2.32159700 |
| H | 1.90608600 | 1.44495000 | -0.71697000 |
| C | 2.05477100 | 1.18297500 | 2.36698600 |
| C | 3.77657400 | 1.97361600 | 0.78257600 |
| C | -0.13426000 | -1.78319600 | 1.70420400 |
| C | -0.37975500 | 0.78571300 | -1.02833400 |
| C | -1.86097600 | 0.42536300 | -0.90085500 |
| C | -2.20334900 | -0.92260000 | -0.49656100 |
| C | -1.11871400 | -1.88044300 | -0.60015000 |
| C | -4.08951300 | 1.33209900 | -0.36539900 |
| C | -4.50829700 | -0.02842500 | 0.21995100 |
| C | -3.60997500 | -1.23698000 | -0.15643200 |
| C | -0.42109400 | 2.34348200 | -0.91376900 |
| C | -1.91487500 | 2.75557400 | -0.81287100 |
| C | -2.59431000 | 1.55504700 | -0.13849600 |
| C | -3.79789700 | -2.41171800 | 0.81502200 |
| C | -2.29254200 | 1.56222700 | 1.37167100 |
| H | 0.00138800 | 0.48892800 | -2.01335000 |
| H | 2.01477000 | -2.51864900 | 0.53043300 |
| H | 0.88810600 | -3.33806900 | -0.52307500 |
| H | 2.68406000 | -2.45308000 | -1.93203400 |
| H | 1.28109600 | -1.45182000 | -2.24565800 |
| H | 4.92507600 | -0.41523700 | -0.46122900 |
| H | 4.20982800 | -1.99415600 | -0.20431200 |
| H | 3.05156400 | -1.24111400 | 1.80281500 |
| H | 4.54158400 | -0.31289900 | 1.91691900 |
| H | 2.49313100 | 0.49365300 | -2.96585500 |
| H | 3.91529000 | 1.08534300 | -2.09626000 |
| H | 3.96335200 | -0.49201500 | -2.89535800 |
| H | 1.43352200 | 2.07284500 | 2.20231800 |
| H | 2.75657100 | 1.42461400 | 3.17132100 |
| H | 1.40997800 | 0.37863400 | 2.73235500 |
| H | 4.39205900 | 1.78188200 | -0.09988400 |
| H | 3.20502200 | 2.89024400 | 0.59762900 |
| H | 4.45238200 | 2.16165800 | 1.62280000 |
| H | -0.95741100 | -1.18087600 | 2.10741400 |
| H | -0.39515700 | -2.84190100 | 1.81059100 |
| H | 0.74700400 | -1.59969800 | 2.32425300 |
| H | -1.39017900 | -2.90620900 | -0.35160300 |
| H | -0.78942400 | -1.83134100 | -1.65258300 |
| H | -4.67925200 | 2.11939500 | 0.11550500 |
| H | -4.31361700 | 1.37731400 | -1.43922300 |
| H | -5.53191300 | -0.26967800 | -0.07880500 |
| H | -4.52341300 | 0.03581000 | 1.31344900 |
| H | -3.93175800 | -1.59257200 | -1.16229900 |
| H | 0.07029600 | 2.82982700 | -1.75875100 |
| H | 0.11935800 | 2.65907900 | -0.01441200 |
| H | -2.05009000 | 3.68491700 | -0.25264600 |
| H | -2.33928700 | 2.90453900 | -1.81353100 |
| H | -3.36080800 | -3.34221300 | 0.44998200 |
| H | -4.86698200 | -2.58024300 | 0.95923400 |
| H | -3.36193900 | -2.17423400 | 1.78991000 |
| H | -2.98127900 | 2.25365500 | 1.86772400 |

| | | | |
|---|---|---|---|
| H | -1.27845600 | 1.90052500 | 1.59292900 |
| H | -2.42057700 | 0.57895000 | 1.83863600 |
| H | -2.30157600 | 0.45255400 | -1.92178300 |

**TS_7-4−7-5**
Sum of electronic and thermal Free Energies
= -976.563534

| | | | |
|---|---|---|---|
| C | 0.22038000 | -1.49762000 | 0.12476000 |
| C | 0.50982000 | 0.05689000 | 0.11355000 |
| C | 1.37907000 | -2.31447000 | -0.47927000 |
| C | 2.05576000 | -1.55902000 | -1.61202000 |
| C | 2.83691000 | -0.34135000 | -1.09128000 |
| C | 1.97601000 | 0.53576000 | -0.11003000 |
| C | 4.05747000 | -0.81564000 | -0.24306000 |
| C | 3.71887000 | -0.52395000 | 1.22258000 |
| C | 2.85607000 | 0.74562000 | 1.17135000 |
| H | 0.19585000 | 0.42938000 | 1.09581000 |
| C | 3.29705000 | 0.47356000 | -2.30624000 |
| H | 1.87172000 | 1.52842000 | -0.56458000 |
| C | 2.09488000 | 0.98616000 | 2.47577000 |
| C | 3.76023000 | 1.97001000 | 0.93975000 |
| C | -0.04780000 | -1.96798000 | 1.55957000 |
| C | -0.41610000 | 0.81415000 | -0.88094000 |
| C | -1.83598000 | 0.40111000 | -0.59283000 |
| C | -2.18436000 | -0.94762000 | -0.47319000 |
| C | -1.04471000 | -1.87511000 | -0.73517000 |
| C | -4.15006000 | 1.20068000 | -0.54362000 |
| C | -4.57528000 | -0.15767000 | 0.05280000 |
| C | -3.59935000 | -1.35266000 | -0.14912000 |
| C | -0.49348000 | 2.36048000 | -0.75810000 |
| C | -1.99974000 | 2.71075000 | -0.85761000 |
| C | -2.71038000 | 1.54395000 | -0.14133000 |
| C | -3.61275000 | -2.27684000 | 1.08487000 |
| C | -2.57907000 | 1.69768000 | 1.38758000 |
| H | -0.13668000 | 0.53622000 | -1.90634000 |
| H | 2.11521000 | -2.53999000 | 0.29907000 |
| H | 0.99852000 | -3.28548000 | -0.81893000 |
| H | 2.74649000 | -2.21463000 | -2.15321000 |
| H | 1.30809000 | -1.23601000 | -2.35330000 |
| H | 4.95077000 | -0.25534000 | -0.53801000 |
| H | 4.28533000 | -1.87242000 | -0.41876000 |
| H | 3.14189000 | -1.34579000 | 1.66614000 |
| H | 4.60885000 | -0.39013000 | 1.84648000 |
| H | 2.43994000 | 0.80982000 | -2.90268000 |
| H | 3.86420000 | 1.36001000 | -2.00894000 |
| H | 3.93850000 | -0.13279000 | -2.95507000 |
| H | 1.43995000 | 1.86417000 | 2.40414000 |
| H | 2.80696000 | 1.18030000 | 3.28387000 |
| H | 1.48732000 | 0.12952000 | 2.78033000 |
| H | 4.36029000 | 1.87977000 | 0.03096000 |
| H | 3.16006000 | 2.88304000 | 0.85409000 |
| H | 4.44983000 | 2.09797000 | 1.78001000 |
| H | -0.88594000 | -1.42567000 | 2.01620000 |
| H | -0.27478000 | -3.03958000 | 1.58401000 |
| H | 0.83116000 | -1.80433000 | 2.18833000 |
| H | -1.32439000 | -2.91635000 | -0.55764000 |
| H | -0.77276000 | -1.79707000 | -1.79678000 |
| H | -4.84346000 | 1.97311000 | -0.19745000 |
| H | -4.22182000 | 1.18449000 | -1.64019000 |
| H | -5.54901000 | -0.44315000 | -0.35397000 |
| H | -4.72888000 | -0.02895000 | 1.12875000 |
| H | -3.92444000 | -1.96465000 | -1.00363000 |
| H | 0.09172000 | 2.86481000 | -1.52900000 |
| H | -0.08552000 | 2.67674000 | 0.20861000 |
| H | -2.23663000 | 3.68182000 | -0.41539000 |
| H | -2.30916000 | 2.73976000 | -1.91031000 |
| H | -2.99735000 | -3.16864000 | 0.95023000 |
| H | -4.64118000 | -2.59924000 | 1.26229000 |
| H | -3.26908000 | -1.74366000 | 1.97590000 |
| H | -3.24937000 | 2.49409000 | 1.72446000 |
| H | -1.56267000 | 1.96748000 | 1.68497000 |
| H | -2.84439000 | 0.77904000 | 1.91868000 |
| H | -2.31586000 | -0.05718000 | -1.61182000 |

**IM7-5**
Sum of electronic and thermal Free Energies
= -976.565152

| | | | |
|---|---|---|---|
| C | 0.20268100 | -1.52199200 | 0.10604800 |
| C | 0.49272100 | 0.02921300 | 0.12004000 |
| C | 1.38735500 | -2.32064900 | -0.47688100 |
| C | 2.06653100 | -1.56085400 | -1.60554800 |
| C | 2.82781800 | -0.33187700 | -1.08385900 |
| C | 1.95124800 | 0.53717400 | -0.10944600 |
| C | 4.05027600 | -0.78559700 | -0.22756500 |
| C | 3.69960200 | -0.49700700 | 1.23565200 |
| C | 2.82002000 | 0.76049800 | 1.17895500 |
| H | 0.18529400 | 0.39628500 | 1.10514200 |
| C | 3.28232900 | 0.48897600 | -2.29718200 |
| H | 1.83814300 | 1.52718300 | -0.56519300 |
| C | 2.05033300 | 0.99236500 | 2.48010500 |
| C | 3.70724100 | 1.99747900 | 0.94802400 |
| C | -0.05575800 | -1.98542100 | 1.54464600 |
| C | -0.44489200 | 0.81287800 | -0.88237400 |
| C | -1.80612000 | 0.40451200 | -0.54761500 |
| C | -2.23449300 | -0.95275300 | -0.70349300 |
| C | -1.02505900 | -1.90342700 | -0.78636500 |
| C | -4.12191300 | 1.16301100 | -0.57242900 |
| C | -4.57616500 | -0.16479300 | 0.05939900 |
| C | -3.59525000 | -1.36927400 | -0.08818300 |
| C | -0.48207000 | 2.35742000 | -0.79742700 |
| C | -1.98637300 | 2.70885700 | -0.86738400 |
| C | -2.68314100 | 1.51764300 | -0.16885400 |

| | | | |
|---|---|---|---|
| C | -3.46885100 | -2.11605600 | 1.24230400 |
| C | -2.55225900 | 1.66572900 | 1.38058800 |
| H | -0.19628100 | 0.45927200 | -1.89484500 |
| H | 2.11874600 | -2.53568100 | 0.30879400 |
| H | 1.02188900 | -3.29596400 | -0.81982000 |
| H | 2.77195400 | -2.20839900 | -2.13733800 |
| H | 1.32281200 | -1.25142400 | -2.35596400 |
| H | 4.93706000 | -0.21309800 | -0.51880000 |
| H | 4.29307800 | -1.83941400 | -0.39966200 |
| H | 3.13143600 | -1.32613200 | 1.67638900 |
| H | 4.58418000 | -0.34957200 | 1.86404600 |
| H | 2.42353300 | 0.81206900 | -2.89877000 |
| H | 3.83636000 | 1.38349000 | -1.99901400 |
| H | 3.93451000 | -0.10955100 | -2.94230600 |
| H | 1.38263800 | 1.86080600 | 2.40519100 |
| H | 2.75743400 | 1.20035500 | 3.28897600 |
| H | 1.45660100 | 0.12676000 | 2.78658600 |
| H | 4.31154200 | 1.91432300 | 0.04146100 |
| H | 3.09505300 | 2.90238600 | 0.85968500 |
| H | 4.39212600 | 2.13574700 | 1.79039200 |
| H | -0.88989900 | -1.44578100 | 2.00693700 |

| | | | |
|---|---|---|---|
| H | -0.27998200 | -3.05685200 | 1.57431400 |
| H | 0.82903300 | -1.81745600 | 2.16480500 |
| H | -1.33883200 | -2.92062600 | -0.53468700 |
| H | -0.68891300 | -1.94501100 | -1.82814400 |
| H | -4.79559900 | 1.96783500 | -0.26212600 |
| H | -4.17496200 | 1.11461200 | -1.66798200 |
| H | -5.54763900 | -0.43837600 | -0.35964400 |
| H | -4.75252000 | 0.00577000 | 1.12650100 |
| H | -4.02111200 | -2.07901600 | -0.80570800 |
| H | 0.08990300 | 2.83204700 | -1.59602400 |
| H | -0.04342800 | 2.68819600 | 0.15081200 |
| H | -2.22810600 | 3.66282300 | -0.39218500 |
| H | -2.31692300 | 2.76321500 | -1.91139200 |
| H | -2.81770300 | -2.99032100 | 1.17188100 |
| H | -4.45906700 | -2.46458000 | 1.54804500 |
| H | -3.08921600 | -1.46821400 | 2.03807700 |
| H | -3.27753700 | 2.42439000 | 1.68890900 |
| H | -1.55484400 | 2.00171900 | 1.67100400 |
| H | -2.77266300 | 0.73835300 | 1.91210000 |
| H | -2.50296800 | -0.71850800 | -1.77738100 |
